## Supplementary Fig 1 - Env reside mapping for "Conformational flexibility of HIV-1 envelope glycoproteins modulates transmitted / founder sensitivity to broadly neutralizing antibodies"

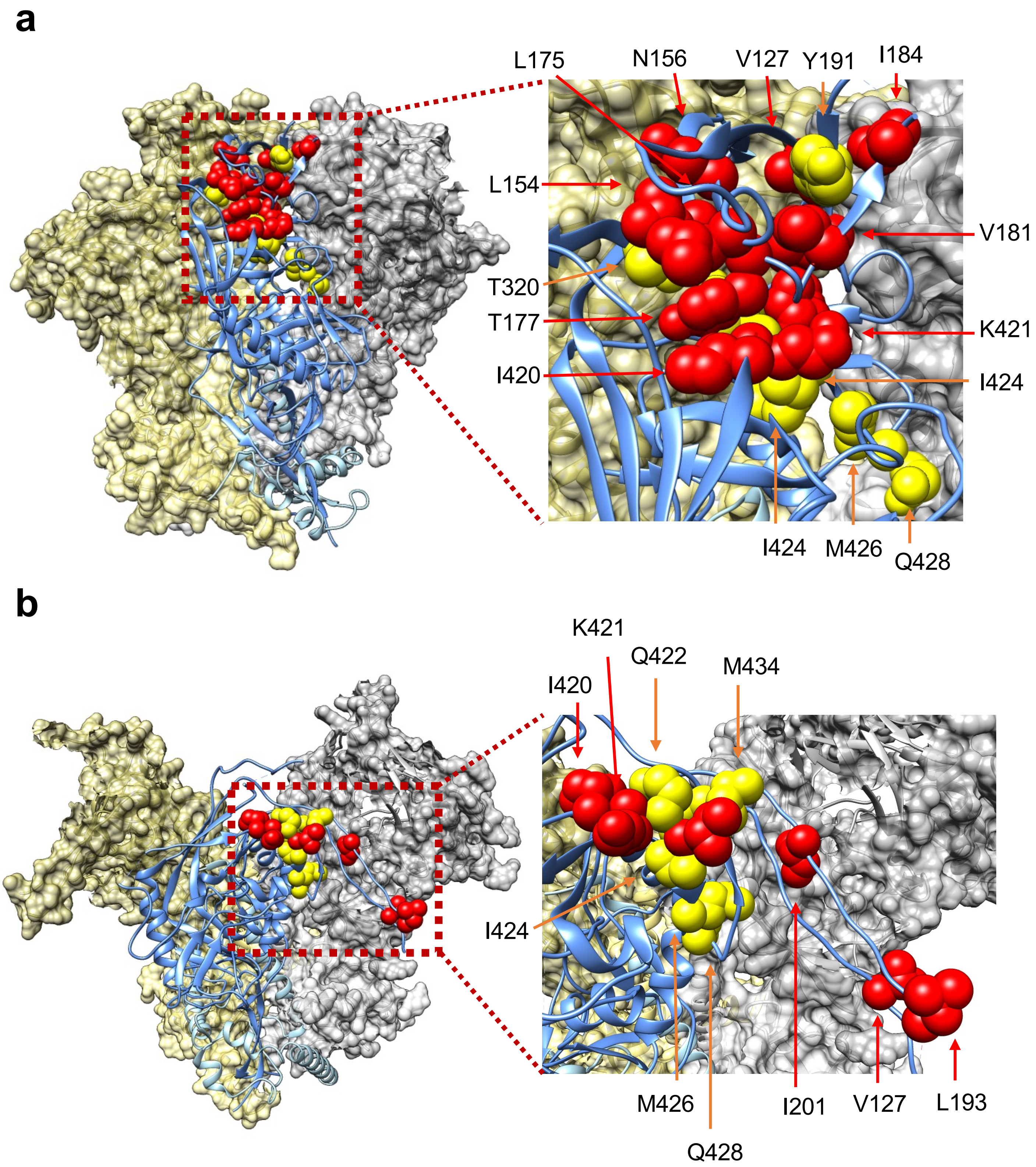


**Supplementary Figure 1.** We mapped Env residues that contribute to maintenance of Env closed conformation on available SOSIP Env structures. Residues that were resolved and in which changes resulted in hypersensitivity to multiple internal-epitope Env ligands (red) or only one internal-epitope Env ligand (yellow) are labeled (neutralization data and ligands tested are shown in Fig. 1 and Supplementary table 1). **a**, We used the crystal structure of BG505 SOSIP bound to BMS806, which blocks conformational changes of HIV-1 Env on virions (pdb entry 6mtj). **b**, We used the model of B41 SOSIP bound to sCD4 and 17b (pdb entry 5vn3). Left panels - SOSIP trimers with 2 of the protomers shown as surfaces and one front protomer shown as ribbon in which specific residues were mapped. Right - enhanced view of specific residues. Figure was prepared using the Chimera program (https://www.cgl.ucsf.edu/chimera/).
