## Supplementary Fig 2 -Flow cytometry gating strategy for "Conformational flexibility of HIV-1 envelope glycoproteins modulates transmitted / founder sensitivity to broadly neutralizing antibodies"

**a** Control

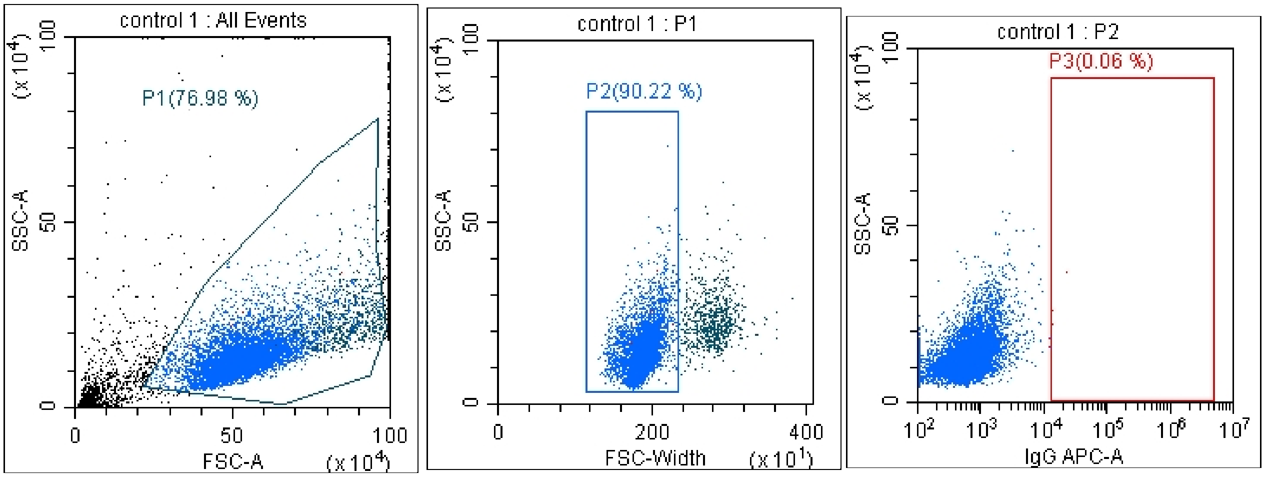


Tube Name: control 1
Sample ID:

| Population | Events | % Total | % Parent | Events/µL(V) |
|---|---|---|---|---|
| All Events | 10000 | 100.00 % | 100.00 % | 1168.90 |
| P1 | 7698 | 76.98 % | 76.98 % | 899.82 |
| P2 | 6945 | 69.45 % | 90.22 % | 811.80 |
| P3 | 4 | 0.04 % | 0.06 % | 0.47 |

**b** N6 bnAb

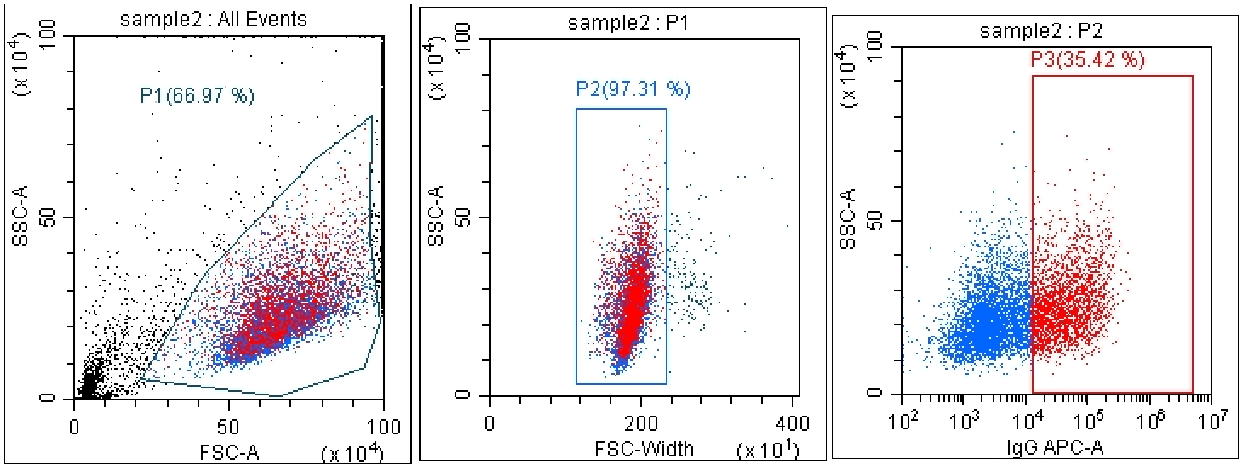


Tube Name: sample2
Sample ID:

| Population | Events | % Total | % Parent | Events/µL(V) |
|---|---|---|---|---|
| All Events | 10000 | 100.00 % | 100.00 % | 845.01 |
| P1 | 6697 | 66.97 % | 66.97 % | 565.90 |
| P2 | 6517 | 65.17 % | 97.31 % | 550.69 |
| P3 | 2308 | 23.08 % | 35.42 % | 195.03 |

**Supplementary Figure 2. Flow cytometry gating strategy.** 293T cells were transfected with Env-expressing plasmid (HIV-1$_{JRFL\Delta CT}$ or HIV-1$_{JRFL\Delta CT}$ L193R) and binding of different bnAbs to transfected cells was detected with allophycocyanin (APC)-conjugated F(ab′)2 fragment donkey anti-human IgG antibody. Cells were gated first according to side and forward scatter (FSC-A & SSC-A) and then according to SSC-A and FSC-width to exclude doublet cells. Gated 293T cells were then analyzed for the level of APC fluorescence. Control 293T cells (**a**) and 293T cells transfected with HIV-1$_{JRFL\Delta CT}$ L193R env plasmid and incubated with N6 bnAb (**b**) are shown.
