## Supplementary Table 1 - Env internal epitopes for "Conformational flexibility of HIV-1 envelope glycoproteins modulates transmitted / founder sensitivity to broadly neutralizing antibodies"

| JR-FL* | Relative infectivity (%) | $IC_{50}$ sCD4 (nM) Average | 19b (μg/ml) Average | 17b (μg/ml) Average | 902090 (μg/ml) Average | T20 (nM) Average | Counts (only single Env element <10%) sCD4<2 | 19b<5 | 17b<15 | 902090<10 | T20<0.23 |
|---|---|---|---|---|---|---|---|---|---|---|---|
| WT | 100 | 19.04 | 50 | 150 | 100 | 2.3 | 0 | 0 | 0 | 0 | 0 |
| V127A | 3 | 1.39 | 0.11 | 76.03 | 100 | 1.29 | 0 | 0 | 0 | 0 | 0 |
| I154A | 51 | 1.09 | 2.33 | 0.33 | 17.89 | 4.16 | 0 | 0 | 0 | 0 | 0 |
| N156A | 19 | 1.02 | 0.56 | 31.2 | ~5 | 85.01 | 0 | 0 | 0 | 0 | 0 |
| L175A | 172 | 0.91 | 2.52 | 60.18 | 5.21 | 4.25 | 0 | 0 | 0 | 0 | 0 |
| Y177A | 56 | 1.66 | 0.67 | 21.26 | 100 | 2.95 | 0 | 0 | 0 | 0 | 0 |
| V181I | 280 | 41.86 | 50 | 150 | 100 | 0.67 | 0 | 0 | 0 | 0 | 0 |
| V181I+L193A | 130 | 0.39 | 0.05 | 0.43 | 17.1 | 1.28 | 0 | 0 | 0 | 0 | 0 |
| I184L+L193A | 60 | 0.82 | 0.018 | 0.44 | 16.6 | 3.29 | 0 | 0 | 0 | 0 | 0 |
| Y191A | 87 | 1 | 6.75 | 65.35 | 26.43 | 3.76 | 1 | 0 | 0 | 0 | 0 |
| Y191A+I423A | 33 | 0.82 | 0.044 | 11.9 | 100 | 1.32 | 0 | 0 | 0 | 0 | 0 |
| Y191A+Q428A | 38 | 0.64 | 0.67 | 19.69 | 2.27 | 0.79 | 0 | 0 | 0 | 0 | 0 |
| L193A | 16 | 1.98 | 0.19 | 0.65 | 7.56 | 5.15 | 0 | 0 | 0 | 0 | 0 |
| L193A + I201W | 36 | 1.27 | 0.13 | 3.67 | 9.26 | 1.22 | 0 | 0 | 0 | 0 | 0 |
| L193A + I423V | 1012 | 0.67 | 0.04 | 0.29 | 3.9 | 2.1 | 0 | 0 | 0 | 0 | 0 |
| L193A + I423V-D674N | 63 | 0.37 | 0.027 | 0.18 | 7.49 | 1.33 | 0 | 0 | 0 | 0 | 0 |
| L193A + I423V-Δ733 | 1571 | 2.00 | 0.38 | 7.39 | 25.6 | 7.16 | 0 | 0 | 0 | 0 | 0 |
| L193A + I423V-Δ765 | 655 | 0.68 | 0.09 | 3.3 | 12.78 | 1.45 | 0 | 0 | 0 | 0 | 0 |
| L193G | 25 | 0.76 | 0.11 | 0.76 | 0.84 | 0.72 | 0 | 0 | 0 | 0 | 0 |
| L193K | 17 | 0.75 | 0.02 | 0.09 | 1.20 | 2.69 | 0 | 0 | 0 | 0 | 0 |
| L193I | 47 | 11.42 | 50.00 | 150 | 100 | 5.48 | 0 | 0 | 0 | 0 | 0 |
| L193V | 70 | 5.21 | 13.25 | 150 | 55.96 | 2.36 | 0 | 0 | 0 | 0 | 0 |
| L193E | 43 | 0.61 | 0.11 | 1.05 | 1.60 | 0.39 | 0 | 0 | 0 | 0 | 0 |
| L193D | 7 | 0.32 | 0.09 | 1.34 | 3.01 | 3.50 | 0 | 0 | 0 | 0 | 0 |
| L193M | 39 | 2.60 | 50.00 | 150 | 43.96 | 0.27 | 0 | 0 | 0 | 0 | 0 |
| L193S | 16 | 0.40 | 0.08 | 0.60 | 2.60 | 1.82 | 0 | 0 | 0 | 0 | 0 |
| L193R | 18 | 0.23 | 0.02 | 0.26 | 0.83 | 0.67 | 0 | 0 | 0 | 0 | 0 |
| L193W | 69 | 0.22 | 1.16 | 2.26 | 13.53 | 3.75 | 0 | 0 | 0 | 0 | 0 |
| L193Q | 18 | 1.17 | 0.12 | 0.66 | 3.58 | 2.12 | 0 | 0 | 0 | 0 | 0 |
| L193F | 70 | 3.20 | 23.84 | 150 | 11.24 | 2.71 | 0 | 0 | 0 | 0 | 0 |
| L193H | 8 | 0.33 | 0.06 | 0.81 | 3.72 | 0.83 | 0 | 0 | 0 | 0 | 0 |
| L193T | 22 | 0.93 | 0.05 | 0.98 | 0.68 | 1.87 | 0 | 0 | 0 | 0 | 0 |
| L193Y | 17 | 0.37 | 0.14 | 0.29 | 4.98 | 1.10 | 0 | 0 | 0 | 0 | 0 |
| L193P | 16 | 0.41 | 0.13 | 0.97 | 5.27 | 4.86 | 0 | 0 | 0 | 0 | 0 |
| T320R | 76 | 3.14 | 0.28 | 41.95 | 100 | 3.92 | 0 | 1 | 0 | 0 | 0 |
| S375W | 9 | 6.24 | 50 | 150 | 100 | 0.33 | 0 | 0 | 0 | 0 | 0 |
| I420A | 159 | 60.24 | 3.08 | 150 | 2.03 | 0.66 | 0 | 0 | 0 | 0 | 0 |
| K421A | 199 | 1.05 | 4.8 | 9.59 | 27.19 | 0.79 | 0 | 0 | 0 | 0 | 0 |
| Q422A | 72 | 1.11 | 2.73 | 150 | 22.76 | 2.14 | 0 | 0 | 0 | 0 | 0 |
| Q422D | 7 | 0.89 | 0.24 | 5.94 | 6.56 | 0.67 | 0 | 0 | 0 | 0 | 0 |
| Q422N | 7 | 6.59 | 1.12 | 45.86 | 28.18 | 0.73 | 0 | 1 | 0 | 0 | 0 |
| I423A | 696 | 1.22 | 0.1 | 39.22 | 1.57 | 0.21 | 0 | 0 | 0 | 0 | 0 |
| I423V | 943 | 19.92 | 47.8 | 150 | 100 | 1.88 | 0 | 0 | 0 | 0 | 0 |
| I424A | 27 | 21.74 | 0.013 | 150 | 16.06 | 2.73 | 0 | 1 | 0 | 0 | 0 |
| N425A | 590 | 335.4 | 50 | 150 | 100 | 0.27 | 0 | 0 | 0 | 0 | 0 |
| M426A | 214 | 254.2 | 50 | 150 | 100 | 0.22 | 0 | 0 | 0 | 0 | 1 |
| Q428A | 532 | 135.4 | 50 | 150 | 100 | 0.18 | 0 | 0 | 0 | 0 | 1 |
| E429A | 1127 | 13.7 | 50 | 150 | 24.65 | 2.9 | 0 | 0 | 0 | 0 | 0 |
| V430A | 18 | 500 | 50 | 150 | 68.32 | 1.35 | 0 | 0 | 0 | 0 | 0 |
| G431A | 752 | 25.27 | 50 | 150 | 59.34 | 2.13 | 0 | 0 | 0 | 0 | 0 |
| K432A | 579 | 29.89 | 50 | 150 | 100 | 2.56 | 0 | 0 | 0 | 0 | 0 |
| A433G | 2084 | 14.46 | 9.49 | 150 | 21.15 | 1.45 | 0 | 0 | 0 | 0 | 0 |
| M434A | 10 | 7.2 | 0.84 | 69.58 | 22.51 | 1.77 | 0 | 1 | 0 | 0 | 0 |
| Y435A | 9 | 46.27 | 0.71 | 150 | 15.87 | 1.29 | 0 | 1 | 0 | 0 | 0 |

| | |
|---|---|
| Average WT infectivity (relative light units) | 1907095.3 |
| Average infectivity (%) | 243.3 |
| Range infectivity (%) | 3 - 2084 |

| % | sCD4 | 19b (V3) | 17b (B20-B21) | 902090 (V1/V2) | T20 (gp41 HR1) | Total |
|---|---|---|---|---|---|---|
| Criteria | **< 2nM** | **< 5 ug/ml** | **< 15 ug/ml** | **< 10 ug/ml** | **< 0.23 nM** | |
| Number of variants out of 53 in each group | 31.0 | 37.0 | 24.0 | 20.0 | 3.0 | 115 |
| % of variants meeting criteria in each group (Number of varaints in each group / 53 *100) | 58.5 | 69.8 | 45.3 | 37.7 | 5.7 | |
| % of variants in each group from the total (Number of varaints in each group / 115 *100) | **27.0** | **32.2** | **20.9** | **17.4** | **2.6** | 100 |

| Number of single site exposures | sCD4 | 19b (V3) | 17b (B20-B21) | 902090 (V1/V2) | T20 (gp41 HR1) |
|---|---|---|---|---|---|
| Criteria (only one of the following) | **< 2nM** | **< 5 ug/ml** | **< 15 ug/ml** | **< 10 ug/ml** | **< 0.23 nM** |
| Number of variants | 1 | 5 | 0 | 0 | 2 |

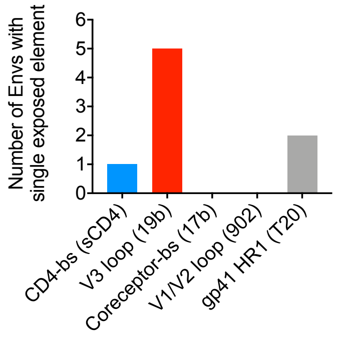


* Distribution of exposed elements among 53 functional Env intermediates. We selected HIV-1$_{JRFL}$ Env variants, which are enriched in intermediates based on prior studies (this is not a random selection). WT and 53 Env mutants were analyzed for sensitivity to ligands that preferentially recognize open Env conformations (17b, 19b, sCD4, 902 directed against the V1/V2 loop, and T20). Env mutants that were > 10 times more sensitive than WT ($IC_{50}$ mutant / $IC_{50}$ WT < 0.1) were identified and their prevalence was calculated. $IC_{50}$ of some Env mutants has been published previously (Herschhorn et al. mBio 2016; Nat Commun 2017).

* Env variants that showed >10 fold hypersensitivity to any of the ligands (17b, 19b, sCD4, 902 directed against the V1/V2 loop, and T20) are in blue.
