## Supplementary Table 2 - T_F Env consensus analysis CH040 d127-d1737 for "Conformational flexibility of HIV-1 envelope glycoproteins modulates transmitted / founder sensitivity to broadly neutralizing antibodies"

### Aligned using an external algorithm

**Consensus**

1. B.US.2006.700010040_C9_4520.EU289193
2. B.US.2006.CH0040_3_d0127_ipe032_27_49.MG900133
3. B.US.2006.CH0040_3_d0127_ipe032_2_02.MG900134
4. B.US.2006.CH0040_3_d0127_ipe032_2_03.MG900135
5. B.US.2006.CH0040_3_d0127_ipe032_2_04.MG900136
6. B.US.2006.CH0040_3_d0127_ipe032_2_06.MG900137
7. B.US.2006.CH0040_3_d0127_ipe032_2_10.MG900138
8. B.US.2006.CH0040_3_d0127_ipe032_2_11.MG900139
9. B.US.2006.CH0040_3_d0127_ipe032_2_13.MG900140
10. B.US.2006.CH0040_3_d0127_ipe032_2_14.MG900141
11. B.US.2006.CH0040_3_d0127_ipe032_2_15.MG900142
12. B.US.2006.CH0040_3_d0127_ipe032_2_17.MG900143
13. B.US.2006.CH0040_3_d0127_ipe032_2_18.MG900144
14. B.US.2006.CH0040_3_d0127_ipe032_2_19.MG900145
15. B.US.2006.CH0040_3_d0127_ipe032_2_20.MG900146
16. B.US.2006.CH0040_3_d0127_ipe032_2_21.MG900147
17. B.US.2006.CH0040_3_d0127_ipe032_2_22.MG900148
18. B.US.2006.CH0040_3_d0127_ipe032_2_24.MG900149
19. B.US.2006.CH0040_3_d0127_ipe032_2_26.MG900150
20. B.US.2006.CH0040_3_d0127_ipe032_2_28.MG900151
21. B.US.2006.CH0040_3_d0127_ipe032_2_29.MG900152
22. B.US.2006.CH0040_3_d0127_ipe032_2_31.MG900153
23. B.US.2006.CH0040_3_d0127_ipe032_2_32.MG900154
24. B.US.2006.CH0040_3_d0127_ipe032_2_36.MG900155
25. B.US.2006.CH0040_3_d0127_ipe032_2_38.MG900156
26. B.US.2006.CH0040_3_d0127_ipe032_2_39.MG900157
27. B.US.2006.CH0040_3_d0127_ipe032_2_41.MG900158
28. B.US.2006.CH0040_3_d0127_ipe032_2_42.MG900159
29. B.US.2006.CH0040_3_d0127_ipe032_2_43.MG900160
30. B.US.2006.CH0040_3_d0127_ipe032_2_46.MG900161
31. B.US.2006.CH0040_3_d0127_ipe032_2_51.MG900162
32. B.US.2006.CH0040_3_d0127_ipe032_2_52.MG900163
33. B.US.2006.CH0040_3_d0127_ipe032_2_54.MG900164
34. B.US.2006.CH0040_3_d0127_ipe032_2_56.MG900165
35. B.US.2006.CH0040_3_d0127_ipe032_2_57.MG900166
36. B.US.2006.CH0040_3_d0127_ipe032_2_58.MG900167
37. B.US.2006.CH0040_3_d0127_ipe032_2_59.MG900168
38. B.US.2006.CH0040_3_d0127_ipe032_2_61.MG900169
39. B.US.2006.CH0040_3_d0127_ipe032_3_01.MG900170
40. B.US.2006.CH0040_3_d0127_ipe032_81_50.MG900171
41. B.US.2006.CH0040_3_d0148_ipe032_2_01.MG900172
42. B.US.2006.CH0040_3_d0148_ipe032_2_02.MG900173
43. B.US.2006.CH0040_3_d0148_ipe032_2_03.MG900174
44. B.US.2006.CH0040_3_d0148_ipe032_2_05.MG900175
45. B.US.2006.CH0040_3_d0148_ipe032_2_08.MG900176
46. B.US.2006.CH0040_3_d0148_ipe032_2_11.MG900177
47. B.US.2006.CH0040_3_d0148_ipe032_2_13.MG900178
48. B.US.2006.CH0040_3_d0148_ipe032_2_14.MG900179
49. B.US.2006.CH0040_3_d0148_ipe032_2_15.MG900180
50. B.US.2006.CH0040_3_d0148_ipe032_2_16.MG900181
51. B.US.2006.CH0040_3_d0148_ipe032_2_18.MG900182
52. B.US.2006.CH0040_3_d0148_ipe032_2_21.MG900183
53. B.US.2006.CH0040_3_d0148_ipe032_2_32.MG900184
54. B.US.2006.CH0040_3_d0148_ipe032_2_35.MG900185
55. B.US.2006.CH0040_3_d0148_ipe032_2_36.MG900186
56. B.US.2006.CH0040_3_d0148_ipe032_2_43.MG900187
57. B.US.2006.CH0040_3_d0148_ipe032_2_46.MG900188
58. B.US.2006.CH0040_3_d0148_ipe032_2_48.MG900189
59. B.US.2006.CH0040_3_d0148_ipe032_2_51.MG900190
60. B.US.2006.CH0040_3_d0148_ipe032_2_52.MG900191
61. B.US.2006.CH0040_3_d0148_ipe032_2_53.MG900192
62. B.US.2006.CH0040_3_d0148_ipe032_2_55.MG900193
63. B.US.2006.CH0040_3_d0148_ipe032_2_58.MG900194
64. B.US.2006.CH0040_3_d0148_ipe032_2_61.MG900195

**65.** B.US.2006.CH0040_3_d0148_ipe032_2_64.MG900196
**66.** B.US.2006.CH0040_3_d0148_ipe032_2_65.MG900197
**67.** B.US.2006.CH0040_3_d0148_ipe032_2_66.MG900198
**68.** B.US.2006.CH0040_3_d0148_ipe032_9_42.MG900199

* Consensus of sequences from day 127 and day 148 contained the identical E147K single change compared with CH040 TF Envs and these sequences were combined for reconstruction

MRVMGIRKNYQHLWREGILLLGILMICSAADNLWVTVYYGVPVWREATTTLFCASDAKAY

| | | |
|---|---|---|
| 1 | MRVMGIRKNYQHLWREGILLLGILMICSAADNLWVTVYYGVPVWREATTTLFCASDAKAY | 60 |
| 2 | MRVMGIRKNYQHLWREGILLLGILMICSAADNLWVTVYYGVPVWREATTTLFCASDAKAY | 60 |
| 3 | MRVMGIRKNYQHLWREGILLLGILMICSAADNLWVTVYYGVPVWREATTTLFCASDAKAY | 60 |
| 4 | MRVMGIRKNYQHLWREGILLLGILMICSAADNLWVTVYYGVPVWREATTTLFCASDAKAY | 60 |
| 5 | MRVMGIRKNYQHLWREGILLLGILMICSAADNLWVTVYYGVPVWREATTTLFCASDAKAY | 60 |
| 6 | MRVMGIRKNYQHLWREGILLLGILMICSAADNLWVTVYYGVPVWREATTTLFCASDAKAY | 60 |
| 7 | MRVMGIRKNYQHLWREGILLLGILMICSAADNLWVTVYYGVPVWREATTTLFCASDAKAY | 60 |
| 8 | MRVMGIRKNYQHLWREGILLLGILMICSAADNLWVTVYYGVPVWREATTTLFCASDAKAY | 60 |
| 9 | MRVMGIRKNYQHLWREGILLLGILMICSAADNLWVTVYYGVPVWREATTTLFCASDAKAY | 60 |
| 10 | MRVMGIRKNYQHLWREGILLLGILMICSAADNLWVTVYYGVPVWREATTTLFCASDAKAY | 60 |
| 11 | MRVMGIRKNYQHLWREGILLLGILMICSAADNLWVTVYYGVPVWREATTTLFCASDAKAY | 60 |
| 12 | MRVMGIRKNYQHLWREGILLLGILMICSAADNLWVTVYYGVPVWREATTTLFCASDAKAY | 60 |
| 13 | MRVMGIRKNYQHLWREGILLLGILMICSAADNLWVTVYYGVPVWREATTTLFCASDAKAY | 60 |
| 14 | MRVMGIRKNYQHLWREGILLLGILMICSAADNLWVTVYYGVPVWREATTTLFCASDAKAY | 60 |
| 15 | MRVMGIRKNYQHLWREGILLLGILMICSAADNLWVTVYYGVPVWREATTTLFCASDAKAY | 60 |
| 16 | MRVMGIRKNYQHLWREGILLLGILMICSAADNLWVTVYYGVPVWKEATTTLFCASDAKAY | 60 |
| 17 | MRVMGIRKNYQHLWREGILLLGILMICSAADNLWVTVYYGVPVWREATTTLFCASDAKAY | 60 |
| 18 | MRVMGIRKNYQHLWREGILLLGILMICSAADNLWVTVYYGVPVWREATTTLFCASDAKAY | 60 |
| 19 | MRVMGIRKNYQHLWREGILLLGILMICSAADNLWVTVYYGVPVWREATTTLFCASDAKAY | 60 |
| 20 | MRVMGIRKNYQHLWREGILLLGILMICSAADNLWVTVYYGVPVWREATTTLFCASDAKAY | 60 |
| 21 | MRVMGIRKNYQHLWREGILLLGILMICSAADNLWVTVYYGVPVWREATTTLFCASDAKAY | 60 |
| 22 | MRVMGIRKNYQHLWREGILLLGILMICSAADNLWVTVYYGVPVWREATTTLFCASDAKAY | 60 |
| 23 | MRVMGIRKNYQHLWREGILLLGILMICSAADNLWVTVYYGVPVWREATTTLFCASDAKAY | 60 |
| 24 | MRVMGIRKNYQHLWREGILLLGILMICSAADNLWVTVYYGVPVWREATTTLFCASDAKAY | 60 |
| 25 | MRVMGIRKNYQHLWREGILLLGILMICSAADNLWVTVYYGVPVWREATTTLFCASDAKAY | 60 |
| 26 | MRVMGIRKNYQHLWREGILLLGILMICSAADNLWVTVYYGVPVWREATTTLFCASDAKAY | 60 |
| 27 | MRVMGIRKNYQHLWREGILLLGILMICSAADNLWVTVYYGVPVWREATTTLFCASDAKAY | 60 |
| 28 | MRVMGIRKNYQHLWREGILLLGILMICSAADNLWVTVYYGVPVWREATTTLFCASDAKAY | 60 |
| 29 | MRVMGIRKNYQHLWREGILLLGILMICSAADNLWVTVYYGVPVWREATTTLFCASDAKAY | 60 |
| 30 | MRVMGIRKNYQHLWREGILLLGILMICSAADNLWVTVYYGVPVWREATTTLFCASDAKAY | 60 |
| 31 | MRVMGIRKNYQHLWREGILLLGILMICSAADNLWVTVYYGVPVWREATTTLFCASDAKAY | 60 |
| 32 | MRVMGIRKNYQHLWREGILLLGILMICSAADNLWVTVYYGVPVWREATTTLFCASDAKAY | 60 |
| 33 | MRVMGIRKNYQHLWREGILLLGILMICSAADNLWVTVYYGVPVWREATTTLFCASDAKAY | 60 |
| 34 | MRVMGIRKNYQHLWREGILLLGILMICSAADNLWVTVYYGVPVWREATTTLFCASDAKAY | 60 |
| 35 | MRVMGIRKNYQHLWREGILLLGILMICSAADNLWVTVYYGVPVWREATTTLFCASDAKAY | 60 |
| 36 | MRVMGIRKNYQHLWREGILLLGILMICSAADNLWVTVYYGVPVWREATTTLFCASDAKAY | 60 |
| 37 | MRVMGIRKNYQHLWREGILLLGILMICSAADNLWVTVYYGVPVWREATTTLFCASDAKAY | 60 |
| 38 | MRVMGIRKNYQHLWREGILLLGILMICSAADNLWVTVYYGVPVWREATTTLFCASDAKAY | 60 |
| 39 | MRVMGIRKNYQHLWREGILLLGILMICSAAGNLWVTVYYGVPVWREATTTLFCASDAKAY | 60 |
| 40 | MRVMGIRKNYQHLWREGILLLGILMICSAADNLWVTVYYGVPVWREATTTLFCASDAKAY | 60 |
| 41 | MRVMGIRKNYQHLWREGILLLGILMICSAADNLWVTVYYGVPVWREATTTLFCASDAKAY | 60 |
| 42 | MRVMGIRKNYQHLWREGILLLGILMICSAADNLWVTVYYGVPVWREATTTLFCASDAKAY | 60 |
| 43 | MRVMGIRKNYQHLWREGILLLGILMICSAADNLWVTVYYGVPVWREATTTLFCASDAKAY | 60 |
| 44 | MRVMGIRKNYQHLWREGILLLGILMICSAADNLWVTVYYGVPVWREATTTLFCASDAKAY | 60 |
| 45 | MRVMGIRKNYQHLWREGILLLGILMICSAADNLWVTVYYGVPVWREATTTLFCASDAKAY | 60 |
| 46 | MRVMGIRKNYQHLWREGILLLGILMICSAADNLWVTVYYGVPVWREATTTLFCASDAKAY | 60 |
| 47 | MRVMGIRKNYQHLWREGILLLGILMICSAADNLWVTVYYGVPVWREATTTLFCASDAKAY | 60 |
| 48 | MRVMGIRKNYQHLWREGILLLGILMICSAADNLWVTVYYGVPVWREATTTLFCASDAKAY | 60 |
| 49 | MRVMGIRKNYQHLWREGILLLGILMICSAADNLWVTVYYGVPVWREATTTLFCASDAKAY | 60 |
| 50 | MRVMGIRKNYQHLWREGILLLGILMICSAADNLWVTVYYGVPVWREATTTLFCASDAKAY | 60 |
| 51 | MRVMGIRKNYQHLWREGILLLGILMICSAADNLWVTVYYGVPVWREATTTLFCASDAKAY | 60 |
| 52 | MRVMGIRKNYQHLWREGILLLGILMICSAADNLWVTVYYGVPVWREATTTLFCASDAKAY | 60 |

| | | |
|---|---|---|
| 53 | MRVMGIRKNYQHLWREGILLLGILMICSAADNLWVTVYYGVPVWREATTTLFCASDAKAY | 60 |
| 54 | MRVMGIRKNYQHLWREGILLLGILMICSAADNLWVTVYYGVPVWREATTTLFCASDAKAY | 60 |
| 55 | MRVMGIRKNYQHLWREGILLLGILMICSAADNLWVTVYYGVPVWREATTTLFCASDAKAY | 60 |
| 56 | MRVMGIRKNYQHLWREGILLLGILMICSAADNLWVTVYYGVPVWREATTTLFCASDAKAY | 60 |
| 57 | MRVMGIRKNYQHLWREGILLLGILMICSAADNLWVTVYYGVPVWREATTTLFCASDAKAY | 60 |
| 58 | MRVMGIRKNYQHLWREGILLLGILMICSAADNLWVTVYYGVPVWREATTTLFCASDAKAY | 60 |
| 59 | MRVMGIRKNYQHLWREGILLLGILMICSAADNLWVTVYYGVPVWREATTTLFCASDAKAY | 60 |
| 60 | MRVMGIRKNYQHLWREGILLLGILMICSAADNLWVTVYYGVPVWREATTTLFCASDAKAY | 60 |
| 61 | MRVMGIRKNYQHLWREGILLLGILMICSAADNLWVTVYYGVPVWREATTTLFCASDAKAY | 60 |
| 62 | MRVMGIRKNYQHLWREGILLLGILMICSAADNLWVTVYYGVPVWREATTTLFCASDAKAY | 60 |
| 63 | MRVMGIRKNYQHLWREGILLLGILMICSAADNLWVTVYYGVPVWREATTTLFCASDAKAY | 60 |
| 64 | MRVMGIRKNYQHLWREGILLLGILMICSAADNLWVTVYYGVPVWREATTTLFCASDAKAY | 60 |
| 65 | MRVMGIRKNYQHLWREGILLLGILMICSAADNLWVTVYYGVPVWREATTTLFCASDAKAY | 60 |
| 66 | MRVMGIRKNYQHLWREGILLLGILMICSAADNLWVTVYYGVPVWREATTTLFCASDAKAY | 60 |
| 67 | MRVMGIRKNYQHLWREGILLLGILMICSAADNLWVTVYYGVPVWREATTTLFCASDAKAY | 60 |
| 68 | MRVMGIRKNYQHLWREGILLLGILMICSAADNLWVTVYYGVPVWREATTTLFCASDAKAY | 60 |

**DTEAHNVWATHACVPTDPNPQEVELKNVTENFNMWENNMVEQMHEDIISLWDQSLKPCVK**

| | | |
|---|---|---|
| 1 | DTEAHNVWATHACVPTDPNPQEVELKNVTENFNMWENNMVEQMHEDIISLWDQSLKPCVK | 120 |
| 2 | DTEAHNVWATHACVPTDPNPQEVELKNVTENFNMWENNMVEQMHEDIISLWDQSLKPCVK | 120 |
| 3 | DTEAHNVWATHACVPTDPNPQEVELKNVTENFNMWENNMVEQMHEDIISLWDQSLKPCVK | 120 |
| 4 | DTEAHNVWATHACVPTDPNPQEVELKNVTENFNMWENNMVEQMHEDIISLWDQSLKPCVK | 120 |
| 5 | DTEAHNVWATHACVPTDPNPQEVELKNVTENFNMWENNMVEQMHEDIISLWDQSLKPCVK | 120 |
| 6 | DTEAHNVWATHACVPTDPNPQEVELKNVTENFNMWENNMVEQMHEDIISLWDQSLKPCVK | 120 |
| 7 | DTEAHNVWATHACVPTDPNPQEVELKNVTENFNMWENNMVEQMHEDIISLWDQSLKPCVK | 120 |
| 8 | DTEAHNVWATHACVPTDPNPQEVELKNVTENFNMWENNMVEQMHEDIISLWDQSLKPCVK | 120 |
| 9 | DTEAHNVWATHACVPTDPNPQEVKLKNVTENFNMWENNMVEQMHEDIISLWDQSLKPCVK | 120 |
| 10 | DTEAHNVWATHACVPTDPNPQEVELKNVTENFNMWENNMVEQMHEDIISLWDQSLKPCVK | 120 |
| 11 | DTEAHNVWATHACVPTDPNPQEVELKNVTENFNMWENNMVEQMHEDIISLWDQSLKPCVK | 120 |
| 12 | DTEAHNVWATHACVPTDPNPQEVELKNVTENFNMWENNMVEQMHEDIISLWDQSLKPCVK | 120 |
| 13 | DTEAHNVWATHACVPTDPNPQEVELKNVTENFNMWENNMVEQMHEDIISLWDQSLKPCVK | 120 |
| 14 | DTEAHNVWATHACVPTDPNPQEVELKNVTENFNMWENNMVEQMHEDIISLWDQSLKPCVK | 120 |
| 15 | DTEAHNVWATHACVPTDPNPQEVELKNVTENFNMWENNMVEQMHEDIISLWDQSLKPCVK | 120 |
| 16 | DTEAHNVWATHACVPTDPNPQEVELKNVTENFNMWENNMVEQMHEDIISLWDQSLKPCVK | 120 |
| 17 | DTEAHNVWATHACVPTDPNPQEVELKNVTENFNMWENNMVEQMHEDIISLWDQSLKPCVK | 120 |
| 18 | DTEAHNVWATHACVPTDPNPQEVELKNVTENFNMWENNMVEQMHEDIISLWDQSLKPCVK | 120 |
| 19 | DTEAHNVWATHACVPTDPNPQEVELKNVTENFNMWENNMVEQMHEDIISLWDQSLKPCVK | 120 |
| 20 | DTEAHNVWATHACVPTDPNPQEVELKNVTENFNMWENNMVEQMHEDIISLWDQSLKPCVK | 120 |
| 21 | DTEAHNVWATHACVPTDPNPQEVELKNVTENFNMWENNMVEQMHEDIISLWDQSLKPCVK | 120 |
| 22 | DTEA---------------PQEVELKNVTENFNMWENNMVEQMHEDIISLWDQSLKPCVK | 105 |
| 23 | DTEAHNVWATHACVPTDPNPQEVELKNVTENFNMWENNMVEQMHEDIISLWDQSLKPCVK | 120 |
| 24 | DTEAHNVWATHACVPTDPNPQEVELKNVTENFNMWENNMVEQMHEDIISLWDQSLKPCVK | 120 |
| 25 | DTEAHNVWATHACVPTDPNPQEVELKNVTENFNMWENNMVEQMHEDIISLWDQSLKPCVK | 120 |
| 26 | DTEAHNVWATHACVPTDPNPQEVELKNVTENFNMWENNMVEQMHEDIISLWDQSLKPCVK | 120 |
| 27 | DTEAHNVWATHACVPTDPNPQEVELKNVTENFNMWENNMVEQMHEDIISLWDQSLKPCVK | 120 |
| 28 | DTEAHNVWATHACVPTDPNPQEVELKNVTENFNMWENNMVEQMHEDIISLWDQSLKPCVK | 120 |
| 29 | DTEAHNVWATHACVPTDPNPQEVELKNVTENFNMWENNMVEQMHEDITSLWDQSLKPCVK | 120 |
| 30 | DTEAHNVWATHACVPTDPNPQEVELKNVTENFNMWENNMVEQMHEDIISVWDQSLKPCVK | 120 |
| 31 | DTEAHNVWATHACVPTDPNPQEVELKNVTENFNMWENNMVEQMHEDIISLWDQSLKPCVK | 120 |
| 32 | DTEAHNVWATHACVPTDPNPQEVELKNVTENFNMWENNMVEQMHEDIISLWDQSLKPCVK | 120 |
| 33 | DTEAHNVWATHACVPTDPNPQEVELKNVTENFNMWENNMVEQMHEDIISLWDQSLKPCVK | 120 |
| 34 | DTEAHNVWATHACVPTDPNPQEVELKNVTENFNMWENNMVEQMHEDIISLWDQSLKPCVK | 120 |

35 DTEAHNVWATHACVPTDPNPQEVELKNVTENFNMWENNMVEQMHEDIISLWDQSLKPCVK 120
36 DTEAHNVWATHACVPTDPNPQEVELKNVTENFNMWENNMVEQMHEDIISLWDQSLKPCVK 120
37 DTEAHNVWATHACVPTDPNPQEVELKNVTENFNMWENNMVEQMHEDIISLWDQSLKPCVK 120
38 DTEAHNVWATHACVPTDPNPQEVELKNVTENFNMWENNMVEQMHEDIISLWDQSLKPCVK 120
39 DTEAHNVWATHACVPTDPNPQEVELKNVTENFNMWENNMVEQMHEDIISLWDQSLKPCVK 120
40 DTEAHNVWATHACVPTDPNPQEVELKNVTENFNMWENNMVEQMHEDIISLWDQSLKPCVK 120
41 DTEAHNVWATHACVPTDPNPQEVELKNVTENFNMWENNMAEQMHEDIISLWDQSLKPCVK 120
42 DTEAHNVWATHACVPTDPNPQEVELKNVTENFNMWENNMVEQMHEDIISLWDQSLKPCVK 120
43 DTEAHNVWATHACVPTDPNPQEVELKNVTENFNMWENNMVEQMHEDIISLWDQSLKPCVK 120
44 DTEAHNVWATHACVPTDPNPQEVGLKNVTENFNMWENNMVEQMHEDIISLWDQSLKPCVK 120
45 DTEAHNVWATHACVPTDPNPQEVELKNVTENFNMWENNMVEQMHEDIISLWDQSLKPCVK 120
46 DTEAHNVWATHACVPTDPNPQEVELKNVTENFNMWENNMVEQMHEDIISLWDQSLKPCVK 120
47 DTEAHNVWATHACVPTDPNPQEVELKNVTENFNMWENNMVEQMHEDIISLWDQSLKPCVK 120
48 DTEAHNVWATHACVPTDPNPQEVELKNVTENFNMWENNMVEQMHEDIISLWDQSLKPCVK 120
49 DTEAHNVWATHACVPTDPNPQEVELKNVTENFNMWENNMVEQMHEDIISLWDQSLKPCVK 120
50 DTEAHNVWATHACVPTDPNPQEVELKNVTENFNMWENNMVEQMHEDIISLWDQSLKPCVK 120
51 DTEAHNVWATHACVPTDPNPQEVELKNVTENFNMWENNMVEQMHEDIISLWDQSLKPCVK 120
52 DTEAHNVWATHACVPTDPNPQEVELKNVTENFNMWENNMVEQMHEDIISLWDQSLKPCVK 120
53 DTEAHNVWATHACVPTDPNPQEVELKNVTENFNMWENNMVEQMHEDIISLWDQSLKPCVK 120
54 DTEAHNVWATHACVPTDPNPQEVELKNVTENFNMWENNMVEQMHEDIISLWDQSLKPCVK 120
55 DTEAHNVWATHACVPTDPNPQEVELKNVTENFNMWENNMVEQMHEDIISLWDQSLKPCVK 120
56 DTEAHNVWATHACVPTDPNPQEVELKNVTENFNMWENNMVEQMHEDIISLWDQSLKPCVK 120
57 DTEAHNVWATHACVPTDPNPQEVELKNVTENFNMWENNMVEQMHEDIISLWDQSLKPCVK 120
58 DTEAHNVWATHACVPTDPNPQEVELKNVTENFNMWENNMVEQMHEDIISLWDQSLKPCVK 120
59 DTEAHNVWATHACVPTDPNPQEVELKNVTENFNMWENNMVEQMHEDIISLWDQSLKPCVK 120
60 DTEAHNVWATHACVPTDPNPQEVELKNVTENFNMWENNMVEQMHEDIISLWDQSLKPCVK 120
61 DTEAHNVWATHACVPTDPNPQEVELKNVTENFNMWENNMVEQMHEDIISLWDQSLKPCVK 120
62 DTEAHNVWATHACVPTDPNPQEVELKNVTENFNMWENNMVEQMHEDIISLWDQSLKPCVK 120
63 DTEAHNVWATHACVPTDPNPQEVELKNVTENFNMWENNMVEQMHEDIISLWDQSLKPCVK 120
64 DTEAHNVWATHACVPTDPNPQEVELKNVTENFNMWENNMVEQMHEDIISLWDQSLKPCVK 120
65 DTEAHNVWATHACVPTDPNPQEVELKNVTENFNMWENNMVEQMHEDIISLWDQSLKPCVK 120
66 DTEAHNVWATHACVPTDPNPQEVELKNVTENFNMWENNMVEQMHEDIISLWDQSLKPCVK 120
67 DTEAHNVWATHACVPTDPNPQEVELKNVTENFNMWENNMVEQMHEDIISLWDQSLKPCVK 120
68 DTEAHNVWATHACVPTDPNPQEVELKNVTENFNMWENNMVEQMHEDIISLWDQSLKPCVK 120

**LTPLCVTLNCTDLGNVTNTT---NSNGKMMEKGEVKNCSFKITTDIKDRTRKEYALFYKL**

1 LTPLCVTLNCTDLGNVTNTT---NSNGEMMEKGEVKNCSFKITTDIKDRTRKEYALFYKL 177
2 LTPLCVTLNCTDLGKVTNTT---NSNGKMMEKGDV-NCSFKITTDIKDRTRKEYALFYKL 176
3 LTPLCVTLNCTDLGNVTNTT---NSNGKMMEKGEVKNCSFKITTDIKDRTRKEYALFYKL 177
4 LTPLCVTLNCTDLGNVTNTT---NSNGKMMEKGEVKNCSFKITTDIKDRTRKEYALFYKL 177
5 LTPLCVTLNCTDLGNVTNTT---NSNGKMMEKGEVKNCSFKITTDIKDRTRKEYALFYKL 177
6 LTPLCVTLNCTDLGNVTNTT---NSNGKMMEKGEVKNCSFKITTDIKDRTRKEYALFYKL 177
7 LTPLCVTLNCTDLGNVTNTT---NSNGKMMEKGEVKNCSFKITTDIKDRTRKEYALFYKL 177
8 LTPLCVTLNCTDLGNVTNTT---NSNGGMMEKGEVKNCSFKITTDIKDRTRKEYALFYKL 177
9 LTPLCVTLNCTDLGNVTNTT---NSNGGMMEKGEVKNCSFKITTDIKDRTRKEYALFYKL 177
10 LTPLCVTLNCTDLGNVTNTT---NSNGKMMEKGEVKNCSFKITTDIKDRTRKEYALFYKL 177
11 LTPLCVTLNCTDLGNVTNTT---NSNGKMMEKGEVKNCSFKITTDIKDRTRKEYALFYKL 177
12 LTPLCVTLNCTDLGNVTNTT---NSNGKMMEKGEVKNCSFKITTDIKDRTRKEYALFYKL 177
13 LTPLCVTLNCTDLGNVTNTT---NSNGKMMEKGEVKNCSFKITTDIKDRTRKEYALFYKL 177
14 LTPLCVTLNCTDLGNVTNTT-----NSKMMEKGEVKNCSFKITTDIKDRTRKEYALFYKL 175
15 LTPLCVTLNCTDLGNVTNTT---NSNGKMMEKGEVKNCSFKITTDIKDRTRKEYALFY-- 175
16 LTPLCVTLNCTDLGNVTNTT---NSNGKMMEKGEVKNCSFKITTDIKDRTRKEYALFYKL 177

| | | |
|---|---|---|
| 17 | LTPLCVTLNCTDLGNVTNTT---NSNGKMMEKGEVKNCSFKITTDIKDRTRKEYALFYKL | 177 |
| 18 | LTPLCVTLNCTDLGNVTNTT---NSNGKMMEKGEVKNCSFKITTDIKDRTRKEYALFYKL | 177 |
| 19 | LTPLCVTLNCTDLGNVTNTT---NSNGKMMEKGEVKNCSFKITTDIKDRTRKEYALFYKL | 177 |
| 20 | LTPLCVTLNCTDLGNVTNTT-----NSKMMEKGEVKNCSFKITTDIKDRTRKEYALFYKL | 175 |
| 21 | LTPLCVTLNCTDLGNVTNTT---NSNGGMMEKGEVKNCSFKITTDIKDRTRKEYALFYKL | 177 |
| 22 | LTPLCVTLNCTDLGNVTNTT-----NSKMMEKGEVKNCSFKITTDIKDRTRKEYALFYKL | 160 |
| 23 | LTPLCVTLNCTDLGNVTNTT---NSNGKMMEKGEVKNCSFKITTDIKDRTRKEYALFYKL | 177 |
| 24 | LTPLCVTLNCTDLGKVTNTT---NSNGKMMEKGEVKNCSFKITTDIKDRTRKEYALFYKL | 177 |
| 25 | LTPLCVTLNCTDLGNVTNTT---NSNGKMMEKGEVKNCSFKITTDIKDRTRKEYALFYKL | 177 |
| 26 | LTPLCVTLNCTDLGNVTNTT---NSNGKMMEKGEVKNCSFKITTDIKDRTRKEYALFYKL | 177 |
| 27 | LTPLCVTLNCTDLGNVTNTT---NSNGKMMEKGEVKNCSFKITTDIKDRTRKEYALFYKL | 177 |
| 28 | LTPLCVTLNCTDLGNVTNTT---NSNGKMMEKGEVKNCSFKITTDIKDRTRKEYALFYKL | 177 |
| 29 | LTPLCVTLNCTDLGNVTNTT---NSNGKMMEKGEVKNCSFKITTDIKDRTRKEYALFYKL | 177 |
| 30 | LTPLCVTLNCTDLGNVTNTT---NSNGKMMEKGEVKNCSFKITTDIKDRTRKEYALFYKL | 177 |
| 31 | LTPLCVTLNCTDLGNVTNTT---NSNGKMMEKGEVKNCSFKITTDIKDRTRKEYALFYKL | 177 |
| 32 | LTPLCVTLNCTDLGNVTNTT---NSNGKMMEKGEVKNCSFKITTDIKDRTRKEYALFYKL | 177 |
| 33 | LTPLCVTLNCTDLGNVTNTT---NSNGKMMEKGEVKNCSFKITTDIKDRTRKEYALFYKL | 177 |
| 34 | LTPLCVTLNCTDLGNVTNTT---NSNGKMMEKGEVKNCSFKITTDIKDRTRKEYALFYKL | 177 |
| 35 | LTPLCVTLNCTDLGNVTNTT---NSNGKMMEKGEVKNCSFKITTDIKDRTRKEYALFYKL | 177 |
| 36 | LTPLCVTLNCTDLGKVTNTT---NSNGKMMEKGEVKNCSFKITTDIKDRTRKEYALFYKL | 177 |
| 37 | LTPLCVTLNCTDLGKVTNTT---NSNGKMMEKGEVKNCSFKITTDIKDRTRKEYALFYKL | 177 |
| 38 | LTPLCVTLNCTDLGNVTNTT-----NSKMMEKGEVKNCSFKITTDIKDRTRKEYALFYKL | 175 |
| 39 | LTPLCVTLNCTDLGNVTNTT---NSNREMMEKGEVKNCSFKITTDIKDRTRKEYALFYKL | 177 |
| 40 | LTPLCVTLNCTDLGNVTNTT---NSNGKMMEKGEVKNCSFKITTDIKDRTRKEYALFYKL | 177 |
| 41 | LTPLCVTLNCTDLGNVTNTT---NSNGKMMEKGEVKNCSFKITTDIKDRTRKEYALFYKL | 177 |
| 42 | LTPLCVTLNCTDLGNVTNTT---NSNGKMMEKGEVKNCSFKITTDIKDRTRKEYALFYKL | 177 |
| 43 | LTPLCVTLNCTDLGNVTNTT---NSNGKMMEKGEVKNCSFKITTDIKDRTRKEYALFYKL | 177 |
| 44 | LTPLCVTLNCTDLGNVTNTT---NSNGKMMEKGEVKNCSFKITTDIKDRTRKEYALFYKL | 177 |
| 45 | LTPLCVTLNCTDLGNVTNTT---NSNGKMMEKGEVKNCSFKITTDIKDRTRKEYALFYKL | 177 |
| 46 | LTPLCVTLNCTDLGNVTNTT---NSNGKMMEKGEVKNCSFKITTDIKDRTRKEYALFYKL | 177 |
| 47 | LTPLCVTLNCTDLGNVTNTT---NSNGGMMEKGEVKNCSFKITTDIKDRTRKEYALFYKL | 177 |
| 48 | LTPLCVTLNCTDLGNVTNTT---NSNGKMMEKGEVKNCSFKITTDIKDRTRKEYALFYKL | 177 |
| 49 | LTPLCVTLNCTDLGNVTNTT---NSNGKMMEKGEVKNCSFKITTDIKDRTRKEYALFYKL | 177 |
| 50 | LTPLCVTLNCTDLGNVTNTT---NSNGGMMEKGEVKNCSFKITTDIKDRTRKEYALFYKL | 177 |
| 51 | LTPLCVTLNCTDLGNVTNTT---NSNGKMMEKGEVKNCSFKITTDIKDRTRKEYALFYKL | 177 |
| 52 | LTPLCVTLNCTDLGNVTNTT---NSNGKMMEKGEVKNCSFKITTDIKDRTRKEYALFYKL | 177 |
| 53 | LTPLCVTLNCTDLGNVTNTT---NSNGGMMEKGEVKNCSFKITTDIKDRTRKEYALFYKL | 177 |
| 54 | LTPLCVTLNCTDLGNVTNTT---NSNGGRMEKGEVKNCSFKITTDIKDRTRKEYALFYKL | 177 |
| 55 | LTPLCVTLNCTDLGNVTNTT---NSNGKMMEKGEVKNCSFKITTDIKDRTRKEYALFYKL | 177 |
| 56 | LTPLCVTLNCTDLGNVTNTT---NSNGKMMEKGEVKNCSFKITTDIKDRTRKEYALFYKL | 177 |
| 57 | LTPLCVTLNCTDLGNVTNTT---NSNGGMMEKGEVKNCSFKITTDIKDRTRKEYALFYKL | 177 |
| 58 | LTPLCVTLNCTDLGNVTNTT---NSNGKMMEKGEVKNCSFKITTDIKDRTRKEYALFYKL | 177 |
| 59 | LTPLCVTLNCTDLGNVTNTT-----NSNGMEKGEVKNCSFKITTDIKDRTRKEYALFYKL | 175 |
| 60 | LTPLCVTLNCTDLGNVTNTT---NSNGKMMEKGEVKNCSFKITTDIKDRTRKEYALFYKL | 177 |
| 61 | LTPLCVTLNCTDLGNVTNTT---NSNGGMMEKGEVKNCSFKITTDIKDRTRKEYALFYKL | 177 |
| 62 | LTPLCVTLNCTDLGNVTNTT---NSNGKMMEKGEVKNCSFKITTDIKDRTRKEYALFYKL | 177 |
| 63 | LTPLCVTLNCTDLGNVTNTT---NSNGKMMEKGEVKNCSFKITTDIKDRTRKEYALFYKL | 177 |
| 64 | LTPLCVTLNCTDLGNVTNTT---NSNGKMMEKGEVKNCSFKITTDIKDRTRKEYALFYKL | 177 |
| 65 | LTPLCVTLNCTDLGNVTNTT---NSNGKMMEKGEVKNCSFKITTDIKDRTRKEYALFYKL | 177 |
| 66 | LTPLCVTLNCTDLGNVTNTT---NSNGKMMEKGEVKNCSFKITTDIKDRTRKEYALFYKL | 177 |
| 67 | LTPLCVTLNCTDLGNVTNTT-----NSNGEMMGEVKNCSFKITTDIKDRTRKEYALFYKL | 175 |
| 68 | LTPLCVTLNCTDLGNVTNTT---NSNGKMMEKGEVKNCSFKITTDIKDRTRKEYALFYKL | 177 |

| | | |
|---|---|---|
|  | DVVPIND-------TRYRLVSCNTSVITQACPKVSFEPIPIHYCAPAGFAILKCNDKQFI |  |
| 1 | DVVPIND-------TRYRLVSCNTSVITQACPKVSFEPIPIHYCAPAGFAILKCNDKQFI | 230 |
| 2 | DVVPIND-------TRYRLVSCNTSVITQACPKVSFEPIPIHYCAPAGFAILKCNDKQFI | 229 |
| 3 | DVVPIND-------TRYRLVSCNTSVITQACPKVSFEPIPIHYCAPAGFAILKCNDKQFI | 230 |
| 4 | DVVPIND-------TRYRLVSCNTSVITQACPKVSFEPIPIHYCAPAGFAILKCNDKQFI | 230 |
| 5 | DVVPIND-------TRYRLVSCNTSVITQACPKVSFEPIPIHYCAPAGFAILKCNDKQFI | 230 |
| 6 | DVVPIND-------TRYRLVSCNTSVITQACPKVSFEPIPIHYCAPAGFAILKCNDKQFI | 230 |
| 7 | DVVPIND-------TRYRLVSCNTSVITQACPKVSFEPIPIHYCAPAGFAILKCNDKQFI | 230 |
| 8 | DVVPIND-------TRYRLVSCNTSVITQACPKVSFEPIPIHYCAPAGFAILKCNDKQFI | 230 |
| 9 | DVVPIND-------TRYRLVSCNTSVITQACPKVSFEPIPIHYCAPAGFAILKCNDKQFI | 230 |
| 10 | DVVPIND-------TRYRLVSCNTSVITQACPKVSFEPIPIHYCAPAGFAILKCNDKQFI | 230 |
| 11 | DVVPIND-------TRYRLVSCNTSVITQACPKVSFEPIPIHYCAPAGFAILKCNDKQFI | 230 |
| 12 | DVVPIND-------TRYRLVSCNTSVITQACPKVSFEPIPIHYCAPAGFAILKCNDKQFI | 230 |
| 13 | DVVPIND-------TRYRLVSCNTSVITQACPKVSFEPIPIHYCAPAGFAILKCNDKQFI | 230 |
| 14 | DVVPIND-------TRYRLVSCNTSVITQACPKVSFEPIPIHYCAPAGFAILKCNDKQFI | 228 |
| 15 | -------------------------------------PIPIHYCAPAGFAILKCNDKQFI | 198 |
| 16 | DVVPIND-------TRYRLVSCNTSVITQACPKVSFEPIPIHYCAPAGFAILKCNDKQFI | 230 |
| 17 | DVVPIND-------TRYRLVSCNTSVITQACPKVSFEPIPIHYCAPAGFAILKCNDKQFI | 230 |
| 18 | DVVPIND-------TRYRLVSCNTSVITQACPKVSFEPIPIHYCAPAGFAILKCNDKQFI | 230 |
| 19 | DVVPIND-------TRYRLVSCNTSVITQACPKVSFEPIPIHYCAPAGFAILKCNDKQFI | 230 |
| 20 | DVVPIND-------TRYRLVSCNTSVITQACPKVSFEPIPIHYCAPAGFAILKCNDKQFI | 228 |
| 21 | DVVPIND-------TRYRLVSCNTSVITQACPKVSFEPIPIHYCAPAGFAILKCNDKQFI | 230 |
| 22 | DVVPIND-------TRYRLVSCNTSVITQACPKVSFEPIPIHYCAPAGFAILKCNDKQFI | 213 |
| 23 | DVVPIND-------TRYRLVSCNTSVITQACPKVSFEPIPIHYCAPAGFAILKCNDKQFI | 230 |
| 24 | DVVPIND-------TRYRLVSCNTSVITQACPKVSFEPIPIHYCAPAGFAILKCNDKQFI | 230 |
| 25 | DVVPIND-------TRYRLVSCNTSVITQACPKVSFEPIPIHYCAPAGFAILKCNDKQFI | 230 |
| 26 | DVVPIND-------TRYRLVSCNTSVITQACPKVSFEPIPIHYCAPAGFAILKCNDKQFI | 230 |
| 27 | DVVPIND-------TRYRLVSCNTSVITQACPKVSFEPIPIHYCAPAGFAILKCNDKQFI | 230 |
| 28 | DVVPIND-------TRYRLVSCNTSVITQACPKVSFEPIPIHYCAPAGFAILKCNDKQFI | 230 |
| 29 | DVVPIND-------TRYRLVSCNTSVITQACPKVSFEPIPIHYCAPAGFAILKCNDKQFI | 230 |
| 30 | DVVPIND-------TRYRLVSCNTSVITQACPKVSFEPIPIHYCAPAGFAILKCNDKQFI | 230 |
| 31 | DVVPIND-------TRYRLVSCNTSVITQACPKVSFEPIPIHYCAPAGFAILKCNDKQFI | 230 |
| 32 | DVVPIND-------TRYRLVSCNTSVITQACPKVSFEPIPIHYCAPAGFAILKCNDKQFI | 230 |
| 33 | DVVPIND-------TRYRLVSCNTSVITQACPKVSFEPIPIHYCAPAGFAILKCNDKQFI | 230 |
| 34 | DVVPIND-------TRYRLVSCNTSVITQACPKVSFEPIPIHYCAPAGFAILKCNDKQFI | 230 |
| 35 | DVVPIND-------TRYRLVSCNTSVITQACPKVSFEPIPIHYCAPAGFAILKCNDKQFI | 230 |
| 36 | DVVPIND-------TRYRLVSCNTSVITQACPKVSFEPIPIHYCAPAGFAILKCNDKQFI | 230 |
| 37 | DVVPIND-------TRYRLVSCNTSVITQACPKVSFEPIPIHYCAPAGFAILKCNDKQFI | 230 |
| 38 | DVVPIND-------TRYRLVSCNTSVITQACPKVSFEPIPIHYCAPAGFAILKCNDKQFI | 228 |
| 39 | DVVPIND-------TRYRLVSCNTSVITQACPKVSFEPIPIHYCAPAGFAILKCNDKQFI | 230 |
| 40 | DVVPIND-------TRYRLVSCNTSVITQACPKVSFEPIPIHYCAPAGFAILKCNDKQFI | 230 |
| 41 | DVVPIND-------TRYRLVSCNTSVITQACPKVSFEPIPIHYCAPAGFAILKCNDKQFI | 230 |
| 42 | DVVPIND-------TRYRLVSCNTSVITQACPKVSFEPIPIHYCAPAGFAILKCNDKQFI | 230 |
| 43 | DVVPIND-------TRYRLVSCNTSVITQACPKVSFEPIPIHYCAPAGFAILKCNDKQFI | 230 |
| 44 | DVVPIND-------TRYRLVSCNTSVITQACPKVSFEPIPIHYCAPAGFAILKCNDKQFI | 230 |
| 45 | DVVPIND-------TRYRLVSCNTSVITQACPKVSFEPIPIHYCAPAGFAILKCNDKQFI | 230 |
| 46 | DVVPIND-------TRYRLVSCNTSVITQACPKVSFEPIPIHYCAPAGFAILKCNDKQFI | 230 |
| 47 | DVVPIND-------TRYRLVSCNTSVITQACPKVSFEPIPIHYCAPAGFAILKCNDKQFI | 230 |
| 48 | DVVPIND-------TRYRLVSCNTSVITQACPKVSFEPIPIHYCAPAGFAILKCNDKQFI | 230 |
| 49 | DVVPIND-------TRYRLVSCNTSVITQACPKVSFEPIPIHYCAPAGFAILKCNDKQFI | 230 |
| 50 | DVVPIDD-------TRYRLVSCNTSVITQACPKVSFEPIPIHYCAPAGFAILKCNDKQFI | 230 |
| 51 | DVVPIND-------TRYRLVSCNTSVITQACPKVSFEPIPIHYCAPAGFAILKCNDKQFI | 230 |
| 52 | DVVPIND-------TRYRLVSCNTSVITQACPKVSFEPIPIHYCAPAGFAILKCNDKQFI | 230 |
| 53 | DVVPIND-------TRYRLVSCNTSVITQACPKVSFEPIPIHYCAPAGFAILKCNDKQFI | 230 |

| | | |
|---|---|---|
| 54 | DVVPIND-------TRYRLVSCNTSVITQACPKVSFEPIPIHYCAPAGFAILKCNDKQFI | 230 |
| 55 | DVVPIND-------TRYRLVSCNTSVITQACPKVSFEPIPIHYCAPAGFAILKCNDKQFI | 230 |
| 56 | DVVPIND-------TRYRLVSCNTSVITQACPKVSFEPIPIHYCAPAGFAILKCNDKQFI | 230 |
| 57 | DVVPIND-------TRYRLVSCNTSVITQACPKVSFEPIPIHYCAPAGFAILKCNDKQFI | 230 |
| 58 | DVVPIND-------TRYRLVSCNTSVITQACPKVSFEPIPIHYCAPAGFAILKCNDKQFI | 230 |
| 59 | DVVPIND-------TRYRLVSCNTSVITQACPKVSFEPIPIHYCAPAGFAILKCNDKQFI | 228 |
| 60 | DVVPIND-------TRYRLVSCNTSVITQACPKVSFEPIPIHYCAPAGFAILKCNDKQFI | 230 |
| 61 | DVVPIND-------TRYRLVSCNTSVITQACPKISFEPIPIHYCAPAGFAILKCNDKQFI | 230 |
| 62 | DVVPIND-------TRYRLVSCNTSVITQACPKVSFEPIPIHYCAPAGFAILKCNDKQFI | 230 |
| 63 | DVVPIND-------TRYRLVSCNTSVITQACPKVSFEPIPIHYCAPAGFAILKCNDKQFI | 230 |
| 64 | DVVPIND-------TRYRLVSCNTSVITQACPKVSFEPIPIHYCAPAGFAILKCNDKQFI | 230 |
| 65 | DVVPIND-------TRYRLVSCNTSVITQACPKVSFEPIPIHYCAPAGFAILKCNDKQFI | 230 |
| 66 | DVVPIND-------TRYRLVSCNTSVITQACPKVSFEPIPIHYCAPAGFAILKCNDKQFI | 230 |
| 67 | DVVPIND-------TRYRLVSCNTSVITQACPKVSFEPIPIHYCAPAGFAILKCNDKQFI | 228 |
| 68 | DVVPIND-------TRYRLVSCNTSVITQACPKVSFEPIPIHYCAPAGFAILKCNDKQFI | 230 |

| | | |
|---|---|---|
| | **GTGPCTNVSTVQCTHGIRPVVSTQLLLNGSLAEEEVVIRSVNFSDNAKTIIVQLNKSVEI** | |
| 1 | GTGPCTNVSTVQCTHGIRPVVSTQLLLNGSLAEEEVVIRSVNFSDNAKTIIVQLNKSVEI | 290 |
| 2 | GTGPCTNVSTVQCTHGIRPVVSTQLLLNGSLAEEEVVIRSVNFSDNAKTIIVQLNKSVEI | 289 |
| 3 | GTGPCTNVSTVQCTHGIRPVVSTQLLLNGSLAEEEVVIRSVNFSDNAKTIIVQLNKSVKI | 290 |
| 4 | GTGPCTNVSTVQCTHGIRPVVSTQLLLNGSLAEEEVVIRSVNFSDNAKTIIVQLNKSVEI | 290 |
| 5 | GTGPCTNVSTVQCTHGIRPVVSTQLLLNGSLAEEEVVIRSVNFSDNAKTIIVQLNKSVKI | 290 |
| 6 | GTGPCTNVSTVQCTHGIRPVVSTQLLLNGSLAEEEVVIRSVNFSDNAKTIIVQLNKSVEI | 290 |
| 7 | GTGPCTNVSTVQCTHGIRPVVSTQLLLNGSLAEEEVVIRSVNFSDNAKTIIVQLNKSVEI | 290 |
| 8 | GTGPCTNVSTVQCTHGIRPVVSTQLLLNGSLAEEEVVIRSVNFSDNAKTIIVQLNKSVEI | 290 |
| 9 | GTGPCTNVSTVQCTHGIRPVVSTQLLLNGSLAEEEVVIRSVNFSDNAKTIIVQLNKSVEI | 290 |
| 10 | GTGPCTNVSTVQCTHGIRPVVSTQLLLNGSLAEEEVVIRSVNFSDNAKTIIVQLNKSVEI | 290 |
| 11 | GTGPCTNVSTVQCTHGIRPVVSTQLLLNGSLAEEEVVIRSVNFSDNAKTIIVQLNKSVEI | 290 |
| 12 | GTGPCTNVSTVQCTHGIRPVVSTQLLLNGSLAEEEVVIRSVNFSDNAKTIIVQLNKSVEI | 290 |
| 13 | GTGPCTNVSTVQCTHGIRPVVSTQLLLNGSLAEEEVVIRSVNFSDNAKTIIVQLNKSVEI | 290 |
| 14 | GTGPCTNVSTVQCTHGIRPVVSTQLLLNGSLAEEEVVIRSVNFSDNAKTIIVQLNKSVEI | 288 |
| 15 | GTGPCTNVSTVQCTHGIRPVVSTQLLLNGSLAEEEVVIRSVNFSDNAKTIIVQLNKSVKI | 258 |
| 16 | GTGPCTNVSTVQCTHGIRPVVSTQLLLNGSLAEEEVVIRSVNFSDNAKTIIVQLNKSVEI | 290 |
| 17 | GTGPCTNVSTVQCTHGIRPVVSTQLLLNGSLAEEEVVIRSVNFSDNAKTIIVQLNKSVKI | 290 |
| 18 | GTGPCTNVSTVQCTHGIRPVVSTQLLLNGSLAEEEVVIRSVNFSDNAKTIIVQLNKSVEI | 290 |
| 19 | GTGPCTNVSTVQCTHGIRPVVSTQLLLNGSLAEEEVVIRSVNFSDNAKTIIVQLNKSVEI | 290 |
| 20 | GTGPCTNVSTVQCTHGIRPVVSTQLLLNGSLAEEEVVIRSVNFSDNAKTIIVQLNKSVEI | 288 |
| 21 | GTGPCTNVSTVQCTHGIRPVVSTQLLLNGSLAEEEVVIRSVNFSDNAKTIIVQLNKSVEI | 290 |
| 22 | GTGPCTNVSTVQCTHGIRPVVSTQLLLNGSLAEEEVVIRSVNFSDNAKTIIVQLNKSVEI | 273 |
| 23 | GTGPCTNVSTVQCTHGIRPVVSTQLLLNGSLAEEEVVIRSVNFSDNAKTIIVQLNKSVEI | 290 |
| 24 | GTGPCTNVSTVQCTHGIRPVVSTQLLLNGSLAEEEVVIRSVNFSDNAKTIIV-LNKSVEI | 289 |
| 25 | GTGPCTNVSTVQCTHGIRPVVSTQLLLNGSLAEEEVVIRSVNFSDNAKTIIVQLNKSVEI | 290 |
| 26 | GTGPCTNVSTVQCTHGIRPVVSTQLLLNGSLAEEEVVIRSVNFSDNAKTIIVQLNKSVEI | 290 |
| 27 | GTGPCTNVSTVQCTHGIRPVVSTQLLLNGSLAEEEVVIRSVNFSDNAKTIIVQLNKSVKI | 290 |
| 28 | GTGPCTNVSTVQCTHGIRPVVSTQLLLNGSLAEEEVVIRSVNFSDNAKTIIVQLNKSVKI | 290 |
| 29 | GTGPCTNVSTVQCTHGIRPVVSTQLLLNGSLAEEEVVIRSVNFSDNAKTIIVQLNKSVKI | 290 |
| 30 | GTGPCTNVSTVQCTHGIRPVVSTQLLLNGSLAEEEVVIRSVNFSDNAKTIIVQLNKSVKI | 290 |
| 31 | GTGPCTNVSTVQCTHGIRPVVSTQLLLNGSLAEEEVVIRSVNFSDNAKTIIVQLNKSVEI | 290 |
| 32 | GTGPCTNVSTVQCTHGIRPVVSTQLLLNGSLAEEEVVIRSVNFSDNAKTIIVQLNKSVEI | 290 |
| 33 | GTGPCTNVSTVQCTHGIRPVVSTQLLLNGSLAEEEVVIRSVNFSDNAKTIIVQLNKSVEI | 290 |
| 34 | GTGPCTNVSTVQCTHGIRPVVSTQLLLNGSLAEEEVVIRSVNFSDNAKTIIVQLNKSVKI | 290 |
| 35 | GTGPCTNVSTVQCTHGIRPVVSTQLLLNGSLAEEEVVIRSVNFSDNAKTIRVQLNKSVEI | 290 |

| | | |
|---|---|---|
| 36 | GTGPCTNVSTVQCTHGIRPVVSTQLLLNGSLAEEEVVIRSVNFSDNAKTIIVQLNKSVEI | 290 |
| 37 | GTGPCTNVSTVQCTHGIRPVVSTQLLLNGSLAEEEVVIRSVNFSDNAKTIIVQLNKSVEI | 290 |
| 38 | GTGPCTNVSTVQCTHGIRPVVSTQLLLNGSLAEEEVVIRSVNFSDNAKTIIVQLNKSVEI | 288 |
| 39 | GTGPCTNVSTVQCTHGIRPVVSTQLLLNGSLAEEEVVIRSVNFSDNAKTIIVQLNKSVEI | 290 |
| 40 | GTGPCTNVSTVQCTHGIRPVVSTQLLLNGSLAEEEVVIRSVNFSDNAKTIIVQLNKSVKI | 290 |
| 41 | GTGPCTNVSTVQCTHGIRPVVSTQLLLNGSLAEEEVVIRSVNFSDNAKTIIVQLNKSVKI | 290 |
| 42 | GTGPCTNVSTVQCTHGIRPVVSTQLLLNGSLAEEEVVIRSVNFSDNAKTIIVQLNKSVEI | 290 |
| 43 | GTGPCTNVSTVQCTHGIRPVVSTQLLLNGSLAEEEVVIRSVNFSDNAKTIIVQLNKSVEI | 290 |
| 44 | GTGPCTNVSTVQCTHGIRPVVSTQLLLNGSLAEEEVVIRSVNFSDNAKTIIVQLNKSVEI | 290 |
| 45 | GTGPCTNVSTVQCTHGIRPVVSTQLLLNGSLAEEEVVIRSVNFSDNAKTIIVQLNKSVEI | 290 |
| 46 | GTGPCTNVSTVQCTHGIRPVVSTQLLLNGSLAEEEVVIRSVNFSDNAKTIIVQLNKSVKI | 290 |
| 47 | GTGPCTNVSTVQCTHGIRPVVSTQLLLNGSLAEEEVVIRSVNFSDNAKTIIVQLNKSVEI | 290 |
| 48 | GTGPCTNVSTVQCTHGIRPVVSTQLLLNGSLAEEEVVIRSVNFSDNAKTIIVQLNKSVEI | 290 |
| 49 | GTGPCTNVSTVQCTHGIRPVVSTQLLLNGSLAEEEVVIRSVNFSDNAKTIIVQLNKSVEI | 290 |
| 50 | GTGPCTNVSTVQCTHGIRPVVSTQLLLNGSLAEEEVVIRSVNFSDNAKTIIVQLNKSVEI | 290 |
| 51 | GTGPCTNVSTVQCTHGIRPVVSTQLLLNGSLAEEEVVIRSVNFSDNAKTIIVQLNKSVEI | 290 |
| 52 | GTGPCTNVSTVQCTHGIRPVVSTQLLLNGSLAEEEVVIRSVNFSDNAKTIIVQLNKSVEI | 290 |
| 53 | GTGPCTNVSTVQCTHGIRPVVSTQLLLNGSLAEEEVVIRSVNFSDNAKTIIVQLNKSVEI | 290 |
| 54 | GTGPCTNVSTVQCTHGIRPVVSTQLLLNGSLAEEEVVIRSVNFSDNAKTIIVQLNKSVEI | 290 |
| 55 | GTGPCTNVSTVQCTHGIRPVVSTQLLLNGSLAEEEVVIRSVNFSDNAKTIIVQLNKSVEI | 290 |
| 56 | GTGPCTNVSTVQCTHGIRPVVSTQLLLNGSLAEGEVVIRSVNFSDNAKTIIVQLNKSVEI | 290 |
| 57 | GTGPCTNVSTVQCTHGIRPVVSTQLLLNGSLAEEEVVIRSVNFSDNAKTIIVQLNKSVEI | 290 |
| 58 | GTGPCTNVSTVQCTHGIRPVVSTQLLLNGSLAEEEVVIRSVNFSDNAKTIIVQLNKSVEI | 290 |
| 59 | GTGPCTNVSTVQCTHGIRPVVSTQLLLNGSLAEEEVVIRSVNFSDNAKTIIVQLNKSVEI | 288 |
| 60 | GTGPCTNVSTVQCTHGIRPVVSTQLLLNGSLAKEEVVIRSVNFSDNAKTIIVQLNKSVEI | 290 |
| 61 | GTGPCTNVSTVQCTHGIRPVVSTQLLLNGSLAEEEVVIRSVNFSDNAKTIIVQLNKSVEI | 290 |
| 62 | GTGPCTNVSTVQCTHGIRPVVSTQLLLNGSLAEEEVVIRSVNFSDNAKTIIVQLNKSVEI | 290 |
| 63 | GTGPCTNVSTVQCTHGIRPVVSTQLLLNGSLAEEEVVIRSVNFSDNAKTIIVQLNKSVEI | 290 |
| 64 | GTGPCTNVSTVQCTHGIRPVVSTQLLLNGSLAEEEVVIRSVNFSDNAKTIIVQLNKSVEI | 290 |
| 65 | GTGPCTNVSTVQCTHGIRPVVSTQLLLNGSLAEEEVVIRSVNFSDNAKTIIVQLNKSVEI | 290 |
| 66 | GTGPCTNVSTVQCTHGIRPVVSTQLLLNGSLAEEEVVIRSVNFSDNAKTIIVQLNKSVEI | 290 |
| 67 | GTGPCTNVSTVQCTHGIRPVVSTQLLLNGSLAEEEVVIRSVNFSDNAKTIIVQLNKSVEI | 288 |
| 68 | GTGPCTNVSTVQCTHGIRPVVSTQLLLNGSLAEEEVVIRSVNFSDNAKTIIVQLNKSVEI | 290 |

| | | |
|---|---|---|
| | **TCTRPNNNTRKSIPMGPGKAFYARGDITGDIRKAYCEINGTEWHSTLKLVVEKLREQY-N** | |
| 1 | TCTRPNNNTRKSIPMGPGKAFYARGDITGDIRKAYCEINGTEWHSTLKLVVEKLREQY-N | 349 |
| 2 | TCTRPNNNTRKSIPMGPGKAFYARGDITGDIRKAYCEINGTEWHSTLKLVVEKLREQY-N | 348 |
| 3 | TCTRPNNNTRKSIPMGPGKAFYARGDITGDIRKAYCEINGTEWHSTLKLVVEKLREQY-N | 349 |
| 4 | TCTRPNNNTRKSIPMGPGKAFYARGDITGDIRKAYCEINGTEWHSTLKLVVEKLREQY-N | 349 |
| 5 | TCTRPNNNTRKSIPMGPGKAFYARGDITGDIRKAYCEINGTEWHSTLKLVVEKLREQY-N | 349 |
| 6 | TCTRPNNNTRKSIPMGPGKAFYARGDITGDIRKAYCEINGTEWHSTLKLVVEKLREQY-N | 349 |
| 7 | TCTRPNNNTRKSIPIGPGKAFYARGDITGDIRKTYCEINGTEWHSTLKLVVEKLREQY-N | 349 |
| 8 | TCTRPNNNTRKSIPMGPGKAFYARGDITGDIRKAYCEINGTEWHSTLKLVVEKLREQY-N | 349 |
| 9 | TCTRPNNNTRKSIPMGPGKAFYARGDITGDIRKAYCEINGTEWHSTLKLVVEKLREQY-N | 349 |
| 10 | TCTRPNNNTRKSIPMGPGKAFYARGDITGDIRKAYCEINGTEWHSTLKLVVEKLREQY-N | 349 |
| 11 | TCTRPNNNTRKSIPMGPGKAFYARGDITGDIRKAYCEINGTEWHSTLKLVVEKLREQY-N | 349 |
| 12 | TCTRPNNNTRKSIPMGPGKAFYARGDITGDIRRAYCEINGTEWHSTLKLVVEKLREQY-N | 349 |
| 13 | TCTRPNNNTRKSIPMGPGKAFYARGDITGDIRKAYCEINGTEWHSTLKLVVEKLREQY-N | 349 |
| 14 | TCTRPNNNTRKSIPMGPGKAFYARGDITGDIRKAYCEINGTEWHSTLKLVVEKLREQY-N | 347 |
| 15 | TCTRPNNNTRKSIPMGPGKAFYARGDITGDIRKAYCEINGTEWHSTLKLVVEKLREQY-N | 317 |
| 16 | TCTRPNNNTRKSIPMGPGKAFYARGDITGDIRKAYCEINGTEWHSTLKLVVEKLREQY-N | 349 |
| 17 | TCTRPNNNTRKSIPMGPGKAFYARGDITGDIRKAYCEINGTEWHSTLKLVVEKLREQY-N | 349 |

| | | |
|---|---|---|
| 18 | TCTRPNNNTRKSIPMGPGKAFYARGDITGDIRKAYCEINGTEWHSTLKLVVEKLREQY-N | 349 |
| 19 | TCTRPNNNTRKSIPMGPGKAFYARGDITGDIRKAYCEINGTEWHSTLKLVVEKLREQY-N | 349 |
| 20 | TCTRPNNNTRKSIPMGPGKAFYARGDITGDIRKAYCEINGTEWHSTLKLVVEKLREQY-N | 347 |
| 21 | TCTRPNNNTRKSIPMGPGKAFYARGDITGDIRKAYCEINGTEWHSTLKLVVEKLREQY-N | 349 |
| 22 | TCTRPNNNTRKSIPMGPGKAFYARGDITGDIRKAYCEINGTEWHSTLKLVVEKLREQY-N | 332 |
| 23 | TCTRPNNNTRKSIPMGPGKAFYARGDITGDIRKAYCEINGTEWHSTLKLVVEKLREQY-N | 349 |
| 24 | TCTRPNNNTRKSIPMGPGKAFYARGDITGDIRKAYCEINGTEWHSTLKLVVEKLREQY-N | 348 |
| 25 | TCTRPNNNTRKSIPMGPGKAFYARGDITGDIRKAYCEINGTEWHSTLKLVVEKLREQY-N | 349 |
| 26 | TCTRPNNNTRKSIPMGPGKAFYARGDITGDIRKAYCEINGTEWHSTLKLVVEKLREQY-N | 349 |
| 27 | TCTRPNNNTRKSIPMGPGKAFYARGDITGDIRKAYCEINGTEWHSTLKLVVEKLREQY-N | 349 |
| 28 | TCTRPNNNTRKSIPMGPGKAFYARGDITGDIRKAYCEINGTEWHSTLKLVVEKLREQY-N | 349 |
| 29 | TCTRPNNNTRKSIPMGPGKAFYARGDITGDIRKAYCEINGTEWHSTLKLVVEKLREQY-N | 349 |
| 30 | TCTRPNNNTRKSIPMGPGKAFYARGDITGDIRKAYCEINGTEWHSTLKLVVEKLREQY-N | 349 |
| 31 | TCTRPNNNTRKSIPMGPGKAFYARGDITGDIRKAYCEINGTEWHSTLKLVVEKLREQY-N | 349 |
| 32 | TCTRPNNNTRKSIPMGPGKAFYARGDITGDIRKAYCEINGTEWHSTLKLVVEKLREQY-N | 349 |
| 33 | TCTRPNNNTRKSIPMGPGKAFYARGDITGDIRKAYCEINGTEWHSTLKLVVEKLREQY-N | 349 |
| 34 | TCTRPNNNTRKSIPMGPGKAFYARGDITGDIRKAYCEINGTEWHSTLKLVVEKLREQY-N | 349 |
| 35 | TCTRPNNNTRKSIPMGPGKAFYARGDITGDIRKAYCEINGTEWHSTLKLVVEKLREQY-N | 349 |
| 36 | TCTRPNNNTRKSIPMGPGKAFYARGDITGDIRKAYCEINGTEWHSTLELVVEKLREQY-N | 349 |
| 37 | TCTRPNNNTRKSIPMGPGKAFYARGDITGDIRKAYCEINGTEWHSTLKLVVEKLREQY-N | 349 |
| 38 | TCTRPNNNTRKSIPMGPGKAFYARGDITGDIRKAYCEINGTEWHSTLKLVVEKLREQY-N | 347 |
| 39 | TCTRPNNNTRKSIPMGPGKAFYARGDITGDIRKASCEINGTEWHSTLKLVVEKLREQY-N | 349 |
| 40 | TCTRPNNNTRKSIPMGPGKAFYARGDITGDIRKAYCEINGTEWHSTLKLVVEKLREQY-N | 349 |
| 41 | TCTRPNNNTRKSIPMGPGKAFYARGDITGDIRKAYCEINGTEWHSTLKLVVEKLREQY-N | 349 |
| 42 | TCTRPNNNTRKSIPMGPGKAFYARGDITGDIRKAYCEINGTEWHSTLKLVVEKLREQY-N | 349 |
| 43 | TCTRPNNNTRKSIPMGPGKAFYARGDITGDIRKAYCEINGTEWHSTLKMVVEKLREQY-N | 349 |
| 44 | TCTRPNNNTRKSIPMGPGKAFYARGDITGDIRKAYCEINGTEWHSTLKLVVEKLREQY-N | 349 |
| 45 | TCTRPNNNTRKSIPMGPGKAFYARGDITGDIRKAYCEINGTEWHSTLKLVVEKLREQY-N | 349 |
| 46 | TCTRPNNNTRKSIPMGPGKAFYARGDITGDIRKAYCEINGTEWHSTLKLVVEKLREQY-N | 349 |
| 47 | TCTRPNNNTRKSIPMGPGKAFYARGDITGDIRKAYCEINGTEWHSTLKLVVEKLREQY-N | 349 |
| 48 | TCTRPNNNTRKSIPMGPGKAFYARGDITGDIRKAYCEINGTEWHSTLKLVVEKLREQY-N | 349 |
| 49 | NCTRPNNNTRKSIPMGPGKAFYARGDITGDIRKAYCEINGTEWHSTLKLVVEKLREQY-N | 349 |
| 50 | TCTRPNNNTRKSIPMGPGKAFYARGDITGDIKKACCEINGTEWHSTLKLVVEKLREQY-N | 349 |
| 51 | TCIRPNNNTRKSIPMGPGKAFYARGDITGDIKKAYCEINGTEWHSTLKLVVEKLREQY-N | 349 |
| 52 | TCTRPNNNTRKSIPMGPGKAFYARGDITGDIRKAYCEINGTEWHSTLKLVVEKLREQY-N | 349 |
| 53 | TCTRPNNNTRKSIPMGPGKAFYARGDITGDIKKAYCEINGTEWHSTLKLVVEKLREQY-N | 349 |
| 54 | TCTRPNNNTRKSIPMGPGKAFYARGDITGDIRKAYCKINGTEWHSTLKLVVKKLREQY-N | 349 |
| 55 | TCTRPNNNTRKSIPMGPGKAFYARGDITGDIRKAYCEINGTEWHSTLKLVVEKLREQY-N | 349 |
| 56 | TCTRPNNNTRKSIPMGPGKAFYARGDITGDIRKAYCEINGTEWHSTLKLVVEKLREQY-N | 349 |
| 57 | TCTRPNNNTRKSIPMGPGKAFYARGDITGDIRKAYCEINGTEWHSTLKLVVEKLREQY-N | 349 |
| 58 | TCTRPNNNTRKSIPMGPGKAFYARGDITGDIRKAYCEINGTEWHSTLKLVVEKLREQY-N | 349 |
| 59 | TCTRPNNNTRKSIPMGPGKAFYARGDITGDIRKAYCEINGTEWHSTLKLVVEKLREQY-N | 347 |
| 60 | TCTRPNNNTRKSIPMGPGKAFYARGDITGDIRKAYCEINGTEWHSTLKLVVEKLREQY-N | 349 |
| 61 | TCTRPNNNTRKSIPMGPGKAFYARGDITGDIRKAYCGINGTEWHSTLKLVVEKLREQY-N | 349 |
| 62 | NCTRPNNNTRKSIPMGPGKAFYARGDITGDIRKAYCEINGTEWHSTLKLVVEKLREQY-N | 349 |
| 63 | TCTRPNNNTRKSIPMGPGKAFYARGDITGDIRKAYCKINGTEWHSTLKLVVEKLREQY-N | 349 |
| 64 | TCTRPNNNTRKSIPMGPGKAFYARGDITGDIRKAYCEINGTEWHSTLKLVVEKLREQY-N | 349 |
| 65 | TCTRPNNNTRKSIPMGPGKAFYARGDITGDIRKAYCEINGTEWHSTLKLVVEKLREQY-N | 349 |
| 66 | TCTRPNNNTRKSIPMGPGKAFYARGDITGDIRKAYCEINGTEWHSTLKLVVEKLREQY-N | 349 |
| 67 | TCTRPNNNTRKSIPMGPGKAFYARGDITGDIRKAYCEINGTEWHSTLKLVVEKLREQY-N | 347 |
| 68 | TCTRPNNNTRKSIPMGPGKAFYARGDITGDIRKAYCEINGTEWHSTLKLVVEKLREQY-N | 349 |

| | | |
|---|---|---|
| | KTIVFNRSSGGDPEIVMYSFNCGGEFFYCNSTKLFNSTWPWND-TKGSHDTNGTLILPCK | |
| 1 | KTIVFNRSSGGDPEIVMYSFNCGGEFFYCNSTKLFNSTWPWND-TKGSHDTNGTLILPCK | 408 |
| 2 | KTIVFNRSSGGDPEIVMYSFNCGGEFFYCNSTRLFNSTWPWND-TKGSHDTNGTLILPCK | 407 |
| 3 | KTIVFNRSSGGDPEIVMYSFNCGGEFFYCNSTKLFNSTWPWND-TKGSHDTNGTLILPCK | 408 |
| 4 | KTIVFNRSSGGDPEIVMYSFNCGGEFFYCNSTKLFNSTWPWND-TKGSHDTNGTLILPCK | 408 |
| 5 | KTIVFNRSSGGDPEIVMYSFNCGGEFFYCNSTKLFNSTWPWND-TKGSHDTNGTLILPCK | 408 |
| 6 | KTIVFNRSSGGDPEIVMYSFNCGGEFFYCNSTKLFNSTWPWND-TKGSHDTNGTLILPCK | 408 |
| 7 | KTIVFNRSSGGDPEIVMYSFNCGGEFFYCNSTKLFNSTWPWND-TKGSHDTNGTLILPCK | 408 |
| 8 | KTIVFNRSSGGDPEIVMYSFNCGGEFFYCNSTKLFNSTWPWND-TKGSHDTNGTLILPCK | 408 |
| 9 | KTIVFNRSSGGDPEIVMYSFNCGGEFFYCNSTKLFNSTWPNDT--KGSHDTNDTLILPCK | 407 |
| 10 | KTIVFNRSSGGDPEIVMYSFNCGGEFFYCNSTKLFNSTWPWND-TKGSHDTNGTLILPCK | 408 |
| 11 | KTIVFNRSSGGDPEIVMYSFNCGGEFFYCNSTKLFNSTWPWND-TKGSHDTNGTLILPCK | 408 |
| 12 | KTIVFNRSSGGDPEIVMYSFNCGGEFFYCNSTKLFNSTWPWND-TKGSHDTNGTLILPCK | 408 |
| 13 | KTIVFNRSSGGDPEIVMYSFNCGGEFFYCNSTKLFNSTWPWND-TKGSHDTNGTLILPCK | 408 |
| 14 | KTIVFNRSSGGDPEIVMYSFNCGGEFFYCNSTKLFNSTWPWND-TKGSHDTNGTLILPCK | 406 |
| 15 | KTIVFNRSSGGDPEIVMYSFNCGGEFFYCNSTKLFNSTWPWND-TKGSHDTNGTLILPCK | 376 |
| 16 | KTIVFNRSSGGDPEIVMYSFNCGGEFFYCNSTKLFNSTWPWND-TKGSHDTNGTLILPCK | 408 |
| 17 | KTIVFNRSSGGDPEIVMYSFNCGGEFFYCNSTKLFNSTWPWND-TKGSHDTNGTLILPCK | 408 |
| 18 | KTIVFNRSSGGDPEIVMYSFNCGGEFFYCNSTKLFNSTWPWND-TKGSHDTNGTLILPCK | 408 |
| 19 | KTIVFNRSSGGDPEIVMYSFNCGGEFFYCNSTKLFNSTWPWND-TKGSHDTNGTLILPCK | 408 |
| 20 | KTIVFNRSSGGDPEIVMYSFNCGGEFFYCNSTKLFNSTWPWND-TKGSHDTNGTLILPCK | 406 |
| 21 | KTIVFNRSSGGDPEIVMYSFNCGGEFFYCNSTKLFNSTWPWND-TKGSHDTNGTLRLPCK | 408 |
| 22 | KTIVFNRSSGGDPEIVMYSFNCGGEFFYCNSTKLFNSTWPWND-TKGSHDTNGTLILPCK | 391 |
| 23 | KTIVFNRSSGGDPEIVMYSFNCGGEFFYCNSTKLFNSTWPWND-TKGSHDTNGTLILPCK | 408 |
| 24 | KTIVFNRSSGGDPEIVMYSFNCGGEFFYCNSTKLFNSTWPWND-TKGSHDTNGTLILPCK | 407 |
| 25 | KTIVFNRSSGGDPEIVMYSFNCGGEFFYCNSTKLFNSTWPWND-TKGSHDTNGTLILPCK | 408 |
| 26 | KTIVFNRSSGGDPEIVMYSFNCGGEFFYCNSTKLFNSTWPWND-TKGSHDTNGTLILPCK | 408 |
| 27 | KTIVFNRSSGGDPEIVMYSFNCGGEFFYCNSTKLFNSTWPWND-TKGSHDTNGTLILPCK | 408 |
| 28 | KTIVFNRSSGGDPEIVMYSFNCGGEFFYCNSTKLFNSTWPWND-TKGSHDTNGTLILPCK | 408 |
| 29 | KTIVFNRSSGGDPEIVMYSFNCGGEFFYCNSTKLFNSTWPWND-TKGSHDTNGTLILPCK | 408 |
| 30 | KTIVFNRSSGGDPEIVMYSFNCGGEFFYCNSTKLFNSTWPWND-TKGSHDTNGTLILPCK | 408 |
| 31 | KTIVFNRSSGGDPEIVMYSFNCGGEFFYCNSTKLFNSTWPWND-TKGSHDTNGTLILPCK | 408 |
| 32 | KTIVFNRSSGGDPEIVMYSFNCGGEFFYCNSTKLFNSTWPWND-TKGSHDTNGTLILPCK | 408 |
| 33 | KTIVFNRSSGGDPEIVMYSFNCGGEFFYCNSTKLFNSTWPWND-TKGSHDTNGTLILPCK | 408 |
| 34 | KTIVFNRSSGGDPEIVMYSFNCGGEFFYCNSTKLFNSTWPWND-TKGSHDTNGTLILPCK | 408 |
| 35 | KTIVFNRSSGGDPEIVMYSFNCGGEFFYCNSTKLFNSTWPWND-TKGSHDTNGTLILPCK | 408 |
| 36 | KTIVFNRSSGGDPEIVMYSFNCGGEFFYCNSTKLFNSTWHWND-TKGSHDTNGTLILPCK | 408 |
| 37 | KTIVFNRSSGGDPEIVMYSFNCGGEFFYCNSTKLFNSTWPWND-TKGSHDTNGTLILPCK | 408 |
| 38 | KTIVFNRSSGGDPEIVMYSFNCGGEFFYCNSTKLFNSTWPWND-TKGSHDTNGTLILPCK | 406 |
| 39 | KTIVFNRSSGGDPEIVMYSFNCGGEFFYCNSTKLFNSTWPWND-TKGSHDTNGTLILPCK | 408 |
| 40 | KTIVFNRSSGGDPEIVMYSFNCGGEFFYCNSTKLFNSTWPWND-TKGSHDTNGTLILPCK | 408 |
| 41 | KTIVFNRSSGGDPEIVMYSFNCGGEFFYCNSTKLFNSTWPWND-TKGSHDTNGTLILPCK | 408 |
| 42 | KTIVFNRSSGGDPEIVMYSFNCGGEFFYCNSTKLFNSTWPWND-TKGSHDTNDTLMLPCK | 408 |
| 43 | KTIVFNRSSGGDPEIVMYSFNCGGEFFYCNSTKLFNSTWPWND-TKGSHDTNDTLMLPCK | 408 |
| 44 | KTIVFNRSSGGDPEIVMYSFNCGGEFFYCNSTKLFNSTWPWND-TKGSHDTNGTLTLPCK | 408 |
| 45 | KTIVFNRSSGGDPEIVMYSFNCGGEFFYCNSTKLFNSTWPWND-TKGSHDTNDTLMLPCK | 408 |
| 46 | KTIVFNRSSGGDPEIVMYSFNCGGEFFYCNSTKLFNSTWPWND-TKGSHDTNGTLILPCK | 408 |
| 47 | KTIVFNRSSGGDPEIVMYSFNCGGEFFYCNSTKLFNSTWPWND-TKGSHDTNGTLTLPCK | 408 |
| 48 | KTIVFNRSSGGDPEIVMYSFNCGGEFFYCNSTKLFNSTWPWND-TKGSHDTNDTLMLPCK | 408 |
| 49 | KTIVFNRSSGGDPEIVMYSFNCGGEFFYCNSTKLFNSTWLWND-TKGSHDTNGTLILPCK | 408 |
| 50 | KTIVFNRSSGGDPEIVMYSFNCGGEFFYCNSTKLFNSTWPWND-TKGSHDTNGTLILPCK | 408 |
| 51 | KTIVFNRSSGGDPEIVMYSFNCGGEFFYCNSTKLFNSTWPWND-TKGSHDTNGTLILPCK | 408 |
| 52 | KTIVFNRSSGGDPEIVMYSFNCGGEFFYCNSTKLFNSTWPWND-TKGSHDTNDTLMLPCK | 408 |
| 53 | KTIVFNRSSGGDPEIVMYSFSCGGEFFYCNSTKLFNSTWPWND-TKGSHDTNGTLRLPCK | 408 |
| 54 | KTIVFNRSSGGDPEIVMYSFNCGGEFFYCNSTKLFNSTWPWND-TKGSHDTNGTLILPCK | 408 |

| | | |
|---|---|---|
| 55 | KTIVFNRSSGGDPEIVMYSFNCGGEFFYCNSTKLFNSTWPWND-TKGSHDTNGTLILPCK | 408 |
| 56 | KTIVFNRSSGGDPEIVMYSFNCGGEFFYCNSTKLFNSTWPWND-TKGSHDTNDTLMLPCK | 408 |
| 57 | KTIVFNRSSGGDPEIVMYSFNCGGEFFYCNSTKLFNSTWPWND-TKGSHDTNGTLTLPCK | 408 |
| 58 | KTIVFNRSSGGDPEIVMYSFNCGGEFFYCNSTKLFNSTWPWND-TKGSHDTNGTLKLPCK | 408 |
| 59 | KTIVFNRSSGGDPEIVMYSFNCGGEFFYCNSTKLFNSTWPWND-TKGSHDTNGTLMLPCK | 406 |
| 60 | KTIVFNRSSGGDPEIVMYSFNCGGEFFYCNSTKLFNSTWPWND-TKGSHDTNDTLMLPCK | 408 |
| 61 | KTIVFNRSSGGDPEIVMYSFNCGGEFFYCNSTKLFNSTWPWND-TKGSHDTNGTLILPCK | 408 |
| 62 | KTIVFNRSSGGDPEIVMYSFNCGGEFFYCNSTKLFNSTWPWND-TKGSHDTNGTLILPCK | 408 |
| 63 | KTIVFNRSSGGDPEIVMYSFNCGGEFFYCNSTKLFNSTWPWND-TKGSHDTNGTLILPCK | 408 |
| 64 | KTIVFNRSSGGDPEIVMYSFNCGGEFFYCNSTKLFNSTWPWND-TKGSHDTNDTLMLPCK | 408 |
| 65 | KTIVFNRSSGGDPEIVMYSFNCGGEFFYCNSTKLFNSTWPWND-TKGSHDTNDTLMLPCK | 408 |
| 66 | KTIVFNRSSGGDPEIVMYSFNCGGEFFYCNSTKLFNSTWPWND-TKGSHDTNDTLMLPCK | 408 |
| 67 | KTIVFNRSSGGDPEIVMYSFNCGGEFFYCNSTKLFNSTWPWND-TKGSHDTNGTLILPCK | 406 |
| 68 | KTIVFNRSSGGDPEIVMYSFNCGGEFFYCNSTKLFNSTWPWND-TKGSHDTNDTLMLPCK | 408 |

| | | |
|---|---|---|
| | **IKQIINMWQGVGKAMYAPPIEGKIRCSSNITGLLLTRDGG-YESNETDEIFRPGGGDMRD** | |
| 1 | IKQIINMWQGVGKAMYAPPIEGKIRCSSNITGLLLTRDGG-YESNETDEIFRPGGGDMRD | 467 |
| 2 | IKQIINMWQGVGKAMYAPPIEGKIRCSSNITGLLLTRDGG-YESNETDEIFRPGGGDMRD | 466 |
| 3 | IKQIINMWQGVGKAMYAPPIEGKIRCSSNITGLLLTRDGG-YESNETDEIFRPGGGDMRD | 467 |
| 4 | IKQIINMWQGVGKAMYAPPIEGKIRCSSNITGLLLTRDGG-YESNETDEIFRPGGGDMRD | 467 |
| 5 | IKQIINMWQGVGKAMYAPPIEGKIRCSSNITGLLLTRDGG-YESNETDEIFRPGGGDMRD | 467 |
| 6 | IKQIINMWQGVGKAMYAPPIEGKIRCSSNITGLLLTRDGG-YESNETDEIFRPGGGDMRD | 467 |
| 7 | IKQIINMWQGVGKAMYAPPIEGKIRCSSNITGLLLTRDGG-YESNETDEIFRPGGGDMRD | 467 |
| 8 | IKQIINMWQGVGKAMYAPPIEGKIRCSSNITGLLLTRDGG-YESNETDEIFRPGGGDMRD | 467 |
| 9 | IKQIINMWQGVGKAMYAPPIEGKIRCSSNITGLLLTRDGG-YESNETDEIFRPGGGDMRD | 466 |
| 10 | IKQIINMWQGVGKAMYAPPIEGKIRCSSNITGLLLTRDGG-YESNETDEIFRPGGGDMRD | 467 |
| 11 | IKQIINMWQGVGKAMYAPPIEGKIRCSSNITGLLLTRDGG-YESNETDEIFRPGGGDMRD | 467 |
| 12 | IKQIINMWQGVGKAMYAPPIEGKIRCSSNITGLLLTRDGG-YESNETDEIFRPGGGDMRD | 467 |
| 13 | IKQIINMWQGVGKAMYAPPIEGKIRCSSNITGLLLTRDGG-YESNETDEIFRPGGGDMRD | 467 |
| 14 | IKQIINMWQGVGKAMYAPPIEGKIRCSSNITGLLLTRDGG-YESNETDEIFRPGGGDMRD | 465 |
| 15 | IKQIINMWQGVGKAMYAPPIEGKIRCSSNITGLLLTRDGG-YESNETDEIFRPGGGDMRD | 435 |
| 16 | IKQIINMWQGVGKAMYAPPIEGKIRCSSNITGLLLTRDGG-YESNETDEIFRPGGGDMRD | 467 |
| 17 | IKQIINMWQGVGKAMYAPPIEGKIRCSSNITGLLLTRDGG-YESNETDEIFRPGGGDMRD | 467 |
| 18 | IKQIINMWQGVGKAMYAPPIEGKIRCSSNITGLLLTRDGG-YESNETDEIFRPGGGDMRD | 467 |
| 19 | IKQIINMWQGVGKAMYAPPIEGKIRCSSNITGLLLTRDGG-YESNETDEIFRPGGGDMRD | 467 |
| 20 | IKQIINMWQGVGKAMYAPPIEGKIRCSSNITGLLLTRDGG-YESNETDEIFRPGGGDMRD | 465 |
| 21 | IKQIINMWQGVGKAMYAPPIEGKIRCSSNITGLLLTRDGG-YESNETDEIFRPGGGDMRD | 467 |
| 22 | IKQIINMWQGVGKAMYAPPIEGKIRCSSNITGLLLTRDGG-YESNETDEIFRPGGGDMRD | 450 |
| 23 | IKQIINMWQGVGKAMYAPPIEGKIRCSSNITGLLLTRDGG-YESNETDEIFRPGGGDMRD | 467 |
| 24 | IKQIINMWQGVGKAMYAPPIEGKIRCSSNITGLLLTRDGG-YESNETDEIFRPGGGDMRD | 466 |
| 25 | IKQIINMWQGVGKAMYAPPIEGKIRCSSNITGLLLTRDGG-YESNETDEIFRPGGGDMRD | 467 |
| 26 | IKQIINMWQGVGKAMYAPPIEGKIRCSSNITGLLLTRDGG-YESNETDEIFRPGGGDMRD | 467 |
| 27 | IKQIINMWQGVGKAMYAPPIEGKIRCSSNITGLLLTRDGG-YESNETDEIFRPGGGDMRD | 467 |
| 28 | IKQIINMWQGVGKAMYAPPIEGKIRCSSNITGLLLTRDGG-YESNETDEIFRPGGGDMRD | 467 |
| 29 | IKQIINMWQGVGKAMYAPPIEGKIRCSSNITGLLLTRDGG-YESNETDEIFRPGGGDMRD | 467 |
| 30 | IKQIINMWQGVGKAMYAPPIEGKIRCSSNITGLLLTRDGG-YESNETDEIFRPGGGDMRD | 467 |
| 31 | IKQIINMWQGVGKAMYAPPIEGKIRCSSNITGLLLTRDGG-YESNETDEIFRPGGGDMRD | 467 |
| 32 | IKQIINMWQGVGKAMYAPPIEGKIRCSSNITGLLLTRDGG-YESNETDEIFRPGGGDMRD | 467 |
| 33 | IKQIINMWQGVGKAMYAPPIEGKIRCSSNITGLLLTRDGG-YESNETDEIFRPGGGDMRD | 467 |
| 34 | IKQIINMWQGVGKAMYAPPIEGKIRCSSNITGLLLTRDGG-YESNETDEIFRPGGGDMRD | 467 |
| 35 | IKQIINMWQGVGKAMYAPPIEGKIRCSSNITGLLLTRDGG-YESNETDEIFRPGGGDMRD | 467 |
| 36 | IKQIINMWQGVGKAMYAPPIEGKIRCSSNITGLLLTRDGG-YESNETDEIFRPGGGDMRD | 467 |

| | | |
|---|---|---|
| 37 | IKQIINMWQGVGKAMYAPPIEGKIRCSSNITGLLLTRDGG-YESNETDEIFRPGGGDMRD | 467 |
| 38 | IKQIINMWQGVGKAMYAPPIEGKIRCSSNITGLLLTRDGG-YESNETDEIFRPGGGDMRD | 465 |
| 39 | IKQIINMWQGVGKAMYAPPIEGKIRCSSNITGLLLTRDGG-YESNETDEIFRPGGGDMRD | 467 |
| 40 | IKQIINMWQGVGKAMYAPPIEGKIRCSSNITGLLLTRDGG-YESNETDEIFRPGGGDMRD | 467 |
| 41 | IKQIINMWQGVGKAMYAPPIEGKIRCSSNITGLLLTRDGG-YESNETDEIFRPGGGDMRD | 467 |
| 42 | IKQIINMWQGVGKAMYAPPIEGKIRCSSNITGLLLTRDGG-YESNETDEIFRPGGGDMRD | 467 |
| 43 | IKQIINMWQGVGKAMYAPPIEGKIRCSSNITGLLLTRDGG-YESNETDEIFRPGGGDMRD | 467 |
| 44 | IKQIINMWQGVGKAMYAPPIEGKIRCSSNITGLLLTRDGG-YESNETDEIFRPGGGDMRD | 467 |
| 45 | IKQIINMWQGVGKAMYAPPIEGKIRCSSNITGLLLTRDGG-YESNETDEIFRPGGGDMRD | 467 |
| 46 | IKQIINMWQGVGKAMYAPPIEGKIRCSSNITGLLLTRDGG-YESNETDEIFRPGGGDMRD | 467 |
| 47 | IKQIINMWQGVGKAMYAPPIEGKIRCSSNITGLLLTRDGG-YESNETDEIFRPGGGDMRD | 467 |
| 48 | IKQIINMWQGVGKAMYAPPIEGKIRCSSNITGLLLTRDGG-YESNETDEIFRPGGGDMRD | 467 |
| 49 | IKQIINMWQGVGKAMYAPPIEGKIRCSSNITGLLLTRDGG-YESNETDEIFRPGGGDMRD | 467 |
| 50 | IKQIINMWQGVGKAMYAPPIEGKIRCSSNITGLLLTRDGG-YESNETDEIFRPGGGDMRD | 467 |
| 51 | IKQIINMWQGVGKAMYAPPIEGKIRCSSNITGLLLTRDGG-YESNETDEIFRPGGGDMRD | 467 |
| 52 | IKQIINMWQGVGKAMYAPPIEGKIRCSSNITGLLLTRDGG-YESNETDEIFRPGGGDMRD | 467 |
| 53 | IKQIINMWQGVGKAMYAPPIEGKIRCSSNITGLLLTRDGG-YESNETDEIFRPGGGDMRD | 467 |
| 54 | IKQIINMWQGVGKAMYAPPIEGKIRCSSNITGLLLTRDGG-YESNETDEIFRPGGGDMRD | 467 |
| 55 | IKQIINMWQGVGKAMYAPPIEGKIRCSSNITGLLLTRDGG-YESNETDEIFRPGGGDMRD | 467 |
| 56 | IKQIINMWQGVGKAMYAPPIEGKIRCSSNITGLLLTRDGG-YESNETDEIFRPGGGDMRD | 467 |
| 57 | IKQIINMWQGVGKAMYAPPIEGKIRCSSNITGLLLTRDGG-YESNETDEIFRPGGGDMRD | 467 |
| 58 | IKQIINMWQGVGKAMYAPPIEGKIRCSSNITGLLLTRDGG-YESNETDEIFRPGGGDMRD | 467 |
| 59 | IKQIINMWQGVGKAMYAPPIEGKIRCSSNITGLLLTRDGG-YESNETDEIFRPGGGDMRD | 465 |
| 60 | IKQIINMWQGVGKAMYAPPIEGKIRCSSNITGLLLTRDGG-YESNETDEIFRPGGGDMRD | 467 |
| 61 | IKQIINMWQGVGKAMYAPPIEGKIRCSSNITGLLLTRDGG-YESNETDEIFRPGGGDMRD | 467 |
| 62 | IKQIINMWQGVGKAMYAPPIEGKIRCSSNITGLLLTRDGG-YESNETDEIFRPGGGDMRD | 467 |
| 63 | IKQIINMWQGVGKAMYAPPIEGKIRCSSNITGLLLTRDGG-YESNETDEIFRPGGGDMRD | 467 |
| 64 | IKQIINMWQGVGKAMYAPPIEGKIRCSSNITGLLLTRDGG-YESNETDEIFRPGGGDMRD | 467 |
| 65 | IKQIINMWQGVGKAMYAPPIEGKIRCSSNITGLLLTRDGG-YESNETDEIFRPGGGDMRD | 467 |
| 66 | IKQIINMWQGVGKAMYAPPIEGKIRCSSNITGLLLTRDGG-YESNETDEIFRPGGGDMRD | 467 |
| 67 | IKQIINMWQGVGKAMYAPPIEGKIRCSSNITGLLLTRDGG-YESNETDEIFRPGGGDMRN | 465 |
| 68 | IKQIINMWQGVGKAMYAPPIEGKIRCSSNITGLLLTRDGG-YESNETDEIFRPGGGDMRD | 467 |

**NWRSELYKYKVVKIEPLGVAPTKAKRRVVQREKRAFG-LGAVFLGFLGAAGSTMGAASIT**

| | | |
|---|---|---|
| 1 | NWRSELYKYKVVKIEPLGVAPTKAKRRVVQREKRAFG-LGAVFLGFLGAAGSTMGAASIT | 526 |
| 2 | NWRSELYKYKVVKIEPLGVAPTKAKRRVVQREKRAFG-LGAVFLGFLGAAGSTMGAASIT | 525 |
| 3 | NWRSELYKYKVVKIEPLGVAPTKAKRRVVQREKRAFG-LGAVFLGFLGAAGSTMGAASIT | 526 |
| 4 | NWRSELYKYKVVKIEPLGVAPTKAKRRVVQREKRAFG-LGAVFLGFLGAAGSTMGAASIT | 526 |
| 5 | NWRSELYKYKVVKIEPLGVAPTKAKRRVVQREKRAFG-LGAVFLGFLGAAGSTMGAASIT | 526 |
| 6 | NWRSELYKYKVVKIEPLGVAPTKAKRRVVQREKRAFG-LGAVFLGFLGAAGSTMGAASIT | 526 |
| 7 | NWRSELYKYKVVKIEPLGVAPTKAKRRVVQREKRAFG-LGAVFLGFLGAAGSTMGAASIT | 526 |
| 8 | NWRSELYKYKVVKIEPLGVAPTKAKRRVVQREKRAFG-LGAVFLGFLGAAGSTMGAASIT | 526 |
| 9 | NWRSELYKYKVVKIEPLGVAPTKAKRRVVQREKRAFG-LGAVFLGFLGAAGSTMGAASIT | 525 |
| 10 | NWRSELYKYKVVKIEPLGVAPTKAKRRVVQREKRAFG-LGAVFLGFLGAAGSTMGAASIT | 526 |
| 11 | NWRSELYKYKVVKIEPLGVAPTKAKRRVVQREKRAFG-LGAVFLGFLGAAGSTMGAASIT | 526 |
| 12 | NWRSELYKYKVVKIEPLGVAPTKAKRRVVQREKRAFG-LGAVFLGFLGAAGSTMGAASIT | 526 |
| 13 | NWRSELYKYKVVKIEPLGVAPTKAKRRVVQREKRAFG-LGAVFLGFLGAAGSTMGAASIT | 526 |
| 14 | NWRSELYKYKVVKIEPLGVAPTKAKRRVVQREKRAFG-LGAVFLGFLGAAGSTMGAASIT | 524 |
| 15 | NWRSELYKYKVVKIEPLGVAPTKAKRRVVQREKRAFG-LGAVFLGFLGAAGSTMGAASIT | 494 |
| 16 | NWRSELYKYKVVKIEPLGVAPTKAKRRVVQREKRAFG-LGAVFLGFLGAAGSTMGAASIT | 526 |
| 17 | NWRSELYKYKVVKIEPLGVAPTKAKRRVVQREKRAFG-LGAVFLGFLGAAGSTMGAASIT | 526 |
| 18 | NWRSELYKYKVVKIEPLGVAPTKAKRRVVQREKRAFG-LGAVFLGFLGAAGSTMGAASIT | 526 |

| | | |
|---|---|---|
| 19 | NWRSELYKYKVVKIEPLGVAPTKAKRRVVQREKRAFG-LGAVFLGFLGAAGSTMSAASIT | 526 |
| 20 | NWRSELYKYKVVKIEPLGVAPTKAKRRVVQREKRAFG-LGAVFLGFLGAAGSTMGAASIT | 524 |
| 21 | NWRSELYKYKVVKIEPLGVAPTKAKRRVVQREKRAFG-LGAVFLGFLGAAGSTMGAASIT | 526 |
| 22 | NWRSELYKYKVVKIEPLGVAPTKAKRRVVQREKRAFG-LGAVFLGFLGAAGSTMGAASIT | 509 |
| 23 | NWRSELYKYKVVKIEPLGVAPTKAKRRVVQREKRAFG-LGAVFLGFLGAAGSTMGAASIT | 526 |
| 24 | NWRSELYKYKVVKIEPLGVAPTKAKRRVVQREKRAFG-LGAVFLGFLGAAGSTMGAASIT | 525 |
| 25 | NWRSELYKYKVVKIEPLGVAPTKAKRRVVQREKRAFG-LGAVFLGFLGAAGSTMGAASIT | 526 |
| 26 | NWRSELYKYKVVKIEPLGVAPTKAKRRVVQREKRAFG-LGAVFLGFLGAAGSTMGAASIT | 526 |
| 27 | NWRSELYKYKVVKIEPLGVAPTKAKRRVVQREKRAFG-LGAVFLGFLGAAGSTMGAASIT | 526 |
| 28 | NWRSELYKYKVVKIEPLGVAPTKAKRRVVQREKRAFG-LGAVFLGFLGAAGSTMGAASIT | 526 |
| 29 | NWRSELYKYKVVKIEPLGVAPTKAKRRVVQREKRAFG-LGAVFLGFLGAAGSTMGAASIT | 526 |
| 30 | NWRSELYKYKVVKIEPLGVAPTKAKRRVVQREKRAFG-LGAVFLGFLGAAGSTMGAASIT | 526 |
| 31 | NWRSELYKYKVVKIEPLGVAPTKAKRRVVQREKRAFG-LGAVFLGFLGAAGSTMGAASIT | 526 |
| 32 | NWRSELYKYKVVKIEPLGVAPTKAKRRVVQREKRAFG-LGAVFLGFLGAAGSTMGAASIT | 526 |
| 33 | NWRSELYKYKVVKIEPLGVAPTKAKRRVVQREKRAFG-LGAVFLGFLGAAGSTMGAASIT | 526 |
| 34 | NWRSELYKYKVVKIEPLGVAPTKAKRRVVQREKRAFG-LGAVFLGFLGAAGSTMGAASIT | 526 |
| 35 | NWRSELYKYKVVKIEPLGVAPTKAKRRVVQREKRAFG-LGAVFLGFLGAAGSTMGAASIT | 526 |
| 36 | NWRSELYKYKVVKIEPLGVAPTKAKRRVVQREKRAFG-LGAVFLGFLGAAGSTMGAASIT | 526 |
| 37 | NWRSELYKYKVVKIEPLGVAPTKAKRRVVQREKRAFG-LGAVFLGFLGAAGSTMGAASIT | 526 |
| 38 | NWRSELYKYKVVKIEPLGVAPTKAKRRVVQREKRAFG-LGAVFLGFLGAAGSTMGAASIT | 524 |
| 39 | NWRSELYKYKVVKIEPLGVAPTKAKRRVVQREKRAFG-LGAVFLGFLGAAGSTMGAASIT | 526 |
| 40 | NWRSELYKYKVVKIEPLGVAPTKAKRRVVQREKRAFG-LGAVFLGFLGAAGSTMGAASIT | 526 |
| 41 | NWRSELYKYKVVKIEPLGVAPTKAKRRVVQREKRAFG-LGAVFLGFLGAAGSTMGAASIT | 526 |
| 42 | NWRSELYKYKVVKIEPLGVAPTKAKRRVVQREKRAFG-LGAVFLGFLGAAGSTMGAASIT | 526 |
| 43 | NWRSELYKYKVVKIEPLGVAPTKAKRRVVQREKRAFG-LGAVFLGFLGAAGSTMGAASIT | 526 |
| 44 | NWRSELYKYKVVKIEPLGVAPTKAKRRVVQREKRAFG-LGAVFLGFLGAAGSTMGAASIT | 526 |
| 45 | NWRSELYKYKVVKIEPLGVAPTKAKRRVVQREKRAFG-LGAVFLGFLGAAGSTMGAASIT | 526 |
| 46 | NWRSELYKYKVVKIEPLGVAPTKAKRRVVQREKRAFG-LGAVFLGFLGAAGSTMGAASIT | 526 |
| 47 | NWRSELYKYKVVKIEPLGVAPTKAKRRVVQREKRAFG-LGAVFLGFLGAAGSTMSAASIT | 526 |
| 48 | NWRSELYKYKVVKIEPLGVAPTKAKRRVVQREKRAFG-LGAVFLGFLGAAGSTMGAASIT | 526 |
| 49 | NWRSELYKYKVVKIEPLGVAPTKAKRRVVQREKRAFG-LGAVFLGFLGAAGSTMGAASIT | 526 |
| 50 | NWRSELYKYKVVKIEPLGVAPTKAKRRVVQREKRAFG-LGAVFLGFLGAAGSTMGAASIT | 526 |
| 51 | NWRSELYKYKVVKIEPLGVAPTKAKRRVVQREKRAFG-LGAVFLGFLGAAGSTMGAASIT | 526 |
| 52 | NWRSELYKYKVVKIEPLGVAPTKAKRRVVQREKRAFG-LGAVFLGFLGAAGSTMGAASIT | 526 |
| 53 | NWRSELYKYKVVKIEPLGVAPTKAKRRVVQREKRAFG-LGAVFLGFLGAAGSTMGAASIT | 526 |
| 54 | NWRSELYKYKVVKIEPLGVAPTKAKRRVVQREKRAFG-LGAVFLGFLGAAGSTMGAASIT | 526 |
| 55 | NWRSELYKYKVVKIEPLGVAPTKAKRRVVQREKRAFG-LGAVFLGFLGAAGSTMGAASIT | 526 |
| 56 | NWRSELYKYKVVKIEPLGVAPTKAKRRVVQREKRAFG-LGAVFLGFLGAAGSTMGAASIT | 526 |
| 57 | NWRSELYKYKVVKIEPLGVAPTKAKRRVVQREKRAFG-LGAVFLGFLGAAGSTMSAASIT | 526 |
| 58 | NWRSELYKYKVVKIEPLGVAPTKAKRRVVQREKRAFG-LGAVFLGFLGAAGSTMGAASIT | 526 |
| 59 | NWRSELYKYKVVKIEPLGVAPTKAKRRVVQREKRAFG-LGAVFLGFLGAAGSTMGAASIT | 524 |
| 60 | NWRSELYKYKVVKIEPLGVAPTKAKRKVVQREKRAFG-LGAVFLGFLGAAGSTMGAASIT | 526 |
| 61 | NWRSELYKYKVVKIEPLGVAPTKAKRRVVQREKRAFG-LGAVFLGFLGAAGSTMGAASIT | 526 |
| 62 | NWRSELYKYKVVKIEPLGVAPTKAKRRVVQREKRAFG-LGAVFLGFLGAAGSTMGAASIT | 526 |
| 63 | NWRSELYKYKVVKIEPLGVAPTKAKRRVVQREKRAFG-LGAVFLGFLGAAGSTMGAASIT | 526 |
| 64 | NWRSELYKYKVVKIEPLGVAPTKAKRRVVQREKRAFG-LGAVFLGFLGAAGSTMGAASIT | 526 |
| 65 | NWRSELYKYKVVKIEPLGVAPTKAKRRVVQREKRAFG-LEAVFLGFLGAAGSTMGAASIT | 526 |
| 66 | NWRSELYKYKVVKIEPLGVAPTKAKRRVVQREKRAFG-LGAVFLGFLGAAGSTMGAASIT | 526 |
| 67 | NWRSELYKYKVVKIEPLGVAPTKAKRRVVQREKRAFG-LGAVFLGFLGAAGSTMGAASIT | 524 |
| 68 | NWRSELYKYKVVKIEPLGVAPTKAKRRVVQREKRAFG-LGAVFLGFLGAAGSTMGAASIT | 526 |

**LTVQARQLLSGIVQQQNNLLRAIEAQQHLLQLTVWGIKQLQARVLAVERYLKDQQLLGIW**

| | | |
|---|---|---|
| 1 | LTVQARQLLSGIVQQQNNLLRAIEAQQHLLQLTVWGIKQLQARVLAVERYLKDQQLLGIW | 586 |
| 2 | LTVQARQLLSGIVQQQNNLLRAIEAQQHLLQLTVWGIKQLQARVLAVERYLKDQQLLGIW | 585 |
| 3 | LTVQARQLLSGIVQQQNNLLRAIEAQQHLLQLTVWGIKQLQARVLAVERYLKDQQLLGIW | 586 |
| 4 | LTVQARQLLSGIVQQQNNLLRAIEAQQHLLQLTVWGIKQLQTRVLAVERYLKDQQLLGIW | 586 |
| 5 | LTVQARQLLSGIVQQQNNLLRAIEAQQHLLQLTVWGIKQLQARVLAVERYLKDQQLLGIW | 586 |
| 6 | LTVQARQLLSGIVQQQNNLLRAIEAQQHLLQLTVWGIKQLQARVLAVERYLKDQQLLGIW | 586 |
| 7 | LTVQARQLLSGIVQQQNNLLRAIEAQQHLLQLTVWGIKQLQARVLAVERYLKDQQLLGIW | 586 |
| 8 | LTVQARQLLSGIVQQQNNLLRAIEAQQHLLQLTVWGIKQLQARVLAVERYLKDQQLLGIW | 586 |
| 9 | LTVQARQLLSGIVQQQNNLLRAIEAQQHLLQLTVWGIKQLQARVLAVERYLKDQQLLGIW | 585 |
| 10 | LTVQARQLLSGIVQQQNNLLRAIEAQQHLLQLTVWGIKQLQARVLAVERYLKDQQLLGIW | 586 |
| 11 | LTVQARQLLSGIVQQQNNLLRAIEAQQHLLQLTVWGIKQLQARVLAVERYLKDQQLLGIW | 586 |
| 12 | LTVQARQLLSGIVQQQNNLLRAIEAQQHLLQLTVWGIKQLQARVLAVERYLKDQQLLGIW | 586 |
| 13 | LTVQARQLLSGIVQQQNNLLRAIEAQQHLLQLTVWGIKQLQARVLAVERYLKDQQLLGIW | 586 |
| 14 | LTVQARQLLSGIVQQQNNLLRAIEAQQHLLQLTVWGIKQLQARVLAVERYLKDQQLLGIW | 584 |
| 15 | LTVQARQLLSGIVQQQNNLLRAIEAQQHLLQLTVWGIKQLQARVLAVERYLKDQQLLGIW | 554 |
| 16 | LTVQARQLLSGIVQQQNNLLRAIEAQQHLLQLTVWGIKQLQARVLAVERYLKDQQLLGIW | 586 |
| 17 | LTVQARQLLSGIVQQQNNLLRAIEAQQHLLQLTVWGIKQLQARVLAVERYLKDQQLLGIW | 586 |
| 18 | LTVQARQLLSGIVQQQNNLLRAIEAQQHLLQLTVWGIKQLQARVLAVERYLKDQQLLGIW | 586 |
| 19 | LTVQARQLLSGIVQQQNNLLRAIEAQQHLLQLTVWGIKQLQARVLAVERYLKDQQLLGIW | 586 |
| 20 | LTVQARQLLSGIVQQQNNLLRAIEAQQHLLQLTVWGIKQLQARVLAVERYLKDQQLLGIW | 584 |
| 21 | LTVQARQLLSGIVQQQNNLLRAIEAQQHLLQLTVWGIKQLQARVLAVERYLKDQQLLGIW | 586 |
| 22 | LTVQARQLLSGIVQQQNNLLRAIEAQQHLLQLTVWGIKQLQARVLAVERYLKDQQLLGIW | 569 |
| 23 | LTVQARQLLSGIVQQQNNLLRAIEAQQHLLQLTVWGIKQLQARVLAVERYLKDQQLLGIW | 586 |
| 24 | LTVQARQLLSGIVQQQNNLLRAIEAQQHLLQLTVWGIKQLQARVLAVERYLKDQQLLGIW | 585 |
| 25 | LTVQARQLLSGIVQQQNNLLRAIEAQQHLLQLTVWGIKQLQARVLAVERYLKDQQLLGIW | 586 |
| 26 | LTVQARQLLSGIVQQQNNLLRAIEAQQHLLQLTVWGIKQLQARVLAVERYLKDQQLLGIW | 586 |
| 27 | LTVQARQLLSGIVQQQNNLLRAIEAQQHLLQLTVWGIKQLQARVLAVERYLKDQQLLGIW | 586 |
| 28 | LTVQARQLLSGIVQQQNNLLRAIEAQQHLLQLTVWGIKQLQARVLAVERYLKDQQLLGIW | 586 |
| 29 | LTVQARQLLSGIVQQQNNLLRAIEAQQHLLQLTVWGIKQLQARVLAVERYLKDQQLLGIW | 586 |
| 30 | LTVQARQLLSGIVQQQNNLLRAIEAQQHLLQLTVWGIKQLQARVLAVERYLKDQQLLGIW | 586 |
| 31 | LTVQARQLLSGIVQQQNNLLRAIEAQQHLLQLTVWGIKQLQARVLAVERYLKDQQLLGIW | 586 |
| 32 | LTVQARQLLSGIVQQQNNLLRAIEAQQHLLQLTVWGIKQLQARVLAVERYLKDQQLLGIW | 586 |
| 33 | LTVQARQLLSGIVQQQNNLLRAIEAQQHLLQLTVWGIKQLQARVLAVERYLKDQQLLGIW | 586 |
| 34 | LTVQARQLLSGIVQQQNNLLRAIEAQQHLLQLTVWGIKQLQARVLAVERYLKDQQLLGIW | 586 |
| 35 | LTVQARQLLSGIVQQQNNLLRAIEAQQHLLQLTVWGIKQLQARVLAVERYLKDQQLLGIW | 586 |
| 36 | LTVQARQLLSGIVQQQNNLLRAIEAQQHLWQLTVWGIKQLQARVLAVERYLKDQQLLGIW | 586 |
| 37 | LTVQARQLLSGIVQQQNNLLRAIEAQQHLLQLTVWGIKQLQARVLAVERYLKDQQLLGIW | 586 |
| 38 | LTVQARQLLSGIVQQQNNLLRAIEAQQHLLQLTVWGIKQLQARVLAVERYLKDQQLLGIW | 584 |
| 39 | LTVQARQLLSGIVQQQNNLLRAIEAQQHLLQLTVWGIKQLQARVLAVERYLKDQQLLGIW | 586 |
| 40 | LTVQARQLLSGIVQQQNNLLRAIEAQQHLLQLTVWGIKQLQARVLAVERYLKDQQLLGIW | 586 |
| 41 | LTVQARQLLSGIVQQQNNLLRAIEAQQHLLQLTVWGIKQLQARVLAVERYLKDQQLLGIW | 586 |
| 42 | LTVQARQLLSGIVQQQNNLLRAIEAQQHLLQLTVWGIKQLQARVLAVERYLKDQQLLGIW | 586 |
| 43 | LTVQARQLLSGIVQQQNNLLRAIEAQQHLLQLTVWGIKQLQARVLAVERYLKDQQLLGIW | 586 |
| 44 | LTVQARQLLSGIVQQQNNLLRAIEAQQHLLQLTVWGIKQLQARVLAVERYLKDQQLLGIW | 586 |
| 45 | LTVQARQLLSGIVQQQNNLLRAIEAQQHLLQLTVWGIKQLQARVLAVERYLKDQQLLGIW | 586 |
| 46 | LTVQARQLLSGIVQQQNNLLRAIEAQQHLLQLTVWGIKQLQARVLAVERYLKDQQLLGIW | 586 |
| 47 | LTVQARQLLSGIVQQQNNLLRAIEAQQHLLQLTVWGIKQLQARVLAVERYLKDQQLLGIW | 586 |
| 48 | LTVQARQLLSGIVQQQNNLLRAIEAQQHLLQLTVWGIKQLQARVLAVERYLKDQQLLGIW | 586 |
| 49 | LTVQARQLLSGIVQQQNNLLRAIEAQQHLLQLTVWGIKQLQARVLAVERYLKDQQLLGIW | 586 |
| 50 | LTVQARQLLSGIVQQQNNLLRAIEAQQHLLQLTVWGIKQLQARVLAVERYLKDQQLLGIW | 586 |
| 51 | LTVQARQLLSGIVQQQNNLLRAIEAQQHLLQLTVWGIKQLQARVLAVERYLKDQQLLGIW | 586 |
| 52 | LTVQARQLLSGIVQQQNNLLRAIEAQQHLLQLTVWGIKQLQARVLAVERYLKDQQLLGIW | 586 |
| 53 | LTVQARQLLSGIVQQQNNLLRAIEAQQHLLQLTVWGIKQLQARVLAVERYLKDQQLLGIW | 586 |
| 54 | LTVQARQLLSGIVQQQNNLLRAIEAQQHLLQLTVWGIKQLQARVLAVERYLKDQQLLGIW | 586 |
| 55 | LTVQARQLLSGIVQQQNNLLRAIEAQQHLLQLTVWGIKQLQARVLAVERYLKDQQLLGIW | 586 |

| | | |
|---|---|---|
| 56 | LTVQARQLLSGIVQQQNNLLRAIEAQQHLLQLTVWGIKQLQARVLAVERYLKDQQLLGIW | 586 |
| 57 | LTVQARQLLSGIVQQQNNLLRAIEAQQHLLQLTVWGIKQLQARVLAVERYLKDQQLLGIW | 586 |
| 58 | LTVQARQLLSGIVQQQNNLLRAIEAQQHLLQLTVWGIKQLQARVLAVERYLKDQQLLGIW | 586 |
| 59 | LTVQARQLLSGIVQQQNNLLRAIEAQQHLLQLTVWGIKQLQARVLAVERYLKDQQLLGIW | 584 |
| 60 | LTVQARQLLSGIVQQQNNLLRAIEAQQHLLQLTVWGIKQLQARVLAVERYLKDQQLLGIW | 586 |
| 61 | LTVQARQLLSGIVQQQNNLLRAIEAQQHLLQLTVWGIKQLQARVLAVERYLKDQQLLGIW | 586 |
| 62 | LTVQARQLLSGIVQQQNNLLRAIEAQQHLLQLTVWGIKQLQARVLAVERYLKDQQLLGIW | 586 |
| 63 | LTVQARQLLSGIVQQQNNLLRAIEAQQHLLQLTVWGIKQLQARVLAVERYLKDQQLLGIW | 586 |
| 64 | LTVQARQLLSGIVQQQNNLLRAIEAQQHLLQLTVWGIKQLQARVLAVERYLKDQQLLGIW | 586 |
| 65 | LTVQARQLLSGIVQQQNNLLRAIEAQQHLLQLTVWGIKQLQARVLAVERYLKDQQLLGIW | 586 |
| 66 | LTVQARQLLSGIVQQQNNLLRAIEAQQHLLQLTVWGIKQLQARVLAVERYLKDQQLLGIW | 586 |
| 67 | LTVQARQLLSGIVQQQNNLLRAIEAQQHLLQLTVWGIKQLQARVLAVERYLKDQQLLGIW | 584 |
| 68 | LTVQARQLLSGIVQQQNNLLRAIEAQQHLLQLTVWGIKQLQARVLAVERYLKDQQLLGIW | 586 |

| | | |
|---|---|---|
| | **GCSGKLICTTTVPWNTSWSNKSLEQIWDNMTWMEWEREIDNYTGYIYQLIEESQNQQEKN** | |
| 1 | GCSGKLICTTTVPWNTSWSNKSLEQIWDNMTWMEWEREIDNYTGYIYQLIEESQNQQEKN | 646 |
| 2 | GCSGKLICTTTVPWNTSWSNKSLEQIWDNMTWMEWEREIDNYTGYIYQLIEESQNQQEKN | 645 |
| 3 | GCSGKLICTTTVPWNTSWSNKSLEQIWDNMTWMEWEREIDNYTGYIYQLIEESQNQQEKN | 646 |
| 4 | GCSGKLICTTTVPWNTSWSNKSLEQIWDNMTWMEWEREIDNYTGYIYQLIEESQNQQEKN | 646 |
| 5 | GCSGKLICTTTVPWNTSWSNKSLEQIWDNMTWMEWEREIDNYTGYIYQLIEESQNQQEKN | 646 |
| 6 | GCSGKLICTTTVPWNTSWSNKSLEQIWDNMTWMEWEREIDNYTGYIYQLIEESQNQQEKN | 646 |
| 7 | GCSGKLICTTTVPWNTSWSNKSLEQIWDNMTWMEWEREIDNYTGYIYQLIEESQNQQEKN | 646 |
| 8 | GCSGKLICTTTVPWNTSWSNKSLEQIWDNMTWMEWEREIDNYTGYIYQLIEESQNQQEKN | 646 |
| 9 | GCSGKLICTTTVPWNTSWSNKSLEQIWDNMTWMEWEREIDNYTGYIYQLIEESQNQQEKN | 645 |
| 10 | GCSGKLICTTTVPWNTSWSNKSLEQIWDNMTWMEWEREIDNYTGYIYQLIEESQNQQEKN | 646 |
| 11 | GCSGKLICTTTVPWNTSWSNKSLEQIWDNMTWMEWEREIDNYTGYIYQLIEESQNQQEKN | 646 |
| 12 | GCSGKLICTTTVPWNTSWSNKSLEQIWDNMTWMEWEREIDNYTGYIYQLIEESQNQQEKN | 646 |
| 13 | GCSGKLICTTTVPWNTSWSNKSLEQIWDNMTWMEWEREIDNYTGYIYQLIEESQNQQEKN | 646 |
| 14 | GCSGKLICTTTVPWNTSWSNKSLEQIWDNMTWMEWEREIDNYTGYIYQLIEESQNQQEKN | 644 |
| 15 | GCSGKLICTTTVPWNTSWSNKSLEQIWDNMTWMEWEREIDNYTGYIYQLIEKSQNQQEKN | 614 |
| 16 | GCSGKLICTTTVPWNTSWSNKSLEQIWDNMTWMEWEREIDNYTGYIYQLIEESQNQQEKN | 646 |
| 17 | GCSGKLICTTTVPWNTSWSNKSLEQIWDNMTWMEWEREIDNYTGYIYQLIEESQNQQEKN | 646 |
| 18 | GCSGKLICTTTVPWNTSWSNKSLEQIWDNMTWMEWEREIDNYTGYIYQLIEESQNQQEKN | 646 |
| 19 | GCSGKLICTTTVPWNTSWSNKSLEQIWDNMTWMEWEREIDNYTGYIYQLIEESQNQQEKN | 646 |
| 20 | GCSGKLICTTTVPWNTSWSNKSLEQIWDNMTWMEWEREIDNYTGYIYQLIEESQNQQEKN | 644 |
| 21 | GCSGKLICTTTVPWNTSWSNKSLEQIWDNMTWMEWEREIDNYTGYIYQLIEESQNQQEKN | 646 |
| 22 | GCSGKLICTTTVPWNTSWSNKSLEQIWDNMTWMEWEREIDNYTGYIYQLIEESQNQQEKN | 629 |
| 23 | GCSGKLICTTTVPWNTSWSNKSLEQIWDNMTWMEWEREIDNYTGYIYQLIEESQNQQEKN | 646 |
| 24 | GCSGKLICTTTVPWNTSWSNKSLEQIWDNMTWMEWEREIDNYTGYIYQLIEESQNQQEKN | 645 |
| 25 | GCSGKLICTTTVPWNTSWSNKSLEQIWDNMTWMEWEREIDNYTGYIYQLIEESQNQQEKN | 646 |
| 26 | GCSGKLICTTTVPWNTSWSNKSLEQIWDNMTWMEWEREIDNYTGYIYQLIEESQNQQEKN | 646 |
| 27 | GCSGKLICTTTVPWNTSWSNKSLEQIWDNMTWMEWEREIDNYTGYIYQLIEESQNQQEKN | 646 |
| 28 | GCSGKLICTTTVPWNTSWSNKSLEQIWDNMTWMEWEREIDNYTGYIYQLIEESQNQQEKN | 646 |
| 29 | GCSGKLICTTTVPWNTSWSNKSLEQIWDNMTWMEWEREIDNYTGYIYQLIEESQNQQEKN | 646 |
| 30 | GCSGKLICTTTVPWNTSWSNKSLEQIWDNMTWMEWEREIDNYTGYIYQLIEESQNQQEKN | 646 |
| 31 | GCSGKLICTTTVPWNTSWSNKSLEQIWDNMTWMEWEREIDNYTGYIYQLIEESQNQQEKN | 646 |
| 32 | GCSGKLICTTTVPWNTSWSNKSLEQIWDNMTWMEWEREIDNYTGYIYQLIEESQNQQEKN | 646 |
| 33 | GCSGKLICTTTVPWNTSWSNKSLEQIWDNMTWMEWEREIDNYTGYIYQLIEESQNQQEKN | 646 |
| 34 | GCSGKLICTTTVPWNTSWSNKSLEQIWDNMTWMEWEREIDNYTGYIYQLIEESQNQQEKN | 646 |
| 35 | GCSGKLICTTTVPWNTSWSNKSLEQIWDNMTWMEWEREIDNYTGYIYQLIEESQNQQEKN | 646 |
| 36 | GCSGKLICTTTVPWNTSWSNKSLEQIWDNMTWMEWEREIDNYTGYIYQLIEESQNQQEKN | 646 |
| 37 | GCSGKLICTTTVPWNTSWSNKSLEQIWDNMTWMEWEREIDNYTGYIYQLIEESQNQQEKN | 646 |

| | | |
|---|---|---|
| 38 | GCSGKLICTTTVPWNTSWSNKSLEQIWDNMTWMEWEREIDNYTGYIYQLIEESQNQQEKN | 644 |
| 39 | GCSGKLICTTTVPWNTSWSNKSLEQIWDNMTWMEWEREIDNYTGYIYQLIEESQNQQEKN | 646 |
| 40 | GCSGKLICTTTVPWNTSWSNKSLEQIWDNMTWMEWEREIDNYTGYIYQLIEESQNQQEKN | 646 |
| 41 | GCSGKLICTTTVPWNTSWSNKSLEQIWDNMTWMEWEREIDNYTGYIYQLIEESQNQQEKN | 646 |
| 42 | GCSGKLICTTTVPWNTSWSNKSLEQIWDNMTWMEWEREIDNYTGYIYQLIEESQNQQEKN | 646 |
| 43 | GCSGKLICTTTVPWNTSWSNKSLEQIWDNMTWMEWEREIDNYTGYIYQLIEESQNQQEKN | 646 |
| 44 | GCSGKLICTTTVPWNTSWSNKSLEQIWDNMTWMEWEREIDNYTGYIYQLIEESQNQQEKN | 646 |
| 45 | GCSGKLICTTTVPWNTSWSNKSLEQIWDNMTWMEWEREIDNYTGYIYQLIEESQNQQEKN | 646 |
| 46 | GCSGKLICTTTVPWNTSWSNKSLEQIWDNMTWMEWEREIDNYTGYIYQLIEESQNQQEKN | 646 |
| 47 | GCSGKLICTTTVPWNTSWSNKSLEQIWDNMTWMEWEREIDNYTGYIYQLIEESQNQQEKN | 646 |
| 48 | GCSGKLICTTTVPWNTSWSNKSLEQIWDNMTWMEWEREIDNYTGYIYQLIEESQNQQEKN | 646 |
| 49 | GCSGKLICTTTVPWNTSWSNKSLEQIWDNMTWMEWEREIDNYTGYIYQLIEESQNQQEKN | 646 |
| 50 | GCSGKLICTTTVPWNTSWSNKSLEQIWDNMTWMEWEREIDNYTGYIYQLIEESQNQQEKN | 646 |
| 51 | GCSGKLICTTTVPWNTSWSNKSLEQIWDNMTWMEWEREIDNYTGYIYQLIEESQNQQEKN | 646 |
| 52 | GCSGKLICTTTVPWNTSWSNKSLEQIWDNMTWMEWEREIDNYTGYIYQLIEESQNQQEKN | 646 |
| 53 | GCSGKLICTTTVPWNTSWSNKSLEQIWDNMTWMEWEREIDNYTGYIYQLIEESQNQQEKN | 646 |
| 54 | GCSGKLICTTTVPWNTSWSNKSLEQIWDNMTWMEWEREIDNYTGYIYQLIEESQNQQEKN | 646 |
| 55 | GCSGKLICTTTVPWNTSWSNKSLEQIWDNMTWMEWEREIDNYTGYIYQLIEESQNQQEKN | 646 |
| 56 | GCSGKLICTTTVPWNTSWSNKSLEQIWDNMTWMEWEREIDNYTGYIYQLIEESQNQQEKN | 646 |
| 57 | GCSGKLICTTTVPWNTSWSNKSLEQIWDNMTWMEWEREIDNYTGYIYQLIEESQNQQEKN | 646 |
| 58 | GCSGKLICTTTVPWNTSWSNKSLEQIWDNMTWMEWEREIDNYTGYIYQLIEESQNQQEKN | 646 |
| 59 | GCSGKLICTTTVPWNTSWSNKSLEQIWDNMTWMEWEREIDNYTGYIYQLIEESQNQQEKN | 644 |
| 60 | GCSGKLICTTTVPWNTSWSNKSLEQIWDNMTWMEWEREIDNYTGYIYQLIEESQNQQEKN | 646 |
| 61 | GCSGKLICTTTVPWNTSWSNKSLEQIWDNMTWMEWEREIDNYTGYIYQLIEESQNQQEKN | 646 |
| 62 | GCSGKLICTTTVPWNTSWSNKSLEQIWDNMTWMEWEREIDNYTGYIYQLIEESQNQQEKN | 646 |
| 63 | GCSGKLICTTTVPWNTSWSNKSLEQIWDNMTWMEWEREIDNYTGYIYQLIEESQNQQEKN | 646 |
| 64 | GCSGKLICTTTVPWNTSWSNKSLEQIWDNMTWMEWEREIDNYTGYIYQLIEESQNQQEKN | 646 |
| 65 | GCSGKLICTTTVPWNTSWSNKSLEQIWDNMTWMEWEREIDNYTGYIYQLIEESQNQQEKN | 646 |
| 66 | GCSGKLICTTTVPWNTSWSNKSLEQIWDNMTWMEWEREIDNYTGYIYQLIEESQNQQEKN | 646 |
| 67 | GCSGKLICTTTVPWNTSWSNKSLEQIWDNMTWMEWEREIDNYTGYIYQLIEESQNQQEKN | 644 |
| 68 | GCSGKLICTTTVPWNTSWSNKSLEQIWDNMTWMEWEREIDNYTGYIYQLIEESQNQQEKN | 646 |

| | | |
|---|---|---|
| | **EQELLALDKWASLWNWFDITNWLWYIKIFIMIVGGLIGLRIVFTVLSIVNRVRQGYSPLS** | |
| 1 | EQELLALDKWASLWNWFDITNWLWYIKIFIMIVGGLIGLRIVFTVLSIVNRVRQGYSPLS | 706 |
| 2 | EQELLALDKWASLWNWFDITNWLWYIKIFIMIVGGLIGLRIVFTVLSIVNRVRQGYSPLS | 705 |
| 3 | EQELLALDKWASLWNWFDITNWLWYIKIFIMIVGGLIGLRIVFTVLSIVNRVRQGYSPLS | 706 |
| 4 | EQELLALDKWASLWNWFDITNWLWYIKIFIMIVGGLIGLRIVFTVLSIVNRVRQGYSPLS | 706 |
| 5 | EQELLALDKWASLWNWFDITNWLWYIKIFIMMVGGLIGLRIVFTVLSIVNRVRQGYSPLS | 706 |
| 6 | EQELLALDKWASLWNWFDITNWLWYIKIFIMIVGGLIGLRIVFTVLSIVNRVRQGYSPLS | 706 |
| 7 | EQELLALDKWASLWNWFDITNWLWYIKIFIMIVGGLIGLRIVFTVLSIVNRVRQGYSPLS | 706 |
| 8 | EQELLALDKWASLWNWFDITNWLWYIKIFIMIVGGLIGLRIVFTVLSIVNRVRQGYSPLS | 706 |
| 9 | EQELLALDKWASLWNWFDITNWLWYIKIFIMIVGGLIGLRIVFTVLSIVNRVRQGYSPLS | 705 |
| 10 | EQELLALDKWASLWNWFDITNWLWYIKIFIMIVGGLIGLRIVFTVLSIVNRVRQGYSPLS | 706 |
| 11 | EQELLALDKWASLWNWFDITNWLWYIKIFIMIVGGLIGLRIVFTVLSIVNRVRQGYSPLS | 706 |
| 12 | EQELLALDKWASLWNWFDITNWLWYIKIFIMIVGGLIGLRIVFTVLSIVNRVRQGYSPLS | 706 |
| 13 | EQELLALDKWASLWNWFDITNWLWYIKIFIMIVGGLIGLRIVFTVLSIVNRVRQGYSPLS | 706 |
| 14 | EQELLALDKWASLWNWFDITNWLWYIKIFIMIVGGLIGLRIVFTVLSIVNRVRQGYSPLS | 704 |
| 15 | EQELLALDKWASLWNWFDITNWLWYIKIFIMIVGGLIGLRIVFTVLSIVNRVRQGYSPLS | 674 |
| 16 | EQELLALDKWASLWNWFDITNWLWYIKIFIMIVGGLIGLRIVFTVLSIVNRVRQGYSPLS | 706 |
| 17 | EQELLALDKWASLWNWFDITNWLWYIKIFIMIVGGLIGLRIVFTVLSIVNRVRQGYSPLS | 706 |
| 18 | EQELLALDEWASLWNWFDITNWLWYIKIFIMIVGGLIGLRIVFTVLSIVNRVRQGYSPLS | 706 |
| 19 | EQELLALDKWASLWNWFDITNWLWYIKIFIMIVGGLIGLRIVFTVLSIVNRVRQGYSPLS | 706 |

20 EQELLALDKWASLWNWFDITNWLWYIKIFIMIVGGLIGLRIVFTVLSIVNRVRQGYSPLS 704
21 EQELLALDKWASLWNWFDITNWLWYIKIFIMIVGGLIGLRIVFTVLSIVNRVRQGYSPLS 706
22 EQELLALDKWASLWNWFDITNWLWYIKIFIMIVGGLIGLRIVFTVLSIVNRVRQGYSPLS 689
23 EQELLALDKWASLWNWFDITNWLWYIKIFIMIVGGLIGLRIVFTVLSIVNRVRQGYSPLS 706
24 EQELLALDKWASLWNWFDITNWLWYIKIFIMIVGGLIGLRIVFTVLSIVNRVRQGYSPLS 705
25 EQELLALDKWASLWNWFDITNWLWYIKIFIMIVGGLIGLRIVFTVLSIVNRVRQGYSPLS 706
26 EQELLALDKWASLWNWFDITNWLWYIKIFIMIVGGLIGLRIVFTVLSIVNRVRQGYSPLS 706
27 EQELLALDKWASLWNWFDITNWLWYIKIFIMIVGGLIGLRIVFTVLSIVNRVRQGYSPLS 706
28 EQELLALDKWASLWNWFDITNWLWYIKIFIMMVGGLIGLRIVFTVLSIVNRVRQGYSPLS 706
29 EQELLALDKWASLWNWFDITNWLWYIKIFIMIVGGLIGLRIVFTVLSIVNRVRQGYSPLS 706
30 EQELLALDKWASLWNWFDITNWLWYIKIFIMIVGGLIGLRIVFTVLSIVNRVRQGYSPLS 706
31 EQELLALDKWASLWNWFDITNWLWYIKIFIMIVGGLIGLRIVFTVLSIVNRVRQGYSPLS 706
32 EQELLALDKWASLWNWFDITNWLWYIKIFIMIVGGLIGLRIVFTVLSIVNRVRQGYSPLS 706
33 EQELLALDKWASLWNWFDITNWLWYIKIFIMIVGGLIGLRIVFTVLSIVNRVRQGYSPLS 706
34 EQELLALDKWASLWNWFDITNWLWYIKIFIMIVGGLIGLRIVFTVLSIVNRVRQGYSPLS 706
35 EQELLALDKWASLWNWFDITNWLWYIKIFIMIVGGLIGLRIVFTVLSIVNRVRQGYSPLS 706
36 EQELLALDKWASLWNWFDITNWLWYIKIFIMIVGGLIGLRIVFTVLSIVNRVRQGYSPLS 706
37 EQELLALDKWASLWNWFDITNWLWYIKIFIMIVGGLIGLRVVFTVLSIVNRVRQGYSPLS 706
38 EQELLALDKWASLWNWFDITNWLWYIKIFIMIVGGLIGLRIVFTVLSIVNRVRQGYSPLS 704
39 EQELLALDKWASLWNWFDITNWLWYIKIFIMIVGGLIGLRIVFTVLSIVNRVRQGYSPLS 706
40 EQELLALDKWASLWNWFDITNWLWYIKIFIMIVGGLIGLRIVFTVLSIVNRVRQGYSPLS 706
41 EQELLALDKWASLWNWFDITNWLWYIKIFIMIVGGLIGLRIVFTVLSIVNRVRQGYSPLS 706
42 EQELLALDKWASLWNWFDITNWLWYIKIFIMIVGGLIGLRIVFTVLSIVNRVRQGYSPLS 706
43 EQELLALDKWASLWNWFDITNWLWYIKIFIMIVGGLIGLRIVFTVLSIVNRVRQGYSPLS 706
44 EQELLALDKWASLWNWFDITNWLWYIKIFIMIVGGLIGLRIVFTVLSIVNRVRQGYSPLS 706
45 EQELLALDKWASLWNWFDITNWLWYIKIFIMIVGGLIGLRIVFTVLSIVNRVRQGYSPLS 706
46 EQELLALDKWASLWNWFDITNWLWYIKIFIMIVGGLIGLRIVFTVLSIVNRVRQGYSPLS 706
47 EQELLALDKWASLWNWFDITNWLWYIKIFIMIVGGLIGLRIVFTVLSIVNRVRQGYSPLS 706
48 EQELLALDKWASLWNWFDITNWLWYIKIFIMIVGGLIGLRIVFTVLSIVNRVRQGYSPLS 706
49 EQELLALDKWASLWNWFDITNWLWYIKIFIMIVGGLIGLRIVFTVLSIVNRVRQGYSPLS 706
50 EQELLALDKWASLWNWFDITDWLWYIKIFIMIVGGLIGLRIVFTVLSIVNRVRQGYSPLS 706
51 EQELLALDKWASLWNWFDITNWLWYIKIFIMIVGGLIGLRIVFTVLSIVNRVRQGYSPLS 706
52 EQELLALDKWASLWNWFDITNWLWYIKIFIMIVGGLIGLRIVFTVLSIVNRVRQGYSPLS 706
53 EQELLALDKWASLWNWFDITNWLWYIKIFIMIVGGLIGLRIVFTVLSIVNRVRQGYSPLS 706
54 EQELLALDKWASLWNWFDITNWLWYIKIFIMIVGGLIGLRIVFTVLSIVNRVRQGYSPLS 706
55 EQELLALDKWASLWNWFDITNWLWYIKIFIMIVGGLIGLRIVFTVLSIVNRVRQGYSPLS 706
56 EQELLALDKWASLWNWFDITNWLWYIKIFIMIVGGLIGLRIVFTVLSIVNRVRQGYSPLS 706
57 EQELLALDKWASLWNWFDITNWLWYIKIFIMIVGGLIGLRIVFTVLSIVNRVRQGYSPLS 706
58 EQELLALDKWASLWNWFDITNWLWYIKIFIMIVGGLIGLRIVFTVLSIVNRVRQGYSPLS 706
59 EQELLALDKWASLWNWFDITNWLWYIKIFIMIVGGLLGLRIVFTVLSIVNRVRQGYSPLS 704
60 EQELLALDKWASLWNWFDITNWLWYIKIFIMIVGGLIGLRIVFTVLSIVNRVRQGYSPLS 706
61 EQELLALDKWASLWNWFDITNWLWYIKIFIMIVGGLIGLRIVFTVLSIVNRVRQGYSPLS 706
62 EQELLALDKWASLWNWFDITNWLWYIKIFIMIVGGLIGLRIVFTVLSIVNRVRQGYSPLS 706
63 EQELLALDKWASLWNWFDITNWLWYIKIFIMIVGGLIGLRIVFTVLSIVNRVRQGYSPLS 706
64 EQELLALDKWASLWNWFDITNWLWYIKIFIMIVGGLIGLRIVFTVLSIVNRVRQGYSPLS 706
65 EQELLALDKWASLWNWFDITNWLWYIKIFIMIVGGLIGLRIVFTVLSIVNRVRQGYSPLS 706
66 EQELLALDKWASLWN-FDITNWLWYIKIFIMIVGGLIGLRIVFTVLSIVNRVRQGYSPLS 705
67 EQELLALDKWASLWNWFDITNWLWYIKIFIMIVGGLIGLRIVFTVLSIVNRVRQGYSPLS 704
68 EQELLALDKWASLWNWFDITNWLWYIKIFIMIVGGLIGLRIVFTVLSIVNRVRQGYSPLS 706

**FQTHLPAQRGPDRPEGIGEEGGERDRDRSDPLVNGFLALIWSDLRSLCLFSYHRLRDLLL**

1 FQTHLPAQRGPDRPEGIGEEGGERDRDRSDPLVNGFLALIWSDLRSLCLFSYHRLRDLLL 766

| | | |
|---|---|---|
| 2 | FQTHLPAQRGPDRPEGIGEEGGERDRDRSDPLVNGFLALIWSDLRSLCLFSYHRLRDLLL | 765 |
| 3 | FQTHLPAQRGPDRPEGIGEEGGERDRDKSDPLVNGFLALIWSDLRSLCLFSYHRLRDLLL | 766 |
| 4 | FQTHLPAQRGPDRPEGIGEEGGERDRDRSDPLVNGFLALIWSDLRSLCLFSYHRLRDLLL | 766 |
| 5 | FQTHLPAQRGPDRPEGIGEEGGERDRDKSDPLVNGFLALIWSDLRSLCLFSYHRLRDLLL | 766 |
| 6 | FQTHLPAQRGPDRPEGIGEEGGERDRDRSDPLVNGFLALIWSDLRSLCLFSYHRLRDLLL | 766 |
| 7 | FQTHLPAQRGPDRPEGIGEEGGERDRDRSDPLVNGFLALIWSDLRSLCLFSYHRLRDLLL | 766 |
| 8 | FQTHLPAQRGPDRPEGIGEEGGERDRDRSDPLVNGFLALIWSDLRSLCLFSYHRLRDLLL | 766 |
| 9 | FQTHLPAQRGPDRPEGIGEEGGERDRDRSDPLVNGFLALIWSDLRSLCLFSYHRLRDLLL | 765 |
| 10 | FQTHLPAQRGPDRPEGIGEEGGERDRDRSDPLVNGFLALIWSDLRSLCLFSYHRLRDLLL | 766 |
| 11 | FQTHLPAQRGPDRPEGIGEEGGERDRDRSDPLVNGFLALIWSDLRSLCLFSYHRLRDLLL | 766 |
| 12 | FQTHLPAQRGPDRPEGIGEEGGERDRDRSDPLVNGFLALIWSDLRSLCLFSYHRLRDLLL | 766 |
| 13 | FQTHLPAQRGPDRPEGIGEEGGERDRDRSDPLVNGFLALIWSDLRSLCLFSYHRLKDLLL | 766 |
| 14 | FQTHLPAQRGPDRPEGIGEEGGERDRDRSDPLVNGFLALIWSDLRSLCLFSYHRLRDLLL | 764 |
| 15 | FQTHLPAQRGPDRPEGIGEEGGERDRDKSDPLVNGFLALIWSDLRSLCLFSYHRLRDLLL | 734 |
| 16 | FQTHLPAQRGPDRPEGIGEEGGERDRDRSDPLVNGFLALIWSDLRSLCLFSYHRLRDLLL | 766 |
| 17 | FQTHLPAQRGPDRPEGIGEEGGERDRDKSDPLVNGFLALIWSDLRSLCLFSYHRLRDLLL | 766 |
| 18 | FQTHLPAQRGPDRPEGIGEEGGERDRDRSDPLVNGFLALIWSDLRSLCLFSYHRLRDLLL | 766 |
| 19 | FQTHLPAQRGPDRPEGIGEEGGERDRDRSDPLVNGFLALIWSDLRSLCLFSYHRLRDLLL | 766 |
| 20 | FQTHLPAQRGPDRPEGIGEEGGERDRDRSDPLVNGFLALIWSDLRSLCLFSYHRLRDLLL | 764 |
| 21 | FQTHLPAQRGPDRPEGIGEEGGERDRDRSDPLVNGFLALIWSDLRSLCLFSYHRLRDLLL | 766 |
| 22 | FQTHLPAQRGPDRPEGIGEEGGERDRDRSDPLVNGFLALIWSDLRSLCLFSYHRLRDLLL | 749 |
| 23 | FQTHLPAQRGPDRPEGIGEEGGERDRDRSDPLVNGFLALIWSDLRSLCLFSYHRLRDLLL | 766 |
| 24 | FQTHLPAQRGPDRPEGIGEEGGERDRDRSDPLVNGFLALIWSDLRSLCLFSYHRLRDLLL | 765 |
| 25 | FQTHLPAQRGPDRPEGIGEEGGERDRDRSDPLVNGFLALIWSDLRSLCLFSYHRLRDLLL | 766 |
| 26 | FQTHLPAQRGPDRPEGIGEEGGERDRDRSDPLVNGFLALIWSDLRSLCLFSYHRLRDLLL | 766 |
| 27 | FQTHLPAQRGPDRPEGIGEEGGERDRDKSDPLVNGFLALIWSDLRSLCLFSYHRLRDLLL | 766 |
| 28 | FQTHLPAQRGPDRPEGIGEEGGERDRDKSDPLVNGFLALIWSDLRSLCLFSCHRLRDLLL | 766 |
| 29 | FQTHLPAQRGPDRPEGIGEEGGERDRDKSDPLVNGFLALIWSDLRSLCLFSYHRLRDLLL | 766 |
| 30 | FQTHLPAQRGPDRPEGIGEEGGERDRDKSDPLVNGFLALIWSDLRSLCLFSYHRLRDLLL | 766 |
| 31 | FQTHLPAQRGPDRPEGIGEEGGERDRDRSDPLVNGFLALIWSDLRSLCLFSYHRLRDLLL | 766 |
| 32 | FQTHLPAQRGPDRPEGIGEEGGERDRDRSDPLVNGFLALIWSDLRSLCLFSYHRLRDLLL | 766 |
| 33 | FQTHLPAQRGPDRPEGIGEEGGERDRDRSDPLVNGFLALIWSDLRSLCLFSYHSLRDLLL | 766 |
| 34 | FQTHLPAQRGPDRPEGIGEEGGERDRDKSDPLVNGFLALIWSDLRSLCLFSYHRLRDLLL | 766 |
| 35 | FQTHLPAQRGPDRPEGIGEEGGERDRDRSDPLVNGFLALIWSDLRSLCLFSYHRLRDLLL | 766 |
| 36 | FQTHLPAQRGPDRPEGIGEEGGERDRDRSDPLVNGFLALIWSDLRSLCLFSYHRLRDLLL | 766 |
| 37 | FQTHLPAQRGPDRPEGIGEEGGERDRDRSDPLVNGFLALIWSDLRSLCLFSYHRLRDLLL | 766 |
| 38 | FQTHLPAQRGPDRPEGIGEEGGERDRDRSDPLVNGFLALIWSDLRSLCLFSYHRLRDLLL | 764 |
| 39 | FQTHLPAQRGPDRPEGIGEEGGERDRDRSDPLVNGFLALIWSDLRSLCLFSYHRLRDLLL | 766 |
| 40 | FQTHLPAQRGPDRPEGIGEEGGERDRDKSDPLVNGFLALIWSDLRSLCLFSYHRLRDLLL | 766 |
| 41 | FQTHLPAQRGPDRPEGIGEEGGERDRDKSDPLVNGFLALIWSDLRSLCLFSYHRLRDLLL | 766 |
| 42 | FQTHLPAQRGPDRPEGIGEEGGERDRDRSDPLVNGFLTLIWSDLRSLCLFSYHRLRDLLL | 766 |
| 43 | FQTHLPAQRGPDRPEGIGEEGGERDRDRSDPLVNGFLTLIWSDLRSLCLFSYHRLRDLLL | 766 |
| 44 | FQTHLPAQRGPDRPEGIGEEGGERDRDRSDPLVNGFLALIWSDLRSLCLFSYHRLRDLLL | 766 |
| 45 | FQTHLPAQRGPDRPEGIGEEGGERDRDRSDPLVNGFLTLIWSDLRSLCLFSYHRLRDLLL | 766 |
| 46 | FQTHLPAQRGPDRPEGIGEEGGERDRDKSDPLVNGFLALIWSDLRSLCLFSYHRLRDLLL | 766 |
| 47 | FQTHLPAQRGPDRPEGIGEEGGERDRDRSDPLVNGFLALIWSDLRSLCLFSYHRLRDLLL | 766 |
| 48 | FQTHLPAQRGPDRPEGIGEEGGERDRDRSDPLVNGFLTLIWSDLRSLCLFSYHRLRDLLL | 766 |
| 49 | FQTHLPAQRGPDRPEGIGEEGGERDRDRSDPLVNGFLALIWSDLRSLCLFSYHRLRDLLL | 766 |
| 50 | FQTHLPAQRGPDRPEGIGEEGGERDRDRSDPLVNGFLALIWSDLRSLCLFSYHRLRDLLL | 766 |
| 51 | FQTHLPAQRGPDRPEGIGEEGGERDRDRSDPLVNGFLALIWSDLRSLCLFSYHRLRDLLL | 766 |
| 52 | FQTHLPAQRGPDRPEGIGEEGGERDRDRSDPLVNGFLTLIWSDLRSLCLFSYHRLRDLLL | 766 |
| 53 | FQTHLPAQRGPDRPEGIGEEGGERDRDRSDPLVNGFLALIWSDLRSLCLFSYHRLRDLLL | 766 |
| 54 | FQTHLPAQRGPDRPEGIGEEGGERDRDRSDPLVNGFLALIWSDLRSLCLFSYHRLRDLLL | 766 |
| 55 | FQTHLPAQRGPDRPEGIGEEGGERDRNRSDPLVNGFLALIWSDLRSLCLFSYHRLRDLLL | 766 |
| 56 | FQTHLPAQRGPDRPEGIGEEGGERDRDRSDPLVNGFLTLIWSDLRSLCLFSYHRLRDLLL | 766 |

```
57  FQTHLPAQRGPDRPEGIGEEGGERDRDRSDPLVNGFLALIWSDLRSLCLFSYHRLRDLLL  766
58  FQTHLPAQRGPDRPERIGEEGGERDRDRSDPLVNGFLALIWSDLRSLCLFSYHRLRDLLL  766
59  FQTHLPAQRGPDRPEGIGEEGGERDRDRSDPLVNGFLALIWSDLRSLCLFSYHRLRDLLL  764
60  FQTHLPAQRGPDRPEGIGEEGGERDRDRSDPLVNGFLTLIWSDLRSLCLFSYHRLRDLLL  766
61  FQTHLPAQRGPDRPEGIGEEGGERDRDRSNPLVNGFLALIWSDLRSLCLFSYHRLRDLLL  766
62  FQTHLPAQRGPDRPEGIGEEGGERDRDRSDPLVNGFLALIWSDLRSLCLFSYHRLRDLLL  766
63  FQTHLPAQRGPDRPEGIGEEGGERDRDRSDPLVNGFLALIWSDLRSLCLFSYHRLRDLLL  766
64  FQTHLPAQRGPDRPEGIGEEGGERDRDRSDPLVNGFLTLIWSDLRSLCLFSYHRLRDLLL  766
65  FQTHLPAQRGPDRPEGIGEEGGERDRDRSDPLVNGFLTLIWSDLRSLCLFSYHRLRDLLL  766
66  FQTHLPAQRGPDRPEGIGEEGGERDRDRSDPLVNGFLTLIWSDLRSLCLFSYHRLRDLLL  765
67  FQTHLPAQRGPDRPEGIGEEGGERDRDRSDPLVNGFLALIWSDLRSLCLFSYHRLRDLLL  764
68  FQTHLPAQRGPDRPEGIGEEGGERDRDRSDPLVNGFLTLIWSDLRSLCLFSYHRLRDLLL  766

    IVTRIVELLGR-------RGWEVLKYWWNLLQYWSQELKNSAVSLLNATAIAVAEGTDRV

1   IVTRIVELLGR-------RGWEVLKYWWNLLQYWSQELKNSAVSLLNATAIAVAEGTDRV  819
2   IVTRIVELLGR-------RGWEVLKYWWNLLQYWSQELKNSAVSLLNATAIAVAEGTDRV  818
3   IVTRIVELLGR-------RGWEVLKYWWNLLQYWSQELKNSAVSLLNATAIAVAEGTDRV  819
4   IVTRIVELLGR-------RGWEVLKYWWNLLQYWSQELKNSAVSLLNATAIAVAEGTDRV  819
5   IVTRIVELLGR-------RGWEVLKYWWNLLQYWSQELKNSAVSLLNATAIAVAEGTDRV  819
6   IVTRIVELLGR-------RGWEVLKYWWNLLQYWSQELKNSAVSLLNATAIAVAEGTDRV  819
7   IVTRIVELLGR-------RGWEVLKYWWNLLQYWSQELKNSAVSLLNATAIAVAEGTDRV  819
8   ---RIVELLGR-------RGWEVLKYWWNLLQYWSQELKNSAVSLLNATAIAVAEGTDRV  816
9   IVTRIVELLGR-------RGWEVLKYWWNLLQYWSQELKNSAVSLLNATAIAVAEGTDRV  818
10  IVTRIVELLGR-------RGWEVLKYWWNLLQYWSQELKNSAVSLLNATAIAVAEGTDRV  819
11  IVTRIVELLGR-------RGWEVLKYWWNLLQYWSQELKNSAVSLLNATAIAVAEGTDRV  819
12  IVTRIVELLGR-------RGWEVLKYWWNLLQYWSQELKNSAVSLLNATAIAVAEGTDRV  819
13  IVTRIVELLGR-------RGWEVLKYWWNLLQYWSQELKNSAVSLLNATAIAVAEGTDRV  819
14  IVTRIVELLGR-------RGWEVLKYWWNLLQYWSQELKNSAVSLLNATAIAVAEGTDRV  817
15  IVTRIVELLGR-------RGWEVLKYWWNLLQYWSQELKNSAVSLLNATAIAVAEGTDRV  787
16  IVTRIVELLGR-------RGWEVLKYWWNLLQYWSQELKNSAVSLLNATAIAVAEGTDRV  819
17  IVTRIVELLGR-------RGWEVLKYWWNLLQYWSQELKNSAVSLLNATAIAVAEGTDRV  819
18  IVTRIVELLGR-------RGWEVLKYWWNLLQYWSQELKNSAVSLLNATAIAVAEGTDRV  819
19  IVTRIVELLGR-------RGWEVLKYWWNLLQYWSQELKNSAVSLLNATAIAVAEGTDRV  819
20  IVTRIVELLGR-------RGWEVLKYWWNLLQYWSQELKNSAVSLLNATAIAVAEGTDRV  817
21  IVTRIVELLGR-------RGWEVLKYWWNLLQYWSQELKNSAVSLLNATAIAVAEGTDRV  819
22  IVTRIVELLGR-------RGWEVLKYWWNLLQYWSQELKNSAVSLLNATAIAVAEGTDRV  802
23  IVTRIVELLGR-------RGWEVLKYWWNLLQYWSQELKNSAVSLLNATAIAVAEGTDRV  819
24  IVTRIVELLGR-------RGWEVLKYWWNLLQYWSQELKNSAVSLLNATAIAVAEGTDRV  818
25  IVTRIVELLGR-------RGWEVLKYWWNLLQYWSQELKNSAVSLLNATAIAVAEGTDRV  819
26  IVTRIVELLGR-------RGWEVLKYWWNLLQYWSQELKNSAVSLLNATAIAVAEGTDRV  819
27  IVTRIVELLGR-------RGWEVLKYWWNLLQYWSQELKNSAVSLLNATAIAVAEGTDRV  819
28  IVTRIVELLGR-------RGWEVLKYWWNLLQYWSQELKNSAVSLLNATAIAVAEGTDRV  819
29  IATRIVELLGR-------RGWEVLKYWWNLLQYWSQELKNSAVSLLNATAIAVAEGTDRV  819
30  IVTRIVELLGR-------RGWEVLKYWWNLLQYWSQELKNSAVSLLNATAIAVAEGTDRV  819
31  IVTRIVELLGR-------RGWEVLKYWWNLLQYWSQELKNSAVSLLNATAIAVAEGTDRV  819
32  IVTRIVELLGR-------RGWEVLKYWWNLLQYWSQELKNSAVSLLNATAIAVAEGTDRV  819
33  IVTRIVELLGR-------RGWEVLKYWWNLLQYWSQELKNSAVSLLNATAIAVAEGTDRV  819
34  IVTRIVELLGR-------RGWEVLKYWWNLLQYWSQELKNSAVSLLNATAIAVAEGTDRV  819
35  IVTRIVELLGR-------RGWEVLKYWWNLLQYWSQELKNSAVSLLNATAIAVAEGTDRV  819
36  IVTRIVELLGR-------RGWEVLKYWWNLLQYWSQELKNSAVSLLNATAIAVAEGTDRV  819
37  IVTRIVELLGR-------RGWEVLKYWWNLLQYWSQELKNSAVSLLNATAIAVAEGTDRV  819
38  IVTRIVELLGR-------RGWEVLKYWWNLLQYWSQELKNSAVSLLNATAIAVAEGTDRV  817
```

| | | |
|---|---|---|
| 39 | IVTRIVELLGR-------RGWEVLKYWWNLLQYWSQELKNSAVSLLNATAIAVAEGTDRV | 819 |
| 40 | IVTRIVELLGR-------RGWEVLKYWWNLLQYWSQELKNSAVSLLNATAIAVAEGTDRV | 819 |
| 41 | IVTRIVELLGR-------RGWEVLKYWWNLLQYWSQELKNSAVSLLNATAIAVAEGTDRV | 819 |
| 42 | IVTRIVELLGR-------RGWEVLKYWWNLLQYWSQELKNSAVSLLNATAIAVAEGTDRV | 819 |
| 43 | IVTRIVELLGR-------RGWEVLKYWWNLLQYWSQELKNSAVSLLNATAIAVAEGTDRV | 819 |
| 44 | IVTRIVELLGR-------RGWEVLKYWWNLLQYWSQELKNSAVSLLNATAIAVAEGTDRV | 819 |
| 45 | IVTRIVELLGR-------RGWEVLKYWWNLLQYWSQELKNSAVSLLNATAIAVAEGTDRV | 819 |
| 46 | IVTRIVELLGR-------RGWEVLKYWWNLLQYWSQELKNSAVSLLNATAIAVAEGTDRV | 819 |
| 47 | IVTRIVELLGR-------RGWEVLKYWWNLLQYWSQELKNSAVSLLNATAIAVAEGTDRV | 819 |
| 48 | IVTRIVELLGR-------RGWEVLKYWWNLLQYWSQELKNSAVSLLNATAIAVAEGTDRV | 819 |
| 49 | IVTRIVELLGR-------RGWEVLKYWWNLLQYWSQELKNSAVSLLNATAIAVAEGTDRV | 819 |
| 50 | IVTRIVELLGR-------RGWEVLKYWWNLLQYWSQELKNSAVSLLNATAIAVAEGTDRV | 819 |
| 51 | IVTRIVELLGR-------RGWEVLKYWWNLLQYWSQKLKNSAVSLLNATAIAVAEGTDRV | 819 |
| 52 | IVTRIVELLGR-------RGWEVLKYWWNLLQYWSQELKNSAVSLLNATAIAVAEGTDRV | 819 |
| 53 | IVTRIVELLGR-------RGWEVLKYWWNLLQYWSQELKNSAVSLLNATAIAVAEGTDRV | 819 |
| 54 | IVTRIVELLGR-------RGWEVLKYWWNLLQYWSQELKNSAVSLLNATAIAVAEGTDRV | 819 |
| 55 | IVTRIVELLGR-------RGWEVLKYWWNLLQYWSQELKNSAVSLLNATAIAVAEGTDRV | 819 |
| 56 | IVTRIVELLGR-------RGWEVLKYWWNLLQYWSQELKNSAVSLLNATAIAVAEGTDRV | 819 |
| 57 | IVTRIVELLGR-------RGWEVLKYWWNLLQYWSQELKNSAVSLLNATAIAVAEGTDRV | 819 |
| 58 | IVTRIVELLGR-------RGWEVLKYWWNLLQYWSQELKNSAVSLLNATAIAVAEGTDRV | 819 |
| 59 | IVTRIVEPLGR-------RGWEVLKYWWNLLQYWSQELKNSAVSLLNATAIAVAEGTDRV | 817 |
| 60 | IVTRIVELLGR-------RGWEVLKYWWNLLQYWSQELKNSAVNLLNATAIAVAEGTDRV | 819 |
| 61 | IVTRIVELLGR-------RGWEVLKYWWNLLQYWSQELKNSAVSLLNATAIAVAEGTDRV | 819 |
| 62 | IVTRIVELLGR-------RGWEVLKYWWNLLQYWSQELKNSAVSLLNATAIAVAEGTDRV | 819 |
| 63 | IVTRIVELLGR-------RGWEVLKYWWNLLQYWSQELKNSAVSLLNATAIAVAEGTDRV | 819 |
| 64 | IVTRIVELLGR-------RGWEILKYWWNLLQYWSQELKNSAVSLLNATAIAVAEGTDRV | 819 |
| 65 | IVTRIVELLGR-------RGWEVLKYWWNLLQYWSQELKNSAVSLLNATAIAVAEGTDRV | 819 |
| 66 | IVTRIVKLLGR-------RGWEVLKYWWNLLQYWSQELKNSAVSLLNATAIAVAEGTDRV | 818 |
| 67 | IVTRIVELLGR-------RGWEVLKYWWNLLQYWSQELKNSAVSLLNATAIAVAEGTDRV | 817 |
| 68 | IVTRIVELLGR-------RGWEVLKYWWNLLQYWSQELKNSAVSLLNATAIAVAEGTDRV | 819 |

**IEVVQRACRAILHIPRRIRQGLERALL**

| | | |
|---|---|---|
| 1 | IEVVQRACRAILHIPRRIRQGLERALL | 846 |
| 2 | IEVVQRACRAILHIPRRIRQGLERALL | 845 |
| 3 | IEVVQRACRAILHIPRRIRQGLERALL | 846 |
| 4 | IEVVQRACRAILHIPRRIGQGLERALL | 846 |
| 5 | IEVVQRACRAILHIPRRIRQGLERALL | 846 |
| 6 | IEVVQRACRAILHIPRRIRQGLERALL | 846 |
| 7 | IEVVQRACRAILHIPRRIRQGLERALL | 846 |
| 8 | IEVVQRACRAILHIPRRIRQGLERALL | 843 |
| 9 | IEVVQRACRAILHIPRRIRQGLERALL | 845 |
| 10 | IEVVQRACRAILHIPRRIRQGLERALL | 846 |
| 11 | IEVVQRACRAILHIPRRIRQGLERALL | 846 |
| 12 | IEVVQRACRAILHIPRRIRQGLERALL | 846 |
| 13 | IEVVQRACRAILHIPRRIRQGLERALL | 846 |
| 14 | IEVVQRACRAILHIPRRIRQGLERALL | 844 |
| 15 | IEVVQRACRAILHIPRRIRQGLERALL | 814 |
| 16 | IEVVQRACRAILHIPRRIRQGLERALL | 846 |
| 17 | IEVVQRACRAILHIPRRIRQGLERALL | 846 |
| 18 | IEVVQRACRATLHIPRRIRQGLERALL | 846 |
| 19 | IEVVQRACRAILHIPRRIRQGLERALL | 846 |
| 20 | IEVVQRACRAILHIPRRIRQGLERALL | 844 |

| | | |
|---|---|---|
| 21 | IEVVQRACRAILHIPRRIRQGLERALL | 846 |
| 22 | IEVVQRACRAILHIPRRIRQGLERALL | 829 |
| 23 | IEVVQRACRAILHIPRRIRQGLERALL | 846 |
| 24 | IEVVQRACRAILHIPRRIRQGLERALL | 845 |
| 25 | IEVVQRACRAILHIPRRIRQGLERALL | 846 |
| 26 | IEVVQRACRAILHIPRRIRQGLERALL | 846 |
| 27 | IEVVQRACRAILHIPRRIRQGLERALL | 846 |
| 28 | IEVVQRACRAILHIPRRIRQGLERALL | 846 |
| 29 | IEVVQRACRAILHIPRRIRQGLERALL | 846 |
| 30 | IEVVQRACRAILHIPRRIRQGLERALL | 846 |
| 31 | IEVVQRACRAILHIPRRIRQGLERALL | 846 |
| 32 | IEVVQRACRAILHIPRRIRQGLERALL | 846 |
| 33 | IEVVQRACRAILHIPRRIRQGLERALL | 846 |
| 34 | IEVVQRACRAILHIPRRIRQGLERALL | 846 |
| 35 | IEVVQRACRAILHTPRRIRQGLERALL | 846 |
| 36 | IEVVQRACRAILHIPRRIRQGLERALL | 846 |
| 37 | IEVVQRACRAILHIPRRIRQGLERALL | 846 |
| 38 | IEVVQRACRAILHIPRRIRQGLERALL | 844 |
| 39 | IEVVQRACRAILHIPRRIRQGLERALL | 846 |
| 40 | IEVVQRACRAILHIPRRIRQGLERALL | 846 |
| 41 | IEVVQRACRAILHIPRRIRQGLERALL | 846 |
| 42 | IEVVQRACRATLHIPRRIRQGLERALL | 846 |
| 43 | IEVVQRACRATLHIPRRIRQGLERALL | 846 |
| 44 | IEVVQRACRAIIHIPRRIRQGLERALL | 846 |
| 45 | IEVVQRACRATLHIPRRIRQGLERALL | 846 |
| 46 | IEVVQRACRAILHIPRRIRQGLERALL | 846 |
| 47 | IEVVQRACRAILHIPRRIRQGLERALL | 846 |
| 48 | IEVVQRACRATLHIPRRIRQGLERALL | 846 |
| 49 | IEVVQRACRAILHIPRRIRQGLERALL | 846 |
| 50 | IEVVQRACRAILHIPRRIRQGLERALL | 846 |
| 51 | IEVVQRACRAILHIPRRIRQGLERALL | 846 |
| 52 | IEVVQRACRATLHIPRRIRQGLERALL | 846 |
| 53 | IEVVQRACRAILHIPRRIRQGLERALL | 846 |
| 54 | IEVVQRACRAILHIPRRIRQGLERALL | 846 |
| 55 | IEVVQRACRAILHIPRRIRQGLERALL | 846 |
| 56 | IEVVQRACRATLHIPRRIRQGLERALL | 846 |
| 57 | IEVVQRACRAILHIPRRIRQGLERALL | 846 |
| 58 | IEVVQRACRAILHIPRRIRQGLERALL | 846 |
| 59 | IEVVQRACRAILHIPRRIRQGLERALL | 844 |
| 60 | IEVVQRACRATLHIPRRIRQGLERALL | 846 |
| 61 | IEVVQRACRAILHIPIRIRQGLERALL | 846 |
| 62 | IEVVQRACRAILHIPRRIRQGLERALL | 846 |
| 63 | IEVVQRACRAILHTPRRIRQGLERALL | 846 |
| 64 | IEVVQRACRATLHIPRRIRQGLERALL | 846 |
| 65 | IEVVQRACRATLHIPRRIRQGLERALL | 846 |
| 66 | IEVVQRACRATLHIPRRIRQGLERALL | 845 |
| 67 | IEVVQRACRAILHIPRRIRQGLERALL | 844 |
| 68 | IEVVQRACRATLHIPRRIRQGLERALL | 846 |

**Consensus**

1. B.US.2006.700010040_C9_4520.EU289193
2. B.US.2007.CH0040_3_d0197_ipe032_27_50.MG900200
3. B.US.2007.CH0040_3_d0197_ipe032_2_03.MG900201
4. B.US.2007.CH0040_3_d0197_ipe032_2_04.MG900202
5. B.US.2007.CH0040_3_d0197_ipe032_2_10.MG900203
6. B.US.2007.CH0040_3_d0197_ipe032_2_12.MG900204
7. B.US.2007.CH0040_3_d0197_ipe032_2_13.MG900205
8. B.US.2007.CH0040_3_d0197_ipe032_2_14.MG900206
9. B.US.2007.CH0040_3_d0197_ipe032_2_17.MG900207
10. B.US.2007.CH0040_3_d0197_ipe032_2_18.MG900208
11. B.US.2007.CH0040_3_d0197_ipe032_2_33.MG900209
12. B.US.2007.CH0040_3_d0197_ipe032_2_35.MG900210
13. B.US.2007.CH0040_3_d0197_ipe032_2_36.MG900211
14. B.US.2007.CH0040_3_d0197_ipe032_2_38.MG900212
15. B.US.2007.CH0040_3_d0197_ipe032_2_39.MG900213
16. B.US.2007.CH0040_3_d0197_ipe032_2_42.MG900214
17. B.US.2007.CH0040_3_d0197_ipe032_2_43.MG900215
18. B.US.2007.CH0040_3_d0197_ipe032_3_46.MG900216
19. B.US.2007.CH0040_3_d0197_ipe032_3_52.MG900217
20. B.US.2007.CH0040_3_d0197_ipe032_3_54.MG900218
21. B.US.2007.CH0040_3_d0197_ipe032_3_55.MG900219
22. B.US.2007.CH0040_3_d0197_ipe032_3_56.MG900220
23. B.US.2007.CH0040_3_d0197_ipe032_3_59.MG900221
24. B.US.2007.CH0040_3_d0197_ipe032_3_60.MG900222
25. B.US.2007.CH0040_3_d0197_ipe032_3_61.MG900223
26. B.US.2007.CH0040_3_d0197_ipe032_3_64.MG900224
27. B.US.2007.CH0040_3_d0197_ipe032_3_65.MG900225
28. B.US.2007.CH0040_3_d0197_ipe032_3_66.MG900226
29. B.US.2007.CH0040_3_d0197_ipe032_3_68.MG900227
30. B.US.2007.CH0040_3_d0197_ipe032_3_69.MG900228
31. B.US.2007.CH0040_3_d0197_ipe032_3_70.MG900229
32. B.US.2007.CH0040_3_d0197_ipe032_3_74.MG900230
33. B.US.2007.CH0040_3_d0197_ipe032_3_75.MG900231

| | | |
|---|---|---|
| | **MRVMGIRKNYQHLWREGILLLGILMICSAADNLWVTVYYGVPVWREATTTLFCASDAKAY** | |
| 1 | MRVMGIRKNYQHLWREGILLLGILMICSAADNLWVTVYYGVPVWREATTTLFCASDAKAY | 60 |
| 2 | MRVMGIRKNYQHLWREGILLLGILMICSAADNLWVTVYYGVPVWREATTTLFCASDAKAY | 60 |
| 3 | MRVMGIRKNYQHLWREGILLLGILMICSAADNLWVTVYYGVPVWREATTTLFCASDAKAY | 60 |
| 4 | MRVMGIRKNYQHLWREGILLLGILMICSAADNLWVTVYYGVPVWREATTTLFCASDAKAY | 60 |
| 5 | MRVMGIRKNYQHLWREGILLLGILMICSAADNLWVTVYYGVPVWREATTTLFCASDAKAY | 60 |
| 6 | MRVMGIRKNYQHLWREGILLLGILMICSAADNLWVTVYYGVPVWREATTTLFCASDAKAY | 60 |
| 7 | MRVMGIRKNYQHLWREGILLLGILMICSAADNLWVTVYYGVPVWREATTTLFCASDAKAY | 60 |
| 8 | MRVMGIRKNYQHLWREGILLLGILMICSAADNLWVTVYYGVPVWREATTTLFCASDAKAY | 60 |
| 9 | MRVMGIRKNYQHLWREGILLLGILMICSAADNLWVTVYYGVPVWREATTTLFCASDAKAY | 60 |
| 10 | MRVMGIRKNYQHLWREGILLLGILMICSAADNLWVTVYYGVPVWREATTTLFCASDAKAY | 60 |
| 11 | MRVMGIRKNYQHLWREGILLLGILMICSAADNLWVTVYYGVPVWREATTTLFCASDAKAY | 60 |
| 12 | MRVMGIRKNYQHLWREGILLLGILMICSAADNLWVTVYYGVPVWREATTTLFCASDAKAY | 60 |
| 13 | MRVMGIRKNYQHLWREGILLLGILMICSAADNLWVTVYYGVPVWREATTTLFCASDAKAY | 60 |
| 14 | MRVMGIRKNYQHLWREGILLLGILMICSAADNLWVTVYYGVPVWREATTTLFCASDAKAY | 60 |
| 15 | MRVMGIRKNYQHLWREGILLLGILMICSAADNLWVTVYYGVPVWREATTTLFCASDAKAY | 60 |
| 16 | MRVMGIRKNYQHLWREGILLLGILMICSAADNLWVTVYYGVPVWREATTTLFCASDAKAY | 60 |
| 17 | MRVMGIRKNYQHLWREGILLLGILMICSAADNLWVTVYYGVPVWREATTTLFCASDAKAY | 60 |
| 18 | MRVMGIRKNYQHLWREGILLLGILMICSAADNLWVTVYYGVPVWREATTTLFCASDAKAY | 60 |
| 19 | MRVMGIRKNYQHLWREGILLLGILMICSAADNLWVTVYYGVPVWREATTTLFCASDAKAY | 60 |
| 20 | MRVMGIRKNYQHLWREGILLLGILMICSAADNLWVTVYYGVPVWREATTTLFCASDAKAY | 60 |
| 21 | MRVMGIRKNYQHLWREGILLLGILMICSAADNLWVTVYYGVPVWREATTTLFCASDAKAY | 60 |
| 22 | MRVMGIRKNYQHLWREGILLLGILMICSAADNLWVTVYYGVPVWREATTTLFCASDAKAY | 60 |
| 23 | MRVMGIRKNYQHLWREGILLLGILMICSAADNLWVTVYYGVPVWREATTTLFCASDAKAY | 60 |
| 24 | MRVMGIRKNYQHLWREGILLLGILMICSAADNLWVTVYYGVPVWREATTTLFCASDAKAY | 60 |
| 25 | MRVMGIRKNYQHLWREGILLLGILMICSAADNLWVTVYYGVPVWREATTTLFCASDAKAY | 60 |
| 26 | MRVMGIRKNYQHLWREGILLLGILMICSAADNLWVTVYYGVPVWREATTTLFCASDAKAY | 60 |
| 27 | MRVMGIRKNYQHLWREGILLLGILMICSAADNLWVTVYYGVPVWREATTTLFCASDAKAY | 60 |
| 28 | MRVMGIRKNYQHLWREGILLLGILMICSAADNLWVTVYYGVPVWREATTTLFCASDAKAY | 60 |
| 29 | MRVMGIRKNYQHLWREGILLLGILMICSAADNLWVTVYYGVPVWREATTTLFCASDAKAY | 60 |
| 30 | MRVMGIRKNYQHLWREGILLLGILMICSAADNLWVTVYYGVPVWREATTTLFCASDAKAY | 60 |
| 31 | MRVMGIRKNYQHLWREGILLLGILMICSAADNLWVTVYYGVPVWREATTTLFCASDAKAY | 60 |
| 32 | MRVMGIRKNYQHLWREGILLLGILMICSAADNLWVTVYYGVPVWREATTTLFCASDAKAY | 60 |
| 33 | MRVMGIRKNYQHLWREGILLLGILMICSAADNLWVTVYYGVPVWREATTTLFCASDAKAY | 60 |

**DTEAHNVWATHACVPTDPNPQEVELKNVTENFNMWENNMVEQMHEDIISLWDQSLKPCVK**

| | | |
|---|---|---|
| 1 | DTEAHNVWATHACVPTDPNPQEVELKNVTENFNMWENNMVEQMHEDIISLWDQSLKPCVK | 120 |
| 2 | DTEAHNVWATHACVPTDPNPQEVELKNVTENFNMWENNMVEQMHEDIISLWDQSLKPCVK | 120 |
| 3 | DTEAHNVWATHACVPTDPNPQEVELKNVTENFNMWENNMVEQMHEDIISLWDQSLKPCVK | 120 |
| 4 | DTEAHNVWATHACVPTDPNPQEVELKNVTENFNMWENNMVEQMHEDIISLWDQSLKPCVK | 120 |
| 5 | DTEAHNVWATHACVPTDPNPQEVELKNVTENFNMWENNMVEQMHEDIISLWDQSLKPCVK | 120 |
| 6 | DTEAHNVWATHACVPTDPNPQEVELKNVTENFNMWENNMVEQMHEDIISLWDQSLKPCVK | 120 |
| 7 | DTEAHNVWATHACVPTDPNPQEVELKNVTENFNMWENNMVEQMHEDIISLWDQSLKPCVK | 120 |
| 8 | DTEAHNVWATHACVPTDPNPQEVELKNVTENFNMWENNMVEQMHEDIISLWDQSLKPCVK | 120 |
| 9 | DTEAHNVWATHACVPTDPNPQEVELKNVTENFNMWENNMVEQMHEDIISLWDQSLKPCVK | 120 |
| 10 | DTEAHNVWATHACVPTDPNPQEVELKNVTENFNMWENNMVEQMHEDIISLWDQSLKPCVK | 120 |
| 11 | DTEAHNVWATHACVPTDPNPQEVELKNVTENFNMWENNMVEQMHEDIISLWDQSLKPCVK | 120 |
| 12 | DTEAHNVWATHACVPTDPNPQEVELKNVTENFNMWENNMVEQMHEDIISLWDQSLKPCVK | 120 |
| 13 | DTEAHNVWATHACVPTDPNPQEVELKNVTENFNMWENNMVEQMHEDIISLWDQSLKPCVK | 120 |
| 14 | DTEAHNVWATHACVPTDPNPQEVELKNVTENFNMWENNMVEQMHEDIISLWDQSLKPCVK | 120 |
| 15 | DTEAHNVWATHACVPTDPNPQEVELKNVTENFNMWENNMVEQMHEDIISLWDQSLKPCVK | 120 |
| 16 | DTEAHNVWATHACVPTDPNPQEVELKNVTENFNMWENNMVEQMHEDIISLWDQSLKPCVK | 120 |
| 17 | DTEAHNVWATHACVPTDPNPQEVELKNVTENFNMWENNMVEQMHEDIISLWDQSLKPCVK | 120 |
| 18 | DTEAHNV-ATHACVPTDPNPQEVELKNVTENFNMWENNMVEQMHEDIISLWDQSLKPCVK | 119 |
| 19 | DTEAHNVWATHACVPTDPNPQEVELKNVTENFNMWENNMVEQMHEDIISLWDQSLKPCVK | 120 |
| 20 | DTEAHNVWATHACVPTDPNPQEVELKNVTENFNMWKNNMVEQMHEDIISLWDQSLKPCVK | 120 |
| 21 | DTEAHNVWATHACVPTDPNPQEVELKNVTENFNMWENNMVEQMHEDIISLWDQSLKPCVK | 120 |
| 22 | DTEAHNVWATHACVPTDPNPQEVELKNVTENFNMWENNMVEQMHEDIISLWDQSLKPCVK | 120 |
| 23 | DTEAHNVWATHACVPTDPNPQEVELKNVTENFNMWENNMVEQMHEDIISLWDQSLKPCVK | 120 |
| 24 | DTEAHNVWATHACVPTDPNPQEVELKNVTENFNMWENNMVEQMHEDIISLWDQSLKPCVK | 120 |
| 25 | DTEAHNVWATHACVPTDPNPQEVELKNVTENFNMWENNMVEQMHEDIISLWDQSLKP-VK | 119 |
| 26 | DTEAHNVWATHACVPTDPNPQEVELKNVTENFNMWENNMVEQMHEDIISLWDQSLKPCVK | 120 |
| 27 | DTEAHNVWATHACVPTDPNPQEVELKNVTENFNMWENNMVEQMHEDIISLWDQSLKPCVK | 120 |
| 28 | DTEAHNVWATHACVPTDPNPQEVELKNVTENFNMWENNMVEQMHEDIISLWDQSLKPCVK | 120 |
| 29 | DTEAHNVWATHACVPTDPNPQEVELKNVTENFNMWENNMVEQMHEDIISLWDQSLKPCVK | 120 |
| 30 | DTEAHNVWATHACVPTDPNPQEVELKNVTENFNMWENNMVEQMHEDIISLWDQSLKPCVK | 120 |
| 31 | DTEAHNVWATHACVPTDPNPQEVELKNVTENFNMWENNMVEQMHEDIISLWDQSLKPCVK | 120 |
| 32 | DTEAHNVWATHACVPTDPNPQEVELKNVTENFNMWENNMVEQMHEDIISLWDQSLKPCVK | 120 |
| 33 | DTEAHNVWATHACVPTDPNPQEVELKNVTENFNMWENNMVEQMHEDIISLWDQSLKPCVK | 120 |

| | | |
|---|---|---|
| | LTPLCVTLNCTDLGNVTNTT---NSNGGMMEKGEVKNCSFKITTDIKDRTRKEYALFYKL | |
| 1 | LTPLCVTLNCTDLGNVTNTT---NSNGEMMEKGEVKNCSFKITTDIKDRTRKEYALFYKL | 177 |
| 2 | LTPLCVTLNCTDLGNVTNTT---NSNGGMMEKGEVKNCSFKITTDIKDRTRKEYALFYKL | 177 |
| 3 | LTPLCVTLNCTDLGNVTNTT---NSNGGMMEKGEVKNCSFKITTDIKDRTRKEYALFYKL | 177 |
| 4 | LTPLCVTLNCTDLGNVTNTT---NSNGGMMEKGEVKNCSFKITTDIKDRTRKEYALFYKL | 177 |
| 5 | LTPLCVTLNCTDLGNVTNTT---NSNGGMMEKGEVKNCSFKITTDIKDRTRKEYALFYKL | 177 |
| 6 | LTPLCVTLNCTDLGNVTNTT---NSNGGMMEKGEVKNCSFKITTDIKDRTRKEYALFYKL | 177 |
| 7 | LTPLCVTLNCTDLGNVTNTT---NSNGKMMEKGEVKNCSFKITTDIKDRTRKEYALFYKL | 177 |
| 8 | LTPLCVTLNCTDLGNVTNTT---NSNGGMMEKGEVKNCSFKITTDIKDRTRKEYALFYKL | 177 |
| 9 | LTPLCVTLNCTDLGNVTNTT---NSNEEMMEKGEVKNCSFKITTDIKDRTRKEYALFYKL | 177 |
| 10 | LTPLCVTLNCTDLGNVTNTT---NSNGGMMEKGEVKNCSFKITTDIKDRTRKEYALFYKL | 177 |
| 11 | LTPLCVTLNCTDLGNVTNTT---NSNGGMMEKGEVKNCSFKITTDIKDRTRKEYALFYKL | 177 |
| 12 | LTPLCVTLNCTDLGNVTNTT---NSNGGMMEKGEVKNCSFKITTDIKDRTRKEYALFYKL | 177 |
| 13 | LTPLCVTLNCTDLGNVTNTT---NSNGKMMEKGEVKNCSFKITTDIKDRTRKEYALFYKL | 177 |
| 14 | LTPLCVTLNCTDLGNVTNTT---NSNGGMMEKGEVKNCSFKITTDIKDRTRKEYALFYKL | 177 |
| 15 | LTPLCVTLNCTDLGNVTNIT---NSNGGMMEKGEVKNCSFKITTDIKDRTRKEYALFYKL | 177 |
| 16 | LTPLCVTLNCTDLGNVTNTT---NSNGGMMEKGEVKNCSFKITTDIKDRTRKEYALFYKL | 177 |
| 17 | LTPLCVTLNCTDLGNVTNTT---NSNGGMMEKGEVKNCSFKITTDIKDRTRKEYALFYKL | 177 |
| 18 | LTPLCVTLNCTDLGNVTNTT---NSKGEMMEKGEVKNCSFKITTDIKDRTRKEYALFYKL | 176 |
| 19 | LTPLCVTLNCTDLGNVTNTT---NSNGGMMEKGEVKNCSFKITTDIKDRTRKEYALFYKL | 177 |
| 20 | LTPLCVTLNCTDLGNVTNTT---NSNGKMMEKGEVKNCSFKITTDIKDRTRKEYALFYKL | 177 |
| 21 | LTPLCVTLNCTDLGNVTNTT---NSNGGMMEKGEVKNCSFKITTDIKDRTRKEYALFYKL | 177 |
| 22 | LTPLCVTLNCTDLGNVTNTT---NSNGGMMEKGEVKNCSFKITTDIKDRTRKEYALFYKL | 177 |
| 23 | LTPLCVTLNCTDLGNVTNTT---NSNGGMMEKGEVKNCSFKITTDIKDRTRKEYALFYKL | 177 |
| 24 | LTPLCVTLNCTDLGNVTNTT---NSNGGMMEKGEVKNCSFKITTDIKDRTRKEYALFYKL | 177 |
| 25 | LTPLCVTLNCTDLGNVTNTT---NSNGGMMEKGEVKNCSFKITTDIKDRTRKEYALFYKL | 176 |
| 26 | LTPLCVTLNCTDLGNVTNTT---NSNGGMMEKGEVKNCSFKITTDIKDRTRKEYALFYKL | 177 |
| 27 | LTPLCVTLNCTDLGNVTNTT---NSNGKMMEKGEVKNCSFKITTDIKDRTRKEYALFYKL | 177 |
| 28 | LTPLCVTLNCTDLGNVTNTT---NSNGGMMEKGEVKNCSFKITTDIKDRTRKEYALFYKL | 177 |
| 29 | LTPLCVTLNCTDLGNVTNTT---NSNGKMMEKGEVKNCSFKITTDIKDRTRKEYALFYKL | 177 |
| 30 | LTPLCVTLNCTDLGNVTNTT---NSKGEMMEKGEVKNCSFKITTDIKDRTRKEYALFYKL | 177 |
| 31 | LTPLCVTLNCTDLGNVTNTT---NSNGKMMEKGEVKNCSFKITTDIKDRTRKEYALFYKL | 177 |
| 32 | LTPLCVTLNCTDLGNVTNTT---NSNGGMMEKGEVKNCSFKITTDIKDRTRKEYALFYKL | 177 |
| 33 | LTPLCVTLNCTDLGNVTNTT---NSNGGMMEKGEVKNCSFKITTDIKDRTRKEYALFYKL | 177 |

| | | |
|---|---|---|
|  | **DVVPIND-------TRYRLVSCNTSVITQACPKVSFEPIPIHYCAPAGFAILKCNDKQFI** |  |
| 1 | DVVPIND-------TRYRLVSCNTSVITQACPKVSFEPIPIHYCAPAGFAILKCNDKQFI | 230 |
| 2 | DVVPIND-------TRYRLVSCNTSVITQACPKVSFEPIPIHYCAPAGFAILKCNDKQFI | 230 |
| 3 | DVVPIND-------TRYRLVSCNTSVITQACPKVSFEPIPIHYCAPAGFAILKCNDKQFI | 230 |
| 4 | DVVPIND-------TRYRLVSCNTSVITQACPKVSFEPIPIHYCAPAGFAILKCNDKQFI | 230 |
| 5 | DVVPIND-------TRYRLVSCNTSVITQACPKVSFEPIPIHYCAPAGFAILKCNDKQFI | 230 |
| 6 | DVVPIND-------TRYRLVSCNTSVITQACPKVSFEPIPIHYCAPAGFAILKCNDKQFI | 230 |
| 7 | DVVPIND-------TRYRLVSCNTSVITQACPKVSFEPIPIHYCAPAGFAILKCNDKQFI | 230 |
| 8 | DVVPIND-------TRYRLVSCNTSVITQACPKVSFEPIPIHYCAPAGFAILKCNGKQFI | 230 |
| 9 | DVVPIND-------TRYRLVSCNTSVITQACPKVSFEPIPIHYCAPAGFAILKCNDKQFI | 230 |
| 10 | DVVPIND-------TRYRLVSCNTSVITQACPKVSFEPIPIHYCAPAGFAILKCNDKQFI | 230 |
| 11 | DVVPIND-------TRYRLVSCNTSVITQACPKVSFEPIPIHYCAPAGFAILKCNDKQFI | 230 |
| 12 | DVVPIND-------TRYRLVSCNTSVITQACPKVSFEPIPIHYCAPAGFAILKCNDKQFI | 230 |
| 13 | DVVPIND-------TRYRLVSCNTSVITQACPKVSFEPIPIHYCAPAGFAILKCNDKQFI | 230 |
| 14 | DVVPIND-------TRYRLVSCNTSVITQACPKVSFEPIPIHYCAPAGFAILKCNDKQFI | 230 |
| 15 | DVVPIND-------TRYRLVSCNTSVITQACPKVSFEPIPIHYCAPAGFAILKCNDKQFI | 230 |
| 16 | DVVPIND-------TRYRLVSCNTSVITQACPKVSFEPIPIHYCAPAGFAILKCNDKQFI | 230 |
| 17 | DVVPIND-------TRYRLVSCNTSVITQACPKVSFEPIPIHYCAPAGFAILKCNDKQFI | 230 |
| 18 | DVVPIND-------TRYRLVSCNTSVITQACPKVSFEPIPIHYCAPAGFAILKCNDKQFI | 229 |
| 19 | DVVPIND-------TRYRLVSCNTSVITQACPKVSFEPIPIHYCAPAGFAILKCNDKQFI | 230 |
| 20 | DVVPIND-------TRYRLVSCNTSVITQACPKVSFEPIPIHYCAPAGFAILKCNDKQFI | 230 |
| 21 | DVVPIND-------TRYRLVSCNTSVITQACPKVSFEPIPIHYCAPAGFAILKCNDKQFI | 230 |
| 22 | DVVPIND-------TRYRLVSCNTSVITQACPKVSFEPIPIHYCAPAGFAILKCNDKQFI | 230 |
| 23 | DVVPIND-------TRYRLVSCNTSVITQACPKVSFEPIPIHYCAPAGFAILKCNDKQFI | 230 |
| 24 | DVVPIND-------TRYRLVSCNTSVITQACPKVSFEPIPIHYCAPAGFAILKCNDKQFI | 230 |
| 25 | DVVPIND-------TRYRLVSCNTSVITQACPKVSFEPIPIHYCAPAGFAILKCNDKQFI | 229 |
| 26 | DVVPIND-------TRYRLVSCNTSVITQACPKVSFEPIPIYYCAPAGFAILKCNDKQFI | 230 |
| 27 | DVVPIND-------TRYRLVSCNTSVITQACPKVSFEPIPIHYCAPAGFAILKCNDKQFI | 230 |
| 28 | DVVPIND-------TRYRLVSCNTSVITQACPKVSFEPIPIHYCAPAGFAILKCNDKQFI | 230 |
| 29 | DVVPIND-------TRYRLVSCNTSVITQACPKVSFEPIPIHYCAPAGFAILKCNDKQFI | 230 |
| 30 | DVVPIND-------TRYRLVSCNTSVITQACPKVSFEPIPIHYCAPAGFAILKCNDKQFI | 230 |
| 31 | DVVPIND-------TRYRLVSCNTSVITQACPKVSFEPIPIHYCAPAGFAILKCNDKQFI | 230 |
| 32 | DVVPIND-------TRYRLVSCNTSVITQACPKVSFEPIPIHYCAPAGFAILKCNDKQFI | 230 |
| 33 | DVVPIND-------TRYRLVSCNTSVITQACPKVSFEPIPIHYCAPAGFAILKCNDKQFI | 230 |

| | | |
|---|---|---|
| | GTGPCTNVSTVQCTHGIRPVVSTQLLLNGSLAEEEVVIRSVNFSDNAKTIIVQLNKSVEI | |
| 1 | GTGPCTNVSTVQCTHGIRPVVSTQLLLNGSLAEEEVVIRSVNFSDNAKTIIVQLNKSVEI | 290 |
| 2 | GTGPCTNVSTVQCTHGIRPVVSTQLLLNGSLAEEEVVIRSVNFSDNAKTIIVQLNKSVEI | 290 |
| 3 | GTGPCTNVSTVQCTHGIRPVVSTQLLLNGSLAEEEVVIRSVNFSDNAKTIIVQLNKSVEI | 290 |
| 4 | GTGPCTNVSTVQCTHGIRPVVSTQLLLNGSLAEEEVVIRSVNFSDNAKTIIVQLNKSVEI | 290 |
| 5 | GTGPCTNVSTVQCTHGIRPVVSTQLLLNGSLAEEEVVIRSVNFSDNAKTIIVQLNKSVEI | 290 |
| 6 | GTGPCTNVSTVQCTHGIRPVVSTQLLLNGSLAEEEVVIRSVNFSDNAKTIIVQLNKSVEI | 290 |
| 7 | GTGPCTNVSTVQCTHGIRPVVSTQLLLNGSLAEEEVVIRSVNFSDNAKTIIVQLNKSVEI | 290 |
| 8 | GTGPCTNVSTVQCTHGIRPVVSTQLLLNGSLAEEEVVIRSVNFSDNAKTIIVQLNKSVEI | 290 |
| 9 | GTGPCTNVSTVQCTHGIRPVVSTQLLLNGSLAEEEVVIRSVNFSDNAKTIIVQLNKSVEI | 290 |
| 10 | GTGPCTNVSTVQCTHGIRPVVSTQLLLNGSLAEEEVVIRSVNFSDNAKTIIVQLNKSVEI | 290 |
| 11 | GTGPCTNVSTVQCTHGIRPVVSTQLLLNGSLAEEEVVIRSVNFSDNAKTIIVQLNKSVEI | 290 |
| 12 | GTGPCTNVSTVQCTHGIRPVVSTQLLLNGSLAEEEVVIRSVNFSDNAKTIIVQLNKSVEI | 290 |
| 13 | GTGPCTNVSTVQCTHGIRPVVSTQLLLNGSLAEEEVVIRSVNFSDNAKTIIVQLNKSVEI | 290 |
| 14 | GTGPCTNVSTVQCTHGIRPVVSTQLLLNGSLAEEEVVIRSVNFSDNAKTIIVQLNKSVEI | 290 |
| 15 | GTGPCTNVSTVQCTHGIRPVVSTQLLLNGSLAEEEVVIRSVNFSDNAKTIIVQLNKSVEI | 290 |
| 16 | GTGPCTNVSTVQCTHGIRPVVSTQLLLNGSLAEEEVVIRSVNFSDNAKTIIVQLNKSVEI | 290 |
| 17 | GTGPCTNVSTVQCTHGIRPVVSTQLLLNGSLAEEEVVIRSVNFSDNAKTIIVQLNKSVEI | 290 |
| 18 | GTGPCTNVSTVQCTHGIRPVVSTQLLLNGSLAEEEVVIRSVNFSDNAKTIIVQLNKSVEI | 289 |
| 19 | GTGPCTNVSTVQCTHGIRPVVSTQLLLNGSLAEEEVVIRSVNFSDNAKTIIVQLNKSVEI | 290 |
| 20 | GTGPCTNVSTVQCTHGIRPVVSTQLLLNGSLAEEEVVIRSVNFSDNAKTIIVQLNKSVEI | 290 |
| 21 | GTGPCTNVSTVQCTHGIRPVVSTQLLLNGSLAEEEVVIRSVNFSDNAKTIIVQLNKSVEI | 290 |
| 22 | GTGPCTNVSTVQCTHGIRPVVSTQLLLNGSLAEEEVVIRSVNFSDNAKTIIVQLNKSVEI | 290 |
| 23 | GTGPCTNVSTVQCTHGIRPVVSTQLLLNGSLAEEEVVIRSVNFSDNAKTIIVQLNKSVEI | 290 |
| 24 | GTGPCTNVSTVQCTHGIRPVVSTQLLLNGSLAEEEVVIRSVNFSDNAKTIIVQLNKSVEI | 290 |
| 25 | GTGPCTNVSTVQCTHGIRPVVSTQLLLNGSLAEEEVVIRSVNFSDNAKTIIVQLNKSVEI | 289 |
| 26 | GTGPCTNVSTVQCTHGIRPVVSTQLLLNGSLAEEEVVIRSVNFSDNAKTIIVQLNKSVEI | 290 |
| 27 | GTGPCTNVSTVQCTHGIRPVVSTQLLLNGSLAEEEVVIRSVNFSDNAKTIIVQLNKSVEI | 290 |
| 28 | GTGPCTNVSTVQCTHGIRPVVSTQLLLNGSLAEEEVVIRSVNFSDNAKTIIVQLNKSVEI | 290 |
| 29 | GTGPCTNVSTVQCTHGIRPVVSTQLLLNGSLAEEEVVIRSVNFSDNAKTIIVQLNKSVEI | 290 |
| 30 | GTGPCTNVSTVQCTHGIRPVVSTQLLLNGSLAEEEVVIRSVNFSDNAKTIIVQLNKSVEI | 290 |
| 31 | GTGPCTNVSTVQCTHGIRPVVSTQLLLNGSLAEEEVVIRSVNFSDNAKTIIVQLNKSVKI | 290 |
| 32 | GTGPCTNVSTVQCTHGIRPVVSTQLLLNGSLAEEEVVIRSVNFSDNAKTIIVQLNKSVEI | 290 |
| 33 | GTGPCTNVSTVQCTHGIRPVVSTQLLLNGSLAEEEVVIRSVNFSDNAKTIIVQLNKSVEI | 290 |

| | TCTRPNNNTRKSIPMGPGKAFYARGDITGDIKKAYCKINGTEWHSTLKLVVEKLREQY-N | |
|---|---|---|
| 1 | TCTRPNNNTRKSIPMGPGKAFYARGDITGDIRKAYCEINGTEWHSTLKLVVEKLREQY-N | 349 |
| 2 | TCTRPNNNTRKSIPMGPGKAFYARGDITGDIKKAYCKINGTEWHSTLKLVVEKLREQY-N | 349 |
| 3 | TCTRPNNNTRKSIPMGPGKAFYARGDITGDIKKAYCKINGTEWHSTLKLVVEKLREQY-N | 349 |
| 4 | TCTRPNNNTRKSIPMGPGKAFYARGDITGDIKKAYCEINGTEWHSTLKLVVEKLREQY-N | 349 |
| 5 | TCTRPNNNTRKSIPMGPGKAFYARGDITGDIKKAYCKINGTEWHSTLKLVVEKLREQY-N | 349 |
| 6 | TCTRPNNNTRKSIPMGPGKAFYARGDITGDIKKAYCKINGTEWHSTLKLVVEKLREQY-N | 349 |
| 7 | TCIRPNNNTRKSIPMGPGKAFYARGDITGDIKKAYCEINGTEWHSTLKLVVEKLREQY-N | 349 |
| 8 | TCTRPNNNTRKSIPMGPGKAFYARGDITGDIKKAYCKINGTEWHSTLKLVVEKLREQY-N | 349 |
| 9 | TCTRPHNNTRKSIPMGPGKAFYARGDITGDIRKAYCEINGTEWHSTLKLVVEKLREQY-N | 349 |
| 10 | TCTRPNNNTRKSIPMGPGKAFYARGDITGDIKKAYCKINGTEWHSTLKLVVEKLREQY-N | 349 |
| 11 | TCTRPNNNTRKSIPMGPGKAFYARGDITGDIKKAYCKINGTEWHSTLKLVVEKLREQY-N | 349 |
| 12 | TCTRPNNNTRKSIPMGPGKAFYARGDITGDIKKAYCKINGTEWHSTLKLVVEKLREQY-N | 349 |
| 13 | TCTRPNNNTRKSIPMGPGKAFYARGDITGDIKKAYCEINGTEWHSTLKLVVEKLREQY-N | 349 |
| 14 | TCTRPNNNTRKSIPMGPGKAFYARGDITGDIKKAYCKINGTEWHSTLKLVVEKLREQY-N | 349 |
| 15 | TCTRPNNNTRKSIPMGPGKAFYARGDITGDIKKAYCKINGTEWHSTLKLVVEKLREQY-N | 349 |
| 16 | TCIRPNNNTRKSIPMGPGKAFYARGDITGDIKKAYCEINGTEWHSTLKLVVEKLREQY-N | 349 |
| 17 | TCTRPNNNTRKSIPMGPGKAFYARGDITGDIKKAYCKINGTEWHSTLKLVVEKLREQY-N | 349 |
| 18 | TCTRPHNNTRKSIPMGPGKAFYARGDITGDIRKAYCEINGTEWHSTLKLVVEKLREQY-N | 348 |
| 19 | TCTRPNNNTRKSIPMGPGKAFYARGDITGDI-KAYCKINGTEWHSTLKLVVEKLREQY-N | 348 |
| 20 | TCTRPNNNTRKSIPMGPGKAFYARGDITGDIRKAYCEINGTEWHSTLKLVVEKLREQY-N | 349 |
| 21 | TCTRPNNNTRKSIPMGPGKAFYARGDITGDIKKAYCKINGTEWHSTLKLVVEKLREQY-N | 349 |
| 22 | TCTRPNNNTRKSIPMGPGKAFYARGDITGDIKKASCEINGTEWHSTLKLVVEKLREQY-N | 349 |
| 23 | TCTRPNNNTRKSIPMGPGKAFYARGDITGDIKKAYCKINGTEWHSTLKLVVEKLREQY-N | 349 |
| 24 | TCTRPNNNTRKSIPMGPGKAFYARGDITGDIKKAYCKINGTEWHSTLKLVVEKLREQY-N | 349 |
| 25 | TCTRPNNNTRKSIPMGPGKAFYARGDITGDIKKAYCKINGTEWHSTLKLVVEKLREQY-N | 348 |
| 26 | TCTRPNNNTRKSIPMGPGKAFYARGDITGDIKKAYCKINGTEWHSTLKLVVEKLREQY-N | 349 |
| 27 | TCTRPNNNTRKSIPMGPGKAFYARGDITGDIRKAYCEINGTEWHSTLKLVVEKLREQY-N | 349 |
| 28 | TCTRPNNNTRKSIPMGPGKAFYARGDITGDIKKAYCKINGTEWHSTLKLVVEKLREQY-N | 349 |
| 29 | TCTRPNNNTRKSIPMGPGKAFYARGDITGDIRKAYCEINGTEWHSTLKLVVEKLREQY-N | 349 |
| 30 | TCTRPHNNTRKSIPMGPGKAFYARGDITGDIRKAYCEINGTEWHSTLKLVVEKLREQY-N | 349 |
| 31 | TCTRPHNNTRKSIPMGPGKAFYARGDITGDIRKAYCEINGTEWHSTLKLVVEKLREQY-N | 349 |
| 32 | TCTRPNNNTRKSIPMGPGKAFYARGDITGDIKKAYCKINGTEWHSTLKLVVEKLREQY-N | 349 |
| 33 | TCTRPNNNTRKSIPMGPGKAFYARGDITGDIKKAYCKINGTEWHSTLKLVVEKLREQY-N | 349 |

| | Sequence | |
|---|---|---|
| | **KTIVFNRSSGGDPEIVMYSFNCGGEFFYCNSTKLFNSTWPWND-TKGSHDTNGTLILPCK** | |
| 1 | KTIVFNRSSGGDPEIVMYSFNCGGEFFYCNSTKLFNSTWPWND-TKGSHDTNGTLILPCK | 408 |
| 2 | KTIVFNRSSGGDPEIVMYSFNCGGEFFYCNSTKLFNSTWPWND-AKGSHDTNGTLILPCK | 408 |
| 3 | KTIVFNRSSGGDPEIVMYSFNCGGEFFYCNSTKLFNSTWPWND-TKGSHDTNGTLILPCK | 408 |
| 4 | KTIVFNRSSGGDPEIVMYSFNCGGEFFYCNSTKLFNSTWPWND-TKGSHDTNGTLRLPCK | 408 |
| 5 | KTIVFNRSSGGDPEIVMYSFNCGGEFFYCNSTKLFNSTWPWND-TKGSHDTNGTLILPCK | 408 |
| 6 | KTIVFNRSSGGDPEIVMYSFNCGGEFFYCNSTKLFNSTWPWND-TKGSHDTNGTLILPCK | 408 |
| 7 | KTIVFNRSSGGDPEIVMYSFNCGGEFFYCNSTKLFNSTWPWND-TKGSHDTNGTLMLPCK | 408 |
| 8 | KTIVFNRSSGGDPEIVMYSFNCGGEFFYCNSTKLFNSTWPWND-TKGSHDTNGTLILPCK | 408 |
| 9 | KTIVFNRSSGGDPEIVMYSFNCGGEFFYCNSTKLFNSTWPWND-TKGSHDTNGTLILPCK | 408 |
| 10 | KTIVFNRSSGGDPEIVMYSFNCGGEFFYCNSTKLFNSTWPWND-TKGSHDTNGTLILPCK | 408 |
| 11 | KTIVFNRSSGGDPEIVMYSFNCGGEFFYCNSTKLFNSTWPWND-TKGSHDTNGTLILPCK | 408 |
| 12 | KTIVFNRSSGGDPEIVMYSFNCGGEFFYCNSTKLFNSTWPWND-TKGSHDTNGTLILPCK | 408 |
| 13 | KTIVFNRSSGGDPEIVMYSFNCGGEFFYCNSTKLFNSTWPWND-TKGSHDTNGTLMLPCK | 408 |
| 14 | KTIVFNRSSGGDPEIVMYSFNCGGEFFYCNSTKLFNSTWPWND-TKGSHDTNGTLILPCK | 408 |
| 15 | KTIVFNRSSGGDPEIVMYSFNCGGEFFYCNSTKLFNSTWPWND-TKGSHDTNGTLILPCK | 408 |
| 16 | KTIVFNRSSGGDPEIVMYSFNCGGEFFYCNSTKLFNSTWPWND-TKGSHDTNDTLRLPCK | 408 |
| 17 | KTIVFNRSSGGDPEIVMYSFNCGGEFFYCNSTKLFNSTWPWND-TKGSHDTNGTLILPCK | 408 |
| 18 | KTIVFNRSSGGDPEIVMYSFNCGGEFFYCNSTKLFNSTWPWND-TKGSHDTNGTLILPCK | 407 |
| 19 | KTIVFNRSSGGDPEIVMYSFNCGGEFFYCNSTKLFNSTWPWND-AKGSHDTNGTLILPCK | 407 |
| 20 | KTIVFNRSSGGDPEIVMYSFNCGGEFFYCNSTKLFNSTWPWND-TKGSHDTNDTLMLPCK | 408 |
| 21 | KTIVFNRSSGGDPEIVMYSFNCGGEFFYCNSTKLFNSTWPWND-TKGSHDTNGTLILPCK | 408 |
| 22 | KTIVFNRSSGGDPEIVMYSFNCGGEFFYCNSTKLFNSTWPWND-TKGSHDTNGTLILPCK | 408 |
| 23 | KTIVFNRSSGGDPEIVMYSFNCGGEFFYCNSTKLFNSTWPWND-AKGSHDTNGTLILPCK | 408 |
| 24 | KTIVFNRSSGGDPEIVMYSFNCGGEFFYCNSTKLFNSTWPWND-TKGSHDTNGTLILPCK | 408 |
| 25 | KTIVFNRSSGGDPEIVMYSFNCGGEFFYCNSTKLFNSTWPWND-TKGSHDTNGTLILPCK | 407 |
| 26 | KTIVFNRSSGGDPEIVMYSFNCGGEFFYCNSTKLFNSTWPWND-TKGSHDTNGTLILPCK | 408 |
| 27 | KTIVFNRSSGGDPEIVMYSFNCGGEFFYCNSTKLFNSTWPWND-TKGSHDTNDTLMLPCK | 408 |
| 28 | KTIVFNRSSGGDPEIVMYSFNCGGEFFYCNSTKLFNSTWPWND-AKGSHDTNGTLILPCK | 408 |
| 29 | KTIVFNRSSGGDPEIVMYSFNCGGEFFYCNSTKLFNSTWPWND-TKGSHDTNDTLMLPCK | 408 |
| 30 | KTIVFNRSSGGDPEIVMYSFNCGGEFFYCNSTKLFNSTWPWND-TKGSHDTNGTLILPCK | 408 |
| 31 | KTIVFNRSSGGDPEIVMYSFNCGGEFFYCNSTKLFNSTWPWND-TKGSHDTNGTLILPCK | 408 |
| 32 | KTIVFNRSSGGDPEIVMYSFNCGGEFFYCNSTKLFNSTWPWND-TKGSHDTNGTLILPCK | 408 |
| 33 | KTIVFNRSSGGDPEIVMYSFNCGGEFFYCNSTKLFNSTWPWND-TKGSHDTNGTLILPCK | 408 |

| | IKQIINMWQGVGKAMYAPPIEGKIRCSSNITGLLLTRDGG-YESNETDEIFRPGGGDMRD | |
|---|---|---|
| 1 | IKQIINMWQGVGKAMYAPPIEGKIRCSSNITGLLLTRDGG-YESNETDEIFRPGGGDMRD | 467 |
| 2 | IKQIINMWQGVGKAMYAPPIEGKIRCSSNITGLLLTRDGG-YESNETDEIFRPGGGDMRD | 467 |
| 3 | IKQIINMWQGVGKAMYAPPIEGKIRCSSNITGLLLTRDGG-YESNETDEIFRPGGGDMRD | 467 |
| 4 | IKQIINMWQGVGKAMYAPPIEGKIRCSSNITGLLLTRDGG-YESNETDEIFRPGGGDMRD | 467 |
| 5 | IKQIINMWQGVGKAMYAPPIEGKIRCSSNITGLLLTRDGG-YESNETDEIFRPGGGDMRD | 467 |
| 6 | IKQIINMWQGVGKAMYAPPIEGKIRCSSNITGLLLTRDGG-YESNETDEIFRPGGGDMRD | 467 |
| 7 | IKQIINMWQGVGKAMYAPPIEGKIRCSSNITGLLLTRDGG-YESNETDEIFRPGGGDMRD | 467 |
| 8 | IKQIINMWQGVGKAMYAPPIEGKIRCSSNITGLLLTRDGG-YESNETDEIFRPGGGDMRD | 467 |
| 9 | IKQIINMWQGVGKAMYAPPIEGKIRCSSNITGLLLTRDGG-YESNETDEIFRPGGGDMRD | 467 |
| 10 | IKQIINMWQGVGKAMYAPPIEGKIRCSSNITGLLLTRDGG-YESNETDEIFRPGGGDMRD | 467 |
| 11 | IKQIINMWQGVGKAMYAPPIEGKIRCSSNITGLLLTRDGG-YESNETDEIFRPGGGDMRD | 467 |
| 12 | IKQIINMWQGVGKAMYAPPIEGKIRCSSNITGLLLTRDGG-YESNETDEIFRPGGGDMRD | 467 |
| 13 | IKQIINMWQGVGKAMYAPPIEGKIRCSSNITGLLLTRDGG-YESNETDEIFRPGGGDMRD | 467 |
| 14 | IKQIINMWQGVGKAMYAPPIEGKIRCSSNITGLLLTRDGG-YESNETDEIFRPGGGDMRD | 467 |
| 15 | IKQIINMWQGVGKAMYAPPIEGKIRCSSNITGLLLTRDGG-YESNETDEIFRPGGGDMRD | 467 |
| 16 | IKQIINMWQGVGKAMYAPPIEGKIRCSSNITGLLLTRDGG-YESNETDEIFRPGGGDMRD | 467 |
| 17 | IKQIINMWQGVGKAMYAPPIEGKIRCSSNITGLLLTRDGG-YESNETDEIFRPGGGDMRD | 467 |
| 18 | IKQIINMWQGVGKAMYAPPIEGKIRCSSNITGLLLARDGG-YESNETDEIFRPGGGDMRD | 466 |
| 19 | IKQIINMWQGVGKAMYAPPIEGKIRCSSNITGLLLTRDGG-YESNETDEIFRPGGGDMRD | 466 |
| 20 | IKQIINMWQGVGKAMYAPPIEGQIRCSSNITGLLLTRDGG-YESNETDEIFRPGGG-MRD | 466 |
| 21 | IKQIINMWQGVGKAMYAPPIEGKIRCSSNITGLLLTRDGG-YESNETDEIFRPGGGDMRD | 467 |
| 22 | IKQIINMWQGVGKAMYAPPIEGKIRCSSNITGLLLTRDGG-YESNETDEIFRPGGGDMRD | 467 |
| 23 | IKQIINMWQGVGKAMYAPPIEGKIRCSSNITGLLLTRDGG-YESNETDEIFRPGGGDMRD | 467 |
| 24 | IKQIINMWQGVGKAMYAPPIEGKIRCSSNITGLLLTRDGG-YESNETDEIFRPGGGDMRD | 467 |
| 25 | IKQIINMWQGVGKAMYAPPIEGKIRCSSNITGLLLTRDGG-YENNETDEIFRPGGGDMRD | 466 |
| 26 | IKQIINMWQGVGKAMYAPPIEGKIRCSSNITGLLLTRDGG-YESNETDEIFRPGGGDMRD | 467 |
| 27 | IKQIINMWQGVGKAMYAPPIEGKIRCSSNITGLLLTRDGG-YESNETDEIFRPGGGDMRD | 467 |
| 28 | IKQIINMWQGVGKAMYAPPIEGKIRCSSNITGLLLTRDGG-YESNETDEIFRPGGGDMRD | 467 |
| 29 | IKQIINMWQGVGKAMYAPPIEGQIRCSSNITGLLLTRDGG-YESNETDEIFRPGGGDMRD | 467 |
| 30 | IKQIINMWQGVGKAMYAPPIEGKIRCSSNITGLLLARDGG-YESNETDEIFRPGGGDMRD | 467 |
| 31 | IKQIINMWQGVGKAMYAPPIEEKIRCSSNITGLLLTRDGG-YESNETDEIFRPGGGDMRD | 467 |
| 32 | IKQIINMWQGVGKAMYAPPIEGKIRCSSNITGLLLTRDGG-YESNETDEIFRPGGGDMRD | 467 |
| 33 | IKQIINMWQGVGKAMYAPPIEGKIRCSSNITGLLLTRDGG-YESNETDEIFRPGGGDMRD | 467 |

| | Sequence | |
|---|---|---|
| | NWRSELYKYKVVKIEPLGVAPTKAKRRVVQREKRAFG-LGAVFLGFLGAAGSTMGAASIT | |
| 1 | NWRSELYKYKVVKIEPLGVAPTKAKRRVVQREKRAFG-LGAVFLGFLGAAGSTMGAASIT | 526 |
| 2 | NWRSELYKYKVVKIEPLGVAPTKAKRRVVQREKRAFG-LGAVFLGFLGAAGSTMGAASIT | 526 |
| 3 | NWRSELYKYKVVKIEPLGVAPTKAKRRVVQREKRAFG-LGAVFLGFLGAAGSTMGAASIT | 526 |
| 4 | NWRSELYKYKVVKIEPLGVAPTKAKRRVVQREKRAFG-LGAVFLGFLGAAGSTMGAASIT | 526 |
| 5 | NWRSELYKYKVVKIEPLGVAPTKAKRRVVQREKRAFG-LGAVFLGFLGAAGSTMGAASIT | 526 |
| 6 | NWRSELYKYKVVKIEPLGVAPTKAKRRVVQREKRAFG-LGAVFLGFLGAAGSTMGAASIT | 526 |
| 7 | NWRSELYKYKVVKIEPLGVAPTKAKRRVVQREKRAFG-LGAVFLGFLGAAGSTMGAASIT | 526 |
| 8 | NWRSELYKYKVVKIEPLGVAPTKAKRRVVQREKRAFG-LGAVFLGFLGAAGSTMGAASIT | 526 |
| 9 | NWRSELYKYKVVKIEPLGVAPTKAKRRVVQREKRAFG-LGAVFLGFLGAAGSTMGAASIT | 526 |
| 10 | NWRSELYKYKVVKIEPLGVAPTKAKRRVVQREKRAFG-LGAVFLGFLGAAGSTMGAASIT | 526 |
| 11 | NWRSELYKYKVVKIEPLGVAPTKAKRRVVQREKRAFG-LGAVFLGFLGAAGSTMGAASIT | 526 |
| 12 | NWRSELYKYKVVKIEPLGVAPTKAKRRVVQREKRAFG-LGAVFLGFLGAAGSTMGAASIT | 526 |
| 13 | NWRSELYKYKVVKIEPLGVAPTKAKRRVVQREKRAFG-LGAVFLGFLGAAGSTMGAASIT | 526 |
| 14 | NWRSELYKYKVVKIEPLGVAPTKAKRRVVQREKRAFG-LGAVFLGFLGAAGSTMGAASIT | 526 |
| 15 | NWRSELYKYKVVKIEPLGVAPTKAKRRVVQREKRAFG-LGAVFLGFLGAAGSTMGAASIT | 526 |
| 16 | NWRSELYKYKVVKIEPLGVAPTKAKRRVVQREKRAFG-LGAVFLGFLGAAGSTMGAASIT | 526 |
| 17 | NWRSELYKYKVVKIEPLGVAPTKAKRRVVQREKRAFG-LGAVFLGFLGAAGSTMGAASIT | 526 |
| 18 | NWRSELYKYKVVKIEPLGVAPTKAKRRVVQREK-AFG-LGAVFLGFLGAAGSTMGAASIT | 524 |
| 19 | NWRSELYKYKVVKIEPLGVAPTKAKRRVVQREKRAFG-LGAVFLGFLGAAGSTMGAASIT | 525 |
| 20 | NWRSELYKYKVVKIEPLGVAPTKAKRRVVQREKRAFG-LGAVFLGFLGAAGSTMGAASIT | 525 |
| 21 | NWRSELYKYKVVKIEPLGVAPTKAKRRVVQREKRAFG-LGAVFLGFLGAAGSTMGAASIT | 526 |
| 22 | NWRSELYKYKVVKIEPLGVAPTKAKRRVVQREKRAFG-LGAVFLGFLGAAGSTMGAASIT | 526 |
| 23 | NWRSELYKYKVVKIEPLGVAPTKAKRRVVQREKRAFG-LGAVFLGFLGAAGSTMGAASIT | 526 |
| 24 | NWRSELYKYKVVKIEPLGVAPTKAKRRVVQREKRAFG-LGAVFLGFLGAAGSTMGAASIT | 526 |
| 25 | NWRSELYKYKVVKIEPLGVAPTKAKRRVVQREKRAFG-LGAVFLGFLGAAGSTMGAASIT | 525 |
| 26 | NWRSELYKYKVVKIEPLGVAPTRAKRRVVQREKRAFG-LGAVFLGFLGAAGSTMGAASIT | 526 |
| 27 | NWRSELYKYKVVKIEPLGVAPTKAKRRVVQREKRAFG-LGAVFLGFLGAAGSTMGAASIT | 526 |
| 28 | NWRSELYKYKVVKIEPLGVAPTKAKRRVVQREKRAFG-LGAVFLGFLGAAGSTMGAASIT | 526 |
| 29 | NWRSELYKYKVVKIEPLGVAPTKAKRRVVQREKRAFG-LGAVFLGFLGAAGSTMGAASIT | 526 |
| 30 | NWRSELYKYKVVKIEPLGVAPTKAKRRVVQREKRAFG-LGAVFLGFLGAAGSTMGAASIT | 526 |
| 31 | NWRSELYKYKVVKIEPLGVAPTKAKRRVVQREKRAFG-LGAVFLGFLGAAGSTMGAASIT | 526 |
| 32 | NWRSELYKYKVVKIEPLGVAPTKAKRRVVQREKRAFG-LGAVFLGFLGAAGSTMGAASIT | 526 |
| 33 | NWRSELYKYKVVKIEPLGVAPTKAKRRVVQREKRAFG-LGAVFLGFLGAAGSTMGAASIT | 526 |

| | | |
|---|---|---|
| | **LTVQARQLLSGIVQQQNNLLRAIEAQQHLLQLTVWGIKQLQARVLAVERYLKDQQLLGIW** | |
| 1 | LTVQARQLLSGIVQQQNNLLRAIEAQQHLLQLTVWGIKQLQARVLAVERYLKDQQLLGIW | 586 |
| 2 | LTVQARQLLSGIVQQQNNLLRAIEAQQHLLQLTVWGIKQLQARVLAVERYLKDQQLLGIW | 586 |
| 3 | LTVQARQLLSGIVQQQNNLLRAIEAQQHLLQLTVWGIKQLQARVLAVERYLKDQQLLGIW | 586 |
| 4 | LTVQARQLLSGIVQQQNNLLRAIEAQQHLLQLTVWGIKQLQARVLAVERYLKDQQLLGIW | 586 |
| 5 | LTVQARQLLSGIVQQQNNLLRAIEAQQHLLQLTVWGIKQLQARVLAVERYLKDQQLLGIW | 586 |
| 6 | LTVQARQLLSGIVQQQNNLLRAIEAQQHLLQLTVWGIKQLQARVLAVERYLKDQQLLGIW | 586 |
| 7 | LTVQARQLLSGIVQQQNNLLRAIEAQQHLLQLTVWGIKQLQARVLAVERYLKDQQLLGIW | 586 |
| 8 | LTVQARQLLSGIVQQQNNLLRAIEAQQHLLQLTVWGIKQLQARVLAVERYLKDQQLLGIW | 586 |
| 9 | LTVQARQLLSGIVQQQNNLLRAIEAQQHLLQLTVWGIKQLQARVLAVERYLKDQQLLGIW | 586 |
| 10 | LTVQARQLLSGIVQQQNNLLRAIEAQQHLLQLTVWGIKQLQARVLAVERYLKDQQLLGIW | 586 |
| 11 | LTVQARQLLSGIVQQQNNLLRAIEAQQHLLQLTVWGIKQLQARVLAVERYLKDQQLLGIW | 586 |
| 12 | LTVQARQLLSGIVQQQNNLLRAIEAQQHLLQLTVWGIKQLQARVLAVERYLKDQQLLGIW | 586 |
| 13 | LTVQARQLLSGIVQQQNNLLRAIEAQQHLLQLTVWGIKQLQARVLAVERYLKDQQLLGIW | 586 |
| 14 | LTVQARQLLSGIVQQQNNLLRAIEAQQHLLQLTVWGIKQLQARVLAVERYLKDQQLLGIW | 586 |
| 15 | LTVQARQLLSGIVQQQNNLLRAIEAQQHLLQLTVWGIKQLQARVLAVERYLKDQQLLGIW | 586 |
| 16 | LTVQARQLLSGIVQQQNNLLRAIEAQQHLLQLTVWGIKQLQARVLAVERYLKDQQLLGIW | 586 |
| 17 | LTVQARQLLSGIVQQQNNLLRAIEAQQHLLQLTVWGIKQLQARVLAVERYLKDQQLLGIW | 586 |
| 18 | LTVQARQLLSGIVQQQNNLLRAIEAQQHLLQLTVWGIKQLQARVLAVERYLKDQQLLGIW | 584 |
| 19 | LTVQARQLLSGIVQQQNNLLRAIEAQQHLLQLTVWGIKQLQARVLAVERYLKDQQLLGIW | 585 |
| 20 | LTVQARQLLSGIVQQQNNLLRAIEAQQHLLQLTVWGIKQLQARVLAVERYLKDQQLLGIW | 585 |
| 21 | LTVQARQLLSGIVQQQNNLLRAIEAQQHLLQLTVWGIKQLQARVLAVERYLKDQQLLGIW | 586 |
| 22 | LTVQARQLLSGIVQQQNNLLRAIEAQQHLLQLTVWGIKQLQARVLAVERYLKDQQLLGIW | 586 |
| 23 | LTVQARQLLSGIVQQQNNLLRAIEAQQHLLQLTVWGIKQLQARVLAVERYLKDQQLLGIW | 586 |
| 24 | LTVQARQLLSGIVQQQNNLLRAIEAQQHLLQLTVWGIKQLQARVLAVERYLKDQQLLGIW | 586 |
| 25 | LTVQARQLLSGIVQQQNNLLRAIEAQQHLLQLTVWGIKQLQARVLAVERYLKDQQLLGIW | 585 |
| 26 | LTVQARQLLSGIVQQQNNLLRAIEAQQHLLQLTVWGIKQLQARVLAVERYLKDQQLLGIW | 586 |
| 27 | LTVQARQLLSGIVQQQNNLLRAIEAQQHLLQLTVWGIKQLQARVLAVERYLKDQQLLGIW | 586 |
| 28 | LTVQARQLLSGIVQQQNNLLRAIEAQQHLLQLTVWGIKQLQARVLAVERYLKDQQLLGIW | 586 |
| 29 | LTVQARQLLSGIVQQQNNLLRAIETQQHLLQLTVWGIKQLQARVLAVERYLKDQQLLGIW | 586 |
| 30 | LTVQARQLLSGIVQQQNNLLRAIEAQQHLLQLTVWGIKQLQARVLAVERYLKDQQLLGIW | 586 |
| 31 | LTVQARQLLSGIVQQQNNLLRAIEAQQHLLQLTVWGIKQLQARVLAVERYLKDQQLLGIW | 586 |
| 32 | LTVQARQLLSGIVQQQNNLLRAIEAQQHLLQLTVWGIKQLQARVLAVERYLKDQQLLGIW | 586 |
| 33 | LTVQARQLLSGIVQQQNNLLRAIEAQQHLLQLTVWGIKQLQARVLAVERYLKDQQLLGIW | 586 |

**GCSGKLICTTTVPWNTSWSNKSLEQIWDNMTWMEWEREIDNYTGYIYQLIEESQNQQEKN**

| | | |
|---|---|---|
| 1 | GCSGKLICTTTVPWNTSWSNKSLEQIWDNMTWMEWEREIDNYTGYIYQLIEESQNQQEKN | 646 |
| 2 | GCSGKLICTTTVPWNTSWSNKSLEQIWDNMTWMEWEREIDNYTGYIYQLIEESQNQQEKN | 646 |
| 3 | GCSGKLICTTTVPWNTSWSNKSLEQIWDNMTWMEWEREIDNYTGYIYQLIEESQNQQEKN | 646 |
| 4 | GCSGKLICTTTVPWNTSWSNKSLEQIWDNMTWMEWEREIDNYTGYIYQLIEESQNQQEKN | 646 |
| 5 | GCSGKLICTTTVPWNTSWSNKSLEQIWDNMTWMEWEREIDNYTGYIYQLIEESQNQQEKN | 646 |
| 6 | GCSGKLICTTTVPWNTSWSNKSLEQIWDNMTWMEWEREIDNYTGYIYQLIEESQNQQEKN | 646 |
| 7 | GCSGKLICTTTVPWNTSWSNKSLEQIWDNMTWMEWEREIDNYTGYIYQLIEESQNQQEKN | 646 |
| 8 | GCSGKLICTTTVPWNTSWSNKSLEQIWDNMTWMEWEREIDNYTGYIYQLIEESQNQQEKN | 646 |
| 9 | GCSGKLICTTTVPWNTSWSNKSLEQIWDNMTWMEWEREIDNYTGYIYQLIEESQNQQEKN | 646 |
| 10 | GCSGKLICTTTVPWNTSWSNKSLEQIWDNMTWMEWEREIDNYTGYIYQLIEESQNQQEKN | 646 |
| 11 | GCSGKLICTTTVPWNTSWSNKSLEQIWDNMTWMEWEREIDNYTGYIYQLIEESQNQQEKN | 646 |
| 12 | GCSGKLICTTTVPWNTSWSNKSLEQIWDNMTWMEWEREIDNYTGYIYQLIEESQNQQEKN | 646 |
| 13 | GCSGKLICTTTVPWNTSWSNKSLEQIWDNMTWMEWEREIDNYTGYIYQLIEESQNQQEKN | 646 |
| 14 | GCSGKLICTTTVPWNTSWSNKSLEQIWDNMTWMEWEREIDNYTGYIYQLIEESQNQQEKN | 646 |
| 15 | GCSGKLICTTTVPWNTSWSNKSLEQIWDNMTWMEWEREIDNYTGYIYQLIEESQNQQEKN | 646 |
| 16 | GCSGKLICTTTVPWNTSWSNKSLEQIWDNMTWMEWEREIDNYTGYIYQLIEESQNQQEKN | 646 |
| 17 | GCSGKLICTTTVPWNTSWSNKSLEQIWDNMTWMEWEREIDNYTGYIYQLIEESQNQQEKN | 646 |
| 18 | GCSGKLICTTTVPWNTSWSNKSLEQIWDNMTWMEWEREIDNYTGYIYQLIEESQNQQEKN | 644 |
| 19 | GCSGKLICTTTVPWNTSWSNKSLEQIWDNMTWMEWEREIDNYTGYIYQLIEESQNQQEKN | 645 |
| 20 | GCSGKLICTTTVPWNTSWSNKSLEQIWDNMTWMEWEREIDNYTGYIYQLIEESQNQQEKN | 645 |
| 21 | GCSGKLICTTTVPWNTSWSNKSLEQIWDNMTWMEWEREIDNYTGYIYQLIEESQNQQEKN | 646 |
| 22 | GCSGKLICTTTVPWNTSWSNKSLEQIWDNMTWMEWEREIDNYTGYIYQLIEESQNQQEKN | 646 |
| 23 | GCSGKLICTTTVPWNTSWSNKSLEQIWDNMTWMEWEREIDNYTGYIYQLIEESQNQQEKN | 646 |
| 24 | GCSGKLICTTTVPWNTSWSNKSLEQIWDNMTWMEWEREIDNYTGYIYQLIEESQNQQEKN | 646 |
| 25 | GCSGKLICTTTVPWNTSWSNKSLEQIWDNMTWMEWEREIDNYTGYIYQLIEESQNQQEKN | 645 |
| 26 | GCSGKLICTTTVPWNTSWSNKSLEQIWDNMTWMEWEREIDNYTGYIYQLIEESQNQQEKN | 646 |
| 27 | GCSGKLICTTTVPWNTSWSNKSLEQIWDNMTWMEWEREIDNYTGYIYQLIEESQNQQEKN | 646 |
| 28 | GCSGKLICTTTVPWNTSWSNKSLEQIWDNMTWMEWEREIDNYTGYIYQLIEESQNQQEKN | 646 |
| 29 | GCSGKLICTTTVPWNTSWSNKSLEQIWDNMTWMEWEREIDNYTGYIYQLIEESQNQQEKN | 646 |
| 30 | GCSGKLICTTTVPWNTSWSNKSLEQIWDNMTWMEWEREIDNYTGYIYQLIEESQNQQEKN | 646 |
| 31 | GCSGKLICTTTVPWNTSWSNKSLEQIWDNMTWMEWEREIDNYTGYIYQLIEESQNQQEKN | 646 |
| 32 | GCSGKLICTTTVPWNTSWSNKSLEQIWDNMTWMEWEREIDNYTGYIYQLIEESQNQQEKN | 646 |
| 33 | GCSGKLICTTTVPWNTSWSNKSLEQIWDNMTWMEWEREIDNYTGYIYQLIEESQNQQEKN | 646 |

| | | |
|---|---|---|
| | **EQELLALDKWASLWNWFDITNWLWYIKIFIMIVGGLIGLRIVFTVLSIVNRVRQGYSPLS** | |
| 1 | EQELLALDKWASLWNWFDITNWLWYIKIFIMIVGGLIGLRIVFTVLSIVNRVRQGYSPLS | 706 |
| 2 | EQELLALDKWASLWNWFDITNWLWYIKIFIMIVGGLISLRIVFTVLSIVNRVRQGYSPLS | 706 |
| 3 | EQELLALDKWASLWNWFDITNWLWYIKIFIMIVGGLIGLRIVFTVLSIVNRVRQGYSPLS | 706 |
| 4 | EQELLALDKWASLWNWFDITNWLWYIKIFIMIVGGLIGLRIVFTVLSIVNRVRQGYSPLS | 706 |
| 5 | EQELLALDKWASLWNWFDITNWLWYIKIFIMIVGGLIGLRIVFTVLSIVNRVRQGYSPLS | 706 |
| 6 | EQELLALDKWASLWNWFDITNWLWYIKIFIMIVGGLIGLRIVFTVLSIVNRVRQGYSPLS | 706 |
| 7 | EQELLALDKWASLWNWFDITNWLWYIKIFIMIVGGLIGLRIVFTVLSIVNRVRQGYSPLS | 706 |
| 8 | EQELLAWDKWASLWNWFDITNWLWYIKIFIMIVGGLIGLRIVFTVLSIVNRVRQGYSPLS | 706 |
| 9 | EQELLALDKWASLWNWFDITNWLWYIKIFIMIVGGLIGLRIVFTVLSIVNRVRQGYSPLS | 706 |
| 10 | EQELLALDKWASLWNWFDITNWLWYIKIFIMIVGGLIGLRIVFTVLSIVNRVRQGYSPLS | 706 |
| 11 | EQELLALDKWASLWNWFDITNWLWYIKIFIMIVGGLIGLRIVFTVLSIVNRVRQGYSPLS | 706 |
| 12 | EQELLALDKWASLWNWFDITNWLWYIKIFIMIVGGLIGLRIVFTVLSIVNRVRQGYSPLS | 706 |
| 13 | EQELLALDKWASLWNWFDITNWLWYIKIFIMIVGGLIGLRIVFTVLSIVNRVRQGYSPLS | 706 |
| 14 | EQELLALDKWASLWNWFDITNWLWYIKIFIMIVGGLIGLRIVFTVLSIVNRVRQGYSPLS | 706 |
| 15 | EQELLALDKWASLWNWFDITNWLWYIKIFIMIVGGLIGLRIVFTVLSIVNRVRQGYSPLS | 706 |
| 16 | EQELLALDKWASLWNWFDITNWLWYIKIFIMIVGGLIGLRIVFTVLSIVNRVRQGYSPLS | 706 |
| 17 | EQELLALDKWAGLWNWFDITNWLWYIKIFIMIVGGLIGLRIVFTVLSIVNRVRQGYSPLS | 706 |
| 18 | EQELLALDKWASLWNWFDITNWLWYIKIFIMIVGGLIGLRIVFTVLSIVNRVRQGYSPLS | 704 |
| 19 | EQELLALDKWASLWNWFDITNWLWYIKIFIMIVGGLIGLRIVFTVLSIVNRVRQGYSPLS | 705 |
| 20 | EQELLALDKWASLWNWFDITNWLWYIKIFIMIVGGLIGLRIVFTVLSIVNRVRQGYSPLS | 705 |
| 21 | EQELLALDKWASLWNWFDITNWLWYIKIFIMIVGGLIGLRIVFTVLSIVNRVRQGYSPLS | 706 |
| 22 | EQELLALDKWASLWNWFDITNWLWYIKIFIMIVGGLIGLRIVFTVLSIVNRVRQGYSPLS | 706 |
| 23 | EQELLALDKWASLWNWFDITNWLWYIKIFIMIVGGLIGLRIVFTVLSIVNRVRQGYSPLS | 706 |
| 24 | EQELLAWDKWASLWNWFDITNWLWYIKIFIMIVGGLIGLRIVFTVLSIVNRVGQGYSPLS | 706 |
| 25 | EQELLAWDKWASLWNWFDITNWLWYIKIFIMIVGGLIGLRIVFTVLSIVNRVRQGYSPLS | 705 |
| 26 | EQELLALDKWASLWNWFDITNWLWYIKIFIMIVGGLIGLRIVFTVLSIVNRVRQGYSPLS | 706 |
| 27 | EQELLALDKWASLWNWFDITNWLWYIKIFIMIVGGLIGLRIVFTVLSIVNRVRQGYSPLS | 706 |
| 28 | EQELLALDKWASLWNWFDITHWLWYIKIFIMIVGGLIGLRIVFTVLSIVNRVRQGYSPLS | 706 |
| 29 | EQELLALDKWASLWNWFDITNWLWYIKIFIMIVGGLIGLRIVFTVLSIVNRVRQGYSPLS | 706 |
| 30 | EQELLALDKWASLWNWFDITNWLWYIKIFIMIVGGLIGLRIVFTVLSIVNRVRQGYSPLS | 706 |
| 31 | EQELLALDKWASLWNWFDITNWLWYIKIFIMIVGGLIGLRIVFTVLSIVNRVRQGYSPLS | 706 |
| 32 | EQELLALDKWASLWNWFDITNWLWYIKIFIMIVGGLIGLRIVFTVLSIVNRVRQGYSPLS | 706 |
| 33 | EQELLALDKWASLWNWFDITNWLWYIKIFIMIVGGLIGLRIVFTVLSIVNRVRQGYSPLS | 706 |

| | | |
|---|---|---|
| | **FQTHLPAQRGPDRPEGIGEEGGERDRDRSDPLVNGFLALIWSDLRSLCLFSYHRLRDLLL** | |
| 1 | FQTHLPAQRGPDRPEGIGEEGGERDRDRSDPLVNGFLALIWSDLRSLCLFSYHRLRDLLL | 766 |
| 2 | FQTHLPAQRGPDRPEGIGEEGGERDRDRSDPLVNGFLALIWSDLRSLCLFSYHRLRDLLL | 766 |
| 3 | FQTHLPAQRGPDRPEGIGEEGGERDRDRSDPLVNGFLALIWSDLRSLCLFSYHRLRDLLL | 766 |
| 4 | FQTHLPAQRGPDRPEGIGEEGGERDRDRSDPLVNGFLALIWSDLRSLCLFSYHRLRDLLL | 766 |
| 5 | FQTHLPAQRGPDRPEGIGEEGGERDRDRSDPLVNGFLALIWSDLRSLCLFSYHRLRDLLL | 766 |
| 6 | FQTHLPAQRGPDRPEGIGEEGGERDRDRSDPLVNGFLALIWSDLRSLCLFSYHRLRDLLL | 766 |
| 7 | FQTHLPAQRGPDRPEGIGEEGGERDRDRSDPLVNGFLTLIWSDLRSLCLFSYHRLRDLLL | 766 |
| 8 | FQTHLPAQRGPDRPEGIGEEGGERDRDRSDPLVNGFLALIWSDLRSLCLFSYHRLRDLLL | 766 |
| 9 | FQTHLPAQRGPDRPEGIGEEGGERDRDRSDPLVNGFLALIWSDLRSLCLFSYHRLRDLLL | 766 |
| 10 | FQTHLPAQRGPDRPEGIGEEGGERDRDRSDPLVNGFLALIWSDLRSLCLFSYHRLRDLLL | 766 |
| 11 | FQTHLPAQRGPDRPEGIGEEGGERDRDRSDPLVNGFLALIWSDLRSLCLFSYHRLRDLLL | 766 |
| 12 | FQTHLPAQRGPDRPEGIGEEGGERDRDRSDPLVNGFLALIWSDLRSLCLFSYHRLRDLLL | 766 |
| 13 | FQTHLPAQRGPDRPEGIGEEGGERDRDRSDPLVNGFLALIWSDLRSLCLFSYHRLRDLLL | 766 |
| 14 | FQTHLPAQRGPDRPEGIGEEGGERDRDRSDPLVNGFLALIWSDLRSLCLFSYHRLRDLLL | 766 |
| 15 | FQTHLPAQRGPDRPEGIGEEGGERDRDRSDPLVNGFLALIWSDLRSLCLFSYHRLRDLLL | 766 |
| 16 | FQTHLPAQRGPDRPEGIGEEGGERDRDRSDPLVNGFLALIWSDLRSLCLFSYHRLRDLLL | 766 |
| 17 | FQTHLPAQRGPDRPEGIGEEGGERDRDRSDPLVNGFLALIWSDLRSLCLFSYHRLRDLLL | 766 |
| 18 | FQTHLPAQRGPDRPEGIGEEGGERDRDRSDPLVNGFLALIWSDLRSLCLFSYHRLRDLLL | 764 |
| 19 | FQTHLPAQRGPDRPEGIGEEGGERDRDRSDPLVNGFLALIWSDLRSLCLFSYHRLRDLLL | 765 |
| 20 | FQTHLPAQRGPDRPEGIGEEGGERDRDRSDPLVNGFLTLIWSDLRSLCLFSYHRLRDLLL | 765 |
| 21 | FQTHLPAQRGPDRPEGIGEEGGKRDRDRSDPLVNGFLALIWSDLRSLCLFSYHRLRDLLL | 766 |
| 22 | FQTHLPAQRGPDRPEGIGEEGGERDRDRSDPLVNGFLALIWSDLRSLCLFSYHRLRDLLL | 766 |
| 23 | FQTHLPAQRGPDRPEGIGEEGGERDRDRSDPLVNGFLALIWSDLRSLCLFSYHRLRDLLL | 766 |
| 24 | FQTHLPAQRGPDRPEGIGEEGGERDRDRSDPLVNGFLALIWSDLRSLCLFSYHRLRDLLL | 766 |
| 25 | FQTHLPAQRGPDRPEGIGEEGGERDRDRSDPLVNGFLALIWSDLRSLCLFSYHRLRDLLL | 765 |
| 26 | FQTHLPAQRGPDRPEGIGEEGGERDRDRSDPLVNGFLALIWSDLRSLCLFSYHRLRDLLL | 766 |
| 27 | FQTHLPAQRGPDRPEGIGEEGGERDRDRSDPLVNGFLTLIWSDLRSLCLFSYHRLRDLLL | 766 |
| 28 | FQTHLPAQRGPDRPEGIGEEGGERDRDRSDPLVNGFLALIWSDLRSLCLFSYHRLRDLLL | 766 |
| 29 | FQTHLPAQRGPDRPEGIGEEGGERDRDRSDPLVNGFLTLIWSDLRSLCLFSYHRLRDLLL | 766 |
| 30 | FQTHLPAQRGPDRPEGIGEEGGERDRDRSDPLVNGFLALIWSDLRSLCLFSYHRLRDLLL | 766 |
| 31 | FQTHLPAQRGPDRPEGIGEEGGERDRDRSDPLVNGFLTLIWSDLRSLCLFSYHRLRDLLL | 766 |
| 32 | FQTHLPAQRGPDRPEGIGEEGGERDRDRSDPLVNGFLALIWSDLRSLCLFSYHRLRDLLL | 766 |
| 33 | FQTHLPAQRGPDRPEGIGEEGGERDRDRSDPLVNGFLALIWSDLRSLCLFSYHRLRDLLL | 766 |

| | | |
|---|---|---|
|  | IVTRIVELLGR-------RGWEVLKYWWNLLQYWSQELKNSAVSLLNATAIAVAEGTDRV |  |
| 1 | IVTRIVELLGR-------RGWEVLKYWWNLLQYWSQELKNSAVSLLNATAIAVAEGTDRV | 819 |
| 2 | IVTRIVELLGR-------RGWEVLKYWWNLLQYWSQELKNSAVSLLNATAIAVAEGTDRV | 819 |
| 3 | IVTRIVELLGR-------RGWEVLKYWWNLLQYWSQELKNSAVSLLNATAIAVAEGTDRV | 819 |
| 4 | IVTRIVELLGR-------RGWEVLKYWWNLLQYWSQELKNSAVSLLNATAIAVAEGTDRV | 819 |
| 5 | IVTRIVELLGR-------RGWEVLKYWWNLLQYWSQELKNSAVSLLNATAIAVAEGTDRV | 819 |
| 6 | IVTRIVELLGR-------RGWEVLKYWWNLLQYWSQELKNSAVSLLNATAIAVAEGTDRV | 819 |
| 7 | IVTRIVELLGR-------RGWEVLKYWWNLLQYWSQELKNSAVSLLNATAIAVAEGTDRV | 819 |
| 8 | IVTRIVELLGR-------RGWEVLKYWWNLLQYWSQELKNSAVSLLNATAIAVAEGTDRV | 819 |
| 9 | IVTRIVELLGR-------RGWEVLKYWWNLLQYWSQELKNSAVSLLNATAIAVAEGTDRV | 819 |
| 10 | IVTRIVELLGR-------RGWEVLKYWWNLLQYWSQELKNSAVSLLNATAIAVAEGTDRV | 819 |
| 11 | IVTRIVELLGR-------RGWEVLKYWWNLLQYWSQELKNSAVSLLNATAIAVAEGTDRV | 819 |
| 12 | IVTRIVELLGR-------RGWEVLKYWWNLLQYWSQELKNSAVSLLNATAIAVAEGTDRV | 819 |
| 13 | IVTRIVELLGR-------RGWEVLKYWWNLLQYWSQELKNSAVSLLNATAIAVAEGTDRV | 819 |
| 14 | IVTRIVELLGR-------RGWEVLKYWWNLLQYWSQELKNSAVSLLNATAIAVAEGTDRV | 819 |
| 15 | IVTRIVELLGR-------RGWEVLKYWWNLLQYWSQELKNSAVSLLNATAIAVAEGTDRV | 819 |
| 16 | IVTRIVELLGR-------RGWEVLKYWWNLLQYWSQELKNSAVSLLNATAIAVAEGTDRV | 819 |
| 17 | IVTRIVELLGR-------RGWEVLKYWWNLLQYWSQELKNSAVSLLNATAIAVAEGTDRV | 819 |
| 18 | IVTRIVELLGR-------RGWEVLKYWWNLLQYWSQELKNSAVSLLNATAIAVAEGTDRV | 817 |
| 19 | IVTRIVELLGR-------RGWEVLKYWWNLLQYWSQELKNSAVSLLNATAIAVAEGTDRV | 818 |
| 20 | IVTRIVELLGR-------RGWEVLKYWWNLLQYWSQELKNSAVSLLNATAIAVAEGTDRV | 818 |
| 21 | IVTRIVELLGR-------RGWEVLKYWWNLLQYWSQELKNSAVSLLNATAIAVAEGTDRV | 819 |
| 22 | IVTRIVELLGR-------RGWEVLKYWWNLLQYWSQELKNSAVSLLNATAIAVAEGTDRV | 819 |
| 23 | IVTRIVELLGR-------RGWEVLKYWWNLLQYWSQELKNSAVSLLNATAIAVAEGTDRV | 819 |
| 24 | IVTRIVELLGR-------RGWEVLKYWWNLLQYWSQELKNSAVSLLNATAIAVAEGTDRV | 819 |
| 25 | IVTRIVELLGR-------RGWEVLKYWWNLLQYWSQELKNSAVSLLNATAIAVAEGTDRV | 818 |
| 26 | IVTRIVELLGR-------RGWEVLKYWWNLLQYWSQELKNSAVSLLNATAIAVAEGTDRV | 819 |
| 27 | IVTRIVELLGR-------RGWEVLKYWWNLLQYWSQELKNSAVNLLNATAIAVAEGTDRV | 819 |
| 28 | IVTRIVELLGR-------RGWEVLKYWWNLLQYWSQELKNSAVSLLNATAIAVAEGTDRV | 819 |
| 29 | IVTRIVELLGR-------RGWEVLKYWWNLLQYWSQELKNSAVSLLNATAIAVAEGTDRV | 819 |
| 30 | IVTRIVELLGR-------RGWEVLKYWWNLLQYWSQELKNSAVSLLNATAIAVAEGTDRV | 819 |
| 31 | IVTRIVELLGR-------RGWEVLKYWWNLLQYWSQELKNSAVSLLNATAIAVAEGTDRV | 819 |
| 32 | IVTRIVELLGR-------RGWEVLKYWWNLLQYWSQELKNSAVSLLNATAIAVAEGTDRV | 819 |
| 33 | IVTRIVELLGR-------RGWEVLKYWWNLLQYWSQELKNSAVSLLNATAIAVAEGTDRV | 819 |

| | | |
|---|---|---|
| | IEVVQRACRAILHIPRRIRQGLERALL | |
| 1 | IEVVQRACRAILHIPRRIRQGLERALL | 846 |
| 2 | IEVVQRACRAILHIPRRIRQGLERALL | 846 |
| 3 | IEVVQRACRAILHIPRRIRQGLERALL | 846 |
| 4 | IEVVQRACRAILHIPRRIRQGLERALL | 846 |
| 5 | VEVVQRACRAILHIPRRIRQGLERALL | 846 |
| 6 | IEVVQRACRAILHIPRRIRQGLERALL | 846 |
| 7 | IEVVQRACRAILHIPRRIRQGLERALL | 846 |
| 8 | IEVVQRACRAILHIPRRIRQGLERALL | 846 |
| 9 | IEVVQRACSAILHIPRRIRQGLERALL | 846 |
| 10 | IEVVQRACRAILHIPRRIRQGLERALL | 846 |
| 11 | IEVVQRACRAILHIPRRIRQGLERALL | 846 |
| 12 | IEVVQRACRAILHIPRRIRQGLERALL | 846 |
| 13 | IEVVQRACRAILHIPRRIRQGLERALL | 846 |
| 14 | IEVVQRACRAILHIPRRIRQGLERALL | 846 |
| 15 | IEVVQRACRAILHIPRRIRQGLERALL | 846 |
| 16 | IEVVQRACRAILHIPRRIRQGLERALL | 846 |
| 17 | IEVVQRACRAILHIPRRIRQGLERALL | 846 |
| 18 | IEVVQRACRAILHIPRRIRQGLERALL | 844 |
| 19 | IEVVQRACRAILHIPRRIRQGLERALL | 845 |
| 20 | IEVVQRACRATLHIPRRIRQGLERALL | 845 |
| 21 | IEVVQRACRAILHIPRRIRQGLERALL | 846 |
| 22 | IEVVQRACRAILHIPRRIRQGLERALL | 846 |
| 23 | IEVVQRACRAILHIPRRIRQGLERALL | 846 |
| 24 | IEVVQRACRAILHIPRRIRQGLERALL | 846 |
| 25 | IEVVQRACRAILHIPRRIRQGLERALL | 845 |
| 26 | IEVVQRACRAILHIPRRIRQGLERALL | 846 |
| 27 | IEVVQRACRATLHIPRRIRQGLERALL | 846 |
| 28 | IEVVQRACRAILHIPRRIRQGLERALL | 846 |
| 29 | IEVVQRACRATLHIPRRIRQGLERALL | 846 |
| 30 | IEVVQRACRAILHIPRRIRQGLERALL | 846 |
| 31 | IEVVQRACRAILHIPRRIRQGLERALL | 846 |
| 32 | IEVVQRACRAILHIPRRIRQGLERALL | 846 |
| 33 | IEVVQRACRAILHIPRRIRQGLERALL | 846 |

**Consensus**

1. B.US.2006.700010040_C9_4520.EU289193
2. B.US.2007.CH0040_3_d0299_ipe032_2_01.MG900232
3. B.US.2007.CH0040_3_d0299_ipe032_2_04.MG900233
4. B.US.2007.CH0040_3_d0299_ipe032_2_05.MG900234
5. B.US.2007.CH0040_3_d0299_ipe032_2_06.MG900235
6. B.US.2007.CH0040_3_d0299_ipe032_2_08.MG900236
7. B.US.2007.CH0040_3_d0299_ipe032_2_09.MG900237
8. B.US.2007.CH0040_3_d0299_ipe032_2_10.MG900238
9. B.US.2007.CH0040_3_d0299_ipe032_2_13.MG900239
10. B.US.2007.CH0040_3_d0299_ipe032_2_16.MG900240
11. B.US.2007.CH0040_3_d0299_ipe032_2_18.MG900241
12. B.US.2007.CH0040_3_d0299_ipe032_2_19.MG900242
13. B.US.2007.CH0040_3_d0299_ipe032_2_20.MG900243
14. B.US.2007.CH0040_3_d0299_ipe032_2_21.MG900244
15. B.US.2007.CH0040_3_d0299_ipe032_2_23.MG900245
16. B.US.2007.CH0040_3_d0299_ipe032_2_24.MG900246
17. B.US.2007.CH0040_3_d0299_ipe032_2_25.MG900247
18. B.US.2007.CH0040_3_d0299_ipe032_2_27.MG900248
19. B.US.2007.CH0040_3_d0299_ipe032_2_29.MG900249
20. B.US.2007.CH0040_3_d0299_ipe032_2_30.MG900250
21. B.US.2007.CH0040_3_d0299_ipe032_2_33.MG900251
22. B.US.2007.CH0040_3_d0299_ipe032_2_35.MG900252
23. B.US.2007.CH0040_3_d0299_ipe032_2_37.MG900253
24. B.US.2007.CH0040_3_d0299_ipe032_2_38.MG900254
25. B.US.2007.CH0040_3_d0299_ipe032_2_39.MG900255
26. B.US.2007.CH0040_3_d0299_ipe032_2_42.MG900256
27. B.US.2007.CH0040_3_d0299_ipe032_2_43.MG900257
28. B.US.2007.CH0040_3_d0299_ipe032_2_45.MG900258
29. B.US.2007.CH0040_3_d0299_ipe032_2_47.MG900259
30. B.US.2007.CH0040_3_d0299_ipe032_2_48.MG900260
31. B.US.2007.CH0040_3_d0299_ipe032_2_49.MG900261
32. B.US.2007.CH0040_3_d0299_ipe032_2_51.MG900262
33. B.US.2007.CH0040_3_d0299_ipe032_2_53.MG900263
34. B.US.2007.CH0040_3_d0299_ipe032_2_55.MG900264
35. B.US.2007.CH0040_3_d0299_ipe032_2_57.MG900265
36. B.US.2007.CH0040_3_d0299_ipe032_2_58.MG900266
37. B.US.2007.CH0040_3_d0299_ipe032_2_59.MG900267
38. B.US.2007.CH0040_3_d0299_ipe032_2_60.MG900268
39. B.US.2007.CH0040_3_d0299_ipe032_2_61.MG900269
40. B.US.2007.CH0040_3_d0299_ipe032_3_40.MG900270

| | | |
|---|---|---|
| | **MRVMGIRKNYQHLWREGILLLGILMICSAADNLWVTVYYGVPVWREATTTLFCASDAKAY** | |
| 1 | MRVMGIRKNYQHLWREGILLLGILMICSAADNLWVTVYYGVPVWREATTTLFCASDAKAY | 60 |
| 2 | MRVMGIRKNYQHLWREGILLLRILMICSAADNLWVTVYYGVPVWREATTTLFCASDAKAY | 60 |
| 3 | MRVMGIRKNYQHLWREGILLLGILMICSAADNLWVTVYYGVPVWREATTTLFCASDAKAY | 60 |
| 4 | MRVMGIRKNYQHLWREGILLLGILMICSAADNLWVTVYYGVPVWREATTTLFCASDAKAY | 60 |
| 5 | MRVMGIRKNYQHLWREGILLLGILMICSAADNLWVTVYYGVPVWREATTTLFCASDAKAY | 60 |
| 6 | MRVMGIRKNYQHLWREGILLLGILMICSAADNLWVTVYYGVPVWREATTTLFCASDAKAY | 60 |
| 7 | MRVMGIRKNYQHLWREGILLLGILMICSAADNLWVTVYYGVPVWREATTTLFCASDAKAY | 60 |
| 8 | MRVMGIRKNYQHLWREGILLLGILMICSAADNLWVTVYYGVPVWREATTTLFCASDAKAY | 60 |
| 9 | MRVMGIRKNYQHLWREGILLLGILMICSAADNLWVTVYYGVPVWREATTTLFCASDAKAY | 60 |
| 10 | MRVMGIRKNYQHLWREGILLLGILMICSAADNLWVTVYYGVPVWREATTTLFCASDAKAY | 60 |
| 11 | MRVMGIRKNYQHLWREGILLLGILMICSAADNLWVTVYYGVPVWREATTTLFCASDAKAY | 60 |
| 12 | MRVMGIRKNYQHLWREGILLLGILMICSAADNLWVTVYYGVPVWREATTTLFCASDAKAY | 60 |
| 13 | MRVMGIRKNYQHLWREGILLLGILMICSAADNLWVTVYYGVPVWREATTTLFCASDAKAY | 60 |
| 14 | MRVMGIRKNYQHLWREGILLLGILMICSAADNLWVTVYYGVPVWREATTTLFCASDAKAY | 60 |
| 15 | MRVMGIRKNYQHLWREGILLLGILMICSAADNLWVTVYYGVPVWREATTTLFCASDAKAY | 60 |
| 16 | MRVMGIRKNYQHLWREGILLLGILMICSAADNLWVTVYYGVPVWREATTTLFCASDAKAY | 60 |
| 17 | MRVMGIRKNYQHLWREGILLLGILMICSAADNLWVTVYYGVPVWREATTTLFCASDAKAY | 60 |
| 18 | MRVMGIRKNYQHLWREGILLLGILMICSAADNLWVTVYYGVPVWREATTTLFCASDAKAY | 60 |
| 19 | MRVMGIRKNYQHLWREGILLLGILMICSAADNLWVTVYYGVPVWREATTTLFCASDAKAY | 60 |
| 20 | MRVMGIRKNYQHLWREGILLLGILMICSAADNLWVTVYYGVPVWREATTTLFCASDAKAY | 60 |
| 21 | MRVMGIRKNYQHLWREGILLLGILMICSAADNLWVTVYYGVPVWREATTTLFCASDAKAY | 60 |
| 22 | MRVMGIRKNYQHLWREGILLLGILMICSAADNLWVTVYYGVPVWREATTTLFCASDAKAY | 60 |
| 23 | MRVMGIRKNYQHLWREGILLLGILMICSAADNLWVTVYYGVPVWREATTTLFCASDAKAY | 60 |
| 24 | MRVMGIRKNYQHLWREGILLLGILMICSAADNLWVTVYYGVPVWREATTTLFCASDAKAY | 60 |
| 25 | MRVMGIRKNYQHLWREGILLLGILMICSAADNLWVTVYYGVPVWREATTTLFCASDAKAY | 60 |
| 26 | MRVMGIRKNYQHLWREGILLLGILMICSAADNLWVTVYYGVPVWREATTTLFCASDAKAY | 60 |
| 27 | MRVMGIRKNYQHLWREGILLLGILMICSAADNLWVTVYYGVPVWREATTTLFCASDAKAY | 60 |
| 28 | MRVMGIRKNYQHLWREGILLLGILMICSAADNLWVTVYYGVPVWREATTTLFCASDAKAY | 60 |
| 29 | MRVMGIRKNYQHLWREGILLLGILMICSAADNLWVTVYYGVPVWREATTTLFCASDAKAY | 60 |
| 30 | MRVMGIRKNYQHLWREGILLLGILMICSAADNLWVTVYYGVPVWREATTTLFCASDAKAY | 60 |
| 31 | MRVMGIRKNYQHLWREGILLLGILMICSAADNLWVTVYYGVPVWREATTTLFCASDAKAY | 60 |
| 32 | MRVMGIRKNYQHLWREGILLLGILMICSAADNLWVTVYYGVPVWREATTTLFCASDAKAY | 60 |
| 33 | MRVMGIRKNYQHLWREGILLLGILMICSAADNLWVTVYYGVPVWREATTTLFCASDAKAY | 60 |
| 34 | MRVMGIRKNYQHLWREGILLLGILMICSAADNLWVTVYYGVPVWREATTTLFCASDAKAY | 60 |
| 35 | MRVMGIRKNYQHLWREGILLLGILMICSAADNLWVTVYYGVPVWREATTTLFCASDAKAY | 60 |
| 36 | MRVMGIRKNYQHLWREGILLLGILMICSAADNLWVTVYYGVPVWREATTTLFCASDAKAY | 60 |
| 37 | MRVMGIRKNYQHLWREGILLLGILMICSAADNLWVTVYYGVPVWREATTTLFCASDAKAY | 60 |
| 38 | MRVMGIRKNYQHLWREGILLLGILMICSAADNLWVTVYYGVPVWREATTTLFCASDAKAY | 60 |
| 39 | MRVMGIRKNYQHLWREGILLLGILMICSAADNLWVTVYYGVPVWREATTTLFCASDAKAY | 60 |
| 40 | MRVMGIRKNYQHLWREGILLLGILMICSAADNLWVTVYYGVPVWREATTTLFCASDAKAY | 60 |

| | Sequence | |
|---|---|---|
| | DTEAHNVWATHACVPTDPNPQEVELKNVTENFNMWENNMVEQMHEDIISLWDQSLKPCVK | |
| 1 | DTEAHNVWATHACVPTDPNPQEVELKNVTENFNMWENNMVEQMHEDIISLWDQSLKPCVK | 120 |
| 2 | DTEAHNV-ATHACVPTDPNPQEVELKNVTENFNMWENNMVEQMHEDIISLWDQSLKPCVK | 119 |
| 3 | DTEAHNVWATHACVPTDPNPQEVELKNVTENFNMWENNMVEQMHEDIISLWDQSLKPCVK | 120 |
| 4 | DTEAHNVWATHACVPTDPNPQEVELKNVTENFNMWENNMVEQMHEDIISLWDQSLKPCVK | 120 |
| 5 | DTEAHNVWATHACVPTDPNPQEVELKNVTENFNMWENNMVEQMHEDIISLWDQSLKPCVK | 120 |
| 6 | DTEAHNVWATHACVPTDPNPQEVELKNVTENFNMWENNMVEQMHEDIISLWDQSLKPCVK | 120 |
| 7 | DTEAHNVWATHACVPTDPNPQEVELKNVTENFNMWENNMVEQMHEDIISLWDQSLKPCVK | 120 |
| 8 | DTEAHNVWATHACVPTDPNPQEVELKNVTENFNMWENNMVEQMHEDIISLWDQSLKPCVK | 120 |
| 9 | DTEAHNVWATHACVPTDPNPQEVELKNVTENFNMWENNMVEQMHEDIISLWDQSLKPCVK | 120 |
| 10 | DTEAHNVWATHACVPTDPNPQEVELKNVTENFNMWENNMVEQMHEDIISLWDQSLKPCVK | 120 |
| 11 | DTEAHNVWATHACVPTDPNPQEVELKNVTENFNMWENNMVEQMHEDIISLWDQSLKPCVK | 120 |
| 12 | DTEAHNVWATHACVPTDPNPREVELKNVTENFNMWENNMVEQMHEDIISLWDQSLKPCVK | 120 |
| 13 | DTEAHNVWATHACVPTDPNPQEVELKNVTENFNMWENNMVEQMHEDIISLWDQSLKPCVK | 120 |
| 14 | DTEAHNVWATHACVPTDPNPQEVELKNVTENFNMWENNMVEQMHEDIISLWDQSLKPCVK | 120 |
| 15 | DTEAHNVWATHACVPTDPNPQEVELKNVTENFNMWENNMVEQMHEDIISLWDQSLKPCVK | 120 |
| 16 | DTEAHNVWATHACVPTDPNPQEVELKNVTENFNMWENNMVEQMHEDIISLWDQSLKPCVK | 120 |
| 17 | DTEAHNVWATHACVPTDPNPQEVELKNVTENFNMWENNMVEQMHEDIISLWDQSLKPCVK | 120 |
| 18 | DTEAHNVWATHACVPTDPNPQEVELKNVTENFNMWENNMVEQMHEDIISLWDQSLKPCVK | 120 |
| 19 | DTEAHNVWATHACVPTDPNPQEVELKNVTENFNMWENNMVEQMHEDIISLWDQSLKPCVK | 120 |
| 20 | DTEAHNVWATHACVPTDPNPQEVELKNVTENFNMWENNMVEQMHEDIISLWDQSLKPCVK | 120 |
| 21 | DTEAHNVWATHACVPTDPNPQEVELKNVTENFNMWENNMVEQMHEDIISLWDQSLKPCVK | 120 |
| 22 | DTEAHNVWATHACVPTDPNPQEVELKNVTENFNMWENNMVEQMHEDIISLWDQSLKPCVK | 120 |
| 23 | DTEAHNVWATHACVPTDPNPQEVELKNVTENFNMWENNMVEQMHEDIISLWDQSLKPCVK | 120 |
| 24 | DTEAHNVWATHACVPTDPNPQEVELKNVTENFNMWENNMVEQMHEDIISLWDQSLKPCVK | 120 |
| 25 | DTEAHNVWATHACVPTDPNPQEVELKNVTENFNMWENNMVEQMHEDIISLWDQSLKPCVK | 120 |
| 26 | DTEAHNVWATHACVPTDPNPQEVELKNVTENFNMWENNMVEQMHEDIISLWDQSLKPCVK | 120 |
| 27 | DTEAHNVWATHACVPTDPNPQEVELKNVTENFNMWENNMVEQMHEDIISLWDQSLKPCVK | 120 |
| 28 | DTEAHNVWATHACVPTDPNPQEVELKNVTENFNMWENNMVEQMHEDIISLWDQSLKPCVK | 120 |
| 29 | DTEAHNVWATHACVPTDPNPQEVELKNVTENFNMWENNMVEQMHEDIISLWDQSLKPCVK | 120 |
| 30 | DTEAHNVWATHACVPTDPNPQEVELKNVTENFNMWENNMVEQMHEDIISLWDQSLKPCVK | 120 |
| 31 | DTEAHNVWATHACVPTDPNPQEVELKNVTENFNMWENNMVEQMHEDIISLWDQSLKPCVK | 120 |
| 32 | DTEAHNVWATHACVPTDPNPQEVELKNVTENFNMWENNMVEQMHEDIISLWDQSLKPCVK | 120 |
| 33 | DTEAHNVWATHACVPTDPNPQEVELKNVTENFNMWENNMVEQMHEDIISLWDQSLKPCVK | 120 |
| 34 | DTEAHNVWATHACVPTDPNPQEVELKNVTENFNMWENNMVEQMHEDIISLWDQSLKPCVK | 120 |
| 35 | DTEAHNVWATHACVPTDPNPQEVELKNVTENFNMWENNMVEQMHEDIISLWDQSLKPCVK | 120 |
| 36 | DTEAHNVWATHACVPTDPNPQEVELKNVTENFNMWENNMVEQMHEDIISLWDQSLKPCVK | 120 |
| 37 | DTEAHNVWATHACVPTDPNPQEVELKNVTENFNMWENNMVEQMHEDIISLWDQSLKPCVK | 120 |
| 38 | DTEAHNVWATHACVPTDPNPQEVELKNVTENFNMWENNMVEQMHEDIISLWDQSLKPCVK | 120 |
| 39 | DTEAHNVWATHACVPTDPNPQEVELKNVTENFNMWENNMVEQMHEDIISLWDQSLKPCVK | 120 |
| 40 | DTEAHNVWATHACVPTDPNPQEVELKNVTENFNMWENNMVEQMHEDIISLWDQSLKPCVK | 120 |

```
    LTPLCVTLNCTDLGNVTNTT---NSKGXMMEKGEVKNCSFKITTDIKDRTRKEYALFYKL
1   LTPLCVTLNCTDLGNVTNTT---NSNGEMMEKGEVKNCSFKITTDIKDRTRKEYALFYKL  177
2   LTPLCVTLNCTDLGNVTNTT---NSKGEMMEKGEVKNCSFKITTDIKDRTRKEYALFYKL  176
3   LTPLCVTLNCTDLGNVTNTT---NSKGEMMEKGEVKNCSFKITTDIKDRTRKEYALFYKL  177
4   LTPLCVTLNCTDLGNVTNTT---NSNGKMMEKGEVKNCSFKITTDIKDRTRKEYALFYKL  177
5   LTPLCVTLNCTDLGNVTNTT---NSNEKMMEKGEVKNCSFKITTDIKDRTRKEYALFYKL  177
6   LTPLCVTLNCTDLGNVTNTTN--SNGERMMEKGEVKNCSFKITTDIKDRTRKEYALFYKL  178
7   LTPLCVTLNCTDLGNVTNTT---NSKGEMMEKGEVKNCSFKITTDIKDRTRKEYALFYKL  177
8   LTPLCVTLNCTDLGNVTNTA---NSNGTMMEKGEVKNCSFKITTDIKDRTRKEYALFYKL  177
9   LTPLCVTLNCTDLENVTNTT---NSTEKMMEKGEVKNCSFKITTDIKDRTRKEYALFYKL  177
10  LTPLCVTLNCTDLENVTNTT---NSTEKMMEKGEVKNCSFKITTDIKDRTRKEYALFYKL  177
11  LTPLCVTLNCTDLGNVTNTT---NSKGEMMEKGEVKNCSFKITTDIKDRTRKEYALFYKL  177
12  LTPLCVTLNCTDLGNVTNTT---NSKGEMMEKGEVKNCSFKITTDIKDRTRKEYALFYKL  177
13  LTPLCVTLNCTDLENVTNTT---NSNEKMMEKGEVKNCSFKITTDIKDRTRKEYALFYKL  177
14  LTPLCVTLNCTDLGNVTNTT---NSKGEMMEKGEVKNCSFKITTDIKDRTRKEYALFYKL  177
15  LTPLCVTLNCTDLGNVTNTTN--SNGERMMEKGEVKNCSFKITTDIKDRTRKEYALFYKL  178
16  LTPLCVTLNCTDLGNVTNTT---NSKGEMMEKGEVKNCSFKITTDIKDRTRKEYALFYKL  177
17  LTPLCVTLNCTDLGNVTNTT---NSKGEMMEKGEVKNCSFKITTDIKDRTRKEYALFYKL  177
18  LTPLCVTLNCTDLGNVTNTT---NSKGEMMEKGEVKNCSFKITTDIKDRTRKEYALFYKL  177
19  LTPLCVTLNCTDLGNVTNTT---NSKGGMMEKGEVKNCSFKITTDIKDRTRKEYALFYKL  177
20  LTPLCVTLNCTDLENVTNTT---NSTEKMMEKGEVKNCSFKITTDIKDRTRKEYALFYKL  177
21  LTPLCVTLNCTDLGNVTNTT---NSKGGMMEKGEVKNCSFKITTDIKDRTRKEYALFYKL  177
22  LTPLCVTLNCTDLGNVTNTT---NSKGEMMEKGEVKNCSFKITTDIKDRTRKEYALFYKL  177
23  LTPLCVTLNCTDLGNVTNTT---NSKGEMMEKGEVKNCSFKITTDIKDRTRKEYALFYKL  177
24  LTPLCVTLNCTDLGNVTNTTN--SNGERMMEKGEVKNCSFKITTDIKDRTRKEYALFYKL  178
25  LTPLCVTLNCTDLGNVTNTT---NSKGEMMEKGEVKNCSFKITTDIKDRTRKEYALFYKL  177
26  LTPLCVTLNCTDLGNVTTTT---NSNGTLMEKGEVKNCSFKITTDIKDRTRKEYALFYKL  177
27  LTPLCVTLNCTDLGNVTNTT---NSKGEMMEKGEVKNCSFKITTDIKDRTRKEYALFYKL  177
28  LTPLCVTLNCTDLGNVTNTT---NSKGGMMEKGEVKNCSFKITTDIKDRTRKEYALFYKL  177
29  LTPLCVTLNCTDLGNVTNTT---NSKGGMMEKGEVKNCSFKITTDIKDRTRKEYALFYKL  177
30  LTPLCVTLNCTDLGNVTTTT---NSNGTLMEKGEVKNCSFKITTDIKDRTRKEYALFYKL  177
31  LTPLCVTLNCTDLGNVTNTT---NSKGGMMEKGEVKNCSFKITTDIKDRTRKEYALFYKL  177
32  LTPLCVTLNCTDLENVTNTT---NSNEKMMEKGEVKNCSFKITTDIKDRTRKEYALFYKL  177
33  LTPLCVTLNCTDLENVTNTT---NSTEKMMEKGEVKNCSFKITTDIKDRTRKEYALFYKL  177
34  LTPLCVTLNCTDLGNVTNTT---NSKGGMMEKGEVKNCSFKITTDIKDRTRKEYALFYKL  177
35  LTPLCVTLNCTDLGNVTNTT---NSNGKMMEKGEVKNCSFKITTDIKDRTRKEYALFYKL  177
36  LTPLCVTLNCTDLGNVTNTT---NSNGRMMEKGEVKNCSFKITTDIKDRTRKEYALFYKL  177
37  LTPLCVTLNCTDLGNVTNTT---NSKGEMMEKGEVKNCSFKITTDIKDRTRKEYALFYKL  177
38  LTPLCVTLNCTDLGNVTNTT---NSKGEMMEKGEVRNCSFKITTDIKDRTRKEYALFYKL  177
39  LTPLCVTLNCTDLGNVTNTT---NSKGEMMEKGEVKNCSFKITTDIKDRTRKEYALFYKL  177
40  LTPLCVTLNCTDLGNVTNTT---NSKGGMMEKGEVKNCSFKITTDIKDRTRKEYALFYKL  177
```

| | Sequence | |
|---|---|---|
|  | DVVPIND-------TRYRLVSCNTSVITQACPKVSFEPIPIHYCAPAGFAILKCNDKQFI |  |
| 1 | DVVPIND-------TRYRLVSCNTSVITQACPKVSFEPIPIHYCAPAGFAILKCNDKQFI | 230 |
| 2 | DVVPIND-------TRYRLVSCNTSVITQACPKVSFEPIPIHYCAPAGFAILKCNDKQFI | 229 |
| 3 | DVVPIND-------TRYRLVSCNTSVITQACPKVSFEPIPIHYCAPAGFAILKCNDKQFI | 230 |
| 4 | DVVPIND-------TRYRLVSCNTSVITQACPKVSFEPIPIHYCAPAGFAILKCNDKQFI | 230 |
| 5 | DVVPIND-------TRYRLVSCNTSVITQACPKVSFEPIPIHYCAPAGFAILKCNDKQFI | 230 |
| 6 | DVVPIND-------TRYRLVSCNTSVITQACPKVSFEPIPIHYCAPAGFAILKCNDKQFI | 231 |
| 7 | DVVPIND-------TRYRLVSCNTSVITQACPKVSFEPIPIHYCAPAGFAILKCNDKQFI | 230 |
| 8 | DVVPIND-------TRYRLVSCNTSVITQACPKVSFEPIPIHYCAPAGFAILKCNDKQFI | 230 |
| 9 | DVVPIND-------TRYRLVSCNTSVITQACPKVSFEPIPIHYCAPAGFAILKCNDKQFI | 230 |
| 10 | DVVPIND-------TRYRLVSCNTSVITQACPKVSFEPIPIHYCAPAGFAILKCNDKQFI | 230 |
| 11 | DVVPIND-------TRYRLVSCNTSVITQACPKVSFEPIPIHYCAPAGFAILKCNDKQFI | 230 |
| 12 | DVVPIND-------TRYRLVSCNTSVITQACPKVSFEPIPIHYCAPAGFAILKCNDKQFI | 230 |
| 13 | DVVPIND-------TRYRLVSCNTSVITQACPKVSFEPIPIHYCAPAGFAILKCNDKQFI | 230 |
| 14 | DVVPIND-------TRYRLVSCNTSVITQACPKVSFEPIPIHYCAPAGFAILKCNDKQFI | 230 |
| 15 | DVVPIND-------TRYRLVSCNTSVITQACPKVSFEPIPIHYCAPAGFAILKCNDKQFI | 231 |
| 16 | DVVPIND-------TRYRLVSCNTSVITQACPKVSFEPIPIHYCAPAGFAILKCNDKQFM | 230 |
| 17 | DVVPIND-------TRYRLVSCNTSVITQACPKVSFEPIPIHYCAPAGFAILKCNDKQFI | 230 |
| 18 | DVVPIND-------TRYRLVSCNTSVITQACPKVSFEPIPIHYCAPAGFAILKCNDKQFI | 230 |
| 19 | DVVPIND-------TRYRLVSCNTSVITQACPKVSFEPIPIHYCAPAGFAILKCNDKQFI | 230 |
| 20 | DVVPIND-------TRYRLVSCNTSVITQACPKVSFEPIPIHYCAPAGFAILKCNDKQFI | 230 |
| 21 | DVVPIND-------TRYRLVSCNTSVITQACPKVSFEPIPIHYCAPAGFAILKCNDKQFI | 230 |
| 22 | DVVPIND-------TRYRLVSCNTSVITQACPKVSFEPIPIHYCAPAGFAILKCNDKQFI | 230 |
| 23 | DVVPIND-------TRYRLVSCNTSVITQACPKVSFEPIPIHYCAPAGFAILKCNDKQFI | 230 |
| 24 | DVVPIND-------TRYRLVSCNTSVITQACPKVSFEPIPIHYCAPAGFAILKCNDKQFI | 231 |
| 25 | DVVPIND-------TRYRLVSCNTSVITQACPKVSFEPIPIHYCAPAGFAILKCNDKQFI | 230 |
| 26 | DVVPIND-------TRYRLVSCNTSVITQACPKVSFEPIPIHYCAPAGFAILKCNDKQFI | 230 |
| 27 | DVVPIND-------TRYRLVSCNTSVITQACPKVSFEPIPIHYCAPAGFAILKCNDKQFI | 230 |
| 28 | DVVPIND-------IRYRLVSCNTSVITQACPKVSFEPIPIHYCAPAGFAILKCNDKQFI | 230 |
| 29 | DVVPIND-------TRYRLVSCNTSVITQACPKVSFEPIPIHYCAPAGFAILKCNDKQFI | 230 |
| 30 | DVVPIND-------TRYRLVSCNTSVITQACPKVSFEPIPIHYCAPAGFAILKCNDKQFI | 230 |
| 31 | DVVPIND-------TRYRLVSCNTSVITQACPKVSFEPIPIHYCAPAGFAILKCNDKQFI | 230 |
| 32 | DVVPIND-------TRYRLVSCNTSVITQACPKVSFEPIPIHYCAPAGFAILKCNDKQFI | 230 |
| 33 | DVVPIND-------TRYRLVSCNTSVITQACPKVSFEPIPIHYCAPAGFAILKCNDKQFI | 230 |
| 34 | DVVPIND-------TRYRLVSCNTSVITQACPKVSFEPIPIHYCAPAGFAILKCNDKQFI | 230 |
| 35 | DVVPIND-------TRYRLVSCNTSVITQACPKVSFEPIPIHYCAPAGFAILKCNDKQFI | 230 |
| 36 | DVVPIND-------TRYRLVSCNTSVITQACPKVSFEPIPIHYCAPAGFAILKCNDKQFI | 230 |
| 37 | DVVPIND-------TRYRLVSCNTSVITQACPKVSFEPIPIHYCAPAGFAILKCNDKQFI | 230 |
| 38 | DVVPIND-------TRYRLVSCNTSVITQACPKVSFEPIPIHYCAPAGFAILKCNDKQFI | 230 |
| 39 | DVVPIND-------TRYRLVSCNTSVITQACPKVSFEPIPIHYCAPAGFAILKCNDKQFI | 230 |
| 40 | DVVPIND-------TRYRLVSCNTSVITQACPKVSFEPIPIHYCAPAGFAILKCNDKQFI | 230 |

| | | |
|---|---|---|
| | **GTGPCTNVSTVQCTHGIRPVVSTQLLLNGSLAEEEVVIRSVNFSDNAKTIIVQLNKSVEI** | |
| 1 | GTGPCTNVSTVQCTHGIRPVVSTQLLLNGSLAEEEVVIRSVNFSDNAKTIIVQLNKSVEI | 290 |
| 2 | GTGPCTNVSTVQCTHGIRPVVSTQLLLNGSLAEEEVVIRSVNFSDNAKTIIVQLNKSVEI | 289 |
| 3 | GTGPCTNVSTVQCTHGIRPVVSTQLLLNGSLAEEEVVIRSVNFSDNAKTIIVQLNKSVEI | 290 |
| 4 | GTGPCTNVSTVQCTHGIRPVVSTQLLLNGSLAEEEVVIRSVNFSDNAKTIIVQLNKSVEI | 290 |
| 5 | GTGPCTNVSTVQCTHGIRPVVSTQLLLNGSLAEEEVVIRSVNFSDNAKTIIVQLNKSVEI | 290 |
| 6 | GTGPCTNVSTVQCTHGIRPVVSTQLLLNGSLAEEEVVIRSVNFSDNAKTIIVQLNKSVEI | 291 |
| 7 | GTGPCTNVSTVQCTHGIRPVVSTQLLLNGSLAEEEVVIRSVNFSDNAKTIIVQLNKSVEI | 290 |
| 8 | GTGPCTNVSTVQCTHGIRPVVSTQLLLNGSLAEEEVVIRSVNFSDNAKTIIVQLNKSVEI | 290 |
| 9 | GTGPCTNVSTVQCTHGIRPGVSTQLLLNGSLAEEEVVIRSVNFSDNAKTIIVQLNKSVEI | 290 |
| 10 | GTGPCTNVSTVQCTHGIRPVVSTQLLLNGSLAEEEVVIRSVNFSDNAKTIIVQLNKSVEI | 290 |
| 11 | GTGPCTNVSTVQCTHGIRPVVSTQLLLNGSLAEEEVVIRSVNFSDNAKTIIVQLNKSVEI | 290 |
| 12 | GTGPCTNVSTVQCTHGIRPVVSTQLLLNGSLAEEEVVIRSVNFSDNAKTIIVQLNKSVEI | 290 |
| 13 | GTGPCTNVSTVQCTHGIRPVVSTQLLLNGSLAEEEVVIRSVNFSDNAKTIIVQLNKSVEI | 290 |
| 14 | GTGPCTNVSTVQCTHGIRPVVSTQLLLNGSLAEEEVVIRSVNFSDNAKTIIVQLNKSVEI | 290 |
| 15 | GTGPCTNVSTVQCTHGIRPVVSTQLLLNGSLAEEEVVIRSVNFSDNAKTIIVQLNKSVEI | 291 |
| 16 | GTGPCTNVSTVQCTHGIRPVVSTQLLLNGSLAEEEVVIRSVNFSDNAKTIIVQLNKSVEI | 290 |
| 17 | GTGPCTNVSTVQCTHGIRPVVSTQLLLNGSLAEEEVVIRSVNFSDNAKTIIVQLNKSVEI | 290 |
| 18 | GTGPCTNVSTVQCTHGIRPVVSTQLLLNGSLAEEEVVIRSVNFSDNAKTIIVQLNKSVEI | 290 |
| 19 | GTGPCTNVSTVQCTHGIRPVVSTQLLLNGSLAEEEVVIRSVNFSDNAKTIIVQLNKSVEI | 290 |
| 20 | GTGPCTNVSTVQCTHGIRPVVSTQLLLNGSLAEEEVVIRSVNFSDNAKTIIVQLNKSVEI | 290 |
| 21 | GTGPCTNVSTVQCTHGIRPVVSTQLLLNGSLAEEEVVIRSVNFSDNAKTIIVQLNKSVEI | 290 |
| 22 | GTGPCTNVSTVQCTHGIRPVVSTQLLLNGSLAEEEVVIRSVNFSDNAKTIIVQLNKSVEI | 290 |
| 23 | GTGPCTNVSTVQCTHGIRPVVSTQLLLNGSLAEEEVVIRSVNFSDNAKTIIVQLNKSVEI | 290 |
| 24 | GTGPCTNVSTVQCTHGIRPVVSTQLLLNGSLAEEEVVIRSVNFSDNAKTIIVQLNKSVEI | 291 |
| 25 | GTGPCTNVSTVQCTHGIRPVVSTQLLLNGSLAEEEVVIRSVNFSDNAKTIIVQLNKSVEI | 290 |
| 26 | GTGPCTNVSTVQCTHGIRPVVSTQLLLNGSLAEEEVVIRSVNFSDNAKTIIVQLNKSVEI | 290 |
| 27 | GTGPCTNVSTVQCTHGIRPVVSTQLLLNGSLAEEEVVIRSVNFSDNAKTIIVQLNKSVEI | 290 |
| 28 | GTGPCTNVSTVQCTHGIRPVVSTQLLLNGSLAEEEVVIRSVNFSDNAKTIIVQLNKSVEI | 290 |
| 29 | GTGPCTNVSTVQCTHGIRPVVSTQLLLNGSLAEEEVVIRSVNFSDNAKTIIVQLNKSVEI | 290 |
| 30 | GTGPCTNVSTVQCTHGIRPVVSTQLLLNGSLAEEEVVIRSVNFSDNAKTIIVQLNKSVEI | 290 |
| 31 | GTGPCTNVSTVQCTHGIRPVVSTQLLLNGSLAEEEVVIRSVNFSDNAKTIIVQLNKSVEI | 290 |
| 32 | GTGPCTNVSTVQCTHGIRPVVSTQLLLNGSLAEEEVVIRSVNFSDNAKTIIVQLNKSVEI | 290 |
| 33 | GTGPCTNVSTVQCTHGIRPVVSTQLLLNGSLAEEEVVIRSVNFSDNAKTIIVQLNKSVEI | 290 |
| 34 | GTGPCTNVSTVQCTHGIRPVVSTQLLLNGSLAEEEVVIRSVNFSDNAKTIIVQLNKSVEI | 290 |
| 35 | GTGPCTNVSTVQCTHGIRPVVSTQLLLNGSLAEEEVVIRSVNFSDNAKTIIVQLNKSVEI | 290 |
| 36 | GTGPCTNVSTVQCTHGIRPVVSTQLLLNGSLAEEEVVIRSVNFSDNAKTIIVQLNKSVEI | 290 |
| 37 | GTGPCTNVSTVQCTHGIRPVVSTQLLLNGSLAEEEVVIRSVNFSDNAKTIIVQLNKSVEI | 290 |
| 38 | GTGPCTNVSTVQCTHGIRPVVSTQLLLNGSLAEEEVVIRSVNFSDNAKTIIVQLNKSVEI | 290 |
| 39 | GTGPCTNVSTVQCTHGIRPVVSTQLLLNGSLAEEEVVIRSVNFSDNAKTIIVQLNKSVEI | 290 |
| 40 | GTGPCTNVSTVQCTHGIRPVVSTQLLLNGSLAEEEVVIRSVNFSDNAKTIIVQLNKSVEI | 290 |

| | | |
|---|---|---|
| | NCTRPHNNTRKSIPMGPGKAFYARGDITGDIRKAYCEINGTEWHSTLKLVVEKLREQY-N | |
| 1 | TCTRPNNNTRKSIPMGPGKAFYARGDITGDIRKAYCEINGTEWHSTLKLVVEKLREQY-N | 349 |
| 2 | NCTRPHNNTRKSIPMGPGKAFYARGDIIGDIRKAYCEINGTEWHSTLKLVVEKLREQY-N | 348 |
| 3 | NCTRPHNNTRKSIPMGPGKAFYARGDIIGDIRKAYCEINGTEWHSTLKLVVEKLREQY-N | 349 |
| 4 | NCTRPHNNTRKSIPMGPGKAFYARGDITGDIRKAYCEINGTEWHSTLKLVVEKLREQY-N | 349 |
| 5 | NCTRPHNNTRKSIPMGPGKAFYARGDITGDIRKAYCEINGTEWHSTLKLVVEKLREQY-N | 349 |
| 6 | TCTRPNNNTRKSIPMGPGKAFYARGDIIGDIRKAYCKINGTEWHSTLKLVVEKLREQY-N | 350 |
| 7 | NCTRPHNNTRKSIPMGPGKAFYARGDIIGDIRKAYCEINGTEWHSTLKLVVEKLREQY-N | 349 |
| 8 | NCTRPNNNTRKSIPMGPGKAFYARGDITGDIRKAYCEINGTEWHSTLKLVVEKLREQY-N | 349 |
| 9 | NCTRPHNNTRKSIPMGPGKAFYARGDITGDIRKAYCEINGTEWHSTLKLVVEKLREQY-N | 349 |
| 10 | NCTRPHNNTRKSIPMGPGKAFYARGDITGDIRKAYCEINGTEWHSTLKLVVEKLREQY-N | 349 |
| 11 | NCTRPHNNTRKSIPMGPGKAFYARGDIIGDIRKAYCEINGTEWHSTLKLVVEKLREQY-N | 349 |
| 12 | NCTRPHNNTRKSIPMGPGKAFYARGDITGDIRKAYCEINGTEWHSTLKLVVEKLREQY-N | 349 |
| 13 | NCTRPHNNTRKSIPMGPGKAFYARGDITGDIRKAYCEINGTEWHSTLKLVVEKLREQY-N | 349 |
| 14 | NCTRPHNNTRKSIPMGPGKAFYARGDITGDIRKAYCEINGTEWHSTLKLVVEKLREQY-N | 349 |
| 15 | TCTRPNNNTRKSIPMGPGKAFYARGDIIGDIRKAYCKINGTEWHSTLKLVVEKLREQY-N | 350 |
| 16 | TCTRPNNNTRKSIPMGPGKAFYARGDIIGDIRKASCKINGTEWHSTLKLVVEKLREQY-N | 349 |
| 17 | NCTRPHNNTRKSIPMGPGKAFYARGDITGDIRKAYCEINGTEWHSTLKLVVEKLREQY-N | 349 |
| 18 | NCTRPHNNTRKSIPMGPGKAFYARGDIIGDIRKAYCEINGTEWHSTLKLVVEKLREQY-N | 349 |
| 19 | NCTRPHNNTRKSIPMGPGKAFYARGDITGDIRKAYCEINGTEWHSTLKLVVEKLREQY-N | 349 |
| 20 | NCTRPHNNTRKSIPMGPGKAFYARGDITGDIRKAYCEINGTEWHSTLKLVVEKLREQY-N | 349 |
| 21 | NCTRPHNNTRKSIPMGPGKAFYARGDITGDIRKAYCEINGTEWHSTLKLVVEKLREQY-N | 349 |
| 22 | NCTRPHNNTRKSIPMGPGKAFYARGDIIGDIRKAYCEINGTEWHSTLKLVVEKLREQY-N | 349 |
| 23 | NCTRPHNNTRKSIPMGPGKAFYARGDIIGDIRKAYCEINGTEWHSTLKLVVEKLREQY-N | 349 |
| 24 | TCTRPNNNTRKSIPMGPGKAFYARGDIIGDIRKAYCKINGTEWHSTLKLVVEKLREQY-N | 350 |
| 25 | NCTRPHNNTRKSIPMGPGKAFYARGDIIGDIRKAYCEINGTEWHSTLKLVVEKLREQY-N | 349 |
| 26 | NCTRPNNNTRKSIPMGPGKAFYARGDITGDIRKAYCEINGTEWHSTLKLVVEKLREQY-N | 349 |
| 27 | NCTRPHNNTRKSIPMGPGKAFYARGDITGDIRKAYCEINGTEWHSTLKLVVEKLREQY-N | 349 |
| 28 | NCTRPHNNTRKSIPMGPGKAFYARGDITGDIRKAYCEINGTEWHSTLKLVVEKLREQY-N | 349 |
| 29 | NCTRPHNNTRKSIPMGPGKAFYARGDITGDIRKAYCEINGTEWHSTLKLVVEKLREQY-N | 349 |
| 30 | NCTRPHNNTRKSIPMGPGKAFYARGDITGDIRKAYCEINGTEWHSTLKLVVEKLREQY-N | 349 |
| 31 | NCTRPHNNTRKSIPMGPGKAFYARGDITGDIRKAYCEINGTEWHSTLKLVVEKLREQY-N | 349 |
| 32 | NCTRPHNNTRKSIPMGPGKAFYARGDITGDIRKAYCEINGTEWHSTLKLVVEKLREQY-N | 349 |
| 33 | NCTRPHNNTRKSIPMGPGKAFYARGDITGDIRKAYCEINGTEWHSTLKLVVEKLREQY-N | 349 |
| 34 | NCTRPHNNTRKSIPMGPGKAFYARGDITGDIRKAYCEINGTEWHSTLKLVVEKLREQY-N | 349 |
| 35 | NCTRPNNNTRKSIPMGPGKAFYARGDITGDIRKAYCEINGTEWHSTLKLVVEKLREQY-N | 349 |
| 36 | TCTRPNNNTRKSIPMGPGKAFYARGDIIGDIRKASCEINGTEWHSTLKLVVEKLREQY-N | 349 |
| 37 | NCTRPHNNTRKSIPMGPGKAFYARGDIIGDIRKAYCEINGTEWHSTLKLVVEKLREQY-N | 349 |
| 38 | NCTRPHNNTRKSIPMGPGKAFYARGDIIGDIRKAYCEINGTEWHSTLKLVVEKLREQY-N | 349 |
| 39 | NCTRPHNNTRKSIPMGPGKAFYARGDITGDIRKAYCEINGTEWHSTLKLVVEKLREQY-N | 349 |
| 40 | NCTRPHNNTRKSIPMGPGKAFYARGDITGDIRKAYCEVNGTEWHSTLKLVVEKLREQY-N | 349 |

| | | |
|---|---|---|
| | KTIVFNRSSGGDPEIVMYSFNCGGEFFYCNSTKLFNSTWPWND-TKGSHDTNGTLILPCK | |
| 1 | KTIVFNRSSGGDPEIVMYSFNCGGEFFYCNSTKLFNSTWPWND-TKGSHDTNGTLILPCK | 408 |
| 2 | KTIVFNRSSGGDPEIVMYSFNCGGEFFYCNSTKLFNSTWPWND-TKGSHDTNGTLILPCK | 407 |
| 3 | KTIVFNRSSGGDPEIVMYSFNCGGEFFYCNSTKLFNSTWPWND-TKGSHDTNGTLILPCK | 408 |
| 4 | KTIVFNRSSGGDPEIVMYSFNCGGEFFYCNSTKLFNSTWPWND-TKGSHDTNDTLMLPCK | 408 |
| 5 | KTIVFNRSSGGDPEIVMYSFNCGGEFFYCNSTKLFNSTWPWND-TKGSHDTNDTLILPCK | 408 |
| 6 | KTIVFNRSSGGDPEIVMYSFNCGGEFFYCNSTKLFNSTWPWND-TKGSHDTNDTLMLPCK | 409 |
| 7 | KTIVFNRSSGGDPEIVMYSFNCGGEFFYCNSTKLFNSTWPWND-TKGSHDTNGTLILPCK | 408 |
| 8 | KTIVFNRSSGGDPEIVMYSFNCGGEFFYCNSTKLFNSTWPWND-TKGSHDTNGTLTLPCK | 408 |
| 9 | KTIVFNRSSGGDPEIVMYSFNCGGEFFYCNSTKLFNSTWPWND-TKGSHDTNGKLILPCK | 408 |
| 10 | KTIVFNRSSGGDPEIVMYSFNCGGEFFYCNSTKLFNSTWPWND-TKGSHDTNGKLILPCK | 408 |
| 11 | KTIVFNRSSGGDPEIVMYSFNCGGEFFYCNSTKLFNSTWPWND-TKGSHDTNGTLILPCK | 408 |
| 12 | KTIVFNRSSGGDPEIVMYSFNCGGEFFYCNSTKLFNSTWPWND-TKGSHDTNGTLILPCK | 408 |
| 13 | KTIVFNRSSGGDPEIVMYSFNCGGEFFYCNSTKLFNSTWPWND-TKGSHDTNGTLILPCK | 408 |
| 14 | KTIVFNRSSGGDPEIVMYSFNCGGEFFYCNSTKLFNSTWPWND-TKGSHDTNGTLILPCK | 408 |
| 15 | KTIVFNRSSGGDPEIVMYSFNCGGEFFYCNSTKLFNSTWPWND-TKGSHDTNDTLMLPCK | 409 |
| 16 | KTIVFNRSSGGDPEIVMYSFNCGGEFFYCNSTKLFNSTWPWND-TKGSHDTNGTLILPCK | 408 |
| 17 | KTIVFNRSSGGDPEIVMYSFNCGGEFFYCNSTKLFNSTWPWND-TKGSHDTNDTLILPCK | 408 |
| 18 | KTIVFNRSSGGDPEIVMYSFNCGGEFFYCNSTKLFNSTWPWND-TKGSHDTNGTLILPCK | 408 |
| 19 | KTIVFNRSSGGDPEIVMYSFNCGGEFFYCNSTKLFNSTWPWND-TKGSHDTNGTLILPCK | 408 |
| 20 | KTIVFNRSSGGDPEIVMHSFNCGGEFFYCNSTKLFNSTWPWND-TKGSHDTNGKLILPCK | 408 |
| 21 | KTIVFNRSSGGDPEIVMYSFNCGGEFFYCNSTKLFNSTWPWND-TKGSHDTNGTLILPCK | 408 |
| 22 | KTIVFNRSSGGDPEIVMYSFNCGGEFFYCNSTKLFNSTWPWND-TKGSHDTNGTLILPCK | 408 |
| 23 | KTIVFNRSSGGDPEIIMYSFNCGGEFFYCNSTKLFNSTWPWND-TKGSHDTNGTLILPCK | 408 |
| 24 | KTIVFNRSSGGDPEIVMYSFTCGGEFFYCNSTKLFNSTWPWND-TKGSHDTNDTLMLPCK | 409 |
| 25 | KTIVFNRSSGGDPEIVMYSFNCGGEFFYCNSTKLFNSTWPWND-TKGSHDTNGTLILPCK | 408 |
| 26 | KTIVFNRSSGGDPEIVMYSFNCGGEFFYCNSTKLFNSTWPWND-TKGSHDTNGTLILPCK | 408 |
| 27 | KTIVFNRSSGGDPEIVMYSFNCGGEFFYCNSTKLFNSTWPWND-TKGSHDTNGTLILPCK | 408 |
| 28 | KTIVFNRSSGGDPEIVMYSFNCGGEFFYCNSTKLFNSTWPWND-TKGSHDTNGTLILPCK | 408 |
| 29 | KTIVFNRSSGGDPEIVMYSFNYGGEFFYCNSTKLFNSTWPWND-TKGSHDTNGTLILPCK | 408 |
| 30 | KTIVFNRSSGGDPEIVMYSFNCGGEFFYCNSTKLFNSTWPWND-TKGSHDTNGTLILPCK | 408 |
| 31 | KTIVFNRSSGGDPEIVMYSFNCGGEFFYCNSTKLFNSTWPWND-TKGSHDTNGTLILPCK | 408 |
| 32 | KTIVFNRSSGGDPEIVMYSFNCGGEFFYCNSTKLFNSTWPWND-TKGSHDTNGTLILPCK | 408 |
| 33 | KTIVFNRSSGGDPEIVMYSFNCGGEFFYCNSTKLFNSTWPWND-TKGSHDTNGKLILPCK | 408 |
| 34 | KTIVFNRSSGGDPEIVMYSFNCGGEFFYCNSTKLFNSTWPWND-TKGSHDTNGTLILPCK | 408 |
| 35 | KTIVFNRSSGGDPEIVMYSFNCGGEFFYCNSTKLFNSTWPWND-TKGSHDTNDTLMLPCK | 408 |
| 36 | KTIVFNRSSGGDPEIVMYSFNCGGEFFYCNSTKLFNSTWPWND-TKGSHDTNGTLILPCK | 408 |
| 37 | KTIVFNRSSGGDPEIVMYSFNCGGEFFYCNSTKLFNSTWPWND-TKGSHDTNGTLILPCK | 408 |
| 38 | KTIVFNRSSGGDPEIVMYSFNCGGEFFYCNSTKLFNSTWPWND-TKGSHDTNGTLILPCK | 408 |
| 39 | KTIVFNRSSGGDPEIVMYSFNCGGEFFYCNSTKLFNSTWPWND-TKGSHDTNGTLILPCK | 408 |
| 40 | KTIVFNRSSGGDPEIVMYSFNCGGEFFYCNSTKLFNSTWPWND-TKGSHDTNGTLILPCK | 408 |

| | | |
|---|---|---|
| | IKQIINMWQGVGKAMYAPPIEGKIRCSSNITGLLLTRDGG-YESNETDEIFRPGGGDMRD | |
| 1 | IKQIINMWQGVGKAMYAPPIEGKIRCSSNITGLLLTRDGG-YESNETDEIFRPGGGDMRD | 467 |
| 2 | IKQIINMWQGVGKAMYAPPIEGKIRCSSNITGLLLTRDGG-YESNETDEIFRPGGGDMRD | 466 |
| 3 | IKQIINMWQGVGKAMYAPPIEGKIRCSSNITGLLLTRDGG-YESNETDEIFRPGGGDMRD | 467 |
| 4 | IKQIINMWQGVGKAMYAPPIEGKIRCSSNITGLLLTRDGG-YESNKTDEIFRPGGGDMRD | 467 |
| 5 | IKQIINMWQGVGKAMYAPPIEGKIRCSSNITGLLLTRDGG-YESNETDEIFRPGGGDMRD | 467 |
| 6 | IKQIINMWQGVGKAMYAPPIEGEIRCSSNITGLLLTRDGG-YESNETDEIFRPGGGDMRD | 468 |
| 7 | I-QIINMWQGVGKAMYAPPIEGKIRCSSNITGLLLTRDGG-YESNETDEIFRPGGGDMRD | 466 |
| 8 | IKQIINMWQGVGKAMYAPPIEGQIRCSSNITGLLLTRDGG-YESNETDEIFRPGGGDMRD | 467 |
| 9 | IKQIINMWQGVGKAMYAPPIEGKIRCSSNITGLLLTRDGGYESNETDDEIFRPGGGDMRD | 468 |
| 10 | IKQIINMWQGVGKAMYAPPIEGKIRCSSNITGLLLTRDGG-YESNETDEIFRPGGGDMRD | 467 |
| 11 | IKQIINMWQGVGKAMYAPPIEGKIRCSSNITGLLLTRDGG-YESNETDEIFRPGGGDMRD | 467 |
| 12 | IKQIINMWQGVGKAMYAPPIEGKIRCSSNITGLLLTRDGG-YESNETDEIFRPGGGDMRD | 467 |
| 13 | IKQIINMWQGVGKAMYAPPIEGKIRCSSNITGLLLTRDGG-YESNETDEIFRPGGGDMRD | 467 |
| 14 | IKQIINMWQGVGKAMYAPPIEGKIRCSSNITGLLLARDGG-YESNETDEIFRPGGGDMRD | 467 |
| 15 | IKQIINMWQGVGKAMYAPPIEGEIRCSSNITGLLLTRDGG-YESNETDEIFRPGGGDMRD | 468 |
| 16 | IKQIINMWQGVGKAMYAPPIEGEIRCSSNITGLLLTRDGG-YESNETDEIFRPGGGDMRD | 467 |
| 17 | IKQIINMWQGVGKAMYAPPIEGKIRCSSNITGLLLTRDGG-YESNETDEIFRPGGGDMRD | 467 |
| 18 | IKQIINMWQGVGKAMYAPPIEGKIRCSSNITGLLLTRDGG-YESNETDEIFRPGGGDMRD | 467 |
| 19 | IKQIINMWQGVGKAMYAPPIEGKIRCSSNITGLLLTRDGG-YESNETDEIFRPGGGDMRD | 467 |
| 20 | IKQIINMWQGVGKAMYAPPIEGKIRCSSNITGLLLTRDGG-YESNETDEIFRPGGGDMRD | 467 |
| 21 | IKQIINMWQGVGKAMYAPPIEGKIRCSSNITGLLLTRDGG-YESNETDEIFRPGGGDMRD | 467 |
| 22 | IKQIINMWQGVGKAMYAPPIEGKIRCSSNITGLLLTRDGG-YESNETDEIFRPGGGDMRD | 467 |
| 23 | IKQIINMWQGVGKAMYAPPIEGKIRCSSNITGLLLTRDGG-YESNETDEIFRPGGGDMRD | 467 |
| 24 | IKQIINMWQGVGKAMYAPPIEGEIRCSSNITGLLLTRDGG-YESNETDEIFRPGGGDMRD | 468 |
| 25 | IKQIINMWQGVGKAMYAPPIEGKIRCSSNITGLLLTRDGG-YESNETDEIFRPGGGDMRD | 467 |
| 26 | IKQIINMWQGVGKAMYAPPIEGKIRCSSNITGLLLTRDGG-YESNKTDEIFRPGGGDMRD | 467 |
| 27 | IKQIINMWQGVGKAMYAPPIEGKIRCSSNITGLLLARDGG-YESNETDEIFRPGGGDMRD | 467 |
| 28 | IKQIINMWQGVGKAMYAPPIEGKIRCSSNITGLLLTRDGG-YESNETDEIFRPGGGDMRD | 467 |
| 29 | IKQIINMWQGVGKAMYAPPIEGKIRCSSNITGLLLTRDGG-YESNETDEIFRPGGGDMRD | 467 |
| 30 | IKQIINMWQGVGKAMYAPPIEGKIRCSSNITGLLLARDGG-YESNETDEIFRPGGGDMRD | 467 |
| 31 | IKQIINMWQGVGKAMYAPPIEGKIRCSSNITGLLSTRDGG-YESNETDEIFRPGGGDMRD | 467 |
| 32 | IKQIINMWQGVGKAMYAPPIEGKIRCSSNITGLLLTRDGG-YESNETDEIFRPGGGDMRD | 467 |
| 33 | IKQIINMWQGVGKAMYAPPIEGKIRCSSNITGLLLTRDGG-YESNETDEIFRPGGGDMRD | 467 |
| 34 | IKQIINMWQGVGKAMYAPPIEGKIRCSSNITGLLSTRDGG-YESNETDEIFRPGGGDMRD | 467 |
| 35 | IKQIINMWQGVGKAMYAPPIEGIIRCSSNITGLLLTRDGG-YESNETDEIFRPGGGDMRD | 467 |
| 36 | IKQIINMWQGVGKAMYAPPIEEKIRCSSNITGLLLTRDGG-YESNETDEIFRPGGGDMRD | 467 |
| 37 | IKQIINMWQGVGKAMYAPPIEGKIRCSSNITGLLLTRDGG-YESNETDEIFRPGGGDMRD | 467 |
| 38 | IKQIINMWQGVGKAMYAPPIEGKIRCSSNITGLLLTRDGG-YESNETDEIFRPGGGDMRD | 467 |
| 39 | IKQIINMWQGVGKAMYAPPIEGKIRCSSNITGLLLARDGG-YESNETDEIFRPGGGDMRD | 467 |
| 40 | IKQIINMWQGVGKAMYAPPIEGKIRCSSNITGLLLTRDGG-YESNETDEIFRPGGGDMRD | 467 |

| | Sequence | |
|---|---|---|
| | **NWRSELYKYKVVKIEPLGVAPTKAKRRVVQREKRAFG-LGAVFLGFLGAAGSTMGAASIT** | |
| 1 | NWRSELYKYKVVKIEPLGVAPTKAKRRVVQREKRAFG-LGAVFLGFLGAAGSTMGAASIT | 526 |
| 2 | NWRSELYKYKVVKIEPLGVAPTKAKRRVVQREKRAFG-LGAVFLGFLGAAGSTMGAASIT | 525 |
| 3 | NWRSELYKYKVVKIEPLGVAPTKAKRRVVQREKRAFG-LGAVFLGFLGAAGSTMGAASIT | 526 |
| 4 | NWRSELYKYKVVKIEPLGVAPTKAKRRVVQREKRAFG-LGAVFLGFLGAAGSTMGAASIT | 526 |
| 5 | NWRSELYKYKVVKIEPLGVAPTKAKRRVVQREKRAFG-LGAVFLGFLGAAGSTMGAASIT | 526 |
| 6 | NWRSELYKYKVVKIEPLGVAPTKAKRRVVQREKRAFG-LGAVFLGFLGAAGSTMGAASIT | 527 |
| 7 | NWRSELYKYKVVKIEPLGVAPTKAKRRVVQREKRAFG-LGAVFLGFLGAAGSTMGAASIT | 525 |
| 8 | NWRSELYKYKVVKIEPLGVAPTKAKRRVVQREKRAFG-LGAVFLGFLGAAGSTMGAASIT | 526 |
| 9 | NWRSELYKYKVVKIEPLGVAPTKAKRRVVQREKRAFG-LGAVFLGFLGAAGSTMGAASIT | 527 |
| 10 | NWRSELYKYKVVKIEPLGVAPTKAKRRVVQREKRAFG-LGAVFLGFLGAAGSTMGAASIT | 526 |
| 11 | NWRSELYKYKVVKIEPLGVAPTKAKRRVVQREKRAFG-LGAVFLGFLGAAGSTMGAASIT | 526 |
| 12 | NWRSELYKYKVVKIEPLGVAPTKAKRRVVQREKRAFG-LGAVFLGFLGAAGSTMGAASIT | 526 |
| 13 | NWRSELYKYKVVKIEPLGVAPTKAKRRVVQREKRAFG-LGAVFLGFLGAAGSTMGAASIT | 526 |
| 14 | NWRSELYKYKVVKIEPLGVAPTKAKRRVVQREKRAFG-LGAVFLGFLGAAGSTMGAASIT | 526 |
| 15 | NWRSELYKYKVVKIEPLGVAPTKAKRRVVQREKRAFG-LGAVFLGFLGAAGSTMGAASIT | 527 |
| 16 | NWRSELYKYKVVKIEPLGVAPTKAKRRVVQREKRAFG-LGAVFLGFLGAAGSTMGAASIT | 526 |
| 17 | NWRSELYKYKVVKIEPLGVAPTKAKRRVVQREKRAFG-LGAVFLGFLGAAGSTMGAASIT | 526 |
| 18 | NWRSELYKYKVVKIEPLGVAPTKAKRRVVQRE--AFG-LGAVFLGFLGAAGSTMGAASIT | 524 |
| 19 | NWRSELYKYKVVKIEPLGVAPTKAKRRVVQREKRAFG-LGAVFLGFLGAAGSTMGAASIT | 526 |
| 20 | NWRSELYKYKVVKIEPLGVAPTKAKRRVVQREKRAFG-LGAVFLGFLGAAGSTMGAASIT | 526 |
| 21 | NWRSELYKYKVVKIEPLGVAPTKAKRRVVQREKRAFG-LGAVFLGFLGAAGSTMGAASIT | 526 |
| 22 | NWRSELYKYKVVKIEPLGVAPTKAKRRVVQREKRAFG-LGAVFLGFLGAAGSTMGAASIT | 526 |
| 23 | NWRSELYKYKVVKIEPLGVAPTKAKRRVVQREKRAFG-LGAVFLGFLGAAGSTMGAASIT | 526 |
| 24 | NWRSELYKYKVVKIEPLGVAPTKAKRRVVQREKRAFG-LGAVFLGFLGAAGSTMGAASIT | 527 |
| 25 | NWRSELYKYKVVKIEPLGVAPTKAKRRVVQREKRAFG-LGAVFLGFLGAAGSTMGAASIT | 526 |
| 26 | NWRSELYKYKVVKIEPLGVAPTKAKRRVVQREKRAFG-LGAVFLGFLGAAGSTMGAASIT | 526 |
| 27 | NWRSELYKYKVVKIEPLGVAPTKAKRRVVQREKRAFG-LGAVFLGFLGAAGSTMGAASIT | 526 |
| 28 | NWRSELYKYKVVKIEPLGVAPTKAKRRVVQREKRAFG-LGAVFLGFLGAAGSTMGAASIT | 526 |
| 29 | NWRSELYKYKVVKIEPLGVAPTKAKRRVVQREKRAFG-LGAVFLGFLGAAGSTMGAASIT | 526 |
| 30 | NWRSELYKYKVVKIEPLGVAPTKAKRRVVQREKRAFG-LGAVFLGFLGAAGSTMGAASIT | 526 |
| 31 | NWRSELYKYKVVKIEPLGVAPTKAKRRVVQREKRAFG-LGAVFLGFLGAAGSTMGAASIT | 526 |
| 32 | NWRSELYKYKVVKIEPLGVAPTKAKRRVVQREKRAFG-LGAVFLGFLGAAGSTMGAASIT | 526 |
| 33 | NWRSELYKYKVVKIEPLGVAPTKAKRRVVQREKRAFG-LGAVFLGFLGAAGSTMGAASIT | 526 |
| 34 | NWRSELYKYKVVKIEPLGVAPTKAKRRVVQREKRAFG-LGAVFLGFLGAAGSTMGAASIT | 526 |
| 35 | NWRSELYKYKVVKIEPLGVAPTKAKRRVVQREKRAFG-LGAVFLGFLGAAGSTMGAASIT | 526 |
| 36 | NWRSELYKYKVVKIEPLGVAPTKAKRRVVQREKRAFG-LGAVFLGFLGAAGSTMGAASIT | 526 |
| 37 | NWRSELYKYKVVKIEPLGVAPTKAKRRVVQREKRAFG-LGAVFLGFLGAAGSTMGAASIT | 526 |
| 38 | NWRSELYKYKVVKIEPLGVAPTKAKRRVVQREKRAFG-LGAVFLGFLGAAGSTMGAASIT | 526 |
| 39 | NWRSELYKYKVVKIEPLGVAPTKAKRRVVQREKRAFG-LGAVFLGFLGAAGSTMGAASIT | 526 |
| 40 | NWRSELYKYKVVKIEPLGVAPTKAKRRVVQREKRAFG-LGAVFLGFLGAAGSTMGAASIT | 526 |

| | | |
|---|---|---|
| | **LTVQARQLLSGIVQQQNNLLRAIEAQQHLLQLTVWGIKQLQARVLAVERYLKDQQLLGIW** | |
| 1 | LTVQARQLLSGIVQQQNNLLRAIEAQQHLLQLTVWGIKQLQARVLAVERYLKDQQLLGIW | 586 |
| 2 | LTVQARQLLSGIVQQQNNLLRAIEAQQHLLQLTVWGIKQLQARVLAVERYLKDQQLLGIW | 585 |
| 3 | LTVQARQLLSGIVQQQNNLLRAIEAQQHLLQLTVWGIKQLQARVLAVERYLKDQQLLGIW | 586 |
| 4 | LTVQARQLLSGIVQQQNNLLRAIEAQQHLLQLTVWGIKQLQARVLAVERYLKDQQLLGIW | 586 |
| 5 | LTVQARQLLSGIVQQQNNLLRAIEAQQHLLQLTVWGIKQLQARVLAVERYLKDQQLLGIW | 586 |
| 6 | LTVQARQLLSGIVQQQNNLLRAIEAQQHLLQLTVWGIKQLQARVLAVERYLKDQQLLGIW | 587 |
| 7 | LTVQARQLLSGIVQQQNNLLRAIEAQQHLLQLTVWGIKQLQARVLAVERYLKDQQLLGIW | 585 |
| 8 | LTVQARQLLSGIVQQQNNLLRAIEAQQHLLQLTVWGIKQLQARVLAVERYLKDQQLLGIW | 586 |
| 9 | LTVQARQLLSGIVQQQNNLLRAIEAQQHLLQLTVWGIKQLQARVLAVERYLKDQQLLGIW | 587 |
| 10 | LTVQARQLLSGIVQQQNNLLRAIEAQQHLLQLTVWGIKQLQARVLAVERYLKDQQLLGIW | 586 |
| 11 | LTVQARQLLSGIVQQQNNLLRAIEAQQHLLQLTVWGIKQLQARVLAVERYLKDQQLLGIW | 586 |
| 12 | LTVQARQLLSGIVQQQNNLLRAIEAQQHLLQLTVWGIKQLQARVLAVERYLKDQQLLGIW | 586 |
| 13 | LTVQARQLLSGIVQQQNNLLRAIEAQQHLLQLTVWGIKQLQARVLAVERYLKDQQLLGIW | 586 |
| 14 | LTVQARQLLSGIVQQQNNLLRAIEAQQHLLQLTVWGIKQLQARVLAVERYLKDQQLLGIW | 586 |
| 15 | LTVQARQLLSGIVQQQNNLLRAIEAQQHLLQLTVWGIKQLQARVLAVERYLKDQQLLGIW | 587 |
| 16 | LTVQARQLLSGIVQQQNNLLRAIEAQQHLLQLTVWGIKQLQARVLAVERYLKDQQLLGIW | 586 |
| 17 | LTVQARQLLSGIVQQQNNLLRAIEAQQHLLQLTVWGIKQLQARVLAVERYLKDQQLLGIW | 586 |
| 18 | LTVQARQLLSGIVQQQNNLLRAIEAQQHLLQLTVWGIKQLQARVLAVERYLKDQQLLGIW | 584 |
| 19 | LTVQARQLLSGIVQQQNNLLRAIEAQQHLLQLTVWGIKQLQARVLAVERYLKDQQLLGIW | 586 |
| 20 | LTVQARQLLSGIVQQQNNLLRAIEAQQHLLQLTVWGIKQLQARVLAVERYLKDQQLLGIW | 586 |
| 21 | LTVQARQLLSGIVQQQNNLLRAIEAQQHLLQLTVWGIKQLQARVLAVERYLKDQQLLGIW | 586 |
| 22 | LTVQARQLLSGIVQQQNNLLRAIEAQQHLLQLTVWGIKQLQARVLAVERYLKDQQLLGIW | 586 |
| 23 | LTVQARQLLSGIVQQQNNLLRAIEAQQHLLQLTVWGIKQLQARVLAVERYLKDQQLLGIW | 586 |
| 24 | LTVQARQLLSGIVQQQNNLLRAIEAQQHLLQLTVWGIKQLQARVLAVERYLKDQQLLGIW | 587 |
| 25 | LTVQARQLLSGIVQQQNNLLRAIEAQQHLLQLTVWGIKQLQARVLAVERYLKDQQLLGIW | 586 |
| 26 | LTVQARQLLSGIVQQQNNLLRAIEAQQHLLQLTVWGIKQLQARVLAVERYLKDQQLLGIW | 586 |
| 27 | LTVQARQLLSGIVQQQNNLLRAIEAQQHLLQLTVWGIKQLQARVLAVERYLKDQQLLGIW | 586 |
| 28 | LTVQARQLLSGIVQQQNNLLRAIEAQQHLLQLTVWGIKQLQARVLAVERYLKDQQLLGIW | 586 |
| 29 | LTVQARQLLSGIVQQQNNLLRAIEAQQHLLQLTVWGIKQLQARVLAVERYLKDQQLLGIW | 586 |
| 30 | LTVQARQLLSGIVQQQNNLLRAIEAQQHLLQLTVWGIKQLQARVLAVERYLKDQQLLGIW | 586 |
| 31 | LTVQARQLLSGIVQQQNNLLRAIEAQQHLLQLTVWGIKQLQARVLAVERYLKDQQLLGIW | 586 |
| 32 | LTVQARQLLSGIVQQQNNLLRAIEAQQHLLQLTVWGIKQLQARVLAVERYLKDQQLLGIW | 586 |
| 33 | LTVQARQLLSGIVQQQNNLLRAIEAQQHLLQLTVWGIKQLQARVLAVERYLKDQQLLGIW | 586 |
| 34 | LTVQARQLLSGIVQQQNNLLRAIEAQQHLLQLTVWGIKQLQARVLAVERYLKDQQLLGIW | 586 |
| 35 | LTVQARQLLSGIVQQQNNLLRAIEAQQHLLQLTVWGIKQLQARVLAVERYLKDQQLLGIW | 586 |
| 36 | LTVQARQLLSGIVQQQNNLLRAIEAQQHLLQLTVWGIKQLQARVLAVERYLKDQQLLGIW | 586 |
| 37 | LTVQARQLLSGIVQQQNNLLRAIEAQQHLLQLTVWGIKQLQARVLAVERYLKDQQLLGIW | 586 |
| 38 | LTVQARQLLSGIVQQQNNLLRAIEAQQHLLQLTVWGIKQLQARVLAVERYLKDQQLLGIW | 586 |
| 39 | LTVQARQLLSGIVQQQNNLLRAIEAQQHLLQLTVWGIKQLQARVLAVERYLKDQQLLGIW | 586 |
| 40 | LTVQARQLLSGIVQQQNNLLRAIEAQQHLLQLTVWGIKQLQARVLAVERYLKDQQLLGIW | 586 |

| | | |
|---|---|---|
| | GCSGKLICTTTVPWNTSWSNKSLEQIWDNMTWMEWEREIDNYTGYIYQLIEESQNQQEKN | |
| 1 | GCSGKLICTTTVPWNTSWSNKSLEQIWDNMTWMEWEREIDNYTGYIYQLIEESQNQQEKN | 646 |
| 2 | GCSGKLICTTTVPWNTSWSNKSLEQIWDNMTWMEWEREIDNYTGYIYQLIEESQNQQEKN | 645 |
| 3 | GCSGKLICTTTVPWNTSWSNKSLEQIWDNMTWMEWEREIDNYTGYIYQLIEESQNQQEKN | 646 |
| 4 | GCSGKLICTTTVPWNTSWSNKSLEQIWDNMTWMEWEREIDNYTGYIYQLIEESQNQQEKN | 646 |
| 5 | GCSGKLICTTTVPWNTSWSNKSLEQIWDNMTWMEWEREIDNYTGYIYQLIEESQNQQEKN | 646 |
| 6 | GCSGKLICTTTVPWNTSWSNKSLEQIWDNMTWMEWEREIDNYTGYIYQLIEESQNQQEKN | 647 |
| 7 | GCSGKLICTTTVPWNTSWSNKSLEQIWDNMTWMEWEREIDNYTGYIYQLIEESQNQQEKN | 645 |
| 8 | GCSGKLICTTTVPWNTSWSNKSLEQIWDNMTWMEWEREIDNYTGYIYQLIEESQNQQEKN | 646 |
| 9 | GCSGKLICTTTVPWNTSWSNKSLEQIWDNMTWMEWEREIDNYTGYIYQLIEESQNQQEKN | 647 |
| 10 | GCSGKLICTTTVPWNTSWSNKSLEQIWDNMTWMEWEREIDNYTGYIYQLIEESQNQQEKN | 646 |
| 11 | GCSGKLICTTTVPWNTSWSNKSLEQIWDNMTWMEWEREIDNYTGYIYQLIEESQNQQEKN | 646 |
| 12 | GCSGKLICTTTVPWNTSWSNKSLEQIWDNMTWMEWEREIDNYTGYIYQLIEESQNQQEKN | 646 |
| 13 | GCSGKLICTTTVPWNTSWSNKSLEQIWDNMTWMEWEREIDNYTGYIYQLIEESQNQQEKN | 646 |
| 14 | GCSGKLICTTTVPWNTSWSNKSLEQIWDNMTWMEWEREIDNYTGYIYQLIEESQNQQEKN | 646 |
| 15 | GCSGKLICTTTVPWNTSWSNKSLEQIWDNMTWMEWEREIDNYTGYIYQLIEESQNQQEKN | 647 |
| 16 | GCSGKLICTTTVPWNTSWSNKSLEQIWDNMTWMEWEREIDNYTGYIYQLIEESQNQQEKN | 646 |
| 17 | GCSGKLICTTTVPWNTSWSNKSLEQIWDNMTWMEWEREIDNYTGYIYQLIEESQNQQEKN | 646 |
| 18 | GCSGKLICTTTVPWNTSWSNKSLEQIWDNMTWMEWEREIDNYTGYIYQLIEESQNQQEKN | 644 |
| 19 | GCSGKLICTTTVPWNTSWSNKSLEQIWDNMTWMEWEREIDNYTGYIYQLIEESQNQQEKN | 646 |
| 20 | GCSGKLICTTTVPWNTSWSNKSLEQIWDNMTWMEWEREIDNYTGYIYQLIEESQNQQEKN | 646 |
| 21 | GCSGKLICTTTVPWNTSWSNKSLEQIWDNMTWMEWEREIDNYTGYIYQLIEESQNQQEKN | 646 |
| 22 | GCSGKLICTTTVPWNTSWSNKSLEQIWDNMTWMEWKREIDNYTGYIYQLIEESQNQQEKN | 646 |
| 23 | GCSGKLICTTTVPWNTSWSNKSLEQIWDNMTWMEWEREIDNYTGYIYQLIEESQNQQEKN | 646 |
| 24 | GCSGKLICTTTVPWNTSWSNKSLEQIWDNMTWMEWEREIDNYTGYIYQLIEESQNQQEKN | 647 |
| 25 | GCSGKLICTTTVPWNTSWSNKSLEQIWDNMTWMEWEREIDNYTGYIYQLIEESQNQQEKN | 646 |
| 26 | GCSGKLICTTTVPWNTSWSNKSLEQIWDNMTWMEWEREIDNYTGYIYQLIEESQNQQEKN | 646 |
| 27 | GCSGKLICTTTVPWNTSWSNKSLEQIWDNMTWMEWEREIDNYTGYIYQLIEESQNQQEKN | 646 |
| 28 | GCSGKLICTTTVPWNTSWSNKSLEQIWDNMTWMEWEREIDNYTGYIYQLIEESQNQQEKN | 646 |
| 29 | GCSGKLICTTTVPWNTSWSNKSLEQIWDNMTWMEWEREIDNYTGYIYQLIEESQNQQEKN | 646 |
| 30 | GCSGKLICTTTVPWNTSWSNKSLEQIWDNMTWMEWEREIDNYTGYIYQLIEESQNQQEKN | 646 |
| 31 | GCSGKLICTTTVPWNTSWSNKSLEQIWDNMTWMEWEREIDNYTGYIYQLIEESQNQQEKN | 646 |
| 32 | GCSGKLICTTTVPWNTSWSNKSLEQIWDNMTWMEWEREIDNYTGYIYQLIEESQNQQEKN | 646 |
| 33 | GCSGKLICTTTVPWNTSWSNKSLEQIWDNMTWMEWEREIDNYTGYIYQLIEESQNQQEKN | 646 |
| 34 | GCSGKLICTTTVPWNTSWSNKSLEQIWDNMTWMEWEREIDNYTGYIYQLIEESQNQQEKN | 646 |
| 35 | GCSGKLICTTTVPWNTSWSNKSLEQIWDNMTWMEWEREIDNYTGYIYQLIEESQNQQEKN | 646 |
| 36 | GCSGKLICTTTVPWNTSWSNKSLEQIWDNMTWMEWEREIDNYTGYIYQLIEESQNQQEKN | 646 |
| 37 | GCSGKLICTTTVPWNTSWSNKSLEQIWDNMTWMEWEREIDNYTGYIYQLIEESQNQQEKN | 646 |
| 38 | GCSGKLICTTTVPWNTSWSNKSLEQIWDNMTWMEWEREIDNYTGYIYQLIEESQNQQEKN | 646 |
| 39 | GCSGKLICTTTVPWNTSWSNKSLEQIWDNMTWMEWEREIDNYTGYIYQLIEESQNQQEKN | 646 |
| 40 | GCSGKLICTTTVPWNTSWSNKSLEQIWDNMTWMEWEREIDNYTGYIYQLIEESQNQQEKN | 646 |

| | Sequence | |
|---|---|---|
| | **EQELLALDKWASLWNWFDITNWLWYIKIFIMIVGGLIGLRIVFTVLSIVNRVRQGYSPLS** | |
| 1 | EQELLALDKWASLWNWFDITNWLWYIKIFIMIVGGLIGLRIVFTVLSIVNRVRQGYSPLS | 706 |
| 2 | EQELLALDKWASLWNWFDITNWLWYIKIFIMIVGGLIGLRIVFTVLSIVNRVRQGYSPLS | 705 |
| 3 | EQELLALDKWASLWNWFDITNWLWYIKIFIMIVGGLIGLRIVFTVLSIVNRVRQGYSPLS | 706 |
| 4 | EQELLALDKWASLWNWFDITNWLWYIKIFIMIVGGLIGLRIVFTVLSIVNRVRQGYSPLS | 706 |
| 5 | EQELLALDKWASLWNWFDITNWLWYIKIFIMIVGGLIGLRIVFTVLSIVNRVRQGYSPLS | 706 |
| 6 | EQELLALDKWASLWNWFDITNWLWYIKIFIMIVGGLIGLRIVFTVLSIVNRVRQGYSPLS | 707 |
| 7 | EQELLALDKWASLWNWFDITNWLWYIKIFIMIVGGLIGLRIVFTVLSIVNRVRQGYSPLS | 705 |
| 8 | EQELLALDKWASLWNWFDITNWLWYIKIFIMIVGGLIGLRIVFTVLSIVNRVRQGYSPLS | 706 |
| 9 | EQELLALDKWASLWNWFDITNWLWYIKIFIMIVGGLIGLRIVFTVLSIVNRVRQGYSPLS | 707 |
| 10 | EQELLALDKWASLWNWFDITNWLWYIKIFIMIVGGLIGLRIVFTVLSIVNRVRQGYSPLS | 706 |
| 11 | EQELLALDKWASLWNWFDITNWLWYIKIFIMIVGGLIGLRIVFTVLSIVNRVRQGYSPLS | 706 |
| 12 | EQELLALDKWASLWNWFDITNWLWYIKIFIMIVGGLIGLRIVFTVLSIVNRVRQGYSPLS | 706 |
| 13 | EQELLALDKWASLWNWFDITNWLWYIKIFIMIVGGLIGLRIVFTVLSIVNRVRQGYSPLS | 706 |
| 14 | EQELLALDKWASLWNWFDITNWLWYIKIFIMIVGGLIGLRIVFTVLSIVNRVRQGYSPLS | 706 |
| 15 | EQELLALDKWASLWNWFDITNWLWYIKIFIMIVGGLIGLRIVFTVLSIVNRVRQGYSPLS | 707 |
| 16 | EQELLALDKWASLWNWFDITNWLWYIKIFIMIVGGLIGLRIVFTVLSIVNRVRQGYSPLS | 706 |
| 17 | EQELLALDKWASLWNWFDITNWLWYIKIFIMIVGGLIGLRIVFTVLSIVNRVRQGYSPLS | 706 |
| 18 | EQELLALDKWASLWNWFDITNWLWYIKIFIMIVGGLIGLRIVFTVLSIVNRVRQGYSPLS | 704 |
| 19 | EQELLALDKWASLWNWFDITNWLWYIKIFIMIVGGLIGLRIVFTVLSIVNRVRQGYSPLS | 706 |
| 20 | EQELLALDKWASLWNWFDITNWLWYIKIFIMIVGGLIGLRIVFTVLSIVNRVRQGYSPLS | 706 |
| 21 | EQELLALDKWASLWNWFDITNWLWYIKIFIMIVGGLIGLRIVFTVLSIVNRVRQGYSPLS | 706 |
| 22 | EQELLALDKWASLWNWFDITNWLWYIKIFIMIVGGLIGLRIVFTVLSIVNRVRQGYSPLS | 706 |
| 23 | EQELLALDKWASLWNWFDITNWLWYIKIFIMIVGGLIGLRIVFTVLSIVNRVRQGYSPLS | 706 |
| 24 | EQELLALDKWASLWNWFDITNWLWYIKIFIMIVGGLIGLRIVFTVLSIVNRVRQGYSPLS | 707 |
| 25 | EQELLALDKWASLWNWFDITNWLWYIKIFIMIVGGLIGLRIVFTVLSIVNRVRQGYSPLS | 706 |
| 26 | EQELLALDKWASLWNWFDITNWLWYIKIFIMIVGGLIGLRIVFTVLSIVKRVRQGYSPLS | 706 |
| 27 | EQELLALDKWASLWNWFDITNWLWYIKIFIMIVGGLIGLRIVFTVLSIVNRVRQGYSPLS | 706 |
| 28 | EQELLALDKWASLWNWFDITNWLWYIKIFIMIVGGLIGLRIVFTVLSIVNRVRQGYSPLS | 706 |
| 29 | EQELLALDKWAGLWNWFDITNWLWYIKIFIMIVGGLIGLRIVFTVLSIVNRVRQGYSPLS | 706 |
| 30 | EQELLALDKWASLWNWFDITNWLWYIKIFIMIVGGLIGLRIVFTVLSIVNRVRQGYSPLS | 706 |
| 31 | EQELLALDKWASLWNWFDITNWLWYIKIFIMIVGGLIGLRIVFTVLSIVNRVRQGYSPLS | 706 |
| 32 | EQELLALDKWASLWNWFDITNWLWYIKIFIMIVGGLIGLRIVFTVLSIVNRVRQGYSPLS | 706 |
| 33 | EQELLALDKWASLWNWFDITNWLWYIKIFIMIVGGLIGLRIVFTVLSIVNRVRQGYSPLS | 706 |
| 34 | EQELLALDKWASLWNWFDITNWLWYIKIFIMIVGGLIGLRIVFTVLSIVNRVRQGYSPLS | 706 |
| 35 | EQELLALDKWASLWNWFDITNWLWYIKIFIMIVGGLIGLRIVFTVLSIVNRVRQGYSPLS | 706 |
| 36 | EQELLALDKWASLWNWFDITNWLWYIKIFIMIVGGLIGLRIVFTVLSIVNRVRQGYSPLS | 706 |
| 37 | EQELLALDKWASLWNWFDITNWLWYIKIFIMIIGGLIGLRIVFTVLSIVNRVRQGYSPLS | 706 |
| 38 | EQELLALDKWASLWNWFDITNWLWYIKIFIMIVGGLIGLRIVFTVLSIVNRVRQGYSPLS | 706 |
| 39 | EQELLALDKWASLWNWFDITNWLWYIKIFIMIVGGLIGLRIVFTVLSIVNRVRQGYSPLS | 706 |
| 40 | EQELLALDKWASLWNWFDITNWLWYIKIFIMIVGGLIGLRIVFTVLSIVNRVRQGYSPLS | 706 |

| | Sequence | |
|---|---|---|
| | **FQTHLPAQRGPDRPEGIGEEGGERDRDRSDPLVNGFLTLIWSDLRSLCLFSYHRLRDLLL** | |
| 1 | FQTHLPAQRGPDRPEGIGEEGGERDRDRSDPLVNGFLALIWSDLRSLCLFSYHRLRDLLL | 766 |
| 2 | FQTHLPAQRGPDRPEGIGEEGGERDRDRSDPLVNGFLTLIWSDLRSLCLFSYHRLRDLLL | 765 |
| 3 | FQTHLPAQRGPDRPEGIGEEGGERDRDRSDPLVNGFLTLIWSDLRSLCLFSYHRLRDLLL | 766 |
| 4 | FQTHLPAQRGPDRPEGIGEEGGERDRDRSDPLVNGFLTLIWSDLRSLCLFSYHRLRDLLL | 766 |
| 5 | FQTHLPAQRGPDRPEGIGEEGGERDRDRSDPLVNGFLTLIWSDLRSLCLFSYHRLRDLLL | 766 |
| 6 | FQTHLPAQRGPDRPEGIGEEGGERDRDRSDPLVNGFLTLIWSDLRSLCLFSYHRLRDLLL | 767 |
| 7 | FQTHLPAQRGPDRPEGIGEEGGERDRDRSDPLVNGFLTLIWSDLRSLCLFSYHRLRDLLL | 765 |
| 8 | FQTHLPAQRGPDRPEGIGEEGGERDRDRSDPLVNGFLALIWSELRSLCLFSYHRLRDLLL | 766 |
| 9 | FQTHLPAQRGPDRPEGIGEEGGERDRDRSDPLVNGFLTLIWSDLRSLCLFSYHRLRDLLL | 767 |
| 10 | FQTHLPAQRGPDRPEGIGEEGGERDRDRSDPLVNGFLTLIWSDLRSLCLFSYHRLRDLLL | 766 |
| 11 | FQTHLPAQRGPDRPEGIGEEGGERDRDRSDPLVNGFLTLIWSDLRSLCLFSYHRLRDLLL | 766 |
| 12 | FQTHLPAQRGPDRPEGIGEEGGERDRDRSDPLVNGFLTLIWSDLRSLCLFSYHRLRDLLS | 766 |
| 13 | FQTHLPAQRGPDRPEGIGEEGGERDRDRSDPLVNGFLTLIWSDLRSLCLFSYHRLRDLLL | 766 |
| 14 | FQTHLPAQRGPDRPEGTGEEGGERDRDRSDPLVNGFLTLIWSDLRSLCLFSYHRLRDLLL | 766 |
| 15 | FQTHLPAQRGPDRPEGIGEEGGERDRDRSDPLVNGFLTLIWSDLRSLCLFSYHRLRDLLL | 767 |
| 16 | FQTHLPAQRGPDRPEGIGEEGGERDRDRSDPLVNGFLTLIWSDLRSLCLFSYHRLRDLLL | 766 |
| 17 | FQTHLPAQRGPDRPEGIGEEGGERDRDRSDPLVNGFLTLIWSDLRSLCLFSYHRLRDLLL | 766 |
| 18 | FQTHLPAQRGPDRPEGIGEEGGERDRDRSDPLVNGFLTLIWSDLRSLCLFSYHRLRDLLL | 764 |
| 19 | FQTHLPAQRGPDRPEGIGEEGGERDRDRSDPLVNGFLTLIWSDLRSLCLFSYHRLRDLLL | 766 |
| 20 | FQTHLPAQRGPDRPEGIGEEGGERDRDRSDPLVNGFLTLIWSDLRSLCLFSYHRLRDLLL | 766 |
| 21 | FQTHLPAQRGPDRPEGIGEEGGERDRDRSDPLVNGFLTLIWSDLRSLCLFSYHRLRDLLL | 766 |
| 22 | FQTHLPAQRGPDRPEGIGEEGGERDRDRSDPLVNGFLTLIWSDLRSLCLFSYHRLRDLLL | 766 |
| 23 | FQTHLPAQRGPDRPEGIGEEGGERDRDRSDPLVNGFLTLIWSDLRSLCLFSYHRLRDLLL | 766 |
| 24 | FQTHLPAQRGPDRPEGIGEEGGERDRDRSDPLVNGFLTLIWSDLRSLCLFSYHRLRDLLL | 767 |
| 25 | FQTHLPAQRGPDRPEGIGEEGGERDRDRSDPLVNGFLTLIWSDLRSLCLFSYHRLRDLLL | 766 |
| 26 | FQTHLPAQRGPDRPEGIGEEGGARDRDRSDPLVNGFLALIWSDLRSLCLFSYHRLRDLLL | 766 |
| 27 | FQTHLPAQRGPDRPEGIGEEGGERDRDRSDPLVNGFLALIWSDLRSLCLFSYHRLRDLLL | 766 |
| 28 | FQTHLPAQRGPDRPEGIGEEGGERDRDRSDPLVNGFLTLIWSDLRSLCLFSYHRLRDLLL | 766 |
| 29 | FQTHLPAQRGPDRPEGIGEEGGERDRDRSDPLVNGFLTLIWSDLRSLCLFSYHRLRDLLL | 766 |
| 30 | FQTHLPAQRGPDRPEGIGEEGGERDRDRSDPLVNGFLTLIWSDLRSLCLFSYHRLRDLLL | 766 |
| 31 | FQTHLPAQRGPDRPEGIGEEGGERDRDRSDPLVNGFLTLIWSDLRSLCLFSYHRLRDLLL | 766 |
| 32 | FQTHLPAQRGPDRPEGIGEEGGERDRDRSDPLVNGFLTLIWSDLRSLCLFSYHRLRDLLL | 766 |
| 33 | FQTHLPAQRGPDRPEGIGEEGGERDRDRSDPLVNGFLTLIWSDLRSLCLFSYHRLRDLLL | 766 |
| 34 | FQTHLPAQRGPDRPEGIGEEGGERDRDRSDPLVNGFLTLIWSDLRSLCLFSYHRLRDLLL | 766 |
| 35 | FQTHLPAQRGPDRPEGIGEEGGERDRDRSDPLVNGFLTLIWSDLRSLCLFSYHRLRDLLL | 766 |
| 36 | FQTHLPAQRGPDRPEGIGEEGGERDRDRSDPLVNGFLTLIWSDLRSLCLFSYHRLRDLLL | 766 |
| 37 | FQTHLPAQRGPDRPEGIGEEGGERDRDRSDPLVNGFLTLIWSDLRSLCLFSYHRLRDLLL | 766 |
| 38 | FQTHLPAQRGPDRPEGIGEEGGERDRDRSDPLVNGFLTLIWSDLRSLCLFSYHRLRDLLL | 766 |
| 39 | FQTHLPAQRGPDRPEGIGEEGGERDRDRSDPLVNGFLTLIWSDLRSLCLFSYHRLRDLLL | 766 |
| 40 | FQTHLPAQRGPDRPEGIGEEGGERDRDRSDPLVNGFLTLIWSDLRSLCLFSYHRLRDLLL | 766 |

|  | IVTRIVELLGR-------RGWEVLKYWWNLLQYWSQELKNSAVSLLNATATAVAEGTDRV |  |
|---|---|---|
| 1 | IVTRIVELLGR-------RGWEVLKYWWNLLQYWSQELKNSAVSLLNATAIAVAEGTDRV | 819 |
| 2 | IVTRIVELLGR-------RGWEVLKYWWNLLQYWSQELKNSAVSLLNATATAVAEGTDRV | 818 |
| 3 | IVTRIVELLGR-------RGWEVLKYWWNLLQYWSQELKNSAVSLLNATATAVAEGTDRV | 819 |
| 4 | IVTRIVELLGR-------RGWEVLKYWWNLLQYWSQELKNSAVSLLNATATAVAEGTDRV | 819 |
| 5 | IVTRIVELLGR-------RGWEVLKYWWNLLQYWSQELKNSAVSLLNATATAVAEGTDRV | 819 |
| 6 | IVTRIVELLGR-------RGWEVLKYWWNLLQYWSQELKNSAVSLLNATAIAVAEGTDRV | 820 |
| 7 | IVTRIVELLGR-------RGWEVLKYWWNLLQYWSQELKNSAVSLLNATATAVAEGTDRV | 818 |
| 8 | IVTRIVELLGR-------RGWEVLKYWWNLLQYWSQELKNSAVSLLNATAIAVAEGTDRV | 819 |
| 9 | IVTRIVELLGR-------RGWEVLKYWWNLLQYWSQELKNSAVSLLNATATAVAEGTDRV | 820 |
| 10 | IVTRIVELLGR-------RGWEVLKYWWNLLQYWSQELKNSAVSLLNATATAVAEGTDRV | 819 |
| 11 | IVTRIVELLGR-------RGWEVLKYWWNLLQYWSQELKNSAVSLLNATATAVAEGTDRV | 819 |
| 12 | IVTRIVELLGR-------RGWEVLKYWWNLLQYWSQELKNSAVSLLNATAIAVAEGTDRV | 819 |
| 13 | IVTRIVELLGR-------RGWEVLKYWWNLLQYWSQELKNSAVSLLNATAIAVAEGTDRV | 819 |
| 14 | IVTRIVELLGR-------RGWEVLKYWWNLLQYWSQELKNSAVSLLNATATAVAEGTDRV | 819 |
| 15 | IVTRIVELLGR-------RGWEVLKYWWNLLQYWSQELKNSAVSLLNATAIAVAEGTDRV | 820 |
| 16 | IVTRIVELLGR-------RGWEVLKYWWNLLQYWSQELKNSAVSLLNATATAVAEGTDRV | 819 |
| 17 | IVTRIVELLGR-------RGWEVLKYWWNLLQYWSQELKNSAVSLLNATATAVAEGTDRV | 819 |
| 18 | IVTRIVELLGR-------RGWEVLKYWWNLLQYWSQELKNSAVSLLNATATAVAEGTDRV | 817 |
| 19 | IVTRIVELLGR-------RGWEVLKYWWNLLQYWSQELKNSAVSLLNATATAVAEGTDRV | 819 |
| 20 | IVTRIVELLGR-------RGWEVLKYWWNLLQYWSQELKNSAVSLLNATATAVAEGTDRV | 819 |
| 21 | IVTRIVELLGR-------RGWEVLKYWWNLLQYWSQELKNSAVSLLNATATAVAEGTDRV | 819 |
| 22 | IVTRIVELLGR-------RGWEVLKYWWNLLQYWSQELKNSAVSLLNATATAVAEGTDRV | 819 |
| 23 | IVTRIVELLGR-------RGWEVLKYWWNLLQYWSQELKNSAVSLLNATATAVAEGTDRV | 819 |
| 24 | IVTRIVELLGR-------RGWEVLKYWWNLLQYWSQELKNSAVSLLNATAIAVAEGTDRV | 820 |
| 25 | IVTRIVELLGR-------RGWEVLKYWWNLLQYWSQELKNSAVSLLNATATAVAEGTDRV | 819 |
| 26 | IVTRIVELLGR-------RGWEVLKYWWNLLQYWSQELKNSAVSLLNATAIAVAEGTDRV | 819 |
| 27 | IVTRIVELLGR-------RGWEVLKYWWNLLQYWSQELKNSAVSLLNATAIAVAEGTDRV | 819 |
| 28 | IVTRIVELLGR-------RGWEVLKYWWNLLQYWSQELKNSAVSLLNATATAVAEGTDRV | 819 |
| 29 | IVTRIVELLGR-------RGWEVLKYWWNLLQYWSQELKNSAVSLLNATATAVAEGTDRV | 819 |
| 30 | IVTRIVELLGR-------RGWEVLKYWWNLLQYWSQELKNSAVSLLNATAIAVAEGTDRV | 819 |
| 31 | IVTRIVELLGR-------RGWEVLKYWWNLLQYWSQELKNSAVSLLNATATAVAEGTDRV | 819 |
| 32 | IVTRIVELLGR-------RGWEVLKYWWNLLQYWSQELKNSAVSLLNATATAVAEGTDRV | 819 |
| 33 | IVTRIVELLGR-------RGWEVLKYWWNLLQYWSQELKNSAVSLLNATATAVAEGTDRV | 819 |
| 34 | IVTRIVELLGR-------RGWEVLKYWWNLLQYWSQELKNSAVSLLNATATAVAEGTDRV | 819 |
| 35 | IVTRIVELLGR-------RGWEVLKYWWNLLQYWSQELKNSAVSLLNATAIAVAEGTDRV | 819 |
| 36 | IVTRIVELLGR-------RGWEVLKYWWNLLQYWSQELKNSAVSLLNATAIAVAEGTDRV | 819 |
| 37 | IVTRIVELLGR-------RGWEVLKYWWNLLQYWSQELKNSAVSLLNATATAVAEGTDRV | 819 |
| 38 | IVTRIVELLGR-------RGWEVLKYWWNLLQYWSQELKNSAVSLLNATATAVAEGTDRV | 819 |
| 39 | IVTRIVELLGR-------RGWEVLKYWWNLLQYWSQELKNSAVSLLNATATAVAEGTDRV | 819 |
| 40 | IVTRIVELLGR-------RGWEVLKYWWNLLQYWSQELKNSAVSLLNATATAVAEGTDRV | 819 |

**IEVVQRACRAILHIPRRIRQGLERALL**

| | | |
|---|---|---|
| 1 | IEVVQRACRAILHIPRRIRQGLERALL | 846 |
| 2 | IEVVQRACRAILHIPRRIRQGLERALL | 845 |
| 3 | IEVVQRACRAILHIPRRIRQGLERALL | 846 |
| 4 | IEVVQRACRAILHIPRRIRQGLERALL | 846 |
| 5 | IEVVQRACRAILHIPRRIRQGLERALL | 846 |
| 6 | IEVVQGACRAILHIPRRIRQGLERALL | 847 |
| 7 | IEVVQRACRAILHIPRRIRQGLERALL | 845 |
| 8 | IEVVQRACRAILHIPRRIRQGLERALL | 846 |
| 9 | IEVVQRACRAILHIPRRIRQGLERALL | 847 |
| 10 | IEVVQRACRAILHIPRRIRQGLERALL | 846 |
| 11 | IEVVQRACRAILHIPRRIRQGLERALL | 846 |
| 12 | IEVVQRACRAILHIPRRIRQGLERALL | 846 |
| 13 | IEVVQRACRAILHIPRRIRQGLERALL | 846 |
| 14 | IEVVQRACRAILHIPRRIRQGLERALL | 846 |
| 15 | IEVVQGACRAILHIPRRIRQGLERALL | 847 |
| 16 | IEVVQRACRAILHIPRRIRQGLERALL | 846 |
| 17 | IEVVQRACRAILHIPRRIRQGLERALL | 846 |
| 18 | IEVVQRACRAILHIPRRIRQGLERALL | 844 |
| 19 | IEVVQRACRAILHIPRRIRQGLERALL | 846 |
| 20 | IEVVQRACRAILHIPRRIRQGLERALL | 846 |
| 21 | IEVVQRACRAILHIPRRIRQGLERALL | 846 |
| 22 | IEVVQRACRAILHIPRRIRQGLERALL | 846 |
| 23 | IEVVQRACRAILHIPRRIRQGLERALL | 846 |
| 24 | IEVVQGACRAILHIPRRIRQGLERALL | 847 |
| 25 | IEVVQRACRAILHIPRRIRQGLERALL | 846 |
| 26 | IEVVQRACRAILHIPRRIRQGLERALL | 846 |
| 27 | IEVVQRACRAILHIPRRIRQGLERALL | 846 |
| 28 | IEVVQRACRAILHIPRRIRQGLERALL | 846 |
| 29 | IEVVQRACRAILHIPRRIRQGLERALL | 846 |
| 30 | IEVVQRACRAIRHIPRRIRQGLERALL | 846 |
| 31 | IEVVQRACRAILHIPRRIRQGLERALL | 846 |
| 32 | IEVVQRACRAILHIPRRIRQGLERALL | 846 |
| 33 | IEVVQRACRAILHIPRRIRQGLERALL | 846 |
| 34 | IEVVQRACRAILHIPRRIRQGLERALL | 846 |
| 35 | IEVVQRACRAIPHIPRRIRQGLERALL | 846 |
| 36 | IEVVQRVGRAILHIPRRIRQGLERALL | 846 |
| 37 | IEVVQRACRAILHIPRRIRQGLERALL | 846 |
| 38 | IEVVQRACRAILHIPRRIRQGLERALL | 846 |
| 39 | IEVVQRACRAILHIPRRIRQGLERALL | 846 |
| 40 | IEVVQRACRAILHIPRRIRQGLERALL | 846 |

### Aligned using an external algorithm

**Consensus**

1. B.US.2006.700010040_C9_4520.EU289193
2. B.US.2007.CH0040_3_d0428_ipe032_10_17.MG900271
3. B.US.2007.CH0040_3_d0428_ipe032_10_19.MG900272
4. B.US.2007.CH0040_3_d0428_ipe032_10_20.MG900273
5. B.US.2007.CH0040_3_d0428_ipe032_10_22.MG900274
6. B.US.2007.CH0040_3_d0428_ipe032_10_23.MG900275
7. B.US.2007.CH0040_3_d0428_ipe032_10_25.MG900276
8. B.US.2007.CH0040_3_d0428_ipe032_10_26.MG900277
9. B.US.2007.CH0040_3_d0428_ipe032_10_27.MG900278
10. B.US.2007.CH0040_3_d0428_ipe032_10_28.MG900279
11. B.US.2007.CH0040_3_d0428_ipe032_10_29.MG900280
12. B.US.2007.CH0040_3_d0428_ipe032_10_32.MG900281
13. B.US.2007.CH0040_3_d0428_ipe032_10_33.MG900282
14. B.US.2007.CH0040_3_d0428_ipe032_10_34.MG900283
15. B.US.2007.CH0040_3_d0428_ipe032_10_35.MG900284
16. B.US.2007.CH0040_3_d0428_ipe032_10_36.MG900285
17. B.US.2007.CH0040_3_d0428_ipe032_10_37.MG900286
18. B.US.2007.CH0040_3_d0428_ipe032_10_39.MG900287
19. B.US.2007.CH0040_3_d0428_ipe032_10_40.MG900288
20. B.US.2007.CH0040_3_d0428_ipe032_10_41.MG900289
21. B.US.2007.CH0040_3_d0428_ipe032_10_45.MG900290
22. B.US.2007.CH0040_3_d0428_ipe032_10_46.MG900291
23. B.US.2007.CH0040_3_d0428_ipe032_10_48.MG900292
24. B.US.2007.CH0040_3_d0428_ipe032_10_49.MG900293
25. B.US.2007.CH0040_3_d0428_ipe032_10_50.MG900294
26. B.US.2007.CH0040_3_d0428_ipe032_27_15.MG900295
27. B.US.2007.CH0040_3_d0428_ipe032_27_16.MG900296
28. B.US.2007.CH0040_3_d0428_ipe032_2_01.MG900297
29. B.US.2007.CH0040_3_d0428_ipe032_2_05.MG900298
30. B.US.2007.CH0040_3_d0428_ipe032_3_01.MG900299
31. B.US.2007.CH0040_3_d0428_ipe032_3_06.MG900300
32. B.US.2007.CH0040_3_d0428_ipe032_3_07.MG900301
33. B.US.2007.CH0040_3_d0428_ipe032_3_08.MG900302
34. B.US.2007.CH0040_3_d0428_ipe032_3_10.MG900303
35. B.US.2007.CH0040_3_d0428_ipe032_3_11.MG900304
36. B.US.2007.CH0040_3_d0428_ipe032_3_12.MG900305
37. B.US.2007.CH0040_3_d0428_ipe032_3_13.MG900306
38. B.US.2007.CH0040_3_d0428_ipe032_3_14.MG900307

**MRVMGIRKNYQHLWREGILLLGILMICSAADNLWVTVYYGVPVWREATTTLFCASDAKAY**

| | | |
|---|---|---|
| 1 | MRVMGIRKNYQHLWREGILLLGILMICSAADNLWVTVYYGVPVWREATTTLFCASDAKAY | 60 |
| 2 | MRVMGIRKNYQHLWREGILLLGILMICSAADNLWVTVYYGVPVWREATTTLFCASDAKAY | 60 |
| 3 | MRVMGIRKNYQHLWREGILLLGILMICSAADNLWVTVYYGVPVWREATTTLFCASDAKAY | 60 |
| 4 | MRVMGIRKNYQHLWREGILLLGILMICSVADNLWVTVYYGVPVWREATTTLFCASDAKAY | 60 |
| 5 | MRVMGIRKNYQHLWREGILLLGILMICSAADNLWVTVYYGVPVWREATTTLFCASDAKAY | 60 |
| 6 | MRVMGIRKNYQHLWREGILLLGILMICSAADNLWVTVYYGVPVWREATTTLFCASDAKAY | 60 |
| 7 | MRVMGIRKNYQHLWREGILLLGILMICSAADNLWVTVYYGVPVWREATTTLFCASDAKAY | 60 |
| 8 | MRVMGIRKNYQHLWREGILLLGILMICSAADNLWVTVYYGVPVWREATTTLFCASDAKAY | 60 |
| 9 | MRVMGIRKNYQHLWREGILLLGILMICSAADNLWVTVYYGVPVWREATTTLFCASDAKAY | 60 |
| 10 | MRVMGIRKNYQHLWREGILLLGILMICSAADNLWVTVYYGVPVWREATTTLFCASDAKAY | 60 |
| 11 | MRVMGIRKNYQHLWREGILLLGILMICSAADNLWVTVYYGVPVWREATTTLFCASDAKAY | 60 |
| 12 | MRVMGIRKNYQHLWREGILLLGILMICSAADNLWVTVYYGVPVWREATTTLFCASDAKAY | 60 |
| 13 | MRVMGIRKNYQHLWREGILLLGILMICSAADNLWVTVYYGVPVWREATTTLFCASDAKAY | 60 |
| 14 | MRVMGIRKNYQHLWREGILLLGILMICSAADNLWVTVYYGVPVWREATTTLFCASDAKAY | 60 |
| 15 | MRVMGIRKNYQHLWREGILLLGILMICSAADNLWVTVYYGVPVWREATTTLFCASDAKAY | 60 |
| 16 | MRVMGIRKNYQHLWREGILLLGILMICSAADNLWVTVYYGVPVWREATTTLFCASDAKAY | 60 |
| 17 | MRVMGIRKNYQHLWREGILLLGILMICSAADNLWVTVYYGVPVWREATTTLFCASDAKAY | 60 |
| 18 | MRVMGIRKNYQHLWREGILLLGILMICSAADNLWVTVYYGVPVWREATTTLFCASDAKAY | 60 |
| 19 | MRVMGIRKNYQHLWREGILLLGILMICSAADNLWVTVYYGVPVWREATTTLFCASDAKAY | 60 |
| 20 | MRVMGIRKNYQHLWREGILLLGILMICSAADNLWVTVYYGVPVWREATTTLFCASDAKAY | 60 |
| 21 | MRVMGIRKNYQHLWREGILLLGILMICSAADNLWVTVYYGVPVWREATTTLFCASDAKAY | 60 |
| 22 | MRVMGIRKNYQHLWREGILLLGILMICSAADNLWVTVYYGVPVWREATTTLFCASDAKAY | 60 |
| 23 | MRVMGIRKNYQHLWREGILLLGILMICSAADNLWVTVYYGVPVWREATTTLFCASDAKAY | 60 |
| 24 | MRVMGIRKNYQHLWREGILLLGILMICSAADNLWVTVYYGVPVWREATTTLFCASDAKAY | 60 |
| 25 | MRVMGIRKNYQHLWREGILLLGILMICSAADNLWVTVYYGVPVWREATTTLFCASDAKAY | 60 |
| 26 | MRVMGIRKNYQHLWREGILLLGILMICSAADNLWVTVYYGVPVWREATTTLFCASDAKAY | 60 |
| 27 | MRVMGIRKNYQHLWREGILLLGILMICSAADNLWVTVYYGVPVWREATTTLFCASDAKAY | 60 |
| 28 | MRVMGIRKNYQHLWREGILLLGILMICSAADNLWVTVYYGVPVWREATTTLFCASDAKAY | 60 |
| 29 | MRVMGIRKNYQHLWREGILLLGILMICSAADNLWVTVYYGVPVWREATTTLFCASDAKAY | 60 |
| 30 | MRVMGIRKNYQHLWREGILLLGILMICSAADNLWVTVYYGVPVWREATTTLFCASDAKAY | 60 |
| 31 | MRVMGIRKNYQHLWREGILLLGILMICSAADNLWVTVYYGVPVWREATTTLFCASDAKAY | 60 |
| 32 | MRVMGIRKNYQHLWREGILLLGILMICSAADNLWVTVYYGVPVWREATTTLFCASDAKAY | 60 |
| 33 | MRVMGIRKNYQHLWREGILLLGILMICSAADNLWVTVYYGVPVWREATTTLFCASDAKAY | 60 |
| 34 | MRVMGIRKNYQHLWREGILLLGILMICSAADNLWVTVYYGVPVWREATTTLFCASDAKAY | 60 |
| 35 | MRVMGIRKNYQHLWREGILLLGILMICSAADNLWVTVYYGVPVWREATTTLFCASDAKAY | 60 |
| 36 | MRVMGIRKNYQHLWREGILLLGILMICSAADNLWVTVYYGVPVWREATTTLFCASDAKAY | 60 |
| 37 | MRVMGIRKNYQHLWREGILLLGILMICSAADNLWVTVYYGVPVWREATTTLFCASDAKAY | 60 |
| 38 | MRVMGIRKNYQHLWREGILLLGILMICSAADNLWVTVYYGVPVWREATTTLFCASDAKAY | 60 |

| | Sequence | |
|---|---|---|
| | **DTEAHNVWATHACVPTDPNPQEVELKNVTENFNMWENNMVEQMHEDIISLWDQSLKPCVK** | |
| 1 | DTEAHNVWATHACVPTDPNPQEVELKNVTENFNMWENNMVEQMHEDIISLWDQSLKPCVK | 120 |
| 2 | DTEAHNVWATHACVPTDPNPQEVELKNVTENFNMWENNMVEQMHEDIISLWDQSLKPCVK | 120 |
| 3 | DTEAHNVWATHACVPTDPNPQEVELKNVTENFNMWENNMVEQMHEDIISLWDQSLKPCVK | 120 |
| 4 | DTEAHNVWATHACVPTDPNPQEVELKNVTENFNMWENNMVEQMHEDIISLWDQSLKPCVK | 120 |
| 5 | DTEAHNVWATHACVPTDPNPQEVELKNVTENFNMWENNMVEQMHEDIISLWDQSLKPCVK | 120 |
| 6 | DTEAHNVWATHACVPTDPNPQEVELKNVTENFNMWENNMVEQMHEDIISLWDQSLKPCVK | 120 |
| 7 | DTEAHNVWATHACVPTDPNPQEVELKNVTENFNMWENNMVEQMHEDIISLWDQSLKPCVK | 120 |
| 8 | DTEAHNVWATHACVPTDPNPQEVELKNVTENFNMWENNMVEQMHEDIISLWDQSLKPCVK | 120 |
| 9 | DTEAHNVWATHACVPTDPNPQEVELKNVTENFNMWENNMVEQMHEDIISLWDQSLKPCVK | 120 |
| 10 | DTEAHNVWATHACVPTDPNPQEVELKNVTENFNMWENNMVEQMHEDIISLWDQSLKPCVK | 120 |
| 11 | DTEAHNVWATHACVPTDPNPQEVELKNVTENFNMWENNMVEQMHEDIISLWDQSLKPCVK | 120 |
| 12 | DTEAHNVWATHACVPTDPNPQEVELKNVTENFNMWENNMVEQMHEDIISLWDQSLKPCVK | 120 |
| 13 | DTEAHNVWATHACVPTDPNPQEVELKNVTENFNMWENNMVEQMHEDIISLWDQSLKPCVK | 120 |
| 14 | DTEAHNVWATHACVPTDPNPQEVELKNVTENFNMWENNMVEQMHEDIISLWDQSLKPCVK | 120 |
| 15 | DTEAHNVWATHACVPTDPNPQEVELKNVTENFNMWENNMVEQMHEDIISLWDQSLKPCVK | 120 |
| 16 | DTEAHNVWATHACVPTDPNPQEVELKNVTENFNMWENNMVEQMHEDIISLWDQSLKPCVK | 120 |
| 17 | DTEAHNVWATHACVPTDPNPQEVELKNVTENFNMWENNMVEQMHEDIISLWDQSLKPCVK | 120 |
| 18 | DTEAHNVWATHACVPTDPNPQEVELKNVTENFNMWENNMVEQMHEDIISLWDQSLKPCVK | 120 |
| 19 | DTEAHNVWATHACVPTDPNPQEVELKNVTENFNMWENNMVEQMHEDIISLWDQSLKPCVK | 120 |
| 20 | DTEAHNVWATHACVPTDPNPQEVELKNVTENFNMWENNMVEQMHEDIISLWDQSLKPCVK | 120 |
| 21 | DTEAHNVWATHACVPTDPNPQEVELKNVTENFNMWENNMVEQMHEDIISLWDQSLKPCVK | 120 |
| 22 | DTEAHNVWATHACVPTDPNPQEVELKNVTENFNMWENNMVEQMHEDIISLWDQSLKPCVK | 120 |
| 23 | DTEAHNVWATHACVPTDPNPQEVELKNVTENFNMWENNMVEQMHEDIISLWDQSLKPCVK | 120 |
| 24 | DTEAHNVWATHACVPTDPNPQEVELKNVTENFNMWENNMVEQMHEDIISLWDQSLKPCVK | 120 |
| 25 | DTEAHNVWATHACVPTDPNPQEVELKNVTENFNMWENNMVEQMHDDIISLWDQSLKPCVK | 120 |
| 26 | DTEAHNVWATHACVPTDPNPQEVELKNVTENFNMWENNMVEQMHEDIISLWDQSLKPCVK | 120 |
| 27 | DTEAHNVWATHACVPTDPNPQEVELKNVTENFNMWENNMVEQMHEDIISLWDQSLKPCVK | 120 |
| 28 | DTEAHNVWATHACVPTDPNPQEVELKNVTENFNMWENNMVEQMHEDIISLWDQSLKPCVK | 120 |
| 29 | DTEAHNVWATHACVPTDPNPQEVELKNVTENFNMWENNMVEQMHEDIISLWDQSLKPCVK | 120 |
| 30 | DTEAHNVWATHACVPTDPNPQEVELKNVTENFNMWENNMVEQMHEDIISLWDQSLKPCVK | 120 |
| 31 | DTEAHNVWATHACVPTDPNPQEVELKNVTENFNMWKNNMVEQMHEDIISLWDQSLKPCVK | 120 |
| 32 | DTEAHNVWATHACVPTDPNPQEVELKNVTENFNMWENNMVEQMHEDIISLWDQSLKPCVK | 120 |
| 33 | DTEAHNVWATHACVPTDPNPQEVELKNVTENFNMWENNMVEQMHEDIISLWDQSLKPCVK | 120 |
| 34 | DTEAHNVWATHACVPTDPNPQEVELKNVTENFNMWENNMVEQMHEDIISLWDQSLKPCVK | 120 |
| 35 | DTEAHNVWATHACVPTDPNPQEVELKNVTENFNMWENNMVEQMHEDIISLWDQSLKPCVK | 120 |
| 36 | DTEAHNVWATHACVPTDPNPQEVELKNVTENFNMWENNMVEQMHEDIISLWDQSLKPCVK | 120 |
| 37 | DTEAHNVWATHACVPTDPNPQEVELKNVTENFNMWENNMVEQMHEDIISLWDQSLKPCVK | 120 |
| 38 | DTEAHNVWATHACVPTDPNPQEVELKNVTENFNMWENNMVEQMHEDIISLWDQSLKPCVK | 120 |

| | | |
|---|---|---|
| | LTPLCVTLNCTDLGNVTTTT---NSNGTLMEKGEVKNCSFKITTDIKDRTRKEYALFYKL | |
| 1 | LTPLCVTLNCTDLGNVTNTT---NSNGEMMEKGEVKNCSFKITTDIKDRTRKEYALFYKL | 177 |
| 2 | LTPLCVTLNCTDLGNVTTTT---NSNGTLMEKGEVKNCSFKITTDIKDRTRKEYALFYKL | 177 |
| 3 | LTPLCVTLNCTDLGNVTTTT---NSNGTLMEKGEVKNCSFKITTDIKDRTRKEYALFYKL | 177 |
| 4 | LTPLCVTLNCTDLGNVTTTT---NSNGTLMEKGEVKNCSFKITTDIKDRTRKEYALFYKL | 177 |
| 5 | LTPLCVTLNCTDLGNVTTTT---NSNGTLMEKGEVKNCSFKITTDIKDRTRKEYALFYKL | 177 |
| 6 | LTPLCVTLNCTDLGNVTNTT---NSNGGMMEKGEVKNCSFKITTDIKDRTRKEYALFYKL | 177 |
| 7 | LTPLCVTLNCTDLGNVTTTT---NSNGTLMEKGEVKNCSFKITTDIKDRTRKEYALFYKL | 177 |
| 8 | LTPLCVTLNCTDLGNVTTTT---NSNGTLMEKGEVKNCSFKITTDIKDRTRKEYALFYKL | 177 |
| 9 | LTPLCVTLNCTDLGNVTTTT---NSNGTLMEKGEVKNCSFKITTDIKDRTRKEYALFYKL | 177 |
| 10 | LTPLCVTLNCTDLGNVTTTT---NSNGTLMEKGEVKNCSFKITTDIKDRTRKEYALFYKL | 177 |
| 11 | LTPLCVTLNCTDLGNVTTTT---NSNGTLMEKGEVKNCSFKITTDIKDRTRKEYALFYKL | 177 |
| 12 | LTPLCVTLNCTDLGNVTTTT---NSNGTLMEKGEVKNCSFKITTDIKDRTRKEYALFYKL | 177 |
| 13 | LTPLCVTLNCTDLENVTNTT---NSNETMMEKGEVKNCSFKITTDIKDRTRKEYALFYKL | 177 |
| 14 | LTPLCVTLNCTDLGNVTTTT---NSNGTLMEKGEVKNCSFKITTDIKDRTRKEYALFYKL | 177 |
| 15 | LTPLCVTLNCTDLGNVTTTT---NSNGTLMEKGEVKNCSFKITTDIKDRTRKEYALFYKL | 177 |
| 16 | LTPLCVTLNCTDLGNVTTTT---NSNGTLMEKGEVKNCSFKITTDIKDRTRKEYALFYKL | 177 |
| 17 | LTPLCVTLNCTDLGNVTTTT---NSNGTLMEKGEVKNCSFKITTDIKDRTRKEYALFYKL | 177 |
| 18 | LTPLCVTLNCTDLGNVTNTT---NSNGGMMEKGEVKNCSFKITTDIKDRTRKEYALFYKL | 177 |
| 19 | LTPLCVTLNCTDLGNVTTTT---NSNGTLMEKGEVKNCSFKITTDIKDRTRKEYALFYKL | 177 |
| 20 | LTPLCVTLNCTDLGNVTTTT---NSNGTLMEKGEVKNCSFKITTDIKDRTRKEYALFYKL | 177 |
| 21 | LTPLCVTLNCTDLGNVTTTT---NSNGTLMEKGEVKNCSFKITTDIKDRTRKEYALFYKL | 177 |
| 22 | LTPLCVTLNCTDLGNVTNTT---NSNGGMMEKGEVKNCSFKITTDIKDRTRKEYALFYKL | 177 |
| 23 | LTPLCVTLNCTDLGNVTTTT---NSNGTLMEKGEVKNCSFKITTDIKDRTRKEYALFYKL | 177 |
| 24 | LTPLCVTLNCTDLGNVTTTT---NSNGTLMEKGEVKNCSFKITTDIKDRTRKEYALFYKL | 177 |
| 25 | LTPLCVTLNCTDLGNVTTTT---NSNGTLMEKGEVKNCSFKITTDIKDRTRKEYALFYKL | 177 |
| 26 | LTPLCVTLNCTDLGNVTTTT---NSNGTLMEKGEVKNCSFKIATDIKDRTRKEYALFYKL | 177 |
| 27 | LTPLCVTLNCTDLGNVTTTT---NSNGTLMEKGEVKNCSFKITTDIKDRTRKEYALFYKL | 177 |
| 28 | LTPLCVTLNCTDLGNVTTTT---NSNGTLMEKGEVKNCSFKITTDIKDRTRKEYALFYKL | 177 |
| 29 | LTPLCVTLNCTDLGNVTTTT---NSNGTLMEKGEVKNCSFKITTDIKDRTRKEYALFYKL | 177 |
| 30 | LTPLCVTLNCTDLGNVTTTT---NSNGTLMEKGEVKNCSFKITTDIKDRTRKEYALFYKL | 177 |
| 31 | LTPLCVTLNCTDLGNVTNTT---NSNGGMMEKGEVKNCSFKITTDIKDRTRKEYALFYKL | 177 |
| 32 | LTPLCVTLNCTDLGNVTTTT---NSNGTLMEKGEVKNCSFKITTDIKDRTRKEYALFYKL | 177 |
| 33 | LTPLCVTLNCTDLGNVTTTT---NSNGTLMEKGEVKNCSFKITTDIKDRTRKEYALFYKL | 177 |
| 34 | LTPLCVTLNCTDLGNVTTTT---NSNGTLMEKGEVKNCSFKITTDIKDRTRKEYALFYKL | 177 |
| 35 | LTPLCVTLNCTDLGNVTNTT---NSNGGMMEKGEVKNCSFKITTDIKDRTRKEYALFYKL | 177 |
| 36 | LTPLCVTLNCTDLGNVTTTT---NSNGTLMEKGEVKNCSFKITTDIKDRTRKEYALFYKL | 177 |
| 37 | LTPLCVTLNCTDLGNVTTTT---NSNGTLMEKGEVKNCSFKITTDIKDRTRKEYALFYKL | 177 |
| 38 | LTPLCVTLNCTDLGNVTTTT---NSNGTLMEKGEVKNCSFKITTDIKDRMRKEYALFYKL | 177 |

| | | |
|---|---|---|
|  | DVVPIND-------TRYRLVSCNTSVITQACPKVSFEPIPIHYCAPAGFAILKCNDKQFI |  |
| 1 | DVVPIND-------TRYRLVSCNTSVITQACPKVSFEPIPIHYCAPAGFAILKCNDKQFI | 230 |
| 2 | DVVPIND-------TRYRLVSCNTSVITQACPKVSFEPIPIHYCAPAGFAILKCNDKQFI | 230 |
| 3 | DVVPIND-------TRYRLVSCNTSVITQACPKVSFEPIPIHYCAPAGFAILKCNDKQFM | 230 |
| 4 | DVVPIND-------TRYRLVSCNTSVITQACPKVSFEPIPIHYCAPAGFAILKCNDKQFM | 230 |
| 5 | DVVPIND-------TRYRLVSCNTSVITQACPKVSFEPIPIHYCAPAGFAILKCNDKQFI | 230 |
| 6 | DVVPIND-------TRYRLVSCNTSVITQACPKVSFEPIPIHYCAPAGFAILKCNDKQFI | 230 |
| 7 | DVVPIND-------TRYRLVSCNTSVITQACPKVSFEPIPIHYCAPAGFAILKCNDKQFM | 230 |
| 8 | DVVPIND-------TRYRLVSCNTSVITQACPKVSFEPIPIHYCAPAGFAILKCNDKQFI | 230 |
| 9 | DVVPIND-------TRYRLVSCNTSVITQACPKVSFEPIPIHYCAPAGFAILKCNDKQFM | 230 |
| 10 | DVVPIND-------TRYRLVSCNTSVITQACPKVSFEPIPIHYCAPAGFAILKCNDKQFM | 230 |
| 11 | DVVPIND-------TRYRLVSCNTSVITQACPKVSFEPIPIHYCAPAGFAILKCNDKQFM | 230 |
| 12 | DVVPIND-------TRYRLVSCNTSVITQACPKVSFEPIPIHYCAPAGFAILKCNDKQFI | 230 |
| 13 | DVVPIND-------TRYRLVSCNTSVITQACPKVSFEPIPIHYCAPAGFAILKCNDKQFI | 230 |
| 14 | DVVPIND-------TRYRLVSCNTSVITQACPKVSFEPIPIHYCAPAGFAILKCNDKQFI | 230 |
| 15 | DVVPIND-------TRYRLVSCNTSVIIQACPKVSFEPIPIHYCAPAGFAILKCNDKQFI | 230 |
| 16 | DVVPIND-------TRYRLVSCNTSVITQACPKVSFEPIPIHYCAPAGFAILKCNDKQFM | 230 |
| 17 | DVVPIND-------TRYRLVSCNTSVITQACPKVSFEPIPIHYCAPAGFAILKCNDKQFM | 230 |
| 18 | DVVPIND-------TRYRLVSCNTSVITQACPKVSFEPIPIHYCAPAGFAILKCNDKQFI | 230 |
| 19 | DVVPIND-------TRYRLVSCNTSVITQACPKVSFEPIPIHYCAPAGFAILKCNDKQFI | 230 |
| 20 | DVVPIND-------TRYRLVSCNTSVITQACPKVSFEPIPIHYCAPAGFAILKCNDKQFI | 230 |
| 21 | DVVPIND-------TRYRLVSCNTSVITQACPKVSFEPIPIHYCAPAGFAILKCNDKQFM | 230 |
| 22 | DVVPIND-------TRYRLVSCNTSVITQACPKVSFEPIPIHYCAPAGFAILKCNDKQFI | 230 |
| 23 | DVVPIND-------TRYRLVSCNTSVITQACPKVSFEPIPIHYCAPAGFAILKCNDKQFI | 230 |
| 24 | DVVPIND-------TRYRLVSCNTSVITQACPKVSFEPIPIHYCAPAGFAILKCNDKQFI | 230 |
| 25 | DVVPIND-------TRYRLVSCNTSVITQACPKVSFEPIPIHYCAPAGFAILKCNDKQFI | 230 |
| 26 | DVVPIND-------TRYRLVSCNTSVITQACPKVSFEPIPIHYCAPAGFAILKCNDKQFI | 230 |
| 27 | DVVPIND-------TRYRLVSCNTSVITQACPKVSFEPIPIHYCAPAGFAILKCNDKQFI | 230 |
| 28 | DVVPIND-------TRYRLVSCNTSVITQACPKVSFEPIPIHYCAPAGFAILKCNDKQFI | 230 |
| 29 | DVVPIND-------TRYRLVSCNTSVITQACPKVSFEPIPIHYCAPAGFAILKCNDKQFM | 230 |
| 30 | DVVPIND-------TRYRLVSCNTSVITQACPKVSFEPIPIHYCAPAGFAILKCNDKQFM | 230 |
| 31 | DVVPIND-------TRYRLVSCNTSVITQACPKVSFEPIPIHYCAPAGFAILKCNDKQFI | 230 |
| 32 | DVVPIND-------TRYRLVSCNTSVITQACPKVSFEPIPIHYCAPAGFAILKCNDKQFM | 230 |
| 33 | DVVPIND-------TRYRLVSCNTSVITQACPKVSFEPIPIHYCAPAGFAILKCNDKQFM | 230 |
| 34 | DVVPIND-------TRYRLVSCNTSVITQACPKVSFEPIPIHYCAPAGFAILKCNDKQFI | 230 |
| 35 | DVVPIND-------TRYRLVSCNTSVITQACPKVSFEPIPIHYCAPAGFAILKCNDKQFI | 230 |
| 36 | DVVPIND-------TRYRLVSCNTSVITQACPKVSFEPIPIHYCAPAGFAILKCNDKQFM | 230 |
| 37 | DVVPIND-------TRYRLVSCNTSVITQACPKVSFEPIPIHYCAPAGFAILKCNDKQFM | 230 |
| 38 | DVVPIND-------TRYRLVSCNTSVITQACPKVSFEPIPIHYCAPAGFAILKCNDKQFM | 230 |

| | Sequence | |
|---|---|---|
| | **GTGPCTNVSTVQCTHGIRPVVSTQLLLNGSLAEEEVVIRSVNFSDNAKTIIVQLNKSVEI** | |
| 1 | GTGPCTNVSTVQCTHGIRPVVSTQLLLNGSLAEEEVVIRSVNFSDNAKTIIVQLNKSVEI | 290 |
| 2 | GTGPCTNVSTVQCTHGIRPVVSTQLLLNGSLAEEEVVIRSVNFSDNAKTIIVQLNKSVEI | 290 |
| 3 | GTGPCTNVSTVQCTHGIRPVVSTQLLLNGSLAEEEVVIRSVNFSDNAKTIIVQLNKSVEI | 290 |
| 4 | GTGPCTNVSTVQCTHGIRPVVSTQLLLNGSLAEEEVVIRSVNFSDNAKTIIVQLNKSVEI | 290 |
| 5 | GTGPCTNVSTVQCTHGIRPVVSTQLLLNGSLAEEEVVIRSVNFSDNAKTIIVQLNKSVEI | 290 |
| 6 | GTGPCTNVSTVQCTHGIRPVVSTQLLLNGSLAEEEVVIRSVNFSDNAKTIIVQLNKSVEI | 290 |
| 7 | GTGPCTNVSTVQCTHGIRPVVSTQLLLNGSLAEEEVVIRSVNFSDNAKTIIVQLNKSVEI | 290 |
| 8 | GTGPCTNVSTVQCTHGIRPVVSTQLLLNGSLAEEEVVIRSVNFSDNAKTIIVQLNKSVEI | 290 |
| 9 | GTGPCTNVSTVQCTHGIRPVVSTQLLLNGSLAEEEVVIRSVNFSDNAKTIIVQLNKSVEI | 290 |
| 10 | GTGPCTNVSTVQCTHGIRPVVSTQLLLNGSLAEEEVVIRSVNFSDNAKTIIVQLNKSVEI | 290 |
| 11 | GTGPCTNVSTVQCTHGIRPVVSTQLLLNGSLAEEEVVIRSVNFSDNAKTIIVQLNKSVEI | 290 |
| 12 | GTGPCTNVSTVQCTHGIRPVVSTQLLLNGSLAEEEVVIRSVNFSDNAKTIIVQLNKSVEI | 290 |
| 13 | GTGPCTNVSTVQCTHGIRPVVSTQLLLNGSLAEEEVVIRSVNFSDNAKTIIVQLNKSVEI | 290 |
| 14 | GTGPCTNVSTVQCTHGIRPVVSTQLLLNGSLAEEEVVIRSVNFSDNAKTIIVQLNKSVEI | 290 |
| 15 | GTGPCTNVSTVQCTHGIRPVVSTQLLLNGSLAEEEVVIRSVNFSDNAKTIIVQLNKSVEI | 290 |
| 16 | GTGPCTNVSTVQCTHGIRPVVSTQLLLNGSLAEEEVVIRSVNFSDNAKTIIVQLNKSVEI | 290 |
| 17 | GTGPCTNVSTVQCTHGIRPVVSTQLLLNGSLAEEEVVIRSVNFSDNAKTIIVQLNKSVEI | 290 |
| 18 | GTGPCTNVSTVQCTHGIRPVVSTQLLLNGSLAEEEVVIRSGNFSDNAKTIIVQLNKSVEI | 290 |
| 19 | GTGPCTNVSTVQCTHGIRPVVSTQLLLNGSLAEEEVVIRSVNFSDNAKTIIVQLNKSVEI | 290 |
| 20 | GTGPCTNVSTVQCTHGIRPVVSTQLLLNGSLAEEEVVIRSVNFSDNAKTIIVQLNKSVEI | 290 |
| 21 | GTGPCTNVSTVQCTHGIRPVVSTQLLLNGSLAEEEVVIRSVNFSDNAKTIIVQLNKSVEI | 290 |
| 22 | GTGPCTNVSTVQCTHGIRPVVSTQLLLNGSLAEEEVVIRSVNFSDNAKTIIVQLNKSVEI | 290 |
| 23 | GTGPCTNVSTVQCTHGIRPVVSTQLLLNGSLAEEEVVIRSVNFSDNAKTIIVQLNKSVEI | 290 |
| 24 | GTGPCTNVSTVQCTHGIRPVVSTQLLLNGSLAEEEVVIRSVNFSDNAKTIIVQLNKSVEI | 290 |
| 25 | GTGPCTNVSTVQCTHGIRPVVSTQLLLNGSLAEEEVVIRSVNFSDNAKTIIVQLNKSVEI | 290 |
| 26 | GTGPCTNVSTVQCTHGIRPVVSTQLLLNGSLAEEEVVIRSVNFSDNAKTIIVQLNKSVEI | 290 |
| 27 | GTGPCTNVSTVQCTHGIRPVVSTQLLLNGSLAEEEVVIRSVNFSDNAKTIIVQLNKSVEI | 290 |
| 28 | GTGPCTNVSTVQCTHGIRPVVSTQLLLNGSLAEEEVVIRSVNFSDNAKTIIVQLNKSVEI | 290 |
| 29 | GTGPCTNVSTVQCTHGIRPVVSTQLLLNGSLAEEEVVIRSVNFSDNAKTIIVQLNKSVEI | 290 |
| 30 | GTGPCTNVSTVQCTHGIRPVVSTQLLLNGSLAEEEVVIRSVNFSDNAKTIIVQLNKSVEI | 290 |
| 31 | GTGPCTNVSTVQCTHGIRPVVSTQLLLNGSLAEEEVVIRSVNFSDNAKTIIVQLNKSVEI | 290 |
| 32 | GTGPCTNVSTVQCTYGIRPVVSTQLLLNGSLAEEEVVIRSVNFSDNAKTIIVQLNKSVEI | 290 |
| 33 | GTGPCTNVSTVQCTHGIRPVVSTQLLLNGSLAEEEVVIRSVNFSDNAKTIIVQLNKSVEI | 290 |
| 34 | GTGPCTNVSTVQCTHGIRPVVSTQLLLNGSLAEEEVVIRSVNFSDNAKTIIVQLNKSVEI | 290 |
| 35 | GTGPCTNVSTVQCTHGIRPVVSTQLLLNGSLAEEEVVIRSVNFSDNAKTIIVQLNKSVEI | 290 |
| 36 | GTGPCTNVSTVQCTHGIRPVVSTQLLLNGSLAEEEVVIRSVNFSDNAKTIIVQLNKSVEI | 290 |
| 37 | GTGPCTNVSTVQCTHGIRPVVSTQLLLNGSLAEEEVVIRSVNFSDNAKTIIVQLNKSVEI | 290 |
| 38 | GTGPCTNVSTVQCTHGIRPVVSTQLLLNGSLAEEEVVIRSVNFSDNAKTIIVQLNKSVEI | 290 |

| | | |
|---|---|---|
| | TCTRPNNNTRKSIPMGPGKAFYARGDIIGDIRKASCKINGTEWHSTLKLVVEKLREQY-N | |
| 1 | TCTRPNNNTRKSIPMGPGKAFYARGDITGDIRKAYCEINGTEWHSTLKLVVEKLREQY-N | 349 |
| 2 | TCTRPNNNTRKSIPMGPGKAFYARGDIIGDIRKASCKINGTEWHSTLKLVVEKLREQY-N | 349 |
| 3 | TCTRPNNNTRKSIPMGPGKAFYARGDIIGDIRKASCKINGTEWHSTLKLVVEKLREQY-N | 349 |
| 4 | TCTRPNNNTRKSIPMGPGKAFYARGDIIGDIRKASCKINGTEWHSTLKLVVEKLREQY-N | 349 |
| 5 | NCTRPNNNTRKSIPMGPGKAFYARGDIIGDIRKASCELNGTEWHSTLKLVVEKLREQY-N | 349 |
| 6 | NCTRPHNNTRKSIPMGPGKAFYARGDIIGDIRKAYCKINGTEWHSTLKLVVEKLREQY-N | 349 |
| 7 | TCTRPNNNTRKSIPMGPGKAFYARGDIIGDIRKASCKINGTEWHSTLKLVVEKLREQY-N | 349 |
| 8 | TCTRPNNNTRKSIPMGPGKAFYARGDIIGDIRKASCKINGTEWHSTLKLVVEKLREQY-N | 349 |
| 9 | TCTRPNNNTRKSIPMGPGKAFYARGDIIGDIRKASCKINGTEWHSTLKLVVEKLREQY-N | 349 |
| 10 | TCTRPNNNTRKSIPMGPGKAFYARGDIIGDIRKASCKINGTEWHSTLKLVVEKLREQY-N | 349 |
| 11 | TCTRPNNNTRKSIPMGPGKAFYARGDIIGDIRKASCKINGTEWHSTLKLVVEKLREQY-N | 349 |
| 12 | TCTRPNNNTRKSIPMGPGKAFYARGDIIGDIRKASCKINGTEWHSTLKLVVEKLREQY-N | 349 |
| 13 | NCTRPHNNTRKSIPMGPGKAFYARGDITGDIRKAYCEINGTEWHSTLKLVVEKLRGQY-N | 349 |
| 14 | TCTRPNNNTRKSIPMGPGKAFYARGDIIGDIRKASCKINGTEWHSTLKLVVEKLREQY-N | 349 |
| 15 | TCTRPNNNTRKSIPMGPGKAFYARGDIIGDIRKASCEINGTEWHSTLKLVVEKLREQY-N | 349 |
| 16 | TCTRPNNNTRKSIPMGPGKAFYARGDIIGDIRKASCKINGTEWHSTLKLVVEKLREQY-N | 349 |
| 17 | TCTRPNNNTRKSIPMGPGKAFYARGDIIGDIRKASCKINGTEWHSTLKLVVEKLREQY-N | 349 |
| 18 | NCTRPHNNTRKSIPMGPGKAFYARGDIIGDIRKAYCKINGTEWHSTLKLVVGKLREQY-N | 349 |
| 19 | TCTRPNNNTRKSIPMGPGKAFYARGDIIGDIRKASCKINGTEWHSTLKLVVEKLREQY-N | 349 |
| 20 | NCTRPNNNTRKSIPMGPGKAFYARGDIIGDIRKASCELNGTEWHSTLKLVVEKLREQY-N | 349 |
| 21 | TCTRPNNNTRKSIPMGPGKAFYARGDIIGDIRKASCKINGTEWHSTLKLVVEKLREQY-N | 349 |
| 22 | NCTRPHNNTRKSIPMGPGKAFYARGDIIGDIRKAYCKINGTEWHSTLKLVVEKLREQY-N | 349 |
| 23 | TCTRPNNNTRKSIPMGPGKAFYARGDIIGDIRKASCKINGTEWHSTLKLVVEKLREQY-N | 349 |
| 24 | TCTRPNNNTRKSIPMGPGKAFYARGDIIGDIRKASCKINGTEWHSTLKLVVEKLREQY-N | 349 |
| 25 | TCTRPNNNTRKSIPMGPGKAFYARGDIIGDIRKASCKINGTEWHSTLKLVVEKLREQY-N | 349 |
| 26 | TCTRPNNNTRKSIPMGPGKAFYARGDIIGDIRKASCKINGTEWHSTLKLVVEKLREQY-N | 349 |
| 27 | TCTRPNNNTRKSIPMGPGKAFYARGDIIGDIRKASCKINGTEWHSTLKLVVEKLREQY-N | 349 |
| 28 | TCTRPNNNTRKSIPMGPGKAFYARGDIIGDIRKASCKINGTEWHSTLKLVVEKLREQY-N | 349 |
| 29 | TCTRPNNNTRKSIPMGPGKAFYARGDIIGDIRKASCKINGTEWHSTLKLVVEKLREQY-N | 349 |
| 30 | TCTRPNNNTRKSIPMGPGKAFYARGDIIGDIRKASCKINGTEWHSTLKLVVEKLREQY-N | 349 |
| 31 | NCTRPHNNTRKSIPMGPGKAFYARGDIIGDIRKAYCKINGTEWHSTLKLVVEKLREQY-N | 349 |
| 32 | TCTRPNNNTRKSIPMGPGKAFYARGDIIGDIRKASCKINGTEWHSTLKLVVEKLREQY-N | 349 |
| 33 | TCTRPNNNTRKSIPMGPGKAFYARGDIIGDIRKASCKINGTEWHSTLKLVVEKLREQY-N | 349 |
| 34 | TCTRPNNNTRKSIPMGPGKAFYARGDIIGDIRKASCKINGTEWHSTLKLVVEKLREQY-N | 349 |
| 35 | NCTRPHNNTRKSIPMGPGKAFYARGDIIGDIRKAYCKINGTEWHSTLKLVVEKLREQY-N | 349 |
| 36 | TCTRPNNNTRKSIPMGPGKAFYARGDIIGDIRKASCKINGTEWHSTLKLVVEKLREQY-N | 349 |
| 37 | TCTRPNNNTRKSIPMGPGKAFYARGDIIGDIRKASCKINGTEWHSTLKLVVEKLREQY-N | 349 |
| 38 | TCTRPNNNTRKSIPMGPGKAFYARGDIIGDIRKASCKINGTEWHSTLKLVVEKLREQY-N | 349 |

| | | |
|---|---|---|
| | **KTIVFNRSSGGDPEIVMYSFNCGGEFFYCNSTKLFNSTWPWND-TKGSHDTNDTLMLPCK** | |
| 1 | KTIVFNRSSGGDPEIVMYSFNCGGEFFYCNSTKLFNSTWPWND-TKGSHDTNGTLILPCK | 408 |
| 2 | KTIVFNRSSGGDPEIVMYSFNCGGEFFYCNSTKLFNSTWPWND-TKGSHDTNDTLMLPCK | 408 |
| 3 | KTIVFNRSSGGDPEIVMYSFNCGGEFFYCNSTKLFNSTWPWND-TKGSHDTNDTLMLPCK | 408 |
| 4 | KTIVFNRSSGGDPEIVMYSFNCGGEFFYCNSTKLFNSTWPWND-TKGSHDTNDTLMLPCK | 408 |
| 5 | KTIVFNRSSGGDPEIVMYSFNCGGEFFYCNSTKLFNSTWPWND-TKGSHDTNGTLILPCR | 408 |
| 6 | KTIVFNRSSGGDPEIVMYSFNCGGEFFYCNSTKLFNSTWPWND-TKGSHDTNDTLMLPCK | 408 |
| 7 | KTIVFNRSSGGDPEIVMYSFNCGGEFFYCNSTKLFNSTWPWND-TKGSHDTNDTLMLPCK | 408 |
| 8 | KTIVFNRSSGGDPEIVMYSFNCGGEFFYCNSTKLFNSTWPWND-TKGSHDTNDTLMLPCK | 408 |
| 9 | KTIVFNRSSGGDPEIVMYSFNCGGEFFYCNSTKLFNSTWPWND-TKGSHDTNDTLMLPCK | 408 |
| 10 | KTIVFNRSSGGDPEIVMYSFNCGGEFFYCNSTKLFNSTWPWND-TKGSHDTNDTLMLPCK | 408 |
| 11 | KTIVFNRSSGGDPEIVMYSFNCGGEFFYCNSTKLFNSTWPWND-TKGSHDTNDTLMLPCK | 408 |
| 12 | KTIVFNRSSGGDPEIVMYSFNCGGEFFYCNSTKLFNSTWPWND-TKGSHDTNDTLMLPCK | 408 |
| 13 | KTIVFNRSSGGDPEIVMYSFNCGGEFFYCNSTKLFNSTWPWND-TKGSHDTNGTLILPCK | 408 |
| 14 | KTIVFNRSSGGDPEIVMYSFNCGGEFFYCNSTKLFNSTWPWND-TKGSHDTNDTLMLPCK | 408 |
| 15 | KTIVFNRSSGGDPEIVMYSFNCGGEFFYCNSTKLFNSTWPWND-TKGSHDTNGRLILPCR | 408 |
| 16 | KTIVFNRSSGGDPEIVMYSFNCGGEFFYCNSTKLFNSTWPWND-TKGSHDTNDTLMLPCK | 408 |
| 17 | KTIVFNRSSGGDPEIVMYSFNCGGEFFYCNSTKLFNSTWPWND-TKGSHDTNDTLMLPCK | 408 |
| 18 | KTIVFNRSSGGDPEIVMYSFNCGGEFFYCNSTKLFNSTWPWND-TKGSHDTNDTLMLPCK | 408 |
| 19 | KTIVFNRSSGGDPEIVMYSFNCGGEFFYCNSTKLFNSTWPWND-TKGSHDTNDTLMLPCK | 408 |
| 20 | KTIVFNRSSGGDPEIVMYSFNCGGEFFYCNSTKLFNSTWPWND-TKGSHDTNGTLILPCR | 408 |
| 21 | KTIVFNRSSGGDPEIVMYSFNCGGEFFYCNSTKLFNSTWPWND-TKGSHDTNDTLMLPCK | 408 |
| 22 | KTIVFNRSSGGDPEIVMYSFNCGGEFFYCNSTKLFNSTWPWND-TKGSHDTNDTLMLPCK | 408 |
| 23 | KTIVFNRSSGGDPEIVMYSFNCGGEFFYCNSTKLFNSTWPWND-TKGSHDTNDTLMLPCK | 408 |
| 24 | KTIVFNRSSGGDPEIVMYSFNCGGEFFYCNSTKLFNSTWPWND-TKGSHDTNDTLMLPCK | 408 |
| 25 | KTIVFNRSSGGDPEIVMYSFNCGGEFFYCNSTKLFNSTWPWND-TKGSHDTNDTLMLPCK | 408 |
| 26 | KTIVFNRSSGGDPEIVMYSFNCGGEFFYCN-TKLFNSTWPWND-TKGSHDTNDTLMLPCK | 407 |
| 27 | KTIVFNRSSGGDPEIVMYSFNCGGEFFYCNSTKLFNSTWPWND-TKGSHDTNDTLMLPCK | 408 |
| 28 | KTIVFNRSSGGDPEIVMYSFNCGGEFFYCNSTKLFNSTWPWND-TKGSHDTNDTLMLPCK | 408 |
| 29 | KTIVFNRSSGGDPEIVMYSFNCGGEFFYCNSTKLFNSTWPWND-TKGSHDTNDTLMLPCK | 408 |
| 30 | KTIVFNRSSGGDPEIVMYSFNCGGEFFYCNSTKLFNSTWPWND-TKGSHDTNDTLMLPCK | 408 |
| 31 | KTIVFNRSSGGDPEIVMYSFNCGGEFFYCNSTKLFNSTWPWND-TKGSHDTNDTLMLPCK | 408 |
| 32 | KTIVFNRSSGGDPEIVMYSFNCGGEFFYCNSTKLFNSTWPWND-TKGSHDTNDTLMLPCK | 408 |
| 33 | KTIVFNRSSGGDPEIVMYSFNCGGEFFYCNSTKLFNSTWPWND-TKGSHDTNDTLMLPCK | 408 |
| 34 | KTIVFNRSSGGDPEIVMYSFNCGGEFFYCNSTKLFNSTWPWND-TKGSHDTNDTLMLPCK | 408 |
| 35 | KTIVFNRSSGGDPEIVMYSFNCGGEFFYCNSTKLFNSTWPWND-TKGSHDTNDTLMLPCK | 408 |
| 36 | KTIVFNRSSGGDPEIVMYSFNCGGEFFYCNSTKLFNSTWPWND-TKGSHDTNDTLMLPCK | 408 |
| 37 | KTIVFNRSSGGDPEIVMYSFNCGGEFFYCNSTKLFNSTWPWND-TKGSHDTNDTLMLPCK | 408 |
| 38 | KTIVFNRSSGGDPEIVMYSFNCGGEFFYCNSTKLFNSTWPWND-TKGSHDTNDTLMLPCK | 408 |

| | Sequence | |
|---|---|---|
| | **IKQIINMWQGVGKAMYAPPIEGIIRCSSNITGLLLTRDGG-YESNETDEIFRPGGGDMRD** | |
| 1 | IKQIINMWQGVGKAMYAPPIEGKIRCSSNITGLLLTRDGG-YESNETDEIFRPGGGDMRD | 467 |
| 2 | IKQIINMWQGVGKAMYAPPIEGIIRCSSNITGLLLTRDGG-YESNETDEIFRPGGGDMRD | 467 |
| 3 | IKQIINMWQGVGKAMYAPPIEGIIRCSSNITGLLLTRDGG-YESNETDEIFRPGGGDMRD | 467 |
| 4 | IKQIINMWQGVGKAMYAPPIEGIIRCSSNITGLLLTRDGG-YESNETDEIFRPGGGDMRD | 467 |
| 5 | IKQIINMWQGVGKAMYAPPIEGEIRCSSNITGLLLTRDGG-YESNETDEIFRPGGGDMRD | 467 |
| 6 | IKQIINMWQGVGKAMYAPPIEGQIRCSSNITGLLLTRDGG-YESNETDEIFRPGGGDMRD | 467 |
| 7 | IKQIINMWQGVGKAMYAPPIEGIIRCSSNITGLLLTRDGG-YESNETDEIFRPGGGDMRD | 467 |
| 8 | IKQIINMWQGVGKAMYAPPIEGIIRCSSNITGLLLTRDGG-YESNETDEIFRPGGGDMRD | 467 |
| 9 | IKQIINMWQGVGKAMYAPPIEGIIRCSSNITGLLLTRDGG-YESNETDEIFRPGGGDMRD | 467 |
| 10 | IKQIINMWQGVGKAMYAPPIEGIIRCSSNITGLLLTRDGG-YESNETDEIFRPGGGDMRD | 467 |
| 11 | IKQIINMWQGVGKAMYAPPIEGIIRCSSNITGLLLTRDGG-YESNETDEIFRPGGGDMRD | 467 |
| 12 | IKQIINMWQGVGKAMYAPPIEGIIRCSSNITGLLLTRDGG-YESNETDEIFRPGGGDMRD | 467 |
| 13 | IKQIINMWQGVGKAMYAPPIEGKIRCSSNITGLLLTRDGG-YESNETDEIFRPGGGDMRD | 467 |
| 14 | IKQIINMWQGVGKAMYAPPIEGIIRCSSNITGLLLTRDGG-YESNETDEIFRPGGGDMRD | 467 |
| 15 | IKQIINMWQGVGKAMYAPPIEGEIRCSSNITGLLLTRDGG-YESNETDEIFRPGGGDMRD | 467 |
| 16 | IKQIINMWQGVGKAMYAPPIEGIIRCSSNITGLLLTRDGG-YESNETDEIFRPGGGDMRD | 467 |
| 17 | IKQIINMWQGVGKAMYAPPIEGIIRCSSNITGLLLTRDGG-YESNETDEIFRPGGGDMRD | 467 |
| 18 | IKQIINMWQGVGKAMYAPPIEGQIRCSSNITGLLLTRDGG-YESNETDEIFRPGGGDMRD | 467 |
| 19 | IKQIINMWQGVGKAMYAPPIEGIIRCSSNITGLLLTRDGG-YESNETDEIFRPGGGDMRD | 467 |
| 20 | IKQIINMWQGVGKAMYAPPIEGEIRCSSNITGLLLTRDGG-YESNETDEIFRPGGGDMRD | 467 |
| 21 | IKQIINMWQGVGKAMYAPPIEGIIRCSSNITGLLLTRDGG-YESNETDEIFRPGGGDMRD | 467 |
| 22 | IKQIINMWQGVGKAMYAPPIEGQIRCSSNITGLLLTRDGG-YESNETDEIFRPGGGDMRD | 467 |
| 23 | IKQIINMWQGVGKAMYAPPIEGIIRCSSNITGLLLTRDGG-YESNETDEIFRPGGGDMRD | 467 |
| 24 | IKQIINMWQGVGKAMYAPPIEGIIRCSSNITGLLLTRDGG-YESNETDEIFRPGGGDMRD | 467 |
| 25 | IKQIINMWQGVGKAMYAPPIEGIIRCSSNITGLLLTRDGG-YESNETDEIFRPGGGDMRD | 467 |
| 26 | IKQIINMWQGVGKAMYAPPIEGIIRCSSNITGLLLTRDGG-YESNETDEIFRPGGGDMRD | 466 |
| 27 | IKQIINMWQGVGKAMYAPPIEGIIRCSSNITGLLLTRDGG-YESNETDEIFRPGGGDMRD | 467 |
| 28 | IKQIINMWQGVGKAMYAPPIEGIIRCSSNITGLLLTRDGG-YESNETDEIFRPGGGDMRD | 467 |
| 29 | IKQIINMWQGVGKAMYAPPIEGIIRCSSNITGLLLTRDGG-YESNETDEIFRPGGGDMRD | 467 |
| 30 | IKQIINMWQGVGKAMYAPPIEGIIRCSSNITGLLLTRDGG-YESNETDEIFRPGGGDMRD | 467 |
| 31 | IKQIINMWQGVGKAMYAPPIEGQIRCSSNITGLLLTRDGG-YESNETDEIFRPGGGDMRD | 467 |
| 32 | IKQIINMWQGVGKAMYAPPIEGIIRCSSNITGLLLTRDGG-YESNETEEIFRPGGGDMRD | 467 |
| 33 | IKQIINMWQGVGKAMYAPPIEGIIRCSSNITGLLLTRDGG-YESNETDEIFRPGGGDMRD | 467 |
| 34 | IKQIINMWQGVGKAMYAPPIEGIIRCSSNITGLLLTRDGG-YESNETDEIFRPGGGDMRD | 467 |
| 35 | IKQIINMWQGVGKAMYAPPIEGQIRCSSNITGLLLTRDGG-YESNETDEIFRPGGGDMRD | 467 |
| 36 | IKQIINMWQGVGKAMYAPPIEGIIRCSSNITGLLLTRDGG-YESNETDEIFRPGGGDMRD | 467 |
| 37 | IKQIINMWQGVGKAMYAPPIEGIIRCSSNITGLLLTRDGG-YESNETDEIFRPGGGDMRD | 467 |
| 38 | IKQIINMWQGVGKAMYAPPIEGIIRCSSNITGLLLTRDGG-YESNETDEIFRPGGGDMRD | 467 |

| | Sequence | Length |
|---|---|---|
| | NWRSELYKYKVVKIEPLGVAPTKAKRRVVQREKRAFG-LGAVFLGFLGAAGSTMGAASIT | |
| 1 | NWRSELYKYKVVKIEPLGVAPTKAKRRVVQREKRAFG-LGAVFLGFLGAAGSTMGAASIT | 526 |
| 2 | NWRSELYKYKVVKIEPLGVAPTKAKRRVVQREKRAFG-LGAVFLGFLGAAGSTMGAASIT | 526 |
| 3 | NWRSELYKYKVVKIEPLGVAPTRAKRRVVQREKRAFG-LGAVFLGFLGAAGSTMGAASIT | 526 |
| 4 | NWRSELYKYKVVKIEPLGVAPTKAKRRVVQREKRAFG-LGAVFLGFLGAAGSTMGAASIT | 526 |
| 5 | NWRSELYKYKVVKIEPLGVAPTKAKRRVVQREKRAFG-LGAVFLGFLGAAGSTMGAASIT | 526 |
| 6 | NWRSELYKYKVVKIEPLGVAPTKAKRRVVQREKRAFG-LGAVFLGFLGAAGSTMGAASIT | 526 |
| 7 | NWRSELYKYKVVKIEPLGVAPTRAKRRVVQREKRAFG-LGAVFLGFLGAAGSTMGAASIT | 526 |
| 8 | NWRSELYKYKVVKIEPLGVAPTKAKRRVVQREKRAFG-LGAVFLGFLGAAGSTMGAASIT | 526 |
| 9 | NWRSELYKYKVVKIEPLGVAPTRAKRRVVQREKRAFG-LGAVFLGFLGAAGSTMGAASIT | 526 |
| 10 | NWRSELYKYKVVKIEPLGVAPTRAKRRVVQREKRAFG-LGAVFLGFLGAAGSTMGAASIT | 526 |
| 11 | NWRSELYKYKVVKIEPLGVAPTRAKRRVVQREKRAFG-LGAVFLGFLGAAGSTMGAASIT | 526 |
| 12 | NWRSELYKYKVVKIEPLGVAPTKAKRRVVQREKRAFG-LGAVFLGFLGAAGSTMGAASIT | 526 |
| 13 | NWRSELYKYKVVKIEPLGVAPTKAKRRVVQREKRAFG-LGAVFLGFLGAAGSTMGAASIT | 526 |
| 14 | NWRSELYKYKVVKIEPLGVAPTKAKRRVVQREKRAFG-LGAVFLGFLGAAGSTMGAASIT | 526 |
| 15 | NWRSELYKYKVVKIEPLGVAPTKAKRRVVQREKRAFG-LGAVFLGFLGAAGSTMGAASIT | 526 |
| 16 | NWRSELYKYKVVKIEPLGVAPTRAKRRVVQREKRAFG-LGAVFLGFLGAAGSTMGAASIT | 526 |
| 17 | NWRSELYKYKVVKIEPLGVAPTRAKRRVVQREKRAFG-LGAVFLGFLGAAGSTMGAASIT | 526 |
| 18 | NWRSELYKYKVVKIEPLGVAPTKAKRRVVQREKRAFG-LGAVFLGFLGAAGSTMGAASIT | 526 |
| 19 | NWRSELYKYKVVKIEPLGVAPTKAKRRVVQREKRAFG-LGAVFLGFLGAAGSTMGAASIT | 526 |
| 20 | NWRSELYKYKVVKIEPLGVAPTKAKRRVVQREKRAFG-LGAVFLGFLGAAGSTMGAASIT | 526 |
| 21 | NWRSELYKYKVVKIEPLGVAPTKAKRRVVQREKRAFG-LGAVFLGFLGAAGSTMGAASIT | 526 |
| 22 | NWRSELYKYKVVKIEPLGVAPTKAKRRVVQREKRAFG-LGAVFLGFLGAAGSTMGAASIT | 526 |
| 23 | NWRSELYKYKVVKIEPLGVAPTKAKRRVVQREKRAFG-LGAVFLGFLGAAGSTMGAASIT | 526 |
| 24 | NWRSELYKYKVVKIEPLGVAPTKAKRRVVQREKRAFG-LGAVFLGFLGAAGSTMGAASIT | 526 |
| 25 | NWRSELYKYKVVKIEPLGVAPTKAKRRVVQREKRAFG-LGAVFLGFLGAAGSTMGAASIT | 526 |
| 26 | NWRSELYKYKVVKIEPLGVAPTKAKRRVVQREKRAFG-LGAVFLGFLRAAGSTMGAASIT | 525 |
| 27 | NWRSELYKYKVVKIEPLGVAPTKAKRRVVQREKRAFG-LGAVFLGFLGAAGSTMGAASIT | 526 |
| 28 | NWRSELYKYKVVKIEPLGVAPTKAKRRVVQREKRAFG-LGAVFLGFLGAAGSTMGAASIT | 526 |
| 29 | NWRSELYKYKVVKIEPLGVAPTRAKRRVVQREKRAFG-LGAVFLGFLGAAGSTMGAASIT | 526 |
| 30 | NWRSELYKYKVVKIEPLGVAPTRAKRRVVQREKRAFG-LGAVFLGFLGAAGSTMGAASIT | 526 |
| 31 | NWRSELYKYKVVKIEPLGVAPTKAKRRVVQREKRAFG-LGAVFLGFLGAAGSTMGAASIT | 526 |
| 32 | NWRSELYKYKVVKIEPLGVAPTRAKRRVVQREKRAFG-LGAVFLGFLGAAGSTMGAASIT | 526 |
| 33 | NWRSELYKYKVVKIEPLGVAPTRAKRRVVQREKRAFG-LGAVFLGFLGAAGSTMGAASIT | 526 |
| 34 | NWRSELYKYKVVKIEPLGVAPTKAKRRVVQREKRAFG-LGAVFLGFLGAAGSTMGAASIT | 526 |
| 35 | NWRSELYKYKVVKIEPLGVAPTKAKRRVVQREKRAFG-LGAVFLGFLGAAGSTMGAASIT | 526 |
| 36 | NWRSELYKYKVVKIEPLGVAPTRAKRRVVQREKRAFG-LGAVFLGFLGAAGSTMGAASIT | 526 |
| 37 | NWRSELYKYKVVKIEPLGVAPTRAKRRVVQREKRAFG-LGAVFLGFLGAAGSTMGAASIT | 526 |
| 38 | NWRSELYKYKVVKIEPLGVAPTRAKRRVVQREKRAFG-LGAVFLGFLGAAGSTMGAASIT | 526 |

| | | |
|---|---|---|
| | LTVQARQLLSGIVQQQNNLLRAIEAQQHLLQLTVWGIKQLQARVLAVERYLKDQQLLGIW | |
| 1 | LTVQARQLLSGIVQQQNNLLRAIEAQQHLLQLTVWGIKQLQARVLAVERYLKDQQLLGIW | 586 |
| 2 | LTVQARQLLSGIVQQQNNLLRAIEAQQHLLQLTVWGIKQLQARVLAVERYLKDQQLLGIW | 586 |
| 3 | LTVQARQLLSGIVQQQNNLLRAIEAQQHLLQLTVWGIKQLQARVLAVERYLKDQQLLGIW | 586 |
| 4 | LTVQARQLLSGIVQQQNNLLRAIEAQQHLLQLTVWGIKQLQARVLAVERYLKDQQLLGIW | 586 |
| 5 | LTVQARQLLSGIVQQQNNLLRAIEAQQHLLQLTVWGIKQLQARVLAVERYLKDQQLLGIW | 586 |
| 6 | LTVQARQLLSGIVQQQNNLLRAIEAQQHLLQLTVWGIKQLQARVLAVERYLKDQQLLGIW | 586 |
| 7 | LTVQARQLLSGIVQQQNNLLRAIEAQQHLLQLTVWGIKQLQARVLAVERYLKDQQLLGIW | 586 |
| 8 | LTVQARQLLSGIVQQQNNLLRAIEAQQHLLQLTVWGIKQLQARVLAVERYLKDQQLLGIW | 586 |
| 9 | LTVQARQLLSGIVQQQNNLLRAIEAQQHLLQLTVWGIKQLQARVLAVERYLKDQQLLGIW | 586 |
| 10 | LTVQARQLLSGIVQQQNNLLRAIEAQQHLLQLTVWGIKQLQARVLAVERYLKDQQLLGIW | 586 |
| 11 | LTVQARQLLSGIVQQQNNLLRAIEAQQHLLQLTVWGIKQLQARVLAVERYLKDQQLLGIW | 586 |
| 12 | LTVQARQLLSGIVQQQNNLLRAIEAQQHLLQLTVWGIKQLQARVLAVERYLKDQQLLGIW | 586 |
| 13 | LTVQARQLLSGIVQQQNNLLRAIEAQQHLLQLTVWGIKQLQARVLAVERYLKDQQLLGIW | 586 |
| 14 | LTVQARQLLSGIVQQQNNLLRAIEAQQHLLQLTVWGIKQLQARVLAVERYLKDQQLLGIW | 586 |
| 15 | LTVQARQLLSGIVQQQNNLLRAIEAQQHLLQLTVWGIKQLQARVLAVERYLKDQQLLGIW | 586 |
| 16 | LTVQARQLLSGIVQQQNNLLRAIEAQQHLLQLTVWGIKQLQARVLAVERYLKDQQLLGIW | 586 |
| 17 | LTVQARQLLSGIVQQQNNLLRAIEAQQHLLQLTVWGIKQLQARVLAVERYLKDQQLLGIW | 586 |
| 18 | LTVQARQLLSGIVQQQNNLLRAIEAQQHLLQLTVWGIKQLQARVLAVERYLKDQQLLGIW | 586 |
| 19 | LTVQARQLLSGIVQQQNNLLRAIEAQQHLLQLTVWGIKQLQARVLAVERYLKDQQLLGIW | 586 |
| 20 | LTVQARQLLSGIVQQQNNLLRAIEAQQHLLQLTVWGIKQLQARVLAVERYLKDQQLLGIW | 586 |
| 21 | LTVQARQLLSGIVQQQNNLLRAIEAQQHLLQLTVWGIKQLQARVLAVERYLKDQQLLGIW | 586 |
| 22 | LTVQARQLLSGIVQQQNNLLRAIEAQQHLLQLTVWGIKQLQARVLAVERYLKDQQLLGIW | 586 |
| 23 | LTVQARQLLSGIVQQQNNLLRAIEAQQHLLQLTVWGIKQLQARVLAVERYLKDQQLLGIW | 586 |
| 24 | LTVQARQLLSGIVQQQNNLLRAIEAQQHLLQLTVWGIKQLQARVLAVERYLKDQQLLGIW | 586 |
| 25 | LTVQARQLLSGIVQQQNNLLRAIEAQQHLLQLTVWGIKQLQARVLAVERYLKDQQLLGIW | 586 |
| 26 | LTVQARQLLSGIVQQQNNLLRAIEAQQHLLQLTVWGIKQLQARVLAVERYLKDQQLLGIW | 585 |
| 27 | LTVQARQLLSGIVQQQNNLLRAIEAQQHLLQLTVWGIKQLQARVLAVERYLKDQQLLGIW | 586 |
| 28 | LTVQARQLLSGIVQQQNNLLRAIEAQQHLLQLTVWGIKQLQARVLAVERYLKDQQLLGIW | 586 |
| 29 | LTVQARQLLSGIVQQQNNLLRAIEAQQHLLQLTVWGIKQLQARVLAVERYLKDQQLLGIW | 586 |
| 30 | LTVQARQLLSGIVQQQNNLLRAIEAQQHLLQLTVWGIKQLQARVLAVERYLKDQQLLGIW | 586 |
| 31 | LTVQARQLLSGIVQQQNNLLRAIEAQQHLLQLTVWGIKQLQARVLAVERYLKDQQLLGIW | 586 |
| 32 | LTVQARQLLSGIVQQQNNLLRAIEAQQHLLQLTVWGIKQLQARVLAVERYLKDQQLLGIW | 586 |
| 33 | LTVQARQLLSGIVQQQNNLLRAIEAQQHLLQLTVWGIKQLQARVLAVERYLKDQQLLGIW | 586 |
| 34 | LTVQARQLLSGIVQQQNNLLRAIEAQQHLLQLTVWGIKQLQARVLAVERYLKDQQLLGIW | 586 |
| 35 | LTVQARQLLSGIVQQQNNLLRAIEAQQHLLQLTVWGIKQLQARVLAVERYLKDQQLLGIW | 586 |
| 36 | LTVQARQLLSGIVQQQNNLLRAIEAQQHLLQLTVWGIKQLQARVLAVERYLKDQQLLGIW | 586 |
| 37 | LTVQARQLLSGIVQQQNNLLRAIEAQQHLLQLTVWGIKQLQARVLAVERYLKDQQLLGIW | 586 |
| 38 | LTVQARQLLSGIVQQQNNLLRAIEAQQHLLQLTVWGIKQLQARVLAVERYLKDQQLLGIW | 586 |

| | Sequence | |
|---|---|---|
| | **GCSGKLICTTTVPWNTSWSNKSLEQIWDNMTWMEWEREIDNYTGYIYQLIEESQNQQEKN** | |
| 1 | GCSGKLICTTTVPWNTSWSNKSLEQIWDNMTWMEWEREIDNYTGYIYQLIEESQNQQEKN | 646 |
| 2 | GCSGKLICTTTVPWNTSWSNKSLEQIWDNMTWMEWEREIDNYTGYIYQLIEESQNQQEKN | 646 |
| 3 | GCSGKLICTTTVPWNTSWSNKSLEQIWDNMTWMEWEREIDNYTGYIYQLIEESQNQQEKN | 646 |
| 4 | GCSGKLICTTTVPWNTSWSNKSLEQIWDNMTWMEWEREIDNYTGYIYQLIEESQNQQEKN | 646 |
| 5 | GCSGKLICPTTVPWNTSWSNKSLEQIWDNMTWMEWEREIDNYTGYIYQLIEESQNQQEKN | 646 |
| 6 | GCSGKLICTTTVPWNTSWSNKSLEQIWDNMTWMEWEREIDNYTDYIYQLIEESQNQQEKN | 646 |
| 7 | GCSGKLICTTTVPWNTSWSNKSLEQIWDNMTWMEWEREIDNYTGYIYQLIEESQNQQEKN | 646 |
| 8 | GCSGKLICTTTVPWNTSWSNKSLEQIWDNMTWMEWEREIDNYTGYIYQLIEESQNQQEKN | 646 |
| 9 | GCSGKLICTTTVPWNTSWSNKSLEQIWDNMTWMEWEREIDNYTGYIYQLIEESQNQQEKN | 646 |
| 10 | GCSGKLICTTTVPWNTSWSNKSLEQIWDNMTWMEWEREIDNYTGYIYQLIEESQNQQEKN | 646 |
| 11 | GCSGKLICTTTVPWNTSWSNKSLEQIWDNMTWMEWEREIDNYTGYIYQLIEESQNQQEKN | 646 |
| 12 | GCSGKLICTTTVPWNTSWSNKSLEQIWDNMTWMEWEREIDNYTGYIYQLIEESQNQQEKN | 646 |
| 13 | GCSGKLICTTTVPWNTSWSNKSLEQIWDNMTWMEWEREIDNYTGYIYQLIEESQNQQEKN | 646 |
| 14 | GCSGKLICTTTVPWNTSWSNKSLEQIWDNMTWMEWEREIDNYTGYIYQLIEESQNQQEKN | 646 |
| 15 | GCSGKLICTTTVPWNTSWSNKSLEQIWDNMTWMEWEREIDNYTGYIYQLIEESQNQQEKN | 646 |
| 16 | GCSGKLICTTTVPWNTSWSNKSLEQIWDNMTWMEWEREIDNYTGYIYQLIEESQNQQEKN | 646 |
| 17 | GCSGKLICTTTVPWNTSWSNKSLEQIWDNMTWMEWEREIDNYTGYIYQLIEESQNQQEKN | 646 |
| 18 | GCSGKLICTTTVPWNTSWSNKSLEQIWDNMTWMEWEREIDNYTGYIYQLIEESQNQQEKN | 646 |
| 19 | GCSGKLICTTTVPWNTSWSNKSLEQIWDNMTWMEWEREIDNYTGYIYQLIEESQNQQEKN | 646 |
| 20 | GCSGKLICPTTVPWNTSWSNKSLEQIWDNMTWMEWEREIDNYTGYIYQLIEESQNQQEKN | 646 |
| 21 | GCSGKLICTTAVPWNTSWSNKSLEQIWDNMTWMEWEREIDNYTGYIYQLIEESQNQQEKN | 646 |
| 22 | GCSGKLICTTTVPWNTSWSNKSLEQIWDNMTWMEWEREIDNYTDYIYQLIEESQNQQEKN | 646 |
| 23 | GCSGKLICTTTVPWNTSWSNKSLEQIWDNMTWMEWEREIDNYTGYIYQLIEESQNQQEKN | 646 |
| 24 | GCSGKLICTTTVPWNTSWSNKSLEQIWDNMTWMEWEREIDNYTGYIYQLIEESQNQQEKN | 646 |
| 25 | GCSGKLICTTTVPWNTSWSNKSLEQIWDNMTWMEWEREIDNYTGYIYQLIEESQNQQEKN | 646 |
| 26 | GCSGKLICTTTVPWNTSWSNKSLEQIWDNMTWMEWEREIDNYTGYIYQLIEESQNQQEKN | 645 |
| 27 | GCSGKLICTTTVPWNTSWSNKSLEQIWDNMTWMEWEREIDNYTGYIYQLIEESQNQQEKN | 646 |
| 28 | GCSGKLICTTTVPWNTSWSNKSLEQIWDNMTWMEWEREIDNYTGYIYQLIEESQNQQEKN | 646 |
| 29 | GCSGKLICTTTVPWNTSWSNKSLEQIWDNMTWMEWEREIDNYTGYIYQLIEESQNQQEKN | 646 |
| 30 | GCSGKLICTTTVPWNTSWSNKSLEQIWDNMTWMEWEREIDNYTGYIYQLIEESQNQQEKN | 646 |
| 31 | GCSGKLICTTTVPWNTSWSNKSLEQIWDNMTWMEWEREIDNYTGYIYQLIEESQNQQEKN | 646 |
| 32 | GCSGKLICTTTVPWNTSWSNKSLEQIWDNMTWMEWEREIDNYTGYIYQLIEESQNQQEKN | 646 |
| 33 | GCSGKLICTTTVPWNTSWSNKSLEQIWDNMTWMEWEREIDNYTGYIYQLIEESQNQQEKN | 646 |
| 34 | GCSGKLICTTTVPWNTSWSNKSLEQIWDNMTWMEWEREIDNYTGYIYQLIEESQNQQEKN | 646 |
| 35 | GCSGKLICTTTVPWNTSWSNKSLEQIWDNMTWMEWEREIDNYTGYIYQLIEESQNQQEKN | 646 |
| 36 | GCSGKLICTTTVPWNTSWSNKSLEQIWDNMTWMEWEREIDNYTGYIYQLIEESQNQQEKN | 646 |
| 37 | GCSGKLICTTTVPWNTSWSNKSLEQIWDNMTWMEWEREIDNYTGYIYQLIEESQNQQEKN | 646 |
| 38 | GCSGKLICTTTVPWNTSWSNKSLEQIWDNMTWMEWEREIDNYTGYIYQLIEESQNQQEKN | 646 |

| | | |
|---|---|---|
| | **EQELLALDKWASLWNWFDITNWLWYIKIFIMIVGGLIGLRIVFTVLSIVNRVRQGYSPLS** | |
| 1 | EQELLALDKWASLWNWFDITNWLWYIKIFIMIVGGLIGLRIVFTVLSIVNRVRQGYSPLS | 706 |
| 2 | EQELLALDKWASLWNWFDITNWLWYIKIFIMIVGGLIGLRIVFTVLSIVNRVRQGYSPLS | 706 |
| 3 | EQELLALDKWASLWNWFDITNWLWYIKIFIMIVGGLIGLRIVFTVLSIVNRVRQGYSPLS | 706 |
| 4 | EQELLALDKWASLWNWFDITNWLWYIKIFIMIVGGLIGLRIVFTVLSIVNRVRQGYSPLS | 706 |
| 5 | EQELLALDKWASLWNWFDITNWLWYIKIFIMIVGGLIGLRIVFTVLSIVNRVRQGYSPLS | 706 |
| 6 | EQELLALDKWASLWNWFDITNWLWYIKIFIMIVGGLIGLRIVFTVLSIVNRVRQGYSPLS | 706 |
| 7 | EQELLALDKWASLWNWFDITNWLWYIKIFIMIVGGLIGLRIVFTVLSIVNRVRQGYSPLS | 706 |
| 8 | EQELLALDKWASLWNWFDITNWLWYIKIFIMIVGGLIGLRIVFTVLSIVNRVRQGYSPLS | 706 |
| 9 | EQELLALDKWASLWNWFDITNWLWYIKIFIMIVGGLIGLRIVFTVLSIVNRVRQGYSPLS | 706 |
| 10 | EQELLALDKWASLWNWFDITNWLWYIKIFIMIVGGLIGLRIVFTVLSIVNRVRQGYSPLS | 706 |
| 11 | EQELLALDKWASLWNWFDITNWLWYIKIFIMIVGGLIGLRIVFTVLSIVNRVRQGYSPLS | 706 |
| 12 | EQELLALDKWASLWNWFDITNWLWYIKIFIMIVGGLIGLRIVFTVLSIVNRVRQGYSPLS | 706 |
| 13 | EQELLALDKWASLWNWFDITNWLWYIKIFIMIVGGLIGLRIVFTVLSIVNRVRQGYSPLS | 706 |
| 14 | EQELLALDKWASLWNWFDITNWLWYIKIFIMIVGGLIGLRIVFTVLSIVNRVRQGYSPLS | 706 |
| 15 | EQELLALDKWASLWNWFDITNWLWYIKIFIMIVGGLIGLRIVFTVLSIVNRVRQGYSPLS | 706 |
| 16 | EQELLALDKWASLWNWFDITNWLWYIKIFIMIVGGLIGLRIVFTVLSIVNRVRQGYSPLS | 706 |
| 17 | EQELLALDKWASLWNWFDITNWLWYIKIFIMIVGGLIGLRIVFTVLSIVNRVRQGYSPLS | 706 |
| 18 | EQELLALDKWASLWNWFDITNWLWYIKIFIMIVGGLIGLRIVFTVLSIVNRVRQGYSPLS | 706 |
| 19 | EQELLALDKWASLWNWFDITNWLWYIKIFIMIVGGLIGLRIVFTVLSIVNRVRQGYSPLS | 706 |
| 20 | EQELLALDKWASLWNWFDITNWLWYIKIFIMIVGGLIGLRIVFTVLSIVNRVRQGYSPLS | 706 |
| 21 | EQELLALDKWASLWNWFDITNWLWYIKIFIMIVGGLIGLRIVFTVLSIVNRVRQGYSPLS | 706 |
| 22 | EQELLALDKWASLWNWFDITNWLWYIKIFIMIVGGLIGLRIVFTVLSIVNRVRQGYSPLS | 706 |
| 23 | EQELLALDKWASLWNWFDITNWLWYIKIFIMIVGGLIGLRIVFTVLSIVNRVRQGYSPLS | 706 |
| 24 | EQELLALDKWASLWNWFDITNWLWYIKIFIMIVGGLIGLRIVFTVLSIVNRVRQGYSPLS | 706 |
| 25 | EQELLALDKWASLWNWFDITNWLWYIKIFIMIVGGLIGLRIVFTVLSIVNRVRQGYSPLS | 706 |
| 26 | EQELLALDKWASLWNWFDITNWLWYIKIFIMIVGGLIGLRIVFTVLSIVNRVRQGYSPLS | 705 |
| 27 | EQELLALDKWASLWNWFDITNWLWYIKIFIMIVGGLIGLRIVFTVLSIVNRVRQGYSPLS | 706 |
| 28 | EQELLALDKWASLWNWFDITNWLWYIKIFIMIVGGLIGLRIVFTVLSIVNRVRQGYSPLS | 706 |
| 29 | EQELLALDKWASLWNWFDITNWLWYIKIFIMIVGGLIGLRIVFTVLSIVNRVRQGYSPLS | 706 |
| 30 | EQELLALDKWASLWNWFDITNWLWYIKIFIMIVGGLIGLRIVFTVLSIVNRVRQGYSPLS | 706 |
| 31 | EQELLALDKWASLWNWFDITNWLWYIKIFIMIVGGLIGLRIVFTVLSIVNRVRQGYSPLS | 706 |
| 32 | EQELLALDKWASLWNWFDITNWLWYIKIFIMIVGGLIGLRIVFTVLSIVNRVRQGYSPLS | 706 |
| 33 | EQELLALDKWASLWNWFDITNWLWYIKIFIMIVGGLIGLRIVFTVLSIVNRVRQGYSPLS | 706 |
| 34 | EQELLALDKWASLWNWFDITNWLWYIKIFIMIVGGLIGLRIVFTVLSIVNRVRQGYSPLS | 706 |
| 35 | EQELLALDKWASLWNWFDITNWLWYIKIFIMIVGGLIGLRIVFTVLSIVNRVRQGYSPLS | 706 |
| 36 | EQELLALDKWASLWNWFDITNWLWYIKIFIMIVGGLIGLRIVFTVLSIVNRVRQGYSPLS | 706 |
| 37 | EQELLALDKWASLWNWFDITNWLWYIKIFIMIVGGLIGLRIVFTVLSIVNRVRQGYSPLS | 706 |
| 38 | EQELLALDKWASLWNWFDITNWLWYIKIFIMIVGGLIGLRIVFTVLSIVNRVRQGYSPLS | 706 |

| | | |
|---|---|---|
| | **FQTHLPAQRGPDRPEGIGEEGGERDRDRSDPLVNGFLTLIWSDLRSLCLFSYHRLRDLLL** | |
| 1 | FQTHLPAQRGPDRPEGIGEEGGERDRDRSDPLVNGFLALIWSDLRSLCLFSYHRLRDLLL | 766 |
| 2 | FQTHLPAQRGPDRPEGIGEEGGERDRDRSDPLVNGFLTLIWSDLRSLCLFSYHRLRDLLL | 766 |
| 3 | FQTHLPAQRGPDRPEGIGEEGGERDRDRSDPLVNGFLTLIWSDLRSLCLFSYHRLRDLLL | 766 |
| 4 | FQTHLPAQRGPDRPEGIGEEGGERDRDRSDPLVNGFLTLIWSDLRSLCLFSYHRLRDLLL | 766 |
| 5 | FQTHLPAQRGPDRPEGIGEEGGERDRDRSDPLVNGFLTLIWSDLRSLCLFSYHRLRDLLL | 766 |
| 6 | FQTHLPAQRGPDRPEGIGEEGGERDRDRSDPLVNGFLTLIWSDLRSLCLFSYHRLRDLLL | 766 |
| 7 | FQTHLPAQRGPDRPEGIGEEGGERDRDRSDPLVNGFLTLIWSDLRSLCLFSYHRLRDLLL | 766 |
| 8 | FQTHLPAQRGPDRPEGIGEEGGERDRDRSDPLVNGFLTLIWSDLRSLCLFSYHRLRDLLL | 766 |
| 9 | FQTHLPAQRGPDRPEGIGEEGGERDRDRSDPLVNGFLTLIWSDLRSLCLFSYHRLRDLLL | 766 |
| 10 | FQTHLPAQRGPDRPEGIGEEGGERDRDRSDPLVNGFLTLIWSDLRSLCLFSYHRLRDLLL | 766 |
| 11 | FQTHLPAQRGPDRPEGIGEEGGERDRDRSDPLVNGFLTLIWSDLRSLCLFSYHRLRDLLL | 766 |
| 12 | FQTHLPAQRGPDRPEGIGEEGGERDRDRSDPLVNGFLTLIWSDLRSLCLFSYHRLRDLLL | 766 |
| 13 | FQTHLPAQRGPDRPEGIGEEGGERDRDRSDPLVNGFLTLIWSDLRSLCLFSYHRLRDLLL | 766 |
| 14 | FQTHLPAQRGPDRPEGIGEEGGERDRDRSDPLVNGFLTLIWSDLRSLCLFSYHRLRDLLL | 766 |
| 15 | FQTHLPAQRGPDRPEGIGEEGGERDRDRSDPLVNGFLTLIWSDLRSLCLFSYHRLRDLLL | 766 |
| 16 | FQTHLPAQRGPDRPEGIGEEGGERDRDRSDPLVNGFLTLIWSDLRSLCLFSYHRLRDLLL | 766 |
| 17 | FQTHLPAQRGPDRPEGIGEEGGERDRDRSDPLVNGFLTLIWSDLRSLCLFSYHRLRDLLL | 766 |
| 18 | FQTHLPAQRGPDRPEGIGEEGGERDRDRSDPLVNGFLTLIWSDLRSLCLFSYHRLRDLLL | 766 |
| 19 | FQTHLPAQRGPDRPEGIGEEGGERDRDRSDPLVNGFLTLIWSDLRSLCLFSYHRLRDLLL | 766 |
| 20 | FQTHLPAQRGPDRPEGIGEEGGERDRDRSDPLVNGFLTLIWSDLRSLCLFSYHRLRDLLL | 766 |
| 21 | FQTHLPAQRGPDRPEGIGEEGGERDRDRSDPLVNGFLTLIWSDLRSLCLFSYHRLRDLLL | 766 |
| 22 | FQTHLPAQRGPDRPEGIGEEGGERDRDRSDPLVNGFLTLIWSDLRSLCLFSYHRLRDLLL | 766 |
| 23 | FQTHLPAQRGPDRPEGIGEEGGERDRDRSDPLVNGFLTLIWSDLRSLCLFSYHRLRDLLL | 766 |
| 24 | FQTHLPAQRGPDRPEGIGEEGGERDRDRSDPLVNGFLTLIWSDLRSLCLFSYHRLRDLLL | 766 |
| 25 | FQTHLPAQRGPDRPEGIGEEGGERDRDRSDPLVNGFLTLIWSDLRSLCLFSYHRLRDLLL | 766 |
| 26 | FQTHLPAQRGPDRPEGIGEEGGERDRDRSDPLVNGFLTLIWSDLRSLCLFSYHRLRDLLL | 765 |
| 27 | FQTHLPAQRGPDRPEGIGEEGGERDRDRSDPLVNGFLTLIWSDLRSLCLFSYHRLRDLLL | 766 |
| 28 | FQTHLPAQRGPDRPEGIGEEGGERDRDRSDPLVNGFLTLIWSDLRSLCLFSYHRLRDLLL | 766 |
| 29 | FQTHLPAQRGPDRPEGIGEEGGERDRDRSDPLVNGFLTLIWSDLRSLCLFSYHRLRDLLL | 766 |
| 30 | FQTHLPAQRGPDRPEGIGEEGGERDRDRSDPLVNGFLTLIWSDLRSLCLFSYHRLRDLLL | 766 |
| 31 | FQTHLPAQRGPDRPEGIGEEGGERDRDRSDPLVNGFLTLI-SDLRSLCLFSYHRLRDLLL | 765 |
| 32 | FQTHLPAQRGPDRPEGIGEEGGERDRDRSDPLVNGFLTLIWSDLRSLCLFSYHRLRDLLL | 766 |
| 33 | FQTHLPAQRGPDRPEGIGEEGGERDRDRSDPLVNGFLTLIWSDLRSLCLFSYHRLRDLLL | 766 |
| 34 | FQTHLPAQRGPDRPEGIGEEGGERDRDRSDPLVNGFLTLIWSDLRSLCLFSYHRLRDLLL | 766 |
| 35 | FQTHLPAQRGPDRPEGIGEEGGERDRDRSDPLVNGFLTLIWSDLRSLCLFSYHRLRDLLL | 766 |
| 36 | FQTHLPAQRGPDRPEGIGEEGGERDRDRSDPLVNGFLTLIWSDLRSLCLFSYHRLRDLLL | 766 |
| 37 | FQTHLPAQRGPDRPEGIGEEGGERDRDRSDPLVNGFLTLIWSDLRSLCLFSYHRLRDLLL | 766 |
| 38 | FQTHLPAQRGPDRPEGIGEEGGERDRDRSDPLVNGFLTLIWSDLRSLCLFSYHRLRDLLL | 766 |

| | Sequence | Length |
| --- | --- | --- |
|  | **IVTRIVELLGR-------RGWEVLKYWWNLLQYWSQELKNSAVSLLNATATAVAEGTDRV** |  |
| 1 | IVTRIVELLGR-------RGWEVLKYWWNLLQYWSQELKNSAVSLLNATAIAVAEGTDRV | 819 |
| 2 | IVTRIVELLGR-------RGWEVLKYWWNLLQYWSQELKNSAVSLLNATATAVAEGTDRV | 819 |
| 3 | IVTRIVELLGR-------RGWEVLKYWWNLLQYWSQELKNSAVSLLNATATAVAEGTDRV | 819 |
| 4 | IVTRIVELLGR-------RGWEILKYWWNLLQYWSQELKNSAVSLLNATATAVAEGTDRV | 819 |
| 5 | IVTRIVELLGR-------RGWEVLKYWWNLLQYWSQELKNSAVSLLNATAIAVAEGTDRV | 819 |
| 6 | IVTRIVELLGR-------RGWEVLKYWWNLLQYWSQELKNSAVSLLNATATAVAEGTDRV | 819 |
| 7 | IVTRIVELLGR-------RGWEVLKYWWNLLQYWSQELKNSAVSLLNATATAVAEGTDRV | 819 |
| 8 | IVTRIVELLGR-------RGWEVLKYWWNLLQYWSQELKNSAVSLLNATATAVAEGTDRV | 819 |
| 9 | IVTRIVELLGR-------RGWEVLKYWWNLLQYWSQELKNSAVSLLNATATAVAEGTDRV | 819 |
| 10 | IVTRIVELLGR-------RGWEVLKYWWNLLQYWSQELKNSAVSLLNATATAVAEGTDRV | 819 |
| 11 | IVTRIVELLGR-------RGWEVLKYWWNLLQYWSQELKNSAVSLLNATATAVAEGTDRV | 819 |
| 12 | IVTRIVELLGR-------RGWEVLKYWWNLLQYWSQELKNSAVSLLNATATAVAEGTDRV | 819 |
| 13 | IVTRIVELLGR-------RGWEVLKYWWNLLQYWSQELKNSAVSLLNATATAVAEGTDRV | 819 |
| 14 | IVTRIVELLGR-------RGWEVLKYWWNLLQYWSQELKNSAVSLLNATATAVAEGTDRV | 819 |
| 15 | IVTRIVELLGR-------RGWEVLKYWWNLLQYWSQELKNSAVSLLNATAIAVAEGTDRV | 819 |
| 16 | IVTRIVELLGR-------RGWEVLKYWWNLLQYWSQELKNSAVSLLNATATAVAEGTDRV | 819 |
| 17 | IVTRIVELLGR-------RGWEVLKYWWNLLQYWSQELKNSAVSLLNATATAVAEGTDRV | 819 |
| 18 | IVTRIVELLGR-------RGWEVLKY-WNLLQYWSQELKNSAVSLLNATATAVAEGTDRV | 818 |
| 19 | IVTRIVELLGR-------RGWEVLKYWWNLLQYWSQELKNSAVSLLNATATAVAEGTDRV | 819 |
| 20 | IVTRIVELLGR-------RGWEVLKYWWNLLQYWSQELKNSAVSLLNATAIAVAEGTDRV | 819 |
| 21 | IVTRIVELLGR-------RGWEVLKYWWNLLQYWSQELKNSAVSLLNATATAVAEGTDRV | 819 |
| 22 | IVTRIVELLGR-------RGWEVLKYWWNLLQYWSQELKNSAVSLLNATATAVAEGTDRV | 819 |
| 23 | IVTRIVELLGR-------RGWEVLKYWWNLLQYWSQELKNSAVSLLNATATAVAEGTDRV | 819 |
| 24 | IVTRIVELLGR-------RGWEVLKYWWNLLQYWSQELKNSAVSLLNATATAVAEGTDRV | 819 |
| 25 | IVTRIVELLGR-------RGWEVLKYWWNLLQYWSQELKNSAVSLLNATATAVAEGTDRV | 819 |
| 26 | IVTRIVELLGR-------RGWEVLKYWWNLLQYWSQELKNSAVSLLNATATAVAEGTDRV | 818 |
| 27 | IVTRIVELLGR-------RGWEVLKYWWNLLQYWSQELKNSAVSLLNATATAVAEGTDRV | 819 |
| 28 | IVTRIVELLGR-------RGWEVLKYWWNLLQYWSQELKNSAVSLLNATATAVAEGTDRV | 819 |
| 29 | IVTRIVELLGR-------RGWEVLKYWWNLLQYWSQELKNSAVSLLNATATAVAEGTDRV | 819 |
| 30 | IVTRIVELLGR-------RGWEVLKYWWNLLQYWSQELKNSAVSLLNATATAVAEGTDRV | 819 |
| 31 | IVTRIVELLGR-------RGWEVLKYWWNLLQYWSQELKNSAVSLLNATATAVAEGTDRV | 818 |
| 32 | IVTRIVELLGR-------RGWEVLKYWWNLLQYWSQELKNSAVSLLNATATAVAEGTDRV | 819 |
| 33 | IVTRIVELLGR-------RGWEVLKYWWNLLQYWSQELKNSAVSLLNATATAVAEGTDRV | 819 |
| 34 | IVTRIVELLGR-------RGWEVLKYWWNLLQYWSQELKNSAVSLLNATATAVAEGTDRV | 819 |
| 35 | IVTRIVELLGR-------RGWEVLKYWWNLLQYWSQELKNSAVSLLNATATAVAEGTDRV | 819 |
| 36 | IVTRIVELLGR-------RGWEVLKYWWNLLQYWSQELKNSAVSLLNATATAVAEGTDRV | 819 |
| 37 | IVTRIVELLGR-------RGWEVLKYWWNLLQYWSQELKNSAVSLLNATATAVAEGTDRV | 819 |
| 38 | IVTRIVELLGR-------RGWEVLKYWWNLLQYWSQELKNSAVSLLNATATAVAEGTDRV | 819 |

| | | |
|---|---|---|
| | IEVVQRACRAILHIPRRIRQGLERALL | |
| 1 | IEVVQRACRAILHIPRRIRQGLERALL | 846 |
| 2 | IEVVQRACRAILHIPRRIRQGLERALL | 846 |
| 3 | IEVVQRACRAILHIPRRIRQGLERALL | 846 |
| 4 | VEVVQRACRAILHIPRRIRQGLERALL | 846 |
| 5 | IEVVQRACRAILHIPRRIRQGLERALL | 846 |
| 6 | IEVVQRACRAILHIPRRIRQGLERALL | 846 |
| 7 | IEVVQRACRAILHIPRRIRQGLERALL | 846 |
| 8 | IEVVQRACRAILHIPRRIRQGLERALL | 846 |
| 9 | IEVVQRACRAILHIPRRIRQGLERALL | 846 |
| 10 | IEVVQRACRAILHIPRRIRQGLERALL | 846 |
| 11 | IEVVQRACRAILHIPRRIRQGLERALL | 846 |
| 12 | IEVVQRACRAILHIPRRIRQGLERALL | 846 |
| 13 | IEVVQRACRAILHIPRRIRQGLERALL | 846 |
| 14 | IEVVQRACRAILHIPRRIRQGLERALL | 846 |
| 15 | IEVVQRACRAIRHIPRRIRQGLERALL | 846 |
| 16 | IEVVQRACRAILHIPRRIRQGLERALL | 846 |
| 17 | IEVVQRACRAILHIPRRIRQGLERALL | 846 |
| 18 | VEVVQRACRAILHIPRRIRQGLERALL | 845 |
| 19 | IEVVQRACRAILHIPRRIRQGLERALL | 846 |
| 20 | IEVVQRACRAILHIPRRIRQGLERALL | 846 |
| 21 | VEVVQRACRAILHIPRRIRQGLERALL | 846 |
| 22 | IEVVQRACRAILHIPRRIRQGLERALL | 846 |
| 23 | IEVVQRACRAILHIPRRIRQGLERALL | 846 |
| 24 | IEVVQRACRAILHIPRRIRQGLERALL | 846 |
| 25 | IEVVQRACRAILHIPRRIRQGLERALL | 846 |
| 26 | IEVVQRACRAILHIPRRIRQGLERALL | 845 |
| 27 | IEVVQRACRAILHIPRRIRQGLERALL | 846 |
| 28 | IEVVQRACRAILHIPRRIRQGLERALL | 846 |
| 29 | IEVVQRACRAILHIPRRIRQGLERALL | 846 |
| 30 | IEVVQRACRAILHIPRRIRQGLERALL | 846 |
| 31 | VEVVQRACRAILHIPRRIRQGLERALL | 845 |
| 32 | IEVVQRACRAILHIPRRIRQGLERALL | 846 |
| 33 | IEVVQRACRAILHIPRRIRQGLERALL | 846 |
| 34 | IEVVQRACRAILHIPRRIRQGLERALL | 846 |
| 35 | VEVVQRACRAILHIPRRIRQGLERALL | 846 |
| 36 | IEVVQRACRAILHIPRRIRQGLERALL | 846 |
| 37 | IEVVQRACRAILHIPRRIRQGLERALL | 846 |
| 38 | IEVVQRACRAILHIPRRIRQGLERALL | 846 |

**Consensus**

1. B.US.2006.700010040_C9_4520.EU289193
2. B.US.2008.CH0040_3_d0568_ipe032_2_01.MG900308
3. B.US.2008.CH0040_3_d0568_ipe032_3_03.MG900309
4. B.US.2008.CH0040_3_d0568_ipe032_3_04.MG900310
5. B.US.2008.CH0040_3_d0568_ipe032_3_06.MG900311
6. B.US.2008.CH0040_3_d0568_ipe032_3_13.MG900312
7. B.US.2008.CH0040_3_d0568_ipe032_3_15.MG900313
8. B.US.2008.CH0040_3_d0568_ipe032_3_17.MG900314
9. B.US.2008.CH0040_3_d0568_ipe032_3_18.MG900315
10. B.US.2008.CH0040_3_d0568_ipe032_3_20.MG900316
11. B.US.2008.CH0040_3_d0568_ipe032_3_21.MG900317
12. B.US.2008.CH0040_3_d0568_ipe032_3_22.MG900318
13. B.US.2008.CH0040_3_d0568_ipe032_3_25.MG900319
14. B.US.2008.CH0040_3_d0568_ipe032_3_26.MG900320
15. B.US.2008.CH0040_3_d0568_ipe032_3_27.MG900321
16. B.US.2008.CH0040_3_d0568_ipe032_3_29.MG900322
17. B.US.2008.CH0040_3_d0568_ipe032_3_30.MG900323
18. B.US.2008.CH0040_3_d0568_ipe032_3_32.MG900324
19. B.US.2008.CH0040_3_d0568_ipe032_3_33.MG900325
20. B.US.2008.CH0040_3_d0568_ipe032_3_34.MG900326
21. B.US.2008.CH0040_3_d0568_ipe032_3_36.MG900327
22. B.US.2008.CH0040_3_d0568_ipe032_3_38.MG900328
23. B.US.2008.CH0040_3_d0568_ipe032_3_39.MG900329
24. B.US.2008.CH0040_3_d0568_ipe032_3_40.MG900330
25. B.US.2008.CH0040_3_d0568_ipe032_3_42.MG900331
26. B.US.2008.CH0040_3_d0568_ipe032_3_43.MG900332
27. B.US.2008.CH0040_3_d0568_ipe032_3_44.MG900333
28. B.US.2008.CH0040_3_d0568_ipe032_3_47.MG900334
29. B.US.2008.CH0040_3_d0568_ipe032_3_49.MG900335
30. B.US.2008.CH0040_3_d0568_ipe032_3_50.MG900336
31. B.US.2008.CH0040_3_d0568_ipe032_3_52.MG900337
32. B.US.2008.CH0040_3_d0568_ipe032_3_53.MG900338
33. B.US.2008.CH0040_3_d0568_ipe032_3_54.MG900339
34. B.US.2008.CH0040_3_d0568_ipe032_3_56.MG900340
35. B.US.2008.CH0040_3_d0568_ipe032_3_58.MG900341

| | | |
|---|---|---|
| | MRVMGIRKNYQHLWREGILLLGILMICSAADNLWVTVYYGVPVWREATTTLFCASDAKAY | |
| 1 | MRVMGIRKNYQHLWREGILLLGILMICSAADNLWVTVYYGVPVWREATTTLFCASDAKAY | 60 |
| 2 | MRVMGIRKNYQHLWREGILLLGILMICSAADNLWVTVYYGVPVWREATTTLFCASDAKAY | 60 |
| 3 | MRVMGIRKNYQHLWREGILLLGILMICSAADNLWVTVYYGVPVWREATTTLFCASDAKAY | 60 |
| 4 | MRVMGIRKNYQHLWREGILLLGILMICSAADNLWVTVYYGVPVWREATTTLFCASDAKAY | 60 |
| 5 | MRVMGIRKNYQHLWREGILLLGILMICSAADNLWVTVYYGVPVWREATTTLFCASDAKAY | 60 |
| 6 | MRVMGIRKNYQHLWREGILLLGILMICSAADNLRVTVYYGVPVWREATTTLFCASDAKAY | 60 |
| 7 | MRVMGIRKNYQHLWREGILLLGILMICSAADNLWVTVYYGVPVWREATTTLFCASDAKAY | 60 |
| 8 | MRVMGIRKNYQHLWREGILLLGILMICSAADNLWVTVYYGVPVWREATTTLFCASDAKAY | 60 |
| 9 | MRVMGIRKNYQHLWREGILLLGILMICSAADNLWVTVYYGVPVWREATTTLFCASDAKAY | 60 |
| 10 | MRVMGIRKNYQHLWREGILLVGILMICSAADNLWVTVYYGVPVWREATTTLFCASDAKAY | 60 |
| 11 | MRVMGIRKNYQHLWREGILLLGLLMICSAADNLWVTVYYGVPVWREATTTLFCASDAKAY | 60 |
| 12 | MRVMGIRKNYQHLWREGILLLGILMICSAADNLWVTVYYGVPVWREATTTLFCASDAKAY | 60 |
| 13 | MRVMGIRKNYQHLWKEGILLLGILMICSAADNLWVTVYYGVPVWREATTTLFCASDAKAY | 60 |
| 14 | MRVMGIRKNYQHLWREGILLLGILMICSAADNLWVTVYYGVPVWREATTTLFCASDAKAY | 60 |
| 15 | MRVMGIRKNYQHLWREGILLLGILMICSAADNLWVTVYYGVPVWREATTTLFCASDAKAY | 60 |
| 16 | MRVMGIRKNYQHLWREGILLLGILMICSAADNLWVTVYYGVPVWREATTTLFCASDAKAY | 60 |
| 17 | MRVMGIRKNYQHLWREGILLLGILMICSAADNLWVTVYYGVPVWREATTTLFCASDAKAY | 60 |
| 18 | MRVMGIRKNYQHLWREGILLLGILMICSAADNLWVTVYYGVPVWREATTTLFCASDAKAY | 60 |
| 19 | MRVMGIRKNYQHLWREGILLLGILMICSAADNLWVTVYYGVPVWREATTTLFCASDAKAY | 60 |
| 20 | MRVMGIRKNYQHLWREGILLLGILMICSAADNLWVTVYYGVPVWREATTTLFCASDAKAY | 60 |
| 21 | MRVMGIRKNYQHLWREGILLLGILMICSAADNLWVTVYYGVPVWREATTTLFCASDAKAY | 60 |
| 22 | MRVMGIRKNYQHLWREGILLLGILMICSAADNLWVTVYYGVPVWREATTTLFCASDAKAY | 60 |
| 23 | MRVMGIRKNYQHLWREGILLLGILMICSAADNLWVTVYYGVPVWREATTTLFCASDAKAY | 60 |
| 24 | MRVMGIRKNYQHLWREGILLLGILMICSAADNLWVTVYYGVPVWREATTTLFCASDAKAY | 60 |
| 25 | MRVMGIRKNYQHLWREGILLLGILMICSAADNLWVTVYYGVPVWREATTTLFCASDAKAY | 60 |
| 26 | MRVMGIRKNYQHLWREGILLLGILMICSAADNLWVTVYYGVPVWREATTTLFCASDAKAY | 60 |
| 27 | MRVMGIRKNYQHLWREGILLLGILMICSAADNLWVTVYYGVPVWREATTTLFCASDAKAY | 60 |
| 28 | MRVMGIRKNYQHLWREGILLLGILMICSAADNLWVTVYYGVPVWREATTTLFCASDAKAY | 60 |
| 29 | MRVMGIRKNYQHLWREGILLLGILMICSAADNLWVTVYYGVPVWREATTTLFCASDAKAY | 60 |
| 30 | MRVMGIRKNYQHLWREGILLLGILMICSAADNLWVTVYYGVPVWREATTTLFCASDAKAY | 60 |
| 31 | MRVMGIRKNYQHLWREGILLLGILMICSAADNLWVTVYYGVPVWREATTTLFCASDAKAY | 60 |
| 32 | MRVMGIRKNYQHLWREGILLLGILMICSAADNLWVTVYYGVPVWREATTTLFCASDAKAY | 60 |
| 33 | MRVMGIRKNYQHLWREGILLLGILMICSAADNLWVTVYYGVPVWREATTTLFCASDAKAY | 60 |
| 34 | MRVMGIRKNYQHLWREGILLLGILMICSAADNLWVTVYYGVPVWR--------------- | 45 |
| 35 | MRVMGIRKNYQHLWREGILLLGILMICSAADNLWVTVYYGVPVWREATTTLFCASDAKAY | 60 |

| | | |
|---|---|---|
| | DTEAHNVWATHACVPTDPNPQEVELKNVTENFNMWENNMVEQMHEDIISLWDQSLKPCVK | |
| 1 | DTEAHNVWATHACVPTDPNPQEVELKNVTENFNMWENNMVEQMHEDIISLWDQSLKPCVK | 120 |
| 2 | DTEAHNVWATHACVPTDPNPQEVELKNVTENFNMWENNMVEQMHEDIISLWDQSLKPCVK | 120 |
| 3 | DTEAHNVWATHACVPTDPNPQEVELKNVTENFNMWENNMVEQMHEDIISLWDQSLKPCVK | 120 |
| 4 | DTEAHNVWATHACVPTDPNPQEVELKNVTENFHMWENNMVEQMHEDIISLWDQSLKPCVK | 120 |
| 5 | DTEAHNVWATHACVPTDPNPQEVELKNVTENFNMWENNMVEQMHEDIISLWDQSLKPCVK | 120 |
| 6 | DTEAHNVWATHACVPTDPNPQEVELKNVTENFNMWENNMVEQMHEDIISLWDQSLKPCVK | 120 |
| 7 | DTEAHNVWATHDCVPTDPNPQEVELKNVTENFNMWENNMVEQMHEDIISLWDQSLKPCVK | 120 |
| 8 | DTEAHNVWATHACVPTDPNPQEVELKNVTENFNMWENNMVEQMHEDIISLWDQSLKPCVK | 120 |
| 9 | DTEAHNVWATHACVPTDPNPQEVELKNVTENFNMWENNMVEQMHEDIISLWDQSLKPCVK | 120 |
| 10 | DTEAHNVWATHACVPTDPNPQEVELKNVTENFNMWENNMVEQMHEDIISLWDQSLKPCVK | 120 |
| 11 | DTEAHNVWATHACVPTDPNPQEVELKNVTENFNMWENNMVEQMHEDIISLWDQSLKPCVK | 120 |
| 12 | DTEAHNVWATHACVPTDPNPQEVELKNVTENFNMWENNMVEQMHEDIISLWDQSLKPCVK | 120 |
| 13 | DTEAHNVWATHACVPTDPNPQEVELKNVTENFNMWENNMVEQMHEDIISLWDQSLKPCVK | 120 |
| 14 | DTEAHNVWATHACVPTDPNPQEVELKNVTENFNMWENNMVEQMHEDIISLWDQSLKPCVK | 120 |
| 15 | DTEAHNVWATHACVPTDPNPQEVELKNVTENFNMWENNMVEQMHEDIISLWDQSLKPCVK | 120 |
| 16 | DTEAHNVWATHACVPTDPNPQEVELKNVTENFNMWENNMVEQMHEDIISLWDQSLKPCVK | 120 |
| 17 | DTEAHNVWATHACVPTDPNPQEVELKNVTENFNMWENNMVEQMHEDIISLWDQSLKPCVK | 120 |
| 18 | DTEAHNVWATHACVPTDPNPQEVELKNVTENFNMWENNMVEQMHEDIISLWDQSLKPCVK | 120 |
| 19 | DTEAHNVWATHACVPTDPNPQEVELKNVTENFNMWENNMVEQMHEDIISLWDQSLKPCVK | 120 |
| 20 | DTEAHNVWATHACVPTDPNPQEVELKNVTENFNMWENNMVEQMHEDIISLWDQSLKPCVK | 120 |
| 21 | DTEAHNVWATHACVPTDPNPQEVELKNVTENFNMWENNMVEQMHEDIISLWDQSLKPCVK | 120 |
| 22 | DTEAHNVWATHACVPTDPNPQEVELKNVTENFNMWENNMVEQMHEDIISLWDQSLKPCVK | 120 |
| 23 | DTEAHNVWATHACVPTDPNPQEVELKNVTENFNMWENNMVEQMHEDIISLWDQSLKPCVK | 120 |
| 24 | DTEAHNVWATHACVPTDPNPQEVELKNVTENFNMWENNMVEQMHEDIISLWDQSLKPCVK | 120 |
| 25 | DTEAHNVWATHACVPTDPNPQEVELKNVTENFNMWENNMVEQMHEDIISLWDQSLKPCVK | 120 |
| 26 | DTEAHNVWATHACVPTDPNPQEVELKNVTENFNMWENNMVEQMHEDIISLWDQSLKPCVK | 120 |
| 27 | DTEAHNVWATHACVPTDPNPQEVELKNVTENFNMWENNMVEQMHEDIISLWDQSLKPCVK | 120 |
| 28 | DTEAHNVWATHACVPTDPNPQEVELKNVTENFNMWENNMVEQMHEDIISLWDQSLKPCVK | 120 |
| 29 | DTEAHNVWATHACVPTDPNPQEVELKNVTENFNMWENNMVEQMHEDIISLWDQSLKPCVK | 120 |
| 30 | DTEAHNVWATHACVPTDPNPQEVELKNVTENFNMWENNMVEQMHEDIISLWDQSLKPCVK | 120 |
| 31 | DTEAHNVWATHACVPTDPNPQEVELKNVTENFNMWENNMVEQMHEDIISLWDQSLKPCVK | 120 |
| 32 | DTEAHNVWATHACVPTDPNPQEVELKNVTENFNMWENNMVEQMHEDIISLWDQSLKPCVK | 120 |
| 33 | DTEAHNVWATHACVPTDPNPQEVELKNVTENFNMWENNMVEQMHEDIISLWDQSLKPCVK | 120 |
| 34 | --------ATHACVPTDPNPQEVELKNVTENFNMWENNMVEQMHEDIISLWDQSLKPCVK | 97 |
| 35 | DTEAHNVWATHACVPTDPNPQEVELKNVTENFNMWENNMVEQMHEDIISLWDQSLKPCVK | 120 |

| | | |
|---|---|---|
| | **LTPLCVTLNCTDLEKVTNTT---NSTEKMMEKGEVKNCSFKITTDVKDRTRKEYALFYKL** | |
| 1 | LTPLCVTLNCTDLGNVTNTT---NSNGEMMEKGEVKNCSFKITTDIKDRTRKEYALFYKL | 177 |
| 2 | LTPLCVTLNCTDLEKVTNTT---NSTEKMMEKGEVKNCSFKITTDVKDRTRKEYALFYKL | 177 |
| 3 | LTPLCVTLNCTDLEKVTNTT---NSTEKMMEKGEVKNCSFKITTDVKDRTRKEYALFYKL | 177 |
| 4 | LTPLCVTLNCTDLEKVTNTT---NSTEKMMEKGEVKNCSFKITTDVKDRTRKEYALFYKL | 177 |
| 5 | LTPLCVTLNCTDLEKVANTT---NSTEKMMEKGEVKNCSFKITTDVKDRTRKEYALFYKL | 177 |
| 6 | LTPLCVTLNCTDLEKVTNTT---NSTEKMMEKGEVKNCSFKITTDVKDRTRKEYALFYKL | 177 |
| 7 | LTPLCVTLNCTDLENVTDTT---NSTEKMMEKGEVKNCSFKITTDLKDRTRKEYALFYKL | 177 |
| 8 | LTPLCVTLNCTDLEKVTNTT---NSTEKMMEKGEVKNCSFKITTDVKDRTRKEYALFYKL | 177 |
| 9 | LTPLCVTLNCTDLGNVTTTT---NSNGTLMEKGEVKNCSFKITTDTKDRTRKEYALFYKL | 177 |
| 10 | LTPLCVTLNCTDLEKVTNTT---NSTEKMMEKGEVKNCSFKITTDVKDRTRKEYALFYKL | 177 |
| 11 | LTPLCVTLNCTDLEKVTNTT---NSTEKMMEKGEVKNCSFKITTDVKDRTRKEYALFYKL | 177 |
| 12 | LTPLCVTLNCTDLEKVTNTT---NSTEKMMEKGEVKNCSFKITTDVKDRTRKEYALFYKL | 177 |
| 13 | LTPLCVTLNCTDLEKVTNTT---NSTEKMMEKGEVKNCSFKITTDMKDRTRKEYALFYKL | 177 |
| 14 | LTPLCVTLNCTDLEKVTNTT---NSTEKMMEKGEVKNCSFKITTDVKDRTRKEYALFYKL | 177 |
| 15 | LTPLCVTLNCTDLGNVTATT---NSNGTLMEKGEVKNCSFKITTDIKDRTRKEYALFYKL | 177 |
| 16 | LTPLCVTLNCTDLEKVTNTT---NSTEKMMEKGEVKNCSFKITTDVKDRTRKEYALFYKL | 177 |
| 17 | LTPLCVTLNCTDLEKVTNTT---NSTEKMMEKGEVKNCSFRITTDIKDRTRKEYALFYKL | 177 |
| 18 | LTPLCVTLNCTDLGNVTTTT---NSNGTLMEKGEVKNCSFKITTEIKDRTRKEYALFYKL | 177 |
| 19 | LTPLCVTLNCTDLGNVTTTT---NSNGTLMEKGEVKNCSFKITTDVKDRTRKEYALF--- | 174 |
| 20 | LTPLCVTLNCTDLEKVTNTT---NSTEKMMEKGEVKNCSFKITTDVKDRTRKEYALFYKL | 177 |
| 21 | LTPLCVTLNCTDLEKVTNTT---NSTEKMMEKGEVKNCSFKITTDRKDRTRKEYALFYKL | 177 |
| 22 | LTPLCVTLNCTDLGNVTTTT---NSNGTLMEKGEVKNCSFKITTEIKDRTRKEYALFYKL | 177 |
| 23 | LTPLCVTLNCTDLEKVTNTT---NSTEKMMEKGEVKNCSFKITTDVKDRTRKEYALFYKL | 177 |
| 24 | LTPLCVTLNCTDLEKVTNTT---NSNETMMEKGEVKNCSFKITTDIKDRTRKEYALFYKL | 177 |
| 25 | LTPLCVTLNCTDLEKVTNTT---NSTEKMMEKGEVKNCSFKITTDVKDRTRKEYALFYKL | 177 |
| 26 | LTPLCVTLNCTDLGNVTTTT---NSNGTLMEKGEVKNCSFKITTDTKDRTRKEYALFYKL | 177 |
| 27 | LTPLCVTLNCTDLGNVTTTT---NSNGTLMEKGEVKNCSFKITTEIKDRTRKEYALFYKL | 177 |
| 28 | LTPLCVTLNCTDLEKVTNTT---NSTEKMMEKGEVKNCSFKITTDVKDRTRKEYALFYKL | 177 |
| 29 | LTPLCVTLNCTDLEKVTNTT---NSTEKMMEKGEVKNCSFKITTDVKDRTRKEYALFYKL | 177 |
| 30 | LTPLCVTLNCTDLEKVTNTT---NSTEKMMEKGEVKNCSFKITTDVKDRTRKEYALFYKL | 177 |
| 31 | LTPLCVTLNCTDLEKVTNTT---NSTEKMMEKGEVKNCSFKITTDVKDRTRKEYALFYKL | 177 |
| 32 | LTPLCVTLNCTDLEKVTNTT---NSTEKMMEKGEVKNCSFKITTDVKDRTRKEYALFYKL | 177 |
| 33 | LTPLCVTLNCTDLEKVTNTT---NSTEKMMEKGEVKNCSFKITTDVKDRTRKEYALFYKL | 177 |
| 34 | LTPLCVTLNCTDLGNVTTTT---NSNGTLMEKGEVKNCSFKITTDTKDRTRKEYALFYKL | 154 |
| 35 | LTPLCVTLNCTDLEKVTNTT---NSTEKMMEKGEVKNCSFKITTDVKDRTRKEYALFYKL | 177 |

| | Sequence | Length |
|---|---|---|
|  | DVVPIND-------TRYRLVSCNTSVITQACPKVSFEPIPIHYCAPAGFAILKCNDKQFI |  |
| 1 | DVVPIND-------TRYRLVSCNTSVITQACPKVSFEPIPIHYCAPAGFAILKCNDKQFI | 230 |
| 2 | DVVPIND-------TRYRLVSCNTSVITQACPKVSFEPIPIHYCAPAGFAILKCNDKQFI | 230 |
| 3 | DVVPIND-------TRYRLVSCNTSVITQACPKVSFEPIPIHYCAPAGFAILKCNDKQFI | 230 |
| 4 | DVVPIND-------TRYRLVSCNTSVITQACPKVSFEPIPIHYCAPAGFAILKCNDKQFI | 230 |
| 5 | DVVPIND-------TRYRLVSCNTSVITQACPKVSFEPIPIHYCAPAGFAILKCNDKQFI | 230 |
| 6 | DVVPIND-------TRYRLVSCNTSVITQACPKVSFEPIPIHYCAPAGFAILKCNDKQFN | 230 |
| 7 | DVVPIND-------TRYRLVSCNTSVITQACPKVSFEPIPIHYCAPAGFAILKCNDKQFI | 230 |
| 8 | DVVPIND-------TRYRLVSCNTSVITQACPKVSFEPIPIHYCAPAGFAILKCNDKQFI | 230 |
| 9 | DVVPIND-------TRYRLVSCNTSVITQACPKVSFEPIPIHYCAPAGFAILKCNDKQFI | 230 |
| 10 | DVVPIND-------TRYRLVSCNTSVITQACPKVSFEPIPIHYCAPAGFAILKCNDKQFI | 230 |
| 11 | DVVPIND-------TRYRLVSCNTSVITQACPKVSFEPIPIHYCAPAGFAILKCNDKQFI | 230 |
| 12 | DVVPIND-------TRYRLVSCNTSVITQACPKVSFEPIPIHYCAPAGFAILKCNDKQFI | 230 |
| 13 | DVVPIND-------TRYRLVSCNTSVITQACPKVSFEPIPIHYCAPAGFAILKCNDKQFI | 230 |
| 14 | DVVPIND-------TRYRLVSCNTSVITQACPKVSFEPIPIHYCAPAGFAILKCNDKQFI | 230 |
| 15 | DVVPIND-------TRYRLVSCNTSVITQACPKVSFEPIPIHYCAPAGFAILKCNDKQFM | 230 |
| 16 | DVVPIND-------TRYRLVSCNTSVITQACPKVSFEPIPIHYCAPAGFAILKCNDKQFI | 230 |
| 17 | DVVPIND-------TRYRLVSCNTSVITQACPKVSFEPIPIHYCAPAGFAILKCNDKQFI | 230 |
| 18 | DVVPIND-------TRYRLVSCNTSVITQACPKVSFEPIPIHYCAPAGFAILKCNDKQFI | 230 |
| 19 | --------------------SCNTSVITQACPKVSFEPIPIHYCAPAGFAILKCNDKQFI | 214 |
| 20 | DVVPIND-------TRYRLVSCNTSVITQACPKVSFEPIPIHYCAPAGFAILKCNDKQFI | 230 |
| 21 | DVVPIND-------TRYRLVSCNTSVITQACPKVSFEPIPIHYCAPAGFAILKCNDKQFM | 230 |
| 22 | DVVPIND-------TRYRLVSCNTSVITQACPKVSFEPIPIHYCAPAGFAILKCNDKQFI | 230 |
| 23 | DVVPIND-------TRYRLVSCNTSVITQACPKVSFEPIPIHYCAPAGFAILKCNDKQFI | 230 |
| 24 | DVVPIND-------TRYRLVSCNTSVITQACPKVSFEPIPIHYCAPAGFAILKCNDKQFI | 230 |
| 25 | DVVPIND-------TRYRLVSCNTSVITQACPKVSFEPIPIHYCAPAGFAILKCNDKQFI | 230 |
| 26 | DVVPIND-------TRYRLVSCNTSVITQACPKVSFEPIPIHYCAPAGFAILKCNDKQFI | 230 |
| 27 | DVVPIND-------TRYRLVSCNTSVITQACPKVSFEPIPIHYCAPAGFAILKCNDKQFI | 230 |
| 28 | DVVPIND-------TRYRLVSCNTSVITQACPKVSFEPIPIHYCAPAGFAILKCNDKQFI | 230 |
| 29 | DVVPIND-------TRYRLVSCNTSVITQACPKVSFEPIPIHYCAPAGFAILKCNDKQFI | 230 |
| 30 | DVVPIND-------TRYRLVSCNTSVITQACPKVSFEPIPIHYCAPAGFAILKCNDKQFI | 230 |
| 31 | DVVPIND-------TRYRLVSCNTSVITQACPKVSFEPIPIHYCAPAGFAILKCNDKQFI | 230 |
| 32 | DVVPIND-------TRYRLVSCNTSVITQACPKVSFEPIPIHYCAPAGFAILKCNDKQFI | 230 |
| 33 | DVVPIND-------TRYRLVSCNTSVITQACPKVSFEPIPIHYCAPAGFAILKCNDKQFI | 230 |
| 34 | DVVPIND-------TRYRLVSCNTSVITQACPKVSFEPIPIHYCAPAGFAILKCNDKQFI | 207 |
| 35 | DVVPIND-------TRYRLVSCNTSVITQACPKVSFEPIPIHYCAPAGFAILKCNDKQFI | 230 |

**GTGPCTNVSTVQCTHGIRPVVSTQLLLNGSLAEEEVVIRSVNFSDNAKTIIVQLNKSVEI**

| | | |
|---|---|---|
| 1 | GTGPCTNVSTVQCTHGIRPVVSTQLLLNGSLAEEEVVIRSVNFSDNAKTIIVQLNKSVEI | 290 |
| 2 | GTGPCTNVSTVQCTHGIRPVVSTQLLLNGSLAEEEVVIRSVNFRDNAKTIIVQLNKSVEI | 290 |
| 3 | GTGPCTNVSTVQCTHGIRPVVSTQLLLNGSLAEEEVVIRSVNFRDNAKTIIVQLNKSVEI | 290 |
| 4 | GTGPCTNVSTVQCTHGIRPVVSTQLLLNGSLAEEEVVIRSVNFRDNAKTIIVQLNKSVEI | 290 |
| 5 | GTGPCTNVSTVQCTHGIRPVVSTQLLLNGSLAEEEVVIRSVNFSDNAKTIIVQLNKSVEI | 290 |
| 6 | GTGPCTNVSTVQCTHGIRPVVSTQLLLNGSLAEEEVVIRSVNFSDNAKTIIVQLNKSVEI | 290 |
| 7 | GTGPCTNVSTVQCTHGIRPVVSTQLLLNGSLAEEEVVIRSVNFRDNAKTIIVQLNKSVEI | 290 |
| 8 | GTGPCTNVSTVQCTHGIRPVVSTQLLLNGSLAEEEVVIRSVNFRDNAKTIIVQLNKSVEI | 290 |
| 9 | GTGPCTNVSTVQCTHGIRPVVSTQLLLNGSLAEEEVVIRSVNFSDNAKTIIVQLNKSVEI | 290 |
| 10 | GTGPCTNVSTVQCTHGIRPVVSTQLLLNGSLAEEEVVIRSVNFSDNAKTIIVQLNKSVEI | 290 |
| 11 | GTGPCTNVSTVQCTHGIRPVVSTQLLLNGSLAEEEVVIRSVNFSDNAKTIIVQLNKSVEI | 290 |
| 12 | GTGPCTNVSTVQCTHGIRPVVSTQLLLNGSLAEEEVVIRSVNFSDNAKTIIVQLNKSVEI | 290 |
| 13 | GTGPCTNVSTVQCTHGIRPVVSTQLLLNGSLAEEEVVIRSVNFRDNAKTIIVQLNKSVEI | 290 |
| 14 | GTGPCTNVSTVQCTHGIRPVVSTQLLLNGSLAEEEVVIRSVNFSDNAKTIIVQLNKSVEI | 290 |
| 15 | GTGPCTNVSTVQCTHGIRPVVSTQLLLNGSLAEEEVVIRSVNFSDNAKTIIVQLNKSVEI | 290 |
| 16 | GTGPCTNVSTVQCTHGIRPVVSTQLLLNGSLAEEEVVIRSVNFRDNAKTIIVQLNKSVEI | 290 |
| 17 | GTGPCTNVSTVQCTHGIRPVVSTQLLLNGSLAEEEVVIRSVNFSDNAKTIIVQLNKSVEI | 290 |
| 18 | GTGPCTNVSTVQCTHGIRPVVSTQLLLNGSLAEEEVVIRSVNFRDNAKTIIVQLNKSVEI | 290 |
| 19 | GTGPCTNVSTVQCTHGIRPVVSTQLLLNGSLAEEEVVIRSVNFSDNAKTIIVQLNKSVEI | 274 |
| 20 | GTGPCTNVSTVQCTHGIRPVVSTQLLLNGSLAEEEVVIRSVNFRDNAKTIIVQLNKSVEI | 290 |
| 21 | GTGPCTNVSTVQCTHGIRPVVSTQLLLNGSLAEEEVVIRSVNFSDNAKTIIVQLNKSVEI | 290 |
| 22 | GTGPCTNVSTVQCTHGIRPVVSTQLLLNGSLAEEEVVIRSVNFRDNAKTIIVQLNKSVEI | 290 |
| 23 | GTGPCTNVSTVQCTHGIRPVVSTQLLLNGSLAEEEVVIRSVNFRDNAKTIIVQLNKSVEI | 290 |
| 24 | GTGPCTNVSTVQCTHGIRPVVSTQLLLNGSLAEEEVVIRSVNFSDNAKTIIVQLNKSVEI | 290 |
| 25 | GTGPCTNVSTVQCTHGIRPVVSTQLLLNGSLAEEEVVIRSVNFRDNAKTIIVQLNKSVEI | 290 |
| 26 | GTGPCTNVSTVQCTHGIRPVVSTQLLLNGSLAEEEVVIRSVNFSDNAKTIIVQLNKSVEI | 290 |
| 27 | GTGPCTNVSTVQCTHGIRPVVSTQLLLNGSLAEEEVVIRSVNFRDNAKTIIVQLNKSVEI | 290 |
| 28 | GTGPCTNVSTVQCTHGIRPVVSTQLLLNGSLAEEEVVIRSVNFRDNAKTIIVQLNKSVEI | 290 |
| 29 | GTGPCTNVSTV--------------------------IRSVNFSDNAKTIIVQLNKSVEI | 264 |
| 30 | GTGPCTNVSTVQCTHGIRPVVSTQLLLNGSLAEEEVVIRSVNFRDNAKTIIVQLNKSVEI | 290 |
| 31 | GTGPCTNVSTVQCTHGIRPVVSTQLLLNGSLAEEEVVIRSVNFSDNAKTIIVQLNKSVEI | 290 |
| 32 | GTGPCTNVSTVQCTHGIRPVVSTQLLLNGSLAEEEVVIRSVNFSDNAKTIIVQLNKSVEI | 290 |
| 33 | GTGPCTNVSTVQCTHGIRPVVSTQLLLNGSLAEEEVVIRSVNFSDNAKTIIVQLNKSVEI | 290 |
| 34 | GTGPCTNVSTVQCTHGIRPVVSTQLLLNGSLAEEEVVIRSVNFSDNAKTIIVQLNKSVEI | 267 |
| 35 | GTGPCTNVSTVQCTHGIRPVVSTQLLLNGSLAEEEVVIRSVNFRDNAKTIIVQLNKSVEI | 290 |

| | | |
|---|---|---|
| | **NCTRPHNNTRKSIPMGPGKAFYARGDIIGDIRKAYCEINGTEWHSTLKLVVEKLREQY-N** | |
| 1 | TCTRPNNNTRKSIPMGPGKAFYARGDITGDIRKAYCEINGTEWHSTLKLVVEKLREQY-N | 349 |
| 2 | NCTRPHNNTRKSIPMGPGKAFYARGDIIGDIRKAYCEVNGTEWHSTLKLVVEKLREQY-N | 349 |
| 3 | NCTRPHNNTRKSIPMGPGKAFYARGDIIGDIRKAYCEINGTEWHSTLKLVVEKLREQY-N | 349 |
| 4 | NCTRPHNNTRKSIPMGPGKAFYARGDIIGDIRKAYCEINGTEWHSTLKLVVEKLREQY-N | 349 |
| 5 | NCTRPHNNTRKSIPMGPGKAFYARGDIIGDIRKAYCEINGTEWHSTLKLVVEKLREKY-N | 349 |
| 6 | NCTRPHNNTRKSIPMGPGKAFYARGDIIGDIRKAYCNINGTEWHSTLKLVVEKLREQY-N | 349 |
| 7 | NCARPHNNTRKSIPMGPGKAFYARGDITGDIRKAYCEINGTEWHSTLKLVVEKLREQY-N | 349 |
| 8 | NCTRPHNNTRKSIPMGPGKAFYARGDIIGDIRKAYCEINGTEWHSTLKLVVEKLREQY-N | 349 |
| 9 | NCTRPNNNTRKSIPMGPGKAFYARGDIIGDIRKASCELNGTEWHSTLKLVVEKLRKQY-N | 349 |
| 10 | NCTRPHNNTRKSIPMGPGKAFYARGDIIGDIRKAYCEINGTEWHSTLKLVVGKLREKY-N | 349 |
| 11 | NCTRPNNNTRKSIPMGPGKAFYARGDIIGDIRKASCKINGTEWHSTLKLVVEKLREQY-N | 349 |
| 12 | NCTRPHNNTRKSIPMGPGKAFYARGDIIGDIRKAYCEINGTEWHSTLKLVVGKLREKY-N | 349 |
| 13 | NCTRPHNNTRKSIPMGPGKAFYARGDIIGDIRKAYCEINGTEWHSTLKLVVEKLREQY-N | 349 |
| 14 | NCTRPHNNTRKSIPMGPGKAFYARGDIIGDIRKAYCEINGTEWHSTLKLVVGKLREKY-N | 349 |
| 15 | TCTRPNNNTRKSIPMGPGKAFYARGDIIGDIRKASCKINGTEWHSTLKLVVEKLREQY-N | 349 |
| 16 | NCTRPHNNTRKSIPMGPGKAFYARGDIIGDIRKAYCEINGTEWHSTLKLVVEKLREQY-N | 349 |
| 17 | NCTRPHNNTRKSIPMGPGKAFYARGDIIGDIRKAYCEINGTEWHSTLKLVVEKLREQY-N | 349 |
| 18 | NCTRPHNNTRKSIPMGPGKAFYARGDIIGDIRKAYCTINGTEWHSTLKLVVEKLREQY-N | 349 |
| 19 | NCTRPHNNTRKSIPMGPGKAFYARGDIIGDIRKAYCKINGTEWHSTLKLVVEKLREQY-N | 333 |
| 20 | NCTRPHNNTRKSIPMGPGKAFYARGDIIGDIRKAYCEINGTEWHSTLKLVVEKLREQY-N | 349 |
| 21 | NCTRPHNNTRKSIPMGPGKAFYARGDITGDIRKAYCEINGTEWHSTLKLVVEKLREQY-N | 349 |
| 22 | NCTRPHNNTRKSIPMGPGKAFYARGDIIGNIRKAYCEINGTEWHSTLKLVVEKLREQY-N | 349 |
| 23 | NCTRPHNNTRKSIPMGPGKAFYARGDIIGDIRKAYCEINGTEWHSTLKLVVEKLREQY-N | 349 |
| 24 | TCTRPNNNTRKSIPMGPGKAFYARGDIIGDIRKASCKINGTEWHSTLKLVVEKLREKY-N | 349 |
| 25 | NCTRPHNNTRKSIPMGPGKAFYARGDIIGDIRKAYCEINGTEWHSTLKLVVEKLREQY-N | 349 |
| 26 | NCTRPNNNTRKSIPMGPGKAFYARGDIIGDIRKASCELDGTEWHSTLKLVVEKLRKQY-N | 349 |
| 27 | NCTRPHNNTRKSIPMGPGKAFYARGDIIGDIRKAYCEINGAEWHSTLKLVVEKLKEQY-N | 349 |
| 28 | NCTRPHNNTRKSIPMGPGKAFYARGDIIGDIRKAYCEINGTEWHSTLKLVVEKLREQY-N | 349 |
| 29 | NCTRPNNNTRKSIPMGPGKAFYARGDIIGDIRKASCKINGTEWHSTLKLVVEKLREQY-N | 323 |
| 30 | NCTRPHNNTRKSIPMGPGKAFYARGDIIGDIRKAYCEVNGTEWHSTLKLVVEKLREQY-N | 349 |
| 31 | NCTRPHNNTRKSIPMGPGKAFYARGDIIGDIRKAYCAINGTEWHSTLKLVVEKLKEQY-N | 349 |
| 32 | NCTRPNNNTRKSIPMGPGKAFYARGDIIGDIRKASCKINGTEWHSTLKLVVEKLREQY-N | 349 |
| 33 | NCTRPNNNTRKSIPMGPGKAFYARGDIIGDIRKASCKINGTEWHSTFKLVVEKLRAQY-N | 349 |
| 34 | NCTRPNNNTRKSIPMGPGKAFYARGDIIGDIRKASCELDGTEWHSTLKLVVEKLRKQY-N | 326 |
| 35 | NCTRPHNNTRKSIPMGPGKAFYARGDIIGDIRKAYCEINGTEWHSTLKLVVEKLREQY-N | 349 |

| | Sequence | |
|---|---|---|
| | KTIVFNRSSGGDPEIVMYSFNCGGEFFYCNSTKLFNSTWPWND-TKGSHDTNGTLILPCR | |
| 1 | KTIVFNRSSGGDPEIVMYSFNCGGEFFYCNSTKLFNSTWPWND-TKGSHDTNGTLILPCK | 408 |
| 2 | KTIVFNRSSGGDPEIVMYSFNCGGEFFYCNSTKLFNSTWPWND-TKGSHDTNGKLILPCR | 408 |
| 3 | KTIVFNRSSGGDPEIVMYSFNCGGEFFYCNSTKLFNSTWPWND-TKGSHDTNGKLILPCR | 408 |
| 4 | KTIVFNRSSGGDPEIVMYSFNCGGEFFYCNSTKLFNSTWPWND-TKGSHDTNGKLILPCR | 408 |
| 5 | KTIVFNRSSGGDPEIVMYSFNCGGEFFYCNSTKLFNSTWPWND-TKGSHDTNDTLMLPCK | 408 |
| 6 | KTIVFNRSSGGDPEIVMYSFNCGGEFFYCNSTKLFNSTWPWND-TKGSHDTNDTLMLPCK | 408 |
| 7 | KTIVFNRSSGGDPEIVMYSFNCGGEFFYCNSTKLFNSTWPWND-TKGSHDTNGKLILPCK | 408 |
| 8 | KTIVFNRSSGGDPEIVMYSFNCGGEFFYCNSTKLFNSTWPWND-TKGSHDTNGKLILPCR | 408 |
| 9 | KTIVFNRSSGGDPEIVMYSFNCGGEFFYCNSTKLFNSTWPWND-TKGSHDTNGTLILPCR | 408 |
| 10 | KTIVFNRSSGGDPEIVMYSFNCGGEFFYCNSTKLFNSTWPWND-TKGSHDTNDTLMLPCK | 408 |
| 11 | KTIVFNRSSGGDPEIVMYSFNCGGEFFYCNSTKLFNSTWPWND-TKGSHDTNDTLMLPCK | 408 |
| 12 | KTIVFNRSSGGDPEIVMYSFNCGGEFFYCNSTKLFNSTWPWND-TKGSHDTNDTLMLPCK | 408 |
| 13 | KTIVFNRSSGGDPEIVMYSFNCGGEFFYCNSTKLFNSTWPWND-TKGSHDTNDTLMLPCK | 408 |
| 14 | KTIVFNRSSGGDPEIVMYSFNCGGEFFYCNSTKLFNSTWPWSD-TKGSHDTNDTLMLPCK | 408 |
| 15 | KTIVFNRSSGGDPEIVMYSFNCGGEFFYCNSTKLFNSTWPWND-TKGSHDTNDTLMLPCK | 408 |
| 16 | KTIVFNRSSGGDPEIVMYSFNCGGEFFYCNSTKLFNSTWPWND-TKGSHDTNGKLILPCR | 408 |
| 17 | KTIVFNRSSGGDPEIVMYSFNCGGEFFYCNSTKLFNSTWPWND-TKGSHDTNDTLMLPCK | 408 |
| 18 | KTIVFNRSSGGDPEIVMYSFNCGGEFFYCNSTKLFNSTWPWND-TKGSHDTNDTLMLPCK | 408 |
| 19 | KTIVFNRSSGGDPEIVMYSFNCGGEFFYCNSTKLFNSTWPWND-TKGSHDTNDTLMLPCR | 392 |
| 20 | KTIVFNRSSGGDPEIVMYSFNCGGEFFYCNSTKLFNSTWPWND-TKGSHDTNGKLILPCR | 408 |
| 21 | KTIVFNRSSGGDPEIVMYSFNCGGEFFYCNSTKLFNSTWPWND-TKGSHDTNGKLILPCR | 408 |
| 22 | KTIVFNRSSGGDPEIVMYSFNCGGEFFYCNSTKLFNSTWPWND-TKGSHDTNGRLILPCR | 408 |
| 23 | KTIVFNRSSGGDPEIVMYSFNCGGEFFYCNSTKLFNSTWPWND-TKGSHDTNGKLILPCR | 408 |
| 24 | KTIVFNRSSGGDPEIVMYSFNCGGEFFYCNSTKLFNSTWPWSD-TKGSHDTNDTLMLPCR | 408 |
| 25 | KTIVFNRSSGGDPEIVMYSFNCGGEFFYCNSTKLFNSTWPWND-TKGSHDTNGSLILPCR | 408 |
| 26 | KTIVFNRSSGGDPEIVMYSFNCGGEFFYCNSTKLFNSTWPWND-TKGSHDTNGTLILPCR | 408 |
| 27 | KTIVFNRSSGGDPEIVMYSFNCGGEFFYCNSTKLFNSTWPWND-TKGSHDINGTLILPCK | 408 |
| 28 | KTIVFNRSSGGDPEIVMYSFNCGGEFFYCNSTKLFNSTWPWND-TKGSHDTNGKLILPCR | 408 |
| 29 | KTIVFNRSSGGDPEIVMYSFNCGGEFFYCNSTKLFNSTWPWND-TKGSHDTNDTLMLPCK | 382 |
| 30 | KTIVFNRSSGGDPEIVMYSFNCGGEFFYCNSTKLFNSTWPWND-TKGSHDTNGKLILPCR | 408 |
| 31 | KTIVFNRSSGGDPEIVMYSFNCGGEFFYCNSTKLFNSTWPWND-TKGSHDTNGTLILPCK | 408 |
| 32 | KTIVFNRSSGGDPEIVMYSFNCGGEFFYCNSTKLFNSTWPWND-TKGSHDTNDTLMLPCK | 408 |
| 33 | KTIVFSRSSGGDPEIVMYSFNCGGDFFYCNSTQLFNSTWPWND-TKGSHDTNDTLMLPCK | 408 |
| 34 | KTIVFNRSSGGDPEIVMYSFNCGGEFFYCNSTKLFNSTWPWND-TKGSHDTNGTLILPCR | 385 |
| 35 | KTIVFNRSSGGDPEIVMYSFNCGGEFFYCNSTKLFNSTWPWND-TKGSHDTNGKLILPCR | 408 |

```
     IKQIINMWQGVGKAMYAPPIEGXIRCSSNITGLLLTRDGG-YESNETDEIFRPGGGDMRD
1    IKQIINMWQGVGKAMYAPPIEGKIRCSSNITGLLLTRDGG-YESNETDEIFRPGGGDMRD  467
2    IKQIINMWQGVGKAMYAPPIEGEIRCSSNITGLLLTRDGG-YESNKTDEIFRPGGGDMRD  467
3    IKQIINMWQGVGKAMYAPPIEGEIRCSSNITGLLLTRDGG-YESNKTDEIFRPGGGDMRD  467
4    IKQIINMWQGVGKAMYAPPIEGEIRCSSNITGLLLTRDGG-YESNKTDEIFRPGGGDMRD  467
5    IKQIINMWQGVGKAMYAPPIEGQIRCSSNITGLLLTRDGG-YESNETDEIFRPGGGDMRD  467
6    IKQIINMWQGVGKAMYAPPIEGQIRCSSNITGLLLTRDGG-YESNETDEIFRPGGGDMRD  467
7    IKQIINMWQGVGKAMYAPPIEGKIRCSSNITGLLLTRDGG-YESNETDEIFRPGGGDMRD  467
8    IKQIINMWQGVGKAMYAPPIEGEIRCSSNITGLLLTRDGG-YESNKTDEIFRPGGGDMRD  467
9    IKQIINMWQGVGKAMYAPPIEGEIRCSSNITGLLLTRDGG-YESNETDEIFRPGGGDMRD  467
10   IKQIINMWQGVGKAMYAPPIEGQIRCSSNITGLLLTRDGG-YESNETDEIFRPGGGDMRD  467
11   IKQIINMWQGVGKAMYAPPIEGIIRCSSNITGLLLTRDGG-YESNETDEIFRPGGGDMRD  467
12   IKQIINMWQGVGKAMYAPPIEGQIRCSSNITGLLLTRDGG-YESNETDEIFRPGGGDMRD  467
13   IKQIINMWQGVGKAMYAPPIEGQIRCSSNITGLLLTRDGG-YESNETDEIFRPGGGDMRD  467
14   IKQIINMWQGVGKAMYAPPIEGQIRCSSNITGLLLTRDGG-YESNETDEIFRPGGGDMRD  467
15   IKQI-NMWQGVGKAMYAPPIEGIISCSSNITGLLLTRDGG-YESNETDEIFRPGGGDMRD  466
16   IKQIINMWQGVGKAMYAPPIEGEIRCSSNITGLLLTRDGG-YESNKTDEIFRPGGGDMRD  467
17   IKQIINMWQGVGKAMYAPPIEGQIRCSSNITGLLLTRDGG-YESNETDEIFRPGGGDMRD  467
18   IKQIINMWQGVGKAMYAPPIEGQIRCSSNITGLLLTRDGG-YESNETDEIFRPGGGDMRD  467
19   IKQIINMWQGVGKAMYAPPIEGIIRCSSNITGLLLTRDGG-YESNETDEIFRPGGGDMRD  451
20   IKQIINMWQGVGKAMYAPPIEGEIRCSSNITGLLLTRDGG-YESNKTDEIFRPGGGDMRD  467
21   IKQIINMWQGVGKAMYAPPIEGQIRCSSNITGLLLTRDGG-YESNETDEIFRPGGGDMRD  467
22   IKQIINMWQGVGKAMYAPPIEGEIRCSSNITGLLLTRDGG-YESNKTDEIFRPGGGDMRD  467
23   IKQIINMWQGVGKAMYAPPIEGEIRCSSNITGLLLTRDGG-YESNKTDEIFRPGGGDMRD  467
24   IKQIINMWQGVGKAMYAPPIEGIIRCSSNITGLLLTRDGG-YESNETDEIFRPGGGDMRD  467
25   IKQIINMWQGVGKAMYAPPIEGEIRCSSNITGLLLTRDGG-YESNKTDEIFRPGGGDMRD  467
26   IKQIINMWQGVGKAMYAPPIEGEIRCSSNITGLLLTRDGG-YESNETDEIFRPGGGDMRD  467
27   IKQIINMWQGVGKAMYAPPIEGQIRCSSNITGLLLTRDGG-YESNKTDEIFRPGGGDMRD  467
28   IKQIINMWQGVGKAMYAPPIEGEIRCSSNITGLLLTRDGG-YESNKTDEIFRPGGGDMRD  467
29   IKQIINMWQGVGKAMYAPPIEGIIRCSSNITGLLLTRDGG-YESNETDEIFRPGGGDMRD  441
30   IKQIINMWQGVGKAMYAPPIEGEIRCSSNITGLLLTRDGG-YESNKTDEIFRPGGGDMRD  467
31   IKQIINMWQGVGKAMYAPPIEGQIRCSSNITGLLLTRDGG-YESNKTDEIFRPGGGDMRD  467
32   IKQIINMWQGVGKAMYAPPIEGIIRCSSNITGLLLTRDGG-YESNETDEIFRPGGGDRRD  467
33   IKQIINMWQGVGKAMYAPPIEGIIRCSSNITGLLLTRDGG-YESNETDEIFRPGGGDMRD  467
34   IKQIINMWQGVGKAMYAPPIEGEIRCSSNITGLLLTRDGG-YESNETDEIFRPGGGDMRD  444
35   IKQIINMWQGVGKAMYAPPIEGEIRCSSNITGLLLTRDGG-YESNKTDEIFRPGGGDMRD  467
```

| | Sequence | |
|---|---|---|
| | NWRSELYKYKVVKIEPLGVAPTKAKRRVVQREKRAFG-LGAVFLGFLGAAGSTMGAASIT | |
| 1 | NWRSELYKYKVVKIEPLGVAPTKAKRRVVQREKRAFG-LGAVFLGFLGAAGSTMGAASIT | 526 |
| 2 | NWRSELYKYKVVKIEPLGVAPTKAKRRVVQREKRAFG-LGAVFLGFLGAAGSTMGAASIT | 526 |
| 3 | NWRSELYKYKVVKIEPLGVAPTKAKRRVVQREKRAFG-LGAVFLGFLGAAGSTMGAASIT | 526 |
| 4 | NWRSELYKYKVVKIEPLGVAPTKAKRRVVQREKRAFG-LGAVFLGFLGAAGSTMGAASIT | 526 |
| 5 | NWRSELYKYKVVKIEPLGVAPTKAKRRVVQREKRAFG-LGAVFLGFLGAAGSTMGAASIT | 526 |
| 6 | NWRSELYKYKVVKIEPLGVAPTKAKRRVVQREKRAFG-LGAVFLGFLGAAGSTMGAASIT | 526 |
| 7 | NWRSELYKYKVVKIEPLGVAPTKAKRRVVQREKRAFG-LGAVFLGFLGAAGSTMGAASIT | 526 |
| 8 | NWRSELYKYKVVKIEPLGVAPTKAKRRVVQREKRAFG-LGAVFLGFLGAAGSTMGAASIT | 526 |
| 9 | NWRSELYKYKVVKIEPLGVAPTKAKRRVVQREKRAFG-LGAVFLGFLGAAGSTMGAASIT | 526 |
| 10 | NWRSELYKYKVVKIEPLGVAPTKAKRRVVQKEKRAFG-LGAVFLGFLGAAGSTMGAASIT | 526 |
| 11 | NWRSELYKYKVVKIEPLGVAPTKAKRRVVQREKRAFG-LGAVFLGFLGAAGSTMGAASIT | 526 |
| 12 | NWRSELYKYKVVKIEPLGVAPTKAKRRVVQREKRAFG-LGAVFLGFLGAAGSTMGAASIT | 526 |
| 13 | NWRSELYKYKVVKIEPLGVAPTKAKRRVVQREKRAFG-LGAVFLGFLGAAGSTMGAASIT | 526 |
| 14 | NWRSELYKYKVVKIKPLGVAPTKAKRRVVQREKRAFG-LGAVFLGFLGAAGSTMGAASIT | 526 |
| 15 | NWRSELYKYKVVKIEPLGVAPTRAKRRVVQREKRAFG-LGAVFLGFLGAAGSTMGAASIT | 525 |
| 16 | NWRSELYKYKVVKIEPLGVAPTKAKRRVVQREKRAFG-LGAVFLGFLGAAGSTMGAASIT | 526 |
| 17 | NWRSELYKYKVVKIEPLGVAPTKAKRRVVQREKRAFG-LGAVFLGFLGAAGSTMGAASIT | 526 |
| 18 | NWRSELYKYKVVKIEPLGVAPTKAKRRVVQREKRAFG-LGAVFLGFLGAAGSTMGAASIT | 526 |
| 19 | NWRSELYKYKVVKIEPLGVAPTKAKRRVVQREKRAFG-LGAVFLGFLGAAGSTMGAASIT | 510 |
| 20 | NWRSELYKYKVVKIEPLGVAPTKAKRRVVQREKRAFG-LGAVFLGFLGAAGSTMGAASIT | 526 |
| 21 | NWRSELYKYKVVKIEPLGVAPTKAKRRVVQREKRAFG-LGAVFLGFLGAAGSTMGAASIT | 526 |
| 22 | NWRSELYKYKVVKIEPLGVAPTKAKRRVVQREKRAFG-LGAVFLGFLGAAGSTMGAASIT | 526 |
| 23 | NWRSELYKYKVVKIEPLGVAPTKAKRRVVQREKRAFG-LGAVFLGFLGAAGSTMGAASIT | 526 |
| 24 | NWRSELYKYKVVKIEPLGVAPTKAKRRVVQREKRAFG-LGAVFLGFLGAAGSTMGAASIT | 526 |
| 25 | NWRSELYKYKVVKIEPLGVAPTKAKRRVVQREKRAFG-LGAVFLGFLGAAGSTMGAASIT | 526 |
| 26 | NWRSELYKYKVVKIEPLGVAPTKAKRRVVQREKRAFG-LGAVFLGFLGAAGSTMGAASIT | 526 |
| 27 | NWRSELYKYKVVKIEPLGVAPTKAKRRVVQREKRAFG-LGAVFLGFLGAAGSTMGAASIT | 526 |
| 28 | NWRSELYKYKVVKIEPLGVAPTKAKRRVVQREKRAFG-LGAVFLGFLGAAGSTMGAASIT | 526 |
| 29 | NWRSELYKYKVVKIEPLGVAPTKAKRRVVQREKRAFG-LGAVFLGFLGAAGSTMGAASIT | 500 |
| 30 | NWRSELYKYKVVKIEPLGVAPTKAKRRVVQREKRAFG-LGAVFLGFLGAAGSTMGAASIT | 526 |
| 31 | NWRSELYKYKVVKIEPLGVAPTKAKRRVVQREKRAFG-LGAVFLGFLGAAGSTMGAASIT | 526 |
| 32 | KWRSELYKHKVVKIEPLGVAPTKAKRRVVQREKRAFG-LGAVFLGFLGAAGSTMGAASIT | 526 |
| 33 | NWRSELYKYKVVKIEPLGVAPTKAKRRVVQREKRAFG-LGAVFLGFLGAAGSTMGAASIT | 526 |
| 34 | NWRSELYKYKVVKIEPLGVAPTKAKRRVVQREKRAFG-LGAVFLGFLGAAGSTMGAASIT | 503 |
| 35 | NWRSELYKYKVVKIEPLGVAPTKAKRRVVQREKRAFG-LGAVFLGFLGAAGSTMGAASIT | 526 |

| | | |
|---|---|---|
| | **LTVQARQLLSGIVQQQNNLLRAIEAQQHLLQLTVWGIKQLQARVLAVERYLKDQQLLGIW** | |
| 1 | LTVQARQLLSGIVQQQNNLLRAIEAQQHLLQLTVWGIKQLQARVLAVERYLKDQQLLGIW | 586 |
| 2 | LTVQARQLLSGIVQQQNNLLRAIEAQQHLLQLTVWGIKQLQARVLAVERYLKDQQLLGIW | 586 |
| 3 | LTVQARQLLSGIVQQQNNLLRAIEAQQHLLQLTVWGIKQLQARVLAVERYLKDQQLLGIW | 586 |
| 4 | LTVQARQLLSGIVQQQNNLLRAIEAQQHLLQLTVWGIKQLQARVLAVERYLKDQQLLGIW | 586 |
| 5 | LTVQARQLLSGIVQQQNNLLRAIEAQQHLLQLTVWGIKQLQARVLAVERYLKDQQLLGIW | 586 |
| 6 | LTVQARQLLSGIVQQQNNLLRAIEAQQHLLQLTVWGIKQLQARVLAVERYLKDQQLLGIW | 586 |
| 7 | LTVQARQLLSGIVQQQNNLLRAIEAQQHLLQLTVWGIKQLQARILAVERYLKDQQLLGIW | 586 |
| 8 | LTVQARQLLSGIVQQQNNLLRAIEAQQHLLQLTVWGIKQLQARVLAVERYLKDQQLLGIW | 586 |
| 9 | LTVQARQLLSGIVQQQNNLLRAIEAQQHLLQLTVWGIKQLQARVLAVERYLKDQQLLGIW | 586 |
| 10 | LTVQARQLLSGIVQQQNNLLRAIEAQQHLLQLTVWGIKQLQARVLAVERYLKDQQLLGIW | 586 |
| 11 | LTVQARQLLSGIVQQQNNLLRAIEAQQHLLQLTVWGIKQLQARVLAVERYLKDQQLLGIW | 586 |
| 12 | LTVQARQLLSGIVQQQNNLLRAIEAQQHLLQLTVWGIKQLQARVLAVERYLKDQQLLGIW | 586 |
| 13 | LTVQARQLLSGIVQQQNNLLRAIEAQQHLLQLTVWGIKQLQARVLAVERYLKDQQLLGIW | 586 |
| 14 | LTVQARQLLSGIVQQQNNLLRAIEAQQHLLQLTVWGIKQLQARVLAVERYLKDQQLLGIW | 586 |
| 15 | LTVQARQLLSGIVQQQNNLLRAIEAQQHLLQLTVWGIKQLQARVLAVERYLKDQQLLGIW | 585 |
| 16 | LTVQARQLLSGIVQQQNNLLRAIEAQQHLLQLTVWGIKQLQARVLAVERYLKDQQLLGIW | 586 |
| 17 | LTVQARQLLSGIVQQQNNLLRAIEAQQHLLQLTVWGIKQLQARVLAVERYLKDQQLLGIW | 586 |
| 18 | LTVQARQLLSGIVQQQNNLLRAIEAQQHLLQLTVWGIKQLQARVLAVERYLKDQQLLGIW | 586 |
| 19 | LTVQARQLLSGIVQQQNNLLRAIEAQQHLLQLTVWGIKQLQARVLAVERYLKDQQLLGIW | 570 |
| 20 | LTVQARQLLSGIVQQQNNLLRAIEAQQHLLQLTVWGIKQLQARVLAVERYLKDQQLLGIW | 586 |
| 21 | LTVQARQLLSGIVQQQNNLLRAIEAQQHLLQLTVWGIKQLQARVLAVERYLKDQQLLGIW | 586 |
| 22 | LTVQARQLLSGIVQQQNNLLRAIEAQQHLLQLTVWGIKQLQARVLAVERYLKDQQLLGIW | 586 |
| 23 | LTVQARQLLSGIVQQQNNLLRAIEAQQHLLQLTVWGIKQLQARVLAVERYLKDQQLLGIW | 586 |
| 24 | LTVQARQLLSGIVQQQNNLLRAIEAQQHLLQLTVWGIKQLQARVLAVERYLKDQQLLGIW | 586 |
| 25 | LTVQARQLLSGIVQQQNNLLRAIEAQQHLLQLTVWGIKQLQARVLAVERYLKDQQLLGIW | 586 |
| 26 | LTVQARQLLSGIVQQQNNLLRAIEAQQHLLQLTVWGIKQLQARVLAVERYLKDQQLLGIW | 586 |
| 27 | LTVQARQLLSGIVQQQNNLLRAIEAQQHLLQLTVWGIKQLQARVLAVERYLKDQQLLGIW | 586 |
| 28 | LTVQARQLLSGIVQQQNNLLRAIEAQQHLLQLTVWGIKQLQARVLAVERYLKDQQLLGIW | 586 |
| 29 | LTVQARQLLSGIVQQQNNLLRAIEAQQHLLQLTVWGIKQLQARVLAVERYLKDQQLLGIW | 560 |
| 30 | LTVQARQLLSGIVQQQNNLLRAIEAQQHLLQLTVWGIKQLQARVLAVERYLKDQQLLGIW | 586 |
| 31 | LTVQARQLLSGIVQQQNNLLRAIEAQQHLLQLTVWGIKQLQARVLAVERYLKDQQLLGIW | 586 |
| 32 | LTVQARQLLSGIVQQQNNLLRAIEAQQHLLQLTVWGIKQLQARVLAVERYLKDQQLLGIW | 586 |
| 33 | LTVQARQLLSGIVQQQNNLLRAIEAQQHLLQLTVWGIKQLQARVLAVERYLKDQQLLGIW | 586 |
| 34 | LTVQARQLLSGIVQQQNNLLRAIEAQQHLLQLTVWGIKQLQARVLAVERYLKDQQLLGIW | 563 |
| 35 | LTVQARQLLSGIVQQQNNLLRAIEAQQHLLQLTVWGIKQLQARVLAVERYLKDQQLLGIW | 586 |

```
      GCSGKLICTTTVPWNTSWSNKSLEQIWDNMTWMEWEREIDNYT----GYIYQLIEESQNQ
1     GCSGKLICTTTVPWNTSWSNKSLEQIWDNMTWMEWEREIDNYT----GYIYQLIEESQNQ  642
2     GCSGKLICTTTVPWNTSWSNKSLEQIWDNMTWMEWEREIDNYT----GYIYQLIEESQNQ  642
3     GCSGKLICTTTVPWNTSWSNKSLEQIWDNMTWMEWEREIDNYT----GYIYQLIEESQNQ  642
4     GCSGKLICTTTVPWNTSWSNKSLEQIWDNMTWMEWEREIDNYTGYIYQLIYQLIEESQNQ  646
5     GCSGKLICTTTVPWNTSWSNKSLEQIWDNMTWMEWEREIDNYT----GYIYQLIEESQNQ  642
6     GCSGKLICTTTVPWNTSWSNKSLEQIWDNMTWMEWEREIDNYT----GYIYQLIEESQNQ  642
7     GCSGKLICTTTVPWNTSWSNKSLEQIWDNMTWMEWEREIDNYT----GYIYQLIEESQNQ  642
8     GCSGKLICTTTVPWNTSWSNKSLEQIWDNMTWMEWEREIDNYT----GYIYQLIEESQNQ  642
9     GCSGKLICPTTVPWNTSWSNKSLEQIWDNMTWMEWEREIDNYT----GYIYQLIEESQNQ  642
10    GCSGKLICTTTVPWNTSWSNKSLEQIWDNMTWMEWEREIDNYT----GYIYQLIEESQNQ  642
11    GCSGKLICTTTVPWNTSWSNKSLEQIWDNMTWMEWEREIDNYT----GYIYQLIEESQNQ  642
12    GCSGKLICTTTVPWNTSWSNKSLEQIWDNMTWMEWEREIDNYT----GYIYQLIEESQNQ  642
13    GCSGKLICTTTVPWNTSWSNKSLEQIWDNMTWMEWEREIDNYT----GYIYQLIEESQNQ  642
14    GCSGKLICTTTVPWNTSWSNKSLEQIWDNMTWMEWEREIDNYT----GYIYQLIEESQNQ  642
15    GCSGKLICTTTVPWNTSWSNKSLEQIWDNMTWMEWEREIDNYT----GYIYQLIEESQNQ  641
16    GCSGKLICTTTVPWNTSWSNKSLEQIWDNMTWMEWEREIDNYT----GYIYQLIEESQNQ  642
17    GCSGKLICTTTVPWNTSWSNKSLEQIWDNMTWMEWEREIDNYT----GYIYQLIEESQNQ  642
18    GCSGKLICTTTVPWNTSWSNKSLEQIWDNMTWMEWEREIDNYT----GYIYQLIEESQNQ  642
19    GCSGKLICTTTVPWNTSWSNKSLEQIWDNMTWMEWEREIDNYT----GYIYQLIEESQNQ  626
20    GCSGKLICTTTVPWNTSWSNKSLEQIWDNMTWMEWEREIDNYT----GYIYQLIEESQNQ  642
21    GCSGKLICTTTVPWNTSWSNKSLEQIWDNMTWMEWEREIDNYT----GYIYQLIEESQNQ  642
22    GCSGKLICTTTVPWNTSWSNKSLEQIWDNMTWMEWEREIDNYT----GYIYQLIEESQNQ  642
23    GCSGKLICTTTVPWNTSWSNKSLEQIWDNMTWMEWEREIDNYT----GYIYQLIEESQNQ  642
24    GCSGKLICTTTVPWNTSWSNKSLEQIWDNMTWMEWEREIDNYT----GYIYQLIEESQNQ  642
25    GCSGKLICTTTVPWNTSWSNKSLEQIWDNMTWMEWEREIDNYT----GYIYQLIEESQNQ  642
26    GCSGKLICPTTVPWNTSWSNKSLEQIWDNMTWMEWEREIDNYT----GYIYQLIEESQNQ  642
27    GCSGKLICTTTVPWNTSWSNKSLEQIWDNMTWMEWEREIDNYT----GYIYQLIEESQNQ  642
28    GCSGKLICTTTVPWNTSWSNKSLEQIWDNMTWMEWEREIDNYT----GYIYQLIEESQNQ  642
29    GCSGKLICTTTVPWNTSWSNKSLEQIWDNMTWMEWEREIDNYT----GYIYQLIEESQNQ  616
30    GCSGKLICTTTVPWNTSWSNKSLEQIWDNMTWMEWEREIDNYT----GYIYQLIEESQNQ  642
31    GCSGKLICTTTVPWNTSWSNKSLEQIWDNMTWMEWEREIDNYT----GYIYQLIEESQNQ  642
32    GCSGKLICTTTVPWNTSWSNKSLEQIWDNMTWMEWEREIDNYT----GYIYQLIEESQNQ  642
33    GCSGKLICTTTVPWNTSWSNKSLEQIWDNMTWMEWEREIDNYT----GYIYQLIEESQNQ  642
34    GCSGKLICPTTVPWNTSWSNKSLEQIWDNMTWMEWEREIDNYT----GYIYQLIEESQNQ  619
35    GCSGKLICTTTVPWNTSWSNKSLEQIWDNMTWMEWEREIDNYT----GYIYQLIEESQNQ  642
```

| | Sequence | |
|---|---|---|
| | **QEKNEQELLALDKWASLWNWFDITNWLWYIKIFIMIVGGLIGLRIVFTVLSIVNRVRQGY** | |
| 1 | QEKNEQELLALDKWASLWNWFDITNWLWYIKIFIMIVGGLIGLRIVFTVLSIVNRVRQGY | 702 |
| 2 | QEKNEQELLALDKWASLWNWFDITNWLWYIKIFIMIVGGLIGLRIVFTVLSIVNRVRQGY | 702 |
| 3 | QEKNEQELLALDKWASLWNWFDITNWLWYIKIFIMIVGGLIGLRIVFTVLSIVNRVRQGY | 702 |
| 4 | QEKNEQELLALDKWASLWNWFDITNWLWYIKIFIMIVGGLIGLRIVFTVLSIVNRVRQGY | 706 |
| 5 | QEKNEQELLALDKWASLWNWFDITNWLWYIKIFIMIVGGLIGLRIVFTVLSIVNRVRQGY | 702 |
| 6 | QEKNEQELLALDKWASLWNWFDITNWLWYIKIFIMIVGGLIGLRIVFTVLSIVNRVRQGY | 702 |
| 7 | QEKNEQELLALDKWASLWNWFDITNWLWYIKIFIMIVGGLIGLRIVFTVLSIVNRVRQGY | 702 |
| 8 | QEKNEQELLALDKWASLWNWFDITNWLWYIKIFIMIVGGLIGLRIVFTVLSIVNRVRQGY | 702 |
| 9 | QEKNEQELLALDKWASLWNWFDITNWLWYIKIFIMIVGGLIGLRIVFTVLSIVNRVRQGY | 702 |
| 10 | QEKNEQELLALDKWASLWNWFDITNWLWYIKIFIMIVGGLIGLRIVFTVLSIVNRVRQGY | 702 |
| 11 | QEKNEQELLALDKWASLWNWFDITNWLWYIKIFIMIVGGLIGLRIVFTVLSIVNRVRQGY | 702 |
| 12 | QEKNEQELLALDKWASLWNWFDITNWLWYIKIFIMIVGGLIGLRIVFTVLSIVNRVRQGY | 702 |
| 13 | QEKNEQELLALDKWASLWNWFDITNWLWYIKIFIMIVGGLIGLRIVFTVLSIVNRVRQGY | 702 |
| 14 | QEKNEQELLALDKWASLWNWFDITNWLWYIKIFIMIVGGLIGLRIVFTVLSIVNRVRQGY | 702 |
| 15 | QEKNEQELLALDKWASLWNWFDITNWLWYIKIFIMIVGGVIGLRIVFTVLSIVNRVRQGY | 701 |
| 16 | QEKNEQELLALDKWASLWNWFDITNWLWYIKIFIMIVGGLIGLRIVFTVLSIVNRVRQGY | 702 |
| 17 | QEKNEQELLALDKWASLWNWFDITNWLWYIKIFIMIVGGLIGLRIVFTVLSIVNRVRQGY | 702 |
| 18 | QEKNEQELLALDKWASLWNWFDITNWLWYIKIFIMIVGGLIGLRIVFTVLSIVNRVRQGY | 702 |
| 19 | QEKNEQELLALDKWASLWNWFDITNWLWYIKIFIMIVGGLIGLRIVFTVLSIVNRVRQGY | 686 |
| 20 | QEKNEQELLALDKWASLWNWFDITNWLWYIKIFIMIVGGLIGLRIVFTVLSIVNRVRQGY | 702 |
| 21 | QEKNEQELLALDKWASLWNWFDITNWLWYIKIFIMIVGGLIGLRIVFTVLSIVNRVRQGY | 702 |
| 22 | QEKNEQELLALDKWASLWNWFDITNWLWYIKIFIMIVGGLIGLRIVFTVLSIVNRVRQGY | 702 |
| 23 | QEKNEQELLALDKWASLWNWFDITNWLWYIKIFIMIVGGLIGLRIVFTVLSIVNRVRQGY | 702 |
| 24 | QEKNEQELLALDKWASLWNWFDITNWLWYIKIFIMIVGGLIGLRIVFTVLSIVNRVRQGY | 702 |
| 25 | QEKNEQELLALDKWASLWNWFDITNWLWYIKIFIMIVGGLIGLRIVFTVLSIVNRVRQGY | 702 |
| 26 | QEKNEQELLALDKWASLWNWFDITNWLWYIKIFIMIVGGLIGLRIVFTVLSIVNRVRQGY | 702 |
| 27 | QEKNEQELLALDKWASLWNWFDITNWLWYIKIFIMIVGGLIGLRIVFTVLSIVNRVRQGY | 702 |
| 28 | QEKNEQELLALDKWAGLWNWFDITNWLWYIKIFIMIVGGLIGLRIVFTVLSIVNRVRQGY | 702 |
| 29 | QEKNEQELLALDKWASLWNWFDITNWLWYIKIFIMIVGGLIGLRIVFTVLSIVNRVRQGY | 676 |
| 30 | QEKNEQELLALDKWASLWNWFDITNWLWYIKIFIMIVGGLIGLRIVFTVLSIVNRVRQGY | 702 |
| 31 | QEKNEQELLALDKWASLWNWFDITNWLWYIKIFIMIVGGLIGLRIVFTVLSIVNRVRQGY | 702 |
| 32 | QEKNEQELLALDKWASLWNWFDITNWLWYIKIFIMIVGGLIGLRIVFTVLSIVNRVRQGY | 702 |
| 33 | QEKNEQELLALDKWASLWNWFDITNWLWYIKIFIMIVGGLIGLRIVFTVLSIVNRVRQGY | 702 |
| 34 | QEKNEQELLALDKWASLWNWFDITNWLWYIKIFIMIVGGLIGLRIVFTVLSIVNRVRQGY | 679 |
| 35 | QEKNEQELLALDKWASLWNWFDITNWLWYIKIFIMIVGGLIGLRIVFTVLSIVNRVRQGY | 702 |

**SPLSFQTHLPAQRGPDRPEGIGEEGGERDRDRSDPLVNGFLTLIWSDLRSLCLFSYHRLR**

| | | |
|---|---|---|
| 1 | SPLSFQTHLPAQRGPDRPEGIGEEGGERDRDRSDPLVNGFLALIWSDLRSLCLFSYHRLR | 762 |
| 2 | SPLSFQTHLPAQRGPDRPEGIGEEGGERDRDRSDPLVNGFLTLIWSDLRSLCLFSYHRLR | 762 |
| 3 | SPLSFQTHLPAQRGPDRPEGIGEEGGERDRDRSDPLVNGFLALIWSDLRSLCLFSYHRLR | 762 |
| 4 | SPLSFQTHLPAQRGPDRPEGIGEEGGERDRDRSDPLVNGFLTLIWSDLRSLCLFSYHRLR | 766 |
| 5 | SPLSFQTHLPAQRGPDRPEGIGEEGGERDRDRSDPLVNGFLTLIWSDLRSLCLFSYHRLR | 762 |
| 6 | SPLSFQTHLPAQRGPDRPEGIGEEGGERDRDRSDPLVNGFLTLIWSDLRSLCLFSYHRLR | 762 |
| 7 | SPLSFQTHLPAQRGPDRPEGIGEEGGERDRDRSDPLVNGFLTLIWSDLRSLCLFSYHRLR | 762 |
| 8 | SPLSFQTHLPAQRGPDRPEGIGEEGGERDRDRSDPLVNGFLTLIWSDLRSLCLFSYHRLR | 762 |
| 9 | SPLSFQTHLPAQRGPDRPEGIGEEGGERDRDRSDPLVNGFLTLIWSDLRSLCLFSYHRLR | 762 |
| 10 | SPLSFQTHLPAQRGPDRPEGIGEEGGERDRDRSDPLVNGFLTLIWSDLRSLCLFSYHRLR | 762 |
| 11 | SPLSFQTHLPAQRGPDRPEGIGEEGGERDRDRSDPLVNGFLTLIWSDLRSLCLFSYHRLR | 762 |
| 12 | SPLSFQTHLPAQRGPDRPEGIGEEGGERDRDRSDPLVNGFLTLIWSDLRSLCLFSYHRLR | 762 |
| 13 | SPLSFQTHLPAQRGPDRPEGIGEEGGERDRDRSDPLVNGFLALIWSDLRSLCLFSYHRLR | 762 |
| 14 | SPLSFQTHLPAQRGPDRPEGIGEEGGERDRDRSDPLVNGFLTLIWSDLRSLCLFSYHRLR | 762 |
| 15 | SPLSFQTHLPAQRGPDRPEGIGEEGGERDRDRSDPLVNGFLTLIWSDLRSLCLFSYHRLR | 761 |
| 16 | SPLSFQTHLPAQRGPDRPEGIGEEGGERDRDRSDPLVNGFLTLIWSDLRSLCLFSYHRLR | 762 |
| 17 | SPLSFQTHLPAQRGPDRPEGIGEEGGERDRDRSDPLVNGFLTLIWSDLRSLCLFSYHRLR | 762 |
| 18 | SPLSFQTHLPAQRGPDRPEGIGEEGGERDRDRSDPLVNGFLTLIWSDLRSLCLFSYHRLR | 762 |
| 19 | SPLSFQTHLPAQRGPDRPEGIGEEGGERDRDRSDPLVNGFLTLIWSDLRSLCLFSYHRLR | 746 |
| 20 | SPLSFQTHLPAQRGPDRPEGIGEEGGERDRDRSDPLVNGFLTLIWSDLRSLCLFSYHRLR | 762 |
| 21 | SPLSFQTHLPAQRGPDRPEGIGEEGGERDRDRSDPLVNGFLTLIWSDLRSLCLFSYHRLR | 762 |
| 22 | SPLSFQTHLPAQRGPDRPEGIGEEGGERDRDRSDPLVNGFLTLIWSDLRSLCLFSYHRLR | 762 |
| 23 | SPLSFQTHLPAQRGPDRPEGIGEEGGERDRDRSDPLVNGFLTLIWSDLRSLCLFSYHRLR | 762 |
| 24 | SPLSFQTHLPAQRGPDRPEGIGEEGGERDRDRSDPLVNGFLTLIWSDLRSLCLFSYHRLR | 762 |
| 25 | SPLSFQTHLPAQRGPDRPEGIGEEGGERDRDRSDPLVNGFLTLIWSDLRSLCLFSYHRLR | 762 |
| 26 | SPLSFQTHLPAQRGPDRPEGIGEEGGERDRDRSDPLVNGFLTLIWSDLRSLCLFSYHRLR | 762 |
| 27 | SPLSFQTHLPAQKGPDRPEGIGEEGGERDRDRSDPLVNGFLTLIWSDLRSLCLFSYHRLR | 762 |
| 28 | SPLSFQTHLPAQRGPDRPEGIGEEGGERDRDRSDPLVNGFLTLIWSDLRSLCLFSYHRLR | 762 |
| 29 | SPLSFQTHLPAQRGPDRPEGIGEEGGERDRDRSDPLVNGFLTLIWSDLRSLCLFSYHRLR | 736 |
| 30 | SPLSFQTHLPAQRGPDRPEGIGEEGGERDRDRSDPLVNGFLTLIWSDLRSLCLFSYHRLR | 762 |
| 31 | SPLSFQTHLPAQRGPDRPEGIGEEGGERDRDRSDPLVNGFLTLIWSDLRSLCLFSYHRLR | 762 |
| 32 | SPLSFQTHLPAQRGPDRPEGIGEEGGERDRDRSDPLVNGFLTLIWSDLRSLCLFSYHRLR | 762 |
| 33 | SPLSFQTHLPAQRGPDRPEGIGEEGGERDRDRSDPLVNGFLTLIWSDLRSLCLFSYHRLR | 762 |
| 34 | SPLSFQTHLPAQRGPDRPEGIGEEGGERDRDRSDPLVNGFLTLIWSDLRSLCLFSYHRLR | 739 |
| 35 | SPLSFQTHLPAQRGPDRPEGIGEEGGERDRDRSDPLVNGFLTLIWSDLRSLCLFSYHRLR | 762 |

| | | |
|---|---|---|
|  | DLLLIVTRIVELLGR-------RGWEVLKYWWNLLQYWSQELKNSAVSLLNATATAVAEG |  |
| 1 | DLLLIVTRIVELLGR-------RGWEVLKYWWNLLQYWSQELKNSAVSLLNATAIAVAEG | 815 |
| 2 | DLLLIVTRIVELLGR-------RGWEVLKYWWNLLQYWSQELKNSAVSLLNATATAVAEG | 815 |
| 3 | DLLLIVTRIVELLGR-------RGWEVLKYWWNLLQYWSQELKNSAVSLLNATATAVAEG | 815 |
| 4 | DLLLIVTRIVELMGR-------RGWEVLKYWWNLLQYWSQELKNSAVSLLNATATAVAEG | 819 |
| 5 | DLLLIVTRIVELLGR-------RGWEVLKYWWNLLQYWSQELKNSAVSLLNATAIAVAEG | 815 |
| 6 | DLLLIVTRIVELLGR-------RGWEVLKYWWNLLQYWSQELKNSAVSLLNATAIAVAEG | 815 |
| 7 | DLLLIVTRIVELLGR-------RGWEVLKYWWNLLQYWSQELKNSAVSLLNATATAVAEG | 815 |
| 8 | DLLLIVTRIVELLGR-------RGWEVLKYWWNLLRYWSQELKNSAVSLLNATATAVAEG | 815 |
| 9 | DLLLIVTRIVELLGR-------RGWEVLKYWWNLLQYWSQELKNSAVSLLNATAIAVAEG | 815 |
| 10 | DLLLIVTRIVELLGR-------RGWEVLKYWWNLLQYWSQELKNSAVSLLNATATAVAEG | 815 |
| 11 | DLLLIVTRIVELLGR-------RGWEVLKYWWNLLQYWSQELKNSAVSLLNATATAVAEG | 815 |
| 12 | DLLLIVTRIVELLGR-------RGWEVLKYWWNLLQYWSQELKNSAVSLLNATATAVAEG | 815 |
| 13 | DLLLIVTRIVELLGR-------RGWEVLKYWWNLLQYWSQELKNSAVSLLNATAIAVAEG | 815 |
| 14 | DLLLIVTRIVELLGR-------RGWEILKYWWNLLQYWSQELKNSAVSLLNATATAVAEG | 815 |
| 15 | DLLLIVTRIVELLGR-------RGWEVLKYWWNLLQYWSQELKNSAVSLLNATATAVAEG | 814 |
| 16 | DLLLIVTRIVELLGR-------RGWEVLKYWWNLLQYWSQELKNSAVSLLNATATAVAEG | 815 |
| 17 | DLLLIVTRIVELLGR-------RGWEVLKYWWNLLQYWSQELKNSAVSLLNATATAVAEG | 815 |
| 18 | DLLLIVTRIVELLGR-------RGWEVLKYWWNLLQYWSQELKNSAVSLLNATAIAVAEG | 815 |
| 19 | DLLLIVTRIVELLGR-------RGWEVLKYWWNLLQYWSQELKNSAVSLLNATAIAVAEG | 799 |
| 20 | DLLLIVTRIVELLGR-------RGWEVLKYWWNLLQYWSQELKNSAVSLLNATATAVAEG | 815 |
| 21 | DLLLIVTRIVELLGR-------RGWEVLKYWWNLLQYWSQELKNSAVSLLNATATAVAEG | 815 |
| 22 | DLLLIVTRIVELLGR-------RGWEVLKYWWNLLQYWSQELKNSAVSLLNATATAVAEG | 815 |
| 23 | DLLLIVTRIVELLGR-------RGWEVLKYWWNLLQYWSQELKNSAVSLLNATATAVAEG | 815 |
| 24 | DLLLIVTRIVELLGR-------RGWEVLKYWWNLLQYWSQELKNSAVSLLNATATAVAEG | 815 |
| 25 | DLLLIVTRIVELLGR-------RGWEVLKYWWNLLQYWSQELKNSAVSLLNATATAVAEG | 815 |
| 26 | DLLLIVTRIVELLGR-------RGWEVLKYWWNLLQYWSQELKNSAVSLLNATAIAVAEG | 815 |
| 27 | DLLLIVTRIVELLGR-------RGWEVLKYWWNLLQYWSQELKNSAVSLLNATATAVAEG | 815 |
| 28 | DLLLIVTRIVELLGR-------RGWEVLKYWWNLLQYWSQELKNSAVSLLNATATAVAEG | 815 |
| 29 | DLLLIVTRIVELLGR-------RGWEVLKYWWNLLQYWSQELKNSAVSLLNATATAVAEG | 789 |
| 30 | DLLLIVTRIVELLGR-------RGWEVLKYWWNLLQYWSQELKNSAVSLLNATATAVAEG | 815 |
| 31 | DLLLIVTRIVELLGR-------RGWEVLKYWWNLLQYWSQELKNSAVSLLNATATAVAEG | 815 |
| 32 | DLLLIVTRIVELLGR-------RGWEVLKYWWNLLQYWSQELKNSAVSLLNATATAVAEG | 815 |
| 33 | DLLLIVTRIVELLGR-------RGWEVLKYWWNLLQYWSQELKNSAVSLLNATATAVAEG | 815 |
| 34 | DLLLIVTRIVELLGR-------RGWEVLKYWWNLLQYWSQELKNSAVSLLNATAIAVAEG | 792 |
| 35 | DLLLIVTRIVELLGR-------RGWEVLKYWWNLLQYWSQELKNSAVSLLNATATAVAEG | 815 |

| | Sequence | |
|---|---|---|
| | **TDRVIEVVQRACRAILHIPRRIRQGLERALL** | |
| 1 | TDRVIEVVQRACRAILHIPRRIRQGLERALL | 846 |
| 2 | TDRVIEVVQRACRAILHIPRRIRQGLERALL | 846 |
| 3 | TDRVIEVVQRACRAILHIPRRIRQGLERALL | 846 |
| 4 | TDRVIEVVQRACRAILHIPRRIRQGLERALL | 850 |
| 5 | TDRVIEVVQRACRAIRHIPRRIRQGLERALL | 846 |
| 6 | TDRVIEVVQGACRAILHIPRRIRQGLERALL | 846 |
| 7 | TDRVIEVVQRACRAILHIPRRIRQGLERALL | 846 |
| 8 | TDRVIEVVQRACRAILHIPRRIRQGLERALL | 846 |
| 9 | TDRVIEVVQRACRAILHIPRRIRQGLERALL | 846 |
| 10 | TDRVIEVVQRACRAILHIPRRIRQGLERALL | 846 |
| 11 | TDRVVEVVQRACRAILHIPRRIRQGLERALL | 846 |
| 12 | TDRVIEVVQRACRAILHIPRRIRQGLERALL | 846 |
| 13 | TDRVIEVVQRTCRAIRHIPRRIRQGLERALL | 846 |
| 14 | TDRVIEVVQRACRAILHIPRRIRQGLERALL | 846 |
| 15 | TDRVIEVVQRACRAILHIPRRIRQGLERALL | 845 |
| 16 | TDRVIEVVQRACRAILHIPRRIRQGLERALL | 846 |
| 17 | TDRVIEVVQRACRAILHIPRRIRQGLERALL | 846 |
| 18 | TDRVIEVVQRACRAIRHIPRRIRQGLERALL | 846 |
| 19 | TDRVIEVVQRACRAIRHIPRRIRQGLERALL | 830 |
| 20 | TDRVIEVVQRACRAILHIPRRIRQGLERALL | 846 |
| 21 | TDRVIEVVQRACRAILHIPRRIRQGLERALL | 846 |
| 22 | TDRVIEVVQRACRAILHIPRRIRQGLERALL | 846 |
| 23 | TDRVIEVVQRACRAILHIPRRIRQGLERALL | 846 |
| 24 | TDRVIEVVQRACRAILHIPRRIRQGLERALL | 846 |
| 25 | TDRVIEVVQRACRAILHIPRRIRQGLERALL | 846 |
| 26 | TDRVIEVVQRACRAILHIPRRIRQGLERALL | 846 |
| 27 | TDRVIEVVQRACRAILHIPRRIRQGLERALL | 846 |
| 28 | TDRVIEVVQRACRAILHIPRRIRQGLERALL | 846 |
| 29 | TDRVIEVVQRACRAILHIPRRIRQGLERALL | 820 |
| 30 | TDRVIEVVQRACRAILHIPRRIRQGLERALL | 846 |
| 31 | TDRVIEVVQRACRAILHIPRRIRQGLERALL | 846 |
| 32 | TDRVIEVVQRACRAILHIPRRIRQGLERALL | 846 |
| 33 | TDRVIEVVQRACRAILHIPRRIRQGLERALL | 846 |
| 34 | TDRVIEVVQRACRAILHIPRRIRQGLERALL | 823 |
| 35 | TDRVIEVVQRACRAILHIPRRIRQGLERALL | 846 |

**Consensus**

1. B.US.2006.700010040_C9_4520.EU289193
2. B.US.2008.CH0040_3_d0666_ipe032_27_9.MG900342
3. B.US.2008.CH0040_3_d0666_ipe032_2_01.MG900343
4. B.US.2008.CH0040_3_d0666_ipe032_3_03.MG900344
5. B.US.2008.CH0040_3_d0666_ipe032_3_04.MG900345
6. B.US.2008.CH0040_3_d0666_ipe032_3_05.MG900346
7. B.US.2008.CH0040_3_d0666_ipe032_3_07.MG900347
8. B.US.2008.CH0040_3_d0666_ipe032_3_08.MG900348
9. B.US.2008.CH0040_3_d0666_ipe032_5_11.MG900349
10. B.US.2008.CH0040_3_d0666_ipe032_5_12.MG900350
11. B.US.2008.CH0040_3_d0666_ipe032_5_13.MG900351
12. B.US.2008.CH0040_3_d0666_ipe032_5_15.MG900352
13. B.US.2008.CH0040_3_d0666_ipe032_5_16.MG900353
14. B.US.2008.CH0040_3_d0666_ipe032_5_17.MG900354
15. B.US.2008.CH0040_3_d0666_ipe032_5_18.MG900355
16. B.US.2008.CH0040_3_d0666_ipe032_5_21.MG900356
17. B.US.2008.CH0040_3_d0666_ipe032_5_22.MG900357
18. B.US.2008.CH0040_3_d0666_ipe032_5_23.MG900358
19. B.US.2008.CH0040_3_d0666_ipe032_5_24.MG900359
20. B.US.2008.CH0040_3_d0666_ipe032_5_33.MG900360
21. B.US.2008.CH0040_3_d0666_ipe032_5_34.MG900361
22. B.US.2008.CH0040_3_d0666_ipe032_5_37.MG900362
23. B.US.2008.CH0040_3_d0666_ipe032_5_38.MG900363
24. B.US.2008.CH0040_3_d0666_ipe032_5_41.MG900364
25. B.US.2008.CH0040_3_d0666_ipe032_5_42.MG900365
26. B.US.2008.CH0040_3_d0666_ipe032_5_43.MG900366
27. B.US.2008.CH0040_3_d0666_ipe032_5_44.MG900367
28. B.US.2008.CH0040_3_d0666_ipe032_5_45.MG900368
29. B.US.2008.CH0040_3_d0666_ipe032_5_48.MG900369
30. B.US.2008.CH0040_3_d0666_ipe032_5_49.MG900370
31. B.US.2008.CH0040_3_d0666_ipe032_5_51.MG900371
32. B.US.2008.CH0040_3_d0666_ipe032_5_52.MG900372
33. B.US.2008.CH0040_3_d0666_ipe032_5_56.MG900373
34. B.US.2008.CH0040_3_d0666_ipe032_5_57.MG900374
35. B.US.2008.CH0040_3_d0666_ipe032_5_58.MG900375
36. B.US.2008.CH0040_3_d0666_ipe032_5_60.MG900376
37. B.US.2008.CH0040_3_d0666_ipe032_5_62.MG900377
38. B.US.2008.CH0040_3_d0666_ipe032_5_63.MG900378
39. B.US.2008.CH0040_3_d0666_ipe032_5_64.MG900379
40. B.US.2008.CH0040_3_d0666_ipe032_5_65.MG900380
41. B.US.2008.CH0040_3_d0666_ipe032_5_66.MG900381
42. B.US.2008.CH0040_3_d0666_ipe032_5_67.MG900382
43. B.US.2008.CH0040_3_d0666_ipe032_5_68.MG900383
44. B.US.2008.CH0040_3_d0666_ipe032_5_73.MG900384
45. B.US.2008.CH0040_3_d0666_ipe032_5_79.MG900385
46. B.US.2008.CH0040_3_d0666_ipe032_5_82.MG900386

| | | |
|---|---|---|
| | **MRVMGIRKNYQHLWREGILLLGILMICSAADNLWVTVYYGVPVWREATTTLFCASDAKAY** | |
| 1 | MRVMGIRKNYQHLWREGILLLGILMICSAADNLWVTVYYGVPVWREATTTLFCASDAKAY | 60 |
| 2 | MRVMGIRKNYQHLWREGILLLGILMICSAADNLWVTVYYGVPVWREATTTLFCASDAKAY | 60 |
| 3 | MRVMGIRKNYQHLWREGILLLGILMICSAADNLWVTVYYGVPVWREATTTLFCASDAKAY | 60 |
| 4 | MRVMGIRKNYQHLWREGILLLGILMICSAADNLWVTVYYGVPVWREATTTLFCASDAKAY | 60 |
| 5 | MRVMGIRKNYQHLWREGILLLGILMICSAADNLWVTVYYGVPVWREATTTLFCASDAKAY | 60 |
| 6 | MRVMGIRKNYQHLWREGILLLGILMICSAADNLWVTVYYGVPVWRKATTTLFCASDAKAY | 60 |
| 7 | MRVMGIRKNYQHLWREGILLLGILMICSAADNLWVTVYYGVPVWREATTTLFCASDAKAY | 60 |
| 8 | MRVMGIRKNYQHLWREGILLLGILMICSAADNLWVTVYYGVPVWREATTTLFCASDAKAY | 60 |
| 9 | MRVMGIRKNYQHLWREGILLLGILMICSAADNLWVTVYYGVPVWREATTTLFCASDAKAY | 60 |
| 10 | MRVMGIRKNYQHLWREGILLLGILMICSAADNLWVTVYYGVPVWREATTTLFCASDAKAY | 60 |
| 11 | MRVMGIRKNYQHLWREGILLLGILMICSAADNLWVTVYYGVPVWREATTTLFCASDAKAY | 60 |
| 12 | MRVMGIRKNYQHLWREGILLLGILMICSAADNLWVTVYYGVPVWREATTTLFCASDAKAY | 60 |
| 13 | MRVMGIRKNYQHLWREGILLLGILMICSAADNLWVTVYYGVPVWREATTTLFCASDAKAY | 60 |
| 14 | MRVMGIRKNYQHLWREGILLLGILMICSAADNLWVTVYYGVPVWREATTTLFCASDAKAY | 60 |
| 15 | MRVMGIRKNYQHLWREGILLLGILMICSAADNLWVTVYYGVPVWREATTTLFCASDAKAY | 60 |
| 16 | MRVMGIRKNYQHLWREGILLLGILMICSAADNLWVTVYYGVPVWREATTTLFCASDAKAY | 60 |
| 17 | MRVMGIRKNYQHLWREGILLLGILMICSAADNLWVTVYYGVPVWREATTTLFCASDAKAY | 60 |
| 18 | MRVMGIRKNYQHLWREGILLLGILMICSAADNLWVTVYYGVPVWREATTTLFCASDAKAY | 60 |
| 19 | MRVMGIRKNYQHLWREGILLLGILMICSAADNLWVTVYYGVPVWREATTTLFCASDAKAY | 60 |
| 20 | MRVMGIRKNYQHLWREGILLLGILMICSAADNLWVTVYYGVPVWREATTTLFCASDAKAY | 60 |
| 21 | MRVMGIRKNYQHLWREGILLLGILMICSAADNLWVTVYYGVPVWREATTTLFCASDAKAY | 60 |
| 22 | MRVMGIRKNYQHLWREGILLLGILMICSAADNLWVTVYYGVPVWREATTTLFCASDAKAY | 60 |
| 23 | MRVMGIRKNYQHLWREGILLLGILMICSAADNLWVTVYYGVPVWREATTTLFCASDAKAY | 60 |
| 24 | MRVMGIRKNYQHLWREGILLLGILMICSAADNLWVTVYYGVPVWREATTTLFCASDAKAY | 60 |
| 25 | MRVMGIRKNYQHLWREGILLLGILMICSAADNLWVTVYYGVPVWREATTTLFCASDAKAY | 60 |
| 26 | MRVMGIRKNYQHLWREGILLLGILMICSAADNLWVTVYYGVPVWREATTTLFCASDAKAY | 60 |
| 27 | MRVMGIRKNYQHLWREGILLLGILMICSAADNLWVTVYYGVPVWREATTTLFCASDAKAY | 60 |
| 28 | MRVMGIRKNYQHLWREGILLLGILMICSAADNLWVTVYYGVPVWREATTTLFCASDAKAY | 60 |
| 29 | MRVMGIRKNYQHLWREGILLLGILMICSAADNLWVTVYYGVPVWREATTTLFCASDAKAY | 60 |
| 30 | MRVMGIRKNYQHLWREGILLLGILMICSAADNLWVTVYYGVPVWREATTTLFCASDAKAY | 60 |
| 31 | MRVMGIRKNYQHLWREGILLLGILMICSAADNLWVTVYYGVPVWREATTTLFCASDAKAY | 60 |
| 32 | MRVMGIRKNYQHLWREGILLLGILMICSAADNLWVTVYYGVPVWREATTTLFCASDAKAY | 60 |
| 33 | MRVMGIRKNYQHLWREGILLLGILMICSAADNLWVTVYYGVPVWREATTTLFCASDAKAY | 60 |
| 34 | MRVMGIRKNYQHLWREGILLLGILMICSAADNLWVTVYYGVPVWREATTTLFCASDAKAY | 60 |
| 35 | MRVMGIRKNYQHLWREGILLLGILMICSAADNLWVTVYYGVPVWREATTTLFCASDAKAY | 60 |
| 36 | MRVMGIRKNYQHLWREGILLLGILMICSAADNLWVTVYYGVPVWREATTTLFCASDAKAY | 60 |
| 37 | MRVMGIRKNYQHLWRKGILLLGILMICSAADNLWVTVYYGVPVWREATTTLFCASDAKAY | 60 |
| 38 | MRVMGIRKNYQHLWREGILLLGILMICSAADNLWVTVYYGVPVWREATTTLFCASDAKAY | 60 |
| 39 | MRVMGIRKNYQHLWREGILLLGILMICSAADNLWVTVYYGVPVWREATTTLFCASDAKAY | 60 |
| 40 | MRVMGIRKNYQHLWREGILLLGILMICSAADNLWVTVYYGVPVWREATTTLFCASDAKAY | 60 |
| 41 | MRVMGIRKNYQHLWREGILLLGILMICSAADNLWVTVYYGVPVWREATTTLFCASDAKAY | 60 |
| 42 | MRVMGIRKNYQHLWREGILLLGILMICSAADNLWVTVYYGVPVWREATTTLFCASDAKAY | 60 |
| 43 | MRVMGIRKNYQHLWREGILLLGILMICSAADNLWVTVYYGVPVWREATTTLFCASDAKAY | 60 |
| 44 | MRVMGIRKNYQHLWREGILLLGILMICSAADNLWVTVYYGVPVWREATTTLFCASDAKAY | 60 |
| 45 | MRVMGIRKNYQHLWREGILLLGILMICSAADNLWVTVYYGVPVWREATTTLFCASDAKAY | 60 |
| 46 | MRVMGIRKNYQHLWREGILLLGILMICSAADNLWVTVYYGVPVWREATTTLFCASDAKAY | 60 |

| | Sequence | |
|---|---|---|
| | **DTEAHNVWATHACVPTDPNPQEVELKNVTENFNMWENNMVEQMHEDIISLWDQSLKPCVK** | |
| 1 | DTEAHNVWATHACVPTDPNPQEVELKNVTENFNMWENNMVEQMHEDIISLWDQSLKPCVK | 120 |
| 2 | DTEAHNVWATHACVPTDPNPQEVELK----NFNMWENNMVEQMHEDIISLWDQSLKPCVK | 116 |
| 3 | DTEAHNVWATHACVPTDPNPQEVELKNVTENFNMWENNMVEQMHEDIISLWDQSLKPCVK | 120 |
| 4 | DTEAHNVWATHACVPTDPNPQEVELKNVTENFNMWENNMVEQMHEDIISLWDQSLKPCVK | 120 |
| 5 | DTEAHNVWATHACVPTDPNPQEVELKNVTENFNMWENNMVEQMHEDIISLWDQSLKPCVK | 120 |
| 6 | DTEAHNVWATHACVPTDPNPQEVELKNVTENFNMWENNMVEQMHEDIISLWDQSLKPCVK | 120 |
| 7 | DTEAHNVWATHACVPTDPNPQEVELKNVTENFNMWENNMVEQMHEDIISLWDQSLKPCVK | 120 |
| 8 | DTEAHNVWATHACVPTDPNPQEVELKNVTENFNMWENNMVEQMHEDIISLWDQSLKPCVK | 120 |
| 9 | DTEAHNVWATHACVPTDPNPQEVELKNVTENFNMWENNMVEQMHEDIISLWDQSLKPCVK | 120 |
| 10 | DTEAHNVWATHACVPTDPNPQEVELKNVTENFNMWENNMVEQMHEDIISLWDQSLKPCVK | 120 |
| 11 | DTEAHNVWATHACVPTDPNPQEVELKNVTENFNMWENNMVEQMHEDIISLWDQSLKPCVK | 120 |
| 12 | DTEAHNVWATHACVPTDPNPQEVELKNVTENFNMWENNMVEQMHEDIISLWDQSLKPCVK | 120 |
| 13 | DTEAHNVWATHACVPTDPNPQEVELKNVTENFNMWENNMVEQMHEDIISLWDQSLKPCVK | 120 |
| 14 | DTEAHNVWATHACVPTDPNPQEVELKNVTENFNMWENNMVEQMHEDIISLWDQSLKPCVK | 120 |
| 15 | DTEAHNVWATHACVPTDPNPQEVELKNVTENFNMWENNMVEQMHEDIISLWDQSLKPCVK | 120 |
| 16 | DTEAHNVWATHACVPTDPNPQEVELKNVTENFNMWENNMVEQMHEDIISLWDQSLKPCVK | 120 |
| 17 | DTEAHNVWATHACVPTDPNPQEVELKNVTENFNMWENNMVEQMHEDIISLWDQSLKPCVK | 120 |
| 18 | DTEAHNVWATHACVPTDPNPQEVELKNVTENFNMWENNMVEQMHEDIISLWDQSLKPCVK | 120 |
| 19 | DTEAHNVWATHACVPTDPNPQEVELKNVTENFNMWENNMVEQMHEDIISLWDQSLKPCVK | 120 |
| 20 | DTEAHNVWATHACVPTDPNPQEVELKNVTENFNMWENNMVEQMHEDIISLWDQSLKPCVK | 120 |
| 21 | DTEAHNVWATHACVPTDPNPQEVELKNVTENFNMWENNMVEQMHEDIISLWDQSLKPCVK | 120 |
| 22 | DTEAHNVWATHACVPTDPNPQEVELKNVTENFNMWENNMVEQMHEDIISLWDQSLKPCVK | 120 |
| 23 | DTEAHNVWATHACVPTDPNPQEVELKNVTENFNMWENNMVEQMHEDIISLWDQSLKPCVK | 120 |
| 24 | DTEAHNVWATHACVPTDPNPQEVELKNVTENFNMWENNMVEQMHEDIISLWDQSLKPCVK | 120 |
| 25 | DTEAHNVWATHACVPTDPNPQEVELKNVTENFNMWENNMVEQMHEDIISLWDQSLKPCVK | 120 |
| 26 | DTEAHNVWATHACVPTDPNPQEVELKNVTENFNMWENNMVEQMHEDIISLWDQSLKPCVK | 120 |
| 27 | DTEAHNVWATHACVPTDPNPQEVELKNVTENFNMWENNMVEQMHEDIISLWDQSLKPCVK | 120 |
| 28 | DTEAHNVWATHACVPTDPNPQEVELKNVTENFNMWENNMVEQMHEDIISLWDQSLKPCVK | 120 |
| 29 | DTEAHNVWATHACVPTDPNPQEVELKNVTENFNMWENNMVEQMHEDIISLWDQSLKPCVK | 120 |
| 30 | DTEAHNVWATHACVPTDPNPQEVELKNVTENFNMWENNMVEQMHEDIISLWDQSLKPCVK | 120 |
| 31 | DTEAHNVWATHACVPTDPNPQEVELKNVTENFNMWENNMVEQMHEDIISLWDQSLKPCVK | 120 |
| 32 | DTEAHNVWATHACVPTDPNPQEVELKNVTENFNMWENNMVEQMHEDIISLWDQSLKPCVK | 120 |
| 33 | DTEAHNVWATHACVPTDPNPQEVELKNVTENFNMWENNMVEQMHEDIISLWDQSLKPCVK | 120 |
| 34 | DTEAHNVWATHACVPTDPNPQEVELKNVTENFNMWENNMVEQMHEDIISLWDQSLKPCVK | 120 |
| 35 | DTEAHNVWATHACVPTDPNPQEVELKNVTENFNMWENNMVEQMHEDIISLWDQSLKPCVK | 120 |
| 36 | DTEAHNVWATHACVPTDPNPQEVELKNVTENFNMWENNMVEQMHEDIISLWDQSLKPCVK | 120 |
| 37 | DTEAHNVWATHACVPTDPNPQEVELKNVTENFNMWENNMVEQMHEDIISLWDQSLKPCVK | 120 |
| 38 | DTEAHNVWATHACVPTDPNPQEVELKNVTENFNMWENNMVEQMHEDIISLWDQSLKPCVK | 120 |
| 39 | DTEAHNVWATHACVPTDPNPQEVELKNVTENFNMWENNMVEQMHEDIISLWDQSLKPCVK | 120 |
| 40 | DTEAHNVWATHACVPTDPNPQEVELKNVTENFNMWENNMVEQMHEDIISLWDQSLKPCVK | 120 |
| 41 | DTEAHNVWATHACVPTDPNPQEVELKNVTENFNMWENNMVEQMHEDIISLWDQSLKPCVK | 120 |
| 42 | DTEAHNVWATHACVPTDPNPQEVELKNVTENFNMWENNMVEQMHEDIISLWDQSLKPCVK | 120 |
| 43 | DTEAHNVWATHACVPTDPNPQEVELKNVTENFNMWENNMVEQMHEDIISLWDQSLKPCVK | 120 |
| 44 | DTEAHNVWATHACVPTDPNPQEVELKNVTENFNMWKNNMVEQMHEDIISLWDQSLKPCVK | 120 |
| 45 | DTEAHNVWATHACVPTDPNPQEVELKNVTENFNMWENNMVEQMHEDIISLWDQSLKPCVK | 120 |
| 46 | DTEAHNVWATHACVPTDPNPQEVELKNVTENFNMWENNMVEQMHEDIISLWDQSLKPCVK | 120 |

| | Sequence | |
|---|---|---|
| | LTPLCVTLNCTDLEKXTNTT---NSTEKMMEKGEVKNCSFKITTDXKDRTRKEYALFYKL | |
| 1 | LTPLCVTLNCTDLGNVTNTT---NSNGEMMEKGEVKNCSFKITTDIKDRTRKEYALFYKL | 177 |
| 2 | LTPLCVTLNCTDLGNVTTTI---NSNGTLMEKGEVKNCSFKITTDTKDRTRKEYALFYKL | 173 |
| 3 | LTPLCVTLNCTDLGNVTTTT---NSNGTLMEKGEVKNCSFKITTDTKDRTRKEYALFYKL | 177 |
| 4 | LTPLCVTLNCTDLEKGTNTT---NSTEKMMEKGEVKNCSFRITTDIKDRTRKEYALFYKL | 177 |
| 5 | LTPLCVTLNCTDLENVTDTT---NSTEKMMEKGEVKNCSFKITTDLKDRTRKEYALFYKL | 177 |
| 6 | LTPLCVTLNCTDLEKGTNTT---NSTEKMMEKGEVKNCSFRITTDIKDRTRKEYALFYKL | 177 |
| 7 | LTPLCVTLNCTDLENVTNTT---NSTEKMMEKGEVKNCSFKITTDITDRTRKEYALFYKL | 177 |
| 8 | LTPLCVTLNCTDLEKGTNTT---NSTEKMMEKGEVKNCSFRITTDIKDRTRKEYALFYKL | 177 |
| 9 | LTPLCVTLNCTDLGNVTTTT---NSNGTLMEKGEVKNCSFKITTDTKDRTRKEYALFYKL | 177 |
| 10 | LTPLCVTLNCTDLEKGTNTT---NSTEKMMEKGEVKNCSFKITTDVKDRTRKEYALFYKL | 177 |
| 11 | LTPLCVTLNCTDLEKGTNTT---NSTEKMMEKGEVKNCSFKITTDMKDRTRKEYALFYKL | 177 |
| 12 | LTPLCVTLNCTDLEKGTNTT---NSTEKMMEKGEVKNCSFKITTDVKDRTRKEYALFYKL | 177 |
| 13 | LTPLCVTLNCTDLGKVTNTT---NSTEKMMEQGEVKNCSFKITTDVKDRTRKEYALFYKL | 177 |
| 14 | LTPLCVTLNCTDLEKGTNTT---NSTEKMMEKGEVKNCSFRITTDIKDRTRKEYALFYKL | 177 |
| 15 | LTPLCVTLNCTDLEKDTNTT---NSTEKMMEKGEVKNCSFKITTDMKDRTRKEYALFYKL | 177 |
| 16 | LTPLCVTLNCTDLEKGTNTT---NSTEKMMEKGEVKNCSFRITTDIKDRTRKEYALFYKL | 177 |
| 17 | LTPLCVTLNCTDLEKGTNTT---NSTEKMMEKGEVKNCSFRITTDIKDRTRKEYALFYKL | 177 |
| 18 | LTPLCVTLNCTDLGNVTTTT---NSNGTLMEKGEVKNCSFKITTDTKDRTRKEYALFYKL | 177 |
| 19 | LTPLCVTLNCTDLGNVTTTT---NSNGTLMEKGEVKNCSFKITTDTKDRTRKEYALFYKL | 177 |
| 20 | LTPLCVTLNCTDLGNVTTTT---NSNGTLMEKGEVKNCSFKITTDTKDRTRKEYALFYKL | 177 |
| 21 | LTPLCVTLNCTDLEKGTNTT---NSTEKMMEKGEVKNCSFRITTDIKDRTRKEYALFYKL | 177 |
| 22 | LTPLCVTLNCTDLGNVTTTT---NSNGTLMEKGEVKNCSFKITTEIKDRTRKEYALFYKL | 177 |
| 23 | LTPLCVTLKCTDLEKGTNTT---NSTEKMMEKGEVKNCSFKITTDVKDRTRKEYALFYKL | 177 |
| 24 | LTPLCVTLNCTDLEKGTNTT---NSTEKMMEKGEVKNCSFKITTDVKDRTRKEYALFYKL | 177 |
| 25 | LTPLCVTLNCTDLEKGTNTT---NSTEKMMEKGEVKNCSFRITTDIKDRTRKEYALFYKL | 177 |
| 26 | LTPLCVTLNCTDLEKGTNTT---NSTEKMMEKGEVKNCSFRITTDIKDRTRKEYALFYKL | 177 |
| 27 | LTPLCVTLNCTDLGNVTTTT---NSNGTLMEKGEVKNCSFKITTDTKDRTRKEYALFYKL | 177 |
| 28 | LTPLCVTLNCTDLEKDTNTT---NSTEKMMEKGEVKNCSFRITTDIKDRTRKEYALFYKL | 177 |
| 29 | LTPLCVTLNCTDLGNVTTTT---NSNGTLMEKGEVKNCSFKITTDTKDRTRKEYALFYKL | 177 |
| 30 | LTPLCVTLNCTDLGNVTTTT---NSNGTLMEKGEVKNCSFKITTDTKDRTRKEYALFYKL | 177 |
| 31 | LTPLCVTLNCTDLGNVTTTT---NSNGTLMEKGEVKNCSFKITTDTKDRTRKEYALFYKL | 177 |
| 32 | LTPLCVTLNCTDLGNVTTTT---NSNGTLMEKGEVKNCSFKITTDTKDRTRKEYALFYKL | 177 |
| 33 | LTPLCVTLNCTDLGNVTTTT---NSNGTLMEKGEVKNCSFKITTDTKDRTRKEYALFYKL | 177 |
| 34 | LTPLCVTLNCTDLEKGTNTT---NSTEKMMEKGEVKNCSFRITTDIKDRTRKEYALFYKL | 177 |
| 35 | LTPLCVTLNCTDLEKGTNTT---NSTEKMMEKGEVKNCSFRITTDIKDRTRKEYALFYKL | 177 |
| 36 | LTPLCVTLNCTDLGNVTTTT---NSNGTLMEKGEVKNCSFKITTEIKDRTRKEYALFYKL | 177 |
| 37 | LTPLCVTLNCTDLGNVTTTT---NSNGTLMEKGEVKNCSFKITTDVKDRTRKEYALFYKL | 177 |
| 38 | LTPLCVTLNCTDLEKGTNTT---NSTEKMMEKGEVKNCSFRITTDIKDRTRKEYALFYKL | 177 |
| 39 | LTPLCVTLNCTDLEKGTNTT---NSTEKMMEKGEVKNCSFRITTDIKDRTRKEYALFYKL | 177 |
| 40 | LTPLCVTLNCTDLEKGTNTT---NSTEKMMEKGEVKNCSFRITTDIKDRTRKEYALFYKL | 177 |
| 41 | LTPLCVTLNCTDLGNVTTTI---NSNGTLMEKGEVKNCSFKITTDTKDRTRKEYALFYKL | 177 |
| 42 | LTPLCVTLNCTDLGNVTTTT---NSNGTLMEKGEVKNCSFKITTDTKDRTRKEYALFYKL | 177 |
| 43 | LTPLCVTLNCTDLEKGTNTT---NSTEKMMEKGEVKNCSFRITTDIKDRTRKEYALFYKL | 177 |
| 44 | LTPLCVTLNCTDLGNVTTTT---NSNGTLMEKGEVKNCSFKITTEIKDRTRKEYALFYKL | 177 |
| 45 | LTPLCVTLNCTDLEKGTNTT---NSTEKMMEKGEVKNCSFKITTDVKDRTRKEYALFYKL | 177 |
| 46 | LTPLCVTLNCTDLEKGTNTT---NSTEKMMEKGEVKNCSFRITTDIKDRTRKEYALFYKL | 177 |

| | Sequence | |
|---|---|---|
|  | **DVVPIND-------TRYRLVSCNTSVITQACPKVSFEPIPIHYCAPAGFAILKCNDKQFI** |  |
| 1 | DVVPIND-------TRYRLVSCNTSVITQACPKVSFEPIPIHYCAPAGFAILKCNDKQFI | 230 |
| 2 | DVVPIND-------TRYRLVSCNTSVITQACPKVSFEPIPIHYCAPAGFAILKCNDKQFI | 226 |
| 3 | DVVPIND-------TRYRLVSCNTSVITQACPKVSFEPIPIHYCAPAGFAILKCNDKQFI | 230 |
| 4 | DVVPIND-------TRYRLVSCNTSVITQACPKVSFEPIPIHYCAPAGFAILKCNDKQFI | 230 |
| 5 | DVVPIND-------TRYRLVSCNTSVITQACPKVSFEPIPIHYCAPAGFAILKCNDKQFI | 230 |
| 6 | DVVPIND-------TRYRLVSCNTSVITQACPKVSFEPIPIHYCAPAGFAILKCNDKQFI | 230 |
| 7 | DVVPIND-------TRYRLVSCNTSVITQACPKVSFEPIPIHYCAPAGFAILKCNDKQFI | 230 |
| 8 | DVVPIND-------TRYRLVSCNTSVITQACPKVSFEPIPIHYCAPAGFAILKCNDKQFI | 230 |
| 9 | DVVPIND-------TRYRLVSCNTSVITQACPKVSFEPIPIHYCAPAGFAILKCNDKQFI | 230 |
| 10 | DVVPIND-------TRYRLVSCNTSVITQACPKVSFEPIPIHYCAPAGFAILKCNDKQFI | 230 |
| 11 | DVVPIND-------TRYRLVSCNTSVITQACPKVSFEPIPIHYCAPAGFAILKCNDKQFI | 230 |
| 12 | DVVPIND-------TRYRLVSCNTSVITQACPKVSFEPIPIHYCAPAGFAILKCNDKQFI | 230 |
| 13 | DVVPIND-------TRYRLVSCNTSVITQACPKVSFEPIPIHYCAPAGFAILKCNDKQFN | 230 |
| 14 | DVVPIND-------TRYRLVSCNTSVITQACPKVSFEPIPIHYCAPAGFAILKCNDKQFI | 230 |
| 15 | DVVPIND-------TRYRLVSCNTSVITQACPKVSFEPIPIHYCAPAGFAILKCNDKQFI | 230 |
| 16 | DVVPIND-------TRYRLVSCNTSVITQACPKVSFEPIPIHYCAPAGFAILKCNDKQFI | 230 |
| 17 | DVVPIND-------TRYRLVSCNTSVITQACPKVSFEPIPIHYCAPAGFAILKCNDKQFI | 230 |
| 18 | DVVPIND-------TRYRLVSCNTSVITQACPKVSFEPIPIHYCAPAGFAILKCNDKQFI | 230 |
| 19 | DVVPIND-------TRYRLVSCNTSVITRACPKVSFEPIPIHYCAPAGFAILKCNDKQFI | 230 |
| 20 | DVVPIND-------TRYRLVSCNTSVITQACPKVSFEPIPIHYCAPAGFAILKCNDKQFI | 230 |
| 21 | DVVPIND-------TRYRLVSCNTSVITQACPKVSFEPIPIHYCAPAGFAILKCNDKQFI | 230 |
| 22 | DVVPIND-------TGYRLVSCNTSVITQACPKVSFEPIPIHYCAPAGFAILKCNDKQFI | 230 |
| 23 | DVVPIND-------TRYRLVSCNTSVITQACPKVSFEPIPIHYCAPAGFAILKCNDKQFI | 230 |
| 24 | DVVPIND-------TRYRLVSCNTSVITQACPKVSFEPIPIHYCAPAGFAILKCNDKQFI | 230 |
| 25 | DVVPIND-------TRYRLVSCNTSVITQACPKVSFEPIPIHYCAPAGFAILKCNDKQFI | 230 |
| 26 | DVVPINE-------------SCNTSVITQACPKVSFEPIPIHYCAPAGFAILKCNDKQFI | 224 |
| 27 | DVVPIND-------TRYRLVSCNTSVITQACPKVSFEPIPIHYCAPAGFAILKCNDKQFI | 230 |
| 28 | DVVPIND-------TRYRLVSCNTSVITQACPKVSFEPIPIHYCAPAGFAILKCNDKQFI | 230 |
| 29 | DVVPIND-------TRYRLVSCNTSVITQACPKVSFEPIPIHYCAPAGFAILKCNDKQFI | 230 |
| 30 | DVVPIND-------TRYRLVSCNTSVITQACPKVSFEPIPIHYCAPAGFAILKCNDKQFI | 230 |
| 31 | DVVPIND-------TRYRLVSCNTSVITQACPKVSFEPIPIHYCAPAGFAILKCNDKQFI | 230 |
| 32 | DVVPIND-------TRYRLVSCNTSVITQACPKVSFEPIPIHYCAPAGFAILKCNDKQFI | 230 |
| 33 | DVVPIND-------TRYRLVSCNTSVITQACPKVSFEPIPIHYCAPAGFAILKCNDKQFI | 230 |
| 34 | DVVPIND-------TRYRLVSCNTSVITQACPKVSFEPIPIHYCAPAGFAILKCNDKQFI | 230 |
| 35 | DVVPIND-------TRYRLVSCNTSVITQACPKVSFEPIPIHYCAPAGFAILKCNDKQFI | 230 |
| 36 | DVVPIND-------TRYRLVSCNTSVITQACPKVSFEPIPIHYCAPAGFAILKCNDKQFI | 230 |
| 37 | DVVPIND-------TRYRLVSCNTSVITQACPKVSFEPIPIHYCAPAGFAILKCNDKQFI | 230 |
| 38 | DVVPIND-------TRYRLVSCNTSVITQACPKVSFEPIPIHYCAPAGFAILKCNDKQFI | 230 |
| 39 | DVVPIND-------TRYRLVSCNTSVITQACPKVSFEPIPIHYCAPAGFAILKCNDKQFI | 230 |
| 40 | DVVPIND-------TRYRLVSCNTSVITQACPKVSFEPIPIHYCAPAGFAILKCNDKQFI | 230 |
| 41 | DVVPIND-------TRYRLVSCNTSVITQACPKVSFEPIPIHYCAPAGFAILKCNDKQFI | 230 |
| 42 | DVVPIND-------TRYRLVSCNTSVITQACPKVSFEPIPIHYCAPAGFAILKCNDKQFI | 230 |
| 43 | DVVPIND-------TRYRLVSCNTSVITQACPKVSFEPIPIHYCAPAGFAILKCNDKQFI | 230 |
| 44 | DVVPIND-------TRYRLVSCNTSVITQACPKVSFEPIPIHYCAPAGFAILKCNDKQFI | 230 |
| 45 | DVVPIND-------TRYRLVSCNTSVITQACPKVSFEPIPIHYCAPAGFAILKCNDKQFI | 230 |
| 46 | DVVPIND-------TRYRLVSCNTSVITQACPKVSFEPIPIHYCAPAGFAILKCNDKQFI | 230 |

| | | |
|---|---|---|
| | GTGPCTNVSTVQCTHGIRPVVSTQLLLNGSLAEEEVVIRSVNFSDNAKTIIVQLNKSVEI | |
| 1 | GTGPCTNVSTVQCTHGIRPVVSTQLLLNGSLAEEEVVIRSVNFSDNAKTIIVQLNKSVEI | 290 |
| 2 | GTGPCTNVSTVQCTHGIRPVVSTQLLLNGSLAEEEVVIRSVNFSDNAKTIIVQLNKSVEI | 286 |
| 3 | GTGPCTNVSTVQCTHGIRPVVSTQLLLNGSLAEEEVVIRSVNFSDNAKTIIVQLNKSVEI | 290 |
| 4 | GTGPCTNVSTVQCTHGIRPVVSTQLLLNGSLAEEEVVIRSVNFSDNAKTIIVQLNKSVEI | 290 |
| 5 | GTGPCTNVSTVQCTHGIRPVVSTQLLLNGSLAEEEVVIRSVNFSDNAKTIIVQLNKSVEI | 290 |
| 6 | GTGPCTNVSTVQCTHGIRPVVSTQLLLNGSLAEEEVVIRSVNFSDNAKTIIVQLNKSVEI | 290 |
| 7 | GTGPCTNVSTVQCTHGIRPVVSTQLLLNGSLAEEEVVIRSVNSSDNAKTIIVQLNKSVEI | 290 |
| 8 | GTGPCTNVSTVQCTHGIRPVVSTQLLLNGSLAEEEVVIRSVNFSDNAKTIIVQLNNSVEI | 290 |
| 9 | GTGPCTNVSTVQCTHGIRPVVSTQLLLNGSLAEEEVVIRSVNFSDNAKTIIVQLNKSVEI | 290 |
| 10 | GTGPCTNVSTVQCTHGIRPVVSTQLLLNGSLAEEEVVIRSVNFRDNAKTIIVQLNKSVEI | 290 |
| 11 | GTGPCTNVSTVQCTHGIRPVVSTQLLLNGSLAEEEVVIRSVNFRDNAKTIIVQLNKSVEI | 290 |
| 12 | GTGPCTNVSTVQCTHGIRPVVSTQLLLNGSLAEEEVVIRSVNFRDNAKTIIVQLNKSVEI | 290 |
| 13 | GTGPCTNVSTVQCTHGIRPVVSTQLLLNGSLAEEEVVIRSVNFSDNAKTIIVQLNKSVEI | 290 |
| 14 | GTGPCTNVSTVQCTHGIRPVVSTQLLLNGSLAEEEVVIRSVNFSDNAKTIIVQLNKSVEI | 290 |
| 15 | GTGPCTNVSTVQCTHGIRPVVSTQLLLNGSLAEEEVVIRSVNFSDNAKTIIVQLNKSVEI | 290 |
| 16 | GTGPCTNVSTVQCTHGIRPVVSTQLLLNGSLAEEEVVIRSVNFSDNAKTIIVQLNKSVEI | 290 |
| 17 | GTGPCTNVSTVQCTHGIRPVVSTQLLLNGSLAEEEVVIRSVNFSDNAKTIIVQLNKSVEI | 290 |
| 18 | GTGPCTNVSTVQCTHGIRPVVSTQLLLNGSLAEEEVVIRSVNFSDNAKTIIVQLNKSVEI | 290 |
| 19 | GTGPCTNVSTVQCTHGIRPVVSTQLLLNGSLAEEEVVIRSVNFSDNAKTIIVQLNKSVEI | 290 |
| 20 | GTGPCTNVSTVQCTHGIRPVVSTQLLLNGSLAEEEVVIRSVNFSDNAKTIIVQLNKSVEI | 290 |
| 21 | GTGPCTNVSTVQCTHGIRPVVSTQLLLNGSLAEEEVVIRSVNFSDNAKTIIVQLNKSVEI | 290 |
| 22 | GTGPCTNVSTVQCTHGIRPVVSTQLLLNGSLAEEEVVIRSVNFNDNAKTIIVQLNKSVEI | 290 |
| 23 | GTGPCTNVSTVQCTHGIRPVVSTQLLLNGSLAEEEVVIRSVNFRDNAKTIIVQLNKSVEI | 290 |
| 24 | GTGPCTNVSTVQCTHGIRPVVSTQLLLNGSLAEEEVVIRSVNFRDNAKTIIVQLNKSVEI | 290 |
| 25 | GTGPCTNVSTVQCTHGIRPVVSTQLLLNGSLAEEEVVIRSVNFSDNAKTIIVQLNKSVEI | 290 |
| 26 | GTGPCTNVSTVQCTHGIRPVVSTQLLLNGSLAEEEVVIRSVNFSDNAKTIIVQLNKSVEI | 284 |
| 27 | GTGPCTNVSTVQCTHGIRPVVSTQLLLNGSLAEEEVVIRSVNFSDNAKTIIVQLNKSVEI | 290 |
| 28 | GTGPCTNVSTVQCTHGIRPVVSTQLLLNGSLAEEEVVIRSVNFSDNAKTIIVQLNKSVEI | 290 |
| 29 | GTGPCTNVSTVQCTHGIRPVVSTQLLLNGSLAEEEVVIRSVNFSDNAKTIIVQLNKSVEI | 290 |
| 30 | GTGPCTNVSTVQCTHGIRPVVSTQLLLNGSLAEEEVVIRSVNFSDNAKTIIVQLNKSVEI | 290 |
| 31 | GTGPCTNVSTVQCTHGIRPVVSTQLLLNGSLAEEEVVIRSVNFSDNAKTIIVQLNKSVEI | 290 |
| 32 | GTGPCTNVSTVQCTHGIRPVVSTQLLLNGSLAEEEVVIRSVNFSDNAKTIIVQLNKSVEI | 290 |
| 33 | GTGPCTNVSTVQCTHGIRPVVSTQLLLNGSLAEEEVVIRSVNFSDNAKTIIVQLNKSVEI | 290 |
| 34 | GTGPCTNVSTVQCTHGIRPVVSTQLLLNGSLAEEEVVIRSVNFSDNAKTIIVQLNKSVEI | 290 |
| 35 | GTGPCTNVSTVQCTHGIRPVVSTQLLLNGSLAEEEVVIRSVNFSDNAKTIIVQLNKSVEI | 290 |
| 36 | GTGPCTNVSTVQCTHGIRPVVSTQLLLNGSLAEEEVVIRSVNFNDNAKTIIVQLNKSVEI | 290 |
| 37 | GTGPCTNVSTVQCTHGIRPVVSTQLLLNGSLAEEEVVIRSVNFSDNAKTIIVQLNKSVEI | 290 |
| 38 | GTGPCTNVSTVQCTHGIRPVVSTQLLLNGSLAEEEVVIRSVNFSDNAKTIIVQLNKSVEI | 290 |
| 39 | GTGPCTNVSTVQCTHGIRPVVSTQLLLNGSLAEEEVVIRSVNFSDNAKTIIVQLNKSVEI | 290 |
| 40 | GTGPCTNVSTVQCTHGIRPVVSTQLLLNGSLAEEEVVIRSVNFSDNAKTIIVQLNKSVEI | 290 |
| 41 | GTGPCTNVSTVQCTHGIRPVVSTQLLLNGSLAEEEVVIRSVNFSDNAKTIIVQLNKSVEI | 290 |
| 42 | GTGPCTNVSTVQCTHGIRPVVSTQLLLNGSLAEEEVVIRSVNFSDNAKTIIVQLNKSVEI | 290 |
| 43 | GTGPCTNVSTVQCTHGIRPVVSTQLLLNGSLAEEEVVIRSVNFSDNAKTIIVQLNKSVEI | 290 |
| 44 | GTGPCTNVSTVQCTHGIRPVVSTQLLLNGSLAEEEVVIRSVNFRDNAKTIIVQLNKSVEI | 290 |
| 45 | GTGPCTNVSTVQCTHGIRPVVSTQLLLNGSLAEEEVVIRSVNFRDNAKTIIVQLNKSVEI | 290 |
| 46 | GTGPCTNVSTVQCTHGIRPVVSTQLLLNGSLAEEEVVIRSVNFSDNAKTIIVQLNKSVEI | 290 |

| | Sequence | |
|---|---|---|
| | NCTRPHNNTRKSIPMGPGKAFYARGDIIGDIRKAYCEINGTEWHSTLKLVVGKLREKY-N | |
| 1 | TCTRPNNNTRKSIPMGPGKAFYARGDITGDIRKAYCEINGTEWHSTLKLVVEKLREQY-N | 349 |
| 2 | NCTRPNNNTRKSIPMGPGKAFYARGDIIGDIRKASCELDGTEWHSTLKLVVEKLRKQY-N | 345 |
| 3 | NCTRPNNNTRKSIPMGPGKAFYARGDIIGDIRKASCELDGTEWHSTLKLVVEKLRKQY-N | 349 |
| 4 | NCTRPHNNTRKSIPMGPGKAFYARGDIIGDIRKAYCEINGTEWHSTLKLVVGKLREKY-N | 349 |
| 5 | NCTRPHNNTRKSIPMGPGKAFYARGDIIGDIRKAYCEINGTEWHSTLKLVVGKLREKY-N | 349 |
| 6 | NCTRPHNNTRKSIPMGPGKAFYARGDIIGDIRKAYCEINGTEWHSTLKLVVGKLREKY-N | 349 |
| 7 | NCTRPHNNTRKSIPMGPGKAFYARGDIIGDIRKAYCEINGTEWHSTLKLVVEKLREQY-N | 349 |
| 8 | NCTRPHNNTRKSIPMGPGKAFYARGDIIGDIRKAYCKINGTEWHSTLKLVVGKLREKY-N | 349 |
| 9 | NCTRPNNNTRKSIPMGPGKAFYARGDIIGDIRKASCELDGTEWHSTLKLVVEKLRKQY-N | 349 |
| 10 | NCTRPHNNTRKSIPMGPGKAFYARGDIIGDIRKAYCEINGTEWHSTLKLVVGKLREKY-N | 349 |
| 11 | NCTRPHNNTRKSIPMGPGKAFYARGDIIGDIRKAYCTINGTEWHSTLKLVVGKLREKY-N | 349 |
| 12 | NCTRPHNNTRKSIPMGPGKAFYARGDIIGDIRKAYCEINGTEWHSTLKLVVGKLREKY-N | 349 |
| 13 | NCTRPHNNTRKSIPMGPGKAFYARGDIIGDIRKAYCNINGTEWHSTLKLVVEKLREQY-N | 349 |
| 14 | NCTRPHNNTRKSIPMGPGKAFYARGDIIGDIRKAYCEINGTEWHSTLKLVVGKLREKY-N | 349 |
| 15 | NCTRPHNNTRKSIPMGPGKAFYARGDIIGDIRKAYCEINGTEWHSTLKLVVEKLREKY-N | 349 |
| 16 | NCTRPHNNTRKSIPMGPGKAFYARGDIIGDIRKAYCEINGTEWHSTLKLVVGKLREKY-N | 349 |
| 17 | NCTRPHNNTRKSIPMGPGKAFYARGDIIGDIRKAYCEINGTEWHSTLKLVVGKLREKY-N | 349 |
| 18 | NCTRPNNNTRKSIPMGPGKAFYARGDIIGDIRKASCELDGTEWHSTLKLVVEKLRKQY-N | 349 |
| 19 | NCTRPNNNTRKSIPMGPGKAFYARGDIIGDIRKASCELNGTEWHSTLKLVVEKLRKQY-N | 349 |
| 20 | NCTRPNNNTRKSIPMGPGKAFYARGDIIGDIRKASCELDGTEWHSTLKLVVEKLRKQY-N | 349 |
| 21 | NCTRPHNNTRKSIPMGPGKAFYARGDIIGDIRKAYCEINGTEWHSTLKLVVGKLRKI--- | 347 |
| 22 | NCTRPHNNTRKSIPMGPGKAFYAKGDIIGDIRKAYCEINGTEWHSTLKLVVEKLRERY-N | 349 |
| 23 | NCTRPHNNTRKSIPMGPGKAFYARGDIIGDIRKAYCEINGTEWHSTLKLVVGKLREKY-N | 349 |
| 24 | NCTRPHNNTRKSIPMGPGKAFYARGDIIGDIRKAYCEINGTEWHSTLKLVVGKLREKY-N | 349 |
| 25 | NCTRPHNNTRKSIPMGPGKAFYARGDIIGDIRKAYCEINGTEWHSTLKLVVGKLREKY-N | 349 |
| 26 | NCTRPHNNTRKSIPMGPGKAFYARGDIIGDIRKAYCEINGTEWHSTLKLVVGKLREKY-N | 343 |
| 27 | SCTRPNNNTRKSIPMGPGKAFYARGDIIGDIRKASCELDGTEWHSTLKLVVEKLRKQY-N | 349 |
| 28 | NCTRPHNNTRKSIPMGPGKAFYARGDIIGDIRKAYCEINGTEWHSTLKLVVGKLREKY-N | 349 |
| 29 | NCTRPNNNTRKSIPMGPGKAFYARGDIIGDIRKASCELDGTEWHSTLKLVVEKLRKQY-N | 349 |
| 30 | NCTRPNNNTRKSIPMGPGKAFYARGDIIGDIRKASCELDGTEWHSTLKLVAEKLRKQY-N | 349 |
| 31 | NCTRPNNNTRKSIPMGPGKAFYARGDIIGDIRKASCELDGTEWHSTLKLVVEKLRKQY-N | 349 |
| 32 | NCTRPNNNTRKSIPMGPGKAFYARGDIIGDIRKASCELDGTEWHSTLKLVVEKLRKQY-N | 349 |
| 33 | NCTRPNNNTRKSIPMGPGKAFYARGDIIGDIRKASCELDGTEWHSTLKLVVEKLRKQY-N | 349 |
| 34 | NCTRPHNNTRKSIPMGPGKAFYARGDIIGDIRKAYCEINGTEWHSTLKLVVGKLREKY-N | 349 |
| 35 | NCTRPHNNTRKSIPMGPGKAFYARGDIIGDIRKAYCEINGTEWHSTLKLVVGKLREKY-N | 349 |
| 36 | NCTRPHNNTRKSIPMGPGKAFYAKGDIIGDIRKAYCEINGTEWHSTLKLVVEKLRERY-N | 349 |
| 37 | NCTRPHNNTRKSIPMGPGKAFYARGDIIGDIRKAYCKINGTEWHSTLKLVVEKLREQY-N | 349 |
| 38 | NCTRPHNNTRKSIPMGPGKAFYARGDIIGDIRKAYCEINGTEWHSTLKLVVGKLREKY-N | 349 |
| 39 | NCTRPHNNTRKSIPMGPGKAFYARGDIIGDIRKAYCEINGTEWHSTLKLVVGKLREKY-N | 349 |
| 40 | NCTRPHNNTRKSIPMGPGKAFYARGDIIGDIRKAYCEINGTEWHSTLKLVVGKLREKY-N | 349 |
| 41 | NCTRPNNNTRKSIPMGPGKAFYARGDIIGDIRKASCELDGTEWHSTLKLVVEKLRKQY-N | 349 |
| 42 | NCTRPNNNTRKSIPMGPGKAFYARGDIIGDIRKASCKLDGTEWHSTLKLVVEKLRKQY-N | 349 |
| 43 | NCTRPHNNTRKSIPMGPGKAFYARGDIIGDIRKAYCEINGTEWHSTLKLVVGKLREKY-N | 349 |
| 44 | NCTRPHNNTRKSIPMGPGKAFYARGDIIGNIRKAYCEINGTEWHSTLKLVVEKLREKY-N | 349 |
| 45 | NCTRPHNNTRKSIPMGPGKAFYARGDIIGDIRKAYCEINGTEWHSTLKLVVGKLREKY-N | 349 |
| 46 | NCTRPHNNTRKSIPMGPGKAFYARGDIIGDIRKAYCEINGTEWHSTLKLVVGKLREKY-N | 349 |

| | KTIVFNRSSGGDPEIVMYSFNCGGEFFYCNSTKLFNSTWPWND-TKGSHDTNDTLMLPCR | |
|---|---|---|
| 1 | KTIVFNRSSGGDPEIVMYSFNCGGEFFYCNSTKLFNSTWPWND-TKGSHDTNGTLILPCK | 408 |
| 2 | KTIIFNRSSGGDPEIVMYSFNCGGEFFYCNSTKLFNSTWPWND-TKGSHDTNGTLILPCR | 404 |
| 3 | KTIVFNRSSGGDPEIVMYSFNCGGEFFYCNSTKLFNSTWPWND-TKGSHDTNGTLILPCR | 408 |
| 4 | KTIVFNRSSGGDPEIVMYSFNCGGEFFYCNSTKLFNSTWPWND-TKGSHDTNDTLMLPCK | 408 |
| 5 | KTIVFNRSSGGDPEIVMYSFNCGGEFFYCNSTKLFNSTWPWND-TKGSHDTNDTLMLPCK | 408 |
| 6 | KTIVFNRSSGGDPEIVMYSFNCGGEFFYCNSTKLFNSTWPWND-TKGSHDTNDTLMLPCK | 408 |
| 7 | KTIVFNRSSGGDPEIVMYSFNCGGEFFYCNSTKLFNSTWPWND-TKGSHDTNGTLILPCR | 408 |
| 8 | KTIVFNRSSGGDPEIVMYSFNCGGEFFYCNSTKLFNSTWPWND-TKGSHDTNDTLMLPCK | 408 |
| 9 | KTIVFNRSSGGDPEIVMYSFNCGGEFFYCNSTKLFNSTWPWND-TKGSHDTNGTLILPCR | 408 |
| 10 | KTIVFNRSSGGDPEIVMYSFNCGGEFFYCNSTKLFNSTWPWND-TKGSHDTNDTLMLPCR | 408 |
| 11 | KTIVFKRSSGGDPEIVMYSFNCGGEFFYCNSTKLFNSTWPWND-TKGSHDTNDTLMLPCK | 408 |
| 12 | KTIVFNRSSGGDPEIVMYSFNCGGEFFYCNSTKLFNSTWPWND-TKGSHDTNDTLMLPCR | 408 |
| 13 | KTIVFNRSSGGDPEIVMYSFNCGGEFFYCNSTKLFNSTWPWND-TKGSHDTNDTLMLPCK | 408 |
| 14 | KTIVFNRSSGGDPEIVMYSFNCGGEFFYCNSTKLFNSTWPWND-TKGSHDTNDTLMLPCK | 408 |
| 15 | KTIVFNRSSGGDPEIVMYSFNCGGEFFYCNSTKLFNSTWPWND-TKGSHDTNDTLMLPCK | 408 |
| 16 | KTIVFNRSSGGDPEIVMYSFNCGGEFFYCNSTKLFNSTWPWND-TKGSHDTNDTLMLPCK | 408 |
| 17 | KTIVFNRSSGGDPEIVMYSFNCGGEFFYCNSTKLFNSTWPWND-TKGSHDTNDTLMLPCK | 408 |
| 18 | KTTVFNRSSGGDPEIVMYSFNCGGEFFYCNSTKLFNSTWPWND-TKGSHDTNGTLILPCR | 408 |
| 19 | KTIVFNRSSGGDPEIVMYSFNCGGEFFYCNSTKLFNSTPWNDT--KGSHDTNGTLILPCR | 407 |
| 20 | KTIVFNRSSGGDPEIVMYSFNCGGEFFYCNSTKLFNSTWPWND-TKGSHDTNGTLILPCR | 408 |
| 21 | -------SSGGDPEIVMYSFNCGGEFFYCNSTKLFNSTWPWND-TKGSHDTNDTLMLPCK | 399 |
| 22 | KTIVFNRSSGGDPEIVMYSFNCGGEFFYCNSTKLFNSTWPWND-TKGSHDTNGTLTLPCR | 408 |
| 23 | KTIVFNRSSGGDPEIVMYSFNCGGEFFYCNSTKLFNSTWPWND-TKGSHDTNDTLMLPCR | 408 |
| 24 | KTIVFNRSSGGDPEIVMYSFNCGGEFFYCNSTKLFNSTWPWND-TKGSHDTNDTLMLPCR | 408 |
| 25 | KTIVFNRSSGGDPEIVMYSFNCGGEFFYCNSTKLFNSTWPWND-TKGSHDTNDTLMLPCK | 408 |
| 26 | KTIVFNRSSGGDPEIVMYSFNCGGEFFYCNSTKLFNSTWPWND-TKGSHDTNDTLMLPCK | 402 |
| 27 | KTIVFNRSSGGDPEIVMYSFNCGGEFFYCNSTKLFNSTWPWND-TKGSHDTNGTLILPCR | 408 |
| 28 | KTIVFNRSSGGDPEIVMYSFNCGGEFFYCNSTKLFNSTWPWND-TKGSHDTNDTLMLPCK | 408 |
| 29 | KTIVFNRSSGGDPEIVMYSFNCGGEFFYCNSTKLFNSTWPWND-TKGSHDTNGTLILPCR | 408 |
| 30 | KTIVFNRSSGGDPEIVMYSFNCGGEFFYCNSTKLFNSTWPWND-TKGSHDTNGTLILPCR | 408 |
| 31 | KTIVFNRSSGGDPEIVMYSFNCGGEFFYCNSTKLFNSTWPWND-TKGSHDTNGTLILPCR | 408 |
| 32 | KTIVFNRSSGGDPEIVMYSFNCGGEFFYCNSTKLFNSTWPWND-TKGSHDTNGTLILPCR | 408 |
| 33 | KTIVFNRSSGGDPEIVMYSFNCGGEFFYCNSTKLFNSTWPWND-TKGSHDTNGTLILPCR | 408 |
| 34 | KTIVFNRSSGGDPEIVMYSFNCGGEFFYCNSTKLFNSTWPWND-TKGSHDTNDTLMLPCK | 408 |
| 35 | KTIVFNRSSGGDPEIVMYSFNCGGEFFYCNSTKLFNSTWPWND-TKGSHDTNDTLMLPCK | 408 |
| 36 | KTIVFNRSSGGDPEIVMYSFNCGGEFFYCNSTKLFNSTWPWND-TKGSHDTNGTLTLPCR | 408 |
| 37 | KTIVFNRSSGGDPEIVMYSFNCGGEFFYCNSTKLFNSTWPWND-TKGSHDTNDTLMLPCR | 408 |
| 38 | KTIVFNRSSGGDPEIVMYSFNCGGEFFYCNSTKLFNSTWPWND-TKGSHDTNDTLMLPCK | 408 |
| 39 | KTIVFNRSSGGDPEIVMYSFNCGGEFFYCNSTKLFNSTWPWND-TKGSHDTNDTLMLPCK | 408 |
| 40 | KTIVFNRSSGGDPEIVMYSFNCGGEFFYCNSTKLFNSTWPWND-TKGSHDTNDTLMLPCK | 408 |
| 41 | KTIIFNRSSGGDPEIVMYSFNCGGEFFYCNSTKLFNSTWPWND-TKGSHDTNGTLILPCR | 408 |
| 42 | KTIVFNRSSGGDPEIVMYSFNCGGEFFYCNSTKLFNSTWPWND-TKGSHDTNGTLILPCR | 408 |
| 43 | KTIVFNRSSGGDPEIVMYSFNCGGEFFYCNSTKLFNSTWPWND-TKGSHDTNDTLMLPCK | 408 |
| 44 | KTIVFNRSSGGDPEIVMYSFNCGGEFFYCNSTKLFNSTWPWND-TKGSHDTNGTLILPCR | 408 |
| 45 | KTIVFNRSSGGDPEIVMYSFNCGGEFFYCNSTKLFNSTWPWND-TKGSHDTNDTLMLPCR | 408 |
| 46 | KTIVFNRSSGGDPEIVM-SFNCGGEFFYCNSTKLFNSTWPWND-TKGSHDTNDTLMLPCK | 407 |

| # | Sequence | End |
|---|---|---|
| | IKQIINMWQGVGKAMYAPPIEGXIRCSSNITGLLLTRDGG-YESNETDEIFRPGGGDMRD | |
| 1 | IKQIINMWQGVGKAMYAPPIEGKIRCSSNITGLLLTRDGG-YESNETDEIFRPGGGDMRD | 467 |
| 2 | IKQIINMWQGVGKAMYAPPIEGEIRCSSNITGLLLTRDGG-YESNETDEIFRPGGGDMRD | 463 |
| 3 | IKQIINMWQGVGKAMYAPPIEGEIRCSSNITGLLLTRDGG-YESNETDEIFRPGGGDMRD | 467 |
| 4 | IKQIINMWQGVGKAMYAPPIEGQIRCSSNITGLLLTRDGG-YESNETDEIFRPGGGDMRD | 467 |
| 5 | IKQIINMWQGVGKAMYAPPIEGQIRCSSNITGLLLTRDGG-YESNETDEIFRPGGGDMRD | 467 |
| 6 | IKQIINMWQGVGKAMYAPPIEGQIRCSSNITGLLLTRDGG-YESNETDEIFRPGGGDMRD | 467 |
| 7 | IKQIINMWQGVGKAMYAPPIEGEIRCSSNITGLLLTRDGG-YESNKTDEIFRPGGGDMRD | 467 |
| 8 | IKQIINMWQGVGKAMYAPPIEGQIRCSSNITGLLLTRDGG-YESNETDEIFRPGGGDMRD | 467 |
| 9 | IKQIINMWQGVGKAMYAPPIEGEIRCS-----LLLTRDGG-YESNETDEIFRPGGGDMRD | 462 |
| 10 | IKQIINMWQGVGKAMYAPPIEGIIRCSSNITGLLLTRDGG-YESNETDEIFRPGGGDMRD | 467 |
| 11 | IKQIINMWQGVGKAMYAPPIEGQIRCSSNITGLLLTRDGG-YESNETDEIFRPGGGDMRD | 467 |
| 12 | IKQIINMWQGVGKVMYAPPIEGIIRCSSNITGLLLTRDGG-YESNETDEIFRPGGGDMRD | 467 |
| 13 | IKQIINMWQGVGKAMYAPPIEGQIRCSSNITGLLLTRDGG-YESNKTDEIFRPGGGDMRD | 467 |
| 14 | IKQIINMWQGVGKAMYAPPIEGQIRCSSNITGLLLTRDGG-YESNETDEIFRPGGGDMRD | 467 |
| 15 | IKQIINMWQGVGKAMYAPPIEGQIRCSSNITGLLLTRDGG-YESNETDEIFRPGGGDMRD | 467 |
| 16 | IKQIINMWQGVGKAMYAPPIEGQIRCSSNITGLLLTRDGG-YESNETDEIFRPGGGDMRD | 467 |
| 17 | IKQIINMWQGVGKAMYAPPIEGQIRCSSNITGLLLTRDGG-YESNETDEIFRPGGGDMRD | 467 |
| 18 | IKQIINMWQGVGKAMYAPPIEGEIRCSSNITGLLLTRDGG-YESNETDEIFRPGGGDMRD | 467 |
| 19 | IKQIINMWQGVGKAMYAPPIEGEIRCSSNITGLLLTRDGG-YESNETDEIFRPGGGDMRD | 466 |
| 20 | IKQIINMWQGVGKAMYAPPIEGEIRCSSNITGLLLTRDGG-YESNETDEIFRPGGGDMRD | 467 |
| 21 | IKQIINMWQGVGKAMYAPPIEGQIRCSSNITGLLLTRDGG-YESNETDEIFRPGGGDMRD | 458 |
| 22 | IKQIINMWQGVGKAMYAPPIEGKIRCSSNITGLLLTRDGG-YESNKTDEIFRPGGGDMRD | 467 |
| 23 | IKQIINMWQGVGKAMYAPPIEGIIRCSSNITGLLLTRDGG-YESNETDEIFRPGGGDMRD | 467 |
| 24 | IKQIINMWQGVGKAMYAPPIEGIIRCSSNITGLLLTRDGG-YESNETDEIFRPGGGDMRD | 467 |
| 25 | IKQIINMWQGVGKAMYAPPIEGQIRCSSNITGLLLTRDGG-YESNETDEIFRPGGGDMRD | 467 |
| 26 | IKQIINMWQGVGKAMYAPPIEGQIRCSSNITGLLLTRDGG-YESNETDEIFRPGGGDMRD | 461 |
| 27 | IKQIINMWQGVGKAMYAPPIEGEIRCSSNITGLLLTRDGG-YESNETDEIFRPGGGDMRD | 467 |
| 28 | IKQIINMWQGVGKAMYAPPIEGQIRCSSNITGLLLTRDGG-YESNETDEIFRPGGGDMRD | 467 |
| 29 | IKQIINMWQGVGKAMYAPPIEGEIRCSSNITGLLLTRDGG-YESNETDEIFRPGGGDMRD | 467 |
| 30 | IKQIINMWQGVGKAMYAPPIEGEIRCSSNITGLLLTRDGG-YESNETDEIFRPGGGDMRD | 467 |
| 31 | IKQIINMWQGVGKAMYAPPIEGEIRCSSNITGLLLTRDGG-YESNETDEIFRPGGGDMRD | 467 |
| 32 | IKQIINMWQGVGKAMYAPPIEGEIRCSSNITGLLLTRDGG-YESNETDEIFRPGGGDMRD | 467 |
| 33 | IKQIINMWQGVGKAMYAPPIEGEIRCSSNITGLLLTRDGG-YESNETDEIFRPGGGDMRD | 467 |
| 34 | IKQIINMWQGVGKAMYAPPIEGQIRCSSNITGLLLTRDGG-YESNETDEIFRPGG----- | 462 |
| 35 | IKQIINMWQGVGKAMYAPPIEGQIRCSSNITGLLLTRDGG-YESNETDEIFRPGGGDMRD | 467 |
| 36 | IKQIINMWQGVGKAMYAPPIEGKIRCSSNITGLLLTRDGG-YESNKTDEIFRPGGGDMRD | 467 |
| 37 | IKQIINMWQGVGKAMYAPPIEGIIRCSSNITGLLLTRDGG-YESNETDEIFRPGGGDMRD | 467 |
| 38 | IKQIINMWQGVGKAMYAPPIEGQIRCSSNITGLLLTRDGG-YESNETDEIFRPGGGDMRD | 467 |
| 39 | IKQIINMWQGVGKAMYAPPIEGQIRCSSNITGLLLTRDGG-YESNETDEIFRPGGGDMRD | 467 |
| 40 | IKQIINMWQGVGKAMYAPPIEGQIRCSSNITGLLLTRDGG-YESNETDEIFRPGGGDMRD | 467 |
| 41 | IKQIINMWQGVGKAMYAPPIEGEIRCSSNITGLLLTRDGG-YESNETDEIFRPGGGDMRD | 467 |
| 42 | IKQIINMWQGVGKAMYAPPIEGEIRCSSNITGLLLTRDGG-YESNETDEIFRPGGGDMRD | 467 |
| 43 | IKQIINMWQGVGKAMYAPPIEGQIRCSSNITGLLLTRDGG-YESNETDEIFRPGGGDMRD | 467 |
| 44 | IKQIINMWQGVGKAMYAPPIEGEIRCSSNITGLLLTRDGG-YESNETDEIFRPGGGDMRD | 467 |
| 45 | IKQIINMWQGVGKAMYAPPIEGIIRCSSNITGLLLTRDGG-YESNETDEIFRPGGGDMRD | 467 |
| 46 | IKQIINMWQGVGKAMYAPPIEGQIRCSSNITGLLLTRDGG-YESNETDEIFRPGGGDMRD | 466 |

| | | |
|---|---|---|
| | NWRSELYKYKVVKIEPLGVAPTKAKRRVVQREKRAFG-LGAVFLGFLGAAGSTMGAASIT | |
| 1 | NWRSELYKYKVVKIEPLGVAPTKAKRRVVQREKRAFG-LGAVFLGFLGAAGSTMGAASIT | 526 |
| 2 | NWRSELYKYKVVKIEPLGVAPTKAKRRVVQREKRAFG-LGAVFLGFLGAAGSTMGAASIT | 522 |
| 3 | NWRSELYKYKVVKIEPLGVAPTKAKRRVVQREKRAFG-LGAVFLGFLGAAGSTMGAASIT | 526 |
| 4 | NWRSELYKYKVVKIEPLGVAPTKAKRRVVQREKRAFG-LGAVFLGFLGAAGSTMGAASIT | 526 |
| 5 | NWRSELYKYKVVKIEPLGVAPTKAKRRVVQREKRAFG-LGAVFLGFLGAAGSTMGAASIT | 526 |
| 6 | NWRSELYKYKVVKIEPLGVAPTKAKRRVVQREKRAFG-LGAVFLGFLGAAGSTMGAASIT | 526 |
| 7 | NWRSELYKYKVVKIEPLGVAPTKAKRRVVQREKRAFG-LGAVFLGFLGAAGSTMGAASIT | 526 |
| 8 | NWRSKLYKYKVVKIEPLGVAPTKAKRRVVQREKRAFE-LGAVFLGFLGAAGSTMGAASIT | 526 |
| 9 | NWRSELYKYKVVKIEPLGVAPTKAKRRVVQREKRAFG-LGAVFLGFLGAAGSTMGAASIT | 521 |
| 10 | NWRSELYKYKVVKIEPLGVAPTKAKRRVVQREKRAFG-LGAVFLGFLGAAGSTMGAASIT | 526 |
| 11 | NWRSELYKYKVVKIEPLGVAPTKAKRRVVQREKRAFG-LGAVFLGFLGAAGSTMGAASIT | 526 |
| 12 | NWRSELYKYKVVKIEPLGVAPTKAKRRVVQREKRAFG-LGAVFLGFLGAAGSTMGAASIT | 526 |
| 13 | NWRSELYKYKVVKIEPLGVAPTKAKRRVVQREKRAFG-LGAVFLGFLGAAGSTMGAASIT | 526 |
| 14 | NWRSELYKYKVVKIEPLGVAPTKAKRRVVQREKRAFG-LGAVFLGFLGAAGSTMGAASIT | 526 |
| 15 | NWRSELYKYKVVKIEPLGVAPTKAKRRVVQREKRAFG-LGAVFLGFLGAAGSTMGAASIT | 526 |
| 16 | NWRSELYKYKVVKIEPLGVAPTKAKRRVVQREKRAFG-LGAVFLGFLGAAGSTMGAASIT | 526 |
| 17 | NWRSELYKYKVVKIEPLGVAPTKAKRRVVQREKRAFG-LGAVFLGFLGAAGSTMGAASIT | 526 |
| 18 | NWRSELYKYKVVKIEPLGVAPTKAKRRVVQREKRAFG-LGAVFLGFLGAAGSTMGAASIT | 526 |
| 19 | NWRSELYKYKVVKIEPLGVAPTKAKRRVVQREKRAFG-LGAVFLGFLGAAGSTMGAASIT | 525 |
| 20 | NWRSELYKYKVVKIEPLGVAPTKAKRRVVQREKRAFG-LGAVFLGFLGAAGSTMGAASIT | 526 |
| 21 | NWRSELYKYKVVKIEPLGVAPTKAKRRVVQREKRAFG-LGAVFLGFLGAAGSTMGAASIT | 517 |
| 22 | NWRSELYKYKVVKIEPLGVAPTKAKRRVVQREKRAFG-LGAVFLGFLGAAGSTMGAASIT | 526 |
| 23 | NWRSELYKYKVVKIEPLGVAPTKAKRRVVQREKRAFG-LGAVFLGFLGAAGSTMGAASIT | 526 |
| 24 | NWRSELYKYKVVKIEPLGVAPTKAKRRVVQREKRAFG-LGAVFLGFLGAAGSTMGAASIT | 526 |
| 25 | NWRSELYKYKVVKIEPLGVAPTKAKRRVVQREKRAFG-LGAVFLGFLGAAGSTMGAASIT | 526 |
| 26 | NWRSELYKYKVVKIEPLGVAPTKAKRRVVQREKRAFG-LGAVFLGFLGAAGSTMGAASIT | 520 |
| 27 | NWRSELYKYKVVKIEPLGVAPTKAKRRVVQREKRAFG-LGAVFLGFLGAAGSTMGAASIT | 526 |
| 28 | NWRSELYKYKVVKIEPLGVAPTKAKRRVVQREKRAFG-LGAVFLGFLGAAGSTMGAASIT | 526 |
| 29 | NWRSELYKYKVVKIEPLGVAPTKAKRRVVQREKRAFG-LGAVFLGFLGAAGSTMGAASIT | 526 |
| 30 | NWRSELYKYKVVKIEPLGVAPTKAKRRVVQREKRAFG-LGAVFLGFLGAAGSTMGAASIT | 526 |
| 31 | NWRSELYKYKVVKIEPLGVAPTKAKRRVVQREKRAFG-LGAVFLGFLGAAGSTMGAASIT | 526 |
| 32 | NWRSELYKYKVVKIEPLGVAPTKAKRRVVQREKRAFG-LGAVFLGFLGAAGSTMGAASIT | 526 |
| 33 | NWRSELYKYKVVKIEPLGVAPTKAKRRVVQREKRAFG-LGAVFLGFLGAAGSTMGAASIT | 526 |
| 34 | ---SELYKYKVVKIEPLGVAPTKAKRRVVQREKRAFG-LGAVFLGFLGAAGSTMGAASIT | 518 |
| 35 | NWRSELYKYKVVKIEPLGVAPTKAKRRVVQREKRAFG-LGAVFLGFLGAAGSTMGAASIT | 526 |
| 36 | NWRSELYKYKVVKIEPLGVAPTKAKRRVVQREKRAFG-LGAVFLGFLGAAGSTMGAASIT | 526 |
| 37 | NWRSELYKYKVVKIEPLGVAPTRAKRRVVQREKRAFG-LGAVFLGFLGAAGSTMGAASIT | 526 |
| 38 | NWRSELYKYKVVKIEPLGIAPTKAKRRVVQREKRAFG-LGAVFLGFLGAAGSTMGAASIT | 526 |
| 39 | NWRSELYKYKVVKIEPLGVAPTKAKRRVVQREK-AFG-LGAVFLGFLGAAGSTMGAASIT | 525 |
| 40 | NWRSELYKYKVVKIEPLGVAPTKAKRRVVQREKRAFG-LGAVFLGFLGAAGSTMGAASIT | 526 |
| 41 | NWRSELYKYKVVKIEPLGVAPTKAKRRVVQREKRAFG-LGAVFLGFLGAAGSTMGAASIT | 526 |
| 42 | NWRSELYKYKVVKIEPLGVAPTKAKRRVVQREKRAFG-LGAVFLGFLGAAGSTMGAASIT | 526 |
| 43 | NWRSELYKYKVVKIEPLGVAPTKAKRRVVQREKRAFG-LGAVFLGFLGAAGSTMGAASIT | 526 |
| 44 | NWRSELYKYKVVKIEPLGVAPTKAKRRVVQREKRAFG-LGAVFLGFLGAAGSTMGAASIT | 526 |
| 45 | NWRSELYKYKVVKIEPLGVAPTKAKRRVVQREKRAFG-LGAVFLGFLGAAGSTMGAASIT | 526 |
| 46 | NWRSELYKYKVVKIEPLGVAPTKAKRRVVQREKRAFG-LGAVFLGFLGAAGSTMGAASIT | 525 |

| | Sequence | |
|---|---|---|
| | LTVQARQLLSGIVQQQNNLLRAIEAQQHLLQLTVWGIKQLQARVLAVERYLKDQQLLGIW | |
| 1 | LTVQARQLLSGIVQQQNNLLRAIEAQQHLLQLTVWGIKQLQARVLAVERYLKDQQLLGIW | 586 |
| 2 | LTVQARQLLSGIVQQQNNLLRAIEAQQHLLQLTVWGIKQLQARVLAVERYLKDQQLLGIW | 582 |
| 3 | LTVQARQLLSGIVQQQNNLLRAIEAQQHLLQLTVWGIKQLQARVLAVERYLKDQQLLGIW | 586 |
| 4 | LTVQARQLLSGIVQQQNNLLRAIEAQQHLLQLTVWGIKQLQARVLAVERYLKDQQLLGIW | 586 |
| 5 | LTVQARQLLSGIVQQQNNLLRAIEAQQHLLQLTVWGIKQLQARVLAVERYLKDQQLLGIW | 586 |
| 6 | LTVQARQLLSGIVQQQNNLLRAIEAQQHLLQLTVWGIKQLQARVLAVERYLKDQQLLGIW | 586 |
| 7 | LTVQARQLLSGIVQQQNNLLRAIEAQQHLLQLTVWGIKQLQARVLAVERYLKDQQLLGIW | 586 |
| 8 | LTVQARQLLSGIVQQQNNLLRAIEAQQHLLQLTVWGIKQLQARVLAVERYLKDQQLLGIW | 586 |
| 9 | LTVQARQLLSGIVQQQNNLLRAIEAQQHLLQLTVWGIKQLQARVLAVERYLKDQQLLGIW | 581 |
| 10 | LTVQARQLLSGIVQQQNNLLRAIEAQQHLLQLTVWGIKQLQARVLAVERYLKDQQLLGIW | 586 |
| 11 | LTVQARQLLSGIVQQQNNLLRAIEAQQHLLQLTVWGIKQLQARVLAVERYLKDQQLLGIW | 586 |
| 12 | LTVQARQLLSGIVQQQNNLLRAIEAQQHLLQLTVWGIKQLQARVLAVERYLKDQQLLGIW | 586 |
| 13 | LTVQARQLLSGIVQQQNNLLRAIEAQQHLLQLTVWGIKQLQARVLAVERYLKDQQLLGIW | 586 |
| 14 | LTVQARQLLSGIVQQQNNLLRAIEAQQHLLQLTVWGIKQLQARVLAVERYLKDQQLLGIW | 586 |
| 15 | LTVQARQLLSGIVQQQNNLLRAIEAQQHLLQLTVWGIKQLQARVLAVERYLKDQQLLGIW | 586 |
| 16 | LTVQARQLLSGIVQQQNNLLRAIEAQQHLLQLTVWGIKQLQARVLAVERYLKDQQLLGIW | 586 |
| 17 | LTVQARQLLSGIVQQQNNLLRAIEAQQHLLQLTVWGIKQLQARVLAVERYLKDQQLLGIW | 586 |
| 18 | LTVQARQLLSGIVQQQNNLLRAIEAQQHLLQLTVWGIKQLQARVLAVERYLKDQQLLGIW | 586 |
| 19 | LTVQARQLLSGIVQQQNNLLRAIEAQQHLLQLTVWGIKQLQARVLAVERYLKDQQLLGIW | 585 |
| 20 | LTVQARQLLSGIVQQQNNLLRAIEAQQHLLQLTVWGIKQLQARVLAVERYLKDQQLLGIW | 586 |
| 21 | LTVQARQLLSGIVQQQNNLLRAIEAQQHLLQLTVWGIKQLQARVLAVERYLKDQQLLGIW | 577 |
| 22 | LTVQARQLLSGIVQQQNNLLRAIEAQQHLLQLTVWGIKQLQARVLAVERYLKDQQLLGIW | 586 |
| 23 | LTVQARQLLSGIVQQQNNLLRAIEAQQHLLQLTVWGIKQLQARVLAVERYLKDQQLLGIW | 586 |
| 24 | LTVQARQLLSGIVQQQNNLLRAIEAQQHLLQLTVWGIKQLQARVLAVERYLKDQQLLGIW | 586 |
| 25 | LTVQARQLLSGIVQQQNNLLRAIEAQQHLLQLTVWGIKQLQARVLAVERYLKDQQLLGIW | 586 |
| 26 | LTVQARQLLSGIVQQQNNLLRAIEAQQHLLQLTVWGIKQLQARVLAVERYLKDQQLLGIW | 580 |
| 27 | LTVQARQLLSGIVQQQNNLLRAIEAQQHLLQLTVWGIKQLQARVLAVERYLKDQQLLGIW | 586 |
| 28 | LTVQARQLLSGIVQQQNNLLRAIEAQQHLLQLTVWGIKQLQARVLAVERYLKDQQLLGIW | 586 |
| 29 | LTVQARQLLSGIVQQQNNLLRAIEAQQHLLQLTVWGIKQLQARVLAVERYLKDQQLLGIW | 586 |
| 30 | LTVQARQLLSGIVQQQNNLLRAIEAQQHLLQLTVWGIKQLQARVLAVERYLKDQQLLGIW | 586 |
| 31 | LTVQARQLLSGIVQQQNNLLRAIEAQQHLLQLTVWGIKQLQARVLAVERYLKDQQLLGIW | 586 |
| 32 | LTVQARQLLSGIVQQQNNLLRAIEAQQHLLQLTVWGIKQLQARVLAVERYLKDQQLLGIW | 586 |
| 33 | LTVQARQLLSGIVQQQNNLLRAIEAQQHLLQLTVWGIKQLQARVLAVERYLKDQQLLGIW | 586 |
| 34 | LTVQARQLLSGIVQQQNNLLRAIEAQQHLLQLTVWGIKQLQARVLAVERYLKDQQLLGIW | 578 |
| 35 | LTVQARQLLSGIVQQQNNLLRAIEAQQHLLQLTVWGIKQLQARVLAVERYLKDQQLLGIW | 586 |
| 36 | LTVQARQLLSGIVQQQNNLLRAIEAQQHLLQLTVWGIKQLQARVLAVERYLKDQQLLGIW | 586 |
| 37 | LTVQARQLLSGIVQQQNNLLRAIEAQQHLLQLTVWGIKQLQARVLAVERYLKDQQLLGIW | 586 |
| 38 | LTVQARQLLSGIVQQQNNLLRAIEAQQHLLQLTVWGIKQLQARVLAVERYLKDQQLLGIW | 586 |
| 39 | LTVQARQLLSGIVQQQNNLLRAIEAQQHLLQLTVWGIKQLQARVLAVERYLKDQQLLGIW | 585 |
| 40 | LTVQARQLLSGIVQQQNNLLRAIEAQQHLLQLTVWGIKQLQARVLAVERYLKDQQLLGIW | 586 |
| 41 | LTVQARQLLSGIVQQQNNLLRAIEAQQHLLQLTVWGIKQLQARVLAVERYLKDQQLLGIW | 586 |
| 42 | LTVQARQLLSGIVQQQNNLLRAIEAQQHLLQLTVWGIKQLQARVLAVERYLKDQQLLGIW | 586 |
| 43 | LTVQARQLLSGIVQQQNNLLRAIEAQQHLLQLTVWGIKQLQARVLAVERYLKDQQLLGIW | 586 |
| 44 | LTVQARQLLSGIVQQQNNLLRAIEAQQHLLQLTVWGIKQLQARVLAVERYLKDQQLLGIW | 586 |
| 45 | LTVQARQLLSGIVQQQNNLLRAIEAQQHLLQLTVWGIKQLQARVLAVERYLKDQQLLGIW | 586 |
| 46 | LTVQARQLLSGIVQQQNNLLRAIEAQQHLLQLTVWGIKQLQARVLAVERYLKDQQLLGIW | 585 |

| | | |
|---|---|---|
| | GCSGKLICTTTVPWNTSWSNKSLEQIWDNMTWMEWEREIDNYTGYIYQLIEESQNQQEKN | |
| 1 | GCSGKLICTTTVPWNTSWSNKSLEQIWDNMTWMEWEREIDNYTGYIYQLIEESQNQQEKN | 646 |
| 2 | GCSGKLICPTTVPWNTSWSNKSLEQIWDNMTWMEWEREIDNYTGYIYQLIEESQNQQEKN | 642 |
| 3 | GCSGKLICPTTVPWNTSWSNKSLEQIWDNMTWMEWEREIDNYTGYIYQLIEESQNQQEKN | 646 |
| 4 | GCSGKLICTTTVPWNTSWSNKSLEQIWDNMTWMEWEREIDNYTGYIYQLIEESQNQQEKN | 646 |
| 5 | GCSGKLICTTTVPWNTSWSNKSLEQIWDNMTWREWEREIDNYTGYIYQLIEESQNQQEKN | 646 |
| 6 | GCSGKLICTTTVPWNTSWSNKSLEQIWDNMTWMEWEREIDNYTGYIYQLIEESQNQQEKN | 646 |
| 7 | GCSGKLICTTTVPWNTSWSNKSLEQIWDNMTWMEWEREIDNYTGYIYQLIEESQNQQEKN | 646 |
| 8 | GCSGKLICTTTVPWNTSWSNKSLEQIWDNMTWMEWEREIDNYTGYIYQLIEESQNQQEKN | 646 |
| 9 | GCSGKLICPTTVPWNTSWSNKSLEQIWDNMTWMEWEREIDNYTGYIYQLIEESQNQQEKN | 641 |
| 10 | GCSGKLICTTTVPWNTSWSNKSLEQIWDNMTWMEWEREIDNYTGYIYQLIEESQNQQEKN | 646 |
| 11 | GCSGKLICTTTVPWNTSWSNKSLEQIWDNMTWMEWEREIDNYTGYIYQLIEESQNQQEKN | 646 |
| 12 | GCSGKLICTTTVPWNTSWSNKSLEQIWDNMTWMEWEREIDNYTGYIYQLIEESQNQQEKN | 646 |
| 13 | GCSGKLICTTTVPWNTSWSNKSLEQIWDNMTWMEWEREIDNYTGYIYQLIEESQNQQEKN | 646 |
| 14 | GCSGKLICTTTVPWNTSWSNKSLEQIWDNMTWMEWEREIDNYTGYIYQLIEESQNQQEKN | 646 |
| 15 | GCSGKLICTTTVPWNTSWSNKSLEQIWDNMTWREWEREIDNYTGYIYQLIEESQNQQEKN | 646 |
| 16 | GCSGKLICTTTVPWNTSWSNKSLEQIWDNMTWMEWEREIDNYTGYIYQLIEESQNQQEKN | 646 |
| 17 | GCSGKLICTTTVPWNTSWSNKSLEQIWDNMTWMEWEREIDNYTGYIYQLIEESQNQQEKN | 646 |
| 18 | GCSGKLICPTTVPWNTSWSNKSLEQIWDNMTWMEWEREIDNYTGYIYQLIEESQNQQEKN | 646 |
| 19 | GCSGKLICPTTVPWNTSWSNKSLEQIWDNMTWMEWEREIDNYTGYIYQLIEESQNQQEKN | 645 |
| 20 | GCSGKLICPTTVPWNTSWSNKSLEQIWDNMTWMEWEREIDNYTGYIYQLIEESQNQQEKN | 646 |
| 21 | GCSGKLICTTTVPWNTSWSNKSLEQIWDNMTWMEWEREIDNYTGYIYQLIEESQNQQEKN | 637 |
| 22 | GCSGKLICTTTVPWNTSWSNKSLEQIWDNMTWMEWEREIDNYTGYIYQLIEESQNQQEKN | 646 |
| 23 | GCSGKLICTTTVPWNTSWSNKSLEQIWDNMTWMEWEREIDNYTGYIYQLIEESQNQQEKN | 646 |
| 24 | GCSGKLICTTTVPWNTSWSNKSLEQIWDNMTWMEWEREIDNYTGYIYQLIEESQNQQEKN | 646 |
| 25 | GCSGKLICTTTVPWNTSWSNKSLEQIWDNMTWMEWEREIDNYTGYIYQLIEESQNQQEKN | 646 |
| 26 | GCSGKLICTTTVPWNTSWSNKSLEQIWDNMTWMEWEREIDNYTGYIYQLIEESQNQQEKN | 640 |
| 27 | GCSGKLICPTTVPWNTSWSNKSLEQIWDNMTWMEWEREIDNYTGYIYQLIEELQNQQEKN | 646 |
| 28 | GCSGKLICTTTVPWNTSWSNKSLEQIWDNMTWMEWEREIDNYTGYIYQLIEESQNQQEKN | 646 |
| 29 | GCSGKLICPTTVPWNTSWSNKSLEQIWDNMTWMEWEREIDNYTGYIYQLIEESQNQQEKN | 646 |
| 30 | GCSGKLICPTTVPWNTSWSNKSLEQIWDNMTWMEWEREIDNYTGYIYQLIEESQNQQEKN | 646 |
| 31 | GCSGKLICPTTVPWNTSWSNKSLEQIWDNMTWMEWEREIDNYTGYIYQLIEESQNQQEKN | 646 |
| 32 | GCSGKLICPTTVPWNTSWSNKSLEQIWDNMTWMEWEREIDNYTGYIYQLIEESQNQQEKN | 646 |
| 33 | GCSGKLICPTTVPWNTSWSNKSLEQIWDNMTWMEWEREIDNYTGYIYQLIEESQNQQEKN | 646 |
| 34 | GCSGKLICTTTVPWNTSWSNKSLEQIWDNMTWMEWEREIDNYTGYIYQLIEESQNQQEKN | 638 |
| 35 | GCSGKLICTTTVPWNTSWSNKSLEQIWDNMTWMEWEREIDNYTGYIYQLIEESQNQQEKN | 646 |
| 36 | GCSGKLICTTTVPWNTSWSNKSLEQIWDNMTWMEWEREIDNYTGYIYQLIEESQNQQEKN | 646 |
| 37 | GCSGKLICTTTVPWNTSWSNKSLEQIWDNMTWMEWEREIDNYTGYIYQLIEESQNQQEKN | 646 |
| 38 | GCSGKLICTTTVPWNTSWSNKSLEQIWDNMTWMEWEREIDNYTGYIYQLIEESQNQQEKN | 646 |
| 39 | GCSGKLICTTTVPWNTSWSNKSLEQIWDNMTWMEWEREIDNYTGYIYQLIEESQNQQEKN | 645 |
| 40 | GCSGKLICTTTVPWNTSWSNKSLEQIWDNMTWMEWEREIDNYTGYIYQLIEESQNQQEKN | 646 |
| 41 | GCSGKLICPTTVPWNTSWSNKSLEQIWDNMTWMEWEREIDNYTGYIYQLIEESQNQQEKN | 646 |
| 42 | GCSGKLICPTTVPWNTSWSNKSLEQIWDNMTWMEWEREIDNYTGYIYQLIEESQNQQEKN | 646 |
| 43 | GCSGKLICTTTVPWNTSWSNKSLEQIWDNMTWMEWEREIDNYTGYIYQLIEESQNQQEKN | 646 |
| 44 | GCSGKLICTTTVPWNTSWSNKSLEQIWDNMTWMEWEREIDNYTGYIYQLIEESQNQQEKN | 646 |
| 45 | GCSGKLICTTTVPWNTSWSNKSLEQIWDNMTWMEWEREIDNYTGYIYQLIEESQNQQEKN | 646 |
| 46 | GCSGKLICTTTVPWNTSWSNKSLEQIWDNMTWMEWEREIDNYTGYIYQLIEESQNQQEKN | 645 |

| | | |
|---|---|---|
| | EQELLALDKWASLWNWFDITNWLWYIKIFIMIVGGLIGLRIVFTVLSIVNRVRQGYSPLS | |
| 1 | EQELLALDKWASLWNWFDITNWLWYIKIFIMIVGGLIGLRIVFTVLSIVNRVRQGYSPLS | 706 |
| 2 | EQELLALDKWASLWNWFDITNWLWYIKIFIMIVGGLIGLRIVFTVLSIVNRVRQGYSPLS | 702 |
| 3 | GQELLALDKWASLWNWFDITNWLWYIKIFIMIVGGLIGLRIVFTVLSIVNRVRQGYSPLS | 706 |
| 4 | EQELLALDKWASLWNWFDITNWLWYIKIFIMIVGGLIGLRIVFTVLSIVNRVRQGYSPLS | 706 |
| 5 | EQELLALDKWAGLWNWFDITNWLWYIKIFIMIVGGLIGLRIVFTVLSIVNRVRQGYSPLS | 706 |
| 6 | EQELLALDKWAGLWNWFDITNWLWYIKIFIMIVGGLIGLRIVFTVLSIVNRVRQGYSPLS | 706 |
| 7 | EQELLALDKWASLWNWFDITNWLWYIKIFIMIVGGLIGLRIVFTVLSIVNRVRQGYSPLS | 706 |
| 8 | EQELLALDKWASLWNWFDITNWLWYIKIFIMIVGGLIGLRIVFTVLSIVNRVRQGYSPLS | 706 |
| 9 | EQELLALDKWASLWNWFDITNWLWYIKIFIMIVGGLIGLRIVFTVLSIVNRVRQGYSPLS | 701 |
| 10 | EQELLALDKWAGLWNWFDITNWLWYIKIFIMIVGGLIGLRIVFTVLSIVNRVRQGYSPLS | 706 |
| 11 | EQELLALDKWASLWNWFDITNWLWYIKIFIMIVGGLIGLRIVFTVLSIVNRVRQGYSPLS | 706 |
| 12 | EQELLALDKWAGLWNWFDITNWLWYIKIFIMIVGGLIGLRIVFTVLSIVNRVRQGYSPLS | 706 |
| 13 | EQELLALDKWASLWNWFDITNWLWYIKIFIMIVGGLIGLRIVFTVLSIVNRVRQGYSPLS | 706 |
| 14 | EQELLALDKWASLWNWFDITNWLWYIKIFIMIVGGLIGLRIVFTVLSIVNRVRQGYSPLS | 706 |
| 15 | EQELLALDKWASLWNWFDITNWLWYIKIFIMIVGGLIGLRIVFTVLSIVNRVRQGYSPLS | 706 |
| 16 | EQELLALDKWASLWNWFDITNWLWYIKIFIMIVGGLIGLRIVFTVLSIVNRVRQGYSPLS | 706 |
| 17 | EQELLALDKWASLWNWFDITNWLWYIKIFIMIVGGLIGLRIVFTVLSIVNRVRQGYSPLS | 706 |
| 18 | EQELLALDKWASLWNWFDITNWLWYIKIFIMIVGGLIGLRIVFTVLSIVNRVRQGYSPLS | 706 |
| 19 | EQELLALDKWASLWNWFDITNWLWYIKIFIMIVGGLIGLRIVFTVLSIVNRVRQGYSPLS | 705 |
| 20 | EQELLALDKWASLWNWFDITNWLWYIKIFIMIVGGLIGLRIVFTVLSIVNRVRQGYSPLS | 706 |
| 21 | EQELLALDKWASLWNWFDITNWLWYIKIFIMIVGGLIGLRIVFTVLSIVNRVRQGYSPLS | 697 |
| 22 | EQELLALDKWASLWNWFDITNWLWYIKIFIMIVGGLIGLRIVFTVLSIVNRVRQGYSPLS | 706 |
| 23 | EQELLALDKWAGLWNWFDITNWLWYIKIFIMIVGGLIGLRIVFTVLSIVNRVRQGYSPLS | 706 |
| 24 | EQELLALDKWAGLWNWFDITNWLWYIKIFIMIVGGLIGLRIVFTVLSIVNRVRQGYSPLS | 706 |
| 25 | EQELLALDKWASLWNWFDITNWLWYIKIFIMIVGGLIGLRIVFTVLSIVNRVRQGYSPLS | 706 |
| 26 | EQELLALDKWASLWNWFDITNWLWYIKIFIMIVGGLIGLRIVFTVLSIVNRVRQGYSPLS | 700 |
| 27 | EQELLALDKWASLWNWFDITNWLWYIKIFIMIVGGLIGLRIVFTVLSIVNRVRQGYSPLS | 706 |
| 28 | EQELLALDKWASLWNWFDITNWLWYIKIFIMIVGGLIGLRIVFTVLSIVNRVRQGYSPLS | 706 |
| 29 | EQELLALDKWASLWNWFDITNWLWYIKIFIMIVGGLIGLRIVFTVLSIVNRVRQGYSPLS | 706 |
| 30 | EQELLALDKWASLWNWFDITNWLWYIKIFIMIVGGLIGLRIVFTVLSIVNRVRQGYSPLS | 706 |
| 31 | EQELLALDKWASLWNWFDITNWLWYIKIFIMIVGGLIGLRIVFTVLSIVNRVRQGYSPLS | 706 |
| 32 | EQELLALDKWASLWNWFDITNWLWYIKIFIMIVGGLIGLRIVFTVLSIVNRVRQGYSPLS | 706 |
| 33 | EQELLALDKWASLWNWFDITNWLWYIKIFIMIVGGLIGLRIVFTVLSIVNRVRQGYSPLS | 706 |
| 34 | EQELLALDKWASLWNWFDITNWLWYIKIFIMIVGGLIGLRIVFTVLSIVNRVRQGYSPLS | 698 |
| 35 | EQELLALDKWASLWNWFDITNWLWYIKIFIMIVGGLIGLRIVFTVLSIVNRVRQGYSPLS | 706 |
| 36 | EQELLALDKWASLWNWFDITNWLWYIKIFIMIVGGLIGLRIVFTVLSIVNRVRQGYSPLS | 706 |
| 37 | EQELLALDEWASLWNWFDITNWLWYIKIFIMIVGGLIGLRIVFTVLSIVNRVRQGYSPLS | 706 |
| 38 | EQELLALDKWASLWNWFDITNWLWYIKIFIMIVGGLIGLRIVFTVLSIVNRVRQGYSPLS | 706 |
| 39 | EQELLALDKWASLWNWFDITNWLWYIKIFIMIVGGLIGLRIVFTVLSIVNRVRQGYSPLS | 705 |
| 40 | EQELLALDKWASLWNWFDITNWLWYIKIFIMIVGGLIGLRIVFTVLSIVNRVRQGYSPLS | 706 |
| 41 | EQELLALDKWASLWNWFDITNWLWYIKIFIMIVG---GLRIVFTVLSIVNRVRQGYSPLS | 703 |
| 42 | EQELLALDKWASL-NWFDITNWLWYIKIFIMIVGGLIGLRIVFTVLSIVNRVRQGYSPLS | 705 |
| 43 | EQELLALDKWASLWNWFDITNWLWYIKIFIMIVGGLIGLRIVFTVLSIVNRVRQGYSPLS | 706 |
| 44 | EQELLALDKWASLWNWFDITNWLWYIKIFIMIVGGLIGLRIVFTVLSIVNRVRQGYSPLS | 706 |
| 45 | EQELLALDKWAGLWNWFDITNWLWYIKIFIMIVGGLIGLRIVFTVLSIVNRVRQGYSPLS | 706 |
| 46 | EQELLALDKWASLWNWFDITNWLWYIKIFIMIVGGLIGLRIVFTVLSIVNRVRQGYSPLS | 705 |

| | | |
|---|---|---|
| | FQTHLPAQRGPDRPEGIGEEGGERDRDRSDPLVNGFLTLIWSDLRSLCLFSYHRLRDLLL | |
| 1 | FQTHLPAQRGPDRPEGIGEEGGERDRDRSDPLVNGFLALIWSDLRSLCLFSYHRLRDLLL | 766 |
| 2 | FQTHLPAQRGPDRPEGIGEEGGERDRDRSDPLVNGFLTLIWSDLRSLCLFSYHRLRDLLL | 762 |
| 3 | FQTHLPAQRGPDRPEGIGEEGGERDRDRSDPLVNGFLTLIWSDLRSLCLFSYHRLRDLLL | 766 |
| 4 | FQTHLPAQRGPDRPEGIGEEGGERDRDRSDPLVNGFLTLIWSDLRSLCLFSYHRLRDLLL | 766 |
| 5 | FQTHLPAQRGPDRPEGIGEEGGERDRDRSDPLVNGFLTLIWSDLRSLCLFSYHRLRDLLL | 766 |
| 6 | FQTHLPAQRGPDRPEGIGEEGGERDRDRSDPLVNGFLTLIWSDLRSLCLFSYHRLRDLLL | 766 |
| 7 | FQTHLPAQRGPDRPEGIGEEGGERDRDRSDPLVNGFLTLIWSDLRSLCLFSYHRLRDLLL | 766 |
| 8 | FQTHLPAQRGPDRPEGIGEEGGERDRDRSDPLVNGFLTLIWSDLRSLCLFSYHRLRDLLL | 766 |
| 9 | FQTHLPAQRGPDRPEGIGEEGGERDRDRSDPLVNGFLTLIWSDLRSLCLFSYHRLRDLLL | 761 |
| 10 | FQTHLPAQRGPDRPEGIGEEGGERDRDRSDPLVNGFLTLIWSDLRSLCLFSYHRLRDLLL | 766 |
| 11 | FQTHLPAQRGPDRPEGIGEEGGERDRDRSDPLVNGFLTLIWSDLRSLCLFSYHRLRDLLL | 766 |
| 12 | FQTHLPAQRGPDRPEGIGEEGGERDRDRSDPLVNGFLTLIWSDLRSLCLFSYHRLRDLLL | 766 |
| 13 | FQTHLPAQRGPDRPEGIGEEGGERDRDRSDPLVNGFLTLIWSDLRSLCLFSYHRLRDLLL | 766 |
| 14 | FQTHLPAQRGPDRPEGIGEEGGERDRDRSDPLVNGFLTLIWSDLRSLCLFSYHRLRDLLL | 766 |
| 15 | FQTHLPAQRGPDRPEGIGEEGGERDRDRSDPLVNGFLTLIWSDLRSLCLFSYHRLRDLLL | 766 |
| 16 | FQTHLPAQRGPDRPEGIGEEGGERDRDRSDPLVNGFLTLIWSDLRSLCLFSYHRLRDLLL | 766 |
| 17 | FQTHLPAQRGPDRPEGIGEEGGERDRDRSDPLVNGFLTLIWSDLRSLCLFSYHRLRDLLL | 766 |
| 18 | FQTHLPAQRGPDRPEGIGEEGGERDRDRSDPLVNGFLTLIWSDLRSLCLFSYHRLRDLLL | 766 |
| 19 | FQTHLPAQRGPDRPEGIGEEGGERDRDRSDPLVNGFLTLIWSDLRSLCLFSYHRLRDLLL | 765 |
| 20 | FQTHLPAQRGPDRPEGIGEEGGERDRDRSDPLVNGFLTLIWSDLRSLCLFSYHRLRDLLL | 766 |
| 21 | FQTHLPAQRGPDRPEGIGEEGGERDRDRSDPLVNGFLTLIWSDLRSLCLFSYHRLRDLLL | 757 |
| 22 | FQTHLPAQRGPDRPEGIGEEGGERDRDRSDPLVNGFLTLIWSDLRSLCLFSYHRLRDLLL | 766 |
| 23 | FQTHLPAQRGPDRPEGIGEEGGERDRDRSDPLVNGFLTLIWSDLRSLCLFSYHRLRDLLL | 766 |
| 24 | FQTHLPAQRGPDRPEGIGEEGGERDRDRSDPLVNGFLTLIWSDLRSLCLFSYHRLRDLLL | 766 |
| 25 | FQTHLPAQRGPDRPEGIGEEGGERDRDRSDPLVNGFLTLIWSDLRSLCLFSYHRLRDLLL | 766 |
| 26 | FQTHLPAQRGPDRPEGIGEEGGERDRDRSDPLVNGFLTLIWSDLRSLCLFSYHRLRDLLL | 760 |
| 27 | FQTHLPAQRGPDRPEGIGEEGGERDRDRSDPLVNGFLTLIWSDLRSLCLLSYHRLRDLLL | 766 |
| 28 | FQTHLPAQRGPDRPEGIGEEGGERDRDRSDPLVNGFLTLIWSDLRSLCLFSYHRLRDLLL | 766 |
| 29 | FQTHLPAQRGPDRPEGIGEEGGERDRDRSDPLVNGFLTLIWSDLRSLCLFSYHRLRDLLL | 766 |
| 30 | FQTHLPAQRGPDRPEGIGEEGGERDRDRSDPLVNGFLTLIWSDLRSLCLFSYHRLRDLLL | 766 |
| 31 | FQTHLPAQRGPDRPEGIGEEGGERDRDRSDPLVNGFLTLIWSDLRSLCLFSYHRLRDLLL | 766 |
| 32 | FQTHLPAQRGPDRPEGIGEEGGERDRDRSDPLVNGFLTLIWSDLRSLCLFSYHRLRDLLL | 766 |
| 33 | FQTHLPAQRGPDRPEGIGEEGGERDRDRSDPLVNGFLTLIWSDLRSLCLFSYHRLRDLLL | 766 |
| 34 | FQTHLPAQRGPDRPEGIGEEGGERDRDRSDPLVNGFLTLIWSDLRSLCLFSYHRLRDLLL | 758 |
| 35 | FQTHLPAQRGPDRPEGIGEEGGERDRDRSDPLVNGFLTLIWSDLRSLCLFSYHRLRDLLL | 766 |
| 36 | FQTHLPAQRGPDRPEGIGEEGGERDRDRSDPLVNGFLTLIWSDLRSLCLFSYHRLRDLLL | 766 |
| 37 | FQTHLPAQRGPDRPEGIGEEGGERDRDRSDPLVNGFLTLIWSDLRSLCLFSYHRLRDLLL | 766 |
| 38 | FQTHLPAQRGPDRPEGIGEEGGERDRDRSDPLVNGFLTLIWSDLRSLCLFSYHRLRDLLL | 766 |
| 39 | FQTHLPAQRGPDRPEGIGEEGGERDRDRSDPLVNGFLTLIWSDLRSLCLFSYHRLRDLLL | 765 |
| 40 | FQTHLPAQRGPDRPEGIGEEGGERDRDRSDPLVNGFLTLIWSDLRSLCLFSYHRLRDLLL | 766 |
| 41 | FQTHLPAQRGPDRPEGIGEEGGERDRDRSDPLVNGFLTLIWSDLRSLCLFSYHRLRDLLL | 763 |
| 42 | FQTHLPAQRGPDRPEGIGEEGGERDRDRSDPLVNGFLTLIWSDLRSLCLFSYHRLRDLLL | 765 |
| 43 | FQTHLPAQRGPDRPEGIGEEGGERDRDRSDPLVNGFLTLIWSDLRSLCLFSYHRLRDLLL | 766 |
| 44 | FQTHLPAQRGPDRPEGIGEEGGERDRDRSDPLVNGFLTLIWSDLRSLCLFSYHRLRDLLL | 766 |
| 45 | FQTHLPAQRGPDRPEGIGEEGGERDRDRSDPLVNGFLTLIWSDLRSLCLFSYHRLRDLLL | 766 |
| 46 | FQTHLPAQRGPDRPEGIGEEGGERDRDRSDPLVNGFLTLIWSDLRSLCLFSYHRLRDLLL | 765 |

| | | |
|---|---|---|
|  | IVTRIVELLGR-------RGWEVLKYWWNLLQYWSQELKNSAVSLLNATATAVAEGTDRV |  |
| 1 | IVTRIVELLGR-------RGWEVLKYWWNLLQYWSQELKNSAVSLLNATAIAVAEGTDRV | 819 |
| 2 | I----VELLGR-------RGWEVLKYWWNLLQYWSQELKNSAVSLLNATAIAVAEGTDRV | 811 |
| 3 | IVTRIVELLGR-------RGWEVLKYWWNLLQYWSQELKNSAVSLLNATAIAVAEGTDRV | 819 |
| 4 | IVTRIVELLGR-------RGWEVLKYWWNLLQYWSQELKNSAVSLLNATATAVAEGTDRV | 819 |
| 5 | IVTRIVELLGR-------RGWEVLKYWWNLLQYWSQELKNSAVSLLNATATAVAEGTDRV | 819 |
| 6 | IVTRIVELLGR-------RGWEVLKYWWNLLQYWSQELKNSAVSLLNATATAVAEGTDRV | 819 |
| 7 | IVTRIVELLGR-------RGWEVLKYWWNLLQYWSQELKNSAVSLLNATATAVAEGTDRV | 819 |
| 8 | IVTRIVELLGR-------RGWEVLKYWWNLLQYWSQELKNSAVSLLNATATAVAEGTDRV | 819 |
| 9 | IVTRIVELLGR-------RGWEVLKYWWNLLQYWSQELKNSAVSLLNATAIAVAEGTDRV | 814 |
| 10 | IVTRIVELLGR-------RGWEVLKYWWNLLQYWSQELKNSAVSLLNATATAVAEGTDRV | 819 |
| 11 | IVTRIVELLGR-------RGWEVLKYWWNLLQYWSQELKNSAVSLLNATATAVAEGTDRV | 819 |
| 12 | IVTRIVELLGR-------RGWEVLKYWWNLLQYWSQELKNSAVSLLNATATAVAEGTDRV | 819 |
| 13 | IVTRIVELLGR-------RGWEVLKYWWNLLQYWSQELKNSAVSLLNATAIAVAEGTDRV | 819 |
| 14 | IVTRIVELLGR-------RGWEVLKYWWNLLQYWSQELKNSAVSLLNATATAVAEGTDRV | 819 |
| 15 | IVTRIVELLGR-------RGWEVLKYWWNLLQYWSQELKNSAVSLLNATATAVAEGTDRV | 819 |
| 16 | IVTRIVELLGR-------RGWEVLKYWWNLLQYWSQELKNSAVSLLNATATAVAEGTDRV | 819 |
| 17 | IVTRIVELLGR-------RGWEVLKYWWNLLQYWSQELKNSAVSLLNATATAVAEGTDRV | 819 |
| 18 | IVTRIVELLGR-------RGWEVLKYWWNLLQYWSQELKNSAVSLLNATAIAVAEGTDRV | 819 |
| 19 | IVTRIVELLGR-------RGWEVLKYWWNLLQYWSQELKNSAVSLLNATAIAVAEGTDRV | 818 |
| 20 | IVTRIVELLGR-------RGWEVLKYWWNLLQYWSQELKNSAVSLLNATAIAVAEGTDRV | 819 |
| 21 | IVTRIVELLGR-------RGWEVLKYWWNLLQYWSQELKNSAVSLLNATATAVAEGTDRV | 810 |
| 22 | IVTRIVELLGR-------RGWEVLKYWWNLLQYWSQELKNSAVSLLNATATAVAEGTDRV | 819 |
| 23 | IVTRIVELLGR-------RGWEVLKYWWNLLQYWSQELKNSAVSLLNATATAVAEGTDRV | 819 |
| 24 | IVTRIVELLGR-------RGWEVLKYWWNLLQYWSQELKNSAVSLLNATATAVAEGTDRV | 819 |
| 25 | IVTRIVELLGR-------RGWEVLKYWWNLLQYWSQELKNSAVSLLNATATAVAEGTDRV | 819 |
| 26 | IVTRIVELLGR-------RGWEVLKYWWNLLQYWSQELKNSAVSLLNATATAVAEGTDRV | 813 |
| 27 | IVTRIVELLGR-------RGWEVLKYWWNLLQYWSQELKNSAVSLLNATAIAVAEGTDRV | 819 |
| 28 | IVTRIVELLGR-------RGWEVLKYWWNLLQYWSQELKNSAVSLLNATATAVAEGTDRV | 819 |
| 29 | IVTRIVELLGR-------RGWEVLKYWWNLLQYWSQELKNSAVSLLNATAIAVAEGTDRV | 819 |
| 30 | IVTRIVELLGR-------RGWEVLKYWWNLLQYWSQELKNSAVSLLNATAIAVAEGTDRV | 819 |
| 31 | IVTRIVELLGR-------RGWEVLKYWWNLLQYWSQELKNSAVSLLNATAIAVAEGTDRV | 819 |
| 32 | IATRIVELLGR-------RGWEVLKYWWNLLQYWSQELKNSAVSLLNATAIAVAEGTDRV | 819 |
| 33 | IVTRIVELLGR-------RGWEVLKYWWNLLQYWSQELKNSAVSLLNATAIAVAEGTDRV | 819 |
| 34 | IVTRIVELLGR-------RGWEVLKYWWNLLQYWSQELKNSAVSLLNATATAVAEGTDRV | 811 |
| 35 | IVTRIVELLGR-------RGWEVLKYWWNLLQYWSQELKNSAVSLLNATATAVAEGTDRV | 819 |
| 36 | IVTRIVELLGR-------RGWEVLKYWWNLLQYWSQELKNSAVSLLNATATAVAEGTDRV | 819 |
| 37 | IVTRIVELLGR-------RGWEVLKYWWNLLQYWSQELKNSAVSLLNATAIAVAEGTDRV | 819 |
| 38 | IVTRIVELLGR-------RGWEVLKYWWNLLQYWSQELKNSAVSLLNATATAVAEGTDRV | 819 |
| 39 | IVTRIVELLGR-------RGWEVLKYWWNLLQYWSQELKNSAVSLLNATATAVAEGTDRV | 818 |
| 40 | IVTRIVELLGR-------RGWEVLKYWWNLLQYWSQELKNSAVSLLNATATAVAEGTDRV | 819 |
| 41 | I----VELLGR-------RGWEVLKYWWNLLQYWSQELKNSAVSLLNATAIAVAEGTDRV | 812 |
| 42 | IVTRIVELLGR-------RGWEVLKYWWNLLQYWSQELKNSAVSLLNATAIAVAEGTDRV | 818 |
| 43 | IVTRIVELLGR-------RGWEVLKYWWNLLQYWSQELKNSAVSLLNATATAVAEGTDRV | 819 |
| 44 | IVTRIVELLGR-------RGWEVLKYWWNLLQYWSQELKNSAVSLLNATATAVAEGTDRV | 819 |
| 45 | IVTRIVELLGR-------RGWEVLKYWWNLLQYWSQELKNSAVSLLNATATAVAEGTDRV | 819 |
| 46 | IVTRIVELLGR-------RGWEVLKYWWNLLQYWSQELKNSAVSLLNATATAVAEGTDRV | 818 |

**IEVVQRACRAILHIPRRIRQGLERALL**

| | | |
|---|---|---|
| 1 | IEVVQRACRAILHIPRRIRQGLERALL | 846 |
| 2 | IEVVQRACRAILHIPRRIRQGLERALL | 838 |
| 3 | IEVVQRACRAILHIPRRIRQGLERALL | 846 |
| 4 | IEVVQRACRAILHIPRRIRQGLERALL | 846 |
| 5 | IEVVQRACRAILHIPRRIRQGLERALL | 846 |
| 6 | IEVVQRACRAILHIPRRIRQGLERALL | 846 |
| 7 | IEVVQRACRAILHIPRRIRQGLERALL | 846 |
| 8 | IEVVQRACRAILHIPRKIRQGLERALL | 846 |
| 9 | IEVVQRACRAILHIPRRIRQGLERALL | 841 |
| 10 | IEVVQRACRAILHIPRRIRQGLERALL | 846 |
| 11 | IEVVQRACRAILHIPRRIRQGLERALL | 846 |
| 12 | IEVVQRACRAILHIPRRIRQGLERALL | 846 |
| 13 | IEVVQGACRAILHIPRRIRQGLERALL | 846 |
| 14 | IEVVQRACRAILHIPRRIRQGLERALL | 846 |
| 15 | IEVVQRACRAILHIPRRIRQGLERALL | 846 |
| 16 | IEVVQRACRAILHIPRRIRQGLERALL | 846 |
| 17 | IEVVQRACRAILHIPRRIRQGLERALL | 846 |
| 18 | IEVVQRACRAILHIPRRIRQGLERALL | 846 |
| 19 | IEVVQRACRAILHIPRRIRQGLERALL | 845 |
| 20 | IEVVQRACRAILHIPRRIRQGLERALL | 846 |
| 21 | IEVVQRACRAILHIPRRIRQGLERALL | 837 |
| 22 | IEVVQRACRAILHIPRRIRQGLERALL | 846 |
| 23 | IEVVQRACRAILHIPRRIRQGLERALL | 846 |
| 24 | IEVVQRACRAILHIPRRIRQGLERALL | 846 |
| 25 | IEVVQRACRAILHIPRRIRQGLERALL | 846 |
| 26 | IEVVQRACRAILHIPRRIRQGLERALL | 840 |
| 27 | IEVVQRACRAILHIPRRIRQGLERALL | 846 |
| 28 | IEVVQRACRAILHIPRRIRQGLERALL | 846 |
| 29 | IEVVQRACRAILHIPRRIRQGLERALL | 846 |
| 30 | IEVVQRACRAILHIPRRIRQGLERALL | 846 |
| 31 | IEVVQRACRAILHIPRRIRQGLERALL | 846 |
| 32 | IEVVQRACRAILHIPRRIRQGLERALL | 846 |
| 33 | IEVVQRACRAILHIPRRIRQGLERALL | 846 |
| 34 | IEVVQRACRAILHIPRRIRQGLERALL | 838 |
| 35 | IEVVQRACRAILHIPRRIRQGLERALL | 846 |
| 36 | IEVVQRACRAILHIPRRIRQGLERALL | 846 |
| 37 | IEVVQRACRAIRHIPRRIRQGLERALL | 846 |
| 38 | IEVVQRACRAILHIPRRIRQGLERALL | 846 |
| 39 | IEVVQRACRAILHIPRRIRQGLERALL | 845 |
| 40 | IEVVQRACRAILHIPRRIRQGLERALL | 846 |
| 41 | IEVVQRACRAILHIPRRIRQGLERALL | 839 |
| 42 | IEVVQRACRAILHIPRRIRQGLERALL | 845 |
| 43 | IEVVQRACRAILHIPRRIRQGLERALL | 846 |
| 44 | IEVVQRACRAILHIPRRIRQGLERALL | 846 |
| 45 | IEVVQRACRAILHIPRRIRQGLERALL | 846 |
| 46 | IEVVQRACRAILHIPRRIRQGLERALL | 845 |

#### Aligned using an external algorithm

**Consensus**

1. B.US.2006.700010040_C9_4520.EU289193
2. B.US.2008.CH0040_3_d0743_ipe032_2_03.MG900387
3. B.US.2008.CH0040_3_d0743_ipe032_2_04.MG900388
4. B.US.2008.CH0040_3_d0743_ipe032_2_05.MG900389
5. B.US.2008.CH0040_3_d0743_ipe032_2_07.MG900390
6. B.US.2008.CH0040_3_d0743_ipe032_2_08.MG900391
7. B.US.2008.CH0040_3_d0743_ipe032_2_10.MG900392
8. B.US.2008.CH0040_3_d0743_ipe032_2_11.MG900393
9. B.US.2008.CH0040_3_d0743_ipe032_2_12.MG900394
10. B.US.2008.CH0040_3_d0743_ipe032_2_13.MG900395
11. B.US.2008.CH0040_3_d0743_ipe032_2_14.MG900396
12. B.US.2008.CH0040_3_d0743_ipe032_2_15.MG900397
13. B.US.2008.CH0040_3_d0743_ipe032_2_16.MG900398
14. B.US.2008.CH0040_3_d0743_ipe032_2_18.MG900399
15. B.US.2008.CH0040_3_d0743_ipe032_2_19.MG900400
16. B.US.2008.CH0040_3_d0743_ipe032_2_20.MG900401
17. B.US.2008.CH0040_3_d0743_ipe032_2_22.MG900402
18. B.US.2008.CH0040_3_d0743_ipe032_2_25.MG900403
19. B.US.2008.CH0040_3_d0743_ipe032_2_26.MG900404
20. B.US.2008.CH0040_3_d0743_ipe032_2_27.MG900405
21. B.US.2008.CH0040_3_d0743_ipe032_2_28.MG900406
22. B.US.2008.CH0040_3_d0743_ipe032_2_29.MG900407
23. B.US.2008.CH0040_3_d0743_ipe032_2_30.MG900408
24. B.US.2008.CH0040_3_d0743_ipe032_2_32.MG900409
25. B.US.2008.CH0040_3_d0743_ipe032_2_33.MG900410
26. B.US.2008.CH0040_3_d0743_ipe032_3_24.MG900411

| | | |
|---|---|---|
| | **MRVMGIRKNYQHLWREGILLLGILMICSAADNLWVTVYYGVPVWREATTTLFCASDAKAY** | |
| 1 | MRVMGIRKNYQHLWREGILLLGILMICSAADNLWVTVYYGVPVWREATTTLFCASDAKAY | 60 |
| 2 | MRVMGIRKNYQHLWREGILLLGILMICSAADNLWVTVYYGVPVWREATTTLFCASDAKAY | 60 |
| 3 | MRVMGIRKNYQHLWREGILLLGILMICSAADNLWVTVYYGVPVWREATTTLFCASDAKAY | 60 |
| 4 | MRVMGIRKNYQHLWREGILLLGILMICSAADNLWVTVYYGVPVWREATTTLFCASDAKAY | 60 |
| 5 | MRVMGIRKNYQHLWREGILLLGILMICSAADNLWVTVYYGVPVWREATTTLFCASDAKAY | 60 |
| 6 | MRVMGIRKNYQHLWREGILLLGILMICSAADNLWVTVYYGVPVWREATTTLFCASDAKAY | 60 |
| 7 | MRVMGIRKNYQHLWREGILLLGILMICSAADNLWVTVYYGVPVWREATTTLFCASDAKAY | 60 |
| 8 | MRVMGIRKNYQHLWREGILLLGILMICSAADNLWVTVYYGVPVWREATTTLFCASDAKAY | 60 |
| 9 | MRVMGIRKNYQHLWREGILLLGILMICSAADNLWVTVYYGVPVWREATTTLFCASDAKAY | 60 |
| 10 | MRVMGIRKNYQHLWREGILLLGILMICSAADNLWVTVYYGVPVWREATTTLFCASDAKAY | 60 |
| 11 | MRVMGIRKNYQHLWREGILLLGILMICSAADNLWVTVYYGVPVWREATTTLFCASDAKAY | 60 |
| 12 | MRVMGIRKNYQHLWRKGILLLGILMICSAADNLWVTVYYGVPVWREATTTLFCASDAKAY | 60 |
| 13 | MRVMGIRKNYQHLWREGILLLGILMICSAADNLWVTVYYGVPVWREATTTLFCASDAKAY | 60 |
| 14 | MRVMGIRKNYQHLWREGILLLGILMICSAADNLWVTVYYGVPVWREATTTLFCASDAKAY | 60 |
| 15 | MRVMGIRKNYQHLWREGILLLGILMICSAADNLWVTVYYGVPVWREATTTLFCASDAKAY | 60 |
| 16 | MRVMGIRKNYQHLWREGILLLGILMICSAADNLWVTVYYGVPVWREATTTLFCASDAKAY | 60 |
| 17 | MRVMGIRKNYQHLWREGILLLGILMICSAADNLWVTVYYGVPVWREATTTLFCASDAKAY | 60 |
| 18 | MRVMGIRKNYQHLWREGILLLGILMICSAADNLWVTVYYGVPVWREATTTLFCASDAKAY | 60 |
| 19 | MRVMGIRKNYQHLWREGILLLGILMICSAADNLWVTVYYGVPVWREATTTLFCASDAKAY | 60 |
| 20 | MRVMGIRKNYQHLWRKGILLLGILMICSAADNLWVTVYYGVPVWREATTTLFCASDAKAY | 60 |
| 21 | MRVMGIRKNYQHLWREGILLLGILMICSAADNLWVTVYYGVPVWREATTTLFCASDAKAY | 60 |
| 22 | MRVMGIRKNYQHLWREGILLLGILMICSAADNLWVTVYYGVPVWREATTTLFCASDAKAY | 60 |
| 23 | MRVMGIRKNYQHLWREGILLLGILMICSAADNLWVTVYYGVPVWREATTTLFCASDAKAY | 60 |
| 24 | MRVMGIRKNYQHLWREGILLLGILMICSAADNLWVTVYYGVPVWREATTTLFCASDAKAY | 60 |
| 25 | MRVMGIRKNYQHLWREGILLLGILMICSAADNLWVTVYYGVPVWREATTTLFCASDAKAY | 60 |
| 26 | MRVMGIRKNYQHLWREGILLLEILMICSAADNLWVTVYYGVPVWREATTTLFCASDAKAY | 60 |

**DTEAHNVWATHACVPTDPNPQEVELKNVTENFNMWENNMVEQMHEDIISLWDQSLKPCVK**

| | | |
|---|---|---|
| 1 | DTEAHNVWATHACVPTDPNPQEVELKNVTENFNMWENNMVEQMHEDIISLWDQSLKPCVK | 120 |
| 2 | DTEAHNVWATHACVPTDPNPQEVELKNVTENFNMWENNMVEQMHEDIISLWDQSLKPCVK | 120 |
| 3 | DTEAHNVWATHACVPTDPNPQEVELKNVTENFNMWENNMVEQMHEDIISLWDQSLKPCVK | 120 |
| 4 | DTEAHNVWATHACVPTDPNPQEVELKNVTENFNMWENNMVEQMHEDIISLWDQSLKPCVK | 120 |
| 5 | DTEAHNVWATHACVPTDPNPQEVELKNVTENFNMWENNMVEQMHEDIISLWDQSLKPCVK | 120 |
| 6 | DTEAHNVWATHACVPTDPNPQEVELKNVTENFNMWENNMVEQMHEDIISLWDQSLKPCVK | 120 |
| 7 | GTEAHNVWATHACVPTDPNPQEVELKNVTENFNMWENNMVEQMHEDIISLWDQSLKPCVK | 120 |
| 8 | DTEAHNVWATHACVPTDPNPQEVELKNVTENFNMWKNNMVEQMHEDIISLWDQSLKPCVK | 120 |
| 9 | DTEAHNVWATHACVPTDPNPQEVELKNVTENFNMWENNMVEQMHEDIISLWDQSLKPCVK | 120 |
| 10 | DTEAHNVWATHACVPTDPNPQEVELKNVTENFNMWENNMVEQMHEDIISLWDQSLKPCVK | 120 |
| 11 | DTEAHNVWATHACVPTDPNPQEVELKNVTENFNMWENNMVEQMHEDIISLWDQSLKPCVK | 120 |
| 12 | DTEAHNVWATHACVPTDPNPQEVELKNVTENFNMWENNMVEQMHEDIISLWDQSLKPCVK | 120 |
| 13 | DTEAHNVWATHACVPTDPNPQEVELKNVTENFNMWENNMVEQMHEDIISLWDQSLKPCVK | 120 |
| 14 | DTEAHNVWATHACVPTDPNPQEVELKNVTENFNMWENNMVEQMHEDIISLWDQSLKPCVK | 120 |
| 15 | DTEAHNVWATHACVPTDPNPQEVELKNVTENFNMWENNMVEQMHEDIISLWDQSLKPCVK | 120 |
| 16 | DTEAHNVWATHACVPTDPNPQEVELKNVTENFNMWENNMVEQMHEDIISLWDQSLKPCVK | 120 |
| 17 | DTEAHNVWATHACVPTDPNPQEVELKNVTENFNMWENNMVEQMHEDIISLWDQSLKPCVK | 120 |
| 18 | DTEAHNVWATHACVPTDPNPQEVELKNVTENFNMWENNMVEQMHEDIISLWDQSLKPCVK | 120 |
| 19 | DTEAHNVWATHACVPTDPNPQEVELKNVTENFNMWENNMVEQMHEDIISLWDQSLKPCVK | 120 |
| 20 | DTEAHNVWATHACVPTDPNPQEVELKNVTENFNMWENNMVEQMHEDIISLWDQSLKPCVK | 120 |
| 21 | DTEAHNVWATHACVPTDPNPQEVELKNVTENFNMWENNMVEQMHEDIISLWDQSLKPCVK | 120 |
| 22 | DTEAHNVWATHACVPTDPNPQEVELKNVTENFNMWENNMVEQMHEDIISLWDQSLKPCVK | 120 |
| 23 | DTEAHNVWATHACVPTDPNPQEVELKNVTENFNMWENNMVEQMHEDIISLWDQSLKPCVK | 120 |
| 24 | DTEAHNVWATHACVPTDPNPQEVELKNVTENFNMWKNNMVEQMHEDIISLWDQSLKPCVK | 120 |
| 25 | DTEAHNVWATHACVPTDPNPQEVELKNVTENFNMWENNMVEQMHEDIISLWDQSLKPCVK | 120 |
| 26 | DTEAHNVWATHACVPTDPNPQEVELKNVTENFNMWENNMVEQMHEDIISLWDQSLKPCVK | 120 |

```
    LTPLCVTLNCTDLGNVTTTT-----------------NSNGTLMEKGEVKNCSFKITTDI

1   LTPLCVTLNCTDLGNVTNTT-----------------NSNGEMMEKGEVKNCSFKITTDI   163
2   LTPLCVTLNCTDLGNVTTTTNRNGTNCTDLGNVTTTTNRNGTLMEKGEVKNCSFRITTDI   180
3   LTPLCVTLNCTDLGNVTTTT-----------------NNNGTLMEKGEVKNCSFKITTDT   163
4   LTPLCVTLNCTDLGNVTTTTNRNGTNCTDLGNVTTTTNRNGTLMEKGEVKNCSFRITTDI   180
5   LTPLCVTLNCTDLGNVTTTT-----------------NSNGTLMEKGEVKNCSFKITTDT   163
6   LTPLCVTLNCTDLGNVTTTT-----------------NSNGTLMEKGEVKNCSFKITTDT   163
7   LTPLCVTLNCTDLGNVTTTTNRNGTNCTDLGNVTTTTNRNGTLMEKGEVKNCSFRITTDI   180
8   LTPLCVTLNCTDLGNVTTTT-----------------NSNGTLMEKGEVKNCSFKITTEI   163
9   LTPLCVTLNCTDLGNGTNTT-----------------NSNEKMMEKGEIKNCSFKITTDI   163
10  LTPLCVTLNCTDLGNDTTTT-----------------NSNGTLMEKGEVKNCSFKITTEI   163
11  LTPLCVTLNCTDLGNVTTTTNRNGTNCTDLGNVTTTTNRNGTLMEKGEVKNCSFRITTDI   180
12  LTPLCVTLNCTDLEKGTNTT-----------------NSTEKMMEKGEVKNCSFRITTDI   163
13  LTPLCVTLNCTDLGNVTTTTNRNGTNCTDLGNVTTTTNRNGTLMEKGEVKNCSFRITTDI   180
14  LTPLCVTLNCTDLGNVTTTT-----------------NSNGTLMEKGEVKNCSFKITTDT   163
15  LTPLCVTLNCTDLGNVTTTTNRNGTNCTDLGNVTTTTNRNGTLMEKGEVKNCSFKITTDV   180
16  LTPLCVTLNCTDLGNDTTTT-----------------NSNGTLMEKGEVKNCSFKITTEI   163
17  LTPLCVTLNCTDLGNDTTTT-----------------NSNGTLMEKGEVKNCSFKITTEI   163
18  LTPLCVTLNCTDLGNVTTTTNRNGTNCTDLGNVTTTTNRNGTLMEKGEVKNCSFRITTDI   180
19  LTPLCVTLNCTDLGNVTTTTNRNGTNCTDLGNVTTTTNRNGTLMEKGEVKNCSFKITTDV   180
20  LTPLCVTLNCTDLGNVTTTT-----------------NSNGTLMEKGEVKNCSFKITTDI   163
21  LTPLCVTLNCTDLGNDTTTT-----------------NSNGTLMEKGEVKNCSFKITTEI   163
22  LTPLCVTLNCTDLGNVTTTT-----------------NSNGTLMEKGEVKNCSFKITTDT   163
23  LTPLCVTLNCTDLGNDTTTT-----------------NSNGTLMEKGEVKNCSFKITTEI   163
24  LTPLCVTLNCTDLGNVTTTT-----------------NSNGTLMEKGEVKNCSFKITTEI   163
25  LTPLCVTLNCTDLGNVTTTTNRNGTNCTDLGNVTTTTNRNGTLMEKGEVKNCSFRITTDI   180
26  LTPLCVTLNCTDLGNVTNTT-----------------NSNGGMMEKGEVKNCSFKITADI   163
```

| | | |
|---|---|---|
|  | **KDRTRKEYALFYKLDVVPIND-------TRYRLVSCNTSVITQACPKVSFEPIPIHYCAP** |  |
| 1 | KDRTRKEYALFYKLDVVPIND-------TRYRLVSCNTSVITQACPKVSFEPIPIHYCAP | 216 |
| 2 | KDRTRKEYALFYKLDVVPIND-------TRYRLVSCNTSVITQACPKVSFEPIPIHYCAP | 233 |
| 3 | KDRTRKEYALFYKLDVVPIND-------TRYRLVSCNTSVITQACPKVSFEPIPIHYCAP | 216 |
| 4 | KDRTRKEYALFYKLDVVPIND-------TRYRLVSCNTSVITQACPKVSFEPIPIHYCAP | 233 |
| 5 | KDRTRKEYALFYKLDVVPIND-------TRYRLVSCNTSVITQACPKVSFEPIPIHYCAP | 216 |
| 6 | KDRTRKEYALFYKLDVVPINH-------TRYRLVSCNTSVITQACPKVSFEPIPIHYCAP | 216 |
| 7 | KDRTRKEYALFYKLDVVPIND-------TRYRLVSCNTSVITQACPKVSFEPIPIHYCAP | 233 |
| 8 | KDRTRKEYALFYKLDVVPIND-------TRYRLVSCNTSVITQACPKVSFEPIPIHYCAP | 216 |
| 9 | RDRTRKEYALFYKLDVVPIND-------TRYRLVSCNTSVITQACPKVSFEPIPIHYCAP | 216 |
| 10 | TDRTRKEYALFYKLDVVPIND-------TRYRLVSCNTSVITQACPKVSFEPIPIHYCAP | 216 |
| 11 | KDRTRKEYALFYKLDVVPIND-------TRYRLVSCNTSVITQACPKVSFEPIPIHYCAP | 233 |
| 12 | KDRTRKEYALFYKLDVVPIND-------TRYRLVSCNTSVITQACPKVSFEPIPIHYCAP | 216 |
| 13 | KDRTRKEYALFYKLDVVPIND-------TRYRLVSCNTSVITQACPKVSFEPIPIHYCAP | 233 |
| 14 | KDRTRKEYALFYKLDVVPIND-------TRYRLVSCNTSVITQACPKVSFEPIPIHYCAP | 216 |
| 15 | KDRMRKEYALFYKLDVVPIND-------TRYRLVSCNTSVITQACPKVSFEPIPIHYCAP | 233 |
| 16 | TDRTRKEYALFYKLDVVPIND-------TRYRLVSCNTSVITQACPKVSFEPIPIHYCAP | 216 |
| 17 | KDRTRKEYALFYNLDVVPIND-------TRYRLVSCNTSVITQACPKVSFEPIPIHYCAP | 216 |
| 18 | KDRTRKEYALFYKLDVVPIND-------TRYRLVSCNTSVITQACPKVSFEPIPIHYCAP | 233 |
| 19 | KDRMRKEYALFYKLDVVPIND-------TRYRLVSCNTSVITQACPKVSFEPIPIHYCAP | 233 |
| 20 | KDRTRTEYALFYKLDVVPIND-------TRYRLVSCNTSVITQACPKVSFEPIPIHYCAP | 216 |
| 21 | KDRTRKEYALFYKLDVVPIND-------TRYRLVSCNTSVITQACPKVSFEPIPIHYCAP | 216 |
| 22 | KDRTRKEYALFYKLDVVPIND-------TRYRLVSCNTSVITQACPKVSFEPIPIHYCAP | 216 |
| 23 | KDRTRKEYALFYKLDVVPIND-------TRYRLVSCNTSVITQACPKVSFEPIPIHYCAP | 216 |
| 24 | KDRTRKEYALFYKLDVVPIND-------TRYRLVSCNTSVITQACPKVSFEPIPIHYCAP | 216 |
| 25 | KDRTRKEYALFYKLDVVPIND-------TRYRLVSCNTSVITQACPKVSFEPIPIHYCAP | 233 |
| 26 | KDRTRKEYALFYKLDVVPIND-------TRYRLVSCNTSVITQACPKVSFEPIPIHYCAP | 216 |

| | | |
|---|---|---|
| | **AGFAILKCNDKQFIGTGPCTNVSTVQCTHGIRPVVSTQLLLNGSLAEEEVVIRSVNFSDN** | |
| 1 | AGFAILKCNDKQFIGTGPCTNVSTVQCTHGIRPVVSTQLLLNGSLAEEEVVIRSVNFSDN | 276 |
| 2 | AGFAILKCNDKQFNGTGPCTNVSTVQCTHGIRPVVSTQLLLNGSLAEEEVVIRSVNFSDN | 293 |
| 3 | AGFAILKCNDKQFIGTGPCTNVSTVQCTHGIRPVVSTQLLLNGSLAEEEVVIRSVNFSDN | 276 |
| 4 | AGFAILKCNDKQFIGTGPCTNVSTVQCTHGIRPVVSTQLLLNGSLAEEEVVIRSVNFSDN | 293 |
| 5 | AGFAILKCNDKQFIGTGPCTNVSTVQCTHGIRPVVSTQLLLNGSLAEEEVVIRSVNFSDN | 276 |
| 6 | AGFAILKCNDKQFIGTGPCTNVSTVQCTHGIRPVVSTQLLLNGSLAEEEVVIRSVNFSDN | 276 |
| 7 | AGFAILKCNDKQFIGTGPCTNVSTVQCTHGIRPVVSTQLLLNGSLAEEEVVIRSVNFSDN | 293 |
| 8 | AGFAILKCNDKQFNGTGPCTNVSTVQCTHGIRPVVSTQLLLNGSLAEEEVVIRSVNFRDN | 276 |
| 9 | AGFAILKCNDKQFNGTGPCTNVSTVQCTHGIRPVVSTQLLLNGSLAEEEVVIRSVNFSDN | 276 |
| 10 | AGFAILKCNDKQFNGTGPCTNVSTVQCTHGIRPVVSTQLLLNGSLAEEEVVIRSVNFSDN | 276 |
| 11 | AGFAILKCNDKQFNGTGPCTNVSTVQCTHGIRPVVSTQLLLNGSLAEEEVVIRSVNFSDN | 293 |
| 12 | AGFAILKCNDKQFIGTGPCTNVSTVQCTHGIRPVVSTQLLLNGSLAEEEVVIRSVNFSDN | 276 |
| 13 | AGFAILKCNDKQFNGTGPCTNVSTVQCTHGIRPVVSTQLLLNGSLAEEEVVIRSVNFSDN | 293 |
| 14 | AGFAILKCNDKQFIGTGPCTNVSTVQCTHGIRPVVSTQLLLNGSLAEEEVVIRSVNFSDN | 276 |
| 15 | AGFAILKCNDKQFNGTGPCTNVSTVQCTHGIRPVVSTQLLLNGSLAEEEVVIRSVNFSDN | 293 |
| 16 | AGFAILKCNDKQFNGTGPCTNVSTVQCTHGIRPVVSTQLLLNGSLAEEEVVIRSVNFSDN | 276 |
| 17 | AGFAILKCNDKQFNGTGPCTNVSTVQCTHGIRPVVSTQLLLNGSLAEEEVVIRSVNFSDN | 276 |
| 18 | AGFAILKCNDKQFIGTGPCTNVSTVQCTHGIRPVVSTQLLLNGSLAEEEVVIRSVNFSDN | 293 |
| 19 | AGFAILKCNDKQFNGTGPCTNVSTVQCTHGIRPVVSTQLLLNGSLAEEEVVIRSVNFSDN | 293 |
| 20 | AGFAILKCNDKQFIGTGPCTNVSTVQCTHGIRPVVSTQLLLNGSLAEEEVVIRSVNFSDN | 276 |
| 21 | AGFAILKCNDKQFNGTGPCTNVSTVQCTHGIRPVVSTQLLLNGSLAEEEVVIRSVNFSDN | 276 |
| 22 | AGFAILKCNDKQFIGTGPCTNVSTVQCTHGIRPVVSTQLLLNGSLAEEEVVIRSVNFSDN | 276 |
| 23 | AGFAILKCNDKQFNGTGPCTNVSTVQCTHGIRPVVSTQLLLNGSLAEEEVVIRSVNFSDN | 276 |
| 24 | AGFAILKCNDKQFIGTGPCTNVSTVQCTHGIRPVVSTQLLLNGSLAEEEVVIRSVNFNDN | 276 |
| 25 | AGFAILKCNDKQFIGTGPCTNVSTVQCTHGIRPVVSTQLLLNGSLAEEEVVIRSVNFSDN | 293 |
| 26 | AGFAILKCNDKQFIGTGPCTNVSTVQCTHGIRPVVSTQLLLNGSLAEEEVVIRSVNFSDN | 276 |

| | | |
|---|---|---|
| | AKTIIVQLNKSVEINCTRPXNNTRKSIPMGPGKAFYARGDIIGDIRKAYCEINGTEWHST | |
| 1 | AKTIIVQLNKSVEITCTRPNNNTRKSIPMGPGKAFYARGDITGDIRKAYCEINGTEWHST | 336 |
| 2 | AKTIIVQLNKSVEINCTRPNNNTRKSIPMGPGKAFYARGDIIGDIRKASCEINGTEWHST | 353 |
| 3 | AKTIIVQLNKSVEINCTRPNNNTRKSIPMGPGKAFYARGDIIGNIRKASCELDGTEWHST | 336 |
| 4 | AKTIIVQLNKSVEINCTRPHNNTRKSIPMGPGKAFYARGDIIGDIRKAYCEINGTEWHST | 353 |
| 5 | AKTIIVQLNKSVEINCTRPNNNTRKSIPMGPGKAFYARGDIIGDIRKASCELDGTEWHST | 336 |
| 6 | AKTIIVQLNKSVEINCTRPNNNTRKSIPMGPGKAFYARGDIIGNIRKASCELDGTEWHST | 336 |
| 7 | AKTIIVQLNKSVEINCTRPHNNTRKSIPMGPGKAFYARGDIIGDIRKAYCEINGTEWHST | 353 |
| 8 | AKTIIVQLNKSVEINCTRPHNNTRKSIPMGPGKAFYARGDIIGNIRKAYCEINGTEWHST | 336 |
| 9 | AKTIIVQLNKSVEINCTRPHNNTRKSIPMGPGKAFYARGDIIGDIRKAYCNINGTEWHST | 336 |
| 10 | AKTIIVQLNKSVEINCTRPHNNTRKSIPMGPGKAFYARGDIIGNIRKAYCTINGTEWHST | 336 |
| 11 | AKTIIVQLNKSVEITCTRPHNNTRKSIPMGPGKAFYAKGDIIGDIRKAYCEINGTEWHST | 353 |
| 12 | AKTIIVQLNKSVEINCTRPHNNTRKSIPMGPGKAFYARGDIIGDIRKAYCEINGTEWHST | 336 |
| 13 | AKTIIVQLNKSVEINCTRPNNNTRKSIPMGPGKAFYARGDIIGDIRKASCEINGTEWHST | 353 |
| 14 | AKTIIVQLNKSVEINCTRPNNNTRKSIPMGPGKAFYARGDIIGDIRKASCELDGTEWHST | 336 |
| 15 | AKTIIVQLNKSVEMTCTRPNNNTRKSIPMGPGKAFYARGDIIGDIRKASCEINGTEWHST | 353 |
| 16 | AKTIIVQLNKSVEINCTRPHNNTRKSIPMGPGKAFYARGDIIGNIRKASCEINGTEWHST | 336 |
| 17 | AKTIIVQLNKSVEINCTRPHNNTRKSIPMGPGKAFYARGDIIGDIRKAYCTINGTEWHSA | 336 |
| 18 | AKTIIVQLNKSVEITCTRPNNNTRKSIPMGPGKAFYARGDIIGDIRKASCEINGTEWHST | 353 |
| 19 | AKTIIVQLNKSVEMTCTRPNNNTRKSIPMGPGKAFYARGDIIGDIRKASCEINGTEWHST | 353 |
| 20 | AKTIIVQLNKSVEINCTRPHNNTRKSIPMGPGKAFYAKGDIIGDIRKAYCEINGTEWHST | 336 |
| 21 | AKTIIVQLNKSVEINCTRPHNNTRKSIPMGPGKAFYARGDIIGDIRKAYCTINGTEWHST | 336 |
| 22 | AKTIIVQLNKSVEINCTRPNNNTRKSIPMGPGKAFYARGDIIGDIRKASCELDGTEWHST | 336 |
| 23 | AKTIIVQLNKSVEINCTRPHNNTRKSIPMGPGKAFYARGDIIGDIRKAYCTINGTEWHST | 336 |
| 24 | AKTIIVQLNKSVEINCTRPHNNTRKSIPMGPGKAFYARGDIIGNIRKAYCEINGTEWHST | 336 |
| 25 | AKTIIVQLNKSVEITCTRPNNNTRKSIPMGPGKAFYARGDIIGDIRKASCEINGTEWHST | 353 |
| 26 | AKTIIVQLNKSVEITCTRPNNNTRKSIPMGPGKAFYARGDITGDIKKAYCKINGTEWHST | 336 |

| | Sequence | |
|---|---|---|
| | **LKLVVEKLREQY-NKTIVFNRSSGGDPEIVMYSFNCGGEFFYCNSTKLFNSTWPWND-TK** | |
| 1 | LKLVVEKLREQY-NKTIVFNRSSGGDPEIVMYSFNCGGEFFYCNSTKLFNSTWPWND-TK | 394 |
| 2 | LKLVVEKLREQY-NKTIVFNRSSGGDPEIVMYSFNCGGEFFYCNSTKLFNSTWPWND-TK | 411 |
| 3 | LKLVVEQLRKQY-NKTIVFNRSSGGDPEIVMYSFNCGGEFFYCNSTKLFNSTWPWND-TK | 394 |
| 4 | LKLVVGKLREKY-NKTIVFNRSSGGDPEIVMYSFNCGGEFFYCNSTKLFNSTWPWND-NK | 411 |
| 5 | LKLVVEQLRKQY-NKTIVFNRSSGGDPEIVMYSFNCGGEFFYCNSTKLFNSTWPWND-TK | 394 |
| 6 | LKLVVEKLRKQY-NKTIVFNRSSGGDPEIVMYSFNCGGEFFYCNSTKLFNSTWPWND-TK | 394 |
| 7 | LKLVVGKLREKY-NKTIVFNRSSGGDPEIVMYSFNCGGEFFYCNSTKLFNSTWPWND-TK | 411 |
| 8 | LKLVVEKLREKY-NKTIVFNRSSGGDPEIVMYSFNCGGEFFYCNSTKLFNSTWPWND-TK | 394 |
| 9 | LKLVVEKLREQY-NKTIVFNRSSGGDPEIVMYSFNCGGEFFYCNSTKLFNSTWPWND-TK | 394 |
| 10 | LKLVVEKLREQY-NKTIVFNRSSGGDPEIVMYSFNCGGEFFYCNSTKLFNSTWPWND-TK | 394 |
| 11 | LKLVVEKLRERY-NKTIVFNRSSGGDPEIVMYSFNCGGEFFYCNSTKLFNSTWPWND-TK | 411 |
| 12 | LKLVVGKLREKY-NKTIVFNRSSGGDPEIVMYSFNCGGEFFYCNSTKLFNSTWPWND-TK | 394 |
| 13 | LKLVVEKLREQY-NKTIVFNRSSGGDPEIVMYSFNCGGEFFYCNSTKLFNSTWPWND-TK | 411 |
| 14 | LKLVVEKLRKQY-NKTIVFNRSSGGDPEIVMYSFNCGGEFFYCNSTKLFNSTWPWND-TK | 394 |
| 15 | LKMVVEKLREQY-NKTIVFNRSSGGDPEIVMYSFNCGGEFFYCNSTKLFNSTWPWND-TK | 411 |
| 16 | LKLVVEKLREQY-NKTIVFNRSSGGDPEIVMYSFNCGGEFFYCNSTKLFNSTWPWND-TK | 394 |
| 17 | LKLVVEKLREQY-NKTIVFNRSSGGDPEIVMYSFNCGGEFFYCNSTKLFNSTWPWND-TK | 394 |
| 18 | LKLVVEKLREQY-NKTIVFNRSSGGDPEIVMYSFNCGGEFFYCNSTKLFNSTWPWND-TK | 411 |
| 19 | LKMVVEKLREQY-NKTIVFNRSSGGDPEIVMYSFNCGGEFFYCNSTKLFNSTWPWND-TK | 411 |
| 20 | LKLVVGKLRERY-NKTIVFNRSSGGDPEIVMYSFNCGGEFFYCNSTKLFNSTWPWND-TK | 394 |
| 21 | LKLVVEKLREQY-NKTIVFNRSSGGDPEIVMYSFNCGGEFFYCNSTKLFNSTWPWND-TK | 394 |
| 22 | LKLVVEKLRKQY-NKTIVFNRSSGGDPEIVMYSFNCGGEFFYCNSTKLFNSTWPWND-TK | 394 |
| 23 | LKLVVEKLREQY-NKTIVFNRSSGGDPEIVMYSFNCGGEFFYCNSTKLFNSTWPWND-TK | 394 |
| 24 | LKLVVEKLREKY-NKTIVFNRSSGGDPEIVMYSFNCGGEFFYCNSTKLFNSTWPWND-TK | 394 |
| 25 | LKLVVEKLREQY-NKTIVFNRSSGGDPEIVMYSFNCGGEFFYCNSTKLFNSTWPWND-TK | 411 |
| 26 | LKLVVEKLREQY-NKTIVFNRSSGGDPEIVMYSFNCGGEFFYCNSTKLFNSTWPWND-TK | 394 |

| | Sequence | |
|---|---|---|
| | GSHDTNGTLILPCRIKQIINMWQGVGKAMYAPPIEGXIRCSSNITGLLLTRDGG-YESNE | |
| 1 | GSHDTNGTLILPCKIKQIINMWQGVGKAMYAPPIEGKIRCSSNITGLLLTRDGG-YESNE | 453 |
| 2 | GSHDTNGKLILPCRIKQIINMWQGVGKAMYAPPIEGIIRCSSNITGLLLTRDGG-YESNE | 470 |
| 3 | GSHDTNGTLILPCRIKQIINMWQGVGKAMYAPPIEGEIRCSSNITGLLLTRDGG-YESNE | 453 |
| 4 | GSHDTNGKLILPCRIKQIINMWQGVGKAMYAPPIEGIIRCSSNITGLLLTRDGG-YESNE | 470 |
| 5 | GSHDTNGTLILPCRIKQIINMWQGVGKAMYAPPIEGEIRCSSSITGLLLTRDGG-YESNE | 453 |
| 6 | GSHDTNGTLILPCRIKQIINMWQGVGKAMYAPPIEGEIRCSSNITGLLLTRDGG-YESNE | 453 |
| 7 | GSHDTNGKLILPCRIKQIINMWQGVGKAMYAPPIEGIIRCSSNITGLLLTRDGG-YESNE | 470 |
| 8 | GSHDTNGTLILPCRIKQIINMWQGVGKAMYAPPIEGEIRCSSNITGLLLTRDGG-YESNE | 453 |
| 9 | GSHDTNDTLMLPCKIKQIINMWQGVGKAMYAPPIEGQISCSSNITGLLLTRDGG-YESNK | 453 |
| 10 | GSHDTNDTLMLPCKIKQIINMWQGVGKAMYAPPIEGQIRCSSNITGLLLTRDGG-YESNE | 453 |
| 11 | GSHDTNGTLTLPCRIKQIINMWQGVGKAMYAPPIEGKIRCSSNITGLLLTRDGG-YESNK | 470 |
| 12 | GSHDTNDTLMLPCKIKQIINMWQGVGKAMYAPPIEGQIRCSSNITGLLLTRDGG-YESNE | 453 |
| 13 | GSHDTNGKLILPCRIKQIINMWQGVGKAMYAPPIEGIIRCSSNITGLLLTRDGG-YESNE | 470 |
| 14 | GSHDTNGTLILPCRIKQIINMWQGVGKAMYAPPIEGEIRCSSNITGLLLTRDGG-YESNE | 453 |
| 15 | GSHDTNGKLILPCRIKQIINMWQGVGKAMYAPPIEGIIRCSSNITGLLLTRDGG-YESNE | 470 |
| 16 | GSHDTNGKLILPCRIKQIINMWQGVGKAMYAPPIEGIIRCSSNITGLLLTRDGG-YESNE | 453 |
| 17 | GSHDTNDTLMLPCKIKQIINMWQGVGKAMYAPPIEGQIRCSSNITGLLLTRDGG-YESNE | 453 |
| 18 | GSHDTNGKLILPCRIKQIINMWQGVGKAMYAPPIEGIIRCSSNITGLLLTRDGG-YESNE | 470 |
| 19 | GSHDTNGKLILPCRIKQIINMWQGVGKAMYAPPIEGIIRCSSNITGLLLTRDGG-YESNE | 470 |
| 20 | GSHDTNGTLTLPCRIKQIINMWQGVGKAMYAPPIEGKIRCSSNITGLLLTRDGG-YESNK | 453 |
| 21 | GSHDTNDTLMLPCKIKQIINMWQGVGKAMYAPPIEGQIRCSSNITGLLLTRDGG-YESNE | 453 |
| 22 | GSHDTNGTLILPCRIKQIINMWQGVGKAMYAPPIEGEIRCSSNITGLLLTRDGG-YESNE | 453 |
| 23 | GSHDTNDTLMLPCKIKQIINMWQGVGKAMYAPPIEGQIRCSSNITGLLLTRDGG-YESNE | 453 |
| 24 | GSHDTNGTLILPCRIKQIINMWQGVGKAMYAPPIEGEIRCSSNITGLLLTRDGG-YESNE | 453 |
| 25 | GSHDTNGKLILPCRIKQIINMWQGVGKAMYAPPIEGIIRCSSNITGLLLTRDGG-YESNE | 470 |
| 26 | GSHDTNGTLILPCKIKQIINMWQGVGKAMYAPPIEEKIRCSSNITGLLLTRDGG-YESNE | 453 |

```
     TDEIFRPGGGDMRDNWRSELYKYKVVKIEPLGVAPTKAKRRVVQREKRAFG-LGAVFLGF

1    TDEIFRPGGGDMRDNWRSELYKYKVVKIEPLGVAPTKAKRRVVQREKRAFG-LGAVFLGF   512
2    TDEIFRPGGGDMRDNWRSELYKYKVVKIEPLGVAPTKAKRRVVQREKRAFG-LGALFLGF   529
3    TDEIFRPGGGDMRDNWRSELYKYKVVKIEPLGVAPTKAKRRVVQREKRAFG-LGAVFLGF   512
4    TDKIFRPGGGDMRDNWRSELYKYKVVKIEPLGVAPTKAKRRVVQREKRAFG-LGAVFLGF   529
5    TDEIFRPGGGDMRDNWRSELYKYKVVKIEPLGVAPTKAKRRVVQREKRAFG-LGAVFLGF   512
6    TDEIFRPGGGDMRDNWRSELYKYKVVKIEPLGVAPTKAKRRVVQREKRAFG-LGAVFLGF   512
7    TDEIFRPGGGDMRDNWRSELYKYKVVKIEPLGVAPTKAKRRVVQREKRAFG-LGAVFLGF   529
8    TDEIFRPGGGDMRDNWRSELYKYKVVKIEPLGVAPTKAKRRVVQREKRAFG-LGAVFLGF   512
9    TDEIFRPGGGDMRDNWRSELYKYKVVKIEPLGVAPTKAKRRVVQREKRAFG-LGAVFLGF   512
10   TDEIFRPGGGDMRDNWRSELYKYKVVKIEPLGVAPTKAKRRMVQREKRAFG-LGAVFLGF   512
11   TDEIFRPGGGDMRDNWRSELYKYKVVKIEPLGVAPTKAKRRVVQREKRAFG-LGAVFLGF   529
12   TDEIFRPGGGDMRDNWRSELYKDKVVKIEPLGVAPTKAKRRVVQREKRAFG-LGAVFLGF   512
13   TDEIFRPGGGDMRDNWRSELYKYKVVKIEPLGVAPTKAKRRVVQREKRAFG-LGALFLGF   529
14   TDEIFRPGGGDMRDNWRSELYKYKVVKIEPLGVAPTKAKRRVVQREKRAFG-LGAVFLGF   512
15   TDEIFRPGGGDMRDNWRSELYKYKVVKIEPLGVAPTKAKRRVVQREKRAFG-LGAVFLGF   529
16   TDEIFRPGGGDMRDNWRSELYKYKVVKIEPLGVAPTKAKRRVVQREKRAFG-LGAVFLGF   512
17   TDEIFRPGGGDMRDNWRSELYKYKVVKIEPLGVAPTKAKRRVVQREKRAFG-LGAVFLGF   512
18   TDEIFRPGGGDMRDNWRSELYKYKVVKIEPLGVAPTKAKRRVVQREKRAFG-LGAVFLGF   529
19   TDEIFRPGGGDMRDNWRSELYKYKVVKIEPLGVAPTKAKRRVVQREKRAFG-LGAVFLGF   529
20   TDEIFRPGGGDMRDNWRSELYKYKVVKIEPLGVAPTKAKRRVVQREKRAFG-LGAVFLGF   512
21   TDEIFRPGGGDMRDNWRSELYKYKVVKIEPLGVAPTKAKRRVVQREKRAFG-LGAVFLGF   512
22   TDEIFRPGGGDMRDNWRSELYKYKVVKIEPLGVAPTKAKRRVVQREKRAFG-LGAVFLGF   512
23   TDEIFRPGGGDMRDNWRSELYKYKVVKIEPLGVAPTKAKRRVVQREKRAFG-LGAVFLGF   512
24   TDEIFRPGGGDMRDNWRSELYKYKVVKIEPLGVAPTKAKRRVVQREKRAFG-LGAVFLGF   512
25   TDEIFRPGGGDMRDNWRSELYKYKVVKIEPLGVAPTKAKRRVVQREKRAFG-LGAVFLGF   529
26   TDEIFRPGGGDMRDNWRSELYKYKVVKIEPLGVAPTKAKRRVVQREKRAFG-LGAVFLGF   512
```

**LGAAGSTMGAASITLTVQARQLLSGIVQQQNNLLRAIEAQQHLLQLTVWGIKQLQARVLA**

| | | |
|---|---|---|
| 1 | LGAAGSTMGAASITLTVQARQLLSGIVQQQNNLLRAIEAQQHLLQLTVWGIKQLQARVLA | 572 |
| 2 | LGAAGSTMGAASITLTVQARQLLSGIVQQQNNLLRAIEAQQHLLQLTVWGIKQLQARVLA | 589 |
| 3 | LGAAGSTMGAASITLTVQARQLLSGIVQQQNNLLRAIEAQQHLLQLTVWGIKQLQARVLA | 572 |
| 4 | LGAAGSTMGAASITLTVQARQLLSGIVQQQNNLLRAIEAQQHLLQLTVWGIKQLQARVLA | 589 |
| 5 | LGAAGSTMGAASITLTVQARQLLSGIVQQQNNLLRAIEAQQHLLQLTVWGIKQLQARVLA | 572 |
| 6 | LGAAGSTMGAASITLTVQARQLLSGIVQQQNNLLRAIEAQQHLLQLTVWGIKQLQARVLA | 572 |
| 7 | LGAAGSTMGAASITLTVQARQLLSGIVQQQNNLLRAIEAQQHLLQLTVWGIKQLQARVLA | 589 |
| 8 | LGAAGSTMGAASITLTVQARQLLSGIVQQQNNLLRAIEAQQHLLQLTVWGIKQLQARVLA | 572 |
| 9 | LGAAGSTMGAASITLTVQARQLLSGIVQQQNNLLRAIEAQQHLLQLTVWGIKQLQARVLA | 572 |
| 10 | LGAAGSTMGAASITLTVQARQLLSGIVQQQNNLLRAIEAQQHLLQLTVWGIKQLQARVLA | 572 |
| 11 | LGAAGSTMGAASITLTVQARQLLSGIVQQQNNLLRAIEAQQHLLQLTVWGIKQLQARVLA | 589 |
| 12 | LGAAGSTMGAASITLTVQARQLLSGIVQQQNNLLRAIEAQQHLLQLTVWGIKQLQARVLA | 572 |
| 13 | LGAAGSTMGAASITLTVQARQLLSGIVQQQNNLLRAIEAQQHLLQLTVWGIKQLQARVLA | 589 |
| 14 | LGAAGSTMGAASITLTVQARQLLSGIVQQQNNLLRAIEAQQHLLQLTVWGIKQLQARVLA | 572 |
| 15 | LGAAGSTMGAASITLTVQARQLLSGIVQQQNNLLRAIEAQQHLLQLTVWGIKQLQARVLA | 589 |
| 16 | LGAAGSTMGAASITLTVQARQLLSGIVQQQNNLLRAIEAQQHLLQLTVWGIKQLQARVLA | 572 |
| 17 | LGAAGSTMGAASITLTVQARQLLSGIVQQQNNLLRAIEAQQHLLQLTVWGIKQLQARVLA | 572 |
| 18 | LGAAGSTMGAASITLTVQARQLLSGIVQQQNNLLRAIEAQQHLLQLTVWGIKQLQARVLA | 589 |
| 19 | LGAAGSTMGAASITLTVQARQLLSGIVQQQNNLLRAIEAQQHLLQLTVWGIKQLQARVLA | 589 |
| 20 | LGAAGSTMGAASITLTVQARQLLSGIVQQQNNLLRAIEAQQHLLQLTVWGIKQLQARVLA | 572 |
| 21 | LGAAGSTMGAASITLTVQARQLLSGIVQQQNNLLRAIEAQQHLLQLTVWGIKQLQARVLA | 572 |
| 22 | LGAAGSTMGAASITLTVQARQLLSGIVQQQNNLLRAIEAQQHLLQLTVWGIKQLQARVLA | 572 |
| 23 | LGAAGSTMGAASITLTVQARQLLSGIVQQQNNLLRAIEAQQHLLQLTVWGIKQLQARVLA | 572 |
| 24 | LGAAGSTMGAASITLTVQARQLLSGIVQQQNNLLRAIEAQQHLLQLTVWGIKQLQARVLA | 572 |
| 25 | LGAAGSTMGAASITLTVQARQLLSGIVQQQNNLLRAIEAQQHLLQLTVWGIKQLQARVLA | 589 |
| 26 | LGAAGSTMGAASITLTVQARQLLSGIVQQQNNLLRAIEAQQHLLQLTVWGIKQLQARVLA | 572 |

| | | |
|---|---|---|
| | **VERYLKDQQLLGIWGCSGKLICTTTVPWNTSWSNKSLEQIWDNMTWMEWEREIDNYTGYI** | |
| 1 | VERYLKDQQLLGIWGCSGKLICTTTVPWNTSWSNKSLEQIWDNMTWMEWEREIDNYTGYI | 632 |
| 2 | VERYLKDQQLLGIWGCSGKLICTTTVPWNNSWSNKSLEQIWDNMTWMEWEREIDNYTGYI | 649 |
| 3 | VERYLKDQQLLGIWGCSGKLICPTTVPWNTSWSNKSLEQIWDNMTWMEWEREIDNYTGYI | 632 |
| 4 | VERYLKDQQLLGIWGCSGKLICTTTVPWNTSWSNKSLEQIWDNMTWMEWEREIDNYTGYI | 649 |
| 5 | VERYLKDQQLLGIWGCSGKLICPTTVPWNTSWSNKSLEQIWDNMTWMEWEREIDNYTGYI | 632 |
| 6 | VERYLKDQQLLGIWGCSGKLICPTTVPWNTSWSNKSLEQIWDNMTWMEWEREIDNYTGYI | 632 |
| 7 | VERYLKDQQLLGIWGCSGKLICTTTVPWNTSWSNKSLEQIWDNMTWMEWEREIDNYTGYI | 649 |
| 8 | VERYLKDQQLLGIWGCSGKLICTTTVPWNTSWSNKSLEQIWDNMTWMEWEREIDNYTGYI | 632 |
| 9 | VERYLKDQQLLGIWGCSGKLICTTTVPWNTSWSNKSLEQIWDNMTWMEWEREIDNYTGYI | 632 |
| 10 | VERYLKDQQLLGIWGCSGKLICTTTVPWNTSWSNKSLEQIWDNMTWMEWEREIDNYTGYI | 632 |
| 11 | VERYLKDQQLLGIWGCSGKLICTTTVPWNTSWSNKSLEQIWDNMTWMEWEREIDNYTGYI | 649 |
| 12 | VERYLKDQQLLGIWGCSGKLICPTTVPWNTSWSNKSLEQIWDNMTWMEWEREIDNYTGYI | 632 |
| 13 | VERYLKDQQLLGIWGCSGKLICTTTVPWNNSWSNKSLEQIWDNMTWMEWEREIDNYTGYI | 649 |
| 14 | VERYLKDQQLLGIWGCSGKLICPTTVPWNTSWSNKSLEQIWDNMTWMEWEREIDNYTGYI | 632 |
| 15 | VERYLKDQQLLGIWGCSGKLICTTTVPWNTSWSNKSLEQIWDNMTWMEWEREIDNYTGYI | 649 |
| 16 | VERYLKDQQLLGIWGCSGKLICTTTVPWNTSWSNKSLEQIWDNMTWMEWEREIDNYTGYI | 632 |
| 17 | VERYLKDQQLLGIWGCSGKLICTTTVPWNTSWSNKSLEQIWDNMTWMEWEREIDNYTGYI | 632 |
| 18 | VERYLKDQQLLGIWGCSGKLICTTTVPWNTSWSNKSLEQIWDNMTWMEWEREIDNYTGYI | 649 |
| 19 | VERYLKDQQLLGIWGCSGKLICTTTVPWNTSWSNKSLEQIWDNMTWMEWEREIDNYTGYI | 649 |
| 20 | VERYLKDQQLLGIWGCSGKLICTTTVPWNTSWSNKSLEQIWDNMTWMEWEREIDNYTGYI | 632 |
| 21 | VERYLKDQQLLGIWGCSGKLICTTTVPWNTSWSNKSLEQIWDNMTWMEWEREIDNYTGYI | 632 |
| 22 | VERYLKDQQLLGIWGCSGKLICPTTVPWNTSWSNKSLEQIWDNMTWMEWEREIDNYTGYI | 632 |
| 23 | VERYLKDQQLLGIWGCSGKLICTTTVPWNTSWSNKSLEQIWDNMTWMEWEREIDNYTGYI | 632 |
| 24 | VERYLKDQQLLGIWGCSGKLICTTTVPWNTSWSNKSLEQIWDNMTWMEWEREIDNYTGYI | 632 |
| 25 | VERYLKDQQLLGIWGCSGKLICTTTVPWNTSWSNKSLEQIWDNMTWMEWEREIDNYTGYI | 649 |
| 26 | VERYLKDQQLLGIWGCSGKLICTTTVPWNTSWSNKSLEQIWDNMTWMEWEREIDNYTGYI | 632 |

| | Sequence | |
|---|---|---|
| | **YQLIEESQNQQEKNEQELLALDKWASLWNWFDITNWLWYIKIFIMIVGGLIGLRIVFTVL** | |
| 1 | YQLIEESQNQQEKNEQELLALDKWASLWNWFDITNWLWYIKIFIMIVGGLIGLRIVFTVL | 692 |
| 2 | YQLIEESQNQQEKNEQELLALDKWASLWNWFDITNWLWYIKIFIMIVGGLIGLRIVFTVL | 709 |
| 3 | YQLIEESQNQQEKNEQELLALDKWASLWNWFDITNWLWYIKIFIMIVGGLIGLRIVFTVL | 692 |
| 4 | YQLIEESQNQQEKNEQELLALDKWASLWNWFDITNWLWYIKIFIMIVGGLIGLRIVFTVL | 709 |
| 5 | YQLIEESQNQQEKNEQELLALDKWASLWNWFDITNWLWYIKIFIMIVGGLIGLRIVFTVL | 692 |
| 6 | YQLIEESQNQQEKNEQELLALDKWASLWNWFDITNWLWYIKIFIMIVGGLIGLRIVFTVL | 692 |
| 7 | YQLIEESQNQQEKNEQELLALDKWASLWNWFDITNWLWYIKIFIMIVGGLIGLRIVFTVL | 709 |
| 8 | YQLIEESQNQQEKNEQELLALDKWASLWNWFDITNWLWYIKIFIMIVGGLIGLRIVFTVL | 692 |
| 9 | YQLIEESQNQQEKNEQELLALDKWASLWNWFDITNWLWYIKIFIMIVGGLIGLRIVFTVL | 692 |
| 10 | YQLIEESQNQQEKNEQELLALDKWASLWNWFDITNWLWYIKIFIMIVGGLIGLRIVFTVL | 692 |
| 11 | YQLIEESQNQQEKNEQELLALDKWASLWNWFDITNWLWYIKIFIMIVGGLIGLRIVFTVL | 709 |
| 12 | YQLIEESQNQQEKNEQELLALDKWASLWNWFDITNWLWYIKIFIMIVGGLIGLRIVFTVL | 692 |
| 13 | YQLIEESQNQQEKNEQELLALDKWASLWNWFDITNWLWYIKIFIMIVGGLIGLRIVFTVL | 709 |
| 14 | YQLIEESQNQQEKNEQELLALDKWASLWNWFDITNWLWYIKIFIMIVGGLIGLRIVFTVL | 692 |
| 15 | YQLIEESQNQQEKNEQELLALDKWASLWNWFDITNWLWYIKIFIMIVGGLIGLRIVFTVL | 709 |
| 16 | YQLIEESQNQQEKNEQELLALDKWASLWNWFDITNWLWYIKIFIMIVGGLIGLRIVFTVL | 692 |
| 17 | YQLIEESQNQQEKNEQELLALDKWASLWNWFDITNWLWYIKIFIMIVGGLIGLRIVFTVL | 692 |
| 18 | YQLIEESQNQQEKNEQELLALDKWASLWNWFDITNWLWYIKIFIMIVGGLIGLRIVFTVL | 709 |
| 19 | YQLIEESQNQQEKNEQELLALDKWASLWNWFDITNWLWYIKIFIMIVGGLIGLRLVFTVL | 709 |
| 20 | YQLIEESQNQQEKNEQELLALDKWASLWNWFDITNWLWYIKIFIMIVGGLIGLRIVFTVL | 692 |
| 21 | YQLIEESQNQQEKNEQELLALDKWASLWNWFDITNWLWYIKIFIMIVGGLIGLRIVFTVL | 692 |
| 22 | YQLIEESQNQQEKNEQELLVLDKWASLWNWFDITNWLWYIKIFIMIVGGLIGLRIVFTVL | 692 |
| 23 | YQLIEESQNQQEKNEQELLALDKWASLWNWFDITNWLWYIKIFIMIVGGLIGLRIVFTVL | 692 |
| 24 | YQLIEESQNQQEKNEQELLALDKWASLWNWFDITNWLWYIKIFIMIVGGLIGLRIVFTVL | 692 |
| 25 | YQLIEESQNQQEKNEQELLALDKWASLWNWFDITNWLWYIKIFIMIVGGLIGLRIVFTVL | 709 |
| 26 | YQLIEESQNQQEKNEQELLALDKWASLWNWFDITNWLWYIKIFIMIVGGLIGLRIVFTVL | 692 |

| | | |
|---|---|---|
| | **SIVNRVRQGYSPLSFQTHLPAQRGPDRPEGIGEEGGERDRDRSDPLVNGFLTLIWSDLRS** | |
| 1 | SIVNRVRQGYSPLSFQTHLPAQRGPDRPEGIGEEGGERDRDRSDPLVNGFLALIWSDLRS | 752 |
| 2 | SIVNRVRQGYSPLSFQTHLPAQRGPDRPEGIGEEGGERDRDRSDPLVNGFLTLIWSDLRS | 769 |
| 3 | SIVNRVRQGYSPLSFQTHLPAQRGPDRPEGIGEEGGERDRDRSDPLVNGFLTLIWSDLRS | 752 |
| 4 | SIVNRVRQGYSPLSFQTHLPAQRGPDRPEGIGEEGGERDRDRSDPLVNGFLTLIWSDLRS | 769 |
| 5 | SIVNRVRQGYSPLSFQTHLPAQRGPDRPEGIGEEGGERDRDRSDPLVNGFLTLIWSDLRS | 752 |
| 6 | SIVNRVRQGYSPLSFQTHLPAQRGPDRPEGIGEEGGERDRDRSDPLVNGFLTLIWSDLRS | 752 |
| 7 | SIVNRVRQGYSPLSFQTHLPAQRGPDRPEGIGEEGGERDRDRSDPLVNGFLTLIWSDLRS | 769 |
| 8 | SIVNRVRQGYSPLSFQTHLPAQRGPDRPEGIGEEGGERDRDRSDPLVNGFLTLIWSDLRS | 752 |
| 9 | SIVNRVRQGYSPLSFQTHLPAQRGPDRPEGIGEEGGERDRDRSDPLVNGFLTLIWSDLRS | 752 |
| 10 | SIVNRVRQGYSPLSFQTHLPAQRGPDRPEGIGEEGGERDRDRSDPLVNGFLTLIWSDLRS | 752 |
| 11 | SIVNRVRQGYSPLSFQTHLPAQRGPDRPEGIGEEGGERDRDRSDPLVNGFLTLIWSDLRS | 769 |
| 12 | SIVNRVRQGYSPLSFQTHLPAQRGPDRPEGIGEEGGERDRDRSDPLVNGFLTLIWSDLRS | 752 |
| 13 | SIVNRVRQGYSPLSFQTHLPAQRGPDRPEGIGEEGGERDRDRSDPLVNGFLTLIWSDLRS | 769 |
| 14 | AIVNRVRQGYAPLSFQTHLPAQRGPDRPEGIGEEGGERDRDRSDPLVNGFLTLIWSDLRS | 752 |
| 15 | SIVNRVRQGYSPLSFQTHLPAQRGPDRPEGIGEEGGERDRDRSDPLVNGFLTLIWSDLRS | 769 |
| 16 | SIVNRVRQGYSPLSFQTHLPAQRGPDRPEGIGEEGGERDRDRSDPLVNGFLTLIWSDLRS | 752 |
| 17 | SIVNRVRQGYSPLSFQTHLPAQRGPDRPEGIGEEGGERDRDRSDPLVNGFLTLIWSDLRS | 752 |
| 18 | SIVNRVRQGYSPLSFQTHLLAQRGPDRPEGIGEEGGERDRDRSDPLVNGFLTLIWSDLRS | 769 |
| 19 | SIVNRVRQGYSPLSFQTHLPAQRGPDRPEGIGEEGGERDRDRSDPLVNGFLTLIWSDLRS | 769 |
| 20 | SIVNRVRQGYSPLSFQTHLPAQRGPDRPEGIGEEGGERDRDRSDPLVNGFLTLIWSDLRS | 752 |
| 21 | SIVNRVRQGYSPLSFQTHLPAQRGPDRPEGIGEEGGERDRDRSDPLVNGFLTLIWSDLRS | 752 |
| 22 | SIVNRVRQGYSPLSFQTHLPAQRGPDRPEGIGEEGGERDRDRSDPLVNGFLTLIWSDLRS | 752 |
| 23 | SIVNRVRQGYSPLSFQTHLPAQRGPDRPEGIGEEGGERDRDRSDPLVNGFLTLIWSDLRS | 752 |
| 24 | SIVNRVRQGYSPLSFQTHLPAQRGPDRPEGIGEEGGERDRDRSDPLVNGFLTLIWSDLRS | 752 |
| 25 | SIVNRVRQGYSPLSFQTHLPAQRGPDRPEGIGEEGGERDRDRSDPLVNGFLTLIWSDLRS | 769 |
| 26 | SIVNRVRQGYSPLSFQTHLPAQRGPDRPEGIGEEGGKRDRDRSDPLVNGFLALIWSDLRS | 752 |

| | | |
|---|---|---|
|  | LCLFSYHRLRDLLLIVTRIVELLGR-------RGWEVLKYWWNLLQYWSQELKNSAVSLL |  |
| 1 | LCLFSYHRLRDLLLIVTRIVELLGR-------RGWEVLKYWWNLLQYWSQELKNSAVSLL | 805 |
| 2 | LCLFSYHRLRDLLLIVTRIVELLGR-------RGWEVLKYWWNLLQYWSQELKNSAVSLL | 822 |
| 3 | LCLFSYHRLRDLLLIVTRIVELLGR-------RGWEVLKYWWNLLQYWSQELKNSAVSLL | 805 |
| 4 | LCLFSYHRLRDLLLIVTRIVELLGR-------RGWEVLKYWWNLLQYWSQELKNSAVSLL | 822 |
| 5 | LCLFSYHRLRDLLLIVTRIVELLGR-------RGWEVLKYWWNLLQYWSQELKNSAVSLL | 805 |
| 6 | LCLFSYHRLRDLLLI----VELLGR-------RGWEVLKYWWNLLQYWSQELKNSAVSLL | 801 |
| 7 | LCLFSYHRLRDLLLIVTRIVELLGR-------RGWEVLKYWWNLLQYWSQELKNSAVSLL | 822 |
| 8 | LCLFSYHRLRDLLLIVTRIVELLGR-------RGWEVLKYWWNLLQYWSQELKNSAVSLL | 805 |
| 9 | LCLFSYHRLRDLLLIVTRIVELLGR-------RGWEVLKYWWNLLQYWSQELKNSAVSLL | 805 |
| 10 | LCLFSYHRLRDLLLIVTRIVELLGR-------RGWEVLKYWWNLLQYWSQELKNSAVSLL | 805 |
| 11 | LCLFSYHRLRDLLLIVTRIVELLGR-------RGWEVLKYWWNLLQYWSQELKNSAVSLL | 822 |
| 12 | LCLFSYHHLRDLLLIVTRIVELLGR-------RGWEVLKYWWNLLQYWSQELKNSAVSLL | 805 |
| 13 | LCLFSYHRLRDLLLIVTRIVELLGR-------RGWEVLKYWWNLLQYWSQELKNSAVSLL | 822 |
| 14 | LCLFSYHRLRDLLLIVTRIVELLGR-------RGWEVLKYWWNLLQYWSQELKNSAVSLL | 805 |
| 15 | LCLFSYHRLRDLLLIVTRIVELLGR-------RGWEVLKYWWNLLQYWSQELKNSAVSLL | 822 |
| 16 | LCLFSYHSLRDLLLIVTRIVELLGR-------RGWEVLKYWWNLLQYWSQELKNSAVSLL | 805 |
| 17 | LCLFSYHRLRDLLLIVTRIVELLGR-------RGWEVLKYWWNLLQYWSQELKNSAVSLL | 805 |
| 18 | LCLFSYHRLRDLLLIVTRIVELLGR-------RGWEVLKYWWNLLQYWSQELKNSAVSLL | 822 |
| 19 | LCLFSYHRLRDLLLIVTRIVELLGR-------RGWEVLKYWWNLLQYWSQELKNSAVSLL | 822 |
| 20 | LCLFSYHRLRDLLLIVTRIVELLGR-------RGWEVLKYWWNLLQYWSQELKNSAVSLL | 805 |
| 21 | LCLFSYHRLRDLLLIVTRIVELLGR-------RGWEVLKYWWNLLQYWSQELKNSAVSLL | 805 |
| 22 | LCLFSYHRLRDLLLIVTRIVELLGR-------RGWEVLKYWWNLLQYWSQELKNSAVSLL | 805 |
| 23 | LCLFSYHRLRDLLLIVTRIVELLGR-------RGWEVLKYWWNLLQYWSQELKNSAVSLL | 805 |
| 24 | LCLFSYHRLRDLLLIVTRIVELLGR-------RGWEVLKYWWNLLQYWSQELKNSAVSLL | 805 |
| 25 | LCLFSYHRLGDLLLIVTRIVELLGR-------RGWEVLKYWWNLLQYWSQELKNSAVSLL | 822 |
| 26 | LCLFSYHRLRDLLLIVTRIVELLGR-------RGWEVLKYWWNLLQYWSQELKNSAVSLL | 805 |

| | | |
|---|---|---|
| | **NATAIAVAEGTDRVIEVVQRACRAILHIPRRIRQGLERALL** | |
| 1 | NATAIAVAEGTDRVIEVVQRACRAILHIPRRIRQGLERALL | 846 |
| 2 | NATAIAVAEGTDRVIEVVQGACRAILHIPRRIRQGLERALL | 863 |
| 3 | NATAIAVAEGTDRVIEVVQRACRAILHIPRRIRQGLERALL | 846 |
| 4 | NATATAVAEGTDRVIEVVQRACRAILHIPRRIRQGLERALL | 863 |
| 5 | NATAIAVAEGTDRVIEVVQRACRAILHIPRRIRQGLERALL | 846 |
| 6 | NATAIAVAEGTDRVIEVVQRACRAILHIPRRIRQGLERALL | 842 |
| 7 | NATAIAVAEGTDRVIEVVQRACRAIRHIPRRIRQGLERALL | 863 |
| 8 | NATATAVAEGTDRVIEVVQRACRAILHIPRRIRQGLERALL | 846 |
| 9 | NATAIAVAEGTDRVIEVVQGACRAILHIPRRIRQGLERALL | 846 |
| 10 | NATAIAVAEGTDRVIEVVQRAGRAIRHIPRRIRQGLERALL | 846 |
| 11 | NTTAIAVAEGTDRVIEVVQGACRAILHIPRRIRQGLERALL | 863 |
| 12 | NATAIAVAEGTDRVIEVVQRACRAILHIPRRIRQGLERALL | 846 |
| 13 | NATAIAVAEGTDRVIEVVQGACRAILHIPRRIRQGLERALL | 863 |
| 14 | NATAIAVAEGTDRVIEVVQRACRAILHIPRRIRQGLERALL | 846 |
| 15 | NATAIAVAEGTDRVIEVVQGACRAILHIPRRIRQGLERALL | 863 |
| 16 | NATAIAVAEGTDRVIEVVQRAGRAIRHIPRRIRQGLERALL | 846 |
| 17 | NATAIAVAEGTDRVIEVVQRAGRAIRHIPRRIRQGLERALL | 846 |
| 18 | NATAIAVAEGTDRVIEVVQGACRAILHIPRRIRQGLERALL | 863 |
| 19 | NATAIAVAEGTDRVIEVVQGACRAILHIPRRIRQGLERALL | 863 |
| 20 | NATATAVAEGTDRVIEVVQRACRAILHIPRRIRQGLERALL | 846 |
| 21 | NATAIAVAEGTDRVIEVVQRAGRAIRHIPRRIRQGLERALL | 846 |
| 22 | NATAIAVAEGTDRVIEVVQRACRAILHIPRRIRQGLERALL | 846 |
| 23 | NATAIAVAEGTDRVIEVVQRAGRAIRHIPRRIRQGLERALL | 846 |
| 24 | NATATAVAEGTDRVIEVVQRACRAILHIPRRIRQGLERALL | 846 |
| 25 | NATAIAVAEGTDRVIEVVQRACRAICHIPRRIRQGLERALL | 863 |
| 26 | NATAIAVAEGTDRVIEVVQRACRAILHIPRRIRQGLERALL | 846 |

**Consensus**

1. B.US.2006.700010040_C9_4520.EU289193
2. B.US.2010.CH0040_3_d1485_ipe032_15_08.MG900412
3. B.US.2010.CH0040_3_d1485_ipe032_15_09.MG900413
4. B.US.2010.CH0040_3_d1485_ipe032_15_10.MG900414
5. B.US.2010.CH0040_3_d1485_ipe032_15_11.MG900415
6. B.US.2010.CH0040_3_d1485_ipe032_15_13.MG900416
7. B.US.2010.CH0040_3_d1485_ipe032_15_16.MG900417
8. B.US.2010.CH0040_3_d1485_ipe032_15_18.MG900418
9. B.US.2010.CH0040_3_d1485_ipe032_15_19.MG900419
10. B.US.2010.CH0040_3_d1485_ipe032_15_24.MG900420
11. B.US.2010.CH0040_3_d1485_ipe032_15_25.MG900421
12. B.US.2010.CH0040_3_d1485_ipe032_15_27.MG900422
13. B.US.2010.CH0040_3_d1485_ipe032_15_28.MG900423
14. B.US.2010.CH0040_3_d1485_ipe032_15_29.MG900424
15. B.US.2010.CH0040_3_d1485_ipe032_15_30.MG900425
16. B.US.2010.CH0040_3_d1485_ipe032_15_31.MG900426
17. B.US.2010.CH0040_3_d1485_ipe032_15_32.MG900427
18. B.US.2010.CH0040_3_d1485_ipe032_15_33.MG900428
19. B.US.2010.CH0040_3_d1485_ipe032_15_37.MG900429
20. B.US.2010.CH0040_3_d1485_ipe032_15_38.MG900430
21. B.US.2010.CH0040_3_d1485_ipe032_15_39.MG900431
22. B.US.2010.CH0040_3_d1485_ipe032_15_41.MG900432
23. B.US.2010.CH0040_3_d1485_ipe032_15_42.MG900433
24. B.US.2010.CH0040_3_d1485_ipe032_15_45.MG900434
25. B.US.2010.CH0040_3_d1485_ipe032_15_46.MG900435
26. B.US.2010.CH0040_3_d1485_ipe032_15_47.MG900436
27. B.US.2010.CH0040_3_d1485_ipe032_15_48.MG900437
28. B.US.2010.CH0040_3_d1485_ipe032_27_06.MG900438
29. B.US.2010.CH0040_3_d1485_ipe032_2_49.MG900439
30. B.US.2010.CH0040_3_d1485_ipe032_2_50.MG900440
31. B.US.2010.CH0040_3_d1485_ipe032_9_03.MG900441

**MRVMGIRKNYQHLWREGILLLGILMICSAADNLWVTVYYGVPVWREATTTLFCASDAKAY**

| | | |
|---|---|---|
| 1 | MRVMGIRKNYQHLWREGILLLGILMICSAADNLWVTVYYGVPVWREATTTLFCASDAKAY | 60 |
| 2 | MRVMGIRKNYQHLWREGILLLGILMICSAADNLWVTVYYGVPVWREATTTLFCASDAKAY | 60 |
| 3 | MRVMGIRKNYQHLWREGILLLGILMICSAADNLWVTVYYGVPVWREATTTLFCASDAKAY | 60 |
| 4 | MRAMGIRKNYQHLWREGILLLGILMICSAADNLWVTVYYGVPVWREATTTLFCASDAKAY | 60 |
| 5 | MRVMGIRKNYQHWWREGILLLGILMICSAADKLWVTVYYGVPVWREATTTLFCASDAKAY | 60 |
| 6 | MRVMGIRKNYQHLWREGILLLGILMICSAADNLWVTVYYGVPVWREATTTLFCASDAKAY | 60 |
| 7 | MRVMGIRKNYQHLWREGILLLGILMICSAADNLWVTVYYGVPVWREATTTLFCASDAKAY | 60 |
| 8 | MRVMGIRKNYQHLWREGILLLGILMICSAADNLWVTVYYGVPVWREATTTLFCASDAKAY | 60 |
| 9 | MRVMGIRKNYQHLWREGILLLGILMICSAADNLWVTVYYGVPVWREATTTLFCASDAKAY | 60 |
| 10 | MRVMGIRKNYQHLWREGILLLGILMICSAADNLWVTVYYGVPVWREATTTLFCASDAKAY | 60 |
| 11 | MRVMGIRKNYQHLWREGILLLGILMICSAADNLWVTVYYGVPVWREATTTLFCASDAKAY | 60 |
| 12 | MRVMGIRKNYQHLWRKGILLLGILMICSAADNLWVTVYYGVPVWREATTTLFCASDAKAY | 60 |
| 13 | MRVMGIRKNYQHLWREGILLLGILMICSAADNLWVTVYYGVPVWREATTTLFCASDAKAY | 60 |
| 14 | MRVMGIRKNYQHLWRKGILLLGILMICSAADNLWVTVYYGVPVWREATTTLFCASDAKAY | 60 |
| 15 | MRVMGIRKNYQHLWREGILLLGILMICSAADNLWVTVYYGVPVWREATTTLFCASDAKAY | 60 |
| 16 | MRVMGIRKNYQHLWREGILLLGILMICSAADNLWVTVYYGVPVWREATTTLFCASDAKAY | 60 |
| 17 | MRVMGIRKNYQHLWREGILLLGILMICSAADNLWVTVYYGVPVWREATTTLFCASDAKAY | 60 |
| 18 | MRVMGIRKNYQHLWREGILLLGILMICSAADNLWVTVYYGVPVWREATTTLFCASDAKAY | 60 |
| 19 | MRVMGIRKNYQHLWREGILLLGILMICSAADNLWVTVYYGVPVWREATTTLFCASDAKAY | 60 |
| 20 | MRVMGIRKNYQHLWREGILLLGILMICSAADNLWVTVYYGVPVWKEATTTLFCASDAKAY | 60 |
| 21 | MRVMGIRKNYQHLWREGILLLGILMICSAADNLWVTVYYGVPVWREATTTLFCASDAKAY | 60 |
| 22 | MRVMGIRKNYQHLWREGILLLGILMICSAADNLWVTVYYGVPVWREATTTLFCASDAKAY | 60 |
| 23 | MRVMGIRKNYQHLWREGILLLGILMICSAADKLWVTVYYGVPVWREATTTLFCASDAKAY | 60 |
| 24 | MRVMGIRKNYQHWWREGILLLGILMICSAADKLWVTVYYGVPVWREATTTLFCASDAKAY | 60 |
| 25 | MRVMGIRKNYQHLWREGILLLGILMICSAADNLWVTVYYGVPVWREATTTLFCASDAKAY | 60 |
| 26 | MRVMGIRKNYQHLWREGILLLGILMICSAADNLWVTVYYGVPVWREATTTLFCASDAKAY | 60 |
| 27 | MRVMGIRKNYQHLWREGILLLGILMICSATDNLWVTVYYGVPVWREATTTLFCASDAKAY | 60 |
| 28 | MRVMGIRKNYQHLWKEGILLLGILMICSAADNLWVTVYYGVPVWREATTTLFCASDAKAY | 60 |
| 29 | MRVMGIRKNYQHLWRKGILLLGILMICSAADNLWVTVYYGVPVWREATTTLFCASDAKAY | 60 |
| 30 | MRVMGIRKNYQHLWRKGILLLGILMICSAADNLWVTVYYGVPVWREATTTLFCASDAKAY | 60 |
| 31 | MRVMGIRKNYQHLWREGILLLGILMICSAADNLWVTVYYGVPVWREATTTLFCASDAKAY | 60 |

**DTEAHNVWATHACVPTDPNPQEVELKNVTENFNMWENNMVEQMHEDIISLWDQSLKPCVK**

| | | |
|---|---|---|
| 1 | DTEAHNVWATHACVPTDPNPQEVELKNVTENFNMWENNMVEQMHEDIISLWDQSLKPCVK | 120 |
| 2 | DTEAHNVWATHACVPTDPNPQEVELKNVTENFNMGKNNMVEQMHEDIISLWDQSLKPCVK | 120 |
| 3 | DTEAHNVWATHACVPTDPNPQEVELKNVTENFNMWENNMVEQMHEDIISLWDQSLKPCVK | 120 |
| 4 | DTEAHNVWATHACVPTDPNPQEVELKNVTENFNMWENNMVEQMHEDIISLWDQSLKPCVK | 120 |
| 5 | DTEAHNVWATHACVPTDPNPQEVELKNVTENFNMWENNMVEQMHEDIISLWDQSLKPCVK | 120 |
| 6 | DTEAHNVWATHACVPTDPNPQEVELKNVTENFNMWENNMVEQMHEDIISLWDQSLKPCVK | 120 |
| 7 | DTEAHNVWATHACVPTDPNPQEVELKNVTENFNMWENNMVEQMHEDIISLWDQSLKPCVK | 120 |
| 8 | DTEAHNVWATHACVPTDPNPQEVELKNVTENFNMWENNMVEQMHEDIISLWDQSLKPCVK | 120 |
| 9 | DTEAHNVWATHACVPTDPNPQEVELKNVTENFNMWENNMVEQMHEDIISLWDQSLKPCVK | 120 |
| 10 | DTEAHNVWATHACVPTDPNPQEVELKNVTENFNMWENNMVEQMHEDIISLWDQSLKPCVK | 120 |
| 11 | DTEAHNGWATHACVPTDPNPQEVELKNVTENFNMWENNMVEQMHEDIISLWDQSLKPCVK | 120 |
| 12 | DTEAHNVWATHACVPTDPNPQEVELKNVTENFNMWENNMVEQMHEDIISLWDQSLKPCVK | 120 |
| 13 | DTEAHNVWATHACVPTDPNPQEVELKNVTENFNMWENNMVEQMHEDIISLWDQSLKPCVK | 120 |
| 14 | DTEAHNVWATHACVPTDPNPQEVELKNVTENFNMWENNMVEQMHEDIISLWDQSLKPCVK | 120 |
| 15 | DTEAHNVWATHACVPTDPNPQEVELKNVTENFNMWENNMVEQMHEDIISLWDQSLKPCVK | 120 |
| 16 | DTEAHNVWATHACVPTDPNPQEVELKNVTENFNMWENNMVEQMHEDIISLWDQSLKPCVK | 120 |
| 17 | DTEAHNVWATHACVPTDPNPQEVELKNVTENFNMWENNMVEQMHEDIISLWDQSLKPCVK | 120 |
| 18 | DTEAHNVWATHACVPTDPNPQEVELKNVTENFNMWENNMVEQMHEDIISLWDQSLKPCVK | 120 |
| 19 | DTEAHNVWATHACVPTDPNPQEVELKNVTENFNMWENNMVEQMHEDIISLWDQSLKPCVK | 120 |
| 20 | DTEAHNVWATHACVPTDPNPQEVELKNVTENFNMWENNMVEQMHEDIISLWDQSLKPCVK | 120 |
| 21 | DTEAHNVWATHACVPTDPNPQEVKLKNVTENFNMWENNMVEQMHEDIISLWDQSLKPCVK | 120 |
| 22 | DTEAHNVWATHACVPTDPNPQEVELKNVTENFNMWENNMVEQMHEDIISLWDQSLKPCVK | 120 |
| 23 | DTEAHNVWATHACVPTDPNPQEVELKNVTENFNMWENNMVEQMHEDIISLWDQSLKPCVK | 120 |
| 24 | DTEAHNVWATHACVPTDPNPQEVELKNVTENFNMWENNMVEQMHEDIISLWDQSLKPCVK | 120 |
| 25 | DTEAHNVWATHACVPTDPNPQEVELKNVTENFNMWENNMVEQMHEDIISLWDQSLKPCVK | 120 |
| 26 | DTEAHNVWATHACVPTDPNPQEVELKNVTENFNMWENNMVEQMHEDIISLWDQSLKPCVK | 120 |
| 27 | DTEAHNVWATHACVPTDPNPQEVELKNVTENFNMGKNNMVEQMHEDIISLWDQSLKPCVK | 120 |
| 28 | DTEAHNVWATHACVPTDPNPQEVELKNVTENFNMWENNMVEQMHEDIISLWDQSLKPCVK | 120 |
| 29 | DTEAHNVWATHACVPTDPNPQEVELKNVTENFNMWENNMVEQMHEDIISLWDQSLKPCVK | 120 |
| 30 | DTEAHNVWATHACVPTDPNPQEVELKNVTENFNMWENNMVEQMHEDIISLWDQSLKPCVK | 120 |
| 31 | DTEAHNVWATHACVPTDPNPQEVELKNVTENFNMWENNMVEQMHEDIISLWDQSLKPCVK | 120 |

| | | |
|---|---|---|
|  | LTPLCVTLNCTDNVTTTTNRNGTNCTDXXNVTXT---------TNXNETLMEKGEVKNCS |  |
| 1 | LTPLCVTLNCTDLGNVTNTT------------------------NSNGEMMEKGEVKNCS | 156 |
| 2 | LTPLCVTLNCTDNVTTTTNRNGTNCTDLRNVTTT---------TNSNETLMEKGEVKNCS | 171 |
| 3 | LTPLCVTLNCTDNVTTTTNRNGTNCTDSGNVTNT---------TNRNGALMEKGEVKNCS | 171 |
| 4 | LTPLCVTLNCTDSGNGTTTTNRNGTL-----------------MGNETLMMEKGEVKNCS | 163 |
| 5 | LTPLCVTLNCTDNVTTTTNRNGTNCTDLRNVTNT---------TNSNETLMEKGEVKNCS | 171 |
| 6 | LTPLCVTLNCTDNVTTTTNRNGTNCTDLRNVTTT---------TNSNETLMEKGEVKNCS | 171 |
| 7 | LTPLCVTLNCTDLENVTTTTNRNGTNCTDSGNVTPTTNRNGT-LMGNETSMEKGEVKNCS | 179 |
| 8 | LTPLCVTLNCTDLGNVTTTTNRNGTNCTDSGNVTAT-------TNRNETFMEKEEV-NCS | 172 |
| 9 | LTPLCVTLNCTDLRNVTTTTNRNGTNCTDSGNGTTTTNRNGTLMGNETLMMEKGEVKNCS | 180 |
| 10 | LTPLCVTLNCTDNVTTTTNRNGTNCTDSGNVTNT---------TNRNGALMEKGEVKNCS | 171 |
| 11 | LTPLCVTLNCTDNVTTTTNRNGTNCTDLRNVTTT---------TNSNETWMEKGEVKNCS | 171 |
| 12 | LTPLCVTLNCTDNVTTTTNRNGTNCTDSGNVTNT---------TNRNGALMEKGEVKNCS | 171 |
| 13 | LTPLCVTLNCTDNVTTTTNRNGTNCTDSGNVTNT---------TNRNGALMEKGEVKNCS | 171 |
| 14 | LTPLCVTLNCTDNVTTTTNRNGTNCTDLRNVTTT---------TNSNETLMEKGEVKNCS | 171 |
| 15 | LTPLCVTLNCTDNVTTTTNRNGTNCTDLRNVTTT---------TNSNETLMEKGEVKNCS | 171 |
| 16 | LTPLCVTLNCTDNVTTTTNRNGTNCTDSGNVTNT---------TNRNGALMEKGEVKNCS | 171 |
| 17 | LTPLCVTLNCTDNVTTTTNRNGTNCTDLGNVNTT---------TNSNETLMEKGEVKNCS | 171 |
| 18 | LTPLCVTLNCTDNVTTTTNRNGTNCTDLGNVNTT---------TNSNETLMEKGEVKNCS | 171 |
| 19 | LTPLCVTLNCTDNVTTTTNRNGTNCTDSGNVTNT---------TNRNGALMEKGEVKNCS | 171 |
| 20 | LTPLCVTLNCTDVGNVTTIT------------------------NSNGTLMEKGEVKNCS | 156 |
| 21 | LTPLCVTLNCTDDLGNVTTTTNRNGTNCTDSGNVTNT------TNRNGALMEKGEVKNCS | 174 |
| 22 | LTPLCVTLNCTDNVTTTTNRNGTNCTDLRNVTTT---------TNSNETLMEKGEVKNCS | 171 |
| 23 | LTPLCVTLNCTDLENVTTTTNRNGTNCTDSGNVTPTTNRNGT-LMGNETLMEKGEVKNCS | 179 |
| 24 | LTPLCVTLNCTDNVTTTTNRNGTNCTDLRNVTTT---------TNSNETLMEKGEVKNCS | 171 |
| 25 | LTPLCVTLNCTDLENVTTTTNRNGINCTDSGNVTPTTNRNGT-LMGNETLMEKGEVKNCS | 179 |
| 26 | LTPLCVTLNCTDNVTTTTNRNGTNCTDLRNVTTT---------TNSNETLMEKGEVKNCS | 171 |
| 27 | LTPLCVTLNCTDNVTTTTNRNETNCTDLRNVTTT---------TNSNETLMEKGEVKNCS | 171 |
| 28 | LTPLCVTLNCTDNVTTTTNRNGTNCTDSGNVTNT---------TNRNGALMEKGEVKNCS | 171 |
| 29 | LTPLCVTLNCTDNVTTTTNRNGTNCTDLGNVNTT---------TNSNETLMEKGEVKNCS | 171 |
| 30 | LTPLCVTLNCTDNVTTTTNRNGTNCTDSGNVTNT---------TNRNGALMEKGEVKNCS | 171 |
| 31 | LTPLCVTLNCTDLRNVTTTTNRNGTNCTDSGNGTTTTNRNGTLMGNETLMMEKGEVKNCS | 180 |

| | | |
|---|---|---|
|  | FRITTDMQDRTRKEYALFYKLDVVPIND-------TRYRLVSCNTSVITQACPKVSFEPI |  |
| 1 | FKITTDIKDRTRKEYALFYKLDVVPIND-------TRYRLVSCNTSVITQACPKVSFEPI | 209 |
| 2 | FMITTDVKDRMRKEYALFYKLDVVPIND-------TRYRLVSCNTSVITQACPKVSFEPI | 224 |
| 3 | FRITTDMQDRTRKEYALFYKLDVVPIND-------TRYRLVSCNTSVITQACPKVSFEPI | 224 |
| 4 | FRITTDMKDRTRKEYALFYKLDVVPIND-------TSYRLVSCNTSVITQACPKVSFEPI | 216 |
| 5 | FMITTDVKDRMRKEYALFYKLDVVPIND-------TRYRLVSCNTSVITQACPKVSFEPI | 224 |
| 6 | FMITTDVKDRMRKEYALFYKLDVVPIND-------TRYRLVSCNTSVITQACPKVSFEPI | 224 |
| 7 | FRITTDMQDRTRKEYALFYKLDVVPIND-------TRYRLVSCNTSVITQACPKVSFEPI | 232 |
| 8 | FRITTDMQDRTRKEYALFYKLDVVPIND-------TRYRLVSCNTSVITQACPKVSFEPI | 225 |
| 9 | FRITTDMKDRTRKEYALFYKLDVVPIND-------TSYRLVSCNTSVITQACPKVSFEPI | 233 |
| 10 | FRITTDMQDRTRKEYALFYKLDVVPIND-------TRYRLVSCNTSVITQACPKVSFEPI | 224 |
| 11 | FRITTDMQDRTRKEYALFYKLDVVPIND-------TRYRLVSCNTSVITQACPKVSFEPI | 224 |
| 12 | FRITTDMQDRTRKEYALFYKLDVVPIND-------TRYRLVSCNTSVITQACPKVSFEPI | 224 |
| 13 | FRITTDMQDRTRKEYALFYKLDVVPIND-------TRYRLVSCNTSVITQACPKVSFEPI | 224 |
| 14 | FMITTDMQDRTRKEYALFYKLDVVPIND-------TRYRLVSCNTSVITQACPKVSFEPI | 224 |
| 15 | FMITTDVKDRMRKEYALFYKLDVVPIND-------TRYRLVSCNTSVITQACPKVSFEPI | 224 |
| 16 | FRITTDMQDRTRKEYALFYKLDVVPIND-------TRYRLVSCNTSVITQACPKVSFEPI | 224 |
| 17 | FMITTDVKDRMRKEYALFYKLDVVPIND-------TRYRLVSCNTSVITQACPKVSFEPI | 224 |
| 18 | FMITTDVKDRMRKEYALFYKLDVVPIND-------TRYRLVSCNTSVITQACPKVSFEPI | 224 |
| 19 | FRITTDMQDRTRKEYALFYKLDVVPIND-------TRYRLVSCNTSVITQACPKVSFEPI | 224 |
| 20 | FKITTDTRDRMRKEYALFYKLDVVPIND-------TRYRFVSCNTSVITQACPKVSFEPI | 209 |
| 21 | FRITTDMQDRTRKEYALFYKLDVVPIND-------TSYRLVSCNTSVITQACPKVSFEPI | 227 |
| 22 | FMITTDMQDRTRKEYALFYKLDVVPIND-------TRYRLVSCNTSVITQACPKVSFEPI | 224 |
| 23 | FRITTDMQDRTRKEYALFYKLDVVPIND-------TRYRLVSCNTSVITQACPKVSFEPI | 232 |
| 24 | FMITTDVKDRMRKEYALFYKLDVVPIND-------TRYRLVSCNTSVITQACPKVSFEPI | 224 |
| 25 | FRITTDMQDRTRKEYALFYKLDVVPIND-------TRYRLVSCNTSVITQACPKVSFEPI | 232 |
| 26 | FMITTDVKDRMRKEYALFYKLDVVPIND-------TRYRLVSCNTSVITQACPKVSFEPI | 224 |
| 27 | FMITTDVKDRMRKEYALFYKLDVVPIND-------TRYRLVSCNTSVLTQACPKVSFEPI | 224 |
| 28 | FRITTDMQDRTRKEYALFYKLDVVPIND-------TRYRLVSCNTSVITQACPKVSFEPI | 224 |
| 29 | FRITTDMQDRTRKEYALFYKLDVVPIND-------TRYRLVSCNTSVITQACPKVSFEPI | 224 |
| 30 | FRITTDMQDRTRKEYALFYKLDVVPIND-------TRYRLVSCNTSVITQACPKVSFEPI | 224 |
| 31 | FRITTDMKDRTRKEYALFYKLDVVPIND-------TSYRLVSCNTSVITQACPKVSFEPI | 233 |

**PIHYCAPAGFAILKCNDKQFNGTGPCTNVSTVQCTHGIRPVVSTQLLLNGSLAEEEVVIR**

| | | |
|---|---|---|
| 1 | PIHYCAPAGFAILKCNDKQFIGTGPCTNVSTVQCTHGIRPVVSTQLLLNGSLAEEEVVIR | 269 |
| 2 | PIHYCAPAGFAILKCNDKQFNGTGPCTNVSTVQCTHGIRPVVSTQLLLNGSLAEEEVVIR | 284 |
| 3 | PIHYCAPAGFAILKCNDKQFNGTGPCTNVSTVQCTHGIRPVVSTQLLLNGSLAEEEVVIR | 284 |
| 4 | PIHYCAPAGFAILKCNDKQFNGTGPCTNVSTVQCTHGIRPVVSTQLLLNGSLAEEEVVIR | 276 |
| 5 | PIHYCAPAGFAILKCNDKQFNGTGPCTNVSTVQCTHGIRPVVSTQLLLNGSLAEEEVVIR | 284 |
| 6 | PIHYCAPAGFAILKCNDKQFNGTGPCTNVSTVQCTHGIRPVVSTQLLLNGSLAEEEVVIR | 284 |
| 7 | PIHYCAPAGFAILKCNDKQFNGTGPCTNVSTVQCTHGIRPVVSTQLLLNGSLAEEEVVIR | 292 |
| 8 | PIHYCAPAGFAILKCNDKQFNGTGPCTNVSTVQCTHGIRPVVSTQLLLNGSLAEEEVVIR | 285 |
| 9 | PIHYCAPAGFAILKCNDKQFNGTGPCTNVSTVQCTHGIRPVVSTQLLLNGSLAEEEVVIR | 293 |
| 10 | PIHYCAPAGFAILKCNDKQFNGTGPCTNVSTVQCTHGIRPVVSTQLLLNGSLAEEEVVIR | 284 |
| 11 | PIHYCAPAGFAILKCNDKQFNGTGPCTNVSTVQCTHGIRPVVSTQLLLNGSLAEEEVVIR | 284 |
| 12 | PIHYCAPAGFAILKCNDKQFNGTGPCTNVSTVQCTHGIRPVVSTQLLLNGSLAEEEVVIR | 284 |
| 13 | PIHYCAPAGFAILKCNDKQFNGTGPCTNVSTVQCTHGIRPVVSTQLLLNGSLAEEEVVIR | 284 |
| 14 | PIHYCAPAGFAILKCNDKQFNGTGPCTNVSTVQCTHGIRPVVSTQLLLNGSLAEEEVVIR | 284 |
| 15 | PIHYCAPAGFAILKCNDKQFNGTGPCTNVSTVQCTHGIRPVVSTQLLLNGSLAEEEVVIR | 284 |
| 16 | PIHYCAPAGFAILKCNDKQFNGTGPCTNVSTVQCTHGIRPVVSTQLLLNGSLAEEEVVIR | 284 |
| 17 | PIHYCAPAGFAILKCNDKQFNGTGPCTNVSTVQCTHGIRPVVSTQLLLNGSLAEEEVVIR | 284 |
| 18 | PIHYCAPAGFAILKCNDKQFNGTGPCTNVSTVQCTHGIRPVVSTQLLLNGSLAEEEVVIR | 284 |
| 19 | PIHYCAPAGFAILKCNDKQFNGTGPCTNVSTVQCTHGIRPVVSTQLLLNGSLAEEEVVIR | 284 |
| 20 | PIHYCAPAGFAILKCNDKQFNGTGPCTNVSTVQCTHGIRPVVSTQLLLNGSLAEEEVVIR | 269 |
| 21 | PIHYCAPAGFAILKCNDKQFNGTGPCTNVSTVQCTHEIRPVVSTQLLLNGSLAEEEVVIR | 287 |
| 22 | PIHYCAPAGFAILKCNDKQFNGTGPCTNVSTVQCTHGIRPVVSTQLLLNGSLAEEEVVIR | 284 |
| 23 | PIHYCAPAGFAILKCNDKQFNGTGPCTNVSTVQCTHGIRPVVSTQLLLNGSLAEEEVVIR | 292 |
| 24 | PIHYCAPAGFAILKCNDKQFNGTGPCTNVSTVQCTHGIRPVVSTQLLLNGSLAEEEVVIR | 284 |
| 25 | PIHYCAPAGFAILKCNDKQFNGTGPCTNVSTVQCTHGIRPVVSTQLLLNGSLAEEEVVIR | 292 |
| 26 | PIHYCAPAGFAILKCNDKQFNGTGPCTNVSTVQCTHGIRPVVSTQLLLNGSLAEEEVVIR | 284 |
| 27 | PIHYCAPAGFAILKCNDKQFNGTGPCTNVSTVQCTHGIRPVVSTQLLLNGSLAEEEVVIR | 284 |
| 28 | PIHYCAPAGFAILKCNDKQFNGTGPCTNVSTVQCTHGIRPVVSTQLLLNGSLAEEEVVIR | 284 |
| 29 | PIHYCAPAGFAILKCNDKQFNGTGPCTNVSTVQCTHGIRPVVSTQLLLNGSLAEEEVVIR | 284 |
| 30 | PIHYCAPAGFAILKCNDKQFNGTGPCTNVSTVQCTHGIRPVVSTQLLLNGSLAEEEVVIR | 284 |
| 31 | PIHYCAPAGFAILKCNDKQFNGTGPCTNVSTVQCTHGIRPVVSTQLLLNGSLAEEEVVIR | 293 |

| | | |
|---|---|---|
| | **SVNFSDNAKTIIVQLNKSVEINCTRPNNNTRKSIHMGPGKAFYARGAIIGDIRKAHCNIS** | |
| 1 | SVNFSDNAKTIIVQLNKSVEITCTRPNNNTRKSIPMGPGKAFYARGDITGDIRKAYCEIN | 329 |
| 2 | SVNFSDNAKTIIVQLNKSVEINCTRPNNNTRKSIPMGPGKAFYARGAIIGDIRKAHCNIS | 344 |
| 3 | SVNFSDNAKTIIVQLNKSVEINCTRPNNNTRKSIHMGPGKAFYARGGIIGDIRKAHCNIS | 344 |
| 4 | SVNFSDNAKTIIVQLNKSVEINCTRP--NTRKSIPMGPGKAFYARGAIIGDIRKAHCNIS | 334 |
| 5 | SVNFSDNAKTIIVQLNKSVEINCTRPNNNTRKSIPMGPGKAFYARGAIIGDIRKAHCNIS | 344 |
| 6 | SVNFSDNAKTIIVQLNKSVEINCTRPNNNTRKSIPMGPGKAFYARGAIIGDIRKAHCNIS | 344 |
| 7 | SVNFSDNAKTIIVQLNKSVEINCTRPNNNTRKSIHMGPGKAFYARGAIIGDIRKAHCNIS | 352 |
| 8 | SVNFSDNAKTIIVQLNKSVEINCTRPNNNTRKSIHMGPGKAFYARGAIIGDIRKAHCNIS | 345 |
| 9 | SVNFSDNAKTIIVQLNKSVEINCTRPNNNTRKSIHMGPGKAFYARGAIIGDIRKAHCNIS | 353 |
| 10 | SVNFSDNAKTIIVQLNKSVEINCTRPNNNTRKSIHMGPGKAFYARGGIIGDIRKAHCNIS | 344 |
| 11 | SVNFSDNAKTIIVQLNKSVEINCTRPNNNTRKSIPMGPGKAFYARGAIIGDIRKAHCNIS | 344 |
| 12 | SVNFSDNAKTIIVQLNKSVEINCTRPNNNTRKSIHMGPGKAFYARGGIIGDIRKAHCNIS | 344 |
| 13 | SVNFSDNAKTIIVQLNKSVEINCTRPNNNTRKSIHMGPGKAFYARGGIIGDIRKAHCNIS | 344 |
| 14 | SVNFSDNAKTIIVQLNKSVEINCTRPNNNTRKSIHMGPGKAFYARGGIIGDIRKAHCNIS | 344 |
| 15 | SVNFSDNAKTIIVQLNKSVEINCTRPNNNTRKSIPMGPGKAFYARGAIIGDIRKAHCNIS | 344 |
| 16 | SVNFSDNAKTIIVQLNKSVEINCTRPNNNTRKSIHMGPGKAFYARGGIIGDIRKAHCNIS | 344 |
| 17 | SVNFSDNAKTIIVQLNKSVEINCTRPNNNTRKSIPMGPGKAFYARGAIIGDIRKAHCNIS | 344 |
| 18 | SVNFSDNAKTIIVQLNKSVEINCTRPNNNTRKSIPMGPGKAFYARGAIIGDIRKAHCNIS | 344 |
| 19 | SVNFSDNAKTIIVQLNKSVEINCTRPNNNTRKSIHMGPGKAFYARGGIIGDIRKAHCNIS | 344 |
| 20 | SVNFSDNVKTIIVQLNKSVEINCTRPNNNTRKSIPMGPGKAFYARGNIIGDIRKAYCNIS | 329 |
| 21 | SVNFSDNAKTIIVQLNKSVEINCTRPNNNTRKSIHMGPGKAFYARGGIIGDIRKAYCNVS | 347 |
| 22 | SVNFSDNAKTIIVQLNKSVEINCTRPNNNTRKSIHMGPGKAFYARGGIIGDIRKAHCNIS | 344 |
| 23 | SVNFSDNAKTIIVQLNKSVEINCTRPNNNTRKSIHMGPGKAFYARGAIIGDIRKAHCNIS | 352 |
| 24 | SVNFSDNAKTIIVQLNKSVEINCTRPNNNTRKSIPMGPGKAFYARGAIIGDIRKAHCNIS | 344 |
| 25 | SVNFSDNAKTIIVQLNKSVEINCTRPNNNTRKSIHMGPGKAFYARGAIIGDIRKAHCNIS | 352 |
| 26 | SVNFSDNAKTIIVQLNKSVEINCTRPNNNTRKSIPMGPGKAFYARGAIIGDIRKAHCNIS | 344 |
| 27 | SVNFSDNAKTIIVQLNKSVEINCTRPNNNTRKSIHMGPGKAFYARGGIIGDIRKAHCNIS | 344 |
| 28 | SVNFSDNAKTIIVQLNKSVEINCTRPNNNTRKSIHMGPGKAFYARGGIIGDIRKAHCNIS | 344 |
| 29 | SVNFSDNAKTIIVQLNKSVEINCTRPNNNTRKSIHMGPGKAFYARGGIIGDIRKAH-NIS | 343 |
| 30 | SVNFSDNAKTIIVQLNKSVEINCTRPNNNTRKSIHMGPGKAFYARGAIIGDIRKAHCNIS | 344 |
| 31 | SVNFSDNAKTIIVQLNKSVEINCTRPNNNTRKSIHMGPGKAFYARGAIIGDIRKAHCNIS | 353 |

| | | |
|---|---|---|
| | GTEWHSTLKLVVEKLREQY-NKTIVFNHSSGGDPEIVMYSFNCGGEFFYCNSTKLFNSTW | |
| 1 | GTEWHSTLKLVVEKLREQY-NKTIVFNRSSGGDPEIVMYSFNCGGEFFYCNSTKLFNSTW | 388 |
| 2 | RAEWHSTLKLVAEKLREQY-NKTIVFNHSSGGDPEIVMYSFNCGGEFFYCNSTKLFNSTW | 403 |
| 3 | GTEWHSTLKLVVKKLREQY-NKTIVFNHSSGGDPEIVMYSFNCGGEFFYCNSTKLFNSTW | 403 |
| 4 | GTEWHSTLKLVVEKLREKY-NKTIVFNHSSGGDPEIVMYSFNCGGEFFYCNSTKLFNSTW | 393 |
| 5 | GTEWHNTLKLVVEKLREQY-NKTIVFNHSSGGDPEIVMYSFNCGGEFFYCNSTKLFNSTW | 403 |
| 6 | RAEWHSTLKLVAEKLREQY-NKTIVFNHSSGGDPEIVMYSFNCGGEFFYCNSTKLFNSTW | 403 |
| 7 | GTEWHNTLKLVVEKLREQY-NKTIVFNHSSGGDPEIVMYSFNCGGEFFYCNSTKLFNSTW | 411 |
| 8 | RAEWHSTLKLVAEKLREQY-NKTIVFNHSSGGDPEIVMYSFNCGGEFFYCNSTKLFNSTW | 404 |
| 9 | GTEWHNTLKLVVEKLREQY-NKTIVFNHSSGGDPEIVMYSFNCGGEFFYCNSTKLFNSTW | 412 |
| 10 | GTEWHSTLKLVVKKLREQY-NKTIVFNHSSGGDPEIVMYSFNCGGEFFYCNSTKLFNSTW | 403 |
| 11 | RAEWHSTLKLVAEKLREQY-NKTIVFNHSSGGDPEIVMYSFNCGGEFFYCNSTKLFNSTW | 403 |
| 12 | GTEWHSTLKLVVKKLREQY-NKTIVFNHSSGGDPEIVMYSFNCGGEFFYCNSTKLFNSTW | 403 |
| 13 | GTEWHSTLKLVVKKLREQY-NKTIVFNHSSGGDPEIVMYSFNCGGEFFYCNSTKLFNSTW | 403 |
| 14 | GTEWHSTLKLVGEKLREQY-NKTIVFNHSSGGDPEIVMYSFNCGGEFFYCNSTKLFNSTW | 403 |
| 15 | RAEWHSTLKLVAEKLREQY-NKTIVFNHSSGGDPEIVMYSFNCGGEFFYCNSTKLFNSTW | 403 |
| 16 | GTEWHSTLKLVVKKLREQY-NKTIVFNHSSGGDPEIVMYSFNCGGEFFYCNSTKLFNSTW | 403 |
| 17 | GTEWHNTLKLVVEKLREQY-NKTIVFNHSSGGDPEIVMYSFNCGGEFFYCNSTKLFNSTW | 403 |
| 18 | GTEWHNTLKLVVEKLREQY-NKTIVFNHSSGGDPEIVMYSFNCGGEFFYCNSTKLFNSTW | 403 |
| 19 | RTEWHSTLKLVAEKLREQY-NKTIVFNHSSGGDPEIVMYSFNCGGEFFYCNSTKLFNSTW | 403 |
| 20 | GPEWYSTLKLVVEKLREQY-NKTIVFNHSSGGDPEIVMYSFNCGGEFFYCNSTKLFNSTW | 388 |
| 21 | RTEWHSTLKLVVKKLREKY-NKTIVFNHSSGGDPEIVMYSFNCGGEFFYCNSTKLFNSTW | 406 |
| 22 | RAEWHSTLKLVAEKLREQY-NKTIVFNHSSGGDPEIVMYSFNCGGEFFYCNSTKLFNSTW | 403 |
| 23 | GTEWHNTLKLVVEKLREQY-NKTIVFNHSSGGDPEIVMYSFNCGGEFFYCNSTKLFNSTW | 411 |
| 24 | GTEWHNTLKLVVEKLREQY-NKTIVFNHSSGGDPEIVMYSFNCGGEFFYCNSTKLFNSTW | 403 |
| 25 | GTEWHNTLKLVVEKLREQY-NKTIVFNHSSGGDPEIVMYSFNCGGEFFYCNSTKLFNSTW | 411 |
| 26 | GTEWHNTLKLVVEKLREQY-NKTIVFNHSSGGDPEIVMYSFNCGGEFFYCNSTKLFNSTW | 403 |
| 27 | GTEWHSTLKLVGEKLREQY-NKTIVFNHSSGGDPEIVMYSFNCGGEFFYCNSTKLFNSTW | 403 |
| 28 | GTEWHSTLKLVVKKLREQY-NKTIVFNHSSGGDPEIVMYSFNCGGEFFYCNSTKLFNSTW | 403 |
| 29 | GTEWHSTLKLVVKKLREQY-NKTIVFNHSSGGDPEIVMYSFNCGGEFFYCNSTKLFNSTW | 402 |
| 30 | GTEWHNTLKLVVEKLREQY-NKTIVFNHSSGGDPEIVMYSFNCGGEFFYCNSTKLFNSTW | 403 |
| 31 | RAEWHNTLKLVVEKLREQY-NKTIVFNHSSGGDPEIVMYSFNCGGEFFYCNSTKLFNSTW | 412 |

| | Sequence | |
|---|---|---|
| | **PWND--TXGSHDTNGTLTLPCRIKQIINMWQGVGKAMYAPPIEGIIKCSSNITGLLLTRD** | |
| 1 | PWND--TKGSHDTNGTLILPCKIKQIINMWQGVGKAMYAPPIEGKIRCSSNITGLLLTRD | 446 |
| 2 | PWND--TQGSHDTNGTLTLPCRIKQIINMWQEVGKAMYAPPIEGIIKCSSNITGLLLTRD | 461 |
| 3 | PWND--TEGSHDTNGTLTLPCRIKQIINMWQGVGKAMYAPPIEGIIKCSSNITGLLLTRD | 461 |
| 4 | PWND--TEGSHDTNGTLTLPCRIKQIINMWQGVGKAMYAPPIEGIIKCSSNITGLLLTRD | 451 |
| 5 | PWNG--TNGSHDTNGTLTLPCRIKQIINMWQGVGKAMYAPPIEGIIKCSSNITGLLLTRD | 461 |
| 6 | PWND--TNGSHDTNGTLTLPCRIKQIINMWQEVGKAMYAPPIEGIIKCSSNITGLLLTRD | 461 |
| 7 | PGND--TEGSHNTNGTLTLPCRIKQIINMWQEVGKAMYAPPIEGIIKCSSNITGLLLTRD | 469 |
| 8 | --------SGMILKGHMTL---MKQIINMWQEVGKAMYAPPIEGIIKCSSNITGLLLTRD | 453 |
| 9 | PGND--TNGSHDINGTLTLPCRIKQIINMWQGVGKAMYAPPIEGIIKCSSNITGLLLTRD | 470 |
| 10 | PWNDTDTEGSHDTNGTLTLPCRIKQIINMWQGVGKAMYAPPIEGIIKCSSNITGLLLTRD | 463 |
| 11 | PWND--TNGSHDTNGTLTLPCRIKQIINMWQGVGKAMYAPPIEGIIKCSSNITGLLLTRD | 461 |
| 12 | PWNDTDTEGSHDTNGTLTLPCRIKQIINMWQGVGKAMYAPPIEGIIKCSSNITGLLLTRD | 463 |
| 13 | PWNDTDTGGSHDTNGTLTLPCRIKQIINMWQGVGKAMYAPPIEGIIKCSSNITGLLLTRD | 463 |
| 14 | PWND--TNGSHDTNGTLTLPCRIKQIINMWQGVGKAMYAPPIEGIIKCSSNITGLLLTRD | 461 |
| 15 | PWND--TNGSHDTNGTLTLPCRIKQIINMWQGVGKAMYAPPIEGIIKCSSNITGLLLTRD | 461 |
| 16 | PWNDTDTEGSHDTNGTLTLPCRIKQIINMWQGVGKAMYAPPIEGIIKCSSNITGLLLTRD | 463 |
| 17 | PGND--TKGSHDNNGTLTLPCRIKQIINMWQGVGKAMYAPPIEGIIKCSSNITGLLLTRD | 461 |
| 18 | PGND--TNGSHDTNGTLTLPCRIKQIINMWQGVGKAMYAPPIEGIIKCSSNITGLLLTRD | 461 |
| 19 | PWND--TEGSHDTNGTLTLPCRIKQIINMWQGVGKAMYAPPIEGIIKCSSNITGLLLTRD | 461 |
| 20 | PWND--TNGSYDTNDTLMLPCKIKQIINMWQGVGKAMYAPPIEGEIRCSSNITGLLLTRD | 446 |
| 21 | PWND--TNGSHDTNGTLTLPCRIKQIINMWQGVGKAMYAPPIEGIIKCSSNITGLLLTRD | 464 |
| 22 | PWND--TQGSHDTNGTLTLPCRIKQIINMWQEVGKAMYAPPIEGIIKCSSNITGLLLTRD | 461 |
| 23 | PGND--TEGSHNTNGTLTLPCRIKQIINMWQEVGKAMYAPPIEGIIKCSSNITGLLLTRD | 469 |
| 24 | PWNG--TNGSHDTNGTLTLPCRIKQIINMWQGVGKAMYAPPIEGIIKCSSNITGLLLTRD | 461 |
| 25 | PGND--TEGSHNTNGTLTLPCRIKQIINMWQEVGKAMYAPPIEGIIKCSSNITGLLLTRD | 469 |
| 26 | PWNG--TNGSHDTNGTLTLPCRIKQIINMWQGVGKAMYAPPIEGIIKCSSNITGLLLTRD | 461 |
| 27 | PWND--TEGSHDTNGTLTLPCRIKQIINMWQEVGKAMYAPPIEGIIKCSSNITGLLLTRD | 461 |
| 28 | PWNDTDTEGSHDTNGTLTLPCRIKQIINMWQGVGKAMYAPPIEGIIKCSSNITGLLLTRD | 463 |
| 29 | PWNDTDTEGSHDTNGTLTLPCRIKQIINMWQGVGKAMYAPPIEGIIKCSSNITGLLLTRD | 462 |
| 30 | PGND--TEGSHNTNGTLTLPCRIKQIINMWQEVGKAMYAPPIEGIIKCSSNITGLLLTRD | 461 |
| 31 | PGND--TNGSHDINGTLTLPCRIKQIINMWQGVGKAMYAPPIEGIIKCSSNITGLLLTRD | 470 |

| | | |
|---|---|---|
| | GG-YESNKTDEIFRPGGGDMRDNWRSELYKYKVVKIEPLGVAPTKAKRRVVQREKRAFG- | |
| 1 | GG-YESNETDEIFRPGGGDMRDNWRSELYKYKVVKIEPLGVAPTKAKRRVVQREKRAFG- | 504 |
| 2 | GG-YESNKTDEIFRPGGGDMRDNWRSELYKYKVVKIEPLGVAPTKAKRRVVQREKRAFG- | 519 |
| 3 | GG-YESNKTDEIFRPGGGDMRDNWRSELYKYKVVKIEPLGVAPTKAKRRVVQREKRAFG- | 519 |
| 4 | GG-YESNKTDEIFRPGGGDMRDNWRSELYKYKVVKIEPLGVAPTKAKRRVVQREKRAFG- | 509 |
| 5 | GG-YESNKTDEIFRPGGGDMRDNWRSELYKYKVVKIEPLGVAPTKAKRRVVQREKRAFG- | 519 |
| 6 | GG-YESNKTDEIFRPGGGDMRDNWRSELYKYKVVKIEPLGVAPTKAKRRVVQREKRAFG- | 519 |
| 7 | GG-YESNKTDEIFRPGGGDMRDNWRSELYKYKVVKIEPLGVAPTKAKRRVVQREKRAFG- | 527 |
| 8 | GG-YESNKTDEIFRPGGGDMRDNWRSELYKYKVVKIEPLGVAPTKAKRRVVQREKRAFG- | 511 |
| 9 | GG-YESNKTDEIFRPGGGDMRDNWRSELYKYKVVKIEPLGVAPTKAKRRVVQREKRAFG- | 528 |
| 10 | GG-YESNKTDEIFRPGGGDMRDNWRSELYKYKVVKIEPLGVAPTKAKRRVVQREKRAFG- | 521 |
| 11 | GG-YESNKTDEIFRPGGGDMRDNWRSELYKYKVVKIEPLGVAPTKAKRRVVQREKRAFG- | 519 |
| 12 | GG-YESNKTDEIFRPGGGDMRDNWRSELYKYKVVKIEPLGVAPTKAKRRVVQREKRAFG- | 521 |
| 13 | GG-YESNKTDEIFRPGGGDMRDNWRSELYKYKVVKIEPLGVAPTKAKRRVVQREKRAFG- | 521 |
| 14 | GG-YESNKTDEIFRPGGGDMRDNWRSELYKYKVVKIEPLGVAPTKAKRRVVQREKRAFG- | 519 |
| 15 | GG-YESNKTDEIFRPGGGDMRDNWRSELYKYKVV-IEPLGVAPTKAKRRVVQREKRAFG- | 518 |
| 16 | GG-YESNKTDEIFRPGGGDMRDNWRSELYKYKVVKIEPLGVAPTKAKRRVVQREKRAFG- | 521 |
| 17 | GG-YESNKTDEIFRPGGGDMRDNWRSELYKYKVVKIEPLGVAPTKAKRRVVQREKRAFG- | 519 |
| 18 | GG-YESNKTDEIFRPGGGDMRDNWRSELYKYKVVKIEPLGVAPTKAKRRVVQREKRAFG- | 519 |
| 19 | GG-YESNKTDEIFRPGGGDMRDNWRSELYKYKVVKIEPLGVAPTKAKRRVVQREKRAFG- | 519 |
| 20 | GG-YESNETDEIFRPGGGDMRDNWRSELYKYKVVKIEPLGVAPTKAKRRVVQREKRAFG- | 504 |
| 21 | GG-YESNKTDEIFRPGGGDMRDNWRSELYKYKVVKIEPLGVAPAKAKRRVVQREKRAFG- | 522 |
| 22 | GG-YESNKTDEIFRPGGGDMRDNWRSELYKYKVVKIEPLGVAPTKAKRRVVQREKRAFG- | 519 |
| 23 | GG-YESNKTDEIFRPGGGDMRDNWRSELYKYKVVKIEPLGVAPTKAKRRVVQREKRAFG- | 527 |
| 24 | GG-YESNKTDEIFRPGGGDMRDNWRSELYKYKVVKIEPLGVAPTKAKRRVVQREKRAFG- | 519 |
| 25 | GG-YESNKTDEIFRPGGGDMRDNWRSELYKYKVVKIEPLGVAPTKAKRRVVQREKRAFG- | 527 |
| 26 | GG-YESNKTDEIFRPGGGDMRDNWRSELYKYKVVKIEPLGVAPTKAKRRVVQREKRAFG- | 519 |
| 27 | GG-YESNKTDEIFRPGGGDMRDNWRSELYKYKVVKIEPLGVAPTKAKRRVVQREKRAFG- | 519 |
| 28 | GG-YESNKTDEIFRPGGGDMRDNWRSELYKYKVVKIEPLGVAPTKAKRRVVQREKRAFG- | 521 |
| 29 | GG-YESNKTDEIFRPGGGDMRDNWRSELYKYKVVKIEPLGVAPTKAKRRVVQREKRAFG- | 520 |
| 30 | GG-YESNKTDEIFRPGGGDMRDNWRSELYKYKVVKIEPLGVAPTKAKRRVVQREKRAFG- | 519 |
| 31 | GG-YESNKTDEIFRPGGGDMRDNWRSELYKYKVVKIEPLGVAPTKAKRRVVQREKRAFG- | 528 |

| | | |
|---|---|---|
| | **LGAVFLGFLGAAGSTMGAASITLTVQARQLLSGIVQQQNNLLRAIEAQQHLLQLTVWGIK** | |
| 1 | LGAVFLGFLGAAGSTMGAASITLTVQARQLLSGIVQQQNNLLRAIEAQQHLLQLTVWGIK | 564 |
| 2 | LGAVFLGFLGAAGSTMGAASITLTVQARQLLSGIVQQQNNLLRAIEAQQHLLQLTVWGIK | 579 |
| 3 | LGAVFLGFLGAAGSTMGAASITLTVQARQLLSGIVQQQNNLLRAIEAQQHLLQLTVWGIK | 579 |
| 4 | LGAVFLGFLGAAGSTMGAASITLTVQARQLLSGIVQQQNNLLRAIEAQQHLLQLTVWGIK | 569 |
| 5 | LGAVFLGFLGAAGSTMGAASITLTVQARQLLSGIVQQQNNLLRAIEAQQHLLQLTVWGIK | 579 |
| 6 | LGAVFLGFLGAAGSTMGAASITLTVQARQLLSGIVQQQNNLLRAIEAQQHLLQLTVWGIK | 579 |
| 7 | LGAVFLGFLGAAGSTMGAASITLTVQARQLLSGIVQQQNNLLRAIEAQQHLLQLTVWGIK | 587 |
| 8 | LGAVFLGFLGAAGSTMGAASITLTVQARQLLSGIVQQQNNLLRAIEAQQHLLQLTVWGIK | 571 |
| 9 | LGAVFLGFLGAAGSTMGAASITLTVQARQLLSGIVQQQNNLLRAIEAQQHLLQLTVWGIK | 588 |
| 10 | LGAVFLGFLGAAGSTMGAASITLTVQARQLLSGIVQQQNNLLRAIEAQQHLLQLTVWGIK | 581 |
| 11 | LGAVFLGFLGAAGSTMGAASITLTVQARQLLSGIVQQQNNLLRAIEAQQHLLQLTVWGIK | 579 |
| 12 | LGAVFLGFLGAAGSTMGAASITLTVQARQLLSGIVQQQNNLLRAIEAQQHLLQLTVWGIK | 581 |
| 13 | LGAVFLGFLGAAGSTMGAASITLTVQARQLLSGIVQQQNNLLRAIEAQQHLLQLTVWGIK | 581 |
| 14 | LGAVFLGFLGAAGSTMGAASITLTVQARQLLSGIVQQQNNLLRAIEAQQHLLQLTVWGIK | 579 |
| 15 | LGAVFLGFLGAAGSTMGAASITLTVQARQLLSGIVQQQNNLLRAIEAQQHLLQLTVWGIK | 578 |
| 16 | LGAVFLGFLGAAGSTMGAASITLTVQARQLLSGIVQQQNNLLRAIEAQQHLLQLTVWGIK | 581 |
| 17 | LGAVFLGFLGAAGSTMGAASITLTVQARQLLSGIVQQQNNLLRAIEAQQHLLQLTVWGIK | 579 |
| 18 | LGAVFLGFLGAAGSTMGAASITLTVQARQLLSGIVQQQNNLLRAIEAQQHLLQLTVWGIK | 579 |
| 19 | LGAVFLGFLGAAGSTMGAASITLTVQARQLLSGIVQQQNNLLRAIEAQQHLLQLTVWGIK | 579 |
| 20 | LGAVFLGFLGAAGSTMGAASITLTVQARQLLSGIVQQQNNLLRAIEAQQHLLQLTVWGIK | 564 |
| 21 | LGAVFLGFLGAAGSTMGAASITLTVQARQLLSGIVQQQNNLLRTIEAQQHLLQLTVWGIK | 582 |
| 22 | LGAVFLGFLGAAGSTMGAASITLTVQARQLLSGIVQQQNNLLRAIEAQQHLLQLTVWGIK | 579 |
| 23 | LGAVFLGFLGAAGSTMGAASITLTVQARQLLSGIVQQQNNLLRAIEAQQHLLQLTVWGIK | 587 |
| 24 | LGAVFLGFLGAAGSTMGAASITLTVQARQLLSGIVQQQNNLLRAIEAQQHLLQLTVWGIK | 579 |
| 25 | LGAVFLGFLGAAGSTMGAASITLTVQARQLLSGIVQQQNNLLRAIEAQQHLLQLTVWGIK | 587 |
| 26 | LGAVFLGFLGAAGSTMGAASITLTVQARQLLSGIVQQQNNLLRAIEAQQHLLQLTVWGIK | 579 |
| 27 | LGAVFLGFLGAAGSTMGAASITLTVQARQLLSGIVQQQNNLLRAIEAQQHLLQLTVWGIK | 579 |
| 28 | LGAVFLGFLGAAGSTMGAASITLTVQARQLLSGIVQQQNNLLRAIEAQQHLLQLTVWGIK | 581 |
| 29 | LGAVFLGFLGAAGSTMGAASITLTVQARQLLSGIVQQQNNLLRAIEAQQHLLQLTVWGIK | 580 |
| 30 | LGAVFLGFLGAAGSTMGAASITLTVQARQLLSGIVQQQNNLLRAIEAQQHLLQLTVWGIK | 579 |
| 31 | LGAVFLGFLGAAGSTMGAASITLTVQARQLLSGIVQQQNNLLRAIEAQQHLLQLTVWGIK | 588 |

QLQARVLAVERYLKDQQLLGIWGCSGKLICTTTVPWNTSWSNKSLEQIWDNMTWMEWERE

| | | |
|---|---|---|
| 1 | QLQARVLAVERYLKDQQLLGIWGCSGKLICTTTVPWNTSWSNKSLEQIWDNMTWMEWERE | 624 |
| 2 | QLQARVLAVERYLKDQQLLGIWGCSGKLICTTTVPWNTSWSNKSLEQIWDNMTWMEWEKE | 639 |
| 3 | QLQARVLAVERYLKDQQLLGIWGCSGKLICTTTVPWNTSWSNKSLEQIWDNMTWMEWERE | 639 |
| 4 | QLQARVLAVERYLKDQQLLGIWGCSGKLICTTTVPWNTSWSNKSLEQIWDNMTWMEWERE | 629 |
| 5 | QLQARVLAVERYLKDQQLLGIWGCSGKLICTTTVPWNTSWSNKSLEQIWDNMTWMEWERE | 639 |
| 6 | QLQARVLAVERYLKDQQLLGIWGCSGKLICTTTVPWNTSWSNKSLEQIWDNMTWMEWERE | 639 |
| 7 | QLQARVLAVERYLKDQQLLGIWGCSGKLICTTTVPWNTSWSNKSLEQIWDKMTWMEWERE | 647 |
| 8 | QLQARVLAVERYLKDQQLLGIWGCSGKLICTTTVPWNTSWSNKSLEQIWDNMTWMEWERE | 631 |
| 9 | QLQARVLAVERYLKDQQLLGIWGCSGKLICTTTVPWNTSWSNKSLEQIWDKMTWMEWERE | 648 |
| 10 | QLQARVLAVERYLKDQQLLGIWGCSGKLICTTTVPWNTSWSNKSLEQIWDNMTWMEWERE | 641 |
| 11 | QLQARVLAVERYLKDQQLLGIWGCSGKLICTTTVPWNTSWSNKSLEQIWDNMTWMEWERE | 639 |
| 12 | QLQARVLAVERYLKDQQLLGIWGCSGKLICTTTVPWNTSWSNKSLEQIWDKMTWMEWERE | 641 |
| 13 | QLQARVLAVERYLKDQQLLGIWGCSGKLICTTTVPWNTSWSNKSLEQIWDKMTWMEWERE | 641 |
| 14 | QLQARVLAVERYLKDQQLLGIWGCSGKLICTTTVPWNTSWSNKSLEQIWDNMTWMEWERE | 639 |
| 15 | QLQARVLAVERYLKDQQLLGIWGCSGKLICTTTVPWNTSWSNKSLEQIWDNMTWMEWERE | 638 |
| 16 | QLQARVLAVERYLKDQQLLGIWGCSGKLICTTTVPWNTSWSNKSLEQIWDKMTWMEWERE | 641 |
| 17 | QLQARVLAVERYLKDQQLLGIWGCSGKLICTTTVPWNTSWSNKSLEQIWDNMTWMEWERE | 639 |
| 18 | QLQARVLAVERYLKDQQLLGIWGCSGKLICTTTVPWNTSWSNKSLEQIWDNMTWMEWERE | 639 |
| 19 | QLQARVLAVERYLKDQQLLGIWGCSGKLICTTTVPWNTSWSNKSLEQIWDNMTWMEWERE | 639 |
| 20 | QLQARVLAVERYLKDQQLLGIWGCSGKLICTTTVPWNTSWSNKSLEQIWDNMTWMEWERE | 624 |
| 21 | QLQARVLAVERYLKDQQLLGIWGCSGKLICTTTVPWNTSWSNKSLEQIWDNMTWMEWKRE | 642 |
| 22 | QLQARVLAVERYLKDQQLLGIWGCSGKLICTTTVPWNTSWSNKSLEQIWDNMTWMEWERE | 639 |
| 23 | QLQARVLAVERYLKDQQLLGIWGCSGKLICTTTVPWNTSWSNKSLEQIWDNMTWMEWERE | 647 |
| 24 | QLQARVLAVERYLKDQQLLGIWGCSGKLICTTTVPWNTSWSNKSLEQIWDNMTWMEWERE | 639 |
| 25 | QLQARVLAVERYLKDQQLLGIWGCSGKLICTTTVPWNTSWSNKSLEQIWDKMTWMEWERE | 647 |
| 26 | QLQARVLAVERYLKDQQLLGIWGCSGKLICTTTVPWNTSWSNKSLEQIWDKMTWMEWERE | 639 |
| 27 | QLQARVLAVERYLKDQQLLGIWGCSGKLICTTTVPWNTSWSNKSLEQIWDKMTWMEWERE | 639 |
| 28 | QLQARVLAVERYLKDQQLLGIWGCSGKLICTTTVPWNTSWSNKSLEQIWDNMTWMEWERE | 641 |
| 29 | QLQARVLAVERYLKDQQLLGIWGCSGKLICTTTVPWNTSWSNKSLEQIWDKMTWMEWERE | 640 |
| 30 | QLQARVLAVERYLKDQQLLGIWGCSGKLICTTTVPWNTSWSNKSLEQIWDKMTWMEWERE | 639 |
| 31 | QLQARVLAVERYLKDQQLLGIWGCSGKLICTTTVPWNTSWSNKSLEQIWDKMTWMEWERE | 648 |

| | | |
|---|---|---|
| | IDNYTGYIYQLIEESQNQQEKNEQELLALDKWASLWNWFDITNWLWYIKIFIMIVGGLIG | |
| 1 | IDNYTGYIYQLIEESQNQQEKNEQELLALDKWASLWNWFDITNWLWYIKIFIMIVGGLIG | 684 |
| 2 | INNYTGYIYQLIEESQNQQEKNEQELLAWDKWASLWNWFDITNWLWYIKIFIMIVGGLIG | 699 |
| 3 | INNYTGYIYQLIEESQNQQEKNEQELLAWDKWASLWNWFDITNWLWYIKIFIMIVGGLIG | 699 |
| 4 | IDNYTGYIYQLIEESQNQQEKNEQELLALDKWASLWNWFDITNWLWYIKIFIMIVGGLIG | 689 |
| 5 | INNYTGYIYQLIEESQNQQEKNEQELLAWDKWASLWNWFDITNWLWYIKIFIMIVGGLIG | 699 |
| 6 | INNYTGYIYQLIEESQNQQEKNEQELLALDKWASLWNWFDITNWLWYIKIFIMIVGGLIG | 699 |
| 7 | IDNYTDYIYQLIEESQNQQEKNEQELLALDKWASLWNWFDITNWLWYIKIFIMIVGGLIG | 707 |
| 8 | INNYTGYIYQLIEESQNQQEKNEQELLALDKWASLWNWFDITNWLWYIKIFIMIVGGLIG | 691 |
| 9 | IDNYTDYIYQLIEESQNQQEKNEQELLALDKWASLWNWFDITNWLWYIKIFIMIVGGLIG | 708 |
| 10 | IDNYTDYIYQLIEESQNQQEKNEQELLALDKWASLWNWFDITKWLWYIKIFIMIVGGLIG | 701 |
| 11 | INNYTGYIYQLIEESQNQQEKNEQELLALDKWASLWNWFDITNWLWYIKIFIMIVGGLIG | 699 |
| 12 | IDNYTDYIYQLIEESQNQQEKNEQELLALDKWASLWNWFDITNWLWYIKIFIMIVGGLIG | 701 |
| 13 | INNYTGYIYQLIEESQNQQEKNEQELLALDKWASLWNWFDITNWLWYIKIFIMIVGGLIG | 701 |
| 14 | INNYTGYIYQLIEESQNQQEKNEQELLALDKWASLWNWFDITNWLWYIKIFIMIVGGLIG | 699 |
| 15 | INNYTGYIYQLIEESQNQQEKNEQELLAWDKWASLWNWFDITNWLWYIKIFIMIVGGLIG | 698 |
| 16 | IDNYTDYIYQLIEESQNQQEKNEQELLALDKWASLWNWFDITNWLWYIKIFIMIVGGLIG | 701 |
| 17 | INNYTGYIYQLIEESQNQQEKNEQELLAWDKWASLWNWFDITNWLWYIKIFIMIVGGLIG | 699 |
| 18 | INNYTGYIYQLIEESQNQQEKNEQELLAWDKWASLWNWFDITNWLWYIKIFIMIVGGLIG | 699 |
| 19 | INNYTGYIYQLIEESQNQQEKNEQELLAWDKWASLWNWFDITNWLWYIKIFIMIVGGLIG | 699 |
| 20 | IDNYTGYIYQLIEESQNQQEKNEQELLALDKWASLWNWFDITNWLWYIKIFIMIVGGLIG | 684 |
| 21 | INNYTGYIYQLIEESQNQQEKNEQELLAWDKWASL-NWFDITNWLWYIKIFIMIVGGLIG | 701 |
| 22 | INNYTGYIYQLIEESQNQQEKNEQELLALDKWASLWNWFDITNWLWYIKIFIMIVGGLIG | 699 |
| 23 | IDNYTGYIYQLIEESQNQQEKNEQELLALDKWASLWNWFDITNWLWYIKIFIMIVGGLIG | 707 |
| 24 | INNYTGYIYQLIEESQNQQEKNEQELLAWDKWASLWNWFDITNWLWYIKIFIMIVGGLIG | 699 |
| 25 | IDNYTDYIYQLIEESQNQQEKNEQELLALDKWASLWNWFDITNWLWYIKIFIMIVGGLIG | 707 |
| 26 | IDNYTDYIYQLIEESQNQQEKNEQELLALDKWASLWNWFDITNWLWYIKIFIMIVGGLIG | 699 |
| 27 | IDNYTDYIYQLIEESQNQQEKNEQELLALDKWASLWNWFDITNWLWYIKIFIMIVGGLIG | 699 |
| 28 | IDNYTGYIYQLIEESQNQQEKNEQELLAWDKWASLWNWFDITNWLWYIKIFIMIVGGLIG | 701 |
| 29 | IDNYTDYIYQLIEESQNQQEKNEQELLALDKWASLWNWFDITNWLWYIKIFIMIVGGLIG | 700 |
| 30 | IDNYTDYIYQLIEESQNQQEKNEQELLALDKWASLWNWFDITNWLWYIKIFIMIVGGLIG | 699 |
| 31 | IDNYTDYIYQLIEESQNQQEKNEQELLALDKWASLWNWFDITNWLWYIKIFIMIVGGLIG | 708 |

| | | |
|---|---|---|
| | **LRIVFTVLSIVNRVRQGYSPLSFQTHLPAQRGPDRPEGIGEEGGERDRDRSDPLVNGFLT** | |
| 1 | LRIVFTVLSIVNRVRQGYSPLSFQTHLPAQRGPDRPEGIGEEGGERDRDRSDPLVNGFLA | 744 |
| 2 | LRIVFTVLSIVNRVRQGYSPLSFQTHLPAQRGPDRPEGIGEEGGERDRDRSDPLVNGFLT | 759 |
| 3 | LRIVFTVLSIVNRVRQGYSPLSFQTHLPAQRGPDRPEGIGEEGGERDRDRSDPLVNGFLT | 759 |
| 4 | LRIVFTVLSIVNRVRQGYSPLSFQTHLPAQRGLDRPEGIGEEGGERDRDRSDPLVNGFLT | 749 |
| 5 | LRIVFTVLSIVNRVRQGYSPLSFQTHLPAQRGPDRPEGIGEEGGERDRDRSDPLVNGFLT | 759 |
| 6 | LRIVFTVLSIVNRVRQGYSPLSFQTHLPAQRGPDRPEGIGEEGGERDRDRSDPLVNGFLT | 759 |
| 7 | LRIVFTVLSIVNRVRQGYSPLSFQTHLPAQRGPDRPEGIGEEGGERDRDRSDPLVNGFLT | 767 |
| 8 | LRIVFTVLSIVNRVRQGYSPLSFQTHLPAQRGPDRPEGIGEEGGERDRDRSDPLVNGFLT | 751 |
| 9 | LRIVFTVLSIVNRVRQGYSPLSFQTHLPAQRGPDRPEGIGEEGGERDRDRSDPLVNGFLT | 768 |
| 10 | LRIVFTVLSIVNRVRQGYSPLSFQTHFPAQRGPDRPEGIGEEGGERDRDRSDPLVNGFLT | 761 |
| 11 | LRIVFTVLSIVNRVRQGYSPLSFQTHFPAQRGPDRPEGIGEEGGERDRDRSDPLVNGFLT | 759 |
| 12 | LRIVFTVLSIVNRVRQGYSPLSFQTHFPAQRGPDRPEGIGEEGGERDRDRSDPLVNGFLT | 761 |
| 13 | LRIVFTVLSIVNRVRQGYSPLSFQTHLPAQRGPDRPEGIGEEGGERDRDRSDPLVNGFLT | 761 |
| 14 | LRIVFTVLSIVNRVRQGYSPLSFQTHLPAQRGPDRPEGIGEEGGERDRDRSAPLVNGFLT | 759 |
| 15 | LRIVFTILSIVNRVRQGYSPLSFQTHLPAQREPDRPEGIGEEGGERDRDRSDPLVNGFLT | 758 |
| 16 | LRIVFTVLSIVNRVRQGYSPLSFQTHFPAQRGPDRPEGIGEEGGERDRDRSDPLVNGFLT | 761 |
| 17 | LRIVFTVLSIVNRVRQGYSPLSFQTHLPAQRGPDRPEGIGEEGGERDRDRSDPLVNGFLT | 759 |
| 18 | LRIVFTVLSIVNRVRQGYSPLSFQTHLPAQRGPDRPEGIGEEGGERDRDRSDPLVNGFLT | 759 |
| 19 | LRIVFTVLSIVNRVRQGYSPLSFQTHFPAQRGPDRPEGIGEEGGERDRDRSDPLVNGFLT | 759 |
| 20 | LRIVFTVLSIVNRVRQGYSPLSFQTHLPAQRGPDRPEGIGEEGGERDRDRSDPLVNGFLT | 744 |
| 21 | LRIVFTVLSIVNRVRQGYSPLSFQTHFPAQRGPDRPEEIGEDGGERDRDRSDPLVNGFLT | 761 |
| 22 | LRIVFTVLSIVNRVRQGYSPLSFQTHLPAQRGPDRPEGIGEEGGERDRDRSDPLVNGFLT | 759 |
| 23 | LRIVFTVLSIVNRVRQGYSPLSFQTHFPAQRGPDRPEGIGEEGGERDRDRSDPLVNGFLT | 767 |
| 24 | LRIVFTVLSIVNRVRQGYSPLSFQTHLPAQRGPDRPEGIGEEGGERDRDRSDPLVNGFLT | 759 |
| 25 | LRIVFTVLSIVNRVRQGYSPLSFQTHLPAQRGPDRPEGIGEEGGERDRDRSDPLVNGFLT | 767 |
| 26 | LRIVFTVLSIVNRVRQGYSPLSFQTHFPAQRGPDRPEGIGEEGGERDRDRSDPLVNGFLT | 759 |
| 27 | LRIVFTVLSIVNRVRQGYSPLSFQTHFPAQRGPDRPEGIGEEGGERDRDRSDPLVNGFLT | 759 |
| 28 | LRIVFTVLSVVNRVRQGYSPLSFQTHLPAQREPDRPEGIGEEGGERDRDRSDPLVNGFLT | 761 |
| 29 | LRIVFTVLSIVNRVRQGYSPLSFQTHLPAQRGPDRPEGIGEEGGERDRDRSDPLVNGFLT | 760 |
| 30 | LRIVFTVLSIVNRVRQGYSPLSFQTHLPAQRGPDRPEGIGEEGGERDRDRSDPLVNGFLT | 759 |
| 31 | LRIVFTVLSIVNRVRQGYSPLSFQTHLPAQRGPDRPEGIGEEGGERDRDRSDPLVNGFLT | 768 |

```
    LIWSDLRSLCLFSYHHLRDLLLIVTRIVELLGR-------RGWEVLKYWWNLLQYWSQEL
1   LIWSDLRSLCLFSYHRLRDLLLIVTRIVELLGR-------RGWEVLKYWWNLLQYWSQEL  797
2   LIWSDLRSLCLFSYHRLRDLLLIATRIVELLGR-------RGWEVLKYWWNLLQYWSQEL  812
3   LIWSDLRSLCLFSYHHLRDLLLIVTRIVELLGR-------RGWEVLKYWWNLLQYWSQEL  812
4   LIWSDLRSLCLFSYHHLRDLLLIVTRIVELLGR-------RGWEVLKYWWNLLQYWSQEL  802
5   LIWSDLRSLCLFSYHHLRDLLLIVTRIVELLGR-------RGWEVLKYWWNLLQYWSQEL  812
6   LIWSDLRSLCLFSYHRLRDLLLIATRIVELLGR-------RGWEVLKYWWNLLQYWSQEL  812
7   LIWSDLRSLCLFSYHRLRDLLLIATRIVELLGR-------RGWEVLKYWWNLLQYWSQEL  820
8   LIWSDLRSLCLFSYHHLRDLLLIVTRIVELLGR-------RGWEVLKYWWNLLQYWSQEL  804
9   LIWSDLRSLCLFSYHHLRDLLLIVTRIVELLGR-------RGWEVLKYWWNLLQYWSQEL  821
10  LIWSDLRSLCLFSYHRLRDLLLIATRIVELLGR-------RGWEVLKYWWNLLQYWSQEL  814
11  LIWSDLRSLCLCSYHRLRDLLLIATRIVELLGR-------RGWEVLKYWWNLLQYWSQEL  812
12  LIWSDLRSLCLFSYHRLRDLLLIATRIVELLGR-------RGWEVLKYWWNLLQYWSQEL  814
13  LIWSDLRSLCLFSYHHLRDLLLIVTRIVELLGR-------RGWEVLKYWWNLLQYWSQEL  814
14  LIWSDLRSLCLFSYHHLRDSLLIVTRIVELLGR-------RGWEVLKYWWNLLQYWSQEI  812
15  LIWSDLRSLCLFSYHRLRDLLLIATRIVELLGR-------RGWEVLKYWWNLLQYWSQEL  811
16  LIWSDLRSLCLFSYHHLRDLLLIVTRIVELLGR-------RGWEVLKYWWNLLQYWSQEL  814
17  LIWSDLRSLCLFSYHRLRDLLLIATRIVELLGR-------RGWEVLKYWWNLLQYWSQEL  812
18  LIWSDLRSLCLFSYHRLRDLLLIATRIVELLGR-------RGWEVLKYWWNLLQYWSQEL  812
19  LIWSDLRSLCLFSYHRLRDLLLIATRIVELLGR-------RGWEVLKYWWNLLQYWSQEL  812
20  LIWSDLRSLCLFSYHHLRDLLLIVTRIVELLGR-------RGWEVLKYWWNLLQYWSQEL  797
21  LIWSDLRSLCLFSYHRLRDLLLIATRIVELLGH-------RGWEVLKYWWNLLQYWSQEL  814
22  LIWSDLRSLCLFSYHHLRDLLLIVTRIVELLGR-------RGWEVLKYWWNLLQYWSQEL  812
23  LIWSDLRSLCLFSYHRLRDLLLIATRIVELLGR-------RGWEVLKYWWNLLQYWSQEL  820
24  LIWSDLRSLCLFSYHHLRDLLLIVTRIVELLGR-------RGWEVLKYWWNLLQYWSQEL  812
25  LIWSDLRSLCLFSYHRLRDLLLIATRIVELLGR-------RGWEVLKYWWNLLQYWSQEL  820
26  LIWSDLRSLCLFSYHHLRDLLLIVTRIVELLGR-------RGWEVLKYWWNLLQYWSQEI  812
27  LIWSDLRSLCLFSYHHLRDLLLIVTRIVELLGR-------RGWEVLKYWWNLLQYWSQEL  812
28  LIWSDLRSLCLFSYHHLRDLLLIVTRIVELLGR-------RGWEVLKYWWNLLQYWSQEI  814
29  LIWSDLRSLCLFSYHRLRDLLLIATRIVELLGR-------RGWEVLKYWWNLLQYWSQEL  813
30  LIWSDLRSLCLFSYHHLRDLLLIVTRIVELLGR-------RGWEVLKYWWNLLQYWSQEL  812
31  LIWSDLRSLCLFSYHHLRDLLLIVTRIVELLGR-------RGWEVLKYWWNLLQYWSQEL  821
```

KNSAVSLLNATAIAVAEGTDRVIEVVQRAGRAIRHIPRRIRQGLERALL

| | | |
|---|---|---|
| 1 | KNSAVSLLNATAIAVAEGTDRVIEVVQRACRAILHIPRRIRQGLERALL | 846 |
| 2 | KKSAVSLLNATAIAVAEGTDRVIEVVQRAGRAIRHIPRRIRQGLERALL | 861 |
| 3 | KNSAISLLNATAIAVAEGTDRVIEVVQRAGRAIRHIPRRIRQGLERALL | 861 |
| 4 | KNSAVSLFNATAIAVAEGTDRVIEVVQRAGRAIRHIPRRIRQGLERALL | 851 |
| 5 | KDSAVSLLNATAIAVAEGTDRVIEVVQRAGRAIRHIPRRIRQGLERALL | 861 |
| 6 | KNSAVSLLNATAIAVAEGTDRVIEVVQRAGRAIRHIPRRIRQGLERALL | 861 |
| 7 | KNSAVSLLNATAIAVAEGTDRVIEVVQRAGRAIRHIPRRIRQGLERALL | 869 |
| 8 | KKSAVSLLNATAIAVAEGTDRVIEVVQRAGRAIRHIPRRIRQGLERALL | 853 |
| 9 | KNSAVSLLNATAIAVAEGTDRVIEVVQRAGRAIRHIPRRIRQGLERALL | 870 |
| 10 | KSSAVSLLNATAIAVAEGTDRVIEVVQRAGRAIRHIPRRIRQGLERALL | 863 |
| 11 | KSSAVSLLNATAIAVAEGTDRVIEVVQRAGRAIRHIPRRIRQGLERALL | 861 |
| 12 | KSSAVSLLNATAIAVAEGTDRVIEVVQRAGRAIRHIPRRIRQGLERALL | 863 |
| 13 | KNSAVSLLNATAIAVAEGTDRVIEVVQRAGRAIRHIPRRIRQGLERALL | 863 |
| 14 | KNSAVSLLNATATAVAEGTDRVIEVVQRAGRAIRHIPRRIRQGLERALL | 861 |
| 15 | KDSAVSLLNATATAVAEGTDRVIEVVQRAGRAIRHIPRRIRQGLERALL | 860 |
| 16 | KNSAVSLLNATAIAVAEGTDRVIEVVQRAGRAIRHIPRRIRQGLERALL | 863 |
| 17 | KNSAVSLFNATAIAVAEGTDRVIEVVQRAGRAIRHIPRRIRQGLERALL | 861 |
| 18 | KNSAVSLLNATAIAVAEGTDRVIEVVQRAGRAIRHIPRRIRQGLERALL | 861 |
| 19 | KNSAGSLLNATAIAVAEGTDRVIEVVQRAGRAIRHIPRRIRQGLERALL | 861 |
| 20 | KNSAVSLLNATAIAVAEGTDRVIEVVQRAGRAIRHIPRRIRQGLERALL | 846 |
| 21 | KKSAVSLLNATAIAVAEGTDRVIEVVQRAGRAIRHIPRRIRQGLERALL | 863 |
| 22 | KNSAVSLLNATAIAVAEGTDRVIEVVQRAGRAIRHIPRRIRQGLERVLL | 861 |
| 23 | RNSAVSLLNATAIAVAEGTDRVIEVVQRAGRAIRHIPRRIRQGLERALL | 869 |
| 24 | KDSAVSLLNATAIAVAEGTDRVIEVVQRAGRAIRHIPRRIRQGLERALL | 861 |
| 25 | KNSAVSLLNATAIAVAEGTDRVIEVVQRAGRAIRHIPRRIRQGLERALL | 869 |
| 26 | KNSAISLLNATAIAVAEGTDRVIEVVQRAGRAIRHIPRRIRQGLERALL | 861 |
| 27 | KNSAVNLLNATAIAVAEGTDRVIEVVQRAGRAIRHIPRRIRQGLERALL | 861 |
| 28 | KNSAVSLLNATAIAVAEGTDRVIEVVQRAGRAIRHIPRRIRQGLERALL | 863 |
| 29 | KSSAVSLLNATAIAVAEGTDRVIEVVQRAGRAIRHIPRRIRQGLERALL | 862 |
| 30 | KNSAVNLLNATAIAIAEGTDRVIEVVQRAGRAIRHIPRRIRQGLERALL | 861 |
| 31 | KNSAVSLLNATAIAVAEGTDRVIEVVQRAGRAIRHIPRRIRQGLERALL | 870 |

### Aligned using an external algorithm

**Consensus**

1. B.US.2006.700010040_C9_4520.EU289193
2. B.US.2010.CH0040_3_d1597_ipe032_15_28.MG900512
3. B.US.2010.CH0040_3_d1597_ipe032_15_29.MG900513
4. B.US.2010.CH0040_3_d1597_ipe032_15_34.MG900514
5. B.US.2010.CH0040_3_d1597_ipe032_15_38.MG900515
6. B.US.2010.CH0040_3_d1597_ipe032_15_40.MG900516
7. B.US.2010.CH0040_3_d1597_ipe032_15_42.MG900517
8. B.US.2010.CH0040_3_d1597_ipe032_15_43.MG900518
9. B.US.2010.CH0040_3_d1597_ipe032_15_44.MG900519
10. B.US.2010.CH0040_3_d1597_ipe032_15_46.MG900520
11. B.US.2010.CH0040_3_d1597_ipe032_15_49.MG900521
12. B.US.2010.CH0040_3_d1597_ipe032_15_50.MG900522
13. B.US.2010.CH0040_3_d1597_ipe032_15_52.MG900523
14. B.US.2010.CH0040_3_d1597_ipe032_15_53.MG900524
15. B.US.2010.CH0040_3_d1597_ipe032_15_54.MG900525
16. B.US.2010.CH0040_3_d1597_ipe032_15_55.MG900526
17. B.US.2010.CH0040_3_d1597_ipe032_15_56.MG900527
18. B.US.2010.CH0040_3_d1597_ipe032_15_57.MG900528
19. B.US.2010.CH0040_3_d1597_ipe032_15_60.MG900529
20. B.US.2010.CH0040_3_d1597_ipe032_15_65.MG900530
21. B.US.2010.CH0040_3_d1597_ipe032_15_66.MG900531
22. B.US.2010.CH0040_3_d1597_ipe032_15_68.MG900532
23. B.US.2010.CH0040_3_d1597_ipe032_15_69.MG900533
24. B.US.2010.CH0040_3_d1597_ipe032_15_70.MG900534
25. B.US.2010.CH0040_3_d1597_ipe032_25_10.MG900535
26. B.US.2010.CH0040_3_d1597_ipe032_25_13.MG900536
27. B.US.2010.CH0040_3_d1597_ipe032_25_21.MG900537
28. B.US.2010.CH0040_3_d1597_ipe032_25_22.MG900538
29. B.US.2010.CH0040_3_d1597_ipe032_25_26.MG900539
30. B.US.2010.CH0040_3_d1597_ipe032_25_27.MG900540
31. B.US.2010.CH0040_3_d1597_ipe032_25_4.MG900541
32. B.US.2010.CH0040_3_d1597_ipe032_25_5.MG900542
33. B.US.2010.CH0040_3_d1597_ipe032_25_6.MG900543
34. B.US.2010.CH0040_3_d1597_ipe032_25_8.MG900544
35. B.US.2010.CH0040_3_d1597_ipe032_27_01.MG900545

| | | |
|---|---|---|
| | **MRVMGIRKNYQHLWREGILLLGILMICSAADNLWVTVYYGVPVWREATTTLFCASDAKAY** | |
| 1 | MRVMGIRKNYQHLWREGILLLGILMICSAADNLWVTVYYGVPVWREATTTLFCASDAKAY | 60 |
| 2 | MRVMGIRKNYQHWWREGILLLGILMICSAADKLWVTVYYGVPVWREATTTLFCASDAKAY | 60 |
| 3 | MRVMGIRKNYQHLWREGILLLGILMICSAADNLWVTVYYGVPVWREATTTLFCASDAKAY | 60 |
| 4 | MRVMGIRKNYQHLWREGILLLGILMICSAADNLWVTVYYGVPVWREATTTLFCASDAKAY | 60 |
| 5 | MRVMGIRKNYQHWWREGILLLGILMICSAADKLWVTVYYGVPVWREATTTLFCASDAKAY | 60 |
| 6 | MRVMGIRKNYQHLWREGILLLGILMICSAADNLWVTVYYGVPVWREATTTLFCASDAKAY | 60 |
| 7 | MRVMGIRKNYQHLWREGILLLGILMICSAADNLWVTVYYGVPVWREATTTLFCASDAKAY | 60 |
| 8 | MRVMGIRKNYQHLWREGILLLGILMICSAADNLWVTVYYGVPVWREATTTLFCASDAKAY | 60 |
| 9 | MRVMGIRKNYQHLWREGILLLGILMICSAADNLWVTVYYGVPVWREATTTLFCASDAKAY | 60 |
| 10 | MRVMGIRKNYQHLWREGILLLGILMICSAADNLWVTVYYGVPVWREATTTLFCASDAKAY | 60 |
| 11 | MRVMGIRKNYQHWWREGILLLGILMICSAADKLWVTVYYGVPVWREATTTLFCASDAKAY | 60 |
| 12 | MRVMGIRKNYQHWWREGILLLGILMICSAADKLWVTVYYGVPVWREATTTLFCASDAKAY | 60 |
| 13 | MRVMGIRKNYQHLWREGILLLGILMICSAADNLWVTVYYGVPVWREATTTLFCASDAKAY | 60 |
| 14 | MRVMGIRKNYQHLWREGILLLGILMICSATDNLWVTVYYGVPVWREATTTLFCASDAKAY | 60 |
| 15 | MRVMGIRKNYQHLWREGILLLGILMICSAADDLWVTVYYGVPVWREATTTLFCASDAKAY | 60 |
| 16 | MRVMGIRKNYQHLWRKGILLLGILMICSATNNLWVTVYYGVPVWREATTTLFCASDAKAY | 60 |
| 17 | MRVMGIRKNYQHLWREGILLLGILMICSAADNLWVTVYYGVPVWREATTTLFCASDAKAY | 60 |
| 18 | MRVMGIRKNYQHWWREGILLLGILMICSAADKLWVTVYYGVPVWREATTTLFCASDAKAY | 60 |
| 19 | MRVMGIRKNYQHLWREGILLLGILMICSAADNLWVTVYYGVPVWREATTTLFCASDAKAY | 60 |
| 20 | MRVMGIRKNYQHLWREGILLLGILMICSAADNLWVTVYYGVPVWREATTTLFCASDAKAY | 60 |
| 21 | MRVMGIRKNYQHLWRKGILLLGILMICSATNNLWVTVYYGVPVWREATTTLFCASDAKAY | 60 |
| 22 | MRVMGIRKNYQHLWREGILLLGILMICSAADNLWVTVYYGVPVWREATTTLFCASDAKAY | 60 |
| 23 | MRVMGIRKNYQHLWREGILLLGILMICSAADNLWVTVYYGVPVWREATTTLFCASDAKAY | 60 |
| 24 | MRVMGIRKNYQHLWREGILLLGILMICSAADNLWVTVYYGVPVWREATTTLFCASDAKAY | 60 |
| 25 | MRVMGIRKNYQHLWRKGILLLGILMICSAADKLWVTVYYGVPVWREATTTLFCASDAKAY | 60 |
| 26 | MRVMGIRKNYQHLWREGILLLGILMICSATDNLWVTVYYGVPVWREATTTLFCASDAKAY | 60 |
| 27 | MRVMGIRKNYQHLWREGILLLGILMICSAADNLWVTVYYGVPVWREATTTLFCASDAKAY | 60 |
| 28 | MRVMGIRKNYQHLWREGILLLGILMICSAADNLWVTVYYGVPVWREATTTLFCASDAKAY | 60 |
| 29 | MRVMGIRKNYQHLWREGILLLGILMICSAADNLWVTVYYGVPVWREATTTLFCASDAKAY | 60 |
| 30 | MRVMGIRKNYQHLWREGILLLGILMICSAADNLWVTVYYGVPVWREATTTLFCASDAKAY | 60 |
| 31 | MRVMGIRKNYQHLWREGILLLGILMICSAADNLWVTVYYGVPVWREATTTLFCASDAKAY | 60 |
| 32 | MRVMGIRKNYQHLWREGILLLGILMICSAADNLWVTVYYGVPVWREATTTLFCASDAKAY | 60 |
| 33 | MRVMGIRKNYQHLWREGILLLGILMICSAADDLWVTVYYGVPVWREATTTLFCASDAKAY | 60 |
| 34 | MRVMGIRKNYQHLWREGILLLGILMICSAADNLWVTVYYGVPVWREATTTLFCASDAKAY | 60 |
| 35 | MRVMGIRKNYQHLWRGGILLLGILMICSAADNLWVTVYYGVPVWREATTTLFCASDAKAY | 60 |

| | | |
|---|---|---|
| | **DTEAHNVWATHACVPTDPNPQEVELKNVTENFNMWENNMVEQMHEDIISLWDQSLKPCVK** | |
| 1 | DTEAHNVWATHACVPTDPNPQEVELKNVTENFNMWENNMVEQMHEDIISLWDQSLKPCVK | 120 |
| 2 | DTEAHNVWATHACVPTDPNPQEVE--NVTENFNMWENNMVEQMHEDIISLWDQSLKPCVK | 118 |
| 3 | DTEAHNVWATHACVPTDPNPQEVELKNVTENFNMWENNMVEQMHEDIISLWDQSLKPCVK | 120 |
| 4 | DTEAHNVWATHACVPTDPNPQEVELKNVTENFNMWENNMVEQMHEDIISLWDQSLKPCVK | 120 |
| 5 | DTEAHNVWATHACVPTDPNPQEVELKNVTENFNMWENNMVEQMHEDIISLWDQSLKPCVK | 120 |
| 6 | DTEAHNVWATHACVPTDPNPQEVELKNVTENFNMWENNMVEQMHEDIISLWDQSLKPCVK | 120 |
| 7 | DTEAHNVWATHACVPTDPNPQEVELKNVTENFNMWENNMVEQMHEDIISLWDQSLKPCVK | 120 |
| 8 | DTEAHNVWATHACVPTDPNPQEVELKNVTENFNMWENNMVEQMHEDIISLWDQSLKPCVK | 120 |
| 9 | DTEAHNVWATHACVPTDPNPQEVELKNVTENFNMWENNMVEQMHEDIISLWDQSLKPCVK | 120 |
| 10 | DTEAHNVWATHACVPTDPNPQEVELKNVTENFNMWENNMVEQMHEDIISLWDQSLKPCVK | 120 |
| 11 | DTEAHNVWATHACVPTDPNPQEVELKNVTENFNMWENNMVEQMHEDIISLWDQSLKPCVK | 120 |
| 12 | DTEAHNVWATHACVPTDPNPQEVELKNVTENFNMWENNMVEQMHEDIISLWDQSLKPCVK | 120 |
| 13 | DTEAHNVWATHACVPTDPNPQEVELKNVTENFNMWENNMVEQMHEDIISLWDQSLKPCVK | 120 |
| 14 | DTEAHNVWATHACVPTDPNPQEVELKNVTENFNMWENNMVEQMHEDIISLWDQSLKPCVK | 120 |
| 15 | DTEAHNVWATHACVPTDPNPQEVELKNVTENFNMWENNMVEQMHEDIISLWDQSLKPCVK | 120 |
| 16 | DTEAHNVWATHACVPTDPNPQEVELKNVTENFNMWENNMVEQMHEDIISLWDQSLKPCVK | 120 |
| 17 | DTEAHNVWATHACVPTDPNPQEVELKNVTENFNMWENNMVEQMHEDIISLWDQSLKPCVK | 120 |
| 18 | DTEAHNVWATHACVPTDPNPQEVELKNVTENFNMWENNMVEQMHEDIISLWDQSLKPCVK | 120 |
| 19 | DTEAHNVWATHACVPTDPNPQEVELKNVTENFNMWENNMVEQMHEDIISLWDQSLKPCVK | 120 |
| 20 | DTEAHNVWATHACVPTDPNPQEVELKNVTENFNMWENNMVEQMHEDIISLWDQSLKPCVK | 120 |
| 21 | DTEAHNVWATHACVPTDPNPQEVELKNVTENFNMWENNMVEQMHEDIISLWDQSLKPCVK | 120 |
| 22 | DTEAHNVWATHACVPTDPNPQEVELKNVTENFNMWENNMVEQMHEDIISLWDQSLKPCVK | 120 |
| 23 | DTEAHNVWATHACVPTDPNPQEVELKNVTENFNMWENNMVEQMHEDIISLWEQSLKPCVK | 120 |
| 24 | DTEAHNVWATHACVPTDPNPQEVELKNVTENFNMWENNMVEQMHEDIISLWDQSLKPCVK | 120 |
| 25 | DTEAHNVWATHACVPTDPNPQEVELKNVTENFNMWENNMVEQMHEDIISLWDQSLKPCVK | 120 |
| 26 | DTEAHNVWATHACVPTDPNPQEVELKNVTENFNMWENNMVEQMHEDIISLWDQSLKPCVK | 120 |
| 27 | DTEAHNVWATHACVPTDPNPQEVELKNVTENFNMWENNMVEQMHEDIISLWDQSLKPCVK | 120 |
| 28 | DTEAHNVWATHACVPTDPNPQEVELKNVTENFNMWENNMVEQMHEDIISLWDQSLKPCVK | 120 |
| 29 | DTEAHNVWATHACVPTDPNPQEVELKNVTENFNMWENNMVEQMHEDIISLWDQSLKPCVK | 120 |
| 30 | DTEAHNVWATHACVPTDPNPQEVELKNVTENFNMWENNMVEQMHEDIISLWDQSLKPCVK | 120 |
| 31 | DTEAHNVWATHACVPTDPNPQEVELKNVTENFNMGKNNMVEQMHEDIISLWDQSLKPCVK | 120 |
| 32 | DTEAHNVWATHACVPTDPNPQEVELKNVTENFNMWENNMVEQMHEDIISLWDQSLKPCVK | 120 |
| 33 | DTEAHNVWATHACVPTDPNPQEVELKNVTENFNMWENNMVEQMHEDIISLWDQSLKPCVK | 120 |
| 34 | DTEAHNVWATHACVPTDPNPQEVELKNVTENFNMWENNMVEQMHEDIISLWDQSLKPCVK | 120 |
| 35 | DTEAHNVWATHACVPTDPNPQEVELKNVTENFNMGKNNMVEQMHEDIISLWDQSLKPCVK | 120 |

```
    LTPLCVTLNCTDNVTTTTNRNGTNCTDSGNVTNTTNRNGTLMGNETLMMEKGEVKNCSFR
1   LTPLCVTLNCTDLGNVTNTT----------------------NSNGEMMEKGEVKNCSFK   158
2   LTPLCVTLNCTDNVTTTTNRNGTNCTDSGNVTNTTNRNGTLMGNETLMMEKGEVKNCSFR   178
3   LTPLCVTLNCTDNVTTTTNRNGTNCTDSGNVTNTTNRNGTLMGNETLMMEKGEVKNCSFR   180
4   LTPLCVTLNCTDNVTTTTNRNGTNCTDSGNVTNTTNRNGTLMGNETLMMEKGEVKNCSFR   180
5   LTPLCVTLNCTDNVTTTTNRNGTNCTDSGNVTNTTNRNGTLMGNETLMMEKGEVKNCSFR   180
6   LTPLCVTLNCTDNVTTTTNRNGTNCTDSGNVTNTTNRNGTLMGNETLMMEKGEVKNCSFR   180
7   LTPLCVTLNCTDNVTTTTNRNGTNCTDSGNVTNTTNRNGTLMGNETLMMEKGEVKNCSFR   180
8   LTPLCVTLNCTDNVTTTTNRNGTNCTDSGNVTNTTNRNGTLMGNETLMMEKGEVKNCSFR   180
9   LTPLCVTLNCTDNVTTTTNRNGTNCTDSGNVTNTTNRNGTLMGNETLMMEKGEVKNCSFR   180
10  LTPLCVTLNCTDNVTTTTNRNGTNCTDSGNVTNTTNRNGTLMGNETLMMEKGEVKNCSFR   180
11  LTPLCVTLNCTDNVTTTTNRNGTNCTDSGNVTNTTNRNGTLMGNETLMMEKGEVKNCSFR   180
12  LTPLCVTLNCTDNVTNTTNRNGTL-----------------MGNETLMMEKGEVKNCSFR   163
13  LTPLCVTLNCTDNVTTTTNRNGTNCTDSGNVTNTTNRNGTLMGNETLMMEKGEVKNCSFR   180
14  LTPLCVTLNCTDNVTTTTNRNGTNCTDSGNVTNTTNRNGTLMGNETLMMEKGEVKNCSFR   180
15  LTPLCVTLNCTDNVTTTTNRNGTNCTDSGNVTNTTNRNGTLMGNETLMMEKGEVKNCSFR   180
16  LTPLCVTLNCTDNVTTTTNRNGTNCTDSGNVTNTTNRNGTLMGNETLMMEKGEVKNCSFR   180
17  LTPLCVTLNCTDNVTTTTNRNGTNCTDSGNVTNTTNRNGTLMGNETLMMEKGEVKNCSFR   180
18  LTPLCVTLNCTDNVTTTTNRNGTNCTDSGNVTNT-------TNRNGALMEKGEVKNCSFR   173
19  LTPLCVTLNCTDNVTTTTNRNGTNCTDSGNVTNTTNRNGTLMGNETLMMEKGEVKNCSFR   180
20  LTPLCVTLNCTDNVTTTTNRNGTNCTDSGNVTNTTNRNGTLMGNETLMMEKGEVKNCSFR   180
21  LTPLCVTLNCTDNVTTTTNRNGTNCTDSGNVTNTTNRNGTLMGNETLMMEKGEVKNCSFR   180
22  LTPLCVTLNCTDNVTTTTNRNGTNCTDSGNVTNTTNRNGTLMGNETLMMEKGEVKNCSFR   180
23  LTPLCVTLNCTDNVTTTTNRNGTNCTDSGNVTNTTNRNGTLMGNETLMMEKGEVKNCSFR   180
24  LTPLCVTLNCTDNVTTTTNRNGTNCTDSGNVTNTTNRNGTLMGNETLMMEKGEVKNCSFR   180
25  LTPLCVTLNCTDNVTTTTNRNGTNCTDSGNVTNTTNRNGTLMGNETLMMEKGEVKNCSFR   180
26  LTPLCVTLNCTDNVTTTTNRNGTNCTDSGNVTNTTNRNGTLMGNETLMMEKGEVKNCSFR   180
27  LTPLCVTLNCTDNVTTTTNRNGTNCTDSGNVTNTTNRNGTLMGNETLMMEKGEVKNCSFR   180
28  LTPLCVTLNCTDNVTTTTNRNGTNCTDSGNVTNTTNRNGTLMGNETLMMEKGEVKNCSFR   180
29  LTPLCVTLNCTDNVTTTTNRNGTNCTDSGNVTNTTNRNGTLMGNETLMMEKGEVKNCSFR   180
30  LTPLCVTLNCTDNVTTTTNRNGTNCTDSGNVTNTTNRNGTLMGNETLMMEKGEVKNCSFR   180
31  LTPLCVTLNCTDNVTTTTNRNGTNYTDSGNVTNT-------TNRNGALMEKGEVKNCSFR   173
32  LTPLCVTLNCTDNVTTTTNRNGTNCTDSGNVTNTTNRNGTLMGNETLMMEKGEVKNCSFR   180
33  LTPLCVTLNCTDNVTTTTNRNGTNCTDSGNVTNTTNRNGTLMGNETLMMEKGEVKNCSFR   180
34  LTPLCVTLNCTDNVTTTTNRNGTNCTDSGNVTNTTNRNGTLMGNETLMMEKGEVKNCSFR   180
35  LTPLCVTLNCTDLGNVTTTTNRNGTNCTDSGNVTAT-----TNRNETFMEKEEV-NCSFR   174
```

| | Sequence | |
|---|---|---|
|  | ITTDMQDRTRKEYALFYKLDVVPIND-------TSYRLVSCNTSVITQACPKVSFEPIPI |  |
| 1 | ITTDIKDRTRKEYALFYKLDVVPIND-------TRYRLVSCNTSVITQACPKVSFEPIPI | 211 |
| 2 | ITTDMQDRTRKEYALFYKLDVVPIND-------TRYRLVSCNTSVITQACPKVSFEPIPI | 231 |
| 3 | ITTDMQDRTRKEYALFYKLDVVPIND-------TSYRLVSCNTSVITQACPKVSFEPIPI | 233 |
| 4 | ITTDMQDRTRKEYALFYKLDVVPIND-------TSYRLVSCNTSVITQACPKVSFEPIPI | 233 |
| 5 | ITTDMQDRTRKEYALFYKLDVVPIND-------TSYRLVSCNTSVITQACPKVSFEPIPI | 233 |
| 6 | ITTDMQDRTRKEYALFYKLDVVPIND-------TSYRLVSCNTSVITQACPKVSFEPIPI | 233 |
| 7 | ITTDMQDRTRKEYALFYKLDVVPIND-------TSYRLVSCNTSVITQACPKVSFEPIPI | 233 |
| 8 | ITTDMQDRTRKEYALFYKLDVVPIND-------TSYRLVSCNTSVITQACPKVSFEPIPI | 233 |
| 9 | ITTDMQDRTRKEYALFYKLDVVPIND-------TSYRLVSCNTSVITQACPKVSFEPIPI | 233 |
| 10 | ITTDMQDRTRKEYALFYKLDVVPIND-------TSYRLVSCNTSVITQACPKVSFEPIPI | 233 |
| 11 | ITTDMQDRTRKEYALFYKLDVVPIND-------TSYRLVSCNTSVITQACPKVSFEPIPI | 233 |
| 12 | ITTDMQDRTRKEYALFYKLDVVPIND-------TSYRLVSCNTSVITQACPKVSFEPIPI | 216 |
| 13 | ITTDMQDRTRKEYALFYKLDVVPIND-------TSYRLVSCNTSVITQACPKVSFEPIPI | 233 |
| 14 | ITTDMQDRTRKEYALFYKLDVVPIND-------TSYRLVSCNTSVITQACPKVSFEPIPI | 233 |
| 15 | ITTDMQDRTRKEYALFYKLDVVPIND-------TSYRLVSCNTSVITQACPKVSFEPIPI | 233 |
| 16 | ITTDMQDRTRKEYALFYKLDVVPIND-------TSYRLVSCNTSVITQACPKVSFEPIPI | 233 |
| 17 | ITTDMQDRTRKEYALFYKLDVVPIND-------TSYRLVSCNTSVITQACPKVSFEPIPI | 233 |
| 18 | ITTDMQDRTRKEYALFYKLDVVPIND-------TRYRLVSCNTSVITQACPKVSFEPIPI | 226 |
| 19 | ITTDMQDRTRKEYALFYKLDVVPIND-------TSYRLVSCNTSVITQACPKVSFEPIPI | 233 |
| 20 | ITTDMQDRTRKEYALFYKLDVVPIND-------TSYRLVSCNTSVITQACPKVSFEPIPI | 233 |
| 21 | ITTDMQDRTRKEYALFYKLDVVPIND-------TSYRLVSCNTSVITQACPKVSFEPIPI | 233 |
| 22 | ITTDMQDRTRKEYALFYKLDVVPIND-------TSYRLVSCNTSVITQACPKVSFEPIPI | 233 |
| 23 | ITTDMQDRTRKEYALFYKLDVVPIND-------TSYRLVSCNTSVIAQACPKVSFEPIPI | 233 |
| 24 | ITTDMQDRTRKEYALFYKLDVVPIND-------TSYRLVSCNTSVITQACPKVSFEPIPI | 233 |
| 25 | ITTDMQDRTRKEYALFYKLDVVPIND-------TSYRLVSCNTSVITQACPKVSFEPIPI | 233 |
| 26 | ITTDMQDRTRKEYALFYKLDVVPIND-------TSYRLVSCNTSVITQACPKVSFEPIPI | 233 |
| 27 | ITTDMQDRTRKEYALFYKLDVVPIND-------TSYRLVSCNTSVITQACPKVSFEPIPI | 233 |
| 28 | ITTDMQDRTRKEYALFYKLDVVPIND-------TSYRLVSCNTSVITQACPKVSFEPIPI | 233 |
| 29 | ITTDMQDRTRKEYALFYKLDVVPIND-------TSYRLVSCNTSVITQACPKVSFEPIPI | 233 |
| 30 | ITTDMQDRTRKEYALFYKLDVVPIND-------TSYRLVSCNTSVITQACPKVSFEPIPI | 233 |
| 31 | ITTDMQDRTRKEYALFYKLDVVPIND-------TRYRLVSCNTSVITQACPKVSFEPIPI | 226 |
| 32 | ITTDMQDRTRKEYALFYKLDVVPIND-------TSYRLVSCNTSVITQACPKVSFEPIPI | 233 |
| 33 | ITTDMQDRTRKEYALFYKLDVVPIND-------TSYRLVSCNTSVITQACPKVSFEPIPI | 233 |
| 34 | ITTDMQDRTRKEYALFYKLDVVPIND-------TSYRLVSCNTSVITQACPKVSFEPIPI | 233 |
| 35 | ITTDMQDRTRKEYALFYKLDVVPIND-------TRYRLVSCNTSVITQACPKVSFEPIPI | 227 |

| | Sequence | |
|---|---|---|
| | HYCAPAGFAILKCNDKQFNGTGPCTNVSTVQCTHGIRPVVSTQLLLNGSLAEEEVVIRSV | |
| 1 | HYCAPAGFAILKCNDKQFIGTGPCTNVSTVQCTHGIRPVVSTQLLLNGSLAEEEVVIRSV | 271 |
| 2 | HYCAPAGFAILKCNDKQFNGTGPCTNVSTVQCTHGIRPVVSTQLLLNGSLAEEEVVIRSV | 291 |
| 3 | HYCAPAGFAILKCNDKQFNGTGPCTNVSTVQCTHGIRPVVSTQLLLNGSLAEEEVVIRSV | 293 |
| 4 | HYCAPAGFAILKCNDKQFNGTGPCTNVSTVQCTHGIRPVVSTQLLLNGSLAEEEVVIRSV | 293 |
| 5 | HYCAPAGFAILKCNDKQFNGTGPCTNVSTVQCTHGIRPVVSTQLLLNGSLAEEEVVIRSV | 293 |
| 6 | HYCAPAGFAILKCNDKQFNGTGPCTNVSTVQCTHGIRPVVSTQLLLNGSLAEEEVVIRSV | 293 |
| 7 | HYCAPAGFAILKCNDKQFNGTGPCTNVSTVQCTHGIRPVVSTQLLLNGSLAEEEVVIRSV | 293 |
| 8 | HYCAPAGFAILKCNDKQFNGTGPCTNVSTVQCTHGIRPVVSTQLLLNGSLAEEEVVIRSV | 293 |
| 9 | HYCAPAGFAILKCNDKQFNGTGPCTNVSTVQCTHGIRPVVSTQLLLNGSLAEEEVVIRSV | 293 |
| 10 | HYCAPAGFAILKCNDKQFNGTGPCTNVSTVQCTHGIRPVVSTQLLLNGSLAEEEVVIRSV | 293 |
| 11 | HYCAPAGFAILKCNDKQFNGTGPCTNVSTVQCTHGIRPVVATQLLLNGSLAEEEVVIRSV | 293 |
| 12 | HYCAPAGFAILKCNDKQFNGTGPCTNVSTVQCTHGIRPVVSTQLLLNGSLAEEEVIIRSV | 276 |
| 13 | HYCAPAGFAILKCNDKQFNGTGPCTNVSTVQCTHGIRPVVSTQLLLNGSLAEEEVVIRSV | 293 |
| 14 | HYCAPAGFAILKCNDKQFNGTGPCTNVSTVQCTHGIRPVVSTQLLLNGSLAEEEVVIRSV | 293 |
| 15 | HYCAPAGFAILKCNDKQFNGTGPCTNVSTVQCTHGIRPVVSTQLLLNGSLAEEEVVIRSV | 293 |
| 16 | HYCAPAGFAILKCNDKQFNGTGPCTNVSTVQCTHGIRPVVSTQLLLNGSLAEEEVVIRSV | 293 |
| 17 | HYCAPAGFAILKCNDKQFNGTGPCTNVSTVQCTHGIRPVVSTQLLLNGSLAEEEVVIRSV | 293 |
| 18 | HYCAPAGFAILKCNDKQFNGTGPCTNVSTVQCTHGIRPVVSTQLLLNGSLAEEEVVIRSV | 286 |
| 19 | HYCAPAGFAILKCNDKQFNGTGPCTNVSTVQCTHGIRPVVSTQLLLNGSLAEEEVVIRSV | 293 |
| 20 | HYCAPAGFAILKCNDKQFNGTGPCTNVSTVQCTHGIRPVVSTQLLLNGSLAEEEVVIRSV | 293 |
| 21 | HYCAPAGFAILKCNDKQFNGTGPCTNVSTVQCTHGIRPVVSTQLLLNGSLAEEEVVIRSV | 293 |
| 22 | HYCAPAGFAILKCNDKQFNGTGPCTNVSTVQCTHGIRPVVSTQLLLNGSLAEEEVVIRSV | 293 |
| 23 | HYCAPAGFAILKCNDKQFNGTGPCTNVSTVQCTHGIRPVVSTQLLLNGSLAEEEVVIRSV | 293 |
| 24 | HYCAPAGFAILKCNDKQFNGTGPCTNVSTVQCTHGIRPVVSTQLLLNGSLAEEEVVIRSV | 293 |
| 25 | HYCAPAGFAILKCNDKQFNGTGPCTNVSTVQCTHGIRPVVSTQLLLNGSLAEEEVVIRSV | 293 |
| 26 | HYCAPAGFAILKCNDKQFNGTGPCTNVSTVQCTHGIRPVVSTQLLLNGSLAEEEVVIRSV | 293 |
| 27 | HYCAPAGFAILKCNDKQFNGTGPCTNVSTVQCTHGIRPVVSTQLLLNGSLAEEEVVIRSV | 293 |
| 28 | HYCAPAGFAILKCNDKQFNGTGPCTNVSTVQCTHGIRPVVSTQLLLNGSLAEEEVVIRSV | 293 |
| 29 | HYCAPAGFAILKCNDKQFNGTGPCTNVSTVQCTHGIRPVVSTQLLLNGSLAEEEVVIRSV | 293 |
| 30 | HYCAPAGFAILKCNDKQFNGTGPCTNVSTVQCTHGIRPVVSTQLLLNGSLAEEEVVIRSV | 293 |
| 31 | HYCAPAGFAILKCNDKQFNGTGPCTNVSTVQCTHGIRPVVSTQLLLNGSLAEEEVVIRSV | 286 |
| 32 | HYCAPAGFAILKCNDKQFNGTGPCTNVSTVQCTHGIRPVVSTQLLLNGSLAEEEVVIRSV | 293 |
| 33 | HYCAPAGFAILKCNDKQFNGTGPCTNVSTVQCTHGIRPVVSTQLLLNGSLAEEEVVIRSV | 293 |
| 34 | HYCAPAGFAILKCNDKQFNGTGPCTNVSTVQCTHGIRPVVSTQLLLNGSLAEEEVVIRSV | 293 |
| 35 | HYCAPAGFAILKCNDKQFNGTGPCTNVSTVQCTHGIRPVVSTQLLLNGSLAEEEVVIRSV | 287 |

| | Sequence | |
|---|---|---|
| | **NFSDNAKTIIVQLNKSVEINCTRPNNNTRKSIHMGPGKAFYARGGIIGDIRKAHCNISGT** | |
| 1 | NFSDNAKTIIVQLNKSVEITCTRPNNNTRKSIPMGPGKAFYARGDITGDIRKAYCEINGT | 331 |
| 2 | NFSDNAKTIIVQLNKSVEINCTRPNNNTRKSIHMGPGKAFYARGAIIGDIRKAHCNISGT | 351 |
| 3 | NFSDNAKTIIVQLNKSVEINCTRPNNNTRKSIHMGPGKAFYARGGIIGDIRKAHCNISGT | 353 |
| 4 | NFSDNAKTIIVQLNKSVEINCTRPNNNTRKSIHMGPGKAFYARGGIIGDIRKAHCNISGT | 353 |
| 5 | NFSDNAKTIIVQLNKSVEINCTRPNNNTRKSIHMGPGKAFYARGGIIGDIRKAHCNISGT | 353 |
| 6 | NFSDNAKTIIVQLNKSVEINCTRPNNNTRKSIHMGPGKAFYARGGIIGDIRKAHCNISGT | 353 |
| 7 | NFSDNAKTIIVQLNKSVEINCTRPNNNTRKSIHMGPGKAFYARGGIIGDIRKAHCNISGT | 353 |
| 8 | NFSDNAKTIIVQLNKSVEINCTRPNNNTRKSIHMGPGKAFYARGGIIGDIRKAHCNISGT | 353 |
| 9 | NFSDNAKTIIVQLNKSVEINCTRPNNNTRKSIHMGPGKAFYARGGIIGDIRKAHCNISGT | 353 |
| 10 | NFSDNAKTIIVQLNKSVEINCTRPNNNTRKSIHMGPGKAFYARGGIIGDIRKAHCNISGT | 353 |
| 11 | NFSDNAKTIIVQLNKSVEINCTRPNNNTRKSIHMGPGKAFYARGAIIGDIRKAHCNISRA | 353 |
| 12 | NFSDNAKTIIVQLNKSVEINCTRPNNNTRKSIHMGPGKAFYARGGIIGDIRKAHCNISGT | 336 |
| 13 | NFSDNAKTIIVQLNKSVEINCTRPNNNTRKSIHMGPGKAFYARGAIIGDIRKAHCNISGT | 353 |
| 14 | NFSDNAKTIIVQLNKSVEINCTRPNNNTRKSIHMGPGKAFYARGGIIGDIRKAHCNISGT | 353 |
| 15 | NFSDNAKTIIVQLNKSVEINCTRPNNNTRKSIHMGPGKAFYARGAIIGDIRKAHCNISGT | 353 |
| 16 | NFSDNAKTIIVQLNKSVEINCTRPNNNTRKSIHMGPGKAFYARGAIIGDIRKAHCNISGT | 353 |
| 17 | NFSDNAKTIIVQLNKSVEINCTRPNNNTRKSIHMGPGKAFYARGGIIGDIRKAHCNISGT | 353 |
| 18 | NFSDNAKTIIVQLNKSVEINCTRPNNNTRKSIHMGPGKAFYARGGIIGDIRKAHCNISGT | 346 |
| 19 | NFSDNAKTIIVQLNKSVEINCTRPNNNTRKSIHMGPGKAFYARGGIIGDIRKAHCNISGT | 353 |
| 20 | NFSDNAKTIIVQLNKSVEINCTRPNNNTRKSIHMGPGKAFYARGGIIGDIRKAHCNISGT | 353 |
| 21 | NFSDNAKTIIVQLNKSVEINCTRPNNNTRKSIHMGPGKAFYARGAIIGDIRKAHCNISGT | 353 |
| 22 | NFSDNAKTIIVQLNKSVEINCTRPNNNTRKSIHMGPGKAFYARGAIIGDIRKAHCNISGT | 353 |
| 23 | NFSDNAKTIIVQLNKSVEINCTRPNNNTRKSIHMGPGKAFYARGGIIGDIRKAHCNISGT | 353 |
| 24 | NFSDNAKTIIVQLNKSVEINCTRPNNNTRKSIHMGPGKAFYARGAIIGDIRKAHCNISGT | 353 |
| 25 | NFSDNAKTIIVQLNKSVEINCTRPNNNTRKSIHMGPGKAFYARGGIIGDIRKAHCNISGT | 353 |
| 26 | NFSDNAKTIIVQLNKSVEINCTRPNNNTRKSIHMGPGKAFYARGGIIGDIRKAHCNISGT | 353 |
| 27 | NFSDNAKTIIVQLNKSVEINCTRPNNNTRKSIHMGPGKAFYARGGIIGDIRKAHCNISGT | 353 |
| 28 | NFSDNAKTIIVQLNKSVEINCTRPNNNTRKSIHMGPGKAFYARGGIIGDIRKAHCNISGT | 353 |
| 29 | NFSDNAKTIIVQLNKSVEINCTRPNNNTRKSIHMGPGKAFYARGGIIGDIRKAHCNISGT | 353 |
| 30 | NFSDNAKTIIVQLNKSVEINCTRPNNNTRKSIHMGPGKAFYARGAIIGDIRKAHCNISGT | 353 |
| 31 | NFSDNAKTIIVQLNKSVEINCTRPNNNTRKSIHMGPGKAFYARGGIIGDIRKAHCNISGT | 346 |
| 32 | NFSDNAKTIIVQLNKSVEINCTRPNNNTRKSIHMGPGKAFYARGGIIGDIRKAHCNISGT | 353 |
| 33 | NFSDNAKTIIVQLNKSVEINCTRPNNNTRKSIHMGPGKAFYARGGIIGDIRKAHCNISGT | 353 |
| 34 | NFSDNAKTIIVQLNKSVEINCTRPNNNTRKSIHMGPGKAFYARGAIIGDIRKAHCNISGT | 353 |
| 35 | NFSDNAKTIIVQLNKSVEINCTRPNNNTRKSIHMGPGKAFYARGGIIGDIRKAHCNISGT | 347 |

| | | |
|---|---|---|
| | EWHNTLKLVVEKLREQY-NKTIVFNHSSGGDPEIVMYSFNCGGEFFYCNSTKLFNSTWPG | |
| 1 | EWHSTLKLVVEKLREQY-NKTIVFNRSSGGDPEIVMYSFNCGGEFFYCNSTKLFNSTWPW | 390 |
| 2 | EWHNTLKLVVEKLREQY-NKTIVFNHSSGGDPEIVMYSFNCGGEFFYCNSTKLFNSTWPG | 410 |
| 3 | EWHNTLKLVVEKLREQY-NKTIVFNHSSGGDPEIVMYSFNCGGEFFYCNSTKLFNSTWPG | 412 |
| 4 | EWHNTLKLVVAKLREQY-NKTIVFNHSSGGDPEIVMYSFNCGGEFFYCNSTKLFNSTWPG | 412 |
| 5 | EWHNTLKLVVEKLREQY-NKTIVFNHSSGGDPEIVMYSFNCGGEFFYCNSTKLFNSTWPG | 412 |
| 6 | EWHNTLKLVVEKLREQY-NKTIVFNHSSGGDPEIVMYSFNCGGEFFYCNSTKLFNSTWPG | 412 |
| 7 | EWHNTLKLVVEKLREQY-NKTIVFNHSSGGDPEIVMYSFNCGGEFFYCNSTKLFNSTWPG | 412 |
| 8 | EWHNTLKLVVKKLREQY-NKTIVFNHSSGGDPEIVMYSFNCGGEFFYCNSTKLFNSTWPG | 412 |
| 9 | EWHNTLKLVVEKLREQY-NKTIVFNHSSGGDPEIVMYSFNCGGEFFYCNSTKLFNSTWPG | 412 |
| 10 | EWHNTLKLVVEKLREQY-NKTIVFNHSSGGDPEIVMYSFNCGGEFFYCNSTKLFNSTWPG | 412 |
| 11 | EWHSTLKLVAEKLREQY-NKTIVFNHSSGGDPEIVMYSFNCGGEFFYCNSTKLFNSTWSW | 412 |
| 12 | EWHNTLKLVVEKLREQY-NKTIVFNHSSGGDPEIVMYSFNCGGEFFYCNSTKLFNSTWPG | 395 |
| 13 | EWHNTLKLVVEKLREQY-NKTIVFNHSSGGDPEIVMYSFNCGGEFFYCNSTKLFNSTWPG | 412 |
| 14 | EWHNTLKLVVEKLREQY-NKTIVFNHSSGGDPEIVMYSFNCGGEFFYCNSTKLFNSTWPW | 412 |
| 15 | EWHNTLKLVVEKLREQY-NKTIVFNHSSGGDPEIVMYSFNCGGEFFYCNSTKLFNSTWPG | 412 |
| 16 | EWHNTLKLVVEKLREQY-NKTIVFNHSSGGDPEIVMYSFNCGGEFFYCNSTKLFNSTWPR | 412 |
| 17 | EWHNTLKLVVEKLREQY-NKTIVFNHSSGGDPEIVMYSFNCGGEFFYCNSTKLFNSTWPG | 412 |
| 18 | EWHSTLKLVVKKLREQY-NKTIVFNHSSGGDPEIVMYSFNCGGEFFYCNSTKLFNSTWPW | 405 |
| 19 | EWHNTLKLVVEKLREQY-NKTIVFNHSSGGDPEIVMYSFNCGGEFFYCNSTKLFNSTWPG | 412 |
| 20 | EWHNTLKLVVEKLREQY-NKTIVFNHSSGGDPEIVMYSFNCGGEFFYCNSTKLFNSTWPG | 412 |
| 21 | EWHNTLKLVVEKLREQY-NKTIVFNHSSGGDPEIVMYSFNCGGEFFYCNSTKLFNSTWPR | 412 |
| 22 | EWHNTLKLVVEKLREQY-NKTIVFNHSSGGDPEIVMYSFNCGGEFFYCNSTKLFNSTWPG | 412 |
| 23 | EWHNTLKLVVEKLREQY-NKTIVFNHSSGGDPEIVMYSFNCGGEFFYCNSTKLFNSTWPG | 412 |
| 24 | EWHNTLKLVVEKLREQY-NKTIVFNHSSGGDPEIVMYSFNCGGEFFYCNSTKLFNSTWPG | 412 |
| 25 | EWHNTLKLVVEKLREQY-NKTIVFNHSSGGDPEIVMYSFNCGGEFFYCNSTKLFNSTWPR | 412 |
| 26 | EWHNTLKLVVEKLREQY-NKTIVFNHSSGGDPEIVMYSFNCGGEFFYCNSTKLFNSTWPG | 412 |
| 27 | EWHNTLKLVVEKLREQY-NKTIVFNHSSGGDPEIVMYSFNCGGEFFYCNSTKLFNSTWPG | 412 |
| 28 | EWHSTLKLVVKKLREQY-NKTIVFNHSSGGDPEIVMYSFNCGGEFFYCNSTKLFNSTWPG | 412 |
| 29 | EWHNTLKLVVEKLREQY-NKTIVFNHSSGGDPEIVMYSFNCGGEFFYCNSTKLFNSTWPG | 412 |
| 30 | EWHNTLKLVVEKLREQY-NKTIVFNHSSGGDPEIVMYSFNCGGEFFYCNSTKLFNSTWPG | 412 |
| 31 | EWHNTLKLVVEKLREQY-NKTIVFNHSSGGDPEIVMYSFNCGGEFFYCNSTKLFNSTWPG | 405 |
| 32 | EWHNTLKLVVEKLREQY-NKTIVFNHSSGGDPEIVMYSFNCGGEFFYCNSTKLFNSTWPG | 412 |
| 33 | EWHNTLKLVVEKLREQY-NKTIVFNHSSGGDPEIVMYSFNCGGEFFYCNSTKLFNSTWPG | 412 |
| 34 | EWHNTLKLVVEKLREQY-NKTIVFNHSSGGDPEIVMYSFNCGGEFFYCNSTKLFNSTWPG | 412 |
| 35 | EWHNTLKLVVEKLREQY-NKTIVFNHSSGGDPEIVMYSFNCGGEFFYCNSTKLFNSTWPG | 406 |

```
    ND--TEGSHNTNGTLTLPCRIKQIINMWQGVGKAMYAPPIEGIIKCSSNITGLLLTRDGG
1   ND--TKGSHDTNGTLILPCKIKQIINMWQGVGKAMYAPPIEGKIRCSSNITGLLLTRDGG  448
2   ND--TEGSHNTNGTLTLPCRIKQIINMWQEVGKAMYAPPIEGIIKCSSNITGLLLTRDGG  468
3   ND--TEGSHDTNGTLTLPCRIKQIINMWQEVGKAMYAPPIEGIIKCSSNITGLLLTRDGG  470
4   ND--TEGSHNTNGTLTLPCRIKQIINMWQGVGKAMYAPPIEGIIKCSSNITGLLLTRDGG  470
5   ND--TEGSHNTNGTLTLPCRIKQIINMWQGVGKAMYAPPIEGIIKCSSNITGLLLTRDGG  470
6   ND--TEGSHNTNGTLTLPCRIKQIINMWQGVGKAMYAPPIEGIIKCSSNITGLLLTRDGG  470
7   ND--TEGSHNTNGTLTLPCRIKQIINMWQGVGKAMYAPPIEGIIKCSSNITGLLLTRDGG  470
8   ND--TEGSHNTNGTLTLPCRIKQIINMWQEVGKAMYAPPIEGIIKCSSNITGLLLTRDGG  470
9   ND--TEGSHNTNGTLTLPCRIKQIINMWQGVGKAMYAPPIEGIIKCSSNITGLLLTRDGG  470
10  ND--TEGSHNTNGTLTLPCRIKQIINMWQGVGKAMYAPPIEGIIKCSSNITGLLLTRDGG  470
11  ND--TQGSHDTNGTLTLPCRIKQIINMWQEVGKAMYAPPIEGIIKCSSNITGLLLTRDGG  470
12  ND--TEGSHNTNGTLTLPCRIKQIINMWQGVGKAMYAPPIEGIIKCSSNITGLLLTRDGG  453
13  ND--TEGSHNTNGTLTLPCRIKQIINMWQEVGKAMYAPPIEGIIKCSSNITGLLLTRDGG  470
14  NND-TEGSHDTNGTLTLPCRIKQIINMWQEVGKAMYAPPIEGIIKCSSNITGLLLTRDGG  471
15  ND--TEGSHNTNGTLTLPCRIKQIINMWQGVGKAMYAPPIEGIIKCSSNITGLLLTRDGG  470
16  NA--TRGSPDTNGTLTLPCRIKQIINMWQEVGKAMYAPPIEGIIKCSSNITGLLLTRDGG  470
17  ND--TEGSHNTNGALTLPCRIKQIINMWQGVGKAMYAPPIEGIIKCSSNITGLLLTRDGG  470
18  NDTDTEGSHDTNGTLTLPCRIKQIINMWQEVGKAMYAPPIEGIIKCSSNITGLLLTRDGG  465
19  ND--TEGSHNTNGTLTLPCRIKQIINMWQGVGKAMYAPPIEGIIKCSSNITGLLLTRDGG  470
20  ND--TEGSHNTNGTLTLPCRIKQIINMWQGVGKAMYAPPIEGIIKCSSNITGLLLTRDGG  470
21  NA--TRGSPDTNGTLTLPCRIKQIINMWQEVGKAMYAPPIEGIIKCSSNITGLLLTRDGG  470
22  NA--TEGSHDTNGTLTLPCRIKQIINMWQEVGKAMYAPPIEGIIKCSSNITGLLLTRDGG  470
23  ND--TEGSHNTNGTLTLPCRIKQIINMWQGVGKAMYAPPIEGIIKCSSNITGLLLTRDGG  470
24  ND--TEGSHNTNGTLTLPCRIKQIINMWQEVGKAMYAPPIEGIIKCSSNITGLLLTRDGG  470
25  ND--TEGSHNTNGTLTLPCRIKQIINMWQEVGKAMYAPPIEGIIKCSSNITGLLLTRDGG  470
26  ND--TEGSHNTNGTLTLPCRIKQIINMWQGVGKAMYAPPIEGIIKCSSNITGLLLTRDGG  470
27  ND--TEGSHNTNGTLTLPCRIKQIINMWQGVGKAMYAPPIEGIIKCSSNITGLLLTRDGG  470
28  ND--TEGSHNTNGTLTLPCRIKQIINMWQGVGKAMYAPPIEGIIKCSSNITGLLLTRDGG  470
29  ND--TEGSHNTNGTLTLPCRIKQIINMWQGVGKAMYAPPIEGIIKCSSNITGLLLTRDGG  470
30  ND--TEGSHNTNGTLTLPCRIKQIINMWQEVGKAMYAPPIEGIIKCSSNITGLLLTRDGG  470
31  ND--TEGSHNTNGTLTLPCRIKQIINMWQGVGKAMYAPPIEGIIKCSSNITGLLLTRDGG  463
32  ND--TEGSHNTNGTLTLPCRIKQIINMWQGVGKAMYAPPIEGIIKCSSNITGLLLTRDGG  470
33  ND--TEGSHNTNGTLTLPCRIKQIINMWQGVGKAMYAPPIEGIIKCSSNITGLLLTRDGG  470
34  ND--TEGSHNTNGTLTLPCRIKQIINMWQEVGKAMYAPPIEGIIKCSSNITGLLLTRDGG  470
35  ND--TEGSHNTNGTLTLPCRIKQIINMWQEVGKAMYAPPIEGIIKC---------TRDGG  455
```

| | | |
|---|---|---|
| | -YESNKTDEIFRPGGGDMRDNWRSELYKYKVVKIEPLGVAPTKAKRRVVQREKRAFG-LG | |
| 1 | -YESNETDEIFRPGGGDMRDNWRSELYKYKVVKIEPLGVAPTKAKRRVVQREKRAFG-LG | 506 |
| 2 | -YESNKTDEIFRPGGGDMRDNWRSELYKYKVVKIEPLGVAPTKAKRRVVQREKRAFG-LG | 526 |
| 3 | -YESNKTDEIFRPGGGDMRDNWRSELYKYKVVKIEPLGVAPTKAKRRVVQREKRAFG-LG | 528 |
| 4 | -YESNKTDEIFRPGGGDMRDNWRSELYKYKVVKIEPLGVAPTKAKRRVVQREKRAFG-LG | 528 |
| 5 | -YESNKTDEIFRPGGGDMRDNWRSELYKYKVVKIEPLGVAPTKAKRRVVQREKRAFG-LG | 528 |
| 6 | -YESNKTDEIFRPGGGDMRDNWRSELYKYKVVKIEPLGVAPTKAKRRVVQREKRAFG-LG | 528 |
| 7 | -YESNKTDEIFRPGGGDMRDNWRSELYKYKVVKIEPLGVAPTKAKRRVVQREKRAFG-LG | 528 |
| 8 | -YESNKTDEIFRPGGGDMRDNWRSELYKYKVVKIEPLGVAPTKAKRRVVQREKRAFG-LG | 528 |
| 9 | -YESNKTDEIFRPGGGDMRDNWRSELYKYKVVKIEPLGVAPTKAKRRVVQREKRAFG-LG | 528 |
| 10 | -YESNKTDEIFRPGGGDMRDNWRSELYKYKVVKIEPLGIAPTKAKRRVVQREKRAFG-LG | 528 |
| 11 | -YESNKTDEIFRPGGGDMRDNWRSELYKYKVVKIEPLGVAPTKAKRRVVQREKRAFG-LG | 528 |
| 12 | -YESNKTDEIFRPGGGDMRDNWRSELYKYKVVKIEPLGVAPTKAKRRVVQREKRAFG-LG | 511 |
| 13 | -YESNKTDEIFRPGGGDMRDNWRSELYKYKVVKIEPLGVAPTKAKRRVVQREKRAFG-LG | 528 |
| 14 | -YESNKTDEIFRPGGGDMRDNWRSELYKYKVVKIEPLGVAPTKAKRRVVQREKRAFG-LG | 529 |
| 15 | -YESNKTDEIFRPGGGDMRDNWRSELYKYKVVKIEPLGVAPTKAKRRVVQREKRAFG-LG | 528 |
| 16 | -YESNKTDEIFRPGGGDMRDNWRSELYKYKVVKIEPLGVAPTKAKRRVVQREKRAFG-LG | 528 |
| 17 | -YESNKTDEIFRPGGGDMRDNWRSELYKYKVVKIEPLGVAPTKAKRRVVQREKRAFG-LG | 528 |
| 18 | -YESNKTDEIFRPGGGDMRDNWRSELYKYKVVKIEPLGVAPTKAKRRVVQREKRAFG-LG | 523 |
| 19 | -YESNKTDEIFRPGGGDMRDNWRSELYKYKVVKIEPLGVAPTKAKRRVVQREKRAFG-LG | 528 |
| 20 | -YESNKTDEIFRPGGGDMRDNWRSELYKYKVVKIEPLGVAPTKAKRRVVQREKRAFG-LG | 528 |
| 21 | -YESNKTDEIFRPGGGDMRDNWRSELYKYKVVKIEPLGVAPTKAKRRVVQREKRAFG-LG | 528 |
| 22 | -YESNKTDEIFRPGGGDMRDNWRSELYKYKVVKIEPLGVAPTKAKRRVVQREKRAFG-LG | 528 |
| 23 | -YESNKTDEIFRPGGGDMRDNWRSELYKYKVVKIEPLGVAPTKAKRRVVQREKRAFG-LG | 528 |
| 24 | -YESNKTDEIFRPGGGDMRDNWRSELYKYKVVKIEPLGVAPTKAKRRVVQREKRAFG-LG | 528 |
| 25 | -YESNKTDEIFRPGGGDMRDNWRSELYKYKVVKIEPLGVAPTKAKRRVVQREKRAFG-LG | 528 |
| 26 | -YESNKTDEIFRPGGGDMRDNWRSELYKYKVVKIEPLGVAPTKAKRRVVQREKRAFG-LG | 528 |
| 27 | -YESNKTDEIFRPGGGDMRDNWRSELYKYKVVKIEPLGVAPTKAKRRVVQREKRAFG-LG | 528 |
| 28 | -YESNKTDEIFRPGGGDMRDNWRSELYKYKVVKIEPLGVAPTKAKRRVVQREKRAFG-LG | 528 |
| 29 | -YESNKTDEIFRPGGGDMRDNWRSELYKYKVVKIEPLGVAPTKAKRRVVQREKRAFG-LG | 528 |
| 30 | -YESNKTDEIFRPGGGDMRDNWRSELYKYKVVKIEPLGVAPTKAKRRVVQREKRAFG-LG | 528 |
| 31 | -YESNKTDEIFRPGGGDMRDNWRSELYKYKVVKIEPLGVAPTKAKRRVVQREKRAFG-LG | 521 |
| 32 | -YESNKTDEIFRPGGGDMRDNWRSELYKYKVVKIEPLGVAPTKAKRRVVQREKRAFG-LG | 528 |
| 33 | -YESNKTDEIFRPGGGDMRDNWRSELYKYKVVKIEPLGVAPTKAKRRVVQREKRAFG-LG | 528 |
| 34 | -YESNKTDEIFRPGGGDMRDNWRSELYKYKVVKIEPLGVAPTKAKRRVVQREKRAFG-LG | 528 |
| 35 | -YESNKTDEIFRPGGGDMRDNWRSELYKYKVVKIEPLGVAPTKAKRRVVQREKRAFG-LG | 513 |

**AVFLGFLGAAGSTMGAASITLTVQARQLLSGIVQQQNNLLRAIEAQQHLLQLTVWGIKQL**

| | | |
|---|---|---|
| 1 | AVFLGFLGAAGSTMGAASITLTVQARQLLSGIVQQQNNLLRAIEAQQHLLQLTVWGIKQL | 566 |
| 2 | AVFLGFLGAAGSTMGAASITLTVQARQLLSGIVQQQNNLLRAIEAQQHLLQLTVWGIKQL | 586 |
| 3 | AVFLGFLGAAGSTMGAASITLTVQARQLLSGIVQQQNNLLRAIEAQQHLLQLTVWGIKQL | 588 |
| 4 | AVFLGFLGAAGSTMGAASITLTVQARQLLSGIVQQQNNLLRAIEAQQHLLQLTVWGIKQL | 588 |
| 5 | AVFLGFLGAAGSTMGAASITLTVQARQLLSGIVQQQNNLLRAIEAQQHLLQLTVWGIKQL | 588 |
| 6 | AVFLGFLGAAGSTMGAASITLTVQARQLLSGIVQQQNNLLRAIEAQQHLLQLTVWGIKQL | 588 |
| 7 | AVFLGFLGAAGSTMGAASITLTVQARQLLSGIVQQQNNLLRAIEAQQHLLQLTVWGIKQL | 588 |
| 8 | AVFLGFLGAAGSTMGAASITLTVQARQLLSGIVQQQNNLLRAIEAQQHLLQLTVWGIKQL | 588 |
| 9 | AVFLGFLGAAGSTMGAASITLTVQARQLLSGIVQQQNNLLRAIEAQQHLLQLTVWGIKQL | 588 |
| 10 | AVFLGFLGAAGSTMGAASITLTVQARQLLSGIVQQQNNLLRAIEAQQHLLQLTVWGIKQL | 588 |
| 11 | AVFLGFLGAAGSTMGAASITLTVQARQLLSGIVQQQNNLLRAIEAQQHLLQLTVWGIKQL | 588 |
| 12 | AVFLGFLGAAGSTMGAASITLTVQARQLLSGIVQQQNNLLRAIEAQQHLLQLTVWGIKQL | 571 |
| 13 | AVFLGFLGAAGSTMGAASITLTVQARQLLSGIVQQQNNLLRAIEAQQHLLQLTVWGIKQL | 588 |
| 14 | AVFLGFLGAAGSTMGAASITLTVQARQLLSGIVQQQNNLLRAIEAQQHLLQLTVWGIKQL | 589 |
| 15 | AVFLGFLGAAGSTMGAASITLTVQARQLLSGIVQQQNNLLRAIEAQQHLLQLTVWGIKQL | 588 |
| 16 | AVFLGFLGAAGSTMGAASITLTVQARQLLSGIVQQQNNLLRAIEAQQHLLQLTVWGIKQL | 588 |
| 17 | AVFLGFLGAAGSTMGAASITLTVQARQLLSGIVQQQNNLLRAIEAQQHLLQLTVWGIKQL | 588 |
| 18 | AVFLGFLGAAGSTMGAASITLTVQARQLLSGIVQQQNNLLRAIEAQQHLLQLTVWGIKQL | 583 |
| 19 | AVFLGFLGAAGSTMGAASITLTVQARQLLSGIVQQQNNLLRAIEAQQHLLQLTVWGIKQL | 588 |
| 20 | AVFLGFLGAAGSTMGAASITLTVQARQLLSGIVQQQNNLLRAIEAQQHLLQLTVWGIKQL | 588 |
| 21 | AVFLGFLGAAGSTMGAASITLTVQARQLLSGIVQQQNNLLRAIEAQQHLLQLTVWGIKQL | 588 |
| 22 | AVFLGFLGAAGSTMGAASITLTVQARQLLSGIVQQQNNLLRAIEAQQHLLQLTVWGIKQL | 588 |
| 23 | AVFLGFLGAAGSTMGAASITLTVQARQLLSGIVQQQNNLLRAIEAQQHLLQLTVWGIKQL | 588 |
| 24 | AVFLGFLGAAGSTMGAASITLTVQARQLLSGIVQQQNNLLRAIEAQQHLLQLTVWGIKQL | 588 |
| 25 | AVFLGFLGAAGSTMGAASITLTVQARQLLSGIVQQQNNLLRAIEAQQHLLQLTVWGIKQL | 588 |
| 26 | AVFLGFLGAAGSTMGAASITLTVQARQLLSGIVQQQNNLLRAIEAQQHLLQLTVWGIKQL | 588 |
| 27 | AVFLGFLGAAGSTMGAASITLTVQARQLLSGIVQQQNNLLRAIEAQQHLLQLTVWGIKQL | 588 |
| 28 | AVFLGFLGAAGSTMGAASITLTVQARQLLSGIVQQQNNLLRAIEAQQHLLQLTVWGIKQL | 588 |
| 29 | AVFLGFLGAAGSTMGAASITLTVQARQLLSGIVQQQNNLLRAIEAQQHLLQLTVWGIKQL | 588 |
| 30 | AVFLGFLGAAGSTMGAASITLTVQARQLLSGIVQQQNNLLRAIEAQQHLLQLTVWGIKQL | 588 |
| 31 | AVFLGFLGAAGSTMGAASITLTVQARQLLSGIVQQQNNLLRAIEAQQHLLQLTVWGIKQL | 581 |
| 32 | AVFLGFLGAAGSTMGAASITLTVQARQLLSGIVQQQNNLLRAIEAQQHLLQLTVWGIKQL | 588 |
| 33 | AVFLGFLGAAGSTMGAASITLTVQARQLLSGIVQQQNNLLRAIEAQQHLLQLTVWGIKQL | 588 |
| 34 | AVFLGFLGAAGSTMGAASITLTVQARQLLSGIVQQQNNLLRAIEAQQHLLQLTVWGIKQL | 588 |
| 35 | AVFLGFLGAAGSTMGAASITLTVQARQLLSGIVQQQNNLLRAIEAQQHLLQLTVWGIKQL | 573 |

| | Sequence | |
|---|---|---|
| | QARVLAVERYLKDQQLLGIWGCSGKLICTTTVPWNTSWSNKSLEQIWDNMTWMEWEREIN | |
| 1 | QARVLAVERYLKDQQLLGIWGCSGKLICTTTVPWNTSWSNKSLEQIWDNMTWMEWEREID | 626 |
| 2 | QARVLAVERYLKDQQLLGIWGCSGKLICTTTVPWNTSWSNKSLEQIWDNMTWMEWEREIN | 646 |
| 3 | QARVLAVERYLKDQQLLGIWGCSGKLICTTTVPWNTSWSNKSLEQIWDNMTWMEWEREIN | 648 |
| 4 | QARVLAVERYLKDQQLLGIWGCSGKLICTTTVPWNTSWSNKSLEQIWDNMTWMEWEREIN | 648 |
| 5 | QARVLAVERYLKDQQLLGIWGCSGKLICTTTVPWNTSWSNKSLEQIWDNMTWMEWEREIN | 648 |
| 6 | QARVLAVERYLKDQQLLGIWGCSGKLICTTTVPWNTSWSNKSLEQIWDNMTWMEWEREIN | 648 |
| 7 | QARVLAVERYLKDQQLLGIWGCSGKLICTTTVPWNTSWSNKSLEQIWDNMTWMEWEREIN | 648 |
| 8 | QARVLAVERYLKDQQLLGIWGCSGKLICTTTVPWNTSWSNKSLEQIWDNMTWMEWEREIN | 648 |
| 9 | QARVLAVERYLKDQQLLGIWGCSGKLICTTTVPWNTSWSNKSLEQIWDNMTWMEWEREIN | 648 |
| 10 | QARVLAVERYLKDQQLLGIWGCSGKLICTTTVPWNTSWSNKSLEQIWDNMTWMEWEREIN | 648 |
| 11 | QARVLAVERYLKDQQLLGIWGCSGKLICTTTVPWNTSWSNKSLEQIWDNMTWMEWEREIN | 648 |
| 12 | QARVLAVERYLKDQQLLGIWGCSGKLICTTTVPWNTSWSNKSLEQIWDNMTWMEWEREIN | 631 |
| 13 | QARVLAVERYLKDQQLLGIWGCSGKLICTTTVPWNTSWSNKSLEQIWDNMTWMEWEREIN | 648 |
| 14 | QARVLAVERYLKDQQLLGIWGCSGKLICTTTVPWNTSWSNKSLEQIWDKMTWMEWEREID | 649 |
| 15 | QARVLAVERYLKDQQLLGIWGCSGKLICTTTVPWNTSWSNKSLEQIWDNMTWMEWEREIN | 648 |
| 16 | QARVLAVERYLKDQQLLGIWGCSGKLICTTTVPWNTSWSNKSLEQIWDNMTWMEWEREIN | 648 |
| 17 | QARVLAVERYLKDQQLLGIWGCSGKLICTTTVPWNTSWSNKSLEQIWDNMTWMEWEREIN | 648 |
| 18 | QARVLAVERYLKDQQLLGIWGCSGKLICTTTVPWNTSWSNKSLEQIWDNMTWMEWEREIN | 643 |
| 19 | QARVLAVERYLKDQQLLGIWGCSGKLICTTTVPWNTSWSNKSLEQIWDNMTWMEWEREIN | 648 |
| 20 | QARVLAVERYLKDQQLLGIWGCSGKLICTTTVPWNTSWSNKSLEQIWDNMTWMEWEREIN | 648 |
| 21 | QARVLAVERYLKDQQLLGIWGCSGKLICTTTVPWNTSWSNKSLEQIWDNMTWMEWEREIN | 648 |
| 22 | QARVLAVERYLKDQQLLGIWGCSGKLICTTTVPWNTSWSNKSLEQIWDNMTWMEWEREIN | 648 |
| 23 | QARVLAVERYLKDQQLLGIWGCSGKLICTTTVPWNTSWSNKSLEQIWDNMTWMEWEREIN | 648 |
| 24 | QARVLAVERYLKDQQLLGIWGCSGKLICTTTVPWNTSWSNKSLEQIWDNMTWMEWEREIN | 648 |
| 25 | QARVLAVERYLKDQQLLGIWGCSGKLICTTTVPWNTSWSNKSLEQIWDNMTWMEWEREIN | 648 |
| 26 | QARVLAVERYLKDQQLLGIWGCSGKLICTTTVPWNTSWSNKSLEQIWDKMTWMEWEREID | 648 |
| 27 | QARVLAVERYLKDQQLLGIWGCSGKLICTTTVPWNTSWSNKSLEQIWDNMTWMEWEREIN | 648 |
| 28 | QARVLAVERYLKDQQLLGIWGCSGKLICTTTVPWNTSWSNKSLEQIWDNMTWMEWEREIN | 648 |
| 29 | QARVLAVERYLKDQQLLGIWGCSGKLICTTTVPWNTSWSNKSLEQIWDNMTWMEWEREIN | 648 |
| 30 | QARVLAVERYLKDQQLLGIWGCSGKLICTTTVPWNTSWSNKSLEQIWDNMTWMEWEREIN | 648 |
| 31 | QARVLAVERYLKDQQLLGIWGCSGKLICTTTVPWNTSWSNKSLEQIWDNMTWMEWEREIN | 641 |
| 32 | QARVLAVERYLKDQQLLGIWGCSGKLICTTTVPWNTSWSNKSLEQIWDNMTWMEWEREIN | 648 |
| 33 | QARVLAVERYLKDQQLLGIWGCSGKLICTTTVPWNTSWSNKSLEQIWDNMTWMEWEREIN | 648 |
| 34 | QARVLAVERYLKDQQLLGIWGCSGKLICTTTVPWNTSWSNKSLEQIWDNMTWMEWEREIN | 648 |
| 35 | QARVLAVERYLKDQQLLGIWGCSGKLICTTTVPWNTSWSNKSLEQIWDNMTWIEWEREIN | 633 |

| | Sequence | |
|---|---|---|
| | NYTGYIYQLIEESQNQQEKNEQELLAWDKWASLWNWFDITNWLWYIKIFIMIVGGLIGLR | |
| 1 | NYTGYIYQLIEESQNQQEKNEQELLALDKWASLWNWFDITNWLWYIKIFIMIVGGLIGLR | 686 |
| 2 | NYTGYIYQLIEESQNQQEQNEQELLAWDKWASLWNWFDITNWLWYIKIFIMIVGGLIGLR | 706 |
| 3 | NYTGYIYQLIEESQNQQEKNEQELLALDKWASLWNWFDITNWLWYIKIFIMIVGGLIGLR | 708 |
| 4 | NYTGYIYQLIEESQNQQEKNEQELLAWDKWASLWNWFDITNWLWYIKIFIMIVGGLIGLR | 708 |
| 5 | NYTGYIYQLIEESQNQQEKNEQELLAWDKWASLWNWFDITNWLWYIKIFIMIVGGLIGLR | 708 |
| 6 | NYTGYIYQLIEESQNQQEKNEQELLAWDKWASLWNWFDITNWLWYIKIFIMIVGGLIGLR | 708 |
| 7 | NYTGYIYQLIEESQNQQEKNEQELLAWDKWASLWNWFDITNWLWYIKIFIMIVGGLIGLR | 708 |
| 8 | NYTGYIYQLIEESQNQQEKNEQELLAWDKWASLWNWFDITNWLWYIKIFIMIVGGLIGLR | 708 |
| 9 | NYTGYIYQLIEESQNQQEKNEQELLAWDKWASLWNWFDITNWLWYIKIFIMIVGGLIGLR | 708 |
| 10 | NYTGYIYQLIEESQNQQEKNEQELLAWDKWASLWNWFDITNWLWYIKIFIMIVGGLIGLR | 708 |
| 11 | NYTGYIYQLIEESQNQQEKNEQELLALDKWASLWNWFDITNWLWYIKIFIMIVGGLIGLR | 708 |
| 12 | NYTGYIYQLIEESQNQQEKNEQELLAWDKWASLWNWFDITNWLWYIKIFIMIVGGLIGLR | 691 |
| 13 | NYTGYIYQLIEESQNQQEKNEQELLAWDKWAGLWNWFDITNWLWYIKIFIMIVGGLIGLR | 708 |
| 14 | NYTDYIYQLIEESQNQQEKNEQELLALDKWASLWNWFDITNWLWYIKIFIMIVGGLIGLR | 709 |
| 15 | NYTGYIYQLIEESQNQQEKNEQELLAWDKWASLWNWFDITNWLWYIKIFIMIVGGLIGLR | 708 |
| 16 | NYTGYIYQLIEESQNQQEKNEQELLAWDKWASLWNWFDITNWLWYIKIFIMIVGGLIGLR | 708 |
| 17 | NYTGYIYQLIEESQNQQEKNEQELLAWDKWASLWNWFDITNWLWYIKIFIMIVGGLIGLR | 708 |
| 18 | NYTGYIYQLIEESQNQQEKNEQELLAWDKWASLWNWFDITNWLWYIKIFIMIVGGLIGLR | 703 |
| 19 | NYTGYIYQLIEESQNQQEKNEQELLAWDKWASLWNWFDITNWLWYIKIFIMIVGGLIGLR | 708 |
| 20 | NYTGYIYQLIEESQNQQEKNEQELLAWDKWASLWNWFDITNWLWYIKIFIMIVGGLIGLR | 708 |
| 21 | NYTGYIYQLIEESQNQQEKNEQELLAWDKWASLWNWFDITNWLWYIKIFIMIVGGLIGLR | 708 |
| 22 | NYTGYIYQLIEESQNQQEKNEQELLAWDKWASLWNWFDITNWLWYIKIFIMIVGGLIGLR | 708 |
| 23 | NYTGYIYQLIEESQNQQEKNEQELLAWDKWASLWNWFDITNWLWYIKIFIMIVGGLIGLR | 708 |
| 24 | NYTGYIYQLIEESQNQQEKNEQELLAWDKWASLWNWFDITNWLWYIKIFIMIVGGLIGLR | 708 |
| 25 | NYTGYIYQLIEESQNQQEKNEQELLAWDKWASLWNWFDITNWLWYIKIFIMIVGGLIGLR | 708 |
| 26 | NYTDYIYQLIEESQNQQEKNEQELLAWDKWASLWNWFDITNWLWYIKIFIMIVGGLIGLR | 708 |
| 27 | NYTGYIYQLIEESQNQQEKNEQELLAWDKWASLWNWFDITNWLWYIKIFIMIVGGLIGLR | 708 |
| 28 | NYTGYIYQLIEESQNQQEKNEQELLAWDKWASLWNWFDITNWLWYIKIFIMIVGGLIGLR | 708 |
| 29 | NYTGYIYQLIEESQNQQEKNEQELLAWDKWASLWNWFDITNWLWYIKIFIMIVGGLIGLR | 708 |
| 30 | NYTGYIYQLIEESQNQQEKNEQELLAWDKWASLWNWFDITNWLWYIKIFIMIVGGLIGLR | 708 |
| 31 | NYTGYIYQLIEESQNQQEKNEQELLAWDKWASLWNWFDITNWLWYIKIFIMIVGGLIGLR | 701 |
| 32 | NYTGYIYQLIEESQNQQEKNEQELLAWDKWASLWNWFDITNWLWYIKIFIMIVGGLIGLR | 708 |
| 33 | NYTGYIYQLIEESQNQQEKNEQELLAWDKWASLWNWFDITNWLWYIKIFIMIVGGLIGLR | 708 |
| 34 | NYTGYIYQLIEESQNQQEKNEQELLAWDKWASLWNWFDITNWLWYIKIFIMIVGGLIGLR | 708 |
| 35 | NYTGYIYQLIEESQNQQEKNEQELLAWDKWASLWNWFDITNWLWYIKIFIMIVGGLIGLR | 693 |

| | | |
|---|---|---|
| | IVFTVLSIVNRVRQGYSPLSFQTHFPAQRGPDRPEGIGEEGGERDRDRSDPLVNGFLTLI | |
| 1 | IVFTVLSIVNRVRQGYSPLSFQTHLPAQRGPDRPEGIGEEGGERDRDRSDPLVNGFLALI | 746 |
| 2 | IVFTVLSIVNRVRQGYSPLSFQTHFPAQRGPDRPEGIGEEGGERDRDRSDPLVNGFLTLI | 766 |
| 3 | IVFTVLSIVNRVRQGYSPLSFQTHFPAQRGPDRPEGIGEEGGERDRDRSEPLVNGFLTLI | 768 |
| 4 | IVFTVLSIVNRVRQGYSPLSFQTHFPAQRGPDRPEGIGEEGGERDRDRSDPLVNGFLTLI | 768 |
| 5 | IVFTVLSIVNRVRQGYSPLSFQTHFPAQRGPDRPEGIGEEGGERDRDRSDPLVNGFLTLI | 768 |
| 6 | IVFTVLSIVNRVRQGYSPLSFQTHFPAQRGPDRPEGIGEEGGERDRDRSDPLVNGFLTLI | 768 |
| 7 | IVFTVLSIVNRVRQGYSPLSFQTHFPAQRGPDRPEGIGEEGGERDRDRSDPLVNGFLTLI | 768 |
| 8 | IVFTVLSIVNRVRQGYSPLSFQTHFPAQRGPDRPEGIGEEGGERDRDRSDPLVNGFLTLI | 768 |
| 9 | IVFTVLSIVNRVRQGYSPLSFQTHFPAQRGPDRPEGIGEEGGERDRDRSDPLVNGFLTLI | 768 |
| 10 | IVFTVLSIVNRVRQGYSPLSFQTHFPAQRGPDRPEGIGEEGGERDRDRSDPLVNGFLTLI | 768 |
| 11 | IVFTVLSIVNRVRQGYSPLSFQTHFPAQRGPDRPEGIGEEGGERDRDRSDPLVNGFLTLI | 768 |
| 12 | IVFTVLSIVNRVRQGYSPLSFQTHLPAQRGPDRPEGIGEEGGERDRDRSDPLVNGFLTLI | 751 |
| 13 | IVFTVLSIVNRVRQGYSPLSFQTHFPAQRGPDRPEGIGEEGGERDRDRSDPLVNGFLTLI | 768 |
| 14 | IVFTVLSIVNRVRQGYSPLSFQTHFPAQRGPDRPEGIGEEGGERDRDRSDPLVNGFLTLI | 769 |
| 15 | IVFTVLSIVNRVRQGYSPLSFQTHFPAQRGPDRPEGIGEEGGERDRDRSDPLVNGFLTLI | 768 |
| 16 | IVFTVLSIVNRVRQGYSPLSFQTHFPAQRGPDRPEGIGEEGGERDRDRSDPLVNGFLTLI | 768 |
| 17 | IVFTVLSIVNRVRQGYSPLSFQTHFPAQRGPDRPEGIGEEGGERDRDRSDPLVNGFLTLI | 768 |
| 18 | IVFTVLSIVNRVRQGYSPLSFQTHLPAQRGPDRPEGIGEEGGERDRDRSDPLVNGFLTLI | 763 |
| 19 | IVFTVLSIVNRVRQGYSPLSFQTHFPAQRGPDRPEGIGEEGGERDRDRSDPLVNGFLTLI | 768 |
| 20 | IVFTVLSIVNRSRQGYSPLSFQTHFPAQRGPDRPEGIGEEGGERDRDRSDPLVNGFLTLI | 768 |
| 21 | IVFTVLSIVNRVRQGYSPLSFQTHFPAQRGPDRPEGIGEEGGERDRDRSDPLVNGFLTLI | 768 |
| 22 | IVFTVLSIVNRVRQGYSPLSFQTHFPAQRGPDRPEGIGEEGGERDRDRSDPLVNGFLTLI | 768 |
| 23 | IVFTVLSIVNRVRQGYSPLSFQTHFPAQRGPDRPEGIGEEGGERDRDRSDPLVNGFLTLI | 768 |
| 24 | IVFTVLSIVNRVRQGYSPLSFQTHFPAQRGPDRPEGIGEEGGERDRDRSDPLVNGFLTLI | 768 |
| 25 | IVFTVLSIVNRVRQGYSPLSFQTHFPAQRGPDRPEGIGEEGGERDRDRSDPLVNGFLTLI | 768 |
| 26 | IVFTVLSIVNRVRQGYSPLSFQTHFPAQRGPDRPEGIGEEGGERDRDRSDPLVNGFLTLI | 768 |
| 27 | IVFTILSIVNRVRQGYSPLSFQTHLPAQRGPDRPEGIGEEGGERDRDRSDPLVNGFLTLI | 768 |
| 28 | IVFTVLSIVNRVRQGYSPLSFQTHFPAQRGPDRPEGIGEEGGERDRDRSDPLVNGFLTLI | 768 |
| 29 | IVFTVLSIVNRVRQGYSPLSFQTHFPAQRGPDRPEGIGEEGGERDRDRSDPLVNGFLTLI | 768 |
| 30 | IVFTVLSIVNRVRQGYSPLSFQTHFPAQRGPDRPEGIGEEGGERDRDRSDPLVNGFLTLI | 768 |
| 31 | IVFTVLSIVNRVRQGYSPLSFQTHFPAQRGPDRPEGIGEEGGERDRDRSDPLVNGFLTLI | 761 |
| 32 | IVFTVLSIVNRVRQGYSPLSFQTHFPAQRGPDRPEGIGEEGGERDRDRSDPLVNGFLTLI | 768 |
| 33 | IVFTVLSIVNRVRQGYSPLSFQTHFPAQRGPDRPEGIGEEGGERDRDRSDPLVNGFLTLI | 768 |
| 34 | IVFTVLSIVNRVRQGYSPLSFQTHFPAQRGPDRPEGIGEEGGERDRDRSDPLVNGFLTLI | 768 |
| 35 | IVFTVLSIVNRVRQGYSPLSFQTHFPAQRGPDRPEGIGEEGGERDRDRSDPLVNGFLTLI | 753 |

| | Sequence | |
|---|---|---|
|  | WSDLRSLCLFSYHRLRDLLLIATRIVELLGR-------RGWEVLKYWWNLLQYWSQELKK |  |
| 1 | WSDLRSLCLFSYHRLRDLLLIVTRIVELLGR-------RGWEVLKYWWNLLQYWSQELKN | 799 |
| 2 | WSDLRSLCLFSYHHLRDLLLIVTRIVELLGR-------RGWEVLKYWWNLLQYWSQELKD | 819 |
| 3 | WSDLRSLCLFSYHHLRDLLLIVTRIVELLGR-------RGWEVLKYWWNLLQYWSQELKK | 821 |
| 4 | WSDLRSLCLFSYHRLRDLLLIATRIVELLGR-------RGWEVLKYWWNLLQYWSQELKK | 821 |
| 5 | WSDLRSLCLFSYHRLRDLLLIATRIVELLGR-------RGWEVLKYWWNLLQYWSQELKK | 821 |
| 6 | WSDLRSLCLFSYHRLRDLLLIATRIVELLGR-------RGWEVLKYWWNLLQYWSQELKK | 821 |
| 7 | WSDLRSLCLFSYHRLRDLLLIATRIVELLGR-------RGWEVLKYWWNLLQYWSQELKK | 821 |
| 8 | WSDLRSLCLFSYHRLRDLLLIATRIVELLGR-------RGWEVLKYWWNLLQYWSQELKK | 821 |
| 9 | WSDLRSLCLFSYHRLRDLLLIATRIVELLGR-------RGWEVLKYWWNLLQYWSQELKK | 821 |
| 10 | WSDLRSLCLFSYHRLRDLLLIATRIVELLGR-------RGWEVLKYWWNLLQYWSQELKK | 821 |
| 11 | WSDLRSLCLFSYHHLRDLLLIVTRIVELLGR-------RGWEVLKYWWNLLQYWSQELKK | 821 |
| 12 | WSDLRSLCLFSYHRLRDLLLIATRIVELLGR-------RGWEVLKYWWNLLQYWSQELKK | 804 |
| 13 | WSDLRSLCLFSYHHLRDLLLIVTRIVELLGR-------RGWEVLKYWWNLLQYWGQELKN | 821 |
| 14 | WSDLRSLCLFSYHRLRDLLLIATRIVELLGR-------RGWEVLKYWWNLLQYWSQELKK | 822 |
| 15 | WSDLRSLCLFSYHRLRDLLLIATRIVELLGR-------RGWEVLKYWWNLLQYWSQELKK | 821 |
| 16 | WSDLRSLCLFSYHRLRDLLLIATRIVELLGR-------RGWEILKYWWNLLQYWSQELKK | 821 |
| 17 | WSDLRSLCLFSYHRLRDLLLIATRIVELLGR-------RGWEVLKYWWNLLQYWSQELKK | 821 |
| 18 | WSDLRSLCLFSYHHLRDLLLIVTRIVELLGR-------RGWEVLKYWWNLLQYWSQELKD | 816 |
| 19 | WSDLRSLCLFSYHRLRDLLLIATRIVELLGR-------RGWEVLKYWWNLLQYWSQELKK | 821 |
| 20 | WSDLRSLCLFSYHRLRDLLLIATRIVELLGR-------RGWEVLKYWWNLLQYWSQELKK | 821 |
| 21 | WSDLRSLCLFSYHRLRDLLLIATRIVELLGR-------RGWEILKYWWNLLQYWSQELKK | 821 |
| 22 | WSDLRSLCLFSYHRLRDLLLIATRIVELLGR-------RGWEVLKYWWNLLQYWSQELKK | 821 |
| 23 | WSDLRSLCLFSYHRLRDLLLIATRIVELLGR-------RGWEVLKYWWNLLQYWSQELKK | 821 |
| 24 | WSDLRSLCLFSYHRLRDLLLIATRIVELLGR-------RGWEVLKYWWNLLQYWSQELKK | 821 |
| 25 | WSDLRSLCLFSYHRLRDLLLIATRIVELLGR-------RGWEVLKYWWNLLQYWSQELKK | 821 |
| 26 | WSDLRSLCLFSYHRLRDLLLIATRIVELLGR-------RGWEVLKYWWNLLQYWSQELKK | 821 |
| 27 | WSDLRSLCLFSYHHLRDLLLIATRIVELLGR-------RGWEVLKYWWNLLQYWSQELKH | 821 |
| 28 | WSDLRSLCLFSYHRLRDLLLIATRIVELLGR-------RGWEVLKYWWNLLQYWSQELKK | 821 |
| 29 | WSDLRSLCLFSYHRLRDLLLIATRIVELLGR-------RGWEVLKYWWNLLQYWSQELKK | 821 |
| 30 | WSDLRSLCLFSYHHLRDLLLIVTRIVELLGR-------RGWEVLKYWWNLLQYWSQELKK | 821 |
| 31 | WSDLRSLCHFSYHRLRDLLLIATRIVELLGR-------RGWEVLKYWWNLLQYWSQELKN | 814 |
| 32 | WSDLRSLCLFSYHRLRDLLLIATRIVELLGR-------RGWEILKYWWNLIQYWSQELKK | 821 |
| 33 | WSDLRSLCLFSYHRLRDLLLIATRIVELLGR-------RGWEVLKYWWNLLQYWGQELKN | 821 |
| 34 | WSDLRSLCLFSYHRLRDLLLIATRIVELLGR-------RGWEVLKYWWNLLQYWSQELKK | 821 |
| 35 | WSDLRSLCLFSYHHLRDLLLIATRIVELLGR-------RGWEVLKYWWNLLQYWSQELKK | 806 |

**SAVSLLNATAIAVAEGTDRVIEVVQRAGRAIRHIPRRIRQGLERALL**

| | | |
|---|---|---|
| 1 | SAVSLLNATAIAVAEGTDRVIEVVQRACRAILHIPRRIRQGLERALL | 846 |
| 2 | SAVSLLNATAIAVAEGTDRVIEVVQRAGRAIRHIPRRIRQGLERALL | 866 |
| 3 | SAVSLLNATAIAVAEGTDRVIEVVQRAGRAIRHIPRRIRQGLERALL | 868 |
| 4 | SAVSLLNATAIAVAEGTDRVIEVVQRAGRAIRHIPRRIRQGLERALL | 868 |
| 5 | SAVSLLNATAIAVAEGTDRVIEVVQRAGRAIRHIPRRIRQGLERALL | 868 |
| 6 | SAVSLLNATAIAVAEGTDRVIEVVQRAGRAIRHIPRRIRQGLERALL | 868 |
| 7 | SAVSLLNATAIAVAEGTDRVIEVVQRAGRAIRHIPRRIRQGLERALL | 868 |
| 8 | SAVSLLNATAIAVAEGTDRVIEVVQRAGRAIRHIPRRIRQGLERALL | 868 |
| 9 | SAVSLLNATAIAVAEGTDRVIEVVQRAGRAIRHIPRRIRQGLERALL | 868 |
| 10 | SAVSLLNATAIAVAEGTDRVIEVVQRAGRTIRHIPRRIRQGLERALL | 868 |
| 11 | SAVSLLNATAIAVAEGTDRVIEVVQRAGRAIRHIPRRIRQGLERALL | 868 |
| 12 | SAVSLLNATAVAVAEGTDRVIEVVQRAGRTIRHIPRRIRQGLERALL | 851 |
| 13 | SAVSLLNATAIAVAEGTDRVIEVVQRAGRAIRHIPRRIRQGLERALL | 868 |
| 14 | SAVSLLNATAIAVAEGTDRVIEVVQRAGRAIRHIPRRIRQGLERALL | 869 |
| 15 | SAVSLLNATAIAVAEGTDRVIEVVQRAGRAIRHIPRRIRQGLERALL | 868 |
| 16 | SAVSLLNATAIAVAEGTDRVIEVVQRAGRAIRHIPRRIRQGLERALL | 868 |
| 17 | SAVSLLNATAIAVAEGTDRVIEVVQRAGRTIRHIPRRIRQGLERALL | 868 |
| 18 | SAVSLLNATAIAVAEGTDRVIEVVQRAGRAIRHIPRRIRQGLERALL | 863 |
| 19 | SAVSLLNATAIAVAEGTDRVIEVVQRAGRAIRHIPRRIRQGLERALL | 868 |
| 20 | SAVSLLNATAIAVAEGTDRVIEVVQRAGRAIRHIPRRIRQGLERALL | 868 |
| 21 | SAVSLLNATAIAVAEGTDRVIEVVQRAGRAIRHIPRRIRQGLERALL | 868 |
| 22 | SAVSLLNATATAVAEGTDRVIEVVQRAGRAIRHIPRRIRQGLERALL | 868 |
| 23 | SAVSLLNATAIAVAEGTDRVIEVVQRAGRAIRHIPRRIRQGLERALL | 868 |
| 24 | SAVSLLNATAIAVAEGTDRVIEVVQRAGRAIRHIPRRIRQGLERALL | 868 |
| 25 | SAVSLLNATAIAVAEGTDRVIEVVQRAGRAIRHIPRRIRQGLERALL | 868 |
| 26 | SAVSLLNATAIAVAEGTDRVIEVVQRAGRAIRHIPRRIRQGLERALL | 868 |
| 27 | SAVSLLNATAIAVAEGTDRVIEVVQRAGRAIRHIPRRIRQGLERALL | 868 |
| 28 | SAVSLLNATAIAVAEGTDRVIEVVQRAGRAIRHIPRRIRQGLERALL | 868 |
| 29 | SAVSLLNATAIAVAEGTDRVIEVVQRAGRAIRHIPRRIRQGLERALL | 868 |
| 30 | SAVSLLNATAIAVAEGTDRVIEVVQRAGRAIRHIPRRIRQGLERALL | 868 |
| 31 | SAVSLLNATAIAVAEGTDRVIEVVQRAGRAIRHIPRRIRQGLERALL | 861 |
| 32 | SAVSLLNATAIAVAEGTDRVIEVVQRAGRAIRHIPRRIRQGLERALL | 868 |
| 33 | SAVSLLNATAIAVAEGTDRIIEVVQRAGRAIRHIPRRIRQGLERALL | 868 |
| 34 | SAVSLLNATAIAVAEGTDRVIEVVQRAGRAIRHIPRRIRQGLERALL | 868 |
| 35 | SAVSLLNATAIAVAEGTDRVIEVVQRAGRAIRHIPRRIRQGLERALL | 853 |

### Aligned using an external algorithm

**Consensus**

1. B.US.2006.700010040_C9_4520.EU289193
2. B.US.2011.CH0040_3_d1737_ipe032_25_13.MG900546
3. B.US.2011.CH0040_3_d1737_ipe032_25_14.MG900547
4. B.US.2011.CH0040_3_d1737_ipe032_25_19.MG900548
5. B.US.2011.CH0040_3_d1737_ipe032_25_20.MG900549
6. B.US.2011.CH0040_3_d1737_ipe032_25_21.MG900550
7. B.US.2011.CH0040_3_d1737_ipe032_25_22.MG900551
8. B.US.2011.CH0040_3_d1737_ipe032_25_24.MG900552
9. B.US.2011.CH0040_3_d1737_ipe032_25_25.MG900553
10. B.US.2011.CH0040_3_d1737_ipe032_25_26.MG900554
11. B.US.2011.CH0040_3_d1737_ipe032_25_27.MG900555
12. B.US.2011.CH0040_3_d1737_ipe032_25_28.MG900556
13. B.US.2011.CH0040_3_d1737_ipe032_25_31.MG900557
14. B.US.2011.CH0040_3_d1737_ipe032_25_33.MG900558
15. B.US.2011.CH0040_3_d1737_ipe032_25_34.MG900559
16. B.US.2011.CH0040_3_d1737_ipe032_25_37.MG900560
17. B.US.2011.CH0040_3_d1737_ipe032_25_38.MG900561
18. B.US.2011.CH0040_3_d1737_ipe032_25_41.MG900562
19. B.US.2011.CH0040_3_d1737_ipe032_25_44.MG900563
20. B.US.2011.CH0040_3_d1737_ipe032_25_45.MG900564
21. B.US.2011.CH0040_3_d1737_ipe032_25_62.MG900565
22. B.US.2011.CH0040_3_d1737_ipe032_25_63.MG900566
23. B.US.2011.CH0040_3_d1737_ipe032_25_66.MG900567
24. B.US.2011.CH0040_3_d1737_ipe032_25_67.MG900568
25. B.US.2011.CH0040_3_d1737_ipe032_25_69.MG900569
26. B.US.2011.CH0040_3_d1737_ipe032_25_71.MG900570
27. B.US.2011.CH0040_3_d1737_ipe032_25_72.MG900571
28. B.US.2011.CH0040_3_d1737_ipe032_25_73.MG900572
29. B.US.2011.CH0040_3_d1737_ipe032_25_74.MG900573
30. B.US.2011.CH0040_3_d1737_ipe032_25_77a.MG900574
31. B.US.2011.CH0040_3_d1737_ipe032_25_78.MG900575
32. B.US.2011.CH0040_3_d1737_ipe032_25_79.MG900576
33. B.US.2011.CH0040_3_d1737_ipe032_25_80.MG900577
34. B.US.2011.CH0040_3_d1737_ipe032_25_81.MG900578
35. B.US.2011.CH0040_3_d1737_ipe032_25_82.MG900579
36. B.US.2011.CH0040_3_d1737_ipe032_25_86.MG900580
37. B.US.2011.CH0040_3_d1737_ipe032_25_87.MG900581
38. B.US.2011.CH0040_3_d1737_ipe032_27_09.MG900582
39. B.US.2011.CH0040_3_d1737_ipe032_27_12.MG900583
40. B.US.2011.CH0040_3_d1737_ipe032_9_01.MG900584
41. B.US.2011.CH0040_3_d1737_ipe032_9_04.MG900585
42. B.US.2011.CH0040_3_d1737_ipe032_9_05.MG900586

```
   MRVMGIRKNYQHLWREGILLLGILMICSAADXLWVTVYYGVPVWREATTTLFCASDAKAY
1  MRVMGIRKNYQHLWREGILLLGILMICSAADNLWVTVYYGVPVWREATTTLFCASDAKAY  60
2  MRVMGIRKNYQHLWREGTLLLGILMICSAEDNLWVTVYYGVPVWREATTTLFCASDAKAY  60
3  MRVMGIRKNYQHLWREGILLLGILMICSAADNLWVTVYYGVPVWREATTTLFCASDAKAY  60
4  MRVMGIRKNYQHLWREGILLLGILMICSAAGNLWVTVYYGVPVWREATTTLFCASDAKAY  60
5  MRVMGIRKNYQHLWREGILFLGILMICSAADNLWVTVYYGVPVWKEATTTLFCASDAKAY  60
6  MRVMGIRKNYQHLWRKGILLLGILMICSAADKLWVTVYYGVPVWREATTTLFCASDAKAY  60
7  MRVMGIRKNYQHWWREGILLLGILMICSAADKLWVTVYYGVPVWREATTTLFCASDAKAY  60
8  MRVMGIRKNYQHLWREGILLLGILMICSAADNLWVTVYYGVPVWREATTTLFCASDAKAY  60
9  MRVMGIRKNYQHLWREGTLLLGILMICSAEDNLWVTVYYGVPVWREATTTLFCASDAKAY  60
10 MRVMGIRKNYQHLWREGILLLGILMICSAADNLWVTVYYGVPVWREATTTLFCASDAKAY  60
11 MRVMGIRKNYQHLWREGILLLGILMICSAADNSWVTVYYGVPVWKEATTTLFCASDAKAY  60
12 MRVMGIRKNYQHWWREGILLLGILMICSAADKLWVTVYYGVPVWREATTTLFCASDAKAY  60
13 MRVMGIRKNYQHLWRKGILLLGILMICSAADKLWVTVYYGVPVWREATTTLFCASDAKAY  60
14 MRVMGIRKNYQHLWRKGILLLGILMICSAADKLWVTVYYGVPVWREATTTLFCASDAKAY  60
15 MRVMGIRKNYQHLWRKGILLLGILMICSAADKLWVTVYYGVPVWREATTTLFCASDAKAY  60
16 MRVMGIRKNYQHLWREGILLLGILMICSAADNLWVTVYYGVPVWKEATTTLFCASDAKAY  60
17 MRVMGIRKNYQHLWREGILLLGILMICSAADNLWVTVYYGVPVWREATTTLFCASDAKAY  60
18 MRVMGIRKNYQHLWREGILLLGILMICSAADDLWVTVYYGVPVWREATTTLFCASDAKAY  60
19 MRVMGIRKNYQHLWREGILLLGILMICSAADNLWVTVYYGVPVWKEATTTLFCASDAKAY  60
20 MRVMGIRKNYQHWWREGILLLGILMICSAADKLWVTVYYGVPVWREATTTLFCASDAKAY  60
21 MRVMGIRKNYQHLWREGILLLGILMICSAADNLWVTVYYGVPVWKEATTTLFCASDAKAY  60
22 MRVMGIRKNYQHLWREGILLLGILMICSAADNLWVTVYYGVPVWREATTTLFCASDAKAY  60
23 MRVMGIRKNYQHLWRKGILLLGILMICSAADKLWVTVYYGVPVWREATTTLFCASDAKAY  60
24 MRVMGIRKNYQHLWREGILLLGILMICSAADNLWVTVYYGVPVWREATTTLFCASDAKAY  60
25 MRVMGIRKNYQHLWRKGILLLGILMICSAADKLWVTVYYGVPVWREATTTLFCASDAKAY  60
26 MRVMGIRKNYQHLWREGILLLGILMICSAADDLWVTVYYGVPVWREATTTLFCASDAKAY  60
27 MRVMGIRKNYQHWWREGILLLGILMICSAADKLWVTVYYGVPVWREATTTLFCASDAKAY  60
28 MRVMGIRKNYQHWWREGILLLGILMICSAADKLWVTVYYGVPVWREATTTLFCASDAKAY  60
29 MRVMGIRKNYQHLWRKGILLLGILMICSAADKLWVTVYYGVPVWREATTTLFCASDAKAY  60
30 MRVMGIRKNYQHWWREGILLLGILMICSAADKLWVTVYYGVPVWREATTTLFCASDAKAY  60
31 MRVMGIRKNYQHLWRKGILLLGILMICSAADKLWVTVYYGVPVWREATTTLFCASDAKAY  60
32 MRVMGIRKNYQHWWREGILLLGILMICSAADKLWVTVYYGVPVWREATTTLFCASDAKAY  60
33 MRVMGIRKNYQHLWREGILLLGILMICSAADNLWVTVYYGVPVWKEATTTLFCASDAKAY  60
34 MRVMEIRKNYQHWWREGILLLGILMICSAADKLWVTVYYGVPVWREATTTLFCASDAKAY  60
35 MRVMGIRKNYQHLWREGILLLGILMICSAADNLWVTVYYGVPVWREATTTLFCASDAKAY  60
36 MRVMGIRKNYQHLWRKGILLLGILMICSAADKLWVTVYYGVPVWREATTTLFCASDAKAY  60
37 MRVMGIRKNYQHWWREGILLLGILMICSAADKLWVTVYYGVPVWREATTTLFCASDAKAY  60
38 MRVMGIRKNYQHWWREGILLLGILMICSAADKLWVTVYYGVPVWREATTTLFCASDAKAY  60
39 MRVMGIRKNYQHLWREGILLLGILMICSAADNLWVTVYYGVPVWREATTTLFCASDAKAY  60
40 MRVMGIRKNYQHLWREGILLLGILMICSAADNLWVTVYYGVPVWREATTTLFCASDAKAY  60
41 MRVMGIRKNYQHLWRKGILLLGILMICSAADKLWVTVYYGVPVWREATTTLFCASDAKAY  60
42 MRVMGIRKNYQHLWREGILLLGILMICSAADNLWVTVYYGVPVWREATTTLFCASDAKAY  60
```

| | | |
|---|---|---|
| | **DTEAHNVWATHACVPTDPNPQEVELKNVTENFNMWENNMVEQMHEDIISLWDQSLKPCVK** | |
| 1 | DTEAHNVWATHACVPTDPNPQEVELKNVTENFNMWENNMVEQMHEDIISLWDQSLKPCVK | 120 |
| 2 | DTEAHNVWATHACVPTDPNPQEVELKNVTENFNMWENNMVEQMHEDIISLWDQSLKPCVK | 120 |
| 3 | DTEAHNVWATHACVPTDPNPQEVELKNVTENFNMWENNMVEQMHEDIISLWDQSLKPCVK | 120 |
| 4 | DTEAHNVWATHACVPTDPNPQEVELKNVTENFNMWENNMVEQMHEDIISLWDQSLKPCVK | 120 |
| 5 | DTEAHNVWATHACVPTDPNPQEVELKNVTENFNMWKNNMVEQMHEDIISLWDQSLKPCVK | 120 |
| 6 | DTEAHNVWATHACVPTDPNPQEVELKNVTENFNMWENNMVEQMHEDIISLWDQSLKPCVK | 120 |
| 7 | DTEAHNVWATHACVPTDPNPQEVELKNVTENFNMWENNMVEQMHEDIISLWDQSLKPCVK | 120 |
| 8 | DTEAHNVWATHACVPTDPNPQEVELKNVTENFNMWENNMVEQMHEDTISLWDQSLKPCVK | 120 |
| 9 | DTEAHNVWATHACVPTDPNPQEVELKNVTENFNMWENNMVEQMHEDIISLWDQSLKPCVK | 120 |
| 10 | DTEAHNVWATHACVPTDPNPQEVELKNVTENFNMWENNMVEQMHEDIISLWDQSLKPCVK | 120 |
| 11 | DTEAHNVWATHACVPTDPNPQEVELKNVTENFNMWKNNMVEQMHEDIISLWDQSLKPCVK | 120 |
| 12 | DTEAHNVWATHACVPTDPNPQEVELKNVTENFNMWENNMVEQMHEDIISLWDQSLKPCVK | 120 |
| 13 | DTEAHNVWATHACVPTDPNPQEVELRNVTENFNMWENNMVEQMHEDIISLWDQSLKPCVK | 120 |
| 14 | DTEAHNVWATHACVPTDPNPQEVELKNVTENFNMWENNMVEQMHEDIISLWDQSLKPCVK | 120 |
| 15 | DTEAHNVWATHACVPTDPNPQEVELKNVTENFNMWENNMVEQMHEDIISLWDQSLKPCVK | 120 |
| 16 | DTEAHNVWATHACVPTDPNPQEVELKNVTENFNMWKNNMVEQMHEDIISLWDQSLKPCVK | 120 |
| 17 | DTEAHNVWATHACVPTDPNPQEVELKNVTENFNMWENNMVEQMHEDIISLWDQSLKPCVK | 120 |
| 18 | DTEAHNVWATHACVPTDPNPQEVELKNVTENFNMWENNMVEQMHEDIISLWDQSLKPCVK | 120 |
| 19 | DAEAHNVWATHACVPTDPNPQEVELKNVTENFNMWKNNMVEQMHEDIISLWDQSLKPCVK | 120 |
| 20 | DTEAHNVWATHACVPTDPNPQEVELKNVTENFNMWENNMVEQMHEDIISLWDQSLKPCVK | 120 |
| 21 | DTEAHNVWATHACVPTDPNPQEVELKNVTENFNMWKNNMVEQMHEDIISLWDQSLKPCVK | 120 |
| 22 | DTEAHNVWATHACVPTDPNPQEVELKNVTENFNMWENNMVEQMHEDIISLWDQSLKPCVK | 120 |
| 23 | DTEAHNVWATHACVPTDPNPQEVELKNVTENFNMWENNMVEQMHEDIISLWDQSLKPCVK | 120 |
| 24 | DTEAHNVWATHACVPTDPNPQEVELKNVTENFNMWENNMVEQMHEDIISLWDQSLKPCVK | 120 |
| 25 | DTEAHNVWATHACVPTDPNPQEVELKNVTENFNMWENNMVEQMHEDIISLWDQSLKPCVK | 120 |
| 26 | DTEAHNVWATHACVPTDPNPQEVELKNVTENFNMWENNMVEQMHEDIISLWDQSLKPCVK | 120 |
| 27 | DTEAHNVWATHACVPTDPNPQEVELKNVTENFNMWENNMVEQMHEDIISLWDQSLKPCVK | 120 |
| 28 | DTEAHNVWATHACVPTDPNPQEVELKNVTENFNMWENNMVEQMHEDIISLWDQSLKPCVK | 120 |
| 29 | DTEAHNVWATHACVPTDPNPQEVELKNVTENFNMWENNMVEQMHEDIISLWDQSLKPCVK | 120 |
| 30 | DTEAHNVWATHACVPTDPNPQEVELKNVTENFNMWENNMVEQMHEDIISLWDQSLKPCVK | 120 |
| 31 | DTEAHNVWATHACVPTDPNPQEVELKNVTENFNMWENNMVEQMHEDIISLWDQSLKPCVK | 120 |
| 32 | DTEAHNVWATHACVPTDPNPQEVELKNVTENFNMWENNMVEQMHEDIISLWDQSLKPCVK | 120 |
| 33 | DTEAHNVWATHACVPTDPNPQEVELKNVTENFNMWKNNMVEQMHEDIISLWDQSLKPCVK | 120 |
| 34 | DTEAHNVWATHACVPTDPNPQEVELKNVTENFNMWENNMVEQMHEDIISLWDQSLKPCVK | 120 |
| 35 | DTEAHNVWATHACVPTDPNPQEVELKNVTENFNMWENNMVEQMHEDIISLWDQSLKPCVK | 120 |
| 36 | DTEAHNVWATHACVPTDPNPQEVELKNVTENFNMWENNMVEQMHEDIISLWDQSLKPCVK | 120 |
| 37 | DTEAHNVWATHACVPTDPNPQEVELKNVTENFNMWENNMVEQMHEDIISLWDQSLKPCVK | 120 |
| 38 | DTEAHNVWATHACVPTDPNPQEVELKNVTENFNMWENNMVEQMHEDIISLWDQSLKPCVK | 120 |
| 39 | DTEAHNVWATHACVPTDPNPQEVELKNVTENFNMWKNNMVEQMHEDIISLWDQSLKPCVK | 120 |
| 40 | DTEAHNVWATHACVPTDPNPQEVELKNVTENFNMWENNMVEQMHEDIISLWDQSLKPCVK | 120 |
| 41 | DTEAHNVWATHACVPTDPNPQEVELKNVTENFNMWENNMVEQMHEDIISLWDQSLKPCVK | 120 |
| 42 | DTEAHNVWATHACVPTDPNPQEVELKNVTENFNMWENNMVEQMHEDIISLWDQSLKPCVK | 120 |

| | Sequence | |
|---|---|---|
| | LTPLCVTLNCTDXXXXTTTTXXXNXTXXXXXXNXTXXXXXX---XXXNEXMMEKGEVKNC | |
| 1 | LTPLCVTLNCTDLGNVTNTT-------------------------NSNGEMMEKGEVKNC | 155 |
| 2 | LTPLCVTLNCTDNVTTTTNRNGTNCTDSGNVTNTTNRNGTL---MGNETLMMEKGEVKNC | 177 |
| 3 | LTPLCVTLNCTDLGNVTTTT-------------------------DRNEKMMEKEEV-NC | 154 |
| 4 | LTPLCVTLNCTDLGNVTTTT-------------------------DRNEKMMEKEEV-NC | 154 |
| 5 | LTPLCVTLNCTDLENVTATTN------------------------SNGTLMEEEGEVKNC | 156 |
| 6 | LTPLCVTLNCTDDLDNVTTTTNRNGTNCTDSGNVTPTTNRNGT-LMGNETLMEKGEVKNC | 179 |
| 7 | LTPLCVTLNCTDNVTTTTNRNGTNCTDSGNVTNTTNRNGTL---MGNETLMMERGEVKNC | 177 |
| 8 | LTPLCVTLNCTDLGNVTTTT-------------------------DRNEKMMEKEEV-NC | 154 |
| 9 | LTPLCVTLNCTDNVTTTTNRNGTNCTDSGNVTNTTNRNGTL---MGNETLMMERGEVKNC | 177 |
| 10 | LTPLCVTLNCTDLGNVTTTT-------------------------DRNEKMMEKEEV-NC | 154 |
| 11 | LTPLCVTLNCTDLENVTATTN------------------------SNGTLMEEEGEVKNC | 156 |
| 12 | LTPLCVTLNCTDDLDNVTTTTNRNGTNCTDSGNVTPTTNRNGT-LMGNETLMEKGEVKNC | 179 |
| 13 | LTPLCVTLNCTDDLDNVTTTTNRNGTNCTDSGNVTPTTNRNGT-LMGNETLMEKGEVKNC | 179 |
| 14 | LTPLCVTLNCTDDLDNVTTTTNRNGTNCTDSGNVTPTTNRNGT-LMGNETLMEKGEVKNC | 179 |
| 15 | LTPLCVTLNCTDDLDNVTTTTNRNGTNCTDSGNVTPTTNRNGT-LMGNETLMEKGEVKNC | 179 |
| 16 | LTPLCVTLNCTDLENDTITT-------------------------NSNGTSMEEGEVKNC | 155 |
| 17 | LTPLCVTLNCTDLGNVTTTT-------------------------DRNEKMMEKEEV-NC | 154 |
| 18 | LTPLCVTLNCTDNVTTTTNRNGTNCTDSGNVTNTTNRNGTL---MGNETLMMEKGEVKNC | 177 |
| 19 | LTPLCVTLNCTDTTTTN-----------------------------SNGTSMEEGEVKNC | 151 |
| 20 | LTPLCVTLNCTDNVTTTTNRNGTNCTDSGNVTNTTNRNGTL---MGNETLMMERGEVKNC | 177 |
| 21 | LTPLCVTLNCTDTTTTN-----------------------------SNGTSMEEGEVKNC | 151 |
| 22 | LTPLCVTLNCTDLGNVTTTT-------------------------DRNEKMMEKEEV-NC | 154 |
| 23 | LTPLCVTLNCTDDLDNVTTTTNRNGTNCTDSGNVTPTTNRNGT-LMGNETLMEKGEVKNC | 179 |
| 24 | LTPLCVTLNCTDLGNVTTTT-------------------------DRNEKMMEKEEV-NC | 154 |
| 25 | LTPLCVTLNCTDNVTTTTNRNGTNCTDSGNVTNTTNRNGTL---MGNETLMMERGEVKNC | 177 |
| 26 | LTPLCVTLNCTDDLDNVTTTTNRNGTNCTDSGNVTPTTNRNGT-LMGNETLMEKGEVKNC | 179 |
| 27 | LTPLCVTLNCTDDLDNVTTTTNRNGTNCTDSGNVTNTTNRNGTLMGNETLMMERGEVKNC | 180 |
| 28 | LTPLCVTLNCTDNVTTTTNRNGTNCTDLRNDTAT----------TNSNETLMEKGEVKNC | 170 |
| 29 | LTPLCVTLNCTDDLDNVTTTTNRNGTNCTDSGNVTPTTNRNGT-LMGNETLMEKGEVKNC | 179 |
| 30 | LTPLCVTLNCTDNVTTTTNRNGTNCTDSGNVTNTTNRNGTL---MGNETLMMEEGEVKNC | 177 |
| 31 | LTPLCVTLNCTDNVTTTTNRNGTNCTDSGNVTNTTNRNGTL---MGNETLMMERGEVKNC | 177 |
| 32 | LTPLCVTLNCTDNVTTTTNRNGTNCTDSGNVTNTTNRNGTL---MGNETLMMERGEVKNC | 177 |
| 33 | LTPLCVTLNCTDLENDTTTT---------------------------NGTSMEEGEVKNC | 153 |
| 34 | LTPLCVTLNCTDNVTTTTNRNGTNCTDSGNVTNTTNRNGTL---MGNETLMMERGEVKNC | 177 |
| 35 | LTPLCVTLNCTDLGNVTTTT-------------------------DRNEKMMEKEEV-NC | 154 |
| 36 | LTPLCVTLNCTDDLDNVTTTTNRNGTNCTDSGNVTPTTNRNGT-LMGNETLMEKGEVKNC | 179 |
| 37 | LTPLCVTLNCTDDLDNVTTTTNRNGTNCTDSGNVTPTTNRNGT-LMGNETLMEKGEVKNC | 179 |
| 38 | LTPLCVTLNCTDDLDNVTTTTNRNGTNCTDSGNVTPTTNRNGT-LMGNETLMEKGEVKNC | 179 |
| 39 | LTPLCVTLNCTDLENDTTTI-------------------------NSNGTSMEEGEVKNC | 155 |
| 40 | LTPLCVTLNCTDLGNVTTTT-------------------------DRNEKMMEKEEV-NC | 154 |
| 41 | LTPLCVTLNCTDDLDNVTTTTNRNGTNCTDSGNVTPTTNRNGT-LMGNETLMEKGEVKNC | 179 |
| 42 | LTPLCVTLNCTDLGNVTTTT-------------------------DRNEKMMEKEEV-NC | 154 |

| | | |
|---|---|---|
|  | SFRITTDMQDRTRKEYALFYKLDVVPIND-------TRYRLVSCNTSVITQACPKVSFEP |  |
| 1 | SFKITTDIKDRTRKEYALFYKLDVVPIND-------TRYRLVSCNTSVITQACPKVSFEP | 208 |
| 2 | SFRITTDMQDRTRKEYALFYKLDVVPIND-------TSYRLVSCNTSVITQACPKVSFEP | 230 |
| 3 | SFKITTDVKDRMRKEYALFYKLDVVPIND-------TRYRLISCNTSVITQACPKVSFEP | 207 |
| 4 | SFKITTDVKDRMRKEYALFYKLDVVPIND-------TRYRLISCNTSVITQACPKVSFEP | 207 |
| 5 | SFKITTEIKDRTRKEYALFYKLDVVPIND-------TRYRLVSCNTSVITQACPKVSFEP | 209 |
| 6 | SFRITTDMQDRTRKEYALFYKLDVVPIND-------TRYRLVSCNTSVITQACPKVSFEP | 232 |
| 7 | SFRITTDMQDRTRKEYALFYKLDVVPIND-------TSYRLVSCNTSVITQACPKVSFEP | 230 |
| 8 | SFKITTDVKDRMRKEYALFYKLDVVPIND-------TRYRLISCNTSVITQACPKVSFEP | 207 |
| 9 | SFRITTDMQDRTRKEYALFYKLDVVPIND-------TSYRLVSCNTSVITQACPKVSFEP | 230 |
| 10 | SFKITTDVKDRMRKEYALFYKLDVVPIND-------TRYRLISCNTSVITQACPKVSFEP | 207 |
| 11 | SFKITTEIKDRTRKEYALFYKLDVVPIND-------TRYRLVSCNTSVITQACPKVSFEP | 209 |
| 12 | SFRITTDMQDRTRKEYALFYKLDVVPIND-------TRYRLVSCNTSVITQACPKVSFEP | 232 |
| 13 | SFRITTDMQDRTRKEYALFYKLDVVPIND-------TRYRLVSCNTSVITQACPKVSFEP | 232 |
| 14 | SFRITTDMQDRTRKEYALFYKLDVVPIND-------TSYRLVSCNTSVITQACPKVSFEP | 232 |
| 15 | SFRITTDMQDRTRKEYALFYKLDVVPIND-------TRYRLVSCNTSVITQACPKVSFEP | 232 |
| 16 | SFKITTEIKDRTRKEYALFYKLDVVPIND-------TRYRLVSCNTSVITQACPKVSFEP | 208 |
| 17 | SFKITTDVKDRMRKEYALFYKLDVVPIND-------TRYRLISCNTSVITQACPKVSFEP | 207 |
| 18 | SFRITTDMQDRTRKEYALFYKLDVVPIND-------TSYRLVSCNTSVITQACPKVSFEP | 230 |
| 19 | SFKITTEMKDRTRKEYALFYKLDVVPIND-------TRYRLVSCNTSVLTQACPKVSFEP | 204 |
| 20 | SFRITTDMQDRTRKEYALFYKLDVVPIND-------TSYRLVSCNTSVITQACPKVSFEP | 230 |
| 21 | SFKITTEMKDRTRKEYALFYKLDVVPIND-------TRYRLVSCNTSVLTQACPKVSFEP | 204 |
| 22 | SFKITTDVKDRMRKEYALFYKLDVVPIND-------TRYRLISCNTSVITQACPKVSFEP | 207 |
| 23 | SFRITTDMQDRTRKEYALFYKLDVVPIND--TPINDTSYRLVSCNTSVITQACPKVSFEP | 237 |
| 24 | SFKITTDVKDRMRKEYALFYKLDVVPIND-------TRYRLISCNTSVITQACPKVSFEP | 207 |
| 25 | SFRITTDMQDRTRKEYALFYKLDVVPIND-------TSYRLVSCNTSVITQACPKVSFEP | 230 |
| 26 | SFRITTDMQDRTRKEYALFYKLDVVPIND-------TRYRLVSCNTSVITQACPKVSFEP | 232 |
| 27 | SFRITTDMQDRTRKEYALFYKLDVVPIND-------TSYRLVSCNTSVITQACPKVSFEP | 233 |
| 28 | SFRITTDMQDRTRKEYALFYKLDVVPIND--TPINDTSYRLVSCNTSVITQACPKVSFEP | 228 |
| 29 | SFMITTDMQDRTRKEYALFYKLDVVPIND-------TRYRLVSCNTSVITQACPKVSFEP | 232 |
| 30 | SFRITTDMQDRTRKEYALFYKLDVVPIND-------TSYRLVSCNTSVITQACPKVSFEP | 230 |
| 31 | SFRITTDMQDRTRKEYALFYKLDVVPIND-------TSYRLVSCNTSVITQACPKVSFEP | 230 |
| 32 | SFRITTDMQDRTRKEYALFYKLDVVPIND-------TSYRLVSCNTSVITQACPKVSFEP | 230 |
| 33 | SFKITTEMKDRTRKEYALFYKLDVVPIND-------TRYRLVSCNTSVLTQACPKVSFEP | 206 |
| 34 | SFRITTDMQDRTRKEYALFYKLDVVPIND-------TSYRLVSCNTSVITQACPKVSFEP | 230 |
| 35 | SFKITTDVKDRMRKEYALFYKLDVVPIND-------TRYRLISCNTSVITQACPKVSFEP | 207 |
| 36 | SFRITTDMQDRTRKEYALFYKLDVVPIND-------TRYRLVSCNTSVITQACPKVSFEP | 232 |
| 37 | SFRITTDMQDRTRKEYALFYKLDVVPIND-------TRYRLVSCNTSVITQACPKVSFEP | 232 |
| 38 | SFRITTDMQDRTRKEYALFYKLDVVPIND-------TRYRLVSCNTSVITQACPKVSFEP | 232 |
| 39 | SFKITTEIRDRTRKEYALFYKLDVVPIND-------TRYRLVSCNTSVITQACPKVSFEP | 208 |
| 40 | SFKITTDVKDRMRKEYALFYKLDVVPIND-------TRYRLISCNTSVITQACPKVSFEP | 207 |
| 41 | SFRITTDMQDRTRKEYALFYKLDVVPIND-------TRYRLVSCNTSVITQACPKVSFEP | 232 |
| 42 | SFKITTDVKDRMRKEYALFYKLDVVPIND-------TRYRLISCNTSVITQACPKVSFEP | 207 |

**IPIHYCAPAGFAILKCNDKQFNGTGPCTNVSTVQCTHGIRPVVSTQLLLNGSLAEEEVVI**

| | | |
|---|---|---|
| 1 | IPIHYCAPAGFAILKCNDKQFIGTGPCTNVSTVQCTHGIRPVVSTQLLLNGSLAEEEVVI | 268 |
| 2 | IPIHYCAPAGFAILKCNDKQFNGTGPCTNVSTVQCTHGIRPVVSTQLLLNGSLAEEEVVI | 290 |
| 3 | IPIHYCAPAGFAILKCNDKQFNGTGPCTNVSAVQCTHGIRPVVSTQLLLNGSLAEEEVVI | 267 |
| 4 | IPIHYCAPAGFAILKCNDKQFNGTGPCTNVSTVQCTHGIRPVVSTQLLLNGSLAEEEVVI | 267 |
| 5 | IPIHYCAPAGFAILKCNDKQFNGTGPCTNVSTVQCTHGIRPVVSTQLLLNGSLAEEEVVI | 269 |
| 6 | IPIHYCAPAGFAILKCNDKQFNGTGPCTNVSTVQCTHGIRPVVSTQLLLNGSLAEEEVVI | 292 |
| 7 | IPIHYCAPAGFAILKCNDKQFNGTGPCTNVSTVQCTHGIRPVVSTQLLLNGSLAEEEVVI | 290 |
| 8 | IPIHYCAPAGFAILKCNDKQFNGTGPCTNVSTVQCTHGIRPVVSTQLLLNGSLAEEEVVI | 267 |
| 9 | IPIHYCAPAGFAILKCNDKQFNGTGPCTNVSTVQCTHGIRPVVSTQLLLNGSLAEEEVVI | 290 |
| 10 | IPIHYCAPAGFAILKCNDKQFNGTGPCTNVSTVQCTHGIRPVVSTQLLLNGSLAEEEVVI | 267 |
| 11 | IPIHYCAPAGFAILKCNDKQFNGTGPCTNVSTVQCTHGIRPVVSTQLLLNGSLAEEEVVI | 269 |
| 12 | IPIHYCAPAGFAILKCNDKQFNGTGPCTNVSTVQCTHGIRPVVSTQLLLNGSLAEEEVVI | 292 |
| 13 | IPIHYCAPAGFAILKCNDKQFNGTGPCTNVSTVQCTHGIRPVVSTQLLLNGSLAEEEVVI | 292 |
| 14 | IPIHYCAPAGFAILKCNDKQFNGTGPCTNVSTVQCTHGIRPVVSTQLLLNGSLAEEEVVI | 292 |
| 15 | IPIHYCAPAGFAILKCNDKQFNGTGPCTNVSTVQCTHGIRPVVSTQLLLNGSLAEEEVVI | 292 |
| 16 | IPIHYCAPAGFAILKCNDKQFNGTGPCTNVSTVQCTHGIRPVVSTQLLLNGSLAEEEVVI | 268 |
| 17 | IPIHYCAPAGFAILKCNDKQFNGTGPCTNVSTVQCTHGIRPVVSTQLLLNGSLAEEEVVI | 267 |
| 18 | IPIHYCAPAGFAILKCNDKQFNGTGPCTNVSTVQCTHGIRPVVSTQLLLNGSLAEEEVVI | 290 |
| 19 | IPIHYCAPAGFAILKCNDKQFNGTGPCTNVSTVQCTHGIRPVVSTQLLLNGSLAEEEVVI | 264 |
| 20 | IPIHYCAPAGFAILKCNDKQFNGTGPCTNVSTVQCTHGIRPVVSTQLLLNGSLAEEEVVI | 290 |
| 21 | IPIHYCAPAGFAILKCNDKQFNGTGPCTNVSTVQCTHGIRPVVSTQLLLNGSLAEEEVVI | 264 |
| 22 | IPIHYCAPAGFAILKCNDKQFNGTGPCTNVSTVQCTHGIRPVVSTQLLLNGSLAEEEVVI | 267 |
| 23 | IPIHYCAPAGFAILKCNDKQFNGTGPCTNVSTVQCTHGIRPVVSTQLLLNGSLAEEEVVI | 297 |
| 24 | IPIHYCAPAGFAILKCNDKQFNGTGPCTNVSTVQCTHGIRPVVSTQLLLNGSLAEEEVVI | 267 |
| 25 | IPIHYCAPAGFAILKCNDKQFNGTGPCTNVSTVQCTHGIRPVVSTQLLLNGSLAEEEVVI | 290 |
| 26 | IPIHYCAPAGFAILKCNDKQFNGTGPCTNVSTVQCTHGIRPVVSTQLLLNGSLAEEEVVI | 292 |
| 27 | IPIHYCAPAGFAILKCNDKQFNGTGPCTNVSTVQCTHGIRPVVSTQLLLNGSLAEEEVVI | 293 |
| 28 | IPIHYCAPAGFAILKCNDKQFNGTGPCTNVSTVQCTHGIRPVVSTQLLLNGSLAEEEVVI | 288 |
| 29 | IPIHYCAPAGFAILKCNDKQFNGTGPCTNVSTVQCTHGIRPVVSTQLLLNGSLAEEEVVI | 292 |
| 30 | IPIHYCAPAGFAILKCNDKQFNGTGPCTNVSTVQCTHGIRPVVSTQLLLNGSLAEEEVVL | 290 |
| 31 | IPIHYCAPAGFAILKCNDKQFNGTGPCTNVSTVQCTHGIRPVVSTQLLLNGSLAEEEVVI | 290 |
| 32 | IPIHYCAPAGFAILKCNDKQFNGTGPCTNVSTVQCTHGIRPVVSTQLLLNGSLAEEEVVI | 290 |
| 33 | IPIHYCAPAGFAILKCNDKQFNGTGPCTNVSTVQCTHGIRPVVSTQLLLNGSLAEEEVVI | 266 |
| 34 | IPIHYCAPAGFAILKCNDKQFNGTGPCTNVSTVQCTHGIRPVVSTQLLLNGSLAEEEVVI | 290 |
| 35 | IPIHYCAPAGFAILKCNDKQFNGTGPCTNVSTVQCTHGIRPVVSTQLLLNGSLAEEEVVI | 267 |
| 36 | IPIHYCAPAGFAILKCNDKQFNGTGPCTNVSTVQCTHGIRPVVSTQLLLNGSLAEEEVVI | 292 |
| 37 | IPIHYCAPAGFAILKCNDKQFNGTGPCTNVSTIQCTHGIRPVVSTQLLLNGSLAEEEVVI | 292 |
| 38 | IPIHYCAPAGFAILKCNDKQFNGTGPCTNVSTVQCTHGIRPVVSTQLLLNGSLAEEEVVI | 292 |
| 39 | IPIHYCAPAGFAILKCNDKQFNGTGPCTNVSTVQCTHGIRPVVSTQLLLNGSLAEEEVVI | 268 |
| 40 | IPIHYCAPAGFAILKCNDKQFNGTGPCTNVSTVQCTHGIRPVVSTQLLLNGSLAEEEVVI | 267 |
| 41 | IPIHYCAPAGFAILKCNDKQFNGTGPCTNVSTVQCTHGIRPVVSTQLLLNGSLAEEEVVI | 292 |
| 42 | IPIHYCAPAGFAILKCNDKQFNGTGPCTNVSTVQCTHGIRPVVSTQLLLNGSLAEEEVVI | 267 |

| | Sequence | Length |
|---|---|---|
| | RSVNFSDNAKTIIVQLNKSVEINCTRPNNNTRKSIHMGPGKAFYARGXIIGDIRKAHCNI | |
| 1 | RSVNFSDNAKTIIVQLNKSVEITCTRPNNNTRKSIPMGPGKAFYARGDITGDIRKAYCEI | 328 |
| 2 | RSVNFSDNAKTIIVQLNKSVEINCTRPNNNTRKSIHMGPGKAFYARGAIIGDIRKAHCNI | 350 |
| 3 | RSVNFSDNAKTIIVQLNKSVEMTCTRPNNNTRKSIPMGPGKVFYARGDIIGDIRKAHCNI | 327 |
| 4 | RSVNFSDNAKTIIVQLNKSVEMTCTRPNNNTRKSIPMGPGKVFYARGDIIGDIRKAHCNI | 327 |
| 5 | RSVNFSDNAKTIIVQLNKSVEINCTRPHNNTRKSIPMGPGKAFYARGNIIGDIRKAYCNI | 329 |
| 6 | RSVNFSDNAKTIIVQLNKSVEINCTRPNNNTRKSIHMGPGKAFYARGAIIGDIRKAHCNI | 352 |
| 7 | RSVNFSDNAKTIIVQLNKSVEINCTRPNNNTRKSIHMGPGKAFYARGGIIGDIRKAHCNI | 350 |
| 8 | RSVNFSDNAKTIIVQLNKSVEMTCTRPNNNTRKSIPMGPGKVFYARGDIIGDIRKAHCNI | 327 |
| 9 | RSVNFSDNAKTIIVQLNKSVEINCTRPNNNTRKSIHMGPGKAFYARGGIIGDIRKAHCNI | 350 |
| 10 | RSVNFSDNAKTIIVQLNKSVEMTCTRPNNNTRKSIPMGPGKVFYARGDIIGDIRKAHCNI | 327 |
| 11 | RSVNFSDNAKTIIVQLNKSVEINCTRPHNNTRKSIPMGPGKAFYARGNIIGDIRKAYCNI | 329 |
| 12 | RSVNFSDNTKTIIVQLNKSVEINCTRPNNNTRKSIHMGPGKAFYARGAIIGDIRKAHCNI | 352 |
| 13 | RSVNFSDNAKTIIVQLNKSVEINCTRPNNNTRKSIHMGPGKAFYARGAIIGDIRKAHCNI | 352 |
| 14 | RSVNFSDNAKTIIVQLNKSVEINCTRPNNNTRKSIHMGPGKAFYARGAIIGDIRKAHCNI | 352 |
| 15 | RSVNFSDNAKTIIVQLNKSVEINCTRPNNNTRKSIHMGPGKAFYARGAIIGDIRKAHCNI | 352 |
| 16 | RSVNFSDNAKTIIVQLNKSVEINCTRPHNNTRKSIPMGPGKAFYARGGIIGDIRKAYCNI | 328 |
| 17 | RSVNFSDNAKTIIVQLNKSVEMTCTRPNNNTRKSIPMGPGKVFYARGDIIGDIRKAHCNI | 327 |
| 18 | RSVNFSDNAKTIIVQLNKSVEINCTRPNNNTRKSIHMGPGKAFYARGGIIGDIRKAHCNI | 350 |
| 19 | RSVNFSDNAKTIIVQLNKSVEINCTRPNNNTRKSIPMGPGKAFYARGGIIGDIRKAYCNI | 324 |
| 20 | RSVNFSDNAKTIIVQLNKSVKINCTRPNNNTRKSIHMGPGKAFYARGGIIGDIRKAHCNI | 350 |
| 21 | RSVNFSDNAKTIIVQLNKSVEINCTRPHNNTRKSIPMGPGKAFYARGNIIGDIRKAYCNI | 324 |
| 22 | RSVNFSDNAKTIIVQLNKSVEMTCTRPNNNTRKSIPMGPGKVFYARGDIIGDIRKAHCNI | 327 |
| 23 | RSVNFSDNAKTIIVQLNKSVEINCTRPNNNTRKSIHMGPGKAFYARGGIIGDIRKAHCNI | 357 |
| 24 | RSVNFSDNAKTIIVQLNKSVEMTCTRPNNNTRKSIPMGPGKVFYARGDIIGDIRKAHCNI | 327 |
| 25 | RSVNFSDNAKTIIVQLNKSVEINCTRPNNNTRKSIHMGPGRAFYARGGIIGDIRKAHCNI | 350 |
| 26 | RSVNFSDNAKTIIVQLNKSVEINCTRPNNNTRKSIHMGPGKAFYARGAIIGDIRKAHCNI | 352 |
| 27 | RSVNFSDNAKTIIVQLNKSVEINCTRPNNNTRKSIHMGPGKAFYARGAIIGDIRKAHCNI | 353 |
| 28 | RSVNFSDNAKTIIVQLNKSVEINCTRPNNNTRKSIHMGPGKAFYARGAIIGDIRKAHCNI | 348 |
| 29 | RSVNFSDNAKTIIVQLNKSVEINCTRPNNNTRKSIHMGPGKAFYARGAIIGDIRKAHCNI | 352 |
| 30 | RSVNFSDNAKTIIVQLNKSVEINCTRPNNNTRKSIHMGPGKAFYARGAIIGDIRKAHCNI | 350 |
| 31 | RSVNFSDNAKTIIVQLNKSVEINCTRPNNNTRKSIHMGPGKAFYARGGIIGDIRKAHCNI | 350 |
| 32 | RSVNFSDNAKTIIVQLNKSVEINCTRPNNNTRKSIHMGPGKAFYARGGIIGDIRKAHCNI | 350 |
| 33 | RSVNFSDNAKTIIVQLNKSVEINCTRPHNNTRKSIPMGPGKAFYARGNIIGDIRKAYCNI | 326 |
| 34 | RSVNFSDNAKTIIVQLNKSVEINCTRPNNNTRKSIHMGPGKAFYARGGIIGDIRKAHCNI | 350 |
| 35 | RSVNFSDNAKTIIVQLNKSVEMTCTRPNNNTRKSIPMGPGKVFYARGDIIGDIRKAHCNI | 327 |
| 36 | RSVNFSDNAKTIIVQLNKSVEINCTRPNNNTRKSIHMGPGKAFYARGAIIGDIRKAHCNI | 352 |
| 37 | RSVNFSDNAKTIIVQLNKSVEINCTRPNNNTRKSIHMGPGKAFYARGAIIGDIRKAHCNI | 352 |
| 38 | RSVNFSDNAKTIIVQLNKSVEINCTRPNNNTRKSIHMGPGKAFYARGAIIGDIRKAHCNI | 352 |
| 39 | RSVNFSDNAKTIIVQLNKSVEINCTRPHNNTRKSIPMGPGKAFYARGGIIGDIRKAYCNI | 328 |
| 40 | RSVNFSDNAKTIIVQLNKSVEMTCTRPNNNTRKSIPMGPGKVFYARGDIIGDIRKAHCNI | 327 |
| 41 | RSVNFSDNAKTIIVQLNKSVEINCTRPNNNTRKSIHMGPGKAFYARGAIIGDIRKAHCNI | 352 |
| 42 | RSVNFSDNAKTIIVQLNKSVEMTCTRPNNNTRKSIPMGPGKVFYARGDIIGDIRKAHCNI | 327 |

```
    SGTEWHSTLKLVVEKLREQY-NKTIVFNHSSGGDPEIVMYSFNCGGEFFYCNSTKLFNST
1   NGTEWHSTLKLVVEKLREQY-NKTIVFNRSSGGDPEIVMYSFNCGGEFFYCNSTKLFNST  387
2   SRAEWYSTLKLVAEKLREQY-HKTIVFNHSSGGDPEIVMYSFNCGGEFFYCNSTKLFNST  409
3   SGTEWHSTLKLVVGKLREQY-NKTIVFNHSSGGDPEIVMYSFNCGGEFFYCNSTKLFNST  386
4   SGTEWHSTLKLVVGKLREQY-NKTIVFNHSSGGDPEIVMYSFNCGGEFFYCNSTKLFNST  386
5   SRPEWHSTLKLVVEKLRKQY-NKTIVFNHSSGGDPEIVMYSFNCGGEFFYCNSTKLFNST  388
6   SGTEWHNTLKLVVEKLKEQY-NKTIVFNHSSGGDPEIVMYSFNCGGEFFYCNSTKLFNST  411
7   SGTEWHNTLKLVVEKLREQY-NKTIVFNHSSGGDPEIVMYSFNCGGEFFYCNSTKLFNST  409
8   SGTEWHSTLKLVVGKLREQY-NKTIVFNHSSGGDPEIVMYSFNCGGEFFYCNSTKLFNST  386
9   SRTEWHSTLKLVAEKLREQY-KKTIVFNHSSGGDPEIVMYSFNCGGEFFYCNSTKLFNST  409
10  SGTEWHSTLKLVVGKLREQY-NKTIVFNHSSGGDPEIVMYSFNCGGEFFYCNSTKLFNST  386
11  SRPEWHSTLKLVVEKLRKQY-NKTIVFNHSSGGDPEIVMYSFNCGGEFFYCNSTKLFNST  388
12  SGTEWHNTLKLVVEKLREQY-NKTIVFNHSSGGDPEIVMYSFNCGGEFFYCNSTKLFNST  411
13  SGTEWHNTLKLVVEKLREQY-NKTIVFNHSSGGDPEIVMYSFNCGGEFFYCNSTKLFNST  411
14  SGTEWHNTLTLVVEKLREQF-NKTIVFNHSSGGDPEIVMYSFNCGGEFFYCNSTKLFNST  411
15  SGTEWHNTLKLVVEKLREQY-NKTIVFNHSSGGDPEIVMYSFNCGGEFFYCNSTKLFNST  411
16  SRAEWHSTLKLVVEKLREQY-NKTIVFNHSSGGDPEIVMYSFNCGGEFFYCNSTKLFNST  387
17  SGTEWHSTLKLVVGKLREQY-NKTIVFNHSSGGDPEIVMYSFNCGGEFFYCNSTKLFNST  386
18  SGTEWHNTLKLVVEKLREQY-NKTIVFNHSSGGDPEIVMYSFNCGGEFFYCNSTKLFNST  409
19  SRAEWHSTLKLVVEKLREQY-NKTIVFNHSSGGDPEIVMYSFNCGGEFFYCNSTKLFNST  383
20  SGTEWHSTLKLVVKKLREQY-NKTIVFNHSSGGDPEIVMYSFNCGGEFFYCNSTKLFNST  409
21  SGPEWHSTLKLVVEKLRAQY-NKTIVFNHSSGGDPEIVMYSFNCGGEFFYCNSTKLFNST  383
22  SGTEWHSTLKLVVGKLREQY-NKTIVFNHSSGGDPEIVMYSFNCGGEFFYCNSTKLFNST  386
23  SGTEWHSTLKLVVEKLREKY-NKTIVFNHSSGGDPEIVMYSFNCGGEFFYCNSTKLFNST  416
24  SGTEWHSTLKLVVGKLREQY-NKTIVFNHSSGGDPEIVMYSFNCGGEFFYCNSTKLFNST  386
25  SGTEWHSTLKLVVKKLREQY-NKTIVFNHSSGGDPEIVMYSFNCGGEFFYCNSTKLFNST  409
26  SGTEWHNTLKLVVEKLREQY-NKTIVFNHSSGGDPEIVMYSFNCGGEFFYCNSTKLFNST  411
27  SGTEWHNTLKLVVEKLREQY-NKTIVFNHSSGGDPEIVMYSFNCGGEFFYCNSTKLFNST  412
28  SGTEWHNTLKLVVKKLREQY-NKTIVFNHSSGGDPEIVMYSFNCGGEFFYCNSTKLFNST  407
29  SGTEWHNTLKLVVKKLREQY-NKTIVFNHSSGGDPEIVMYSFNCGGEFFYCNSTKLFNST  411
30  SGTEWHNTLKLVVKKLREQY-NKTIVFNHSSGGDPEIVMYSFNCGGEFFYCNSTKLFNST  409
31  SGTEWHNTLKLVVEKLREQY-NKTIVFNHSSGGDPEIVMYSFNCGGEFFYCNSTKLFNST  409
32  SGTEWHNTLKLVVEKLREQY-NKTIVFNHSSGGDPEIVMYSFNCGGEFFYCNSTKLFNST  409
33  SGPDWHSTLKLVVEKLREQY-NKTIVFNHSSGGDPEIVMYSFNCGGEFFYCNSTKLFNST  385
34  SGTEWHNTLKLVVKKLREQY-NKTIVFNHSSGGDPEIVMYSFNCGGEFFYCNSTKLFNST  409
35  SGTEWHSTLKLVVGKLREQY-NKTIVFNHSSGGDPEIVMYSFNCGGEFFYCNSTKLFNST  386
36  SGTEWHNTLKLVVEKLREQY-NKTIVFNHSSGGDPEIVMYSFNCGGEFFYCNSTKLFNST  411
37  SGTEWHNTLKLVVEKLREQY-NKTIVFNHSSGGDPEIVMYSFNCGGEFFYCNSTKLFNST  411
38  SGTEWHNTLKLVVEKLREQY-NKTIVFNHSSGGDPEIVMYSFNCGGEFFYCNSTKLFNST  411
39  SRAEWHSTLKLVVEKLREQY-NKTIVFNHSSGGDPEIVMYSFNCGGEFFYCNSTKLFNST  387
40  SGTEWHSTLKLVVGKLREQY-NKTIVFNHSSGGDPEIVMYSFNCGGEFFYCNSTKLFNST  386
41  SGTEWHNTLKLVVKKLREQY-NKTIVFNHSSGGDPEIVMYSFNCGGEFFYCNSTKLFNST  411
42  SGTEWHSTLKLVVGKLREQY-NKTIVFNHSSGGDPEIVMYSFNCGGEFFYCNSTKLFNST  386
```

| | | |
|---|---|---|
| | WPXNX--TXGSHDTNGTLTLPCRIKQIINMWQEVGKAMYAPPIEGIIKCSSNITGLLLTR | |
| 1 | WPWND--TKGSHDTNGTLILPCKIKQIINMWQGVGKAMYAPPIEGKIRCSSNITGLLLTR | 445 |
| 2 | WPGND--TEGSHDTNGTLTLPCRIKQIINMWQEVGKAMYAPPIEGIIKCSSNITGLLLTR | 467 |
| 3 | WPWNG--TNGTHDTNGTLTLPCRIKQIINMWQEVGKAMYAPPIEGIIKCSSNITGLLLTR | 444 |
| 4 | WPWNG--TNGSHDTNGTLTLPCRIKQIINMWQEVGKAMYAPPIEGIIKCSSNITGLLLTR | 444 |
| 5 | WPWND--TNRSHDTNDTLMLPCKIKQII-TWQGVGKAMYAPPIEGEIRCSSNITGLLLTR | 445 |
| 6 | WPRNA--TKGSPDTNGTLTLPCRIKQIINMWQEVGKAMYAPPIEGIIKCSSNITGLLLTR | 469 |
| 7 | WHGND--TEGSHNTNGTLTLPCRIKQIINMWQEVGKAMYAPPIEGIIKCSSNITGLLLTR | 467 |
| 8 | WPWNG--TNGSHDTNGTLTLPCRIKQIINMWQEVGKAMYAPPIEGIIKCSSNITGLLLTR | 444 |
| 9 | WPGSD--TEGSHNTNGTLTLPCRIKQIINMWQEVGKAMYAPPIEGIIKCSSNITGLLLTR | 467 |
| 10 | WPWNG--TNGSHDTNGTLTLPCRIKQIINMWQEVGKAMYAPPIEGIIKCSSNITGLLLTR | 444 |
| 11 | WPWND--TNRSHDTNDTLMLPCKIKQIINMWQGVGKAMYAPPIEGEIRCSSNITGLLLTR | 446 |
| 12 | WPRNA--IRGSPDTNGTLTLPCRIKQIINMWQEVGKAMYAPPIEGIIKCSSNITGLLLTR | 469 |
| 13 | WHGND--TEGSHNTNGTLTLPCRIKQIINMWQEVGKAMYAPPIEGIIKCSSNITGLLLTR | 469 |
| 14 | WHRNA--TRGSPDTNGTLTLPCRIKQIINMWQEVGKAMYAPPIEGIIKCSSNITGLLLTR | 469 |
| 15 | WPGNG--TNGSPDTNGTLTLPCRIKQIINMWQEVGKAMYAPPIEGIIKCSSNITGLLLTR | 469 |
| 16 | WPRND--TQGSHDTNDTLMLPCKIKQIINMWQGVGKAMYAPPIEGEIRCSSNITGLLLTR | 445 |
| 17 | WPWNG--TNGSHDTNGTLTLPCRIKQIINMWQEVGKAMYAPPIKGIIKCSSNITGLLLTR | 444 |
| 18 | WLGND--TEGSHNTNGTLTLPCRIKQIINMWQGVGKAMYAPPIEGIIKCSSNITGLLLTR | 467 |
| 19 | WPRNA--TQGSHDTNDTLMLPCKIKQIINMWQGVGKAMYAPPIEGEIRCSSNITGLLLTR | 441 |
| 20 | WPGND--TEGSHNTNGTLTLPCRIKQIINMWQEVGKAMYAPPIEGIIKCSSNITGLLLTR | 467 |
| 21 | WPWND--TNRSHDTNDTLMLPCKIKQIINMWQGVGKAMYAPPIEGEIRCSSNITGLLLTR | 441 |
| 22 | WPWNG--TNGSHDTNGTLTLPCRIKQIINMWQEVGKAMYAPPIEGIIKCSSNITGLLLTR | 444 |
| 23 | WPRNA--TKGSPDTNGTLTLPCRIKQIINMWQEVGKAMYAPPIEGIIKCSSNITGLLLTR | 474 |
| 24 | WPWNG--TNGSHDTNGTLTLPCRIKQIINMWQEVGKAMYAPPIEGIIKCSSNITGLLLTR | 444 |
| 25 | WPWNDNDTEGSHDTNGTLTLPCRIKQIINMWQGVGKAMYAPPIEGIIKCSSNITGLLLTR | 469 |
| 26 | WHGND--TEGSHNTNGTLTLPCRIKQIINMWQEVGKAMYAPPIEGIIKCSSNITGLLLTR | 469 |
| 27 | WHGND--TEGSHNTNGTLTLPCRIKQIINMWQEVGKAMYAPPIEGIIKCSSNITGLLLTR | 470 |
| 28 | WPWNG--TQGSHDTNGTLTLPCRIKQIINMWQGVGKAMYAPPIEGIIKCSSNITGLLLTR | 465 |
| 29 | WPWNND-TEGSHDTNGTLTLPCRIKQIINMWQEVGKAMYAPPIEGIIKCSSNITGLLLTR | 470 |
| 30 | WPGND--TNGSHDTNGTLTLPCRIKQIINMWQGVGKAMYAPPIEGIIKCSSNITGLLLTR | 467 |
| 31 | WPRNT--TRGSPDTNGTLTLPCRIKQIINMWQEVGKAMYAPPIEGIIKCSSNITGLLLTR | 467 |
| 32 | WHGND--TEGSHNTNGTLTLPCRIKQIINMWQEVGKAMYAPPIEGIIKCSSNITGLLLTR | 467 |
| 33 | WPRND--TKRSHDTNDTLMLPCKIKQIINMWQGVGKAMYAPPIEGEIRCSSNITGLLLTR | 443 |
| 34 | WPGND--TEGSHNTNGTLTLPCRIKQIINMWQGVGKAMYAPPIEGIIKCSSNITGLLLTR | 467 |
| 35 | WPWNG--TNGSHDTNGTLTLPCRIKQIINMWQEVGKAMYAPPIEGIIKCSSNITGLLLTR | 444 |
| 36 | WTRNA--TRGSPDTNGTLTLPCRIKQIINMWQEVGKAMYAPPIEGIIKCSSNITGLLLTR | 469 |
| 37 | WPRNT--TRGSPDTNGTLTLPCRIKQIINMWQEVGKAMYAPPIEGIIKCSSNITGLLLTR | 469 |
| 38 | WPRNT--TRGSPDTNGTLTLPCRIKQIINMWQEVGKAMYAPPIEGIIKCSSNITGLLLTR | 469 |
| 39 | WSWNG--TKGSHDTNDTLMLPCKIKQIINMWQGVGKAMYAPPIEGEIRCSSNITGLLLTR | 445 |
| 40 | WPWNG--TNGSHDTNGTLTLPCRIKQIINMWQEVGKAMYAPPIEGIIKCSSNITGLLLTR | 444 |
| 41 | WPWNG--TNGSHDTNGTLTLPCRIKQIINMWQGVGKAMYAPPIEGIIKCSSNITGLLLTR | 469 |
| 42 | WPWNG--TNGSHDTNGTLTLPCRIKQIINMWQEVGKAMYAPPIEGIIKCSSNITGLLLTR | 444 |

DGG-YESNKTDEIFRPGGGDMRDNWRSELYKYKVVKIEPLGVAPTKAKRRVVQREKRAFG

| | | |
|---|---|---|
| 1 | DGG-YESNETDEIFRPGGGDMRDNWRSELYKYKVVKIEPLGVAPTKAKRRVVQREKRAFG | 504 |
| 2 | DGG-YESNKTDEIFRPGGGDMRDNWRSELYKYKVVKIEPLGVAPTKAKRRVVQREKRAFG | 526 |
| 3 | DGG-YESNETDEIFRPGGGDMRDNWRSELYKYKVVKIEPLGVAPTKAKRRVVQREKRAFG | 503 |
| 4 | DGG-YESNETDEIFRPGGGDMRDNWRSELYKYKVVKIEPLGVAPTKAKRRVVQREKRAFG | 503 |
| 5 | DGG-YESNETDEIFRPGGGDMRDNWRSELYKYKVVKIEPLGVAPTRAKRRVVQREKRAFG | 504 |
| 6 | DGG-YESNKTDEIFRPGGGDMRDNWRSELYKYKVVKIEPLGVAPTKAKRRVVQREKRAFG | 528 |
| 7 | DGG-YESNKTDEIFRPGGGDMRDNWRSELYKYKVVKIEPLGVAPTKAKRRVVQREKRAFG | 526 |
| 8 | DGG-YESNETDEIFRPGGGDMRDNWRSELYKYKVVKIEPLGVAPTKAKRRVVQREKRAFG | 503 |
| 9 | DGG-YESNKTDEIFRPGGGDMRDNWRSELYKYKVVKIEPLGVAPTKAKRRVVQREKRAFG | 526 |
| 10 | DGG-YESNETDEIFRPGGGDMRDNWRSELYKYKVVKIEPLGVAPTKAKRRVVQREKRAFG | 503 |
| 11 | DGG-YESNETDEIFRPGGGDMRDNWRSELYKYKVVKIEPLGVAPTKAKRRVVQREKRAFG | 505 |
| 12 | DGG-YESNKTDEIFRPGGGDMRDNWRSELYKYKVVKIEPLGVAPTKAKRRVVQREKRAFG | 528 |
| 13 | DGG-YESNKTDEIFRPGGGDMRDNWRSELYKYKVVKIEPLGVAPTKAKRRVVQREKRAFG | 528 |
| 14 | DGG-YESNKTDEIFRPGGGDMRDNWRSELYKYKVVKIEPLGVAPTKAKRRVVQREKRAFG | 528 |
| 15 | DGG-YESNKTDEIFRPGGGDMRDNWRSELYKYKVVKIEPLGVAPTKAKRRVVQREKRAFG | 528 |
| 16 | DGG-YESNETDEIFRPGGGDMRDNWRSELYKYKVVKIEPLGVAPTKAKRRVVQREKRAFG | 504 |
| 17 | DGG-YESNETDEIFRPGGGDMRDNWRSELYKYKVVKIEPLGVAPTKAKRRVVQREKRAFG | 503 |
| 18 | DGG-YESNKTDEIFRPGGGDMRDNWRSELYKYKVVKIEPLGVAPTKAKRRVVQREKRAFG | 526 |
| 19 | DGG-YESNETDEIFRPGGGDMRDNWRSELYKYKVVKIEPLGVAPTKAKRRVVQREKRAFG | 500 |
| 20 | DGG-YGSNKTDEIFRPGGGDMRDNWRSELYKYKVVKIEPLGVAPTKAKRRVVQREKRAFG | 526 |
| 21 | DGG-YESNETDEIFRPGGGDMRDNWRSELYKYKVVKIEPLGVAPTKAKRRVVQREKRAFG | 500 |
| 22 | DGG-YESNETDEIFRPGGGDMRDNWRSELYKYKVVKIEPLGVAPTKAKRRVVQREKRAFG | 503 |
| 23 | DGG-YESNKTDEIFRPGGGDMRDNWRSELYKYKVVKIEPLGVAPTKAKRRVVQREKRAFG | 533 |
| 24 | DGG-YESNETDEIFRPGGGDMRDNWRSELYKYKVVKIEPLGVAPTKAKRRVVQREKRAFG | 503 |
| 25 | DGG-YESNKTDEIFRPGGGDMRDNWRSELYKYKVVKIEPLGVAPTKAKRRVVQREKRAFG | 528 |
| 26 | DGG-YESNKTDEIFRPGGGDMRDNWRSELYKYKVVKIEPLGVAPTKAKRRVVQREKRAFG | 528 |
| 27 | DGG-YESNKTDEIFRPGGGDMRDNWRSELYKYKVVKIEPLGVAPTKAKRRVVQREKRAFG | 529 |
| 28 | DGG-YESNKTDEIFRPGGGDMRDNWRSELYKYKVVKIEPLGVAPTKAKRRVVQREKRAFG | 524 |
| 29 | DGG-YESNKTDEIFRPGGGDMRDNWRSELYKYKVVKIEPLGVAPTKAKRRVVQREKRAFG | 529 |
| 30 | DGG-YESNKTDETFRPGGGDMRDNWRSELYKYKVVKIEPLGVAPTKAKRRVVQREKRAFG | 526 |
| 31 | DGG-YESNKTDEVFRPGGGDMRDNWRSELYKYKVVKIEPLGVAPTKAKRRVVQREKRAFG | 526 |
| 32 | DGG-YESNKTDEIFRPGGGDMRDNWRSELYKYKVVKIEPLGVAPTKAKRRVVQREKRAFG | 526 |
| 33 | DGG-YESNETDEIFRPGGGDMRDNWRSELYKYKVVKIEPLGVAPTKAKRRVVQREKRAFG | 502 |
| 34 | DGG-YESNKTDEIFRPGGGDMRDNWRSELYKYKVVKIEPLGVAPTKAKRRVVQREKRAFG | 526 |
| 35 | DGG-YESNETDEIFRPGGGDMRDNWRSELYKYKVVKIEPLGVAPTKAKRRVVQREKRAFG | 503 |
| 36 | DGG-YESNKTDEIFRPGGGDMRDNWRSELYKYKVVKIEPLGVAPTKAKRRVVQREKRAFG | 528 |
| 37 | DGG-YESNKTDEIFRPGGGDMRDNWRSELYKYKVVKIEPLGVAPTKAKRRVVQREKRAFG | 528 |
| 38 | DGG-YESNKTDEIFRPGGGDMRDNWRSELYKYKVVKIEPLGVAPTKAKRRVVQREKRAFG | 528 |
| 39 | DGG-YESNETDEIFRPGGGDMRDNWRSELYKYKVVKIEPLGVAPTKAKRRVVQREKRAFG | 504 |
| 40 | DGG-YESNETDEIFRPGGGDMRDNWRSELYKYKVVKIE-LGVAPTKAKRRVVQREKRAFG | 502 |
| 41 | DGG-YESNKTDEIFRPGGGDMRDNWRSELYKYKVVKIEPLGVAPTKAKRRVVQREKRAFG | 528 |
| 42 | DGG-YESNETDEIFRPGGGDMRDNWRSELYKYKVVKIEPLGVAPTKAKRRVVQREKRAFG | 503 |

| | Sequence | |
|---|---|---|
| | -LGAVFLGFLGAAGSTMGAASITLTVQARQLLSGIVQQQNNLLRAIEAQQHLLQLTVWGI | |
| 1 | -LGAVFLGFLGAAGSTMGAASITLTVQARQLLSGIVQQQNNLLRAIEAQQHLLQLTVWGI | 563 |
| 2 | -LGAVFLGFLGAAGSTMGAASITLTVQARQLLSGIVQQQNNLLRAIEAQQHLLQLTVWGI | 585 |
| 3 | -LGAVFLGFLGAAGSTMGAASITLTVQARQLLSGIVQQQNNLLRAIEAQQHLLQLTVWGI | 562 |
| 4 | -LGAVFLGFLGAAGSTMGAASITLTVQARQLLSGIVQQQNNLLRAIEAQQHLLQLTVWGI | 562 |
| 5 | -LGAVFLGFLGAAGSTMGAASITLTVQARQLLSGIVQQQNNLLRAIEAQQHLLQLTVWGI | 563 |
| 6 | -LGAVFLGFLGAAGSTMGAASITLTVQARQLLSGIVQQQNNLLRAIEAQQHLLQLTVWGI | 587 |
| 7 | -LGAVFLGFLGAAGSTMGAASITLTVQARQLLSGIVQQQNNLLRAIEAQQHLLQLTVWGI | 585 |
| 8 | -LGAVFLGFLGAAGSTMGAASITLTVQARQLLSGIVQQQNNLLRAIEAQQHLLQLTVWGI | 562 |
| 9 | -LGAVFLGFLGAAGSTMGAASITLTVQARQLLSGIVQQQNNLLRAIEAQQHLLQLTVWGI | 585 |
| 10 | -LGAVFLGFLGAAGSTMGAASITLTVQARQLLSGIVQQQNNLLRAIEAQQHLLQLTVWGI | 562 |
| 11 | -LGAVFLGFLGAAGSTMGAASITLTVQARQLLSGIVQQQNNLLRAIEAQQHLLQLTVWGI | 564 |
| 12 | -LGAVFLGFLGAAGSTMGAASITLTVQARQLLSGIVQQQNNLLRAIEAQQHLLQLTVWGI | 587 |
| 13 | -LGAVFLGFLGAAGSTMGAASITLTVQARQLLSGIVQQQNNLLRAIEAQQHLLQLTVWGI | 587 |
| 14 | -LGAVFLGFLGAAGSTMGAASITLTVQARQLLSGIVQQQNNLLRAIEAQQHLLQLTVWGI | 587 |
| 15 | -LGAVFLGFLGAAGSTMGAASITLTVQARQLLSGIVQQQNNLLRAIEAQQHLLQLTVWGI | 587 |
| 16 | -LGAVFLGFLGAAGSTMGAASITLTVQARQLLSGIVQQQNNLLRAIEAQQHLLQLTVWGI | 563 |
| 17 | -LGAVFLGFLGAAGSTMGAASITLTVQARQLLSGIVQQQNNLLRAIEAQQHLLQLTVWGI | 562 |
| 18 | -LGAVFLGFLGAAGSTMGAASITLTVQARQLLSGIVQQQNNLLRAIEAQQHLLQLTVWGI | 585 |
| 19 | -LGAVFLGFLGAAGSTMGAASITLTVQARQLLSGIVQQQNNLLRAIEAQQHLLQLTVWGI | 559 |
| 20 | -LGAVFLGFLGAAGSTMGAASITLTVQARQLLSGIVQQQNNLLRAIEAQQHLLQLTVWGI | 585 |
| 21 | -LGAVFLGFLGAAGSTMGAASITLTVQARQLLSGIVQQQNNLLRAIEAQQHLLQLTVWGI | 559 |
| 22 | -LGAVFLGFLGAAGSTMGAASITLTVQARQLLSGIVQQQNNLLRAIEAQQHLLQLTVWGI | 562 |
| 23 | -LGAVFLGFLGAAGSTMGAASITLTVQARQLLSGIVQQQNNLLRAIEAQQHLLQLTVWGI | 592 |
| 24 | -LGAVFLGFLGAAGSTMGAASITLTVQARQLLSGIVQQQNNLLRAIEAQQHLLQLTVWGI | 562 |
| 25 | -LGAVFLGFLGAAGSTMGAASITLTVQARQLLSGIVQQQNNLLRAIEAQQHLLQLTVWGI | 587 |
| 26 | -LGAVFLGFLGAAGSTMGAASITLTVQARQLLSGIVQQQNNLLRAIEAQQHLLQLTVWGI | 587 |
| 27 | -LGAVFLGFLGAAGSTMGAASITLTVQARQLLSGIVQQQNNLLRAIEAQQHLLQLTVWGI | 588 |
| 28 | -LGAVFLGFLGAAGSTMGAASITLTVQARQLLSGIVQQQNNLLRAIEAQQHLLQLTVWGI | 583 |
| 29 | -LGAVFLGFLGAAGSTMGAASITLTVQARQLLSGIVQQQNNLLRAIEAQQHLLQLTVWGI | 588 |
| 30 | -LGAVFLGFLGAAGSTMGAASITLTVQARQLLSGIVQQQNNLLRAIEAQQHLLQLTVWGI | 585 |
| 31 | -LGAVFLGFLGAAGSTMGAASITLTVQARQLLSGIVQQQNNLLRAIEAQQHLLQLTVWGI | 585 |
| 32 | -LGAVFLGFLGAAGSTMGAASITLTVQARQLLSGIVQQQNNLLRAIEAQQHLLQLTVWGI | 585 |
| 33 | -LGAVFLGFLGAAGSTMGAASITLTVQARQLLSGIVQQQNNLLRAIEAQQHLLQLTAWGI | 561 |
| 34 | -LGAVFLGFLGAAGSTMGAASITLTVQARQLLSGIVQQQNNLLRAIEAQQHLLQLTVWGI | 585 |
| 35 | -LGAVFLGFLGAAGSTMGAASITLTVQARQLLSGIVQQQNNLLRAIEAQQHLLQLTVWGI | 562 |
| 36 | -LGAVFLGFLGAAGSTMGAASITLTVQARQLLSGIVQQQNNLLRAIEAQQHLLQLTVWGI | 587 |
| 37 | -LGAVFLGFLGAAGSTMGAASITLTVQARQLLSGIVQQQNNLLRAIEAQQHLLQLTVWGI | 587 |
| 38 | -LGAVFLGFLGAAGSTMGAASITLTVQARQLLSGIVQQQNNLLRAIEAQQHLLQLTVWGI | 587 |
| 39 | -LGAVFLGFLGAAGSTMGAASITLTVQARQLLSGIVQQQNNLLRAIEAQQHLLQLTVWGI | 563 |
| 40 | -LGAVFLGFLGAAGSTMGAASITLTVQARQLLSGIVQQQNNLLRAIEAQQHLLQLTVWGI | 561 |
| 41 | -LGAVFLGFLGAAGSTMGAASITLTVQARQLLSGIVQQQNNLLRAIEAQQHLLQLTVWGI | 587 |
| 42 | -LGAVFLGFLGAAGSTMGAASITLTVQARQLLSGIVQQQNNLLRAIEAQQHLLQLTVWGI | 562 |

| | | |
|---|---|---|
| | **KQLQARVLAVERYLKDQQLLGIWGCSGKLICTTTVPWNTSWSNKSLEQIWDNMTWMEWER** | |
| 1 | KQLQARVLAVERYLKDQQLLGIWGCSGKLICTTTVPWNTSWSNKSLEQIWDNMTWMEWER | 623 |
| 2 | KQLQARVLAVERYLKDQQLLGIWGCSGKLICTTTVPWNTSWSNKSLEQIWDNMTWMEWER | 645 |
| 3 | KQLQARVLAVERYLKDQQLLGIWGCSGKLICTTTVPWNTSWSNKSLEQIWDNMTWMEWER | 622 |
| 4 | KQLQARVLAVERYLKDQQLLGIWGCSGKLICTTTVPWNTSWSNKSLEQIWDNMTWMEWER | 622 |
| 5 | KQLQARVLAVERYLKDQQLLGIWGCSGKLICTTTVPWNTSWSNKSLEQIWDNMTWMEWER | 623 |
| 6 | KQLQARVLAVERYLKDQQLLGIWGCSGKLICTTTVPWNTSWSNKSLEQIWDNMTWMEWER | 647 |
| 7 | KQLQARVLAVERYLKDQQLLGIWGCSGKLICTTTVPWNTSWSNKSLEQIWDNMTWMEWER | 645 |
| 8 | KQLQARVLAVERYLKDQQLLGIWGCSGKLICTTTVPWNTSWSNKSLEQIWDNMTWMEWER | 622 |
| 9 | KQLQARVLAVERYLKDQQLLGIWGCSGKLICTTTVPWNTSWSNKSLEQIWDNMTWMEWER | 645 |
| 10 | KQLQARVLAVERYLKDQQLLGIWGCSGKLICTTTVPWNTSWSNKSLEQIWDNMTWMEWER | 622 |
| 11 | KQLQARVLAVERYLKDQQLLGIWGCSGKLICTTTVPWNTSWSNKSLEQIWDNMTWMEWER | 624 |
| 12 | KQLQARVLAVERYLKDQQLLGIWGCSGKLICTTTVPWNTSWSNKSLEQIWDNMTWMEWER | 647 |
| 13 | KQLQARVLAVERYLKDQQLLGIWGCSGKLICTTTVPWNTSWSNKSLEQIWDNMTWMEWER | 647 |
| 14 | KQLQARVLAVERYLKDQQLLGIWGCSGKLICTTTVPWNTSWSNKSLEQIWDNMTWMEWER | 647 |
| 15 | KQLQARVLAVERYLKDQQLLGIWGCSGKLICTTTVPWNTSWSNKSLEQIWDNMTWMEWER | 647 |
| 16 | KQLQARVLAVERYLKDQQLLGIWGCSGKLICTTTVPWNTSWSNKSLEQIWDNMTWMEWER | 623 |
| 17 | KQLQARVLAVERYLKDQQLLGIWGCSGKLICTTTVPWNTSWSNKSLEQIWDNMTWMEWER | 622 |
| 18 | KQLQARVLAVERYLKDQQLLGIWGCSGKLICTTTVPWNTSWSNKSLEQIWDNMTWMEWER | 645 |
| 19 | KQLQARVLAVERYLKDQQLLGIWGCSGKLICTTTVPWNTSWSNKSLEQIWDNITWMEWER | 619 |
| 20 | KQLQARVLAVERYLKDQQLLGIWGCSGKLICTTTVPWNTSWSNKSLEQIWDNMTWMEWER | 645 |
| 21 | KQLQARVLAVERYLKDQQLLGIWGCSGKLICTTTVPWNTSWSNKSLEQIWDNMTWMEWER | 619 |
| 22 | KQLQARVLAVERYLKDQQLLGIWGCSGKLICTTTVPWNTSWSNKSLEQIWDNMTWMEWER | 622 |
| 23 | KQLQARVLAVERYLKDQQLLGIWGCSGKLICTTTVPWNTSWSNKSLEQIWDNMTWMEWER | 652 |
| 24 | KQLQARVLAVERYLKDQQLLGIWGCSGKLICTTTVPWNTSWSNKSLEQIWDNMTWMEWER | 622 |
| 25 | KQLQARVLAVERYLKDQQLLGIWGCSGKLICTTTVPWNTSWSNKSLEQIWDNMTWMEWER | 647 |
| 26 | KQLQARVLAVERYLKDQQLLGIWGCSGKLICTTTVPWNTSWSNKSLEQIWDNMTWMEWER | 647 |
| 27 | KQLQARVLAVERYLKDQQLLGIWGCSGKLICTTTVPWNTSWSNKSLEQIWNNMTWMEWER | 648 |
| 28 | KQLQARVLAVERYLKDQQLLGIWGCSGKLICTTTVPWNTSWSNKSLEQIWDNMTWMEWER | 643 |
| 29 | KQLQARVLAVERYLKDQQLLGIWGCSGKLICTTTVPWNTSWSNKSLEQIWDNMTWMEWER | 648 |
| 30 | KQLQARVLAVERYLKDQQLLGIWGCSGKLICTTTVPWNTSWSNKSLEQIWDNMTWMEWER | 645 |
| 31 | KQLQARVLAVERYLKDQQLLGIWGCSGKLICTTTVPWNTSWSNKSLEQIWDNMTWMEWER | 645 |
| 32 | KQLQARVLAVERYLKDQQLLGIWGCSGKLICTTTVPWNTSWSNKSLEQIWDNMTWMEWER | 645 |
| 33 | KQLQARVLAVERYLKDQQLLGIWGCSGKLICTTTVPWNTSWSNKSLEQIWDNMTWMEWER | 621 |
| 34 | KQLQARVLAVERYLKDQQLLGIWGCSGKLICTTTVPWNTSWSNKSLEQIWDNMTWMEWER | 645 |
| 35 | KQLQARVLAVERYLKDQQLLGIWGCSGKLICTTTVPWNTSWSNKSLEQIWDNMTWMEWER | 622 |
| 36 | KQLQARVLAVERYLKDQQLLGIWGCSGKLICTTTVPWNTSWSNKSLEQIWDNMTWMEWER | 647 |
| 37 | KQLQARVLAVERYLKDQQLLGIWGCSGKLICTTTVPWNTSWSNKSLEQIWDNMTWMEWER | 647 |
| 38 | KQLQARVLAVERYLKDQQLLGIWGCSGKLICTTTVPWNTSWSNKSLEQIWDNMTWMEWER | 647 |
| 39 | KQLQARVLAVERYLKDQQLLGIWGCSGKLICTTTVPWNTSWSNKSLEQIWDNMTWMEWER | 623 |
| 40 | KQLQARVLAVERYLKDQQLLGIWGCSGKLICTTTVPWNTSWSNKSLEQIWDNMTWMEWER | 621 |
| 41 | KQLQARVLAVERYLKDQQLLGIWGCSGKLICTTTVPWNTSWSNKSLEQIWDNMTWMEWER | 647 |
| 42 | KQLQARVLAVERYLKDQQLLGIWGCSGKLICTTTVPWNTSWSNKSLEQIWDNMTWMEWER | 622 |

| | | |
|---|---|---|
| | EINNYTGYIYQLIEESQNQQEKNEQELLALDKWASLWNWFDITNWLWYIKIFIMIVGGLI | |
| 1 | EIDNYTGYIYQLIEESQNQQEKNEQELLALDKWASLWNWFDITNWLWYIKIFIMIVGGLI | 683 |
| 2 | EINNYTGYIYQLIEESQNQQEKNEQELLAWDKWASLWNWFDITNWLWYIKIVIMIVGGLI | 705 |
| 3 | EIDNYTGYIYQLIEESQNQQEKNEQELLALDKWASLWNWFDITNWLWYIKIFIMIVGGLI | 682 |
| 4 | EISNYTGYIYQLIEESQNQQEKNEQELLALDKWASLWNWFDITNWLWYIKIFIMIVGGLI | 682 |
| 5 | EINNYTGYIYQLIEESQNQQEKNEQELLALDKWASLWNWFDITNWLWYIKIFIMIVGGLI | 683 |
| 6 | EINNYTGYIYQLIEESQNQQEKNEQELLAWDKWASLWNWFDITNWLWYIKIFIMIVGGLI | 707 |
| 7 | EINNYTGYIYQLIEESQNQQEKNEQELLAWDKWASLWNWFDITNWLWYIKIFIMIVGGLI | 705 |
| 8 | EISNYTGYIYQLIEESQNQQEKNEQELLALDKWASLWNWFDITNWLWYIKIFIMIVGGLI | 682 |
| 9 | EINNYTGYIYQLIEESQNQQEKNEQELLALDKWASLWNWFDITNWLWYIKIFIMIVGGLI | 705 |
| 10 | EISNYTGYIYQLIEESQNQQEKNEQELLALDKWASLWNWFDITNWLWYIKIFIMIVGGLI | 682 |
| 11 | EINNYTGYIYQLIEESQNQQEKNEQELLALDKWASLWNWFDITNWLWYIKIFIMIVGGLI | 684 |
| 12 | EINNYTGYIYQLIEESQNQQEKNEQELLAWDKWASLWNWFDITNWLWYIKIFIMIVGGLI | 707 |
| 13 | EINNYTGYIYQLIEESQNQQEKNEQELLAWDKWASLWNWFDITNWLWYIKIFIMIVGGLI | 707 |
| 14 | EINNYTGYIYQLIEESQNQQEKNEQELLALDKWASLWNWFDITNWLWYIKIFIMIVGGLI | 707 |
| 15 | EINNYTGYIYQLIEESQNQQEKNEQELLAWDKWASLWNWFDITNWLWYIKIFIMIVGGLI | 707 |
| 16 | EINNYTGYIYQLIEESQNQQEKNEQELLALDKWASLWNWFDITNWLWYIKIFIMIVGGLI | 683 |
| 17 | EISNYTGYIYQLIEESQNQQEKNEQELLALDKWASLWNWFDITNWLWYIKIFIMIVGGLI | 682 |
| 18 | EINNYTGYIYQLIEESQNQQE-----LLAWDKWASLWNWFDITNWLWYIKIFIMIVGGLI | 700 |
| 19 | EINNYTGYIYQLIEESQNQQEKNEQELLAWDKWASLWNWFDITNWLWYIKIFIMIVGGLI | 679 |
| 20 | EINNYTGYIYQLIEESQNQQEKNEQELLAWDKWASLWNWFDITNWLWYIKIFIMIVGGLI | 705 |
| 21 | EIDNYTGYIYQLIEESQNQQEKNEQELLALDKWASLWNWFDITNWLWYIKIFIMIVGGLI | 679 |
| 22 | EISNYTGYIYQLIEESQNQQEKNEQELLALDKWASLWNWFDITNWLWYIKIFIMIVGGLI | 682 |
| 23 | EINNYTGYIYQLIEESQNQQEKNEQELLALDKWASLWNWFDITNWLWYIKIFIMIVGGLI | 712 |
| 24 | EISNYTGYIYQLIEESQNQQEKNEQELLALDKWASLWNWFDITNWLWYIKIFIMIVGGLI | 682 |
| 25 | EINNYTGYIYQLIEESQNQQEKNEQELLALDKWASLW-WFDITNWLWYIKIFIMIVGGLI | 706 |
| 26 | EINNYTGYIYQLIEESQNQQEKNEQELLAWDKWASLWNWFDITNWLWYIKIFIMIVGGLI | 707 |
| 27 | EINNYTGYIYQLIEESQNQQEKNEQELLALDKWASLWNWFDITNWLWYIKIFIMIVGGLI | 708 |
| 28 | EINNYTGYIYQLIEESQNQQEKNEQELLAWDKWASLWNWFDITNWLWYIKIFIMIVGGLI | 703 |
| 29 | EINNYTGYIYQLIEESQNQQEKNEQELLAWDKWASLWNWFDITNWLWYIKIFIMIVGGLI | 708 |
| 30 | EINNYTGYIYQLIEESQNQQEKNEQELLA-DKWASLWNWFDITNWLWYIKIFIMIVGGLI | 704 |
| 31 | EINNYTGYIYQLIEESQNQQEKNEQELLAWDKWASLWNWFDITNWLWYIKIFIMIVGGLI | 705 |
| 32 | EINNYTGYIYQLIEESQNQQEKNEQELLAWDKWASLWNWFDITNWLWYIKIFIMIVGGLI | 705 |
| 33 | EINNYTGYIYQLIEESQNQQEKNEQELLALDKWASLWNWFDITNWLWYIKIFIMIVGGLI | 681 |
| 34 | EINNYTGYIYQLIEESQNQQEKNEQELLAWDRWASLWNWFDITNWLWYIKIFIMIVGGLI | 705 |
| 35 | EISNYTGYIYQLIEESQNQQEKNEQELLALDKWASLWNWFDITNWLWYIKIFIMIVGGLI | 682 |
| 36 | EINNYTGYIYQLIEESQNQQEKNEQELLAWDKWASLWNWFDITNWLWYIKIFIMIVGGLI | 707 |
| 37 | EINNYTGYIYQLIEESQNQQEKNEQELLALDKWASLWNWFDITNWLWYIKIFIMIVGGLI | 707 |
| 38 | EINNYTGYIYQLIEESQNQQEKNEQELLALDKWASLWNWFDITNWLWYIKIFIMIVGGLI | 707 |
| 39 | EIDNYTGYIYQLIEESQNQQEKNEQELLALDKWASLWNWFDITNWLWYIKIFIMIVGGLI | 683 |
| 40 | EISNYTGYIYQLIEESQNQQEKNEQELLALDKWASLWNWFDITNWLWYIKIFIMIVGGLI | 681 |
| 41 | EINNYTGYIYQLIEESQNQQEKNEQELLALDKWASLWNWFDITNWLWYIKIFIMIVGGLI | 707 |
| 42 | EISNYTGYIYQLIEESQNQQEKNEQELLALDKWASLWNWFDITNWLWYIKIFIMIVGGLI | 682 |

| | | |
|---|---|---|
| | GLRIVFTVLSIVNRVRQGYSPLSFQTHLPAQRGPDRPEGIGEEGGERDRDRSDPLVNGFL | |
| 1 | GLRIVFTVLSIVNRVRQGYSPLSFQTHLPAQRGPDRPEGIGEEGGERDRDRSDPLVNGFL | 743 |
| 2 | GLRIVFTVLSIVNRVRQGYSPLSFQTHLPAQRGPDRPEGIGEEGGERDRDRSDPLVNGFL | 765 |
| 3 | GLRIVFTVLSIVNRVRQGYSPLSFQTHLPAQRGPDRPEGIGEEGGERDRDRSDPLVNGFL | 742 |
| 4 | GLRIVFTVLSIVNRVRQGYSPLSFQTHLPAQRGPDRPEGIGEEGGERDRDRSDPLVNGFL | 742 |
| 5 | GLRIVFTVLSIVNRVRQGYSPLSFQTHLPAQRGPDRPEGIGEEGGERDRDRSDPLVNGFL | 743 |
| 6 | GLRIVFTVLSIVNRVRQGYSPLSFQTHFPAQRGPDRPEGIGEEGGERDRDRSAPLVNGFL | 767 |
| 7 | GLRIVFTVLSIVNRVRQGYSPLSFQTHFPAQRGPDRPEGIGEEGGERDRDRSDPLVNGFL | 765 |
| 8 | GLRIVFTVLSIVNRVRQGYSPLSFQTHLPAQRGPDRPEGIGEEGGERDRDRSDPLVNGFL | 742 |
| 9 | GLRIVFTVLSIVNRVRQGYSPLSFQTHLPAQRGPDRPEGIGEEGGERDRDRSDPLVNGFL | 765 |
| 10 | GLRIVFTVLSIVNRVRQGYSPLSFQTHLPAQRGPDRPEGIGEEGGERDRDRSDPLVNGFL | 742 |
| 11 | GLRIVFTVLSIVNRVRQGYSPLSFQTHLPAQRGPDRPEGIGEEGGERDRDRSDPLVNGFL | 744 |
| 12 | GLRIVFTVLSIVNRVRQGYSPLSFQTHLPAQRGPDRPEGIGEEGGERDRDRSDPLVTGFL | 767 |
| 13 | GLRIVFTVLSIVNRVRQGYSPLSFQTHLPAQRGPDRPEGIGEEGGERDRDRSAPLVNGFL | 767 |
| 14 | GLRIVFTVLSIVNRVRQGYSPLSFQTHFPAQRGPDRPEGIGEEGGERDRDRSAPLVNGFL | 767 |
| 15 | GLRIVFTVLSIVNRVRQGYSPLSFQTHFPAQRGPDRPEGIGEEGGERDRDRSAPLVNGFL | 767 |
| 16 | GLRIVFTVLSIVNRVRQGYSPLSFQTHLPAQRGLDRPEGIGEEGGERDRDRSDPLVNGFL | 743 |
| 17 | GLRIVFTVLSIVNRVRQGYSPLSFQTHLPAQRGPDRPEGIGEEGGERDRDRSDPLVNGFL | 742 |
| 18 | GLRIVFAVLSIVNRVRQGYSPLSFQTHLPAQRGPDRPEGIGEEGGERDRDRSDPLVNGFL | 760 |
| 19 | GLRIVFTVLSIVNRVRQGYSPLSFQTHLPAQRGPDRPEGIGEEGGERDRDRSDPLVNGFL | 739 |
| 20 | GLRIVFTVLSIVNRVRQGYSPLSFQTHLPAQRGPDRPEGIGEEGGERDRDRSDPLVNGFL | 765 |
| 21 | GLRIVFTVLSIVNRVRQGYSPLSFQTHLPAQRGLDRPEGIGEEGGERDRDRSDPLVNGFL | 739 |
| 22 | GLRIVFTVLSIVNRVRQGYSPLSFQTHLPAQRGPDRPEGIGEEGGERDRDRSDPLVNGFL | 742 |
| 23 | GLRIVFTVLSIVNRVRQGYSPLSFQTHFPAQRGPDRPEGIGEEGGERDRDRSALLVNGFL | 772 |
| 24 | GLRIVFTVLSIVNRVRQGYSPLSFQTHLPAQRGPDRPEGIGEEGGERDRDRSDPLVNGFL | 742 |
| 25 | GLRIVFTVLSIVNRVRQGYSPLSFQTHFPAQRGPDRPEGIGEEGGERDRDRSDPLVNGFL | 766 |
| 26 | GLRIVFTVLSIVNRVRQGYSPLSFQTHFPAQRGPDRPEGIGEEGGERDRDRSAPLVNGFL | 767 |
| 27 | GLRIVFTVLSIVNRVRQGYSPLSFQTHFPAQRGPDRPEGIGEEGGERDRDRSAPLVNGFL | 768 |
| 28 | GLRIVFTVLSIVNRVRQGYSPLSFQTHFPAQRGPDRPEGIGEEGGERDRDRSAPLVNGFL | 763 |
| 29 | GLRIVFTVLSIVNRVRQGYSPLSFQTHLPAQRGPDRPEGIGEEGGERDRDRSDPLVTGFL | 768 |
| 30 | GLRIVFTVLSIVNRVRQGYSPLSFQTHLPAQRGPDRPEGIGEEGGERDRDRSDPLVTGFL | 764 |
| 31 | GLRIVFTVLSIVNRVRQGYSPLSFQTHFPAQRGPDRPEGIGEEGGERDRDRSDPLVNGFL | 765 |
| 32 | GLRIVFTVLSIVNRVRQGYSPLSFQTHFPAQRGPDRPEGIGEEGGERDRDRSDPLVNGFL | 765 |
| 33 | GLRIVFTVLSIVNRVRQGYSPLSFQTHLPAQRGPDRPEGIGEEGGERDRDRSDPLVNGFL | 741 |
| 34 | GLRIVFTVLSIVNRVRQGYSPLSFQTHLPAQRGPDRPEGIGEEGGERDRDRSDPLVTGFL | 765 |
| 35 | GLRIVFTVLSIVNRVRQGYSPLSFQTHLPAQRGPDRPEGIGEEGGERDRDRSDPLVNGFL | 742 |
| 36 | GLRIVFTVLSIVNRVRQGYSPLSFQTHFPAQRGPDRPEGIGEEGGERDRDRSAPLVNGFL | 767 |
| 37 | GLRIVFTVLSIVNRVRQGYSPLSFQTHLPAQRGPDRPEGIGEEGGERDRDRSAPLVNGFL | 767 |
| 38 | GLRIVFTVLSIVNRVRQGYSPLSFQTHFPAQRGPDRPEGIGEEGGERDRDRSDPLVNGFL | 767 |
| 39 | GLRIVFTVLSIVNRVRQGYSPLSFQTHLPAQRGLDRPEGIGEEGGERDRDRSDPLVNGFL | 743 |
| 40 | GLRIVFTVLSIVNRVRQGYSPLSFQTHLPAQRGPDRPEGIGEEGGERDRDRSDPLVNGFL | 741 |
| 41 | GLRIVFTVLSIVNRVRQGYSPLSFQTHFPAQRGPDRPEGIGEEGGERDRDRSAPLVNGFL | 767 |
| 42 | GLRIVFTVLSIVNRVRQGYSPLSFQTHLPAQRGPDRPEGIGEEGGERDRDRSDPLVNGFL | 742 |

| | Sequence | |
|---|---|---|
|  | TLIWSDLRSLCLFSYHRLRDLLLIVTRIVELLGR-------RGWEVLKYWWNLLQYWSQE |  |
| 1 | ALIWSDLRSLCLFSYHRLRDLLLIVTRIVELLGR-------RGWEVLKYWWNLLQYWSQE | 796 |
| 2 | TFIWSDLRSLCLFSYHHLRDLLLIVTRIVELLGR-------RGWEVLKYWWNLLQYWSQE | 818 |
| 3 | TLIWSDLRSLCLFSYHRLRDLLLIATRIVELLGR-------RGWEVLKYWWNLLQYWSQE | 795 |
| 4 | TLIWSDLRSLCLFSYHRLRDLLLIATRIVELLGR-------RGWEVLKYWWNLLQYWSQE | 795 |
| 5 | TLIWSDLRSLCLFSYHHLRDLLLIVTRIVELLGR-------RGWEVLKYWWNLLQYWSQE | 796 |
| 6 | TLIWSDLRSLCLFSYHRLRDLLLIATRIVELLGR-------RGWEVLKYWWNLLQYWSQE | 820 |
| 7 | TLIWSDLRSLCLFSYHHLRDSLLIVTRIVELLGR-------RGWEVLKYWWNLLQYWSQE | 818 |
| 8 | TLIWSDLRSLCLFSYHRLRDLLLIATRIVELLGR-------RGWEVLKYWWNLLQYWSQE | 795 |
| 9 | TLIWSDLRSLCLFSYHHLRDSLLIVTRIVELLGR-------RGWEVLKYWWNLLQYWSQE | 818 |
| 10 | TIIWSDLRSLCLFSYHRLRDLLLIATRIVELLGR-------RGWEVLKYWWNLLQYWSQE | 795 |
| 11 | TLIWSDLRSLCLFSYHHLRDLLLIVTRIVELLGR-------RGWEVLKYWWNLLQYWSQE | 797 |
| 12 | TLIWSDLRSLCLFSYHRLSDLLLIVTRIVELLGR-------RGWEVLKYWWNLLQYWSQE | 820 |
| 13 | TLIWSDLRSLCLFSYHHLRDSLLIVTRIVELLGR-------RGWEVLKYWWNLLQYWSQE | 820 |
| 14 | TLIWSDLRSLCLFSYHHLRDSLLIVTRIVELLGR-------RGWEVLKYWWNLLQYWSQE | 820 |
| 15 | TFIWSDLRSLCLFSYHHLRDSLLIVTRIVELLGR-------RGWEVLKYWWNLLQYWSQE | 820 |
| 16 | TLIWSDLRSLCLFSYHHLRDLLLIVTRIVELLGR-------RGWEVLKYWWNLLQYWSQE | 796 |
| 17 | TLIWSDLRSLCLFSYHRLRDLLLIATRIVELLGR-------RGWEVLKYWWNLLQYWSQE | 795 |
| 18 | TLIWSDLRSLCLFSYHRLRDLLLIATRIVELLGR-------RGWEVLKYWWNLLQYWSQE | 813 |
| 19 | TLIWSDLRSLGLFSYHHLRDLLLIVTRIVELLGR-------RGWEVLKYWWNLLQYWSQE | 792 |
| 20 | TFIWSDLRSLCLFSYHRLSDLLLIVTRIVELLGR-------RGWEVLKYWWNLLQYWSQE | 818 |
| 21 | TLIWSDLRSLCLFSYHHLRDLLLIVTRIVELLGR-------RGWEVLKYWWNLLQYWSQE | 792 |
| 22 | TLIWSDLRSLCLFSYHRLRDLLLIATRIVELLGR-------RGWEVLKYWWNLLQYWSQE | 795 |
| 23 | TLIWSDLRSLCLFSYHHLRDSLLIVTRIVELLGR-------RGWEVLKYWWNLLQYWSQE | 825 |
| 24 | TLIWSDLRSLCLFSYHRLRDLLLIATRIVELLGR-------RGWEVLKYWWNLLQYWSQE | 795 |
| 25 | TLIWSDLRSLCLFSYHRLRDLLLIATRIVELLGR-------RGWEVLKYWWNLLQYWSQE | 819 |
| 26 | TLIWSDLRSLCLFSYHHLRDLLLIVTRIVELLGR-------RGWEVLKYWWSLLQYWSQE | 820 |
| 27 | TLIWSDLRSLCLFSYHRLRDLLLIATRIVELLGR-------RGWEVLKYWWNLLQYWSQE | 821 |
| 28 | TLIWSDLRSLCLFSYHHLRDSLLIVTRIVELLGR-------RGWEVLKYWWNLLQYWSQE | 816 |
| 29 | TLIWSDLRSLCLFSYHRLSDLLLIVTRIVELLGR-------RGWEVLKYWWNLLQYWSQE | 821 |
| 30 | TLIWSDLRSLCLFSYHRLSDLLLIVTRIVELLGR-------RGWEVLKYWWNLLQYWSQE | 817 |
| 31 | TLIWSDLRSLCLFSYHRLSDLLLIITRIVELLGR-------RGWEVLKYWWNLLQYWSQE | 818 |
| 32 | TFIWSDLRSLCLFSYHRLSDLLLIATRIVELLGR-------RGWEVLKYWWNLLQYWSQE | 818 |
| 33 | TLIWSDLRSLCLFSYHHLRDLLLIVTRIVELLGR-------RGWEVLKYWWNLLQYWSQE | 794 |
| 34 | TLIWSDLRSLCLFSYHRLSDLLLIVTRIVELLGR-------RGWEVLKYWWNLLQYWSQE | 818 |
| 35 | TLIWSDLRSLCLFSYHRLRDLLLIATRIVELLGR-------RGWEVLKYWWNLLQYWSQE | 795 |
| 36 | TLIWSDLRSLCLFSYHRLRDLLLIATRIVELLGR-------RGWEVLKYWWNLLQYWSQE | 820 |
| 37 | TLIWSDLRSLCLFSYHRLRDLLLIATRIVELLGR-------RGWEVLKYWWNLLQYWSQE | 820 |
| 38 | TLIWSDLRSLCLFSYHRLRDLLLIATRIVELLGR-------RGWEVLKYWWNLLQYWSQE | 820 |
| 39 | TLIWSDLRSLCLFSYHHLRDLLLIVTRIVELLGR-------RGWEVLKYWWNLLQYWSQE | 796 |
| 40 | TLIWSDLRSLCLFSYHRLRDLLLIATRIVELLGR-------RGWEVLKYWWNLLQYWSQE | 794 |
| 41 | TLIWSDLRSLCLFSYHHLRDSLLIVTRIVELLGR-------RGWEVLKYWWNLLQYWSQE | 820 |
| 42 | TLIWSDLRSLCLFSYHRLRDLLLIATRIVELLGR-------RGWEVLKYWWNLLQYWSQE | 795 |

| | | |
|---|---|---|
| | LKNSAVSLLNATAIAVAEGTDRVIEVVQRAGRAIRHIPRRIRQGLERALL | |
| 1 | LKNSAVSLLNATAIAVAEGTDRVIEVVQRACRAILHIPRRIRQGLERALL | 846 |
| 2 | LKKSAVSLLNATAIAVAEGTDRVIEVVQRAGRAIRHIPRRIRQGLERALL | 868 |
| 3 | LKNSAVSLFNATAIAVAEGTDRVIEVVQRAGRAIRHIPRRIRQGLERALL | 845 |
| 4 | LRNSAVSLLNATAIAVAEGTDRVIEVVQRAGRAIRHIPRRIRQGLERALL | 845 |
| 5 | LKNSAVSLLNATAIAVAEGTDRVIEVVQRAGRAIRHIPRRIRQGLERALL | 846 |
| 6 | LKKSAVSLLNATAIAVAEGTDRVIEVVQRAGRAIRHIPRRIRQGLERALL | 870 |
| 7 | LKKSAVSLLNATAIAVAEGTDRVIEVVQRTGRAIRHIPRRIRQGLERALL | 868 |
| 8 | LRNSAVSLLNATAIAVAEGTDRVIEVVQRAGRAIRHIPRRIRQGLERALL | 845 |
| 9 | LKKSAVSLLNATAIAVAEGTDRVIEVVQRTGRAIRHIPRRIRQGLERALL | 868 |
| 10 | LRNSAVSLLNATAIAVAEGTDRVIEVVQRAGRAIRHIPRRIRQGLERALL | 845 |
| 11 | LKNSAVSLLNATAIAVAEGTDRVIEVVQRAGRAIRHIPRRIRQGLERALL | 847 |
| 12 | LTNSAVSLLNATAIAVAEGTDRVIEVVQRAGRAVRHIPRRIRQGLERALL | 870 |
| 13 | LKKSAVSLLNATAIAVAEGTDRVIEVAQRTGRAIRHIPRRIRQGLERALL | 870 |
| 14 | LKKSAVSLLNATAIAVAEGTDRVIEVVQRTGRAIRHIPRRIRQGLERALL | 870 |
| 15 | LKKSAVSLLNATAIAVAEGTDRVIEVVQRTGRAIRHIPRRIRQGLERALL | 870 |
| 16 | LKNSTVSLLNATAIAVAEGTDRVIEVVQRAGRAIRHIPRRIRQGLERALL | 846 |
| 17 | LRNSAVSLLNATAIAVAEGTDRVIEVVQRAGRAIRHIPRRIRQGLERALL | 845 |
| 18 | LKKSAVSLLNATAIAVAEGTDRVIEVVQRAGRAIRHIPRRIRQGLERALL | 863 |
| 19 | LKNSAVSLLNATAIAVAEGTDRVIEVVQRAGRAIRHIPRRIRQGLERALL | 842 |
| 20 | LKKSAVSLLNATAIAVAEGTDRVIEVVQRTGRAIRHIPRRIRQGLERALL | 868 |
| 21 | LKNSAVSLLNATAIAVAEGTDRVIEVVQRAGRAIRHIPRRIRQGLERALL | 842 |
| 22 | LRNSAVSLLNATAIAVAEGTDRVIEVVQRAGRAIRHIPRRIRQGLERALL | 845 |
| 23 | LKKSAVSLLNATAIAVAEGTDRVIEVVQRAGRAIRHIPRRIRQGLERALL | 875 |
| 24 | LRNSAVSLLNATAIAVAEGTDRVIEVVQRAGRAIRHIPRRIRQGLERALL | 845 |
| 25 | LKKSAVSLLNATAIAVAEGTDRVIEVVQRAGRAIRHIPRRIRQGLERALL | 869 |
| 26 | LKKSAVSLLNATAIAVAEGTDRVIEVVQRTGRAIRHIPRRIRQGLERALL | 870 |
| 27 | LKKSAVSLLNATAIAVAEGTDRVIEVVQRTGRAIRHIPRRIRQGLERALL | 871 |
| 28 | LKKSAVSLLNATAIAVAEGTDRVIEVVQRTGRAIRHIPRRIRQGLERALL | 866 |
| 29 | LTNSAVSLLNATAIAVAEGTDRVIEVVQRAGRAVRHIPRRIRQGLERALL | 871 |
| 30 | LTNSAVSLLNATAIAVAEGTDRVIEVVQRAGRAVRHIPRRIRQGLERALL | 867 |
| 31 | LTNSAVSLLNATAIAVAEGTDRVIEVVQRAGRAVRHIPRRIRQGLERALL | 868 |
| 32 | LKKSAVSLLNATAIAVAEGTDRVIEVVQRAGRAVRHIPRRIRQGLERALL | 868 |
| 33 | LKNSAVSLLNATAIAVAEGTDRVIEVVQRAGRAIRHIPRRIRQGLERALL | 844 |
| 34 | LTNSAVSLLNATAIAVAEGTDRVIEVVQRAGRAVRHIPRRIRQGLERALL | 868 |
| 35 | LRNSAVSLLNATAIAVAEGTDRVIEVVQRAGRAIRHIPRRIRQGLERALL | 845 |
| 36 | LKKSAVSLLNATAIAVAEGTDRVIEVVQRTGRAIRHIPRRIRQGLERALL | 870 |
| 37 | LKKSAVSLLNATAIAVAEGTDRGIEVVQRTGRAIRHIPRRIRQGLERALL | 870 |
| 38 | LKKSAVSLLNATAIAVAEGTDRVIEVVQRAGRAIRHIPRRIRQGLERALL | 870 |
| 39 | LKNSVVSLLNATAIAVAEGTDRVIEVVQRAGRAIRHIPRRIRQGLERALL | 846 |
| 40 | LRNSAVSLLNATAIAVAEGTDRVIEVVQRAGRAIRHIPRRIRQGLERALL | 844 |
| 41 | LKKSAVSLLNATAIAVAEGTDRVIEVVQRTGRAIRHIPRRIRQGLERALL | 870 |
| 42 | LRNSAVSLLNATAIAVAEGTDRVIEVVQRAGRAIRHIPRRIRQGLERALL | 845 |
