## Supplementary Movie 2 BG505 SOSIP+ for "Conformational flexibility of HIV-1 envelope glycoproteins modulates transmitted / founder sensitivity to broadly neutralizing antibodies"

### Slide 1
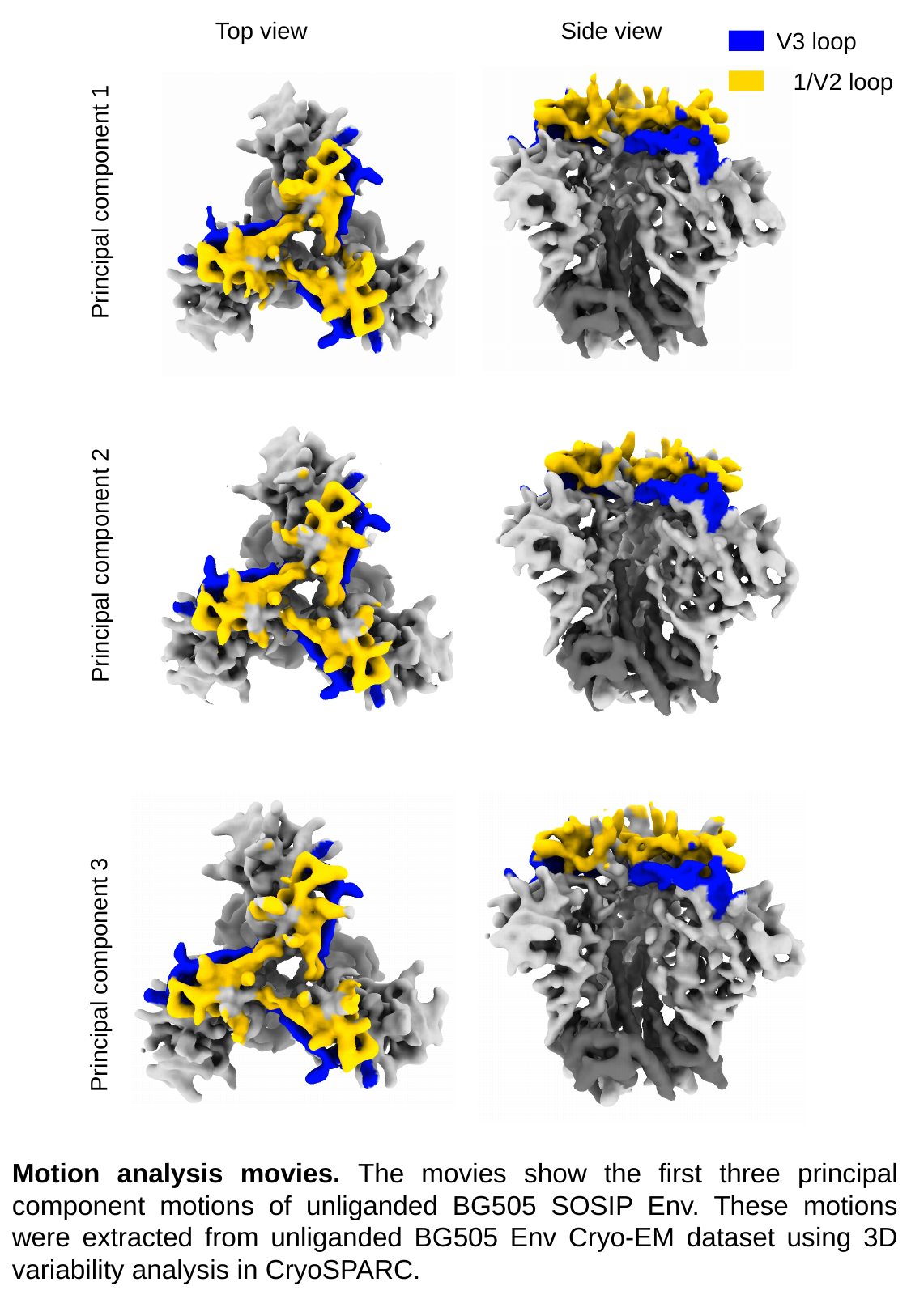

Top view Side view
V3 loop
V1/V2 loop
Principal component 1
Principal component 2
Principal component 3
Motion analysis movies. The movies show the first three principal component motions of unliganded BG505 SOSIP Env. These motions were extracted from unliganded BG505 Env Cryo-EM dataset using 3D variability analysis in CryoSPARC.
