## Supplementary Movie 1 1059 SOSIP for "Conformational flexibility of HIV-1 envelope glycoproteins modulates transmitted / founder sensitivity to broadly neutralizing antibodies"

### Slide 1
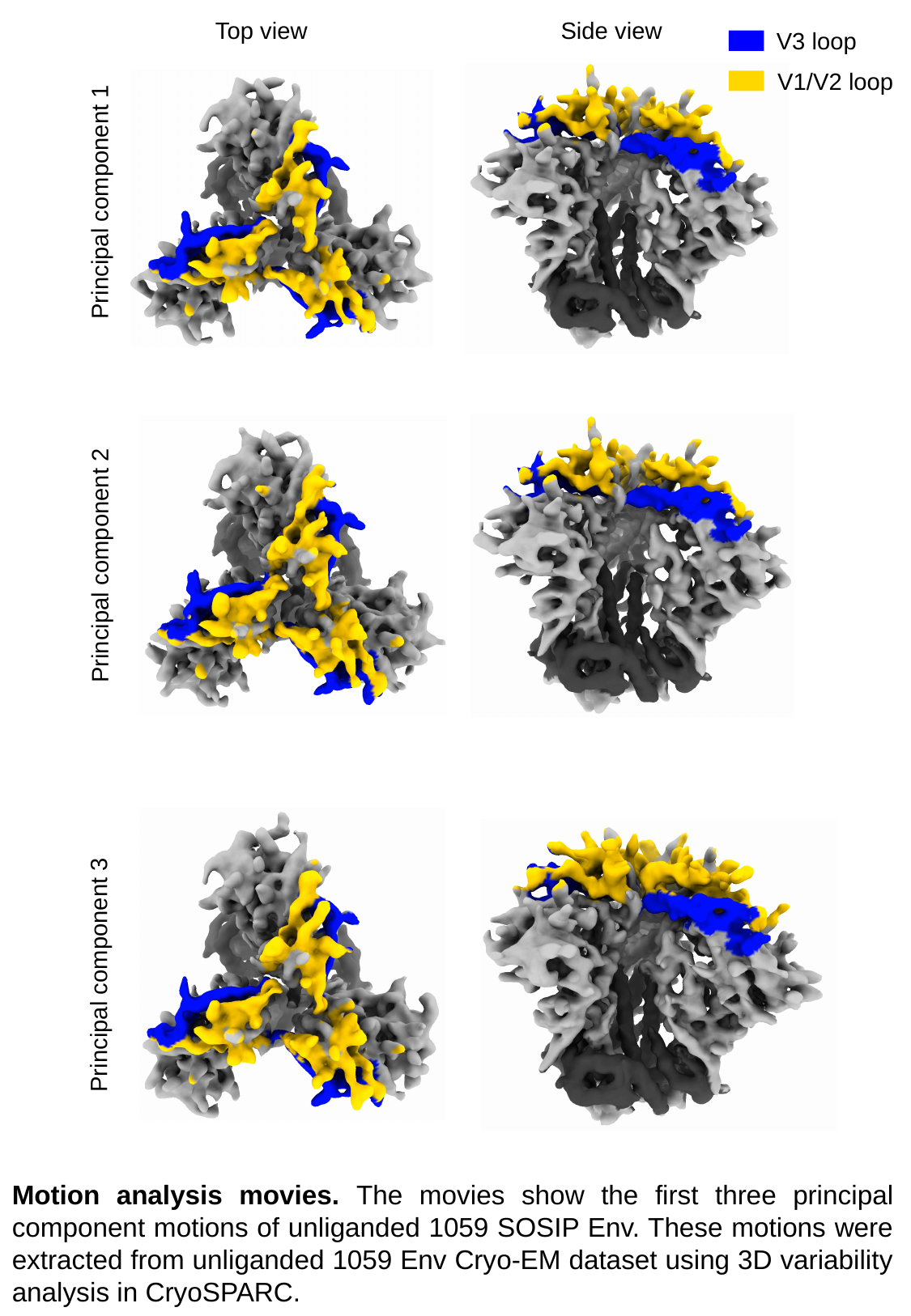

Top view Side view
V3 loop
V1/V2 loop
Principal component 1
Principal component 2
Principal component 3
Motion analysis movies. The movies show the first three principal component motions of unliganded 1059 SOSIP Env. These motions were extracted from unliganded 1059 Env Cryo-EM dataset using 3D variability analysis in CryoSPARC.
